# Explainable machine learning models of major crop traits from satellite-monitored continent-wide field trial data

**DOI:** 10.1101/2021.03.08.434495

**Authors:** Saul Justin Newman, Robert T Furbank

**Affiliations:** ARC Centre of Excellence for Translational Photosynthesis, Research School of Biology, Australian National University; Biological Data Science Institute, Australian National University

## Abstract

Four species of grass generate half of all human-consumed calories^1^. However, abundant biological data on species that produce our food remains largely inaccessible, imposing direct barriers to understanding crop yield and fitness traits. Here, we assemble and analyse a continent-wide database of field experiments spanning ten years and hundreds of thousands of machine-phenotyped populations of ten major crop species. Training an ensemble of machine learning models, using thousands of variables capturing weather, ground-sensor, soil, chemical and fertiliser dosage, management, and satellite data, produces robust cross-continent yield models exceeding R^2^ = 0.8 prediction accuracy. In contrast to ‘black box’ analytics, detailed interrogation of these models reveals fundamental drivers of crop behaviour and complex interactions predicting yield and agronomic traits. These results demonstrate the capacity of machine learning models to build unified, interpretable, and explainable models of crop behaviour, and highlight the powerful role of data in the future of food.

## Introduction

Over two billion people are projected to enter the world population by 2050^2^. Feeding these people sustainably requires an improved understanding of the complex evolutionary interactions driving major crop traits^3,4^, and the application of this knowledge through plant breeding to produce new crop varieties. However, large-scale data on the growth and yield traits of major crops remain generally unavailable or inaccessible to academic scientists^5^ and where available, big data often result in incomprehensible black box models of plant behaviour.

Here, we demonstrate the promise of machine learning (ML) and artificial intelligence algorithms to provide robust prediction of important agronomic traits, including yield, and improve our understanding of crop biology. By linking satellite data to a freely available ‘big’ dataset, the Australian National Variety Trials (NVTs), we develop a framework to train and test accurate ML models and extract meaningful and testable hypotheses from ML models. These findings highlight the power of unified, comprehensive cross-species models for the prediction and understanding of vital agronomic traits and crop species.

The NVT database constitutes one of the largest public experiments on earth (Fig. 1; Table 1). The NVTs capture over a quarter of a million unique variety-year observations, and over a million unique population-averaged phenotypes, aggregated from experimentally replicated plant populations in 6,547 geolocated randomised controlled experiments (Table 1; see database descriptor in Newman & Furbank^6^). Each population contains hundreds of individual plants, sown at controlled densities and replicated across a randomised controlled design trial, each conducted and phenotyped according to highly standardised protocols^7^. As such, the NVTs capture the aggregated phenotypic variation of hundreds of millions of individual organisms, across thousands of trial sites containing millions of plant populations^6^.

**Table 1.** Frequency of a million key agronomic traits by species.

| Phenotype | Wheat | Oat | Triticale | Barley | Canola | Lupin | Chickpea | Faba Bean | Field Pea | Lentil |
| --- | --- | --- | --- | --- | --- | --- | --- | --- | --- | --- |
| Early Growth | 7709 | 1229 | 1350 | 4258 | 5084 | 1168 | 1616 | 1104 | 1882 | 768 |
| Establishment | 9768 | 2561 | 2027 | 6116 | 6768 | 1804 | 1706 | 1347 | 2683 | 1300 |
| Flowering 50pct | 4826 | 748 | 734 | 1848 | 3326 | 805 | 963 | 566 | 864 | 473 |
| % Glucosinolates | 0 | 0 | 0 | 0 | 12460 | 0 | 0 | 0 | 0 | 0 |
| Heading date | 2084 | 561 | 279 | 1581 | 1355 | 453 | 388 | 359 | 313 | 109 |
| Hectolitre weight | 16684 | 4585 | 2473 | 8318 | 15170 | 5140 | 5343 | 3706 | 4460 | 1993 |
| Hectolitre weight (metadata) | 57863 | 3281 | 2212 | 15445 | 0 | 0 | 0 | 0 | 0 | 0 |
| Height cm | 3223 | 668 | 609 | 1691 | 2474 | 571 | 586 | 485 | 516 | 192 |
| Lodging score | 4334 | 614 | 885 | 2152 | 3116 | 756 | 1036 | 849 | 659 | 465 |
| Oil content | 0 | 0 | 0 | 0 | 12611 | 0 | 0 | 0 | 0 | 0 |
| Pct below 2.0 mm | 11839 | 3704 | 1415 | 6745 | 11158 | 4499 | 3987 | 3315 | 3826 | 1327 |
| Pct below 2.2 mm | 4624 | 881 | 1086 | 1477 | 3963 | 577 | 1356 | 389 | 634 | 620 |
| Protein | 16681 | 4585 | 2501 | 8195 | 15192 | 5146 | 5343 | 3706 | 4514 | 2003 |
| Protein (metadata) | 57863 | 3281 | 2212 | 1298 | 0 | 0 | 0 | 0 | 0 | 0 |
| Protein meal | 1994 | 309 | 217 | 833 | 10699 | 343 | 0 | 84 | 605 | 115 |
| Sieve 2.0mm (metadata) | 42305 | 1733 | 1923 | 0 | 0 | 0 | 0 | 0 | 0 | 0 |
| Thousand grain weight | 13537 | 3567 | 1773 | 7281 | 12459 | 3754 | 4318 | 3006 | 3682 | 1652 |
| Thousand grain weight (metadata) | 62872 | 3281 | 2212 | 17135 | 1869 | 0 | 0 | 0 | 0 | 0 |
| Site-Relative Yield | 73536 | 4892 | 2371 | 22145 | 18037 | 2877 | 5405 | 3592 | 4786 | 2709 |
| Yield, t/Ha | 54658 | 3873 | 2081 | 14117 | 14929 | 2747 | 3677 | 2224 | 3896 | 2260 |
| Zadoks Score | 2282 | 200 | 461 | 1130 | 1657 | 162 | 117 | 133 | 345 | 395 |
| Pass-Fail Yield Status | 73536 | 4892 | 2371 | 17377 | 18037 | 2877 | 5405 | 3592 | 4786 | 2709 |

Metadata in brackets indicate metadata-derived measurements that partially overlap reported data.

**Figure 1.**
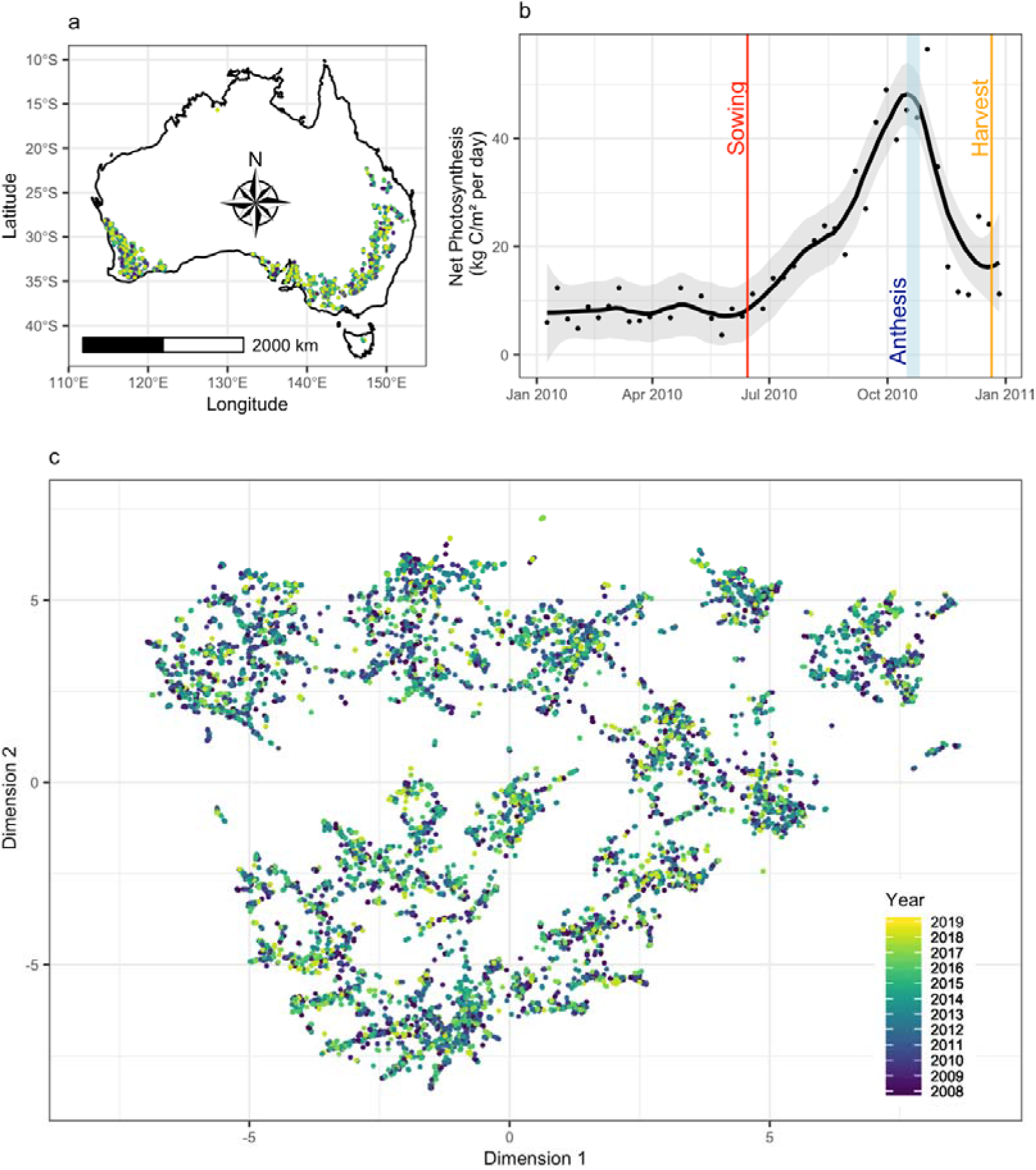
National variety trials and large-scale environmental patterns captured by satellite. Location of 6,547 successful field trial experiments **a**, tracked from 2008-2019 using remote sensing and ground station data. Remote sensing data captures environmental patterns at each location, through variables such as **b** net primary photosynthesis (black points; locally weighted smoothed spline), from pre-sowing, through the sowing date (red line), anthesis (blue shaded region), and harvest (orange), to post-harvest. These remote sensing data reveal environmental diversity across sites **c**, shown here by Uniform Manifold Approximation and Projection clustering, not captured by weather station data.

We linked these data to extensive satellite data characterising vegetation, temperature, and spectral patterns^8–11^ (Supplementary Table 1; Supplementary Fig. 1), weather station data from the Australian Bureau of Meteorology (BOM), over 10,000 standardised soil sample tests, extensive observational and site management data, over 50,000 field-years of stubble burn patterns^12^, the dose and timing of over 350,000 chemical and fertiliser applications, and crop rotation histories for over 10,000 field-years (see Newman & Furbank^6^). Collectively these data capture patterns of management inputs, vegetation, and environment, and include satellite-derived observations (Fig. 1b) that improve the categorisation of growing environments, provide information on canopy level agronomic traits, and reveal the complex environmental diversity of trial sites (Fig. 1c).

Using this open source^6^ environmental and agronomic database, we train a suite of robust ML algorithms for the prediction of key agronomic traits including yield, flowering, and grain protein (Table 1). In addition to providing a catalogue of phenotype prediction models, these models are used to demonstrate the potential for ML algorithms to generate comprehensible outputs and testable hypotheses beyond variable importance rankings and the ‘black box’ paradigm.

Using targeted ML model interrogation and analysis, we generate candidates for the causal drivers of complex traits including yield and grain protein content. We reduce random forests to predictively valuable and readable prediction rules, using an approach pioneered by Deng^13^, with direct and testable outcomes for agronomic research. This model reduction approach reveals cross-domain interactions between variables that robustly predict trait variation. As a result, rather than produce a ‘black box’ prediction model, these analyses reveal pathways to generate both accurate forecast models and potentially useful and biologically informative hypotheses.

## Results

Machine learning provided clear and accurate predictive models across a broad array of challenges. However, to provide a robust estimate of model accuracy it was necessary to extend model evaluation beyond the standard analytical framework. The standard approach to assessing ML accuracy is to use random-sample holdout values, with random observations excluded from the dataset prior to model training, and these values subsequently used as predictive targets to evaluate model accuracy. However, across all models this testing approach generated a misleading picture of model accuracy (Supplementary Table 2).

**Table 2.** Yield prediction accuracy over diverse species and models.

| Random Holdout<br>Trial Prediction | Accuracy R<sup>2</sup> of models trained on all data to: |  |  |
| --- | --- | --- | --- |
| Model | TOS | 100DAS | 200DAS |
| RPRM | 0.60 | 0.66 | 0.76 |
| Naïve BCRF | 0.76 | 0.79 | 0.82 |
| xvBCRF | 0.80 | 0.81 | 0.84 |
| XGBM | 0.64 | 0.73 | 0.78 |
| LSVM | 0.64 | 0.64 | 0.68 |
| PLSR | 0.58 | 0.73 | 0.76 |
| Annual Forecasting Predictions |  |  |  |
| Model | TOS | 100DAS | 200DAS |
| RPRM | 0.58 | 0.42 | 0.59 |
| Naïve BCRF | 0.67 | 0.71 | 0.70 |
| xvBCRF | 0.68 | 0.72 | 0.69 |
| XGBM | 0.52 | 0.61 | 0.69 |
| LSVM | 0.38 | 0.37 | 0.45 |
| PLSR | 0.58 | 0.68 | 0.74 |

DAS (Days After Sowing); TOS (Time of Sowing)

Model accuracy appeared to be systematically over-estimated when using random-sample holdout populations (Supplementary Table 2). We overcame this problem using a more rigorous model evaluation framework (Fig. 2) that tested machine-learning models using unobserved randomly-sampled field trials (hereafter ‘holdout trial prediction’; Fig. 2a-b), or all field trials observed in ‘future’ years hidden from model training (hereafter ‘annual forecast prediction’; Fig. 2a,c). By testing models in locations and years excluded from model training, this approach substantially reduced the reported accuracy of ML models (Supplementary Table 2) while ensuring structural risk minimisation and robust, translatable models. For example, under the standard random holdout data approach LSVM models reported an R^2^ = 0.99 prediction accuracy (Supplementary Table 2; RMSE = 0.17; N = 90,765). When the same models were evaluated using holdout trial prediction, predicting unobserved randomly-sampled field trial locations, model accuracy fell to between R^2^ = 0.64 and 0.68 (RMSE = 1.09-1.50; Supplementary Table 2; Supplementary Code 1).

**Figure 2.**
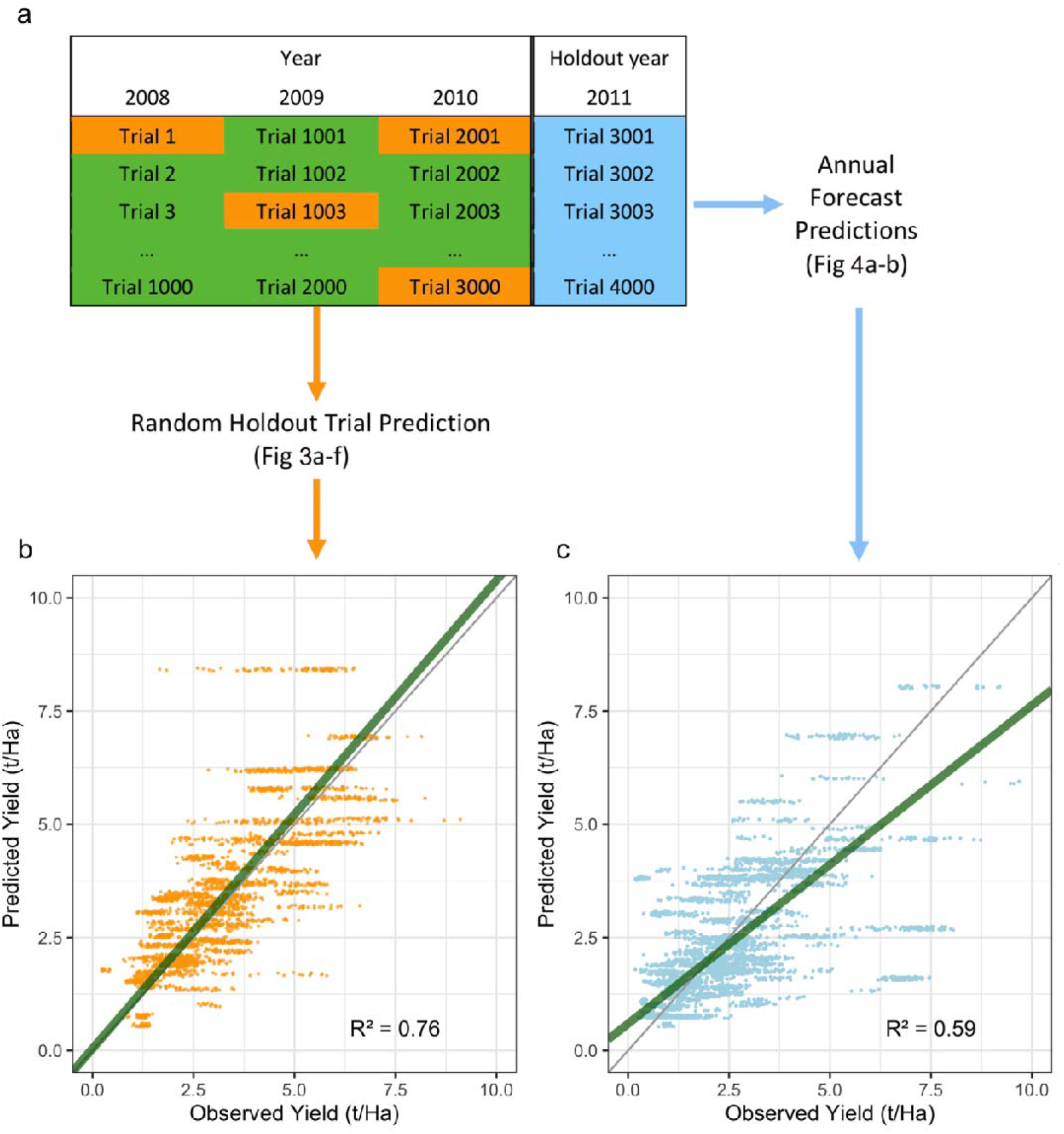
Schematic of model training and evaluation. All machine learning models are trained **a** on data all years before a given date (dark green), after excluding 100 randomly selected trials (orange). Each ML model is then used to predict phenotypic variation in both the **b** randomly selected holdout trials (orange) and **c** all trials in unobserved ‘future’ years (blue) excluded from model training. Model accuracy when predicting these holdout samples, as represented by the linear model fit (green) and residuals in scatterplots **b** and **c** (RPRMs shown), are used for model evaluation.

Despite reduced model accuracy under these more stringent test criteria, the accurate prediction of complex traits was possible under a wide range of ML models (Table 2). For example, when trained for the holdout trial prediction of yield using full-season data across all species (Fig. 3) foundational ML models such as unstratified ‘naïve’ Breiman-Cutler Random Forests^14,15^ (BCRFs; Fig. 3b; R^2^ = 0.82) and BCRFs cross-validated by calendar year (xvBCRFs; Fig. 3c; R^2^ = 0.84) captured variation within years with over R^2^ = 0.8 accuracy (N=3,182; p<10e-16; Supplementary Code 1), while extreme gradient boosting models^16^ (XGBMs; Fig. 3d; R^2^ = 0.78), recursively partitioned regression models^17^ or decision trees (RPRMs; Fig. 3a; R^2^ = 0.76), linear support vector machines^18,19^ (LSVMs; Fig. 3e; R^2^ = 0.68) and partial least squared regression models^20^ (PLSRs; Fig. 3f; R^2^ = 0.76) captured marked yield variation (N=3,101; p<10e-16; Supplementary Code 1).

**Figure 3.**
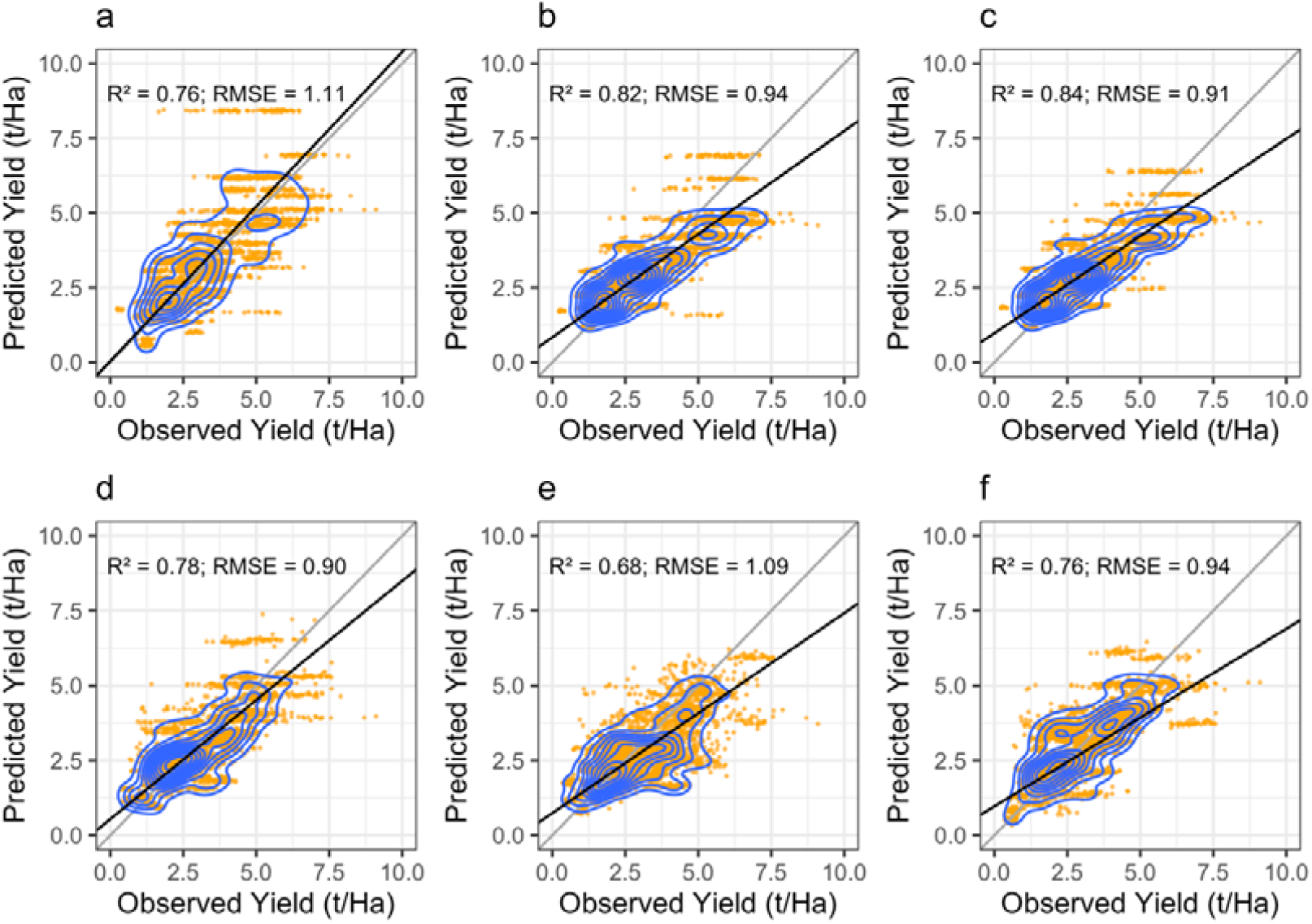
Accuracy of yield models subject to holdout trial prediction of 100 random unobserved field trials. Predictions of yield variation in 100 randomly selected hold-out field trials, by: **a** recursively partitioned regression model (RPRM) decision tree, **b** naïve or unstratified Breiman-Cutler random forests (BCRFs), **c** year-stratified BCRFs, **d** an extreme gradient boosting machine, **e** linear support vector regression, and **f** partial least squares regression (RMSE is root mean squared error; **a-c** sample size N = 3,182; **e-f** sample size N=3,101; all p<2.2e-16). Horizontal banding in tree-based models **a-d** is due to grouping of predictions into terminal decision tree nodes, y-axis is randomly jittered, blue contour lines indicate kernel density.

While baseline accuracy was high, ML models exhibited different performance across prediction challenges. These included agronomically relevant problems, such as using the data available at the time of sowing (TOS) for prediction of end-of-season phenotypic variation in new locations (Fig. 3; Table 2), the projection of end-of-season yield as a season progresses (Supplementary Fig. 2), and annual forecast predictions (Fig. 4; Table 2).

**Figure 4.**
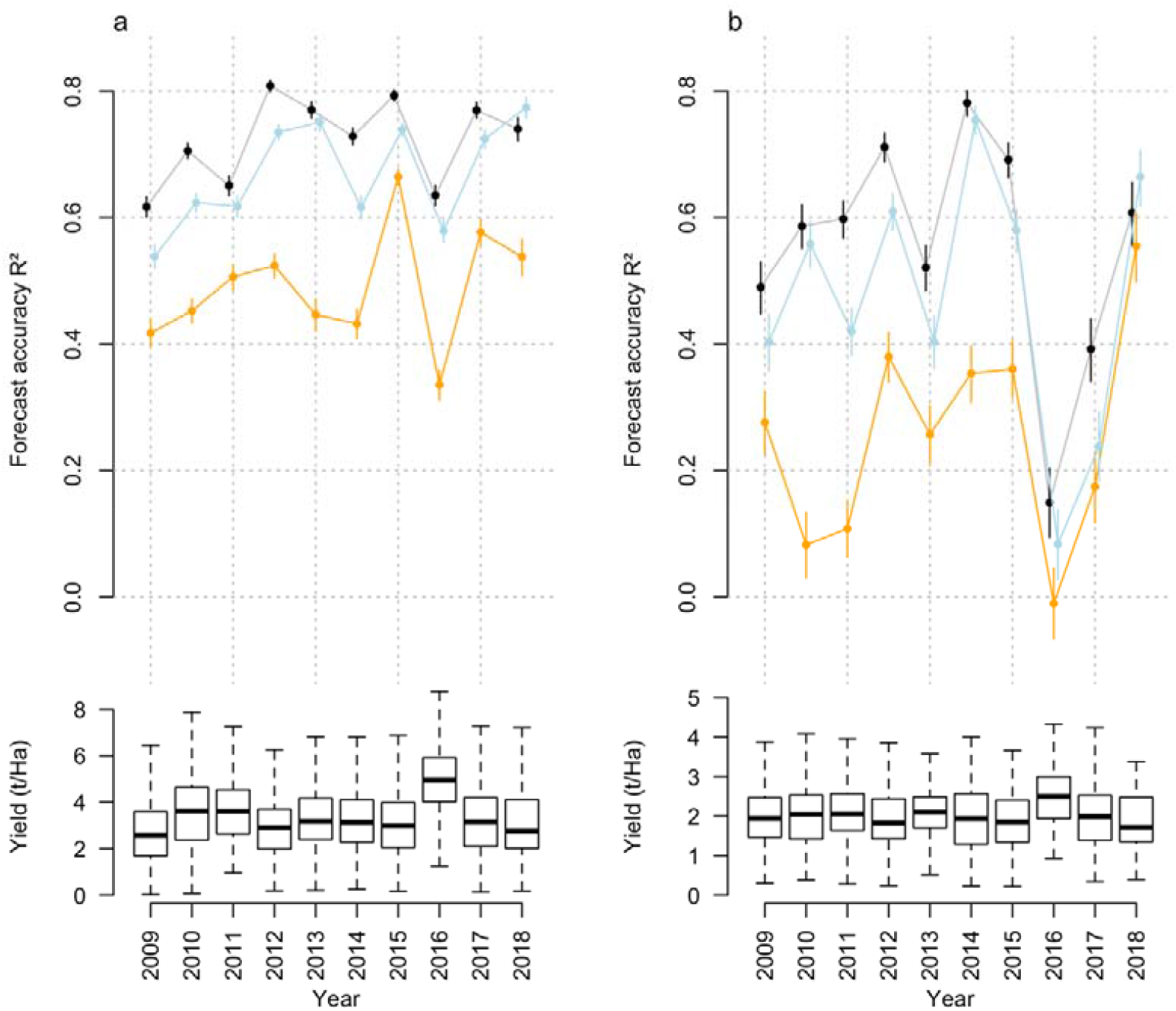
Cross-model and cross-species shifts in accuracy under rolling annual forecast prediction. Across diverse methods and phenotypes, annual forecast predictions of **a**, wheat and **b**, canola phenotypes display substantial annual shifts in accuracy, absent any change in model parameters or target sample size. Forecasting years with high yields, in 2016 and in 2013 for canola (boxplots, bottom), was coincident with reduced model accuracy, but otherwise models display no clear pattern. Colours indicate models trained using naïve BCRFs (black), LSVMs (orange), and PLSR models (blue), whiskers indicate 95% CI.

Model accuracy extended to other agronomically important complex traits. Several traits could be predicted under holdout trial prediction and annual forecasts with accuracy of R^2^=0.5 or more (Supplementary Fig. 3a-c), including complex traits such as protein (R^2^=0.48; Supplementary Fig. 3a), flowering time (R^2^ = 0.53; Supplementary Fig. 3b), and Glucosinolate content (R^2^=0.58; Supplementary Fig. 3c), while prediction of other traits such as proxy metrics of grain volume proved more challenging (Supplementary Fig. 3d-f).

Predictive accuracy was largely retained in models trained using only data available at the TOS. Under holdout trial prediction, yield data could be predicted with up to R^2^=0.80 accuracy under stratified cross-validated BCRFs (Table 2), while methods such as LSVMs (R^2^ = 0.64), XGBMs (R^2^ = 0.64), and PLSRs (R^2^ = 0.58) displayed more limited accuracy (Table 2).

Unsurprisingly, ML model accuracy fell under all annual forecast prediction tests (Table 2). For full-season data, over rolling annual forecasts, BCRF models retained the highest accuracy when predicting yield (black line Fig. 4; Table 2; Supplementary Code 1), followed by the PLSR model (blue line Fig. 4; R^2^ =0.74; Table 2), and the XGBM (R^2^ = 0.69; Supplementary Code 1). Similar patterns in accuracy occurred for ML models trained on data available at the TOS, 100 days after sowing (DAS) and 200 DAS, tested in 2018 only rather than rolling annual forecasts (Table 2). Again, the BCRF and PLSR models had the highest predictive accuracy, with the greatest reduction and lowest absolute accuracy in the LSVMs (from R^2^ = 0.64-0.68 to R^2^ = 0.37-0.45; Table 2).

As expected, ML approaches generally increased in accuracy with the inclusion of more years’ data (Fig. 4), and for predictions made at progressively later points in the growing season (Supplementary Fig. 3). However, rolling forecast data, in which models were used to predict variation in the next calendar year, displayed surprising patterns. Concurrent shifts in forecast accuracy occurred across different models, phenotypes, and species (Fig. 4a-b) with accuracy increasing (*e*.*g*. 2011-2012 and 2013-2014) or decreasing (*e*.*g*. 2015-2016) across diverse models and phenotypes. While observations were limited, changes in model forecast accuracy were correlated across ML models and species (p < 0.02; Supplementary Code 1) despite fixed model training parameters, the relatively constant sample size of target data, and the absence of overfitting (as measured in random holdout trials; Fig. 1b; Table 2). Some years were less predictable, across models and species, independent of model construction and sample size.

Annual forecast and random holdout trial prediction models were constructed both with and without partitioning crops into separate datasets for model training (Supplementary Code 1). For models such as xvBCRFs and LSVMS, incorporating all species in a single ‘omnibus’ dataset often, unsurprisingly, led to less accurate models (Supplementary Table 3). However, in some notable cases implementing a unified approach, where crop types were fit as binary independent variables, generated models of comparable or greater accuracy than models with identical parameters trained on single-species data (Fig. 3; Supplementary Fig. 4; Supplementary Code 1). For example, when predicting wheat yield in 2018, annual forecast models trained using only wheat data to 100 DAS had R^2^=0.66 accuracy under naïve BCRFs, R^2^=0.38 under LSVMs, and R^2^=0.59 under PLSR models (Supplementary Code 1). When these models were re-trained using cross-species data, using identical parameters, accuracy of 2018 wheat yield predictions improved marginally to R^2^ = 0.69 for BCRFs, R^2^ = 0.40 for LSVMs, and R^2^ = 0.65 for PLSRs (Supplementary Code 1). In Canola, construction of ‘omnibus’ cross-species models more substantially modified accuracy: at 100 DAS accuracy increased from R^2^ = 0.28 to 0.60 in naïve BCRF, and from R^2^ = 0.19 to 0.37 in PLSRs, with a low-accuracy model of R^2^ = 0.11 falling to R^2^ = −0.008 in the high prediction variance LSVM models (Supplementary Code 1).

**Table 3.**
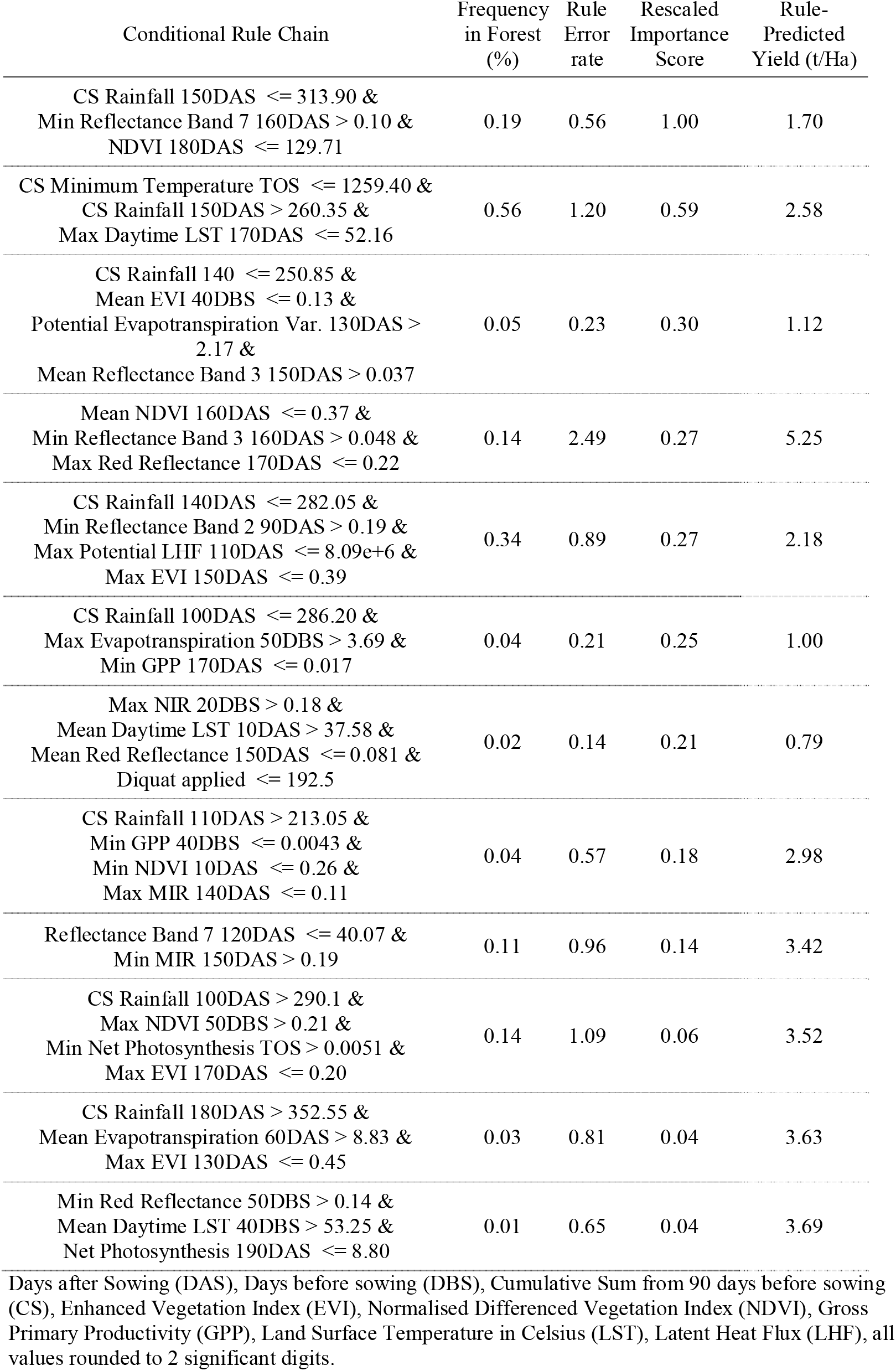
Variable interactions predicting canola yield.

| Conditional Rule Chain | Frequency<br>in Forest<br>(%) | Rule<br>Error<br>rate | Rescaled<br>Importance<br>Score | Rule-Predicted<br>Yield (t/Ha) |
| --- | --- | --- | --- | --- |
| CS Rainfall 150DAS <= 313.90 &<br>Min Reflectance Band 7 160DAS > 0.10 &<br>NDVI 180DAS <= 129.71 | 0.19 | 0.56 | 1.00 | 1.70 |
| CS Minimum Temperature TOS <= 1259.40 &<br>CS Rainfall 150DAS > 260.35 &<br>Max Daytime LST 170DAS <= 52.16 | 0.56 | 1.20 | 0.59 | 2.58 |
| CS Rainfall 140 <= 250.85 &<br>Mean EVI 40DBS <= 0.13 &<br>Potential Evapotranspiration Var. 130DAS ><br>2.17 &<br>Mean Reflectance Band 3 150DAS > 0.037 | 0.05 | 0.23 | 0.30 | 1.12 |
| Mean NDVI 160DAS <= 0.37 &<br>Min Reflectance Band 3 160DAS > 0.048 &<br>Max Red Reflectance 170DAS <= 0.22 | 0.14 | 2.49 | 0.27 | 5.25 |
| CS Rainfall 140DAS <= 282.05 &<br>Min Reflectance Band 2 90DAS > 0.19 &<br>Max Potential LHF 110DAS <= 8.09e+6 &<br>Max EVI 150DAS <= 0.39 | 0.34 | 0.89 | 0.27 | 2.18 |
| CS Rainfall 100DAS <= 286.20 &<br>Max Evapotranspiration 50DBS > 3.69 &<br>Min GPP 170DAS <= 0.017 | 0.04 | 0.21 | 0.25 | 1.00 |
| Max NIR 20DBS > 0.18 &<br>Mean Daytime LST 10DAS > 37.58 &<br>Mean Red Reflectance 150DAS <= 0.081 &<br>Diquat applied <= 192.5 | 0.02 | 0.14 | 0.21 | 0.79 |
| CS Rainfall 110DAS > 213.05 &<br>Min GPP 40DBS <= 0.0043 &<br>Min NDVI 10DAS <= 0.26 &<br>Max MIR 140DAS <= 0.11 | 0.04 | 0.57 | 0.18 | 2.98 |
| Reflectance Band 7 120DAS <= 40.07 &<br>Min MIR 150DAS > 0.19 | 0.11 | 0.96 | 0.14 | 3.42 |
| CS Rainfall 100DAS > 290.1 &<br>Max NDVI 50DBS > 0.21 &<br>Min Net Photosynthesis TOS > 0.0051 &<br>Max EVI 170DAS <= 0.20 | 0.14 | 1.09 | 0.06 | 3.52 |
| CS Rainfall 180DAS > 352.55 &<br>Mean Evapotranspiration 60DAS > 8.83 &<br>Max EVI 130DAS <= 0.45 | 0.03 | 0.81 | 0.04 | 3.63 |
| Min Red Reflectance 50DBS > 0.14 &<br>Mean Daytime LST 40DBS > 53.25 &<br>Net Photosynthesis 190DAS <= 8.80 | 0.01 | 0.65 | 0.04 | 3.69 |

Days after Sowing (DAS), Days before sowing (DBS), Cumulative Sum from 90 days before sowing (CS), Enhanced Vegetation Index (EVI), Normalised Differenced Vegetation Index (NDVI), Gross Primary Productivity (GPP), Land Surface Temperature in Celsius (LST), Latent Heat Flux (LHF), all values rounded to 2 significant digits.

Assessing the features and interactions behind predictively accurate models was approached on several fronts. While ‘black box’ models are generally impenetrable to further analysis, altering model inputs and examining shifts in model accuracy can provide limited insight into model mechanics. Collectively, for example, remote sensing data were a key driver of model accuracy (Supplementary Fig. 5; Supplementary Fig. 6). Under a leave-one-out model testing approach, removal of the remote sensing variables caused the greatest loss in predictive value (Supplementary Fig. 5; Supplementary Code 2) compared to the more marginal effect of removing management data, BOM weather station data, or metadata (Supplementary Fig. 5; Supplementary Code 2). This pattern, where satellite data held the greatest predictive value for model training, was reinforced across holdout trial and annual forecast model assessment (Supplementary Code 1). For example, under RPRM models the removal of satellite data reduces the accuracy of wheat yield annual forecast models, from R^2^ = 0.75 for models containing all input variables, to just R^2^=0.36 for models trained without satellite data (Supplementary Code 2). In contrast, the impact from removal of either weather station data, management data or metadata was, at worst, a marginal reduction in accuracy from R^2^ = 0.75 to 0.71 (Supplementary Code 2).

**Figure 5.**
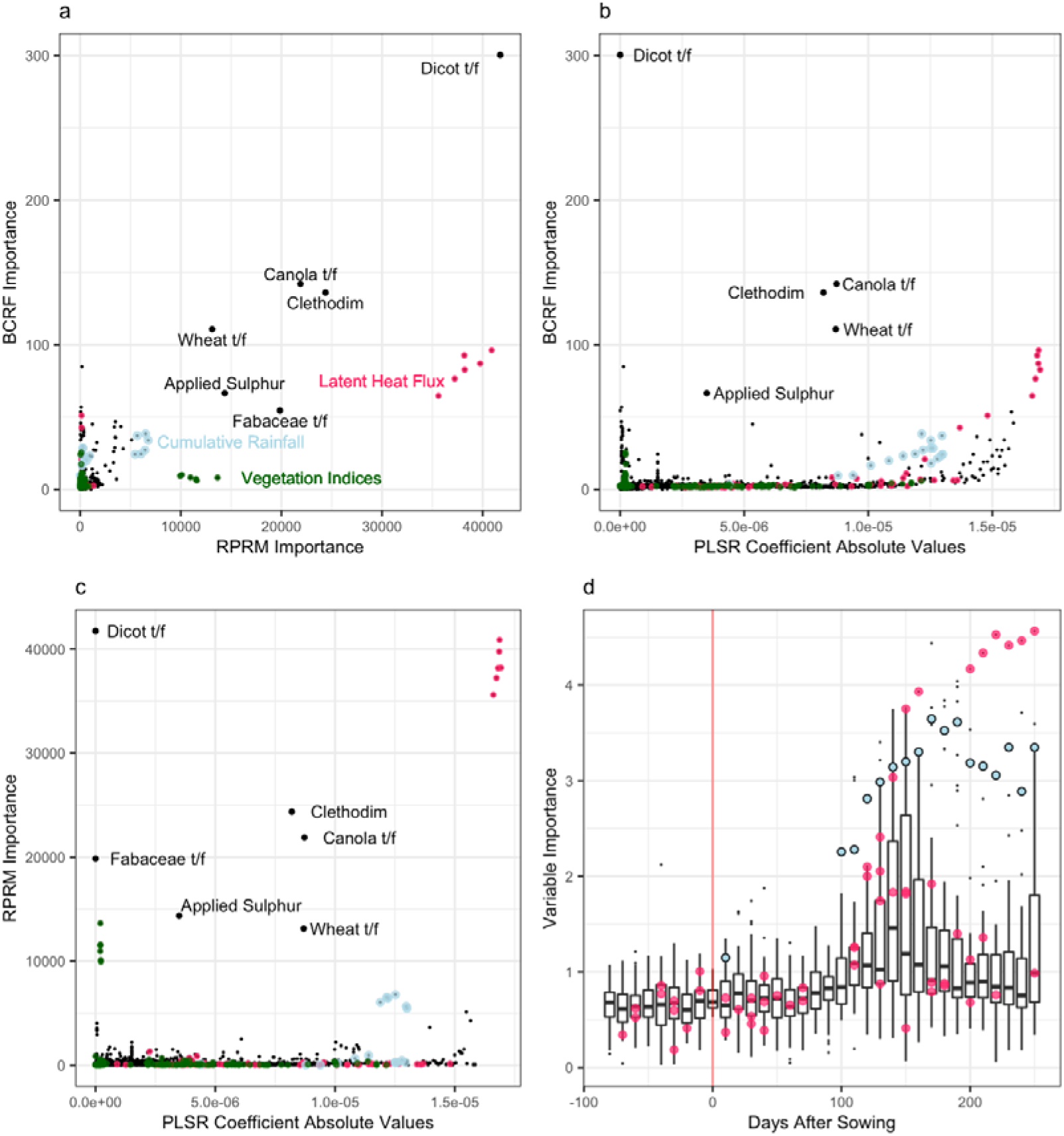
Concordance and pattern of variable importance across machine learning models. Variable importance indicators, as measured by voting heuristics in BCRF models (y-axes in **a** and **b**), the absolute value of coefficients in PLSR models (x-axes in **b** and **c**), or error reduction or shrinkage in RPRM models (x-axis in **a**, y-axis in **c**), showed some concordance between models. Likewise, as shown for BCRF models in **d**, the predictive importance of time-ordered variables increased throughout the season to a maximum around flowering and grain filling. Highly ranked variables across all models included cumulative total rainfall (blue), latent heat flux (fuchsia), dosage of the pesticide Clethodim (labelled), total applied sulphur fertiliser (labelled) and crop taxa (labelled). Models trained using full-season data shown.

Re-training models using only single domains, such as using only management or satellite data for model training, provided insight into the predictive value of domains without cross-domain interactions (Supplementary Fig. 6). Again, satellite data had the greatest contribution to accuracy in all ML models. Models constructed exclusively using satellite data predicted wheat yield variation with R^2^=0.71 accuracy (Supplementary Fig. 6), exceeding the accuracy of metadata-only (R^2^=0.59), management-only (R^2^=0.37), and weather station data only models (R^2^=0.34; Supplementary Fig. 6; Supplementary Code 1). While valuable, interrogating model accuracy in this way was fundamentally constrained: assessing the interactive contributions of smaller variable groupings and individual variables was not combinatorially limited.

Further interrogation of model dynamics, using ML algorithm heuristics that rank features by their importance, revealed the nominal predictive value of individual variables (Fig. 5). These feature detection and scoring methods showed limited concordance (Fig. 5a-c), with some differences likely arising from the diverse scoring methods employed (Supplementary Code 1). However, some variables, such as accumulated rainfall, attained high importance ranks across all models (blue points; Fig. 5d). Across PLSR, RPRM and BCRF models, consistently high importance ranks were assigned to cumulative rainfall, latent heat flux (an indicator of transpiration rates^21^ and stomatal^22^ conductance across a canopy), application rates of Sulphur, and application of the pesticide Clethodim (Fig. 5a-c).

However, variable importance scores lack an indication of the direction of effect, and any indication of whether orthogonal or interactive effects underpin the predictive value of ‘important’ variables. As such, ML models were interrogated to generate interpretable outputs.

Decision trees generated by RPRMs include the direction of observed effects, and reveal possible variable interactions through hierarchical dependencies within decision trees (Fig. S7-S8; Supplementary Data 1). For example, the yield-predictive RPRM in Supplementary Data 1 reveals that, in TOS-only and full-season prediction models, lower soil sodicity, higher soil carbon, phosphorous, and nitrogen, and higher doses of total applied of nitrogen and phosphorous were uniformly predictive of higher yield. Other interactions were context dependent: for example, higher rainfall was predictive of higher yield in eight of the fourteen decision points in the full-season RPRM model above (Data S1). Species-targeted RRPM decision trees also provided insight onto the full-season (Supplementary Fig. 7) and pre-sowing predictors of yield (Supplementary Fig. 8; Supplementary Data 1). Decision points in these trees included well-known agronomic interactions, such as gains in wheat yield from the pre-sowing application of nitrogen fertilisers or monoammonium phosphate (Supplementary Fig. 8), as well as previously unknown interactions, for example the discrimination between lower-yielding populations through satellite reflectance bands and latent heat flux (leftmost branches; Supplementary Fig. 7).

Visual inspection of tree-based models does not work at scale, for example when generating a forest of thousands of decision trees using BCRFs. We overcame this problem by reducing BCRFs to their most common and predictively robust decision sub-trees (the most common sequences of decisions within a random forest of decision trees) using the inTrees analytical approach of Deng^13^. This approach revealed complex yield-predictive dependencies between remote sensing, environmental, and management inputs (Table 3-4). For example, the most common predictively robust decision sub-tree in canola constitutes a complex cross-domain dependency (Table 3) where the effect of the normalised differenced vegetation index on canola yield depends, respectively, on the MODIS band 7 reflectance exceeding 0.10, and the total accumulated rainfall to 150 DAS falling below 310mm. Similar patterns were revealed in wheat: for example, a high enhanced vegetation index at 160 DAS and a high maximum enhanced vegetation index at 150 DAS predicted a high 5.5 t/Ha yield. Low-yield prediction rules included indicators of vegetation stress, such as the prediction of low (1.1t/ha) yields from a combined low fraction of photosynthetically absorbed radiation, low normalised differenced vegetation index, and a high middle infrared reflectance value; or the low yields (0.8 t/Ha) predicted by a combined high MODIS reflectance band 7 value and low observed latent heat flux (an indicator of the extent of stomatal closure, leaf hydraulics, and crop evapotranspiration^21–23^) during the growing season (Table 4).

**Table 4.** Variable interactions predicting wheat yield.

| Conditional Rule Chain | Frequency<br>in Forest<br>(%) | Rule<br>Error<br>rate | Rescaled<br>Importance<br>Score | Rule-Predicted<br>Yield (t/Ha) |
| --- | --- | --- | --- | --- |
| Mean Night-time LST 40DAS <= 27.33 &<br>Evapotranspiration Var. 90DAS <= 1.39e+08 &<br>Min Reflectance Band 6 130DAS > 0.23 &<br>MIR 190DAS > 56.88 | 0.19 | 0.56 | 1.00 | 1.70 |
| FPAR 120DAS > 43.93 &<br>Max Reflectance Band 6 140DAS > 0.21 &<br>Max FPAR 170DAS <= 0.42 | 0.56 | 1.20 | 0.59 | 2.58 |
| Max FPAR 100DAS <= 0.60 &<br>Min NDVI 130DAS <= 0.41 &<br>Mean MIR 180DAS > 0.32 | 0.05 | 0.23 | 0.30 | 1.12 |
| Mean EVI 160DAS > 0.28 &<br>Max NIR 170DAS > 0.34 | 0.14 | 2.49 | 0.27 | 5.25 |
| Emissivity Band 32 Var. 70DAS <= 1.089e-06 &<br>Mean Reflectance Band 6 140DAS > 0.21 &<br>Max MIR 160DAS > 0.21 &<br>GPP 190DAS <= 5.80 | 0.34 | 0.89 | 0.27 | 2.18 |
| Mean Reflectance Band 4 80DBS > 0.096 &<br>Max MIR 130DAS > 0.29 | 0.04 | 0.21 | 0.25 | 1.00 |
| Mean Reflectance Band 7 140DAS > 0.32 &<br>LHF 190DAS <= 3.75e+08 | 0.02 | 0.14 | 0.21 | 0.79 |
| Reflectance Band 6 10DBS > 0.27 &<br>Max Net Photosynthesis 150DAS <= 0.036 &<br>GPP 160DAS <= 4.72 &<br>Min MIR 170DAS <= 0.192 &<br>Max GPP 180DAS <= 0.028 | 0.04 | 0.57 | 0.18 | 2.98 |
| Max Reflectance Band 7 30DAS > 0.14 &<br>Mean Net Photosynthesis 60DAS <= 0.014 &<br>Reflectance Band 6 120DAS > 54.81 &<br>Max NDVI 150DAS > 0.53 &<br>Max Net Photosynthesis 180DAS <= 0.029 | 0.11 | 0.96 | 0.14 | 3.42 |
| Max LAI 160 DAS <= 1.28 &<br>Mean MIR 160 DAS <= 0.19 &<br>Max Net Photosynthesis 170 DAS <= 0.029 | 0.14 | 1.09 | 0.06 | 3.52 |
| Mean Solar Radiation TOS > 12.64 &<br>LHF 80 DAS > 26341250 &<br>Mean NDVI 140 DAS <= 0.59 | 0.03 | 0.81 | 0.04 | 3.63 |
| CS Rainfall 170 DAS > 262.9 &<br>NDVI Variance 20 DAS <= 7.67e-07 &<br>Min Reflectance Band 7 120DAS > 0.12 | 0.01 | 0.65 | 0.04 | 3.69 |
| Max maximum temperature 180 DAS > 25.40 | 0.98 | 2.28 | 0.03 | 3.33 |
| Mean Red Reflectance 130 DAS > 0.11 &<br>Red Reflectance 140 DAS > 23.11 &<br>Net Photosynthesis 180 DAS > 8.63 | 0.06 | 0.47 | 0.02 | 2.09 |

Abbreviations as in Table 3.

In line with their high importance ranks across models and their over-representation in RPART decision trees, and despite each constituting only 2.2% of all predictor variables in the initial model, vegetation indices, gross primary productivity and latent heat flux were common across predictively valuable sub-trees (Table 3-4). For example, NDVI occurs in 13% (11/83) of all decisions in Table 3-4, yet constitutes only 2.2% of all input variables. Likewise, cumulative rainfall (0.04% of the initial predictor variables) is present in nine of the 26 prediction rules and constitutes 11% of the prediction rule chains: a 25-fold overrepresentation.

## Discussion

Foundational crops such as wheat, canola, and oats provide a substantial fraction of total human caloric intake^1^. However, traits underpinning the yield of these species are driven by poorly-understood complex interactions. In particular, environments are largely characterised using ground-station data and, increasingly, highly complex but variable-coverage drone data. These approaches overlook the opportunity of satellites as consistent, regularly-timed, instantaneous, and global instruments for capturing environmental variation.

In this study, low-resolution satellite data is used to characterise environmental diversity across sites, providing several novel insights into crop yield. Even these crude whole-site measures added substantial predictive capacity to our models (Supplementary Fig. 1; Supplementary Fig. 7-8). In particular, our findings suggest the importance of latent heat flux, a proxy for both water availability^21^, stomatal conductance^21,22^, and canopy transpiration rates^22^, as a predictor of yield variation across sites.

These findings suggest the remarkable potential for both existing low-resolution and developing higher resolution satellite resources for agriculture and plant breeding. In contrast with the 250m-1000m pixel daily resolution data used here, sub-40cm resolution data will be available multiple times daily from multiple providers as soon as late 2021. This resolution is sufficient to resolve, for example, individual cotton and canola plants every clear day throughout a field season. Capturing environmental patterns at this spatial and temporal scale, along with the potential for direct satellite observation of key agronomic traits such as growth patterns and phenology^24^, has the capacity to dramatically alter the conduct of large-scale plant breeding trials. If these data can be meaningfully integrated into plant breeding models and used to inform plant biology, high-resolution satellite data has a substantial future in agronomy.

Integration of satellite data, and drone and large ‘omics data, into plant breeding models has largely been constrained by statistical barriers. It is generally easier to generate ‘big’ data, and saturate plant populations with variables, than provide a meaningful analysis of such data. That is not to say such success is not possible: for example, large ‘omics data sets may be reduced to indices or trait values without substantial information loss^25^. However, this dimension reduction approach is not always appropriate and may reveal little or nothing about the interactions between variables. Integrating data into ML pipelines with comprehensible model mechanics, and carefully interpreting the results, provides a pathway to solve these problems and meaningfully integrate advances in data generation with plant breeding platforms^25^.

A major advantage of ML models is the capacity to generate a one-shot model that captures interactions across multiple domains, such as the management-environment interactions shown in Supplementary Fig. 7-8. While our focus has been the capture of environmental interactions, our findings highlight the potential of ML to capture interactions across further data layers: metabolomic, transcriptomic, proteomic and large-scale phenomic data are all suitable for inclusion into a unified ML model.

While the promise of ML models is considerable, gains in predictive accuracy are limited by fundamental challenges. For example, concurrent changes in model accuracy across diverse models and species are unlikely to be a result of overfitting (Fig. 4). Rather, these patterns may arise due to fundamental model limitations when extrapolating in complex systems^26–28^. For example, while ML models generalise well across observed data, they suffer direct constraints on predictive accuracy under pattern-extrapolation. This is epitomised by the ‘checkerboard problem’, where ML models fail to extrapolate the alternating pattern of black and white squares from a smaller to a larger checkerboard^27^. The failure of ML models to extrapolate phenotypes to future years, across a stochastic ‘checkerboard’ of switching between El Niño, neutral, and La Niña climates, may therefore represent a failure of ML algorithms to extrapolate complex patterns.

Likewise, non-stationarity, a shift in mean and variance over time, places fundamental limits on the accuracy of all statistical models^28^. Cross-model changes in accuracy, when projecting new fields and years (Fig. 4), may reflect the independent or combined role of non-stationarity of management practices, soil diversity, hidden genetic diversity over time, or the role of climatic or environmental non-stationarity (Fig 4.; Supplementary Fig. 9) over time.

As such, shifts in predictability of crop behaviour across the NVTs raises important questions on systems dynamics. If non-stationarity in the Australian climate drives collapses in the predictability of a complex system, crop behaviour and yield, within the space of a single year (Fig. 4), this has important ramifications for agronomy under climate change. Climates are becoming increasingly nonstationary^29^, including Australian grain-growing regions^30–32^, causing a dramatic global loss in the predictability of rainfall patterns^29^ and temperatures^33,34^.

Non-stationary rainfall patterns are particularly concerning. Already, a quarter^35^ of the global land surface area has non-stationary rainfall patterns, and this fraction is rising^35^. This increasing climate non-stationarity causes crop breeding targets to become intrinsically less predictable, regardless of advances in models or methodology. Crop breeding pipelines from initial plant crosses to the release of new varieties require development times of around ten years^36^. If the degree of climatic non-stationarity increases within this horizon, and future climatic patterns become less predictable as targets for plant breeding, climate instability may pose a serious challenge to food production systems^37^ given fundamental bounds on model predictability^28^.

There remains considerable cause for optimism from ML algorithms beyond these limitations. As we have shown, ML models can learn and recall the growing season of millions of plants, integrate these data into meaningful models, and accurately forecast phenotypic variation. Furthermore, the promise of ‘big data’ and ML in agriculture is not constrained to gains in prediction accuracy. While improved accuracy from ‘black box’ models has enormous utility for share trading or insurance, such models often have more limited scientific and in-field applications. For the greatest utility for farmers and biologists ML models need to be made accessible and understandable, even at the cost of predictive accuracy.

Predictions from neural networks or deep learning models are often more accurate yet, with some exceptions^38,39^, are not comprehensible by examination of model dynamics. Even when black box models produce variable importance rankings (*e*.*g*. Fig. 5) it is unclear whether high-ranked predictive variables interact with one another or act independently, or in what context they are most important, when generating a predictive outcome. In contrast, approaches such as RPRMs and BCRFs allow the understanding and interrogation of internal model dynamics: it is possible to ask *why* an algorithm generated a specific prediction.

For example, trusting Cauer’s conceptual “black box” to evaluate our models produced may have resulted in a surprising conclusion: that use of herbicides routinely has an inhibitory effect on yield. For example, in the model shown in Supplementary Fig. 7, reduced yield was predicted by the previous application of common herbicides such as Roundup (1.3t/Ha yield loss) and Cadence (1.8t/Ha yield loss). However, the conclusion that herbicides are yield-inhibitory is likely incorrect. Herbicide application is confounded with pathogens, environmental stress, and degraded soils: as such, herbicide use may predict lower yield because herbicide use is more likely under worse growing conditions. Discrimination between these cases depends on careful analysis of model mechanics, a process that is not possible within true ‘black box’ models.

Likewise, across the NVTs zero-yield and extremely low-yield trials are not missing at random, but have been actively removed. This violation of the ‘missing completely at random’ criteria^40,41^ has counterintuitive effects that are independent of the applied predictive model. For example, increasing frost severity is predictive of increasing crop yield across trials (Supplementary Fig. 10). This is not necessarily a result of severe frosts causing better crop yield but, more likely, because the removal of low- and zero-yield trials has generated a non-random survival bias: better-conditioned, higher-yield crops are most likely to survive a severe frost than crops in stressed and suboptimal environments. As a result, higher yields may be positively correlated with a worse environment purely as a statistical artefact. Black box models are not panacea against such subtle issues.

Interpretation of machine learning models should not, therefore, be reliant on the simple scoring of variable importance. Rather, our results suggest that detailed assessment of the internal mechanics of machine learning models is a key analytical challenge for biologists who seek to understand, rather than simply predict, biological systems.

## Methods

Compilation of the NVT data resulted in the location and measurement of 266,033 variety-trial combinations, aggregated from 780,569 field trial plots in 6,547 successful (or non-failed) field trials. Linked to these data were over eight thousand variables, including approximately ten thousand field-years of soil samples and hundreds of thousands of chemical and fertiliser doses. A total of seventy phenotypic traits were available, for over one and a half million unique phenotypic measurements^6^. Of these, seven agronomically important traits, of sufficient data quality and sample size, were selected to train machine learning algorithms: grain protein percentage, days to 50% flowering, percentage Glucosinolate oil content, Hectolitre weight, thousand grain weight, the fraction of grain sieving below 2.0mm, and yield (Table 1; Supplementary Fig. 3-4).

Data used to train PLSRs, XGBMs, and LSVMs were dummy-coded for factors, and transformed to zero mean and unit variance for numeric data (see database descriptor for further details^6^). However, tree-based algorithms partition the input space based on the non-transformed target variable, and do not require rescaling to avoid model fit biases. As such, to preserve the direction and magnitude of effects and facilitate interpretable models, non-transformed data were used to train RPRMs, BCRF, and xvBCRF models.

Missing data were concentrated into missing metadata and field trial comments measured in small subsets of trials, such as disease scores and animal damage scores. Only 0.05% of all satellite data were missing, generally as a result of a pixel failing the quality control screening of a NASA data product algorithm. Broader environmental data, including ground sensor arrays and weather station data, were 0.7% missing. All missing data were imputed using two approaches, described in further detail in the associated Database descriptor^6^, for both untransformed and unit-variance, zero-mean transformed data. The variance in model accuracy arising from imputation noise and error was evaluated by applying machine learning models to all imputed datasets, predicting site-mean yield, and measuring model accuracy post-imputation (Supplementary Fig. 11; Supplementary Code 1).

Models were trained using, as often as appropriate, default tuning and input parameters. Hyperparameters for model training were subjected to minimal training and optimisation, with parameters given in Supplementary Table 4. All models were subjected to 10-fold internal cross-validation, with identical training and target data across models, to allow direct comparisons of accuracy.

### Training targets

Yield models were initially trained to predict two holdout samples: a ‘holdout trial’ set of 100 field trials randomly sampled from the years 2008-2017, and an ‘annual forecast’ sample consisting of all data from 2018 (Fig. 2). Models were trained on data from 2008-2017 excluding the random holdout trials, and used to predict both the holdout trials and the 2018 data. For rolling forecast models presented in Fig. 4, no holdout trial sample was selected: instead, models were trained on all data before each successive cut-off year, and tested on the next year’s data.

In-season rolling forecasts were developed using all data available before a given time, relative to sowing, and using these data to predict end-of-season yield in Canola and Wheat only. The three most functionally distinct methods were used: tree-based Naïve BCRFs trained in ranger, kernel-based LSVMs, and principal components regression-based PLSRs. As in previous models, PLSR models were trained using 10-fold crossvalidation, in 10 segments, using default loss criteria (Supplementary Code 2). Due to the greater sample size constrains, these variety-specific models were subjected to random holdout trial prediction only (Fig. 2).

### Algorithms and hyperparameters

Hyperparameters used in model training tasks are given in Supplementary Table 4 and Supplementary Code 1 and 2.

Random forest models were constructed using three different approaches: naïve Breiman-cutler random forests (BCRFs) trained using the Ranger implementation^42^, BCRFs cross-validated by calendar year such that test data was never in the same year as training data^15^ (xvBCRF), and extreme gradient-boosted forest models (XGBMs). To preserve the magnitude of changes to leaf node averages caused by decision points (*e*.*g*. Supplementary Fig. 7-8), and the heuristics derived from these decisions (Table 3-4), RPRMs, BCRFs, and xvBCRFs were trained using non-scaled data. Unlike other machine learning methods, scaling does not impact the accuracy of these tree-based partitioning methods.

Extreme gradient boosting machines (XGBMs), Linear support vector machines^19^ (LSVMs) and Partial Least Squared Regressions^20^ (PLSRs) were trained using data rescaled to zero mean and unit variance, with factors excluded (*e*.*g*. unstructured crop comment fields, unique trial identifiers) or dummy-coded (*e*.*g*. varieties observed in over N > 1000 plots, breeder names, experimental series, trial operators, trial comments common to > 1000 trials, crop species; see Supplementary Code 1; Supplementary Code 2).

The LSVM regression models were constructed using a grid-based stepwise search using fixed gamma and lambda values defined in Supplementary Table 4. Each LSVM was subject to 10-fold internal cross-validation, tuned over a 10×10 hyperparameter grid, using the “liquidSVM” package^19^.

### Importance and heuristic rule reduction

Variable importance scores were returned using default heuristics from the RPRM and BCRF models, and variable importance was approximated in PLSR models by dividing coefficients by the sum of the absolute value of the coefficient matrix (Fig. 4; Supplementary Code 1). To reveal common interactions predictive of phenotypic variation, xvBCRF models were subjected to the analytical pipeline described in Deng^13^. This approach aggregates the frequency of all sub-trees of decisions within random forests, to reveal the most common predictive decision sequences or paths (Supplementary Code 2). This set of decision paths is then pruned by treating common decision paths as features, re-training a regularised random forest using these and all previous features, and ranking the importance these decision paths in the subsequent model. By treating this interaction as a feature, the importance of complex dependencies between variables may be explicitly stated, in a way that is impossible with black-box models. Therefore, all predictively valuable decision paths were captured for all cross-species and species-specific xvBCRF yield models (Supplementary Code 2).

All data are available in the linked Database descriptor^6^, from the associated figshare repository, or on request from the corresponding author. All code and secondary data generated by this analysis, such as models and decision trees, are available in the supplementary data and from the corresponding author.

## Acknowledgements

The authors acknowledge funding from the Australian Research Council Centre of Excellence for Translational Photosynthesis (CE140100015). We wish to acknowledge the hard work of the many researchers and agronomists who collected the historical agronomic data for the Grains Research and Development Corporation National Variety Trials used in the work described here.

## Author contributions

SJN conceived and designed the study, wrote the code, performed the analysis, designed and plotted the figures, and co-wrote the manuscript. RF co-wrote the manuscript and contributed to the experiment design.

## Competing Financial Interests

The authors declare no competing financial interests.

All data and code are available from the supplementary materials the linked Database Descriptor publication uploaded to *Scientific Data* and the figshare repository^6^.

