## Supplementary Tables and Figures for "Explainable machine learning models of major crop traits from satellite-monitored continent-wide field trial data"

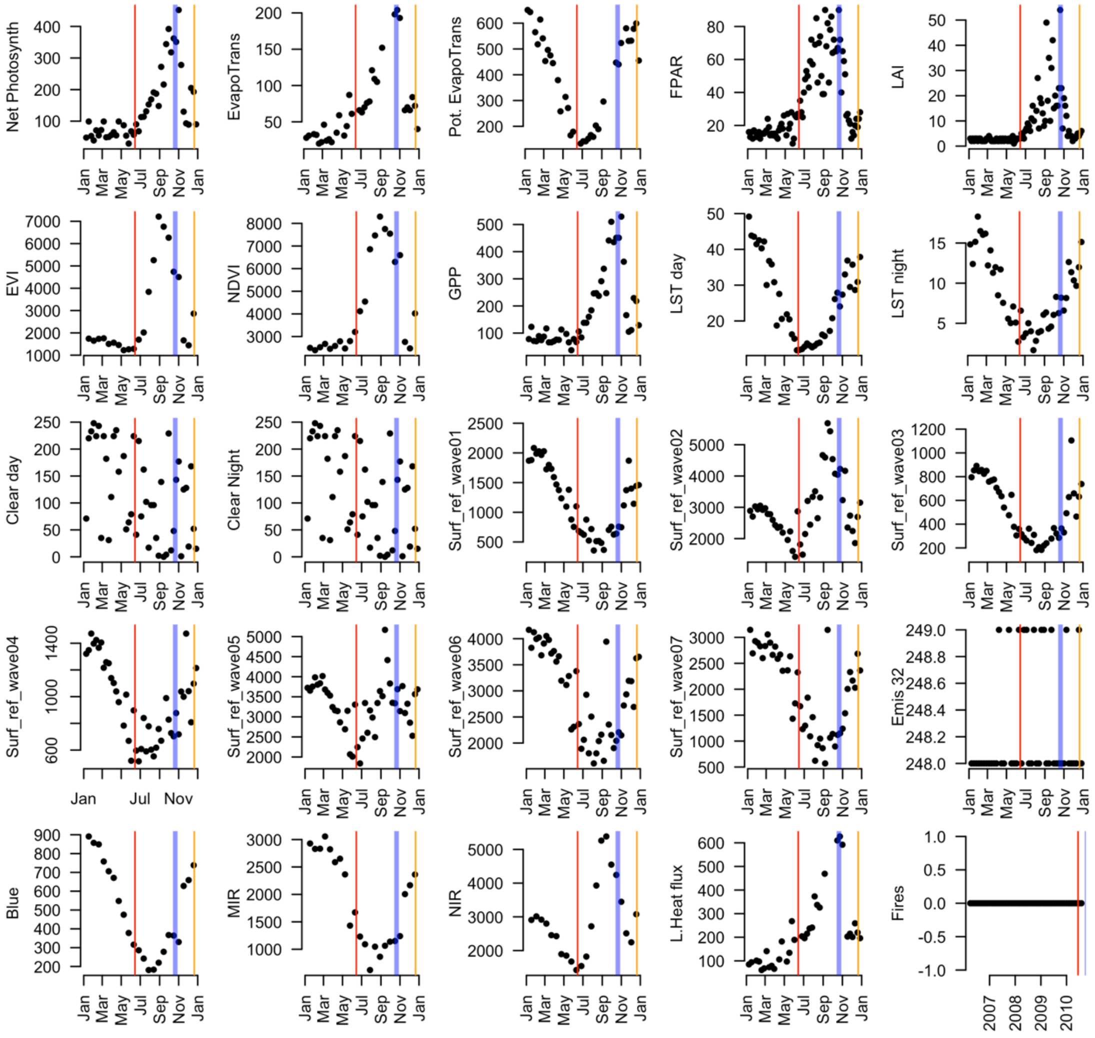

**Figure S1. Environmental and vegetation patterns captured by remote sensing.** Remote sensing arrays capture seasonal patterns across a season, from sowing (red lines) through anthesis or flowering (blue lines), to harvest (orange lines), in diverse variables such as photosynthesis and leaf area (top row), vegetation indices and land surface temperatures (second row), cloud cover and raw reflectance values (third to fifth rows), infrared bands, latent heat flux, and fire events (fifth row). Variation shown for a single site (Turretfield 2010; Main Season planting), abbreviations as in Supplementary Table 1.

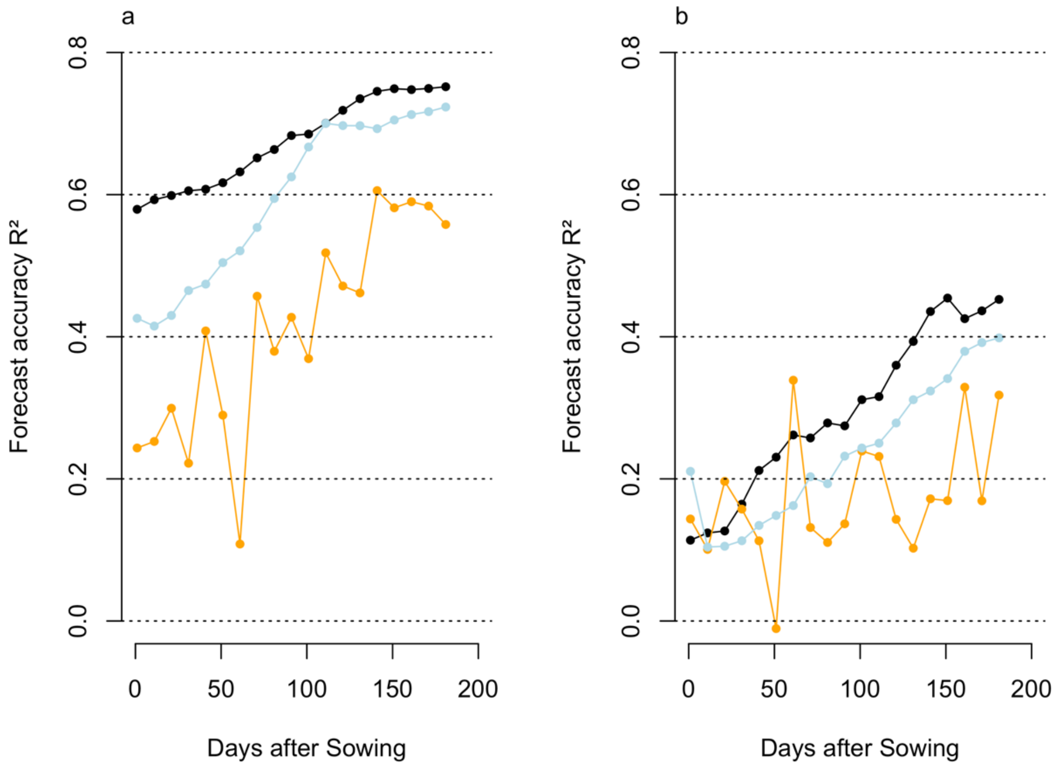


**Figure S2. Learning curves for models constructed using progressive seasonal data.** Models constructed to predict yield from a given point in the season relative to sowing (x-axis), in **a** wheat and **b** canola, trained on 2008-2017 data and used to predict 2018 season data. As the season progresses, later-season data is included and models improve in forecast accuracy (y-axes) with xvBCRF models (black) outperforming PLSR models (blue) and LSVMs (orange) over almost all forecast horizons.

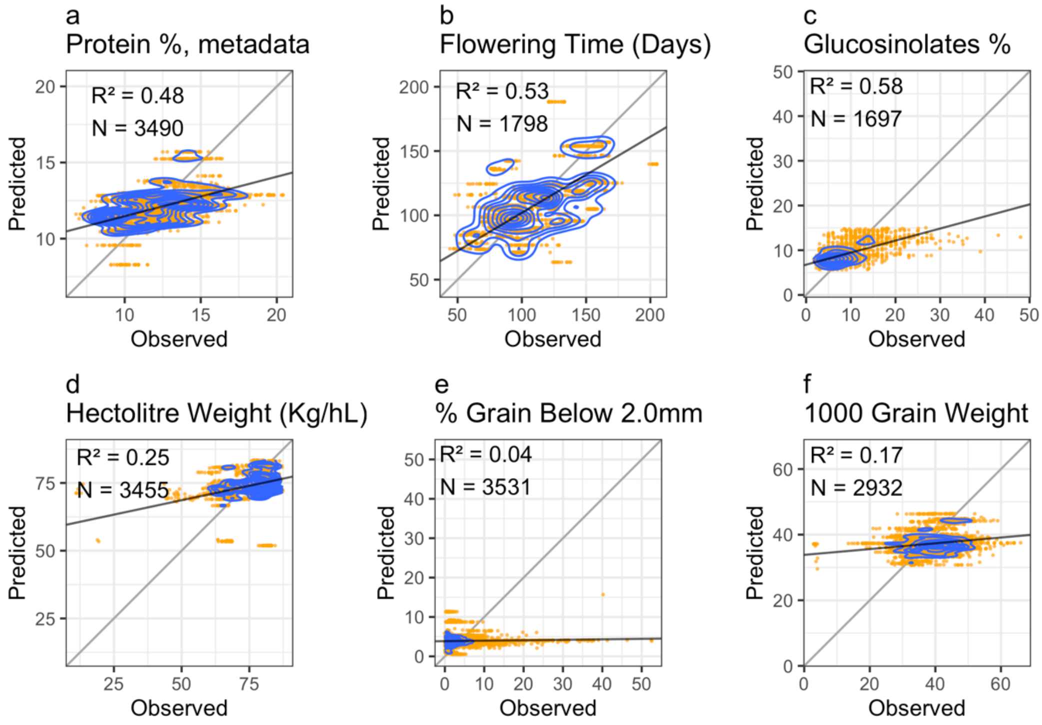


**Figure S3. Prediction accuracy of BCRF models in diverse phenotypes under random holdout trial prediction.** Using complete season data from 90 days before to 270 days after sowing to train BCRF models, tested in 100 random holdout sites, allows the prediction of key agronomic traits such as: **a** grain protein content, **b** time to 50% flowering, and **c** Glucosinolate oil content (Canola only). However, BCRF models trained under the same criteria were of limited or no use for predicting grain size or volume metrics such as: **d** Hectolitre weight, **e** the fraction of grain below 2.0mm and **f** thousand-grain weight. Numbers N indicate sample sizes across predicted holdout sites.

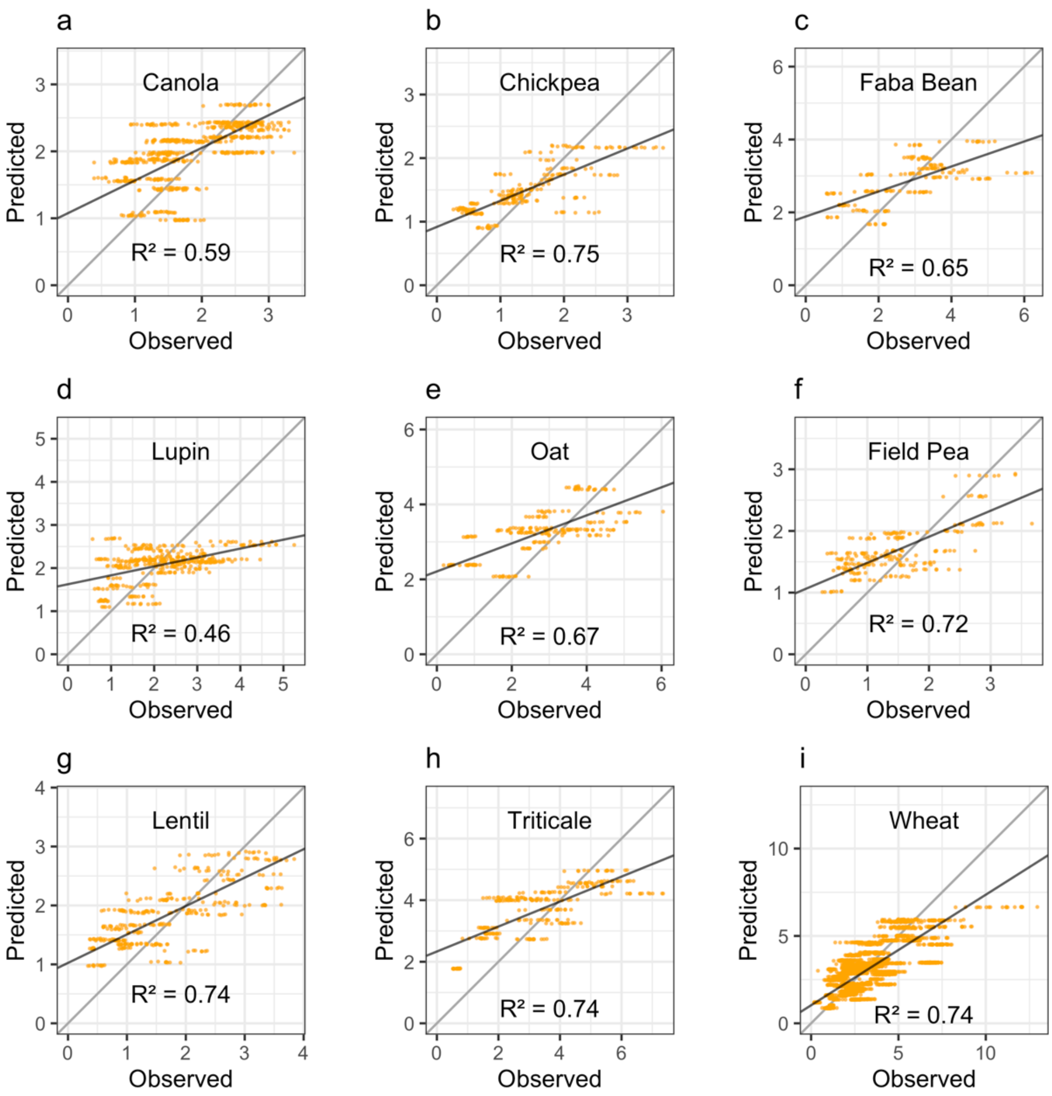


**Figure S4. Prediction accuracy of annual forecast BCRF models trained using species-specific data.** Prediction accuracy of models trained using data to 200DAS from within species, rather than cross-species models, provide good accuracy for projecting yield in **a** Canola (N=608), **b** Chickpeas (N=243), **c** Faba Beans (N=188), **d** Lupins (N=403), **e** Oats (N=216), **f** Field Peas (N=230), **g** Lentils (N=331), **h** Triticale (N=266), and **i** Wheat (N=2296). Test data are for the latest observed year containing successful trials in each species.

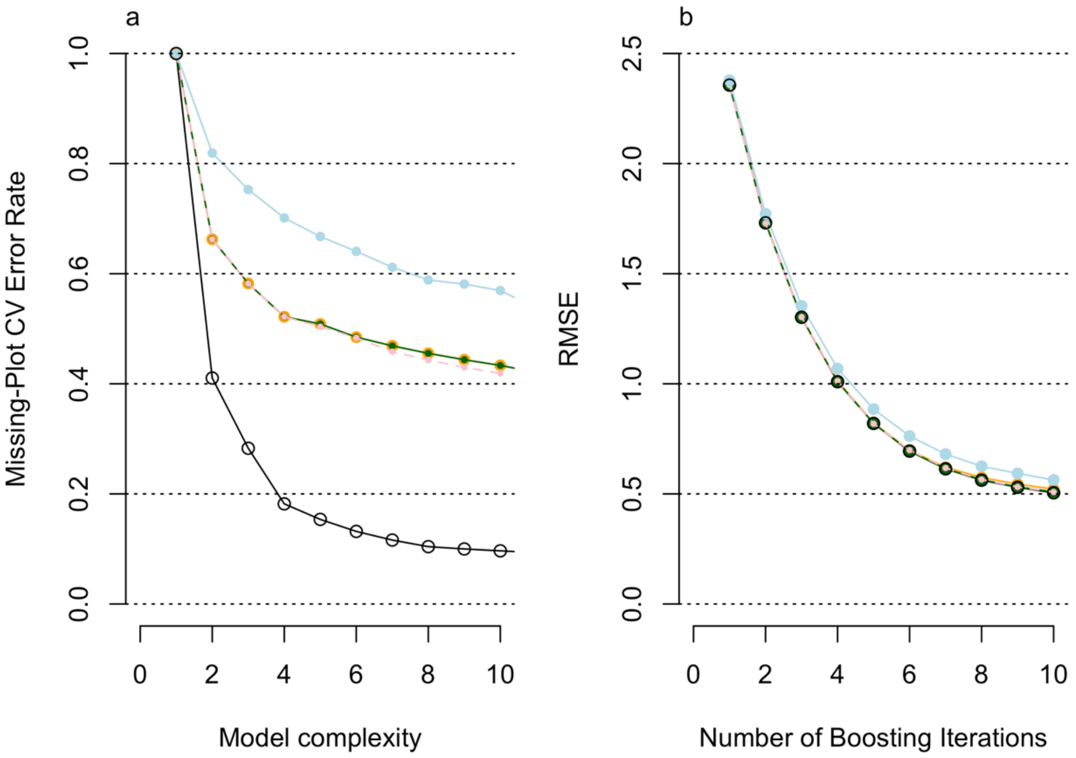


**Figure S5. Model complexity and accuracy excluding individual domains of input data.** Predictive accuracy within models, such as the **a** RPRM and **b** extreme gradient boosting models shown, was reduced by different rates under lave-one-out testing. Across all levels of model complexity (x-axis), removal of satellite data (blue) produced the greatest reduction in accuracy relative to models trained using all available data (black circles). Exclusion of weather station data (orange), metadata (pink), and management data (green) imparted similar and more limited costs in accuracy. Cross-validation error rates (y-axis, **a**) is the inverse of $R^2$ values and RMSE (y-axis, **b**) is the root mean squared error for random holdout observations.

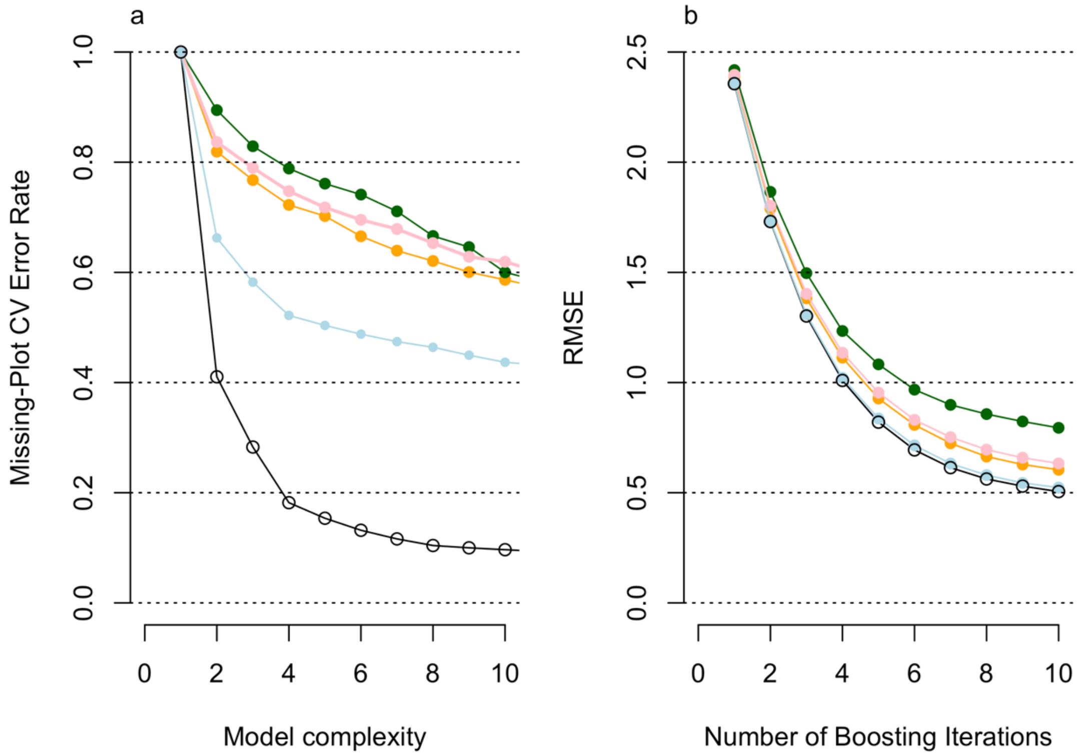


**Figure S6. Model complexity and accuracy using single domains of input data.** Predictive accuracy within models, such as the **a** RPRM and **b** extreme gradient boosting models shown, was driven by inclusion of certain domains. Relative to models trained using all available data (black circles), satellite data alone (blue) produced models of greater accuracy than weather station data alone (orange), metadata alone (pink) and management data alone (green) across all levels of model complexity (x-axis). Cross-validation error rates (y-axis, **a**) is the inverse of $R^2$ values, RMSE (y-axis, **b**) is the root mean squared error for random holdout observations.

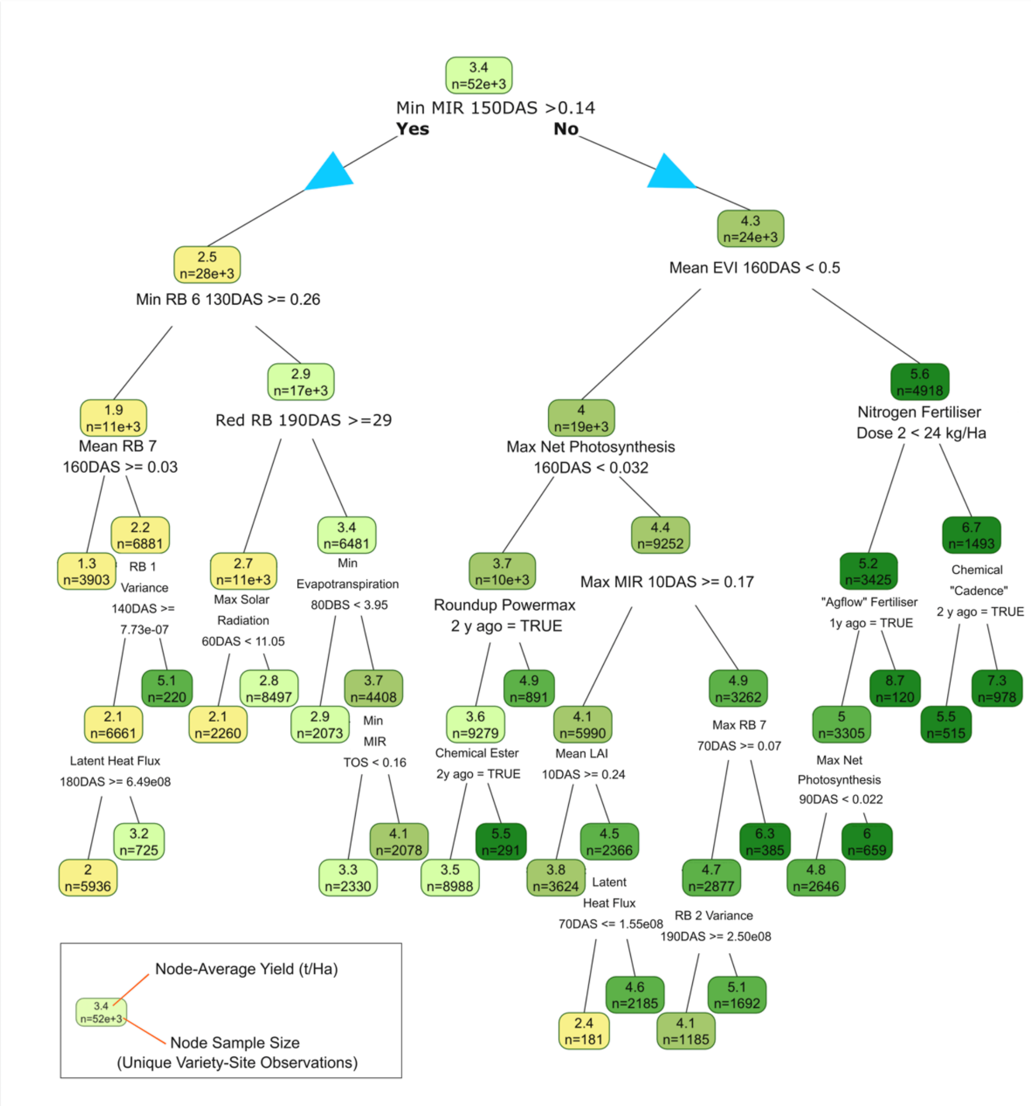


**Figure S7. Pruned decision tree for wheat yield.** Pruned decision tree predicting yield variation (green colour scale, t/Ha shown in boxes) from all input data to 200DAS, illustrating the readable heuristics generated by tree models. Model accuracy under annual forecast prediction is $R^2 = 0.79$; RMSE = 1.13. Days after Sowing (DAS), Time of Sowing (TOS), Days before sowing (DBS), Middle Infrared (MIR), Enhanced Vegetation Index (EVI), Normalised Differenced Vegetation Index (NDVI), Leaf Area Index (LAI), MODIS Reflection Band (RB), all values rounded to 2 significant digits.

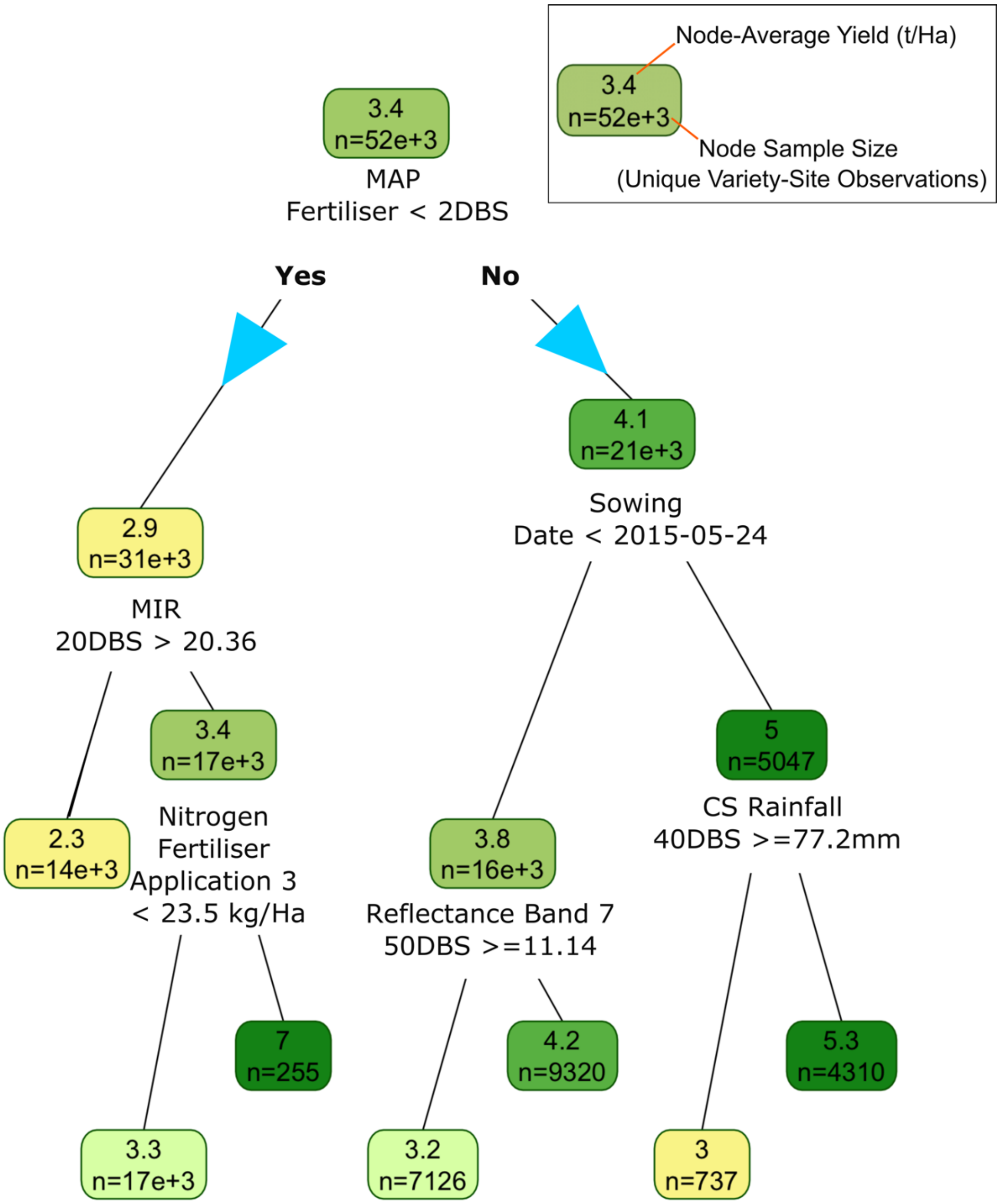


**Figure S8. Pruned decision tree for wheat yield trained on data to time of sowing.** A pruned decision tree for wheat yield, constructed using data available at the time of sowing, illustrating the readable output of decision tree models (annual forecast accuracy $R^2 = 0.3$). Cumulative sum (CS), Monoammonium Phosphate (MAP), MODIS middle infrared reflectance band (MIR).

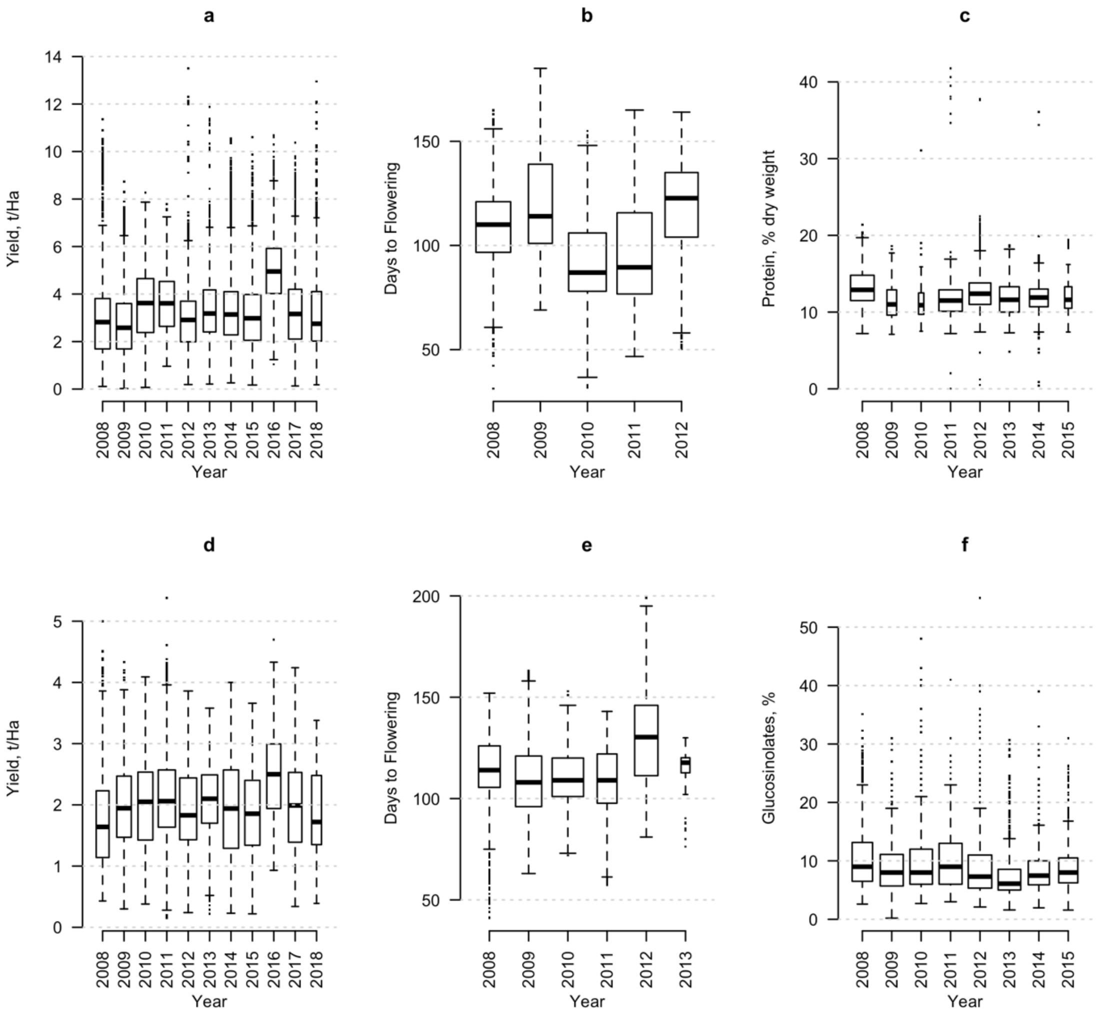


**Figure S9. Annual variation in key agronomic traits.** Annual variation in key agronomic traits, as shown here by shifts in mean and variance of wheat yield **a** (N = 54658 variety-year observations), flowering time **b** (N = 4093), and grain protein content **c** (N = 16681); and Canola yield **d** (N = 14929), flowering time **e** (N = 2841), and Glucosinolate content **f** (N = 9585).

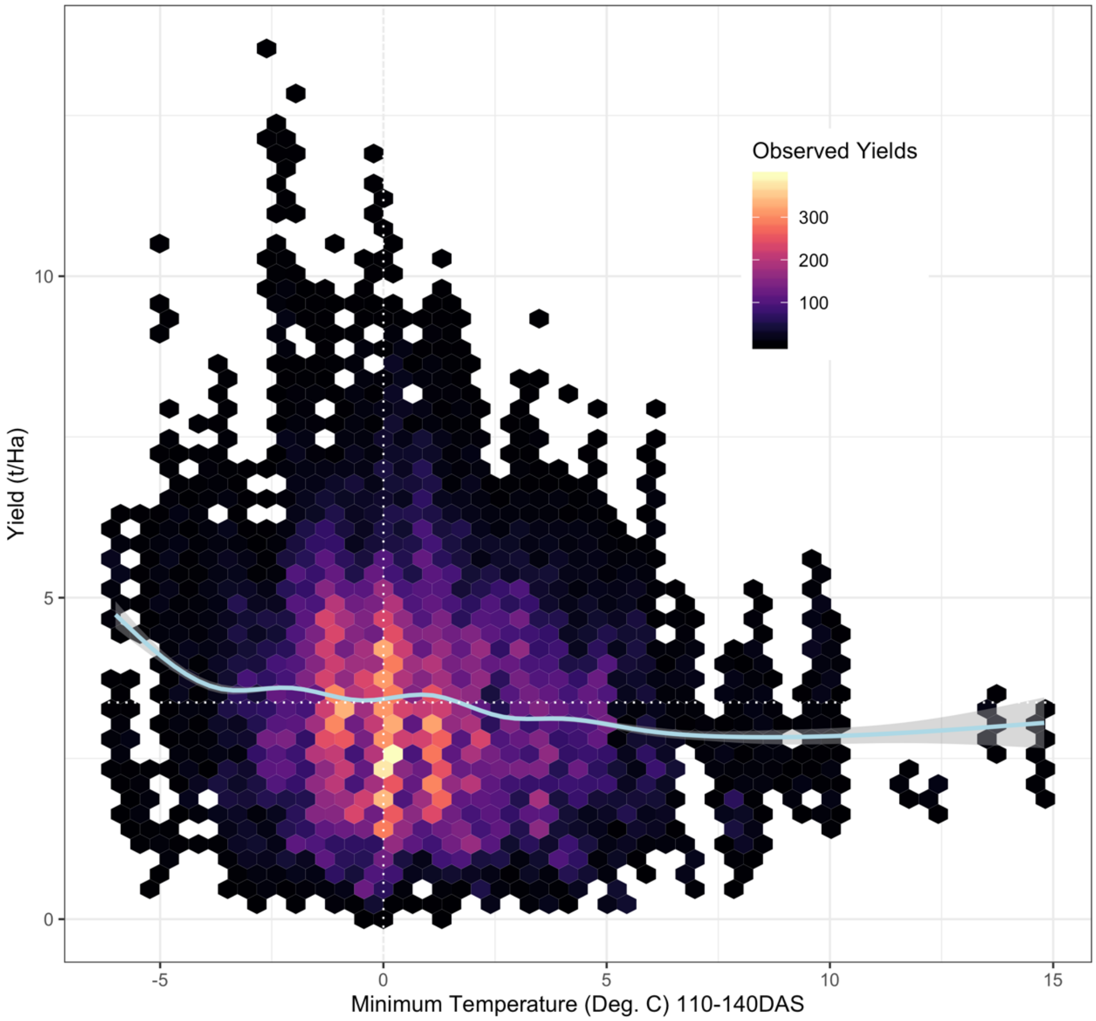


**Figure S10. Flowering-window minimum temperatures and yield.** Average wheat yields (y-axis) observed for a given variety-site combination exhibit a counter-intuitive relationship with frost severity during flowering (here defined as 110-140 Days after Sowing, x-axis): more severe frosts predicting higher yields (blue curve; locally weighted smoothed spline; N= 54,658). This may reflect a survival bias resulting from the absence of reporting zero-yielding trials, where only high-yielding crops survive extreme low temperatures.

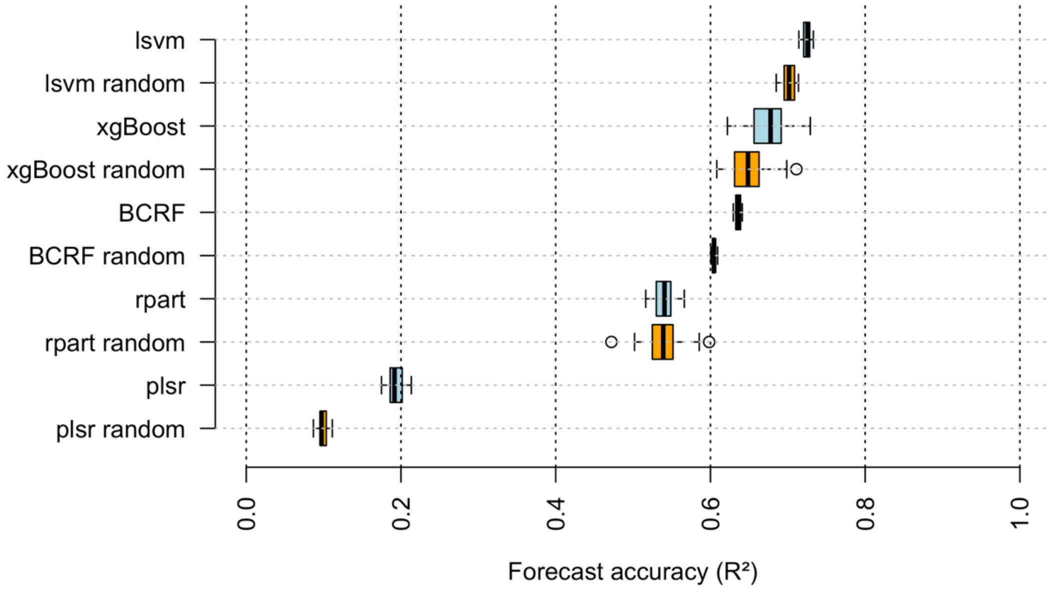


**Figure S11. Distribution of model accuracy from imputation error and noise.** There were substantial differences in the sensitivity of model predictions to imputation noise under both random-sample imputation (orange) and random forest imputation (blue) under forward-projection accuracy. Models with the lowest sensitivity to imputation-based variability were stratified sampling BCRFs and linear support vector machines (LSVMs).

**Supplementary Table 1. NASA data products captured at NVT sites 2009-2018**

| Description (abbreviation) | Product (Product Code) | Spatial resolution | Temporal resolution | Unit |
|---|---|---|---|---|
| Daytime Land Surface Temperature (LST) | LST_Day_1km (MOD11A1.006) | 1000m | Daily | Degrees K |
| Night-time Land Surface Temperature (LST) | LST_Night_1km (MOD11A1.006) | 1000m | Daily | Degrees K |
| Day Cloud Cover | Clear_day_cov (MOD11A1.006) | 1000m | Daily | Integer |
| Night Cloud Cover | Clear_night_cov (MOD11A1.006) | 1000m | Daily | Integer |
| Thermal emissivity band 31 (Emis31) | Emis_31 (MOD11A1.006) | 1000m | Daily | Integer |
| Thermal emissivity band 32 (Emis32) | Emis_32 (MOD11A1.006) | 1000m | Daily | Integer |
| Black Sky Albedo Bands 1-3 (BSA) | Albedo_BSA_I1 (VNP43IA3.001) | 500m | Daily | Integer |
| Fraction of Photosynthetically Absorbed Radiation (FPAR) | Fpar_500m (MCD15A3H.006) | 500m | 4-day | Percent |
| Leaf Area Index (LAI) | Lai_500m (MCD15A3H.006) | 500m | 4-day | $m^2/m^2$ |
| Snow / Ice events | Snow Ice Detected (MCD15A3H_006) | 500m | 4-day | Logical |
| Atmospheric Aerosols | Aerosol Quality (MCD15A3H_006) | 500m | 4-day | Logical |
| Cirrus Cloud | Cirrus Clouds (MCD15A3H_006) | 500m | 4-day | Logical |
| Total Evapotranspiration (EvapoTrans) | ET_500m (MOD16A2.006) | 500m | 8-day | $kg/m^2/8day$ |
| Average latent heat flux | LE_500m (MOD16A2.006) | 500m | 8-day | $J/m^2/day$ |
| Potential Evapotranspiration (PET) | PET_500m (MOD16A2.006) | 500m | 8-day | $kg/m^2/8day$ |
| Average potential latent heat flux (PLE) | PLE_500m (MOD16A2.006) | 500m | 8-day | $J/m^2/day$ |
| Gross Primary Productivity (GPP) | Gpp_500m (MOD17A2H.006) | 500m | 8-day | kg*C/m^2 |
| Net photosynthesis | PsnNet_500m (MOD17A2H.006) | 500m | 8-day | kg*C/m^2 |
| Surface Reflectance Bands 1-7 | sur_refl_b01-sur_refl_b07 (MOD09A1.006) | 500m | 8-day | Integer |
| Enhanced Vegetation Index (EVI) | _250m_16_days_EVI (MOD13Q1.006) | 250m | 16-day | Integer |
| Middle Infrared Reflectance (MIR) | _250m_16_days_MIR_refl ectance (MOD13Q1.006) | 250m | 16-day | Integer |
| Normalised Differenced Vegetation Index (NDVI) | _250m_16_days_NDVI (MOD13Q1.006) | 250m | 16-day | Integer |
| Near Infrared Reflectance (NIR) | _250m_16_days_NIR_refl ectance (MOD13Q1.006) | 250m | 16-day | Integer |

**Supplementary Table 2. Reduced model accuracy when predicting the standard implementation of random-sample holdouts, versus random unobserved trials.**

Accuracy of predictions $R^2$, standard random-sample holdout

| | Models trained on all data at: | | |
|---|---|---|---|
| Model | TOS | 100DAS | 200DAS |
| RPRM | 0.92 | 0.92 | 0.93 |
| Naïve BCRF | 0.97 | 0.96 | 0.96 |
| xvBCRF | 0.95 | 0.95 | 0.95 |
| XGBM | 0.87 | 0.90 | 0.95 |
| LSVM | 0.99 | 0.99 | 0.99 |
| PLSR | 0.81 | 0.82 | 0.85 |

Reduction in accuracy, delta $R^2$ (% reduction), when predicting random holdout trial data

| | | | |
|---|---|---|---|
| RPRM | -0.32 (34.8) | -0.26 (28.3) | -0.17 (18.3) |
| Naïve BCRF | -0.21 (21.6) | -0.17 (17.7) | -0.14 (14.6) |
| xvBCRF | -0.15 (15.8) | -0.14 (14.7) | -0.11 (11.6) |
| XGBM | -0.23 (26.4) | -0.17 (18.9) | -0.17 (17.9) |
| LSVM | -0.35 (35.4) | -0.35 (35.4) | -0.31 (31.3) |
| PLSR | -0.23 (28.4) | -0.09 (11.0) | -0.09 (10.6) |

**Supplementary Table 3. Changes in accuracy of BCRF and LSVM models with targeted (single-species) versus omnibus (all-species) training data.**

**Training Data 200DAS**

| | | Model accuracy $R^2$ | | |
|---|---|---|---|---|
| Species | Test Sample | Targeted xvBCRF | Omnibus xvBCRF | Delta $R^2$ |
| Canola | 608 | 0.59 | 0.58 | 0.00 |
| Chickpea | 243 | 0.75 | 0.64 | -0.11 |
| Faba Bean | 188 | 0.65 | 0.49 | -0.16 |
| Oat | 216 | 0.67 | 0.75 | 0.08 |
| Field Pea | 230 | 0.72 | 0.60 | -0.12 |
| Wheat | 2296 | 0.74 | 0.73 | -0.01 |
| | | Targeted LSVM | Omnibus LSVM | |
| Canola | 608 | 0.55 | 0.59 | 0.04 |
| Chickpea | 243 | 0.52 | 0.44 | -0.08 |
| Faba Bean | 188 | 0.28 | 0.29 | 0.01 |
| Oat | 216 | 0.28 | 0.57 | 0.29 |
| Field Pea | 230 | 0.34 | 0.06 | -0.28 |
| Wheat | 2296 | 0.60 | 0.46 | -0.14 |

**Training Data 100DAS**

| | | Model accuracy $R^2$ | | |
|---|---|---|---|---|
| Species | Test Sample | Targeted xvBCRF | Omnibus xvBCRF | Delta $R^2$ |
| Canola | 608 | 0.55 | 0.60 | 0.06 |
| Chickpea | 243 | 0.62 | 0.65 | 0.02 |
| Faba Bean | 188 | 0.29 | 0.48 | 0.19 |
| Oat | 216 | 0.49 | 0.61 | 0.13 |
| Field Pea | 230 | 0.31 | 0.44 | 0.13 |
| Wheat | 2296 | 0.71 | 0.70 | -0.02 |
| | | Targeted LSVM | Omnibus LSVM | |
| Canola | 608 | 0.40 | 0.32 | -0.08 |
| Chickpea | 243 | 0.48 | 0.61 | 0.13 |
| Faba Bean | 188 | 0.19 | 0.12 | -0.08 |
| Oat | 216 | 0.23 | 0.03 | -0.20 |
| Field Pea | 230 | 0.31 | 0.33 | 0.02 |
| Wheat | 2296 | 0.38 | 0.43 | 0.05 |

**Supplementary Table 4. Machine learning model parameters by predictive target.**

| Model | Training data | Target data | Parameters |
|---|---|---|---|
| RPART | 2008-2017 Raw data | Holdout Trial and Forecast data | 10-fold crossvalidation, minbucket = 300, minsplit = 100, Complexity parameter = 0.0001 |
| Naïve BCRF | 2008-2017 Raw data | Holdout Trial and Forecast data | Model defaults, 10,000 trees |
| xvBCRF | 2008-2017 Raw data | Holdout Trial and Forecast data | 10-fold crossvalidation, minbucket = 300, minsplit = 100, mtry = 100, 1,001 trees, subsample size = 10,000, calendar year as cross-validation strata |
| XGBM | 2008-2017 Rescaled data | Holdout Trial and Forecast data | 10-fold crossvalidation, minbucket = 300, minsplit = 100, |
| LSVM | 2008-2017 Rescaled data | Holdout Trial and Forecast data | 10-fold crossvalidation, gamma steps = 20, lambda steps =20, min-max gamma = 0.05-20.0, min-max lambda = 0.00001-0.01 (grid defaults) |
| PLSR | 2008-2017 Rescaled data | Holdout Trial and Forecast data | 10-fold crossvalidation, ncomp threads = 10 |
| | | Rolling Annual Forecasts | |
| xvBCRF | All data before nth year | All nth year data | 10-fold crossvalidation, minbucket = 300, minsplit = 100, mtry = 100, 1,001 trees, subsample size = 10,000, calendar year as cross-validation strata |
| LSVM | All data before nth year, rescaled | All nth year data, rescaled | 10-fold crossvalidation, gamma steps = 20, lambda steps =20, min-max gamma = 0.05-20.0, min-max lambda = 0.00001-0.01 (grid defaults) |
| PLSR | All data before nth year, rescaled | All nth year data, rescaled | 10-fold crossvalidation, ncomp threads = 10 |
