## Supplementary data for "Explainable machine learning models of major crop traits from satellite-monitored continent-wide field trial data"

**Cross-Species ‘Omnibus’ Regression tree: Predicted at Time of Sowing**

**Variable names are defined in Newman & Furbank, *Scientific Data***

**Leaf values indicate node-average yield, in t/Ha**

**Model Accuracy: Missing Site Prediction R^2^ = 0.67; p<0.0001**

**n= 97333**

**node), split, n, deviance, yval**

*** denotes terminal node**

1) root 97333 2.139798e+05 3.0011290

2) METADom_Dicot_TF>=0.5 27709 2.238398e+04 2.0564560

4) METADom_sub_cropFaba.Bean< 0.5 25673 1.756516e+04 1.9874700

8) SatDom_GPP..20< 0.6096563 9847 7.110657e+03 1.7627480

16) METADom_PROCYMIDONE< 131.3155 9731 6.742395e+03 1.7425200

32) BOMDom_min_temperature_csum_0< 1518.75 8740 5.753470e+03 1.6816650

64) SatDom_Reflect_b6_var..70< 0.0001229021 5209 3.198972e+03 1.5321370

128) METADom_OMETHOATE_repeat_1< 0.2489644 4893 2.653462e+03 1.4795990

256) SatDom_FPAR_min..70< 0.13625 3076 1.447416e+03 1.3202020

512) ENVDom_Any_damage< 0.002503859 2804 1.136177e+03 1.2538940

1024) METADom_Fertiliser_DAP.ZINC.COTE.2.5.< 77.5 2662 9.756675e+02 1.2019420

2048) SatDom_Reflect_b1_mean..70>=0.1345462 2353 7.567105e+02 1.1271310

4096) SatDom_PsnNet_mean.0< 0.0051375 1355 3.467879e+02 0.9597343

8192) SatDom_LST_night_var..20< 7.286894 1064 2.159415e+02 0.8582331

16384) SatDom_Reflect_b2_max.0< 0.20175 493 3.722596e+01 0.6558621 *

16385) SatDom_Reflect_b2_max.0>=0.20175 571 1.410929e+02 1.0329600

32770) SatDom_NIR_mean..40>=0.2137156 471 6.743882e+01 0.9378981 *

32771) SatDom_NIR_mean..40< 0.2137156 100 4.935065e+01 1.4807000 *

8193) SatDom_LST_night_var..20>=7.286894 291 7.980409e+01 1.3308590 *

4097) SatDom_PsnNet_mean.0>=0.0051375 998 3.204014e+02 1.3544090

8194) SatDom_PsnNet_max..10>=0.00549375 791 1.968491e+02 1.2152210

16388) SatDom_Emis32_var..70>=5.736111e-07 488 8.460715e+01 1.0315980

32776) SatDom_Pot_EvapoTrans_mean..30>=42.3225 278 2.849078e+01 0.8357914 *

32777) SatDom_Pot_EvapoTrans_mean..30< 42.3225 210 3.134776e+01 1.2908100 *

16389) SatDom_Emis32_var..70< 5.736111e-07 303 6.928762e+01 1.5109570

32778) SatDom_Emis31_mean..50< 0.983 104 8.912253e+00 1.1093270 *

32779) SatDom_Emis31_mean..50>=0.983 199 3.483215e+01 1.7208540 *

8195) SatDom_PsnNet_max..10< 0.00549375 207 4.967044e+01 1.8862800 *

2049) SatDom_Reflect_b1_mean..70< 0.1345462 309 1.055080e+02 1.7716180

4098) METADom_Fert_rotation_minus_1_SULPHATE>=0.05922024 131 3.344923e+01 1.3944270 *

4099) METADom_Fert_rotation_minus_1_SULPHATE< 0.05922024 178 3.970449e+01 2.0492130 *

1025) METADom_Fertiliser_DAP.ZINC.COTE.2.5.>=77.5 142 1.863462e+01 2.2278170 *

513) ENVDom_Any_damage>=0.002503859 272 1.718218e+02 2.0037500 *

257) SatDom_FPAR_min..70>=0.13625 1817 9.955847e+02 1.7494440

514) METADom_Chem_rotation_minus_2_DUAL_GOLD>=0.002889951 1476 7.038033e+02 1.6045930

1028) SatDom_Latent_heat_min.0>=1036596 495 1.403152e+02 1.2451920

2056) SatDom_Pot_EvapoTrans_min..50< 42.625 247 4.212298e+01 1.0106880 *

2057) SatDom_Pot_EvapoTrans_min..50>=42.625 248 7.108091e+01 1.4787500 *

1029) SatDom_Latent_heat_min.0< 1036596 981 4.672865e+02 1.7859430

2058) SatDom_MIR_var.0< 3.263921e-05 545 1.777288e+02 1.5092110

4116) BOMDom_solar_exposure_min_0>=4.85 425 9.866918e+01 1.3401880

8232) SatDom_LAI_var..20>=0.001333333 114 9.582057e+00 0.9627193 *

8233) SatDom_LAI_var..20< 0.001333333 311 6.689005e+01 1.4785530 *

4117) BOMDom_solar_exposure_min_0< 4.85 120 2.391604e+01 2.1078330 *

2059) SatDom_MIR_var.0>=3.263921e-05 436 1.956508e+02 2.1318580

4118) SatDom_Reflect_b5_mean.0< 0.3058156 274 6.197626e+01 1.8104010 *

4119) SatDom_Reflect_b5_mean.0>=0.3058156 162 5.747260e+01 2.6755560 *

515) METADom_Chem_rotation_minus_2_DUAL_GOLD< 0.002889951 341 1.267648e+02 2.3764220

1030) SatDom_Reflect_b7_var..50< 2.309099e-05 100 1.921085e+01 1.8693000 *

1031) SatDom_Reflect_b7_var..50>=2.309099e-05 241 7.116560e+01 2.5868460 *

129) METADom_OMETHOATE_repeat_1>=0.2489644 316 3.228830e+02 2.3456330

258) SatDom_Emis32_var..60>=1.027778e-06 104 1.695069e+01 1.2227880 *

259) SatDom_Emis32_var..60< 1.027778e-06 212 1.104876e+02 2.8964620 *

65) SatDom_Reflect_b6_var..70>=0.0001229021 3531 2.266219e+03 1.9022510

130) SatDom_Reflect_b6_min..60>=0.4081625 720 3.258849e+02 1.4524580

260) METADom_Chem_rotation_minus_1_BRODAL< 0.001131369 379 9.972405e+01 1.0957780

520) METADom_Organic_C_10cm.test_10cm< 1.415206 216 1.692079e+01 0.8001852 *

521) METADom_Organic_C_10cm.test_10cm>=1.415206 163 3.892047e+01 1.4874850 *

261) METADom_Chem_rotation_minus_1_BRODAL>=0.001131369 341 1.243546e+02 1.8488860

522) SatDom_GPP..60>=0.24485 188 2.678640e+01 1.5301060 *

523) SatDom_GPP..60< 0.24485 153 5.498865e+01 2.2405880 *

131) SatDom_Reflect_b6_min..60< 0.4081625 2811 1.757358e+03 2.0174600

262) SatDom_NIR_var..10< 4.126655e-06 687 3.970925e+02 1.6113680

524) SatDom_Reflect_b5_min.0>=0.2049625 532 2.140570e+02 1.3967860

1048) BOMDom_min_temperature_min_.70< 11.65 411 1.080193e+02 1.2289540

2096) SatDom_PsnNet..70< 0.5603438 164 1.793349e+01 0.8751829 *

2097) SatDom_PsnNet..70>=0.5603438 247 5.593245e+01 1.4638460 *

1049) BOMDom_min_temperature_min_.70>=11.65 121 5.513781e+01 1.9668600 *

525) SatDom_Reflect_b5_min.0< 0.2049625 155 7.446160e+01 2.3478710 *

263) SatDom_NIR_var..10>=4.126655e-06 2124 1.210327e+03 2.1488090

526) SatDom_Emis32_mean..70< 0.9870417 1743 9.014588e+02 2.0383820

1052) SatDom_Pot_EvapoTrans_var..10< 6.04963e+07 1383 6.335769e+02 1.9051480

2104) SatDom_NIR_var..80>=5.337198e-07 1235 4.862838e+02 1.8120320

4208) BOMDom_solar_exposure_var_.10< 4.954722 486 1.344499e+02 1.4765840

8416) BOMDom_min_temperature_min_.70< 11.85 326 4.631271e+01 1.2787730 *

8417) BOMDom_min_temperature_min_.70>=11.85 160 4.939038e+01 1.8796250 *

4209) BOMDom_solar_exposure_var_.10>=4.954722 749 2.616618e+02 2.0296930

8418) SatDom_PsnNet_var..20< 1.606771e-07 206 8.122734e+01 1.6335920 *

8419) SatDom_PsnNet_var..20>=1.606771e-07 543 1.358524e+02 2.1799630

16838) SatDom_EvapoTrans_mean..50>=3.2575 245 5.480112e+01 1.9250610 *

16839) SatDom_EvapoTrans_mean..50< 3.2575 298 5.204473e+01 2.3895300 *

2105) SatDom_NIR_var..80< 5.337198e-07 148 4.722991e+01 2.6821620 *

1053) SatDom_Pot_EvapoTrans_var..10>=6.04963e+07 360 1.490190e+02 2.5502220

2106) SatDom_Emis31_mean..40< 0.98335 232 5.613688e+01 2.2757330 *

2107) SatDom_Emis31_mean..40>=0.98335 128 4.371984e+01 3.0477340 *

527) SatDom_Emis32_mean..70>=0.9870417 381 1.903803e+02 2.6539900

1054) SatDom_Reflect_b4_var..60< 2.00728e-05 277 8.533373e+01 2.4587000 *

1055) SatDom_Reflect_b4_var..60>=2.00728e-05 104 6.634512e+01 3.1741350 *

33) BOMDom_min_temperature_csum_0>=1518.75 991 6.711001e+02 2.2792230

66) SatDom_GPP_min..60< 0.00485 388 1.901659e+02 1.7939950

132) SatDom_Reflect_b1_max..10< 0.1863188 286 1.052877e+02 1.5524830 *

133) SatDom_Reflect_b1_max..10>=0.1863188 102 2.142166e+01 2.4711760 *

67) SatDom_GPP_min..60>=0.00485 603 3.307998e+02 2.5914430

134) METADom_Chem_rotation_minus_2_FASTAC>=0.000776699 358 1.491185e+02 2.2768440

268) SatDom_EVI_max..40< 0.108525 119 2.598569e+01 1.7397480 *

269) SatDom_EVI_max..40>=0.108525 239 7.171245e+01 2.5442680 *

135) METADom_Chem_rotation_minus_2_FASTAC< 0.000776699 245 9.447468e+01 3.0511430 *

17) METADom_PROCYMIDONE>=131.3155 116 3.025899e+01 3.4596550 *

9) SatDom_GPP..20>=0.6096563 15826 9.647824e+03 2.1272930

18) METADom_Chem_rotation_minus_1_TERBYNE< 0.00340273 14864 8.861072e+03 2.0844270

36) METADom_Chem_rotation_minus_1_FORCE< 0.001257611 12934 7.242096e+03 2.0301030

72) METADom_Chem_rotation_minus_2_BONUS>=-1.218643e-17 12474 6.626925e+03 2.0002460

144) SatDom_EvapoTrans_min..70< 4.58125 5727 3.004448e+03 1.8553310

288) BOMDom_solar_exposure_mean_.40< 16.84 2558 1.099319e+03 1.6269000

576) BOMDom_max_temperature_max_0>=22.95 1389 3.856871e+02 1.3834770

1152) SatDom_EVI_min..60>=0.1281844 907 2.116970e+02 1.2231640

2304) SatDom_Reflect_b3_var..60< 5.13892e-06 392 4.585976e+01 0.9533673

4608) SatDom_Latent_heat_max..20< 2096250 286 1.391296e+01 0.8010839 *

4609) SatDom_Latent_heat_max..20>=2096250 106 7.419390e+00 1.3642450 *

2305) SatDom_Reflect_b3_var..60>=5.13892e-06 515 1.155845e+02 1.4285240

4610) SatDom_LAI_var..60< 0.00140625 283 4.147229e+01 1.1786570 *

4611) SatDom_LAI_var..60>=0.00140625 232 3.489074e+01 1.7333190 *

1153) SatDom_EVI_min..60< 0.1281844 482 1.068162e+02 1.6851450

2306) SatDom_LST_night_min..60< 32.525 365 5.539122e+01 1.5344660 *

2307) SatDom_LST_night_min..60>=32.525 117 1.728512e+01 2.1552140 *

577) BOMDom_max_temperature_max_0< 22.95 1169 5.335329e+02 1.9161330

1154) BOMDom_min_temperature_min_.20< 4.65 690 2.689583e+02 1.6972170

2308) SatDom_Reflect_b6_min.0>=0.2875937 254 3.568920e+01 1.3000790 *

2309) SatDom_Reflect_b6_min.0< 0.2875937 436 1.698703e+02 1.9285780

4618) BOMDom_min_temperature_var_.10< 12.81694 327 5.702664e+01 1.7608870 *

4619) BOMDom_min_temperature_var_.10>=12.81694 109 7.606230e+01 2.4316510 *

1155) BOMDom_min_temperature_min_.20>=4.65 479 1.838728e+02 2.2314820

2310) METADom_Fertiliser_time_MAP>=-15.75484 338 6.819616e+01 1.9826330 *

2311) METADom_Fertiliser_time_MAP< -15.75484 141 4.457084e+01 2.8280140 *

289) BOMDom_solar_exposure_mean_.40>=16.84 3169 1.663908e+03 2.0397190

578) METADom_Fertiliser_PIVOT.15_repeat_1>=-3.433018e-16 2938 1.405448e+03 1.9799900

1156) SatDom_Reflect_b7_mean..10>=0.1853894 2477 1.105659e+03 1.8897050

2312) BOMDom_solar_exposure_min_.40>=6.75 2093 9.108137e+02 1.7998180

4624) BOMDom_min_temperature_min_.20< 6.4 801 2.542636e+02 1.5122600

9248) BOMDom_min_temperature_var_.60< 5.167833 317 4.440979e+01 1.1286120 *

9249) BOMDom_min_temperature_var_.60>=5.167833 484 1.326371e+02 1.7635330

18498) BOMDom_max_temperature_max_.20>=26.8 337 4.435272e+01 1.5581310 *

18499) BOMDom_max_temperature_max_.20< 26.8 147 4.147103e+01 2.2344220 *

4625) BOMDom_min_temperature_min_.20>=6.4 1292 5.492519e+02 1.9780960

9250) METADom_Nitrogen_fert< 16.475 180 5.712758e+01 1.3758330 *

9251) METADom_Nitrogen_fert>=16.475 1112 4.162662e+02 2.0755850

18502) BOMDom_rainfall_csum_0< 125.95 895 2.634281e+02 1.9358660

37004) SatDom_NIR_var..70>=1.328692e-05 397 9.041754e+01 1.6567250

74008) SatDom_Reflect_b4_min..10>=0.09125 131 1.357090e+01 1.2929010 *

74009) SatDom_Reflect_b4_min..10< 0.09125 266 5.096663e+01 1.8359020 *

37005) SatDom_NIR_var..70< 1.328692e-05 498 1.174163e+02 2.1583940

74010) SatDom_LST_night_var..70>=4.066871 374 5.399489e+01 2.0198660

148020) SatDom_Latent_heat_var..30< 7.181043e+10 263 1.915329e+01 1.8529660 *

148021) SatDom_Latent_heat_var..30>=7.181043e+10 111 1.015736e+01 2.4153150 *

74011) SatDom_LST_night_var..70< 4.066871 124 3.459772e+01 2.5762100 *

18503) BOMDom_rainfall_csum_0>=125.95 217 6.330646e+01 2.6518430 *

2313) BOMDom_solar_exposure_min_.40< 6.75 384 8.576235e+01 2.3796350

4626) SatDom_Latent_heat_min..50< 481875 136 2.417975e+01 1.9925000 *

4627) SatDom_Latent_heat_min..50>=481875 248 3.002207e+01 2.5919350 *

1157) SatDom_Reflect_b7_mean..10< 0.1853894 461 1.711115e+02 2.4650980

2314) BOMDom_max_temperature_mean_0>=21.26 144 1.987704e+01 1.9427080 *

2315) BOMDom_max_temperature_mean_0< 21.26 317 9.408758e+01 2.7023970

4630) SatDom_FPAR_min..70< 0.12 107 2.121175e+01 2.2620560 *

4631) SatDom_FPAR_min..70>=0.12 210 4.155720e+01 2.9267620 *

579) METADom_Fertiliser_PIVOT.15_repeat_1< -3.433018e-16 231 1.146667e+02 2.7993940 *

145) SatDom_EvapoTrans_min..70>=4.58125 6747 3.400120e+03 2.1232530

290) BOMDom_min_temperature_min_.50< 1.6 171 8.013667e+01 1.1225730 *

291) BOMDom_min_temperature_min_.50>=1.6 6576 3.144298e+03 2.1492750

582) METADom_total_N_10cm.test_10cm< 20.15 1535 5.690283e+02 1.8755500

1164) SatDom_Reflect_b7_min..20>=0.1507438 1219 3.902416e+02 1.7545530

2328) BOMDom_Longitude< 148.1397 910 2.320023e+02 1.6085710

4656) BOMDom_min_temperature_max_.80< 22.25 668 1.279839e+02 1.4692370

9312) SatDom_Reflect_b2_min..60>=0.1886 563 8.306678e+01 1.3828950

18624) METADom_Chem_rotation_minus_1_SPRAY>=0.0001954397 176 1.972587e+01 1.0886360 *

18625) METADom_Chem_rotation_minus_1_SPRAY< 0.0001954397 387 4.117073e+01 1.5167180 *

9313) SatDom_Reflect_b2_min..60< 0.1886 105 1.821580e+01 1.9321900 *

4657) BOMDom_min_temperature_max_.80>=22.25 242 5.525185e+01 1.9931820 *

2329) BOMDom_Longitude>=148.1397 309 8.173564e+01 2.1844660

4658) SatDom_LST_night_var..20>=2.388813 201 3.271999e+01 1.9607460 *

4659) SatDom_LST_night_var..20< 2.388813 108 2.023243e+01 2.6008330 *

1165) SatDom_Reflect_b7_min..20< 0.1507438 316 9.209481e+01 2.3423100

2330) BOMDom_solar_exposure_var_.50< 14.67361 141 2.412772e+01 1.9548230 *

2331) BOMDom_solar_exposure_var_.50>=14.67361 175 2.973893e+01 2.6545140 *

583) METADom_total_N_10cm.test_10cm>=20.15 5041 2.425239e+03 2.2326240

1166) SatDom_Pot_EvapoTrans_max..70>=46.925 4064 1.949929e+03 2.1546830

2332) SatDom_Reflect_b2..60< 9.35905 3862 1.774644e+03 2.1143400

4664) BOMDom_max_temperature_max_.80< 37.85 2990 1.347412e+03 2.0267560

9328) SatDom_LST_night_var..60>=1.820207 2455 1.031708e+03 1.9431080

18656) SatDom_NIR_max..50>=0.1802219 2256 8.458801e+02 1.8812770

37312) BOMDom_rainfall_mean_.20< 0.03 297 8.331856e+01 1.4112120 *

37313) BOMDom_rainfall_mean_.20>=0.03 1959 6.869869e+02 1.9525420

74626) SatDom_Reflect_b3_min..10< 0.03185 248 5.241938e+01 1.4427820 *

74627) SatDom_Reflect_b3_min..10>=0.03185 1711 5.607827e+02 2.0264290

149254) SatDom_Reflect_b2_min..10< 0.2072562 1024 2.786914e+02 1.8765820

298508) BOMDom_max_temperature_max_.40>=27.95 690 1.335624e+02 1.7215070

597016) BOMDom_solar_exposure_mean_0< 9.285 180 1.926017e+01 1.3958330 *

597017) BOMDom_solar_exposure_mean_0>=9.285 510 8.847268e+01 1.8364510 *

298509) BOMDom_max_temperature_max_.40< 27.95 334 9.425629e+01 2.1969460

597018) SatDom_FPAR_min..10>=0.2430728 169 2.350956e+01 1.9069820 *

597019) SatDom_FPAR_min..10< 0.2430728 165 4.198354e+01 2.4939390 *

149255) SatDom_Reflect_b2_min..10>=0.2072562 687 2.248263e+02 2.2497820

298510) SatDom_Reflect_b2_var..20< 7.068567e-05 443 1.179951e+02 2.0688490

597020) SatDom_MIR_var..10< 1.549202e-05 226 5.743828e+01 1.8419030 *

597021) SatDom_MIR_var..10>=1.549202e-05 217 3.679402e+01 2.3052070 *

298511) SatDom_Reflect_b2_var..20>=7.068567e-05 244 6.599868e+01 2.5782790 *

18657) SatDom_NIR_max..50< 0.1802219 199 7.942480e+01 2.6440700 *

9329) SatDom_LST_night_var..60< 1.820207 535 2.197024e+02 2.4105980

18658) SatDom_LST_night_min..70>=30.89 391 1.142928e+02 2.2037850

37316) SatDom_Reflect_b4..60>=2.993737 248 3.688340e+01 2.0151210 *

37317) SatDom_Reflect_b4..60< 2.993737 143 5.327306e+01 2.5309790 *

18659) SatDom_LST_night_min..70< 30.89 144 4.327643e+01 2.9721530 *

4665) BOMDom_max_temperature_max_.80>=37.85 872 3.256505e+02 2.4146560

9330) SatDom_PsnNet_max..10< 0.0192 722 2.004202e+02 2.3064130

18660) SatDom_LST_day_var..80>=10.41232 360 1.022264e+02 2.0746390

37320) SatDom_Reflect_b1.0< 13.4246 259 6.250640e+01 1.9072200 *

37321) SatDom_Reflect_b1.0>=13.4246 101 1.384442e+01 2.5039600 *

18661) SatDom_LST_day_var..80< 10.41232 362 5.962293e+01 2.5369060 *

9331) SatDom_PsnNet_max..10>=0.0192 150 7.605308e+01 2.9356670 *

2333) SatDom_Reflect_b2..60>=9.35905 202 4.882605e+01 2.9259900 *

1167) SatDom_Pot_EvapoTrans_max..70< 46.925 977 3.479255e+02 2.5568370

2334) SatDom_LST_night_max..20< 33.04 739 2.249306e+02 2.4055620

4668) SatDom_blue_band_var..70>=2.206624e-06 311 8.216565e+01 2.1443730

9336) SatDom_Emis32_var..80< 7.234568e-07 182 3.692671e+01 1.9071430 *

9337) SatDom_Emis32_var..80>=7.234568e-07 129 2.054549e+01 2.4790700 *

4669) SatDom_blue_band_var..70< 2.206624e-06 428 1.061322e+02 2.5953500 *

2335) SatDom_LST_night_max..20>=33.04 238 5.357237e+01 3.0265550 *

73) METADom_Chem_rotation_minus_2_BONUS< -1.218643e-17 460 3.025170e+02 2.8397390

146) SatDom_Reflect_b7_var..40>=1.463795e-05 345 1.003336e+02 2.5172750

292) SatDom_LAI_var..40< 0.006684028 104 3.659813e+01 2.0971150 *

293) SatDom_LAI_var..40>=0.006684028 241 3.745312e+01 2.6985890 *

147) SatDom_Reflect_b7_var..40< 1.463795e-05 115 5.868695e+01 3.8071300 *

37) METADom_Chem_rotation_minus_1_FORCE>=0.001257611 1930 1.325005e+03 2.4484870

74) SatDom_Reflect_b3.0< 4.606575 241 4.765872e+01 1.4994190 *

75) SatDom_Reflect_b3.0>=4.606575 1689 1.029296e+03 2.5839080

150) METADom_Chem_rotation_minus_2_ROUNDUP>=0.06626442 1518 7.280901e+02 2.4688600

300) SatDom_EVI_var..70< 3.775988e-06 510 2.444064e+02 2.0694710

600) SatDom_Reflect_b1_min..50>=0.1468813 126 1.212385e+01 1.1969840 *

601) SatDom_Reflect_b1_min..50< 0.1468813 384 1.048950e+02 2.3557550

1202) SatDom_LST_night_var..30< 4.412724 136 4.148019e+01 1.9852940 *

1203) SatDom_LST_night_var..30>=4.412724 248 3.451441e+01 2.5589110 *

301) SatDom_EVI_var..70>=3.775988e-06 1008 3.611727e+02 2.6709330

602) BOMDom_min_temperature_min_.10< 2.55 142 4.412810e+01 2.0363380 *

603) BOMDom_min_temperature_min_.10>=2.55 866 2.504830e+02 2.7749880

1206) BOMDom_solar_exposure_min_.10>=8.25 151 1.823991e+01 2.2311260 *

1207) BOMDom_solar_exposure_min_.10< 8.25 715 1.781469e+02 2.8898460

2414) SatDom_LAI_mean.0>=0.35625 305 4.322271e+01 2.6175080 *

2415) SatDom_LAI_mean.0< 0.35625 410 9.547496e+01 3.0924390

4830) SatDom_Latent_heat_var..40< 1.580372e+10 281 3.143917e+01 2.9328830 *

4831) SatDom_Latent_heat_var..40>=1.580372e+10 129 4.129900e+01 3.4400000 *

151) METADom_Chem_rotation_minus_2_ROUNDUP< 0.06626442 171 1.027527e+02 3.6052050 *

19) METADom_Chem_rotation_minus_1_TERBYNE>=0.00340273 962 3.374380e+02 2.7896150

38) BOMDom_solar_exposure_var_0< 6.430944 475 1.196869e+02 2.4558320

76) SatDom_Reflect_b2_var..50>=2.618084e-05 359 5.131429e+01 2.2774090 *

77) SatDom_Reflect_b2_var..50< 2.618084e-05 116 2.157464e+01 3.0080170 *

39) BOMDom_solar_exposure_var_0>=6.430944 487 1.132140e+02 3.1151750

78) BOMDom_solar_exposure_var_.20>=5.489333 374 5.994392e+01 2.9529140 *

79) BOMDom_solar_exposure_var_.20< 5.489333 113 1.083295e+01 3.6522120 *

5) METADom_sub_cropFaba.Bean>=0.5 2036 3.156017e+03 2.9263360

10) METADom_CHLOROTHALONIL_repeat_3< 3.912885 1864 2.663262e+03 2.8062340

20) SatDom_GPP_max..20< 0.007025 486 7.113529e+02 2.2223250

40) SatDom_Reflect_b7_max..70>=0.2915 193 1.068199e+02 1.5269430 *

41) SatDom_Reflect_b7_max..70< 0.2915 293 4.497323e+02 2.6803750 *

21) SatDom_GPP_max..20>=0.007025 1378 1.727767e+03 3.0121700

42) METADom_SPINETORAM_repeat_1>=-4.071396e-16 1237 1.427219e+03 2.9080520

84) METADom_Organic_C_10cm.test_10cm< 1.523714 1053 1.092792e+03 2.7855560

168) SatDom_Reflect_b6_mean.0>=0.3193106 201 1.280242e+02 2.1284580 *

169) SatDom_Reflect_b6_mean.0< 0.3193106 852 8.575060e+02 2.9405750

338) SatDom_Reflect_b3_max..60< 0.092875 746 6.384346e+02 2.8047720

676) SatDom_EvapoTrans_max..80< 4.3 103 6.238712e+01 1.9957280 *

677) SatDom_EvapoTrans_max..80>=4.3 643 4.978290e+02 2.9343700

1354) SatDom_Reflect_b7_var..60>=1.761313e-05 501 3.480054e+02 2.7579440

2708) SatDom_Pot_EvapoTrans_mean..50>=53.57313 330 1.725076e+02 2.4670910

5416) SatDom_NDVI_min..20>=0.2335953 229 7.764098e+01 2.2541920 *

5417) SatDom_NDVI_min..20< 0.2335953 101 6.095300e+01 2.9498020 *

2709) SatDom_Pot_EvapoTrans_mean..50< 53.57313 171 9.370720e+01 3.3192400 *

1355) SatDom_Reflect_b7_var..60< 1.761313e-05 142 7.921047e+01 3.5568310 *

339) SatDom_Reflect_b3_max..60>=0.092875 106 1.084877e+02 3.8963210 *

85) METADom_Organic_C_10cm.test_10cm>=1.523714 184 2.282019e+02 3.6090760 *

43) METADom_SPINETORAM_repeat_1< -4.071396e-16 141 1.694941e+02 3.9256030 *

11) METADom_CHLOROTHALONIL_repeat_3>=3.912885 172 1.744848e+02 4.2279070 *

3) METADom_Dicot_TF< 0.5 69624 1.570270e+05 3.3770900

6) SatDom_Reflect_b7.0>=24.17471 30344 5.774680e+04 2.8173200

12) METADom_Year< 2015.5 25893 3.792992e+04 2.6310030

24) SatDom_GPP..40< 0.4096125 13597 1.522992e+04 2.2474970

48) BOMDom_solar_exposure_mean_.60< 20.315 8046 7.127546e+03 1.9083160

96) BOMDom_rainfall_csum_.10< 43.3 3132 1.749617e+03 1.4877230

192) BOMDom_min_temperature_max_0>=9.65 2671 1.116392e+03 1.3508650

384) SatDom_NDVI_var..10< 0.0001123014 2242 7.431482e+02 1.2220380

768) METADom_Chem_rotation_minus_2_ATRAZINE< 0.1922788 2015 5.129262e+02 1.1249130

1536) SatDom_LST_day..50>=2340.6 613 4.862163e+01 0.7517455

3072) SatDom_LST_day_var..50< 16.34615 372 1.255180e+01 0.5867204 *

3073) SatDom_LST_day_var..50>=16.34615 241 1.030150e+01 1.0064730 *

1537) SatDom_LST_day..50< 2340.6 1402 3.416184e+02 1.2880740

3074) SatDom_Emis32_var..70>=4.277778e-07 1131 1.868594e+02 1.1486560

6148) SatDom_Pot_EvapoTrans..30< 2818.075 317 3.172011e+01 0.7947950 *

6149) SatDom_Pot_EvapoTrans..30>=2818.075 814 9.998701e+01 1.2864620

12298) SatDom_FPAR_var..70< 6.423611e-05 508 4.210297e+01 1.1324410 *

12299) SatDom_FPAR_var..70>=6.423611e-05 306 2.582678e+01 1.5421570 *

3075) SatDom_Emis32_var..70< 4.277778e-07 271 4.102780e+01 1.8699260 *

769) METADom_Chem_rotation_minus_2_ATRAZINE>=0.1922788 227 4.248552e+01 2.0841850 *

385) SatDom_NDVI_var..10>=0.0001123014 429 1.415780e+02 2.0241260

770) SatDom_NIR_var..80< 6.529154e-07 132 1.120913e+01 1.3966670 *

771) SatDom_NIR_var..80>=6.529154e-07 297 5.530243e+01 2.3029970 *

193) BOMDom_min_temperature_max_0< 9.65 461 2.933341e+02 2.2806720

386) SatDom_PsnNet_var..20< 5.722656e-08 213 5.583169e+01 1.6409860 *

387) SatDom_PsnNet_var..20>=5.722656e-08 248 7.548440e+01 2.8300810 *

97) BOMDom_rainfall_csum_.10>=43.3 4914 4.470758e+03 2.1763860

194) BOMDom_rainfall_var_.70>=0.01383333 3086 2.178925e+03 1.8717660

388) SatDom_blue_band_var..20>=3.444661e-07 2586 1.316779e+03 1.7011330

776) METADom_Chem_rotation_minus_2_MCPA< 0.1377111 2318 9.300912e+02 1.5888400

1552) SatDom_GPP_max..30< 0.00694375 1155 3.377782e+02 1.3136450

3104) SatDom_NDVI_var..50>=1.314553e-07 1043 2.074772e+02 1.2095970

6208) SatDom_NDVI_var..40< 1.306723e-05 641 7.655085e+01 1.0275510

12416) METADom_Chem_rotation_minus_1_BOXERGOLD>=-3.176712e-17 419 4.202529e+01 0.8884487 *

12417) METADom_Chem_rotation_minus_1_BOXERGOLD< -3.176712e-17 222 1.111640e+01 1.2900900 *

6209) SatDom_NDVI_var..40>=1.306723e-05 402 7.580989e+01 1.4998760 *

3105) SatDom_NDVI_var..50< 1.314553e-07 112 1.385795e+01 2.2825890 *

1553) SatDom_GPP_max..30>=0.00694375 1163 4.179738e+02 1.8621410

3106) SatDom_LST_day_max..70>=59.93 906 1.724519e+02 1.6668540

6212) SatDom_Pot_latent_heat_mean..60< 1.332344e+07 188 2.843397e+01 1.1769680 *

6213) SatDom_Pot_latent_heat_mean..60>=1.332344e+07 718 8.708654e+01 1.7951250

12426) SatDom_EVI_var..40< 5.597575e-06 543 3.569265e+01 1.6703130 *

12427) SatDom_EVI_var..40>=5.597575e-06 175 1.668819e+01 2.1824000 *

3107) SatDom_LST_day_max..70< 59.93 257 8.916381e+01 2.5505840 *

777) METADom_Chem_rotation_minus_2_MCPA>=0.1377111 268 1.046443e+02 2.6723880 *

389) SatDom_blue_band_var..20< 3.444661e-07 500 3.974374e+02 2.7542800

778) METADom_Fert_rotation_minus_2_UREA< 0.4275046 222 7.761915e+01 2.0082880 *

779) METADom_Fert_rotation_minus_2_UREA>=0.4275046 278 9.761720e+01 3.3500000 *

195) BOMDom_rainfall_var_.70< 0.01383333 1828 1.522045e+03 2.6906400

390) SatDom_Pot_EvapoTrans_max..60< 51.00625 639 3.161248e+02 2.0780590

780) SatDom_GPP_var..10< 4.744661e-07 411 1.136247e+02 1.7098780

1560) BOMDom_min_temperature_min_.50>=5.9 248 3.248379e+01 1.4598390 *

1561) BOMDom_min_temperature_min_.50< 5.9 163 4.204568e+01 2.0903070 *

781) SatDom_GPP_var..10>=4.744661e-07 228 4.635410e+01 2.7417540 *

391) SatDom_Pot_EvapoTrans_max..60>=51.00625 1189 8.372635e+02 3.0198570

782) SatDom_Emis32_mean..20< 0.9865 628 2.192259e+02 2.5190610

1564) SatDom_Reflect_b4_min..10< 0.08865 298 5.528640e+01 2.1105700 *

1565) SatDom_Reflect_b4_min..10>=0.08865 330 6.931040e+01 2.8879390 *

783) SatDom_Emis32_mean..20>=0.9865 561 2.842261e+02 3.5804630

1566) SatDom_Reflect_b7_min..50< 0.317775 362 5.592281e+01 3.1525140 *

1567) SatDom_Reflect_b7_min..50>=0.317775 199 4.140568e+01 4.3589450 *

49) BOMDom_solar_exposure_mean_.60>=20.315 5551 5.835043e+03 2.7391280

98) SatDom_GPP_min..80< 0.0065375 3915 3.270136e+03 2.4579820

196) METADom_Fert_rotation_minus_1_TWENTY_FOUR_TO_SIXTEEN>=0.0001547988 952 5.373003e+02 1.8678360

392) SatDom_GPP..20< 0.43535 198 4.357373e+01 0.9053535 *

393) SatDom_GPP..20>=0.43535 754 2.621383e+02 2.1205840

786) BOMDom_min_temperature_csum_.80< 207.975 575 1.033302e+02 1.8980520

1572) SatDom_EvapoTrans_min.0< 5 350 2.371625e+01 1.6903710 *

1573) SatDom_EvapoTrans_min.0>=5 225 4.103542e+01 2.2211110 *

787) BOMDom_min_temperature_csum_.80>=207.975 179 3.886684e+01 2.8354190 *

197) METADom_Fert_rotation_minus_1_TWENTY_FOUR_TO_SIXTEEN< 0.0001547988 2963 2.294753e+03 2.6475940

394) SatDom_Emis32_var..10< 6.462963e-07 713 2.257891e+02 1.8980220

788) BOMDom_solar_exposure_max_.30>=18.45 582 9.125362e+01 1.7038830

1576) SatDom_LST_day_var..60< 9.179545 114 1.261334e+01 1.2671050 *

1577) SatDom_LST_day_var..60>=9.179545 468 5.159426e+01 1.8102780 *

789) BOMDom_solar_exposure_max_.30< 18.45 131 1.514546e+01 2.7605340 *

395) SatDom_Emis32_var..10>=6.462963e-07 2250 1.541413e+03 2.8851240

790) SatDom_Reflect_b6_max.0>=0.3514875 1769 9.952865e+02 2.6907010

1580) SatDom_PsnNet_max..30< 0.00655 1302 5.168018e+02 2.4796010

3160) SatDom_MIR_min..80>=0.3438062 699 2.346951e+02 2.1497570

6320) SatDom_Reflect_b5_var..40< 7.854488e-05 443 9.770893e+01 1.9055760

12640) SatDom_MIR_var..30>=3.401872e-05 185 4.008506e+01 1.5795140 *

12641) SatDom_MIR_var..30< 3.401872e-05 258 2.385190e+01 2.1393800 *

6321) SatDom_Reflect_b5_var..40>=7.854488e-05 256 6.486454e+01 2.5723050 *

3161) SatDom_MIR_min..80< 0.3438062 603 1.179013e+02 2.8619570

6322) SatDom_Emis31_var..20>=5.444444e-07 491 6.892988e+01 2.7406720 *

6323) SatDom_Emis31_var..20< 5.444444e-07 112 1.008540e+01 3.3936610 *

1581) SatDom_PsnNet_max..30>=0.00655 467 2.586988e+02 3.2792510

3162) SatDom_Reflect_b6_mean..50< 0.406425 218 5.029002e+01 2.7409170 *

3163) SatDom_Reflect_b6_mean..50>=0.406425 249 8.992032e+01 3.7505620 *

791) SatDom_Reflect_b6_max.0< 0.3514875 481 2.333296e+02 3.6001660

1582) SatDom_LST_day_mean..60< 60.9572 340 6.795570e+01 3.3886760 *

1583) SatDom_LST_day_mean..60>=60.9572 141 1.134958e+02 4.1101420 *

99) SatDom_GPP_min..80>=0.0065375 1636 1.514921e+03 3.4119190

198) SatDom_Latent_heat_max..40>=845000 1265 7.105334e+02 3.1145380

396) SatDom_Latent_heat_var..30< 6.196719e+09 340 6.198470e+01 2.3920290 *

397) SatDom_Latent_heat_var..30>=6.196719e+09 925 4.058244e+02 3.3801080

794) SatDom_Pot_latent_heat_max..70>=1.7785e+07 700 2.121292e+02 3.1708000

1588) SatDom_FPAR..30>=9.41 179 1.652997e+01 2.6474860 *

1589) SatDom_FPAR..30< 9.41 521 1.297367e+02 3.3505950

3178) METADom_Chem_rotation_minus_2_SPRAYSEED< 0.03249206 393 6.868886e+01 3.2288300 *

3179) METADom_Chem_rotation_minus_2_SPRAYSEED>=0.03249206 128 3.733036e+01 3.7244530 *

795) SatDom_Pot_latent_heat_max..70< 1.7785e+07 225 6.762013e+01 4.0312890 *

199) SatDom_Latent_heat_max..40< 845000 371 3.110682e+02 4.4259030

398) METADom_Chem_rotation_minus_1_BOXERGOLD< -3.272122e-17 146 6.923450e+01 3.6739730 *

399) METADom_Chem_rotation_minus_1_BOXERGOLD>=-3.272122e-17 225 1.057207e+02 4.9138220 *

25) SatDom_GPP..40>=0.4096125 12296 1.848880e+04 3.0550870

50) BOMDom_rainfall_csum_.10< 47.4 2472 2.503741e+03 2.1931960

100) SatDom_Reflect_b1_mean.0>=0.1322638 1270 6.380561e+02 1.6617240

200) SatDom_MixClouds..10>=3.5 646 1.396467e+02 1.2283900

400) BOMDom_min_temperature_var_.40< 4.845389 180 1.520407e+01 0.7226111 *

401) BOMDom_min_temperature_var_.40>=4.845389 466 6.061033e+01 1.4237550

802) BOMDom_min_temperature_var_.30>=7.556056 318 2.577899e+01 1.2701890 *

803) BOMDom_min_temperature_var_.30< 7.556056 148 1.121866e+01 1.7537160 *

201) SatDom_MixClouds..10< 3.5 624 2.515226e+02 2.1103370

402) SatDom_GPP..50< 0.3703812 202 4.610604e+01 1.5505450 *

403) SatDom_GPP..50>=0.3703812 422 1.118164e+02 2.3782940

806) METADom_Fert_rotation_minus_2_EIGHTEEN_TO_TWENTY< 0.0009434856 210 3.488245e+01 2.0450950 *

807) METADom_Fert_rotation_minus_2_EIGHTEEN_TO_TWENTY>=0.0009434856 212 3.052492e+01 2.7083490 *

101) SatDom_Reflect_b1_mean.0< 0.1322638 1202 1.127937e+03 2.7547340

202) BOMDom_solar_exposure_max_.50< 20.05 529 2.120157e+02 2.1426650

404) METADom_Chem_rotation_minus_2_ROUNDUP_READY>=-1.31839e-17 407 7.251881e+01 1.8986980

808) SatDom_Reflect_b3_var..40>=7.432414e-06 113 1.318064e+01 1.5015040 *

809) SatDom_Reflect_b3_var..40< 7.432414e-06 294 3.465906e+01 2.0513610 *

405) METADom_Chem_rotation_minus_2_ROUNDUP_READY< -1.31839e-17 122 3.445695e+01 2.9565570 *

203) BOMDom_solar_exposure_max_.50>=20.05 673 5.619692e+02 3.2358400

406) SatDom_Reflect_b7_var..50< 3.774441e-05 135 9.428497e+01 2.2247410 *

407) SatDom_Reflect_b7_var..50>=3.774441e-05 538 2.950393e+02 3.4895540

814) SatDom_Aero_q_csum..20>=14 430 1.265696e+02 3.2834880 *

815) SatDom_Aero_q_csum..20< 14 108 7.751240e+01 4.3100000 *

51) BOMDom_rainfall_csum_.10>=47.4 9824 1.368664e+04 3.2719640

102) METADom_Fertiliser_time_MAP< -3.073846 6581 8.283557e+03 3.0267470

204) SatDom_Reflect_b5_mean..20>=0.2969987 5339 5.996877e+03 2.8371700

408) SatDom_FPAR_var..20< 0.001451667 4605 4.573398e+03 2.6882080

816) SatDom_Reflect_b6_var..70>=2.53066e-05 3752 3.200752e+03 2.5322520

1632) BOMDom_min_temperature_mean_.30< 13.24 2575 1.825639e+03 2.2889860

3264) BOMDom_min_temperature_var_.10>=2.8565 2285 1.186221e+03 2.1332520

6528) SatDom_Emis32..40< 50.32014 1522 4.794346e+02 1.9050130

13056) SatDom_FullClouds..80< 3.5 1375 3.359054e+02 1.8134760

26112) SatDom_Reflect_b7_max..50>=0.3610875 146 1.336145e+01 1.2149320 *

26113) SatDom_Reflect_b7_max..50< 0.3610875 1229 2.640249e+02 1.8845810

52226) SatDom_EVI_min.0< 0.1511562 650 1.049825e+02 1.7134620

104452) BOMDom_max_temperature_min_.20< 18.6 412 5.394620e+01 1.5745150 *

104453) BOMDom_max_temperature_min_.20>=18.6 238 2.931271e+01 1.9539920 *

52227) SatDom_EVI_min.0>=0.1511562 579 1.186420e+02 2.0766840

104454) SatDom_LST_day_var..10< 9.53642 436 6.378071e+01 1.9606650 *

104455) SatDom_LST_day_var..10>=9.53642 143 3.109917e+01 2.4304200 *

13057) SatDom_FullClouds..80>=3.5 147 2.424278e+01 2.7612240 *

6529) SatDom_Emis32..40>=50.32014 763 4.693466e+02 2.5885320

13058) SatDom_GPP_max..80< 0.0125 490 1.400860e+02 2.2248980

26116) SatDom_Latent_heat..30>=7.735354e+07 141 2.154813e+01 1.6123400 *

26117) SatDom_Latent_heat..30< 7.735354e+07 349 4.425593e+01 2.4723780 *

13059) SatDom_GPP_max..80>=0.0125 273 1.481735e+02 3.2412090 *

3265) BOMDom_min_temperature_var_.10< 2.8565 290 1.473367e+02 3.5160690 *

1633) BOMDom_min_temperature_mean_.30>=13.24 1177 8.893489e+02 3.0644600

3266) SatDom_Latent_heat_min..30< 933643.4 564 3.249874e+02 2.5831740

6532) SatDom_PsnNet_var..20>=1.492901e-06 113 9.053711e+01 1.6118580 *

6533) SatDom_PsnNet_var..20< 1.492901e-06 451 1.011284e+02 2.8265410

13066) SatDom_Emis31_var.0< 1.027778e-06 334 5.532820e+01 2.6900600 *

13067) SatDom_Emis31_var.0>=1.027778e-06 117 2.181837e+01 3.2161540 *

3267) SatDom_Latent_heat_min..30>=933643.4 613 3.135180e+02 3.5072760

6534) SatDom_EVI_var..70< 3.236979e-06 416 1.744433e+02 3.2130050

13068) SatDom_EVI_var..50< 1.871707e-06 107 2.056506e+01 2.6346730 *

13069) SatDom_EVI_var..50>=1.871707e-06 309 1.056976e+02 3.4132690 *

6535) SatDom_EVI_var..70>=3.236979e-06 197 2.698066e+01 4.1286800 *

817) SatDom_Reflect_b6_var..70< 2.53066e-05 853 8.799830e+02 3.3741970

1634) METADom_Fert_rotation_minus_1_P< -2.278993e-17 278 8.985526e+01 2.4263310 *

1635) METADom_Fert_rotation_minus_1_P>=-2.278993e-17 575 4.196007e+02 3.8324700

3270) SatDom_MIR_max..60< 0.3378031 425 1.650785e+02 3.4974350

6540) SatDom_blue_band_var..20>=4.031864e-06 146 3.101098e+01 2.9043150 *

6541) SatDom_blue_band_var..20< 4.031864e-06 279 5.582857e+01 3.8078140 *

3271) SatDom_MIR_max..60>=0.3378031 150 7.165155e+01 4.7817330 *

409) SatDom_FPAR_var..20>=0.001451667 734 6.802191e+02 3.7717300

818) SatDom_FPAR_min..70< 0.14625 244 7.122911e+01 2.8534430 *

819) SatDom_FPAR_min..70>=0.14625 490 3.007796e+02 4.2290000

1638) SatDom_MIR..60< 9.164228 141 4.108976e+01 3.5721280 *

1639) SatDom_MIR..60>=9.164228 349 1.742714e+02 4.4943840

3278) METADom_Chem_rotation_minus_1_SPINREPORTED_NAKER>=-1.755323e-17 194 2.627100e+01 4.2700000 *

3279) METADom_Chem_rotation_minus_1_SPINREPORTED_NAKER< -1.755323e-17 155 1.260077e+02 4.7752260 *

205) SatDom_Reflect_b5_mean..20< 0.2969987 1242 1.269962e+03 3.8416830

410) METADom_Chem_rotation_minus_2_TOPIK>=-6.938894e-19 537 2.369657e+02 3.1345070

820) SatDom_NDVI_var..70>=3.573756e-06 279 7.831925e+01 2.7656630 *

821) SatDom_NDVI_var..70< 3.573756e-06 258 7.964357e+01 3.5333720 *

411) METADom_Chem_rotation_minus_2_TOPIK< -6.938894e-19 705 5.598863e+02 4.3803400

822) SatDom_LST_day_var..50>=4.863918 298 1.534954e+02 3.9266440 *

823) SatDom_LST_day_var..50< 4.863918 407 3.001379e+02 4.7125310 *

103) METADom_Fertiliser_time_MAP>=-3.073846 3243 4.204318e+03 3.7695810

206) SatDom_Reflect_b1_max..10>=0.1644 319 2.998063e+02 2.5171160

412) METADom_Chem_rotation_minus_1_TARGA< 0.01113129 205 8.728568e+01 2.0460980 *

413) METADom_Chem_rotation_minus_1_TARGA>=0.01113129 114 8.525376e+01 3.3641230 *

207) SatDom_Reflect_b1_max..10< 0.1644 2924 3.349514e+03 3.9062210

414) METADom_Chem_rotation_minus_1_STARANE< 0.0004098361 2701 2.835669e+03 3.7982670

828) BOMDom_solar_exposure_min_.80>=17.8 778 5.131890e+02 3.1987660

1656) SatDom_Emis32_mean..30>=0.98635 286 6.922542e+01 2.4463640 *

1657) SatDom_Emis32_mean..30< 0.98635 492 1.879395e+02 3.6361380

3314) SatDom_Reflect_b6_var..60>=3.316174e-05 331 6.846487e+01 3.3749240 *

3315) SatDom_Reflect_b6_var..60< 3.316174e-05 161 5.045708e+01 4.1731680 *

829) BOMDom_solar_exposure_min_.80< 17.8 1923 1.929740e+03 4.0408110

1658) SatDom_red_band_max..40< 0.168225 1509 1.327111e+03 3.8249830

3316) SatDom_Reflect_b7_min..70< 0.2733313 255 5.428182e+01 2.9035290 *

3317) SatDom_Reflect_b7_min..70>=0.2733313 1254 1.012286e+03 4.0123600

6634) SatDom_LAI..30>=15.825 1088 8.063762e+02 3.8750000

13268) SatDom_Latent_heat.0< 1.762355e+08 952 6.234498e+02 3.7854940

26536) SatDom_blue_band_mean.0>=0.05709844 126 5.827655e+01 3.2262700 *

26537) SatDom_blue_band_mean.0< 0.05709844 826 5.197583e+02 3.8707990 *

13269) SatDom_Latent_heat.0>=1.762355e+08 136 1.219118e+02 4.5015440 *

6635) SatDom_LAI..30< 15.825 166 5.083523e+01 4.9126510 *

1659) SatDom_red_band_max..40>=0.168225 414 2.761292e+02 4.8274880

3318) METADom_Fert_rotation_minus_2_SOA< 0.007665497 278 1.348937e+02 4.5085250 *

3319) METADom_Fert_rotation_minus_2_SOA>=0.007665497 136 5.513886e+01 5.4794850 *

415) METADom_Chem_rotation_minus_1_STARANE>=0.0004098361 223 1.011094e+02 5.2137670 *

13) METADom_Year>=2015.5 4451 1.368908e+04 3.9011930

26) METADom_Phos_fert< 17.25 3159 5.839820e+03 3.2284170

52) METADom_Fert_rotation_minus_1_EXTRA>=-5.850355e-17 2108 2.876338e+03 2.6792550

104) SatDom_EvapoTrans_mean..20< 7.40875 1360 1.455896e+03 2.2259850

208) SatDom_Aero_q.0< 7 1191 8.699671e+02 2.0042650

416) BOMDom_min_temperature_max_.50>=14.15 1034 4.705249e+02 1.7872340

832) METADom_Chem_rotation_minus_1_HAMMER< 0.005171871 676 2.420715e+02 1.4996750

1664) SatDom_LST_night_var..70>=1.258607 554 9.559899e+01 1.3201440

3328) METADom_Fertiliser_time_MAP< -6.883083 396 3.515474e+01 1.1291920 *

3329) METADom_Fertiliser_time_MAP>=-6.883083 158 9.815347e+00 1.7987340 *

1665) SatDom_LST_night_var..70< 1.258607 122 4.753265e+01 2.3149180 *

833) METADom_Chem_rotation_minus_1_HAMMER>=0.005171871 358 6.700278e+01 2.3302230 *

417) BOMDom_min_temperature_max_.50< 14.15 157 2.997383e+01 3.4336310 *

209) SatDom_Aero_q.0>=7 169 1.147635e+02 3.7885210 *

105) SatDom_EvapoTrans_mean..20>=7.40875 748 6.329939e+02 3.5033820

210) SatDom_Latent_heat_min..70< 496250 395 1.676261e+02 2.9564300

420) SatDom_Reflect_b6_var..30< 6.345502e-05 115 4.170456e+01 2.2578260 *

421) SatDom_Reflect_b6_var..30>=6.345502e-05 280 4.674444e+01 3.2433570 *

211) SatDom_Latent_heat_min..70>=496250 353 2.149748e+02 4.1154110

422) SatDom_Pot_latent_heat_max..40< 1.22425e+07 117 3.763111e+01 3.4053850 *

423) SatDom_Pot_latent_heat_max..40>=1.22425e+07 236 8.911752e+01 4.4674150 *

53) METADom_Fert_rotation_minus_1_EXTRA< -5.850355e-17 1051 1.052668e+03 4.3298760

106) METADom_Chem_rotation_minus_1_DUAL_GOLD>=0.004336695 242 1.233669e+02 3.1424790 *

107) METADom_Chem_rotation_minus_1_DUAL_GOLD< 0.004336695 809 4.860382e+02 4.6850680

214) BOMDom_max_temperature_mean_.30< 27.355 317 1.088147e+02 4.1395580

428) SatDom_Reflect_b1_var..70< 1.758052e-05 130 3.158500e+01 3.6772310 *

429) SatDom_Reflect_b1_var..70>=1.758052e-05 187 3.012543e+01 4.4609630 *

215) BOMDom_max_temperature_mean_.30>=27.355 492 2.221107e+02 5.0365450

430) SatDom_GPP_var..30< 5.947708e-07 345 1.231656e+02 4.8031010 *

431) SatDom_GPP_var..30>=5.947708e-07 147 3.601923e+01 5.5844220 *

27) METADom_Phos_fert>=17.25 1292 2.923359e+03 5.5461610

54) METADom_Year>=2016.5 284 2.053995e+02 3.7417250 *

55) METADom_Year< 2016.5 1008 1.532728e+03 6.0545540

110) SatDom_LST_day_var..40>=6.363743 615 5.096998e+02 5.4639020

220) METADom_Chem_rotation_minus_2_ROUNDUP_POWERMAX< 0.006221173 138 3.227515e+01 4.4227540 *

221) METADom_Chem_rotation_minus_2_ROUNDUP_POWERMAX>=0.006221173 477 2.845561e+02 5.7651150

442) BOMDom_min_temperature_max_.10>=15.75 221 6.305105e+01 5.3708140 *

443) BOMDom_min_temperature_max_.10< 15.75 256 1.574835e+02 6.1055080 *

111) SatDom_LST_day_var..40< 6.363743 393 4.727212e+02 6.9788550

222) METADom_Fert_rotation_minus_2_PLAIN>=2.63678e-17 257 1.831224e+02 6.5207390 *

223) METADom_Fert_rotation_minus_2_PLAIN< 2.63678e-17 136 1.337380e+02 7.8445590 *

7) SatDom_Reflect_b7.0< 24.17471 39280 8.242715e+04 3.8095150

14) METADom_Phos_fert< 17.16 19521 3.061495e+04 3.3814240

28) SatDom_EvapoTrans_min..60>=1.1125 18996 2.613362e+04 3.3188470

56) SatDom_GPP..80< 0.14755 11060 1.374385e+04 3.0680240

112) METADom_ESP_60cm.test_60cm>=10.61598 1725 2.180203e+03 2.3831590

224) SatDom_PsnNet_max..20>=0.0102 283 1.813204e+01 0.9195406 *

225) SatDom_PsnNet_max..20< 0.0102 1442 1.436857e+03 2.6704020

450) SatDom_red_band..80>=1.307647 1209 8.287268e+02 2.4475770

900) METADom_Fert_rotation_minus_2_PLAIN>=0.0006756757 484 1.991418e+02 1.8880370

1800) METADom_Chem_rotation_minus_1_BOXERGOLD< -2.808084e-17 223 2.793041e+01 1.4437220 *

1801) METADom_Chem_rotation_minus_1_BOXERGOLD>=-2.808084e-17 261 8.957347e+01 2.2676630 *

901) METADom_Fert_rotation_minus_2_PLAIN< 0.0006756757 725 3.768910e+02 2.8211170

1802) SatDom_LST_day..60>=1687.064 602 2.311380e+02 2.6373260

3604) SatDom_NDVI_var..60< 5.664914e-06 181 6.300483e+01 2.0414360 *

3605) SatDom_NDVI_var..60>=5.664914e-06 421 7.623140e+01 2.8935150 *

1803) SatDom_LST_day..60< 1687.064 123 2.589115e+01 3.7206500 *

451) SatDom_red_band..80< 1.307647 233 2.366242e+02 3.8266090 *

113) METADom_ESP_60cm.test_60cm< 10.61598 9335 1.060505e+04 3.1945780

226) METADom_Chem_rotation_minus_1_POWER< -2.725684e-17 3718 3.730480e+03 2.8053010

452) SatDom_Reflect_b5_min..10>=0.2338875 2234 1.671930e+03 2.4611680

904) METADom_Chem_rotation_minus_2_LVE< 0.03836598 1576 7.579776e+02 2.1534960

1808) SatDom_Latent_heat_max..40>=601209.7 1452 4.591348e+02 2.0310470

3616) SatDom_MIR..50< 10.86607 694 1.478874e+02 1.7307490

7232) SatDom_Reflect_b5_var..60>=3.940002e-05 496 7.518312e+01 1.5627620

14464) BOMDom_Latitude>=-32.95154 295 2.686261e+01 1.3809830 *

14465) BOMDom_Latitude< -32.95154 201 2.426606e+01 1.8295520 *

7233) SatDom_Reflect_b5_var..60< 3.940002e-05 198 2.364421e+01 2.1515660 *

3617) SatDom_MIR..50>=10.86607 758 1.913636e+02 2.3059890

7234) SatDom_Reflect_b5_mean..30< 0.2643575 156 2.023807e+01 1.7791030 *

7235) SatDom_Reflect_b5_mean..30>=0.2643575 602 1.165960e+02 2.4425250

14470) SatDom_LST_day_var..20>=2.774109 474 6.516580e+01 2.3333760 *

14471) SatDom_LST_day_var..20< 2.774109 128 2.487142e+01 2.8467190 *

1809) SatDom_Latent_heat_max..40< 601209.7 124 2.213962e+01 3.5873390 *

905) METADom_Chem_rotation_minus_2_LVE>=0.03836598 658 4.074400e+02 3.1980850

1810) SatDom_EVI_var..70< 7.854485e-06 371 1.480197e+02 2.7403500

3620) METADom_Chem_rotation_minus_2_LVE< 0.07967318 220 3.814553e+01 2.3825000 *

3621) METADom_Chem_rotation_minus_2_LVE>=0.07967318 151 4.065555e+01 3.2617220 *

1811) SatDom_EVI_var..70>=7.854485e-06 287 8.120479e+01 3.7897910 *

453) SatDom_Reflect_b5_min..10< 0.2338875 1484 1.395707e+03 3.3233560

906) SatDom_PsnNet_var..30>=1.731988e-07 798 5.659926e+02 2.8153630

1812) METADom_Chem_rotation_minus_1_ALLY>=0.04435775 529 2.428400e+02 2.4456900

3624) SatDom_Reflect_b3_var..70< 4.935772e-06 170 2.020772e+01 1.8120000 *

3625) SatDom_Reflect_b3_var..70>=4.935772e-06 359 1.220402e+02 2.7457660

7250) METADom_Fert_rotation_minus_1_TEN_TO_TWENTY_TWO< 0.002063944 218 4.199974e+01 2.4982570 *

7251) METADom_Fert_rotation_minus_1_TEN_TO_TWENTY_TWO>=0.002063944 141 4.603766e+01 3.1284400 *

1813) METADom_Chem_rotation_minus_1_ALLY< 0.04435775 269 1.086944e+02 3.5423420 *

907) SatDom_PsnNet_var..30< 1.731988e-07 686 3.842356e+02 3.9142860

1814) SatDom_Latent_heat_max..70< 951250 177 6.443223e+01 3.0839550 *

1815) SatDom_Latent_heat_max..70>=951250 509 1.553351e+02 4.2030260

3630) BOMDom_min_temperature_var_.10< 18.41139 354 6.332987e+01 3.9811020 *

3631) BOMDom_min_temperature_var_.10>=18.41139 155 3.475240e+01 4.7098710 *

227) METADom_Chem_rotation_minus_1_POWER>=-2.725684e-17 5617 5.938217e+03 3.4522490

454) METADom_HALOXYFOP_R>=7.837484 762 6.468545e+02 2.6466930

908) BOMDom_solar_exposure_min_.10>=5.9 494 2.958281e+02 2.2592910

1816) METADom_MCPA_AMINE>=18.09825 176 1.772702e+01 1.5635230 *

1817) METADom_MCPA_AMINE< 18.09825 318 1.457454e+02 2.6443710

3634) METADom_X2_4_DB< -2.479128e-14 171 2.733425e+01 2.2571930 *

3635) METADom_X2_4_DB>=-2.479128e-14 147 6.295787e+01 3.0947620 *

909) BOMDom_solar_exposure_min_.10< 5.9 268 1.402269e+02 3.3607840 *

455) METADom_HALOXYFOP_R< 7.837484 4855 4.719277e+03 3.5786820

910) SatDom_FPAR..60>=6.32 1230 9.289900e+02 3.0446180

1820) SatDom_NIR_var..80< 3.99681e-05 992 4.057731e+02 2.7900710

3640) METADom_Nitrogen_fert< 20.48267 476 1.462881e+02 2.4740970

7280) SatDom_Pot_latent_heat..20< 8.939127e+08 181 3.167547e+01 2.0703870 *

7281) SatDom_Pot_latent_heat..20>=8.939127e+08 295 6.701315e+01 2.7217970 *

3641) METADom_Nitrogen_fert>=20.48267 516 1.681218e+02 3.0815500

7282) BOMDom_solar_exposure_var_.10>=12.71189 134 2.135208e+01 2.6057460 *

7283) BOMDom_solar_exposure_var_.10< 12.71189 382 1.057920e+02 3.2484550

14566) SatDom_Pot_EvapoTrans_max.0>=37.4125 279 4.453188e+01 3.0979930 *

14567) SatDom_Pot_EvapoTrans_max.0< 37.4125 103 3.783467e+01 3.6560190 *

1821) SatDom_NIR_var..80>=3.99681e-05 238 1.910343e+02 4.1055880 *

911) SatDom_FPAR..60< 6.32 3625 3.320422e+03 3.7598950

1822) SatDom_Reflect_b7_var..50< 1.783232e-05 701 5.444166e+02 3.0471040

3644) BOMDom_rainfall_var_.10< 14.69472 495 2.638354e+02 2.7234550

7288) SatDom_Latent_heat_max..80< 910000 102 5.461285e+01 1.8259800 *

7289) SatDom_Latent_heat_max..80>=910000 393 1.057425e+02 2.9563870

14578) METADom_Chem_rotation_minus_2_ROUNDUP_MAX< 0.0003618699 267 4.433897e+01 2.7356180 *

14579) METADom_Chem_rotation_minus_2_ROUNDUP_MAX>=0.0003618699 126 2.081447e+01 3.4242060 *

3645) BOMDom_rainfall_var_.10>=14.69472 206 1.041375e+02 3.8248060 *

1823) SatDom_Reflect_b7_var..50>=1.783232e-05 2924 2.334463e+03 3.9307800

3646) BOMDom_min_temperature_mean_.20>=7.91 2203 1.578231e+03 3.7181530

7292) SatDom_FPAR_var..70>=5.583333e-05 1838 1.034021e+03 3.5808870

14584) BOMDom_min_temperature_var_.40< 5.109111 523 2.786390e+02 3.1427530

29168) METADom_Fert_rotation_minus_2_N>=0.007494354 140 6.772751e+01 2.4914290 *

29169) METADom_Fert_rotation_minus_2_N< 0.007494354 383 1.298105e+02 3.3808360

58338) SatDom_red_band_min.0< 0.0792375 209 3.730986e+01 3.0958850 *

58339) SatDom_red_band_min.0>=0.0792375 174 5.514692e+01 3.7231030 *

14585) BOMDom_min_temperature_var_.40>=5.109111 1315 6.150570e+02 3.7551410

29170) SatDom_Reflect_b4..60>=3.539431 222 1.092722e+02 3.1682430 *

29171) SatDom_Reflect_b4..60< 3.539431 1093 4.137859e+02 3.8743460

58342) SatDom_Pot_latent_heat..80< 2.154525e+08 960 2.634591e+02 3.7699900

116684) SatDom_Pot_latent_heat_var..20>=1.128443e+12 514 1.173864e+02 3.5912450 *

116685) SatDom_Pot_latent_heat_var..20< 1.128443e+12 446 1.107247e+02 3.9759870 *

58343) SatDom_Pot_latent_heat..80>=2.154525e+08 133 6.441023e+01 4.6275940 *

7293) SatDom_FPAR_var..70< 5.583333e-05 365 3.351886e+02 4.4093700

14586) SatDom_EvapoTrans_min.0>=18433.53 101 2.418696e+01 3.4094060 *

14587) SatDom_EvapoTrans_min.0< 18433.53 264 1.713715e+02 4.7919320 *

3647) BOMDom_min_temperature_mean_.20< 7.91 721 3.523125e+02 4.5804580

7294) SatDom_EvapoTrans_mean..60>=3.928125 568 2.249147e+02 4.4237500

14588) SatDom_GPP_mean..10>=0.004304375 412 1.018511e+02 4.2768200 *

14589) SatDom_GPP_mean..10< 0.004304375 156 9.067890e+01 4.8117950 *

7295) SatDom_EvapoTrans_mean..60< 3.928125 153 6.166644e+01 5.1622220 *

57) SatDom_GPP..80>=0.14755 7936 1.072425e+04 3.6684060

114) SatDom_Reflect_b5_min..40>=0.2708813 3166 4.046318e+03 3.2283800

228) BOMDom_rainfall_mean_.50< 1.75 2450 2.710150e+03 3.0088780

456) BOMDom_rainfall_csum_.80>=9.4 1064 7.344791e+02 2.5029890

912) SatDom_Reflect_b7_var..80< 0.0001872413 660 3.171319e+02 2.1662580

1824) METADom_Fert_rotation_minus_2_EIGHTEEN_TO_TWENTY< 0.000676819 357 8.612206e+01 1.7708120

3648) SatDom_EvapoTrans_min..60>=14.5375 132 2.180955e+01 1.3460610 *

3649) SatDom_EvapoTrans_min..60< 14.5375 225 2.652660e+01 2.0200000 *

1825) METADom_Fert_rotation_minus_2_EIGHTEEN_TO_TWENTY>=0.000676819 303 1.094074e+02 2.6321780

3650) SatDom_Reflect_b5_max..80< 0.34065 156 2.219342e+01 2.2792950 *

3651) SatDom_Reflect_b5_max..80>=0.34065 147 4.717227e+01 3.0066670 *

913) SatDom_Reflect_b7_var..80>=0.0001872413 404 2.202544e+02 3.0530940

1826) SatDom_red_band.0>=11.63981 103 1.510577e+01 2.1949510 *

1827) SatDom_red_band.0< 11.63981 301 1.033432e+02 3.3467440

3654) METADom_Chem_rotation_minus_2_ROUNDUP>=0.1000209 189 2.270448e+01 3.1073540 *

3655) METADom_Chem_rotation_minus_2_ROUNDUP< 0.1000209 112 5.153014e+01 3.7507140 *

457) BOMDom_rainfall_csum_.80< 9.4 1386 1.494328e+03 3.3972370

914) SatDom_Reflect_b6_mean..40< 0.3392569 786 5.466635e+02 2.9664890

1828) SatDom_MIR_mean..40>=0.1837422 568 2.298376e+02 2.6915320

3656) SatDom_Reflect_b6_var..80>=0.0002614661 291 7.313825e+01 2.3411000 *

3657) SatDom_Reflect_b6_var..80< 0.0002614661 277 8.342207e+01 3.0596750 *

1829) SatDom_MIR_mean..40< 0.1837422 218 1.620001e+02 3.6828900 *

915) SatDom_Reflect_b6_mean..40>=0.3392569 600 6.107797e+02 3.9615170

1830) SatDom_Reflect_b6_mean..50>=0.3628369 221 1.370983e+02 3.1683260 *

1831) SatDom_Reflect_b6_mean..50< 0.3628369 379 2.535613e+02 4.4240370

3662) METADom_Chem_rotation_minus_2_SELECT< 0.08917317 226 1.288639e+02 4.1065490 *

3663) METADom_Chem_rotation_minus_2_SELECT>=0.08917317 153 6.826722e+01 4.8930070 *

229) BOMDom_rainfall_mean_.50>=1.75 716 8.142028e+02 3.9794690

458) SatDom_Reflect_b7_mean..70>=0.2572187 304 1.911628e+02 3.1324670

916) SatDom_MIR_var..10>=0.0001109178 128 4.795500e+01 2.5575000 *

917) SatDom_MIR_var..10< 0.0001109178 176 7.011803e+01 3.5506250 *

459) SatDom_Reflect_b7_mean..70< 0.2572187 412 2.440232e+02 4.6044420

918) SatDom_Reflect_b1_mean..80< 0.1024569 204 8.176298e+01 4.1397060 *

919) SatDom_Reflect_b1_mean..80>=0.1024569 208 7.498789e+01 5.0602400 *

115) SatDom_Reflect_b5_min..40< 0.2708813 4770 5.658046e+03 3.9604650

230) METADom_Chem_rotation_minus_2_ROUNDUP_POWERMAX< 0.1117911 4298 4.550797e+03 3.8348930

460) SatDom_Reflect_b7_mean.0< 0.1017831 342 2.159574e+02 2.6074850

920) SatDom_NIR_max..40< 0.1610531 148 1.818996e+01 1.8805410 *

921) SatDom_NIR_max..40>=0.1610531 194 5.989138e+01 3.1620620 *

461) SatDom_Reflect_b7_mean.0>=0.1017831 3956 3.775064e+03 3.9410040

922) METADom_ESP_10cm.test_10cm>=4.15 344 2.010574e+02 2.8788950

1844) BOMDom_rainfall_csum_0>=144.55 115 4.999552e+01 2.2075650 *

1845) BOMDom_rainfall_csum_0< 144.55 229 7.320568e+01 3.2160260 *

923) METADom_ESP_10cm.test_10cm< 4.15 3612 3.148992e+03 4.0421570

1846) SatDom_blue_band_min..40>=0.03363438 3045 2.487301e+03 3.9242530

3692) SatDom_Reflect_b7_var.0>=9.926866e-06 2452 1.895285e+03 3.7873980

7384) SatDom_Reflect_b5_var.0< 0.000104433 1115 7.413650e+02 3.4559100

14768) SatDom_Reflect_b5_min..70>=0.207475 894 4.110960e+02 3.2688590

29536) SatDom_Reflect_b3_mean..70>=0.05193 614 2.292345e+02 3.0430780

59072) SatDom_MIR_max..30< 0.2115406 473 1.424789e+02 2.8770400

118144) BOMDom_Longitude< 151.0605 356 7.572309e+01 2.7160670 *

118145) BOMDom_Longitude>=151.0605 117 2.946253e+01 3.3668380 *

59073) SatDom_MIR_max..30>=0.2115406 141 2.997170e+01 3.6000710 *

29537) SatDom_Reflect_b3_mean..70< 0.05193 280 8.192550e+01 3.7639640 *

14769) SatDom_Reflect_b5_min..70< 0.207475 221 1.724564e+02 4.2125790 *

7385) SatDom_Reflect_b5_var.0>=0.000104433 1337 9.292222e+02 4.0638440

14770) METADom_Fert_rotation_minus_1_PHOS< -1.048424e-17 589 2.142877e+02 3.6558570

29540) SatDom_red_band_var..30>=2.660856e-06 272 6.863758e+01 3.3147060 *

29541) SatDom_red_band_var..30< 2.660856e-06 317 8.683086e+01 3.9485800 *

14771) METADom_Fert_rotation_minus_1_PHOS>=-1.048424e-17 748 5.396927e+02 4.3851070

29542) SatDom_Reflect_b6_max..50< 0.2647625 360 2.879340e+02 4.0453610

59084) METADom_Chem_rotation_minus_2_TWO_FOURD>=0.1220828 105 4.631754e+01 3.5307620 *

59085) METADom_Chem_rotation_minus_2_TWO_FOURD< 0.1220828 255 2.023619e+02 4.2572550 *

29543) SatDom_Reflect_b6_max..50>=0.2647625 388 1.716499e+02 4.7003350

59086) BOMDom_Longitude>=148.7419 254 6.552542e+01 4.4786220 *

59087) BOMDom_Longitude< 148.7419 134 6.997155e+01 5.1205970 *

3693) SatDom_Reflect_b7_var.0< 9.926866e-06 593 3.561996e+02 4.4901350

7386) SatDom_Reflect_b1..60< 3.039756 293 1.064059e+02 4.0650170 *

7387) SatDom_Reflect_b1..60>=3.039756 300 1.451243e+02 4.9053330 *

1847) SatDom_blue_band_min..40< 0.03363438 567 3.920365e+02 4.6753440

3694) SatDom_LST_night_var..20< 5.920159 171 1.036643e+02 3.8585960 *

3695) SatDom_LST_night_var..20>=5.920159 396 1.250447e+02 5.0280300

7390) BOMDom_Latitude< -34.54333 102 1.779020e+01 4.5731370 *

7391) BOMDom_Latitude>=-34.54333 294 7.882514e+01 5.1858500 *

231) METADom_Chem_rotation_minus_2_ROUNDUP_POWERMAX>=0.1117911 472 4.223422e+02 5.1039190

462) SatDom_blue_band_max..30< 0.04949688 278 9.462562e+01 4.4943170 *

463) SatDom_blue_band_max..30>=0.04949688 194 7.636646e+01 5.9774740 *

29) SatDom_EvapoTrans_min..60< 1.1125 525 1.715438e+03 5.6456380

58) METADom_Chem_rotation_minus_3_UNKNOWN>=0.5 424 4.198819e+02 4.9544810

116) METADom_Fertiliser_time_MAP< -2.465005 194 1.657604e+02 4.2901030 *

117) METADom_Fertiliser_time_MAP>=-2.465005 230 9.626215e+01 5.5148700 *

59) METADom_Chem_rotation_minus_3_UNKNOWN< 0.5 101 2.427285e+02 8.5471290 *

15) METADom_Phos_fert>=17.16 19759 4.470037e+04 4.2324500

30) METADom_Nitrogen_fert_repeat_3< 31.05 18967 3.773467e+04 4.1461000

60) METADom_Fert_rotation_minus_2_THIRTY_TWO_TO_NINE< 0.1 18049 3.164273e+04 4.0579420

120) SatDom_Latent_heat_max..10< 2285000 12277 2.100479e+04 3.8503940

240) METADom_total_P_10cm.test_10cm< 19.5 1294 1.141092e+03 2.8276120

480) BOMDom_rainfall_mean_.40< 0.6 635 3.147512e+02 2.2691810

960) METADom_Fert_rotation_minus_2_SOA< 0.003833346 159 2.616088e+01 1.4391190 *

961) METADom_Fert_rotation_minus_2_SOA>=0.003833346 476 1.424451e+02 2.5464500

1922) SatDom_NDVI_min..10< 0.3017125 348 5.681932e+01 2.3723850 *

1923) SatDom_NDVI_min..10>=0.3017125 128 4.641579e+01 3.0196880 *

481) BOMDom_rainfall_mean_.40>=0.6 659 4.375087e+02 3.3657060

962) SatDom_LST_night_var..70>=4.912232 443 1.335089e+02 2.9248980

1924) BOMDom_min_temperature_max_.80>=20.85 152 2.995409e+01 2.4383550 *

1925) BOMDom_min_temperature_max_.80< 20.85 291 4.877793e+01 3.1790380 *

963) SatDom_LST_night_var..70< 4.912232 216 4.137689e+01 4.2697690 *

241) METADom_total_P_10cm.test_10cm>=19.5 10983 1.835058e+04 3.9708970

482) SatDom_LST_day_mean..30< 46.7018 7641 1.152749e+04 3.7180030

964) METADom_Chem_rotation_minus_2_GOAL>=0.01100636 745 6.743295e+02 2.4992890

1928) SatDom_Pot_latent_heat_max.0>=5003750 641 2.394234e+02 2.2344770

3856) METADom_Chem_rotation_minus_2_BALANCE< -1.669671e-17 316 4.582248e+01 1.7952530 *

3857) METADom_Chem_rotation_minus_2_BALANCE>=-1.669671e-17 325 7.336503e+01 2.6615380

7714) SatDom_LST_day..20< 3249.513 203 2.404245e+01 2.4335470 *

7715) SatDom_LST_day..20>=3249.513 122 2.121280e+01 3.0409020 *

1929) SatDom_Pot_latent_heat_max.0< 5003750 104 1.129077e+02 4.1314420 *

965) METADom_Chem_rotation_minus_2_GOAL< 0.01100636 6896 9.627096e+03 3.8496650

1930) SatDom_EVI_var..40< 2.033691e-06 1491 1.496295e+03 3.1258280

3860) SatDom_Emis31_mean..20>=0.98326 965 7.825738e+02 2.7333260

7720) SatDom_LST_night_min..40< 22.4 144 6.129413e+01 1.5881940 *

7721) SatDom_LST_night_min..40>=22.4 821 4.993284e+02 2.9341780

15442) SatDom_LST_day_mean..30>=42.88429 389 1.697293e+02 2.4139850

30884) SatDom_MixClouds_csum..30< 7.5 218 3.939531e+01 2.1079820 *

30885) SatDom_MixClouds_csum..30>=7.5 171 8.389733e+01 2.8040940 *

15443) SatDom_LST_day_mean..30< 42.88429 432 1.295491e+02 3.4025930

30886) SatDom_Pot_latent_heat.0< 6.998649e+08 244 6.767194e+01 3.1532380 *

30887) SatDom_Pot_latent_heat.0>=6.998649e+08 188 2.701522e+01 3.7262230 *

3861) SatDom_Emis31_mean..20< 0.98326 526 2.923129e+02 3.8459130

7722) SatDom_NIR_var..30< 6.837874e-06 213 5.966790e+01 3.2699530 *

7723) SatDom_NIR_var..30>=6.837874e-06 313 1.139029e+02 4.2378590 *

1931) SatDom_EVI_var..40>=2.033691e-06 5405 7.134110e+03 4.0493400

3862) SatDom_blue_band_var..80>=1.754557e-08 5267 6.135971e+03 3.9818550

7724) SatDom_GPP_var..80< 7.224696e-07 2158 1.950667e+03 3.6208900

15448) METADom_Chem_rotation_minus_2_AFFINITY< 0.00832273 953 7.111530e+02 3.1872610

30896) SatDom_blue_band_var..70>=4.827717e-06 172 1.120080e+02 2.3202330 *

30897) SatDom_blue_band_var..70< 4.827717e-06 781 4.413703e+02 3.3782070

61794) SatDom_Emis32_mean.0< 0.98665 596 3.062685e+02 3.2061580

123588) METADom_sub_cropWheat>=0.5 360 1.394973e+02 3.0003890

247176) BOMDom_min_temperature_csum_.80< 152.35 221 5.489085e+01 2.7287330 *

247177) BOMDom_min_temperature_csum_.80>=152.35 139 4.236706e+01 3.4323020 *

123589) METADom_sub_cropWheat< 0.5 236 1.282769e+02 3.5200420 *

61795) SatDom_Emis32_mean.0>=0.98665 185 6.062286e+01 3.9324860 *

15449) METADom_Chem_rotation_minus_2_AFFINITY>=0.00832273 1205 9.185973e+02 3.9638340

30898) SatDom_PsnNet_var..60< 1.65941e-07 337 1.194953e+02 3.2222550

61796) METADom_Chem_rotation_minus_1_LONTREL< 0.03033847 201 6.314699e+01 2.9554230 *

61797) METADom_Chem_rotation_minus_1_LONTREL>=0.03033847 136 2.088624e+01 3.6166180 *

30899) SatDom_PsnNet_var..60>=1.65941e-07 868 5.418185e+02 4.2517510

61798) BOMDom_max_temperature_min_.30>=15.85 449 2.228644e+02 3.9301780

123596) SatDom_LST_night_mean..50>=30.159 245 1.064542e+02 3.5925310 *

123597) SatDom_LST_night_mean..50< 30.159 204 5.493360e+01 4.3356860 *

61799) BOMDom_max_temperature_min_.30< 15.85 419 2.227683e+02 4.5963480

123598) BOMDom_Longitude>=147.8297 110 2.312586e+01 4.1806360 *

123599) BOMDom_Longitude< 147.8297 309 1.738654e+02 4.7443370

247198) SatDom_Reflect_b2_mean..50>=0.2316794 198 6.043973e+01 4.5046970 *

247199) SatDom_Reflect_b2_mean..50< 0.2316794 111 8.177244e+01 5.1718020 *

7725) SatDom_GPP_var..80>=7.224696e-07 3109 3.708955e+03 4.2324060

15450) SatDom_FPAR_mean..30>=0.192125 2635 2.695255e+03 4.0273850

30900) SatDom_Reflect_b6_mean..50>=0.3636656 557 3.662417e+02 3.1250630

61800) METADom_Chem_rotation_minus_1_MAP< 0.003235 253 1.134783e+02 2.5698810 *

61801) METADom_Chem_rotation_minus_1_MAP>=0.003235 304 1.098833e+02 3.5871050

123602) METADom_Chem_rotation_minus_1_BOXERGOLD< -3.729655e-17 187 2.085346e+01 3.2188770 *

123603) METADom_Chem_rotation_minus_1_BOXERGOLD>=-3.729655e-17 117 2.314828e+01 4.1756410 *

30901) SatDom_Reflect_b6_mean..50< 0.3636656 2078 1.753952e+03 4.2692490

61802) SatDom_EvapoTrans_min..30< 7.4875 1511 1.038257e+03 4.0496290

123604) SatDom_FPAR_min..50>=0.265 546 2.461377e+02 3.6442490

247208) SatDom_LST_day_max..50>=49.32 329 8.355860e+01 3.3547110

494416) METADom_Fert_rotation_minus_1_THIRTY_TWO_TO_TEN>=-2.012279e-17 107 2.218136e+01 2.9814950 *

494417) METADom_Fert_rotation_minus_1_THIRTY_TWO_TO_TEN< -2.012279e-17 222 3.928971e+01 3.5345950 *

247209) SatDom_LST_day_max..50< 49.32 217 9.318234e+01 4.0832260 *

123605) SatDom_FPAR_min..50< 0.265 965 6.516261e+02 4.2789950

247210) SatDom_LAI_min..70< 0.325 562 2.362989e+02 3.9916900

494420) SatDom_GPP_var..80>=1.678563e-06 206 3.847716e+01 3.6614560 *

494421) SatDom_GPP_var..80< 1.678563e-06 356 1.623569e+02 4.1827810 *

247211) SatDom_LAI_min..70>=0.325 403 3.042454e+02 4.6796530

494422) SatDom_LST_night_var..40>=3.328452 173 1.739676e+02 4.2600000 *

494423) SatDom_LST_night_var..40< 3.328452 230 7.689473e+01 4.9953040 *

61803) SatDom_EvapoTrans_min..30>=7.4875 567 4.485976e+02 4.8545150

123606) SatDom_Reflect_b1_var..40< 2.635254e-05 376 2.428136e+02 4.5306120

247212) MANDom_Rows_per_plot_6< 0.5 235 1.488603e+02 4.3422550 *

247213) MANDom_Rows_per_plot_6>=0.5 141 7.172030e+01 4.8445390 *

123607) SatDom_Reflect_b1_var..40>=2.635254e-05 191 8.868102e+01 5.4921470 *

15451) SatDom_FPAR_mean..30< 0.192125 474 2.872291e+02 5.3721310

30902) METADom_Chem_rotation_minus_2_SUMMER< -7.589415e-18 222 9.539528e+01 4.8992790 *

30903) METADom_Chem_rotation_minus_2_SUMMER>=-7.589415e-18 252 9.846967e+01 5.7886900 *

3863) SatDom_blue_band_var..80< 1.754557e-08 138 5.865645e+01 6.6250000 *

483) SatDom_LST_day_mean..30>=46.7018 3342 5.217107e+03 4.5491020

966) METADom_Chem_rotation_minus_1_TREFLAN< 0.06314856 1924 2.791629e+03 4.1004830

1932) SatDom_PsnNet..80< 0.2848437 660 6.611385e+02 3.2212730

3864) SatDom_LST_night..70< 739.88 161 2.613328e+01 1.9164600 *

3865) SatDom_LST_night..70>=739.88 499 2.724569e+02 3.6422650

7730) SatDom_blue_band_var.0>=3.08336e-06 284 7.947001e+01 3.1972890 *

7731) SatDom_blue_band_var.0< 3.08336e-06 215 6.247410e+01 4.2300470 *

1933) SatDom_PsnNet..80>=0.2848437 1264 1.353908e+03 4.5595650

3866) SatDom_Reflect_b3_var..80>=1.456987e-06 1141 8.541435e+02 4.3843650

7732) BOMDom_rainfall_mean_.30>=0.45 475 3.077135e+02 3.9087160

15464) SatDom_Emis32..60>=30.5885 156 8.109474e+01 3.3379490 *

15465) SatDom_Emis32..60< 30.5885 319 1.509450e+02 4.1878370

30930) METADom_Chem_rotation_minus_2_PARAQUAT>=0.0001414427 147 8.321966e+01 3.8746940 *

30931) METADom_Chem_rotation_minus_2_PARAQUAT< 0.0001414427 172 4.099126e+01 4.4554650 *

7733) BOMDom_rainfall_mean_.30< 0.45 666 3.623198e+02 4.7236040

15466) SatDom_Reflect_b3_var..20>=2.712584e-05 246 8.555463e+01 4.2243090 *

15467) SatDom_Reflect_b3_var..20< 2.712584e-05 420 1.795186e+02 5.0160480

30934) BOMDom_solar_exposure_min_.20< 4.35 311 1.152438e+02 4.8751770 *

30935) BOMDom_solar_exposure_min_.20>=4.35 109 4.049416e+01 5.4179820 *

3867) SatDom_Reflect_b3_var..80< 1.456987e-06 123 1.398517e+02 6.1847970 *

967) METADom_Chem_rotation_minus_1_TREFLAN>=0.06314856 1418 1.512856e+03 5.1578070

1934) SatDom_Pot_EvapoTrans_min.0>=20.58125 1160 9.652543e+02 4.9125600

3868) SatDom_EVI_mean..40< 0.2052053 1050 7.358187e+02 4.7854190

7736) METADom_Fert_rotation_minus_1_GRANULOCK< -1.03563e-16 122 9.802649e+01 3.7932790 *

7737) METADom_Fert_rotation_minus_1_GRANULOCK>=-1.03563e-16 928 5.019147e+02 4.9158510

15474) SatDom_Reflect_b4_var..40< 3.825216e-06 124 5.509628e+01 4.2409680 *

15475) SatDom_Reflect_b4_var..40>=3.825216e-06 804 3.816299e+02 5.0199380

30950) SatDom_red_band_min..80>=0.15565 430 1.074900e+02 4.8345120 *

30951) SatDom_red_band_min..80< 0.15565 374 2.423568e+02 5.2331280 *

3869) SatDom_EVI_mean..40>=0.2052053 110 5.044600e+01 6.1261820 *

1935) SatDom_Pot_EvapoTrans_min.0< 20.58125 258 1.641419e+02 6.2604650 *

121) SatDom_Latent_heat_max..10>=2285000 5772 8.984252e+03 4.4993940

242) METADom_total_N_10cm.test_10cm< 40.6 3747 4.805004e+03 4.2157620

484) SatDom_LST_day..80>=656.605 695 6.560211e+02 3.4046470

968) SatDom_Reflect_b2_mean..40>=0.2621475 169 1.373339e+02 2.2328990 *

969) SatDom_Reflect_b2_mean..40< 0.2621475 526 2.120998e+02 3.7811220

1938) BOMDom_max_temperature_min_.50< 23.95 366 8.749664e+01 3.5655460 *

1939) BOMDom_max_temperature_min_.50>=23.95 160 6.868611e+01 4.2742500 *

485) SatDom_LST_day..80< 656.605 3052 3.587614e+03 4.4004690

970) SatDom_red_band_var..60< 7.720737e-06 1445 1.300625e+03 3.9622910

1940) SatDom_Reflect_b1_var..40< 2.537063e-05 698 4.402033e+02 3.4728800

3880) SatDom_Latent_heat_min..80< 3371250 542 2.258597e+02 3.2511070

7760) SatDom_LST_day_min.0>=33.78 158 6.233039e+01 2.6488610 *

7761) SatDom_LST_day_min.0< 33.78 384 8.264334e+01 3.4989060 *

3881) SatDom_Latent_heat_min..80>=3371250 156 9.506950e+01 4.2433970 *

1941) SatDom_Reflect_b1_var..40>=2.537063e-05 747 5.370141e+02 4.4195980

3882) SatDom_Emis32.0< 89.7865 574 3.595486e+02 4.1934670

7764) METADom_Fert_rotation_minus_1_DAP>=0.05217489 392 2.073745e+02 3.9639030

15528) SatDom_Reflect_b3.0< 5.222106 168 3.446596e+01 3.6911900 *

15529) SatDom_Reflect_b3.0>=5.222106 224 1.510432e+02 4.1684370 *

7765) METADom_Fert_rotation_minus_1_DAP< 0.05217489 182 8.702121e+01 4.6879120 *

3883) SatDom_Emis32.0>=89.7865 173 5.072700e+01 5.1698840 *

971) SatDom_red_band_var..60>=7.720737e-06 1607 1.760078e+03 4.7944740

1942) SatDom_Latent_heat_var..80>=7.872057e+09 1177 9.209494e+02 4.4854800

3884) SatDom_EvapoTrans..30< 643.5 355 1.445093e+02 3.8543380

7768) METADom_Fert_rotation_minus_2_Z< -3.538836e-17 153 4.500574e+01 3.4757520 *

7769) METADom_Fert_rotation_minus_2_Z>=-3.538836e-17 202 6.096476e+01 4.1410890 *

3885) SatDom_EvapoTrans..30>=643.5 822 5.739577e+02 4.7580540

7770) SatDom_blue_band_var..40>=8.252331e-07 428 1.899998e+02 4.4109350

15540) SatDom_LST_day_var..70>=8.115285 147 7.759947e+01 3.9866670 *

15541) SatDom_LST_day_var..70< 8.115285 281 7.209757e+01 4.6328830 *

7771) SatDom_blue_band_var..40< 8.252331e-07 394 2.763668e+02 5.1351270

15542) SatDom_FPAR_var..10>=0.0007167448 280 1.867618e+02 4.9293930 *

15543) SatDom_FPAR_var..10< 0.0007167448 114 4.864488e+01 5.6404390 *

1943) SatDom_Latent_heat_var..80< 7.872057e+09 430 4.191527e+02 5.6402560

3886) SatDom_Reflect_b6_max..80>=0.3683125 256 2.009067e+02 5.1942970 *

3887) SatDom_Reflect_b6_max..80< 0.3683125 174 9.242622e+01 6.2963790 *

243) METADom_total_N_10cm.test_10cm>=40.6 2025 3.320047e+03 5.0242170

486) METADom_Chem_rotation_minus_2_BRODAL< 0.003576852 864 9.491014e+02 4.3815620

972) SatDom_blue_band_var..30>=7.012467e-06 188 1.125076e+02 3.3569150 *

973) SatDom_blue_band_var..30< 7.012467e-06 676 5.843189e+02 4.6665240

1946) SatDom_Latent_heat_mean..50>=1085750 550 3.826990e+02 4.4734550

3892) SatDom_NIR_var..20>=6.637953e-05 225 1.594500e+02 4.1514670 *

3893) SatDom_NIR_var..20< 6.637953e-05 325 1.837723e+02 4.6963690 *

1947) SatDom_Latent_heat_mean..50< 1085750 126 9.162704e+01 5.5092860 *

487) METADom_Chem_rotation_minus_2_BRODAL>=0.003576852 1161 1.748557e+03 5.5024720

974) SatDom_PsnNet_var..50>=7.794809e-07 438 6.357493e+02 4.7993380

1948) SatDom_Pot_latent_heat_var.0>=2.751216e+11 154 1.458212e+02 3.9161690 *

1949) SatDom_Pot_latent_heat_var.0< 2.751216e+11 284 3.046755e+02 5.2782390 *

975) SatDom_PsnNet_var..50< 7.794809e-07 723 7.650757e+02 5.9284370

1950) BOMDom_min_temperature_var_.10< 13.39406 558 4.167728e+02 5.6173660

3900) SatDom_MIR_var..80>=2.058382e-05 263 1.671217e+02 5.2344110 *

3901) SatDom_MIR_var..80< 2.058382e-05 295 1.766948e+02 5.9587800 *

1951) BOMDom_min_temperature_var_.10>=13.39406 165 1.117061e+02 6.9804240 *

61) METADom_Fert_rotation_minus_2_THIRTY_TWO_TO_NINE>=0.1 918 3.193691e+03 5.8794010

122) METADom_Chem_rotation_minus_1_GOAL>=0.003762028 203 4.776409e+02 3.3966500 *

123) METADom_Chem_rotation_minus_1_GOAL< 0.003762028 715 1.109483e+03 6.5842940

246) METADom_Organic_C_10cm.test_10cm< 1.05 306 1.587312e+02 5.6274510

492) METADom_Chem_rotation_minus_1_LT>=-1.897354e-18 189 5.539891e+01 5.3480950 *

493) METADom_Chem_rotation_minus_1_LT< -1.897354e-18 117 6.475671e+01 6.0787180 *

247) METADom_Organic_C_10cm.test_10cm>=1.05 409 4.609893e+02 7.3001710

494) METADom_Chem_rotation_minus_2_STARANE>=0.0001257862 210 1.482361e+02 6.6666190 *

495) METADom_Chem_rotation_minus_2_STARANE< 0.0001257862 199 1.395108e+02 7.9687440 *

31) METADom_Nitrogen_fert_repeat_3>=31.05 792 3.437467e+03 6.3003660

62) METADom_pH_CaCl2_10cm.test_10cm>=6.075 335 2.998472e+02 4.4923280

124) SatDom_PsnNet_min..30< 0.0041875 152 7.710895e+01 3.8280920 *

125) SatDom_PsnNet_min..30>=0.0041875 183 9.997101e+01 5.0440440 *

63) METADom_pH_CaCl2_10cm.test_10cm< 6.075 457 1.239739e+03 7.6257330

126) SatDom_EVI_var..20< 1.419524e-05 118 2.510349e+02 5.8200850 *

127) SatDom_EVI_var..20>=1.419524e-05 339 4.700659e+02 8.2542480

254) SatDom_EvapoTrans..40< 262359 238 1.285406e+02 7.8323110 *

255) SatDom_EvapoTrans..40>=262359 101 1.993087e+02 9.2485150 *

**Cross-Species ‘Omnibus’ Regression tree: Predicted at End of Season**

**Leaf values indicate node-average yield, in t/Ha**

**Model Accuracy: Missing Site Prediction R^2^ = 0.70; p<0.0001**

**n= 97333**

**node), split, n, deviance, yval**

*** denotes terminal node**

1) root 97333 2.139798e+05 3.0011290

2) SatDom_Latent_heat_250< 7.149511e+08 38822 4.953533e+04 2.2059310

4) BOMDom_rainfall_csum_180< 277.3 18461 1.809311e+04 1.7938420

8) SatDom_Reflect_b6_max_110>=0.246525 10201 7.612290e+03 1.4832390

16) METADom_IODOSULFURON_METHYL_SODIUM< 0.05594712 10043 5.109848e+03 1.4312790

32) SatDom_Reflect_b1_min_150>=0.1634125 2707 6.813167e+02 1.0512860

64) BOMDom_min_temperature_var_160>=5.511389 2275 4.544425e+02 0.9479297

128) SatDom_MIR_min_140>=0.3157031 838 5.340488e+01 0.6644272

256) SatDom_Emis31_var_50>=6.944444e-07 619 2.182237e+01 0.5684814 *

257) SatDom_Emis31_var_50< 6.944444e-07 219 9.778192e+00 0.9356164 *

129) SatDom_MIR_min_140< 0.3157031 1437 2.944070e+02 1.1132570

258) SatDom_FPAR_var_180>=2.392361e-05 978 1.295949e+02 0.9624131

516) SatDom_Latent_heat_var_210< 4.82874e+10 761 6.683769e+01 0.8518528

1032) SatDom_Reflect_b4_var_50< 3.471727e-05 398 2.097565e+01 0.6912814 *

1033) SatDom_Reflect_b4_var_50>=3.471727e-05 363 2.434921e+01 1.0279060 *

517) SatDom_Latent_heat_var_210>=4.82874e+10 217 2.083330e+01 1.3501380 *

259) SatDom_FPAR_var_180< 2.392361e-05 459 9.514342e+01 1.4346620

518) METADom_Chem_rotation_minus_2_LOGRAN>=0.01668498 285 3.457796e+01 1.2031580 *

519) METADom_Chem_rotation_minus_2_LOGRAN< 0.01668498 174 2.027272e+01 1.8138510 *

65) BOMDom_min_temperature_var_160< 5.511389 432 7.458946e+01 1.5955790

130) SatDom_PsnNet_var_160>=1.43362e-07 267 1.786583e+01 1.3453930 *

131) SatDom_PsnNet_var_160< 1.43362e-07 165 1.296787e+01 2.0004240 *

33) SatDom_Reflect_b1_min_150< 0.1634125 7336 3.893418e+03 1.5714980

66) METADom_Dicot_TF>=0.5 1862 6.601364e+02 1.0724920

132) BOMDom_solar_exposure_mean_.40< 20.05 1595 3.673858e+02 0.9559060

264) SatDom_MIR_mean_190>=0.2904969 1157 1.352728e+02 0.8284183

528) SatDom_FPAR_min..30< 0.16125 900 7.692619e+01 0.7494111 *

529) SatDom_FPAR_min..30>=0.16125 257 3.305502e+01 1.1050970 *

265) SatDom_MIR_mean_190< 0.2904969 438 1.636342e+02 1.2926710

530) SatDom_LST_day_max_70< 33.54 222 1.826400e+01 0.8900000 *

531) SatDom_LST_day_max_70>=33.54 216 7.237830e+01 1.7065280 *

133) BOMDom_solar_exposure_mean_.40>=20.05 267 1.415611e+02 1.7689510 *

67) METADom_Dicot_TF< 0.5 5474 2.611918e+03 1.7412370

134) METADom_Fert_rotation_minus_1_TWENTY_EIGHT_TO_THIRTEEN>=-5.55762e-17 5372 2.072713e+03 1.7021630

268) SatDom_Reflect_b7_min_150>=0.2240125 2844 8.482412e+02 1.5024610

536) SatDom_blue_band_var_20< 9.39467e-07 1408 2.994441e+02 1.2490480

1072) SatDom_FPAR_var_130< 0.001527396 937 1.326459e+02 1.0527530

2144) SatDom_Reflect_b5_var_70>=8.130503e-05 655 6.180188e+01 0.9107023

4288) METADom_Fert_rotation_minus_2_LEGUME< 0.0008196721 430 2.394307e+01 0.7602558 *

4289) METADom_Fert_rotation_minus_2_LEGUME>=0.0008196721 225 9.525889e+00 1.1982220 *

2145) SatDom_Reflect_b5_var_70< 8.130503e-05 282 2.692815e+01 1.3826950 *

1073) SatDom_FPAR_var_130>=0.001527396 471 5.886901e+01 1.6395540

2146) SatDom_Reflect_b3_var_110>=6.989695e-06 175 9.435678e+00 1.2866860 *

2147) SatDom_Reflect_b3_var_110< 6.989695e-06 296 1.476021e+01 1.8481760 *

537) SatDom_blue_band_var_20>=9.39467e-07 1436 3.697217e+02 1.7509330

1074) SatDom_blue_band_max_220< 0.09935625 1095 1.951980e+02 1.5924380

2148) SatDom_FPAR_40< 21.73 343 3.340441e+01 1.2681340 *

2149) SatDom_FPAR_40>=21.73 752 1.092650e+02 1.7403590

4298) SatDom_Reflect_b7_var..20>=0.0005621688 214 1.263221e+01 1.4211680 *

4299) SatDom_Reflect_b7_var..20< 0.0005621688 538 6.615735e+01 1.8673230 *

1075) SatDom_blue_band_max_220>=0.09935625 341 5.868760e+01 2.2598830

2150) SatDom_blue_band_max_200>=0.1140656 143 6.181324e+00 1.8855940 *

2151) SatDom_blue_band_max_200< 0.1140656 198 1.800479e+01 2.5302020 *

269) SatDom_Reflect_b7_min_150< 0.2240125 2528 9.834526e+02 1.9268280

538) SatDom_LST_night_mean_100< 25.7995 1818 5.285049e+02 1.7277060

1076) SatDom_Reflect_b4_mean_140>=0.07354563 1403 2.813711e+02 1.5795300

2152) SatDom_Emis31_var_220< 9.166667e-07 1053 1.523225e+02 1.4700090

4304) SatDom_LAI_var_100< 0.01808333 334 4.824481e+01 1.2205090 *

4305) SatDom_LAI_var_100>=0.01808333 719 7.362758e+01 1.5859110 *

2153) SatDom_Emis31_var_220>=9.166667e-07 350 7.841887e+01 1.9090290 *

1077) SatDom_Reflect_b4_mean_140< 0.07354563 415 1.121868e+02 2.2286510

2154) SatDom_GPP..70< 0.1855125 267 3.754014e+01 1.9684270 *

2155) SatDom_GPP..70>=0.1855125 148 2.394867e+01 2.6981080 *

539) SatDom_LST_night_mean_100>=25.7995 710 1.982937e+02 2.4366900

1078) SatDom_MIR_mean_90>=0.1369219 315 4.868371e+01 2.0509520 *

1079) SatDom_MIR_mean_90< 0.1369219 395 6.536268e+01 2.7443040

2158) SatDom_Latent_heat_110>=3.445169e+08 283 2.687364e+01 2.5834980 *

2159) SatDom_Latent_heat_110< 3.445169e+08 112 1.268026e+01 3.1506250 *

135) METADom_Fert_rotation_minus_1_TWENTY_EIGHT_TO_THIRTEEN< -5.55762e-17 102 9.904542e+01 3.7991180 *

17) METADom_IODOSULFURON_METHYL_SODIUM>=0.05594712 158 7.518804e+02 4.7859490 *

9) SatDom_Reflect_b6_max_110< 0.246525 8260 8.281293e+03 2.1774320

18) METADom_Dicot_TF>=0.5 2721 1.659189e+03 1.4852960

36) SatDom_PsnNet_mean_180< 0.014995 2215 6.867679e+02 1.3291020

72) SatDom_LST_day_min_80< 28.97 489 6.093764e+01 0.9039059

144) SatDom_LST_night_var..40< 9.089281 375 2.478385e+01 0.7706400 *

145) SatDom_LST_night_var..40>=9.089281 114 7.586207e+00 1.3422810 *

73) SatDom_LST_day_min_80>=28.97 1726 5.123764e+02 1.4495650

146) SatDom_LST_night_190< 8606.534 1451 3.499572e+02 1.3651620

292) SatDom_FPAR_mean..30< 0.138 429 7.099699e+01 1.0616780

584) BOMDom_max_temperature_max_180< 39.2 262 2.205212e+01 0.8281679 *

585) BOMDom_max_temperature_max_180>=39.2 167 1.224585e+01 1.4280240 *

293) SatDom_FPAR_mean..30>=0.138 1022 2.228626e+02 1.4925540

586) SatDom_Emis31..40>=50.15217 313 3.607416e+01 1.1845370 *

587) SatDom_Emis31..40< 50.15217 709 1.439831e+02 1.6285330

1174) SatDom_Reflect_b3_var..20< 1.831803e-05 467 7.968415e+01 1.4955250 *

1175) SatDom_Reflect_b3_var..20>=1.831803e-05 242 4.009384e+01 1.8852070 *

147) SatDom_LST_night_190>=8606.534 275 9.754127e+01 1.8949090 *

37) SatDom_PsnNet_mean_180>=0.014995 506 6.818300e+02 2.1690320

74) SatDom_MIR_max_40>=0.1773875 286 1.568895e+02 1.5306290 *

75) SatDom_MIR_max_40< 0.1773875 220 2.568493e+02 2.9989550 *

19) METADom_Dicot_TF< 0.5 5539 4.678264e+03 2.5174400

38) SatDom_Reflect_b7_max..40>=0.259275 4426 2.917058e+03 2.3307840

76) SatDom_Emis31_var_160< 9.611111e-07 3611 1.805136e+03 2.1599890

152) SatDom_GPP_140< 3.54865 1436 4.972924e+02 1.7733430

304) BOMDom_rainfall_mean_170>=0.28 717 1.621493e+02 1.4467640

608) BOMDom_rainfall_mean_.20>=0.37 389 6.345188e+01 1.1867870 *

609) BOMDom_rainfall_mean_.20< 0.37 328 4.122400e+01 1.7550910 *

305) BOMDom_rainfall_mean_170< 0.28 719 1.824148e+02 2.0990130

610) SatDom_red_band_mean..40>=0.1598322 286 3.260159e+01 1.6738810 *

611) SatDom_red_band_mean..40< 0.1598322 433 6.398039e+01 2.3798150 *

153) SatDom_GPP_140>=3.54865 2175 9.514336e+02 2.4152640

306) SatDom_Pot_latent_heat_var_210>=1.303697e+12 977 2.559685e+02 2.0457220

612) SatDom_Latent_heat_var_220< 5.920911e+10 739 1.256694e+02 1.8895400

1224) SatDom_PsnNet_min_40< 0.00625 107 2.643661e+01 1.4021500 *

1225) SatDom_PsnNet_min_40>=0.00625 632 6.951173e+01 1.9720570 *

613) SatDom_Latent_heat_var_220>=5.920911e+10 238 5.630069e+01 2.5306720 *

307) SatDom_Pot_latent_heat_var_210< 1.303697e+12 1198 4.532359e+02 2.7166360

614) SatDom_Reflect_b5_var_10>=8.960882e-05 711 1.985072e+02 2.4390300

1228) SatDom_Reflect_b5_var_150< 7.165549e-06 118 2.633485e+01 1.8380510 *

1229) SatDom_Reflect_b5_var_150>=7.165549e-06 593 1.210731e+02 2.5586170

2458) METADom_Chem_rotation_minus_2_ATLANTIS>=-2.094679e-17 449 7.133409e+01 2.4358570 *

2459) METADom_Chem_rotation_minus_2_ATLANTIS< -2.094679e-17 144 2.187452e+01 2.9413890 *

615) SatDom_Reflect_b5_var_10< 8.960882e-05 487 1.199390e+02 3.1219300

1230) SatDom_LST_day_min_50< 30.34 177 1.086045e+01 2.6873450 *

1231) SatDom_LST_day_min_50>=30.34 310 5.656260e+01 3.3700650 *

77) SatDom_Emis31_var_160>=9.611111e-07 815 5.398742e+02 3.0875210

154) SatDom_GPP_mean_180< 0.00903125 214 5.614788e+01 2.0296730 *

155) SatDom_GPP_mean_180>=0.00903125 601 1.589802e+02 3.4641930

310) SatDom_LST_day_mean.0< 37.959 169 1.406887e+01 2.9515980 *

311) SatDom_LST_day_mean.0>=37.959 432 8.313437e+01 3.6647220 *

39) SatDom_Reflect_b7_max..40< 0.259275 1113 9.937890e+02 3.2597040

78) BOMDom_min_temperature_var_130>=8.551556 780 4.817174e+02 2.8771790

156) SatDom_Reflect_b3_mean_210< 0.0818475 309 8.615114e+01 2.2261170

312) SatDom_FPAR_var..10< 0.0004043403 126 9.744537e+00 1.7424600 *

313) SatDom_FPAR_var..10>=0.0004043403 183 2.663846e+01 2.5591260 *

157) SatDom_Reflect_b3_mean_210>=0.0818475 471 1.786570e+02 3.3043100

314) SatDom_Reflect_b6_max_170< 0.3487 286 6.307072e+01 3.0808040 *

315) SatDom_Reflect_b6_max_170>=0.3487 185 7.921210e+01 3.6498380 *

79) BOMDom_min_temperature_var_130< 8.551556 333 1.305994e+02 4.1557060

158) SatDom_FPAR_mean..10< 0.222625 196 4.120047e+01 3.8114800 *

159) SatDom_FPAR_mean..10>=0.222625 137 3.294844e+01 4.6481750 *

5) BOMDom_rainfall_csum_180>=277.3 20361 2.546476e+04 2.5795660

10) METADom_Dicot_TF>=0.5 5745 3.544675e+03 1.9398800

20) SatDom_Pot_latent_heat_max_140>=6127859 5127 2.747265e+03 1.8444590

40) SatDom_PsnNet_mean_190< 0.00935625 2094 9.465647e+02 1.5879370

80) BOMDom_min_temperature_min_.30< 10.15 1493 4.890747e+02 1.4144680

160) SatDom_NDVI_var..70< 1.916068e-05 1234 2.919246e+02 1.3140110

320) SatDom_MixClouds_10< 4.5 966 1.704210e+02 1.2027430

640) SatDom_FPAR_max..80< 0.16875 720 9.266475e+01 1.1037920

1280) SatDom_LST_night_max_250< 45.435 575 5.680996e+01 1.0139650 *

1281) SatDom_LST_night_max_250>=45.435 145 1.281700e+01 1.4600000 *

641) SatDom_FPAR_max..80>=0.16875 246 5.007283e+01 1.4923580 *

321) SatDom_MixClouds_10>=4.5 268 6.643570e+01 1.7150750 *

161) SatDom_NDVI_var..70>=1.916068e-05 259 1.253659e+02 1.8930890 *

81) BOMDom_min_temperature_min_.30>=10.15 601 3.009562e+02 2.0188690

162) SatDom_Latent_heat_mean_170< 594553.9 135 5.070829e+01 1.3068890 *

163) SatDom_Latent_heat_mean_170>=594553.9 466 1.619892e+02 2.2251290

326) SatDom_Reflect_b5_min..20< 0.3358375 328 6.884037e+01 2.0032320

652) SatDom_LST_day_mean_80< 35.318 127 2.112774e+01 1.6572440 *

653) SatDom_LST_day_mean_80>=35.318 201 2.290402e+01 2.2218410 *

327) SatDom_Reflect_b5_min..20>=0.3358375 138 3.861281e+01 2.7525360 *

41) SatDom_PsnNet_mean_190>=0.00935625 3033 1.567775e+03 2.0215630

82) BOMDom_min_temperature_mean_.30< 14.305 2402 1.047789e+03 1.9070650

164) SatDom_Reflect_b2_min_90>=0.2356875 1999 7.497884e+02 1.8094600

328) SatDom_GPP_var_130< 7.704326e-06 1176 4.054023e+02 1.6463520

656) SatDom_Latent_heat_250< 6.920408e+08 1063 2.811154e+02 1.5746570

1312) SatDom_LST_night_max..40< 33.85 343 6.818071e+01 1.2805250 *

1313) SatDom_LST_night_max..40>=33.85 720 1.691242e+02 1.7147780

2626) SatDom_LST_night_max..60>=37.87 427 8.130578e+01 1.5327870

5252) SatDom_LST_day_var_100>=3.313666 126 1.075820e+01 1.1219840 *

5253) SatDom_LST_day_var_100< 3.313666 301 4.038291e+01 1.7047510 *

2627) SatDom_LST_night_max..60< 37.87 293 5.306540e+01 1.9800000 *

657) SatDom_Latent_heat_250>=6.920408e+08 113 6.742183e+01 2.3207960 *

329) SatDom_GPP_var_130>=7.704326e-06 823 2.683939e+02 2.0425270

658) BOMDom_min_temperature_var_220>=23.0945 159 3.436242e+01 1.4359750 *

659) BOMDom_min_temperature_var_220< 23.0945 664 1.615269e+02 2.1877710

1318) SatDom_Reflect_b1_mean_180>=0.1557662 292 4.211989e+01 1.9048290 *

1319) SatDom_Reflect_b1_mean_180< 0.1557662 372 7.768129e+01 2.4098660 *

165) SatDom_Reflect_b2_min_90< 0.2356875 403 1.844925e+02 2.3912160

330) SatDom_PsnNet_mean_40>=0.00390625 296 8.068540e+01 2.1525340 *

331) SatDom_PsnNet_mean_40< 0.00390625 107 4.029556e+01 3.0514950 *

83) BOMDom_min_temperature_mean_.30>=14.305 631 3.686265e+02 2.4574170

166) BOMDom_min_temperature_var_.20>=9.189667 246 9.829239e+01 1.9802440 *

167) BOMDom_min_temperature_var_.20< 9.189667 385 1.785314e+02 2.7623120

334) SatDom_FPAR_var..40< 6.430556e-05 189 3.203592e+01 2.3960320 *

335) SatDom_FPAR_var..40>=6.430556e-05 196 9.668825e+01 3.1155100 *

21) SatDom_Pot_latent_heat_max_140< 6127859 618 3.634459e+02 2.7315050

42) SatDom_Reflect_b7_var_30< 0.0003580819 508 1.902334e+02 2.5096260

84) SatDom_Emis32_var_160< 4.888889e-07 102 2.145766e+01 1.8488240 *

85) SatDom_Emis32_var_160>=4.888889e-07 406 1.130468e+02 2.6756400

170) SatDom_Reflect_b5_max_50>=0.2771062 257 3.726409e+01 2.4798050 *

171) SatDom_Reflect_b5_max_50< 0.2771062 149 4.892595e+01 3.0134230 *

43) SatDom_Reflect_b7_var_30>=0.0003580819 110 3.270760e+01 3.7561820 *

11) METADom_Dicot_TF< 0.5 14616 1.864521e+04 2.8310030

22) SatDom_MIR_mean_150>=0.1676194 10927 9.245363e+03 2.5366650

44) SatDom_GPP_max_130< 0.02310625 1913 8.981248e+02 1.7704390

88) METADom_PROPICONAZOLE_repeat_1< 180.2363 1591 4.564645e+02 1.5892460

176) SatDom_Reflect_b1_min_190>=0.1844875 685 1.031964e+02 1.2465690

352) SatDom_red_band_var_40< 8.387858e-06 407 2.328119e+01 1.0215720 *

353) SatDom_red_band_var_40>=8.387858e-06 278 2.914689e+01 1.5759710 *

177) SatDom_Reflect_b1_min_190< 0.1844875 906 2.120140e+02 1.8483330

354) BOMDom_solar_exposure_var_.20< 3.909556 471 6.665983e+01 1.5833120 *

355) BOMDom_solar_exposure_var_.20>=3.909556 435 7.645384e+01 2.1352870 *

89) METADom_PROPICONAZOLE_repeat_1>=180.2363 322 1.313375e+02 2.6657140

178) METADom_IMAZAPYR>=25.15533 201 2.952800e+01 2.2676620 *

179) METADom_IMAZAPYR< 25.15533 121 1.705797e+01 3.3269420 *

45) SatDom_GPP_max_130>=0.02310625 9014 6.985756e+03 2.6992780

90) METADom_METALAXYL_M< 0.3071952 8180 5.421380e+03 2.6040510

180) SatDom_NIR_mean_110< 0.3337466 5281 3.342761e+03 2.4196000

360) SatDom_LST_night_max_10< 30.83 3851 1.769373e+03 2.2326690

720) SatDom_Reflect_b7_var..30>=0.0001004051 1934 6.253336e+02 1.9752330

1440) SatDom_Emis31_mean_150< 0.98345 1327 3.197776e+02 1.7923440

2880) SatDom_LST_night_var_180< 3.738174 465 8.396056e+01 1.4463660

5760) SatDom_Latent_heat_mean..30< 1130745 243 1.625642e+01 1.1976130 *

5761) SatDom_Latent_heat_mean..30>=1130745 222 3.620919e+01 1.7186490 *

2881) SatDom_LST_night_var_180>=3.738174 862 1.501303e+02 1.9789790

5762) SatDom_EvapoTrans_min..20< 3.775 394 3.501975e+01 1.7278930 *

5763) SatDom_EvapoTrans_min..20>=3.775 468 6.935944e+01 2.1903630 *

1441) SatDom_Emis31_mean_150>=0.98345 607 1.641350e+02 2.3750580

2882) BOMDom_max_temperature_min_170>=21.55 306 4.487567e+01 2.0228760 *

2883) BOMDom_max_temperature_min_170< 21.55 301 4.272123e+01 2.7330900 *

721) SatDom_Reflect_b7_var..30< 0.0001004051 1917 8.865561e+02 2.4923890

1442) SatDom_blue_band_min_100>=0.03840937 290 9.230200e+01 1.6805860 *

1443) SatDom_blue_band_min_100< 0.03840937 1627 5.690720e+02 2.6370870

2886) BOMDom_solar_exposure_var_90< 7.075611 784 1.849916e+02 2.3351280

5772) SatDom_LST_day_max_70>=36.59 573 7.390758e+01 2.1545030

11544) BOMDom_max_temperature_mean_240< 33.13 313 2.565935e+01 1.9510540 *

11545) BOMDom_max_temperature_mean_240>=33.13 260 1.969641e+01 2.3994230 *

5773) SatDom_LST_day_max_70< 36.59 211 4.162259e+01 2.8256400 *

2887) BOMDom_solar_exposure_var_90>=7.075611 843 2.461143e+02 2.9179120

5774) SatDom_Reflect_b3_var..70< 1.232009e-05 635 1.004244e+02 2.7488980

11548) SatDom_GPP_20< 0.7169375 165 1.164720e+01 2.3870910 *

11549) SatDom_GPP_20>=0.7169375 470 5.959536e+01 2.8759150 *

5775) SatDom_Reflect_b3_var..70>=1.232009e-05 208 7.217315e+01 3.4338940 *

361) SatDom_LST_night_max_10>=30.83 1430 1.076434e+03 2.9230070

722) SatDom_Reflect_b2_var_190< 0.0002736447 978 3.616979e+02 2.6040080

1444) SatDom_Emis31_var_20< 1.790123e-06 749 2.059087e+02 2.4097330

2888) SatDom_GPP_mean_180>=0.01034312 205 3.058192e+01 1.8772200 *

2889) SatDom_GPP_mean_180< 0.01034312 544 9.528851e+01 2.6104040

5778) BOMDom_rainfall_var_50< 22.37917 371 4.283213e+01 2.4371700 *

5779) BOMDom_rainfall_var_50>=22.37917 173 1.744607e+01 2.9819080 *

1445) SatDom_Emis31_var_20>=1.790123e-06 229 3.505783e+01 3.2394320 *

723) SatDom_Reflect_b2_var_190>=0.0002736447 452 3.998783e+02 3.6132300

1446) SatDom_Reflect_b2_max..80>=0.2848375 327 6.614199e+01 3.2587460 *

1447) SatDom_Reflect_b2_max..80< 0.2848375 125 1.851533e+02 4.5405600 *

181) SatDom_NIR_mean_110>=0.3337466 2899 1.571648e+03 2.9400590

362) SatDom_Reflect_b3_var_60>=1.317713e-05 720 3.257958e+02 2.3283330

724) SatDom_Reflect_b6_var..40< 5.366458e-05 380 6.787550e+01 1.9585530

1448) METADom_Fert_rotation_minus_2_TWENTY_EIGHT_TO_THIRTEEN< -1.513546e-17 120 1.506259e+01 1.5352500 *

1449) METADom_Fert_rotation_minus_2_TWENTY_EIGHT_TO_THIRTEEN>=-1.513546e-17 260 2.138660e+01 2.1539230 *

725) SatDom_Reflect_b6_var..40>=5.366458e-05 340 1.478866e+02 2.7416180

1450) SatDom_Pot_EvapoTrans_mean..10>=18034.27 229 2.697982e+01 2.4522710 *

1451) SatDom_Pot_EvapoTrans_mean..10< 18034.27 111 6.218097e+01 3.3385590 *

363) SatDom_Reflect_b3_var_60< 1.317713e-05 2179 8.873963e+02 3.1421890

726) SatDom_PsnNet_mean_160< 0.02201313 2033 7.118480e+02 3.0711210

1452) SatDom_Aero_q_250< 7.5 1832 5.699321e+02 2.9977570

2904) SatDom_Aero_q..60>=4.5 209 8.863532e+01 2.4668420 *

2905) SatDom_Aero_q..60< 4.5 1623 4.147997e+02 3.0661240

5810) SatDom_EVI_230>=75.92181 968 2.449664e+02 2.9360230

11620) METADom_Chem_rotation_minus_2_ATTACK>=-9.324139e-18 550 9.565570e+01 2.7600180 *

11621) METADom_Chem_rotation_minus_2_ATTACK< -9.324139e-18 418 1.098550e+02 3.1676080 *

5811) SatDom_EVI_230< 75.92181 655 1.292340e+02 3.2583970

11622) METADom_Chem_rotation_minus_2_GLEAN< -4.97432e-17 120 1.284870e+01 2.8659170 *

11623) METADom_Chem_rotation_minus_2_GLEAN>=-4.97432e-17 535 9.375428e+01 3.3464300 *

1453) SatDom_Aero_q_250>=7.5 201 4.218179e+01 3.7398010 *

727) SatDom_PsnNet_mean_160>=0.02201313 146 2.230374e+01 4.1317810 *

91) METADom_METALAXYL_M>=0.3071952 834 7.626606e+02 3.6332730

182) SatDom_EVI_230< 65.92012 388 1.984253e+02 2.9209540

364) METADom_PROPICONAZOLE< 2157.036 242 5.668413e+01 2.5128510 *

365) METADom_PROPICONAZOLE>=2157.036 146 3.463061e+01 3.5973970 *

183) SatDom_EVI_230>=65.92012 446 1.960953e+02 4.2529600

366) SatDom_Reflect_b7_max_100< 0.138 298 7.276664e+01 3.9792620 *

367) SatDom_Reflect_b7_max_100>=0.138 148 5.605697e+01 4.8040540 *

23) SatDom_MIR_mean_150< 0.1676194 3689 5.649135e+03 3.7028460

46) SatDom_GPP_min_160< 0.0376 3477 4.155498e+03 3.5531780

92) SatDom_Latent_heat_mean_40< 2474114 3119 2.809112e+03 3.3817730

184) SatDom_LST_night_min_210>=28.225 2275 1.711638e+03 3.1282150

368) SatDom_FPAR_mean_200< 0.1295 559 2.277190e+02 2.4203760

736) METADom_Fert_rotation_minus_2_EIGHTEEN_TO_TWENTY< 0.000676819 233 5.376524e+01 1.9741200 *

737) METADom_Fert_rotation_minus_2_EIGHTEEN_TO_TWENTY>=0.000676819 326 9.438965e+01 2.7393250

1474) METADom_Chem_rotation_minus_2_SELECT>=0.03212549 138 3.105675e+01 2.4158700 *

1475) METADom_Chem_rotation_minus_2_SELECT< 0.03212549 188 3.829672e+01 2.9767550 *

369) SatDom_FPAR_mean_200>=0.1295 1716 1.112601e+03 3.3588000

738) BOMDom_min_temperature_var_80>=12.05111 211 1.125691e+02 2.4549290 *

739) BOMDom_min_temperature_var_80< 12.05111 1505 8.034810e+02 3.4855220

1478) BOMDom_max_temperature_var_170< 31.13561 1201 5.004054e+02 3.3403080

2956) BOMDom_solar_exposure_mean_170< 25.81 1013 3.190308e+02 3.2129320

5912) SatDom_GPP_max..40< 0.012825 729 1.808875e+02 3.0485050

11824) BOMDom_min_temperature_min_180< 7.2 408 7.533276e+01 2.8697060 *

11825) BOMDom_min_temperature_min_180>=7.2 321 7.593284e+01 3.2757630 *

5913) SatDom_GPP_max..40>=0.012825 284 6.784170e+01 3.6350000 *

2957) BOMDom_solar_exposure_mean_170>=25.81 188 7.637899e+01 4.0266490 *

1479) BOMDom_max_temperature_var_170>=31.13561 304 1.776980e+02 4.0592110

2958) SatDom_EvapoTrans_mean..20>=21912.86 117 4.229611e+01 3.5891450 *

2959) SatDom_EvapoTrans_mean..20< 21912.86 187 9.337434e+01 4.3533160 *

185) SatDom_LST_night_min_210< 28.225 844 5.569593e+02 4.0652370

370) SatDom_NIR_var_180< 3.084636e-05 262 1.342814e+02 3.3183590 *

371) SatDom_NIR_var_180>=3.084636e-05 582 2.107343e+02 4.4014600

742) SatDom_Reflect_b3_max_100>=0.02871875 464 1.278581e+02 4.2674140 *

743) SatDom_Reflect_b3_max_100< 0.02871875 118 4.175446e+01 4.9285590 *

93) SatDom_Latent_heat_mean_40>=2474114 358 4.563975e+02 5.0465080

186) SatDom_Reflect_b3_max_70< 0.0351125 132 6.781208e+01 3.9746210 *

187) SatDom_Reflect_b3_max_70>=0.0351125 226 1.483447e+02 5.6725660 *

47) SatDom_GPP_min_160>=0.0376 212 1.383323e+02 6.1575470 *

3) SatDom_Latent_heat_250>=7.149511e+08 58511 1.236078e+05 3.5287410

6) METADom_Dicot_TF>=0.5 17101 1.227262e+04 2.3097950

12) METADom_sub_cropFaba.Bean< 0.5 15569 9.206321e+03 2.2342540

24) BOMDom_rainfall_csum_160< 231.65 1423 5.737222e+02 1.5278290

48) SatDom_Latent_heat_max_30>=1915000 1041 3.233517e+02 1.3687030

96) METADom_ALPHA_CYPERMETHRIN_repeat_1< 1.844547 792 1.519292e+02 1.2004550

192) SatDom_EVI_var..80< 4.153499e-06 351 5.076187e+01 0.9388889 *

193) SatDom_EVI_var..80>=4.153499e-06 441 5.803978e+01 1.4086390 *

97) METADom_ALPHA_CYPERMETHRIN_repeat_1>=1.844547 249 7.769230e+01 1.9038550 *

49) SatDom_Latent_heat_max_30< 1915000 382 1.521796e+02 1.9614660

98) SatDom_Reflect_b6_max_250>=0.404325 265 3.116888e+01 1.7333210 *

99) SatDom_Reflect_b6_max_250< 0.404325 117 7.597612e+01 2.4782050 *

25) BOMDom_rainfall_csum_160>=231.65 14146 7.851035e+03 2.3053160

50) SatDom_LST_day_max_170>=52.42 7798 4.371157e+03 2.1380660

100) SatDom_LST_night_max_90< 29.92 5987 2.905211e+03 2.0299930

200) SatDom_Reflect_b2_min_120< 0.407125 3676 1.449976e+03 1.8882920

400) SatDom_LST_night_mean_60< 24.16655 2061 6.852891e+02 1.7177920

800) SatDom_LST_night_min_190< 35.2 1887 5.467795e+02 1.6596660

1600) SatDom_NIR_var..50>=9.242079e-07 1657 3.787093e+02 1.5783830

3200) SatDom_Pot_EvapoTrans..40>=525549.3 139 1.839851e+01 1.0566190 *

3201) SatDom_Pot_EvapoTrans..40< 525549.3 1518 3.190047e+02 1.6261590

6402) SatDom_red_band_min..70>=0.1619375 203 2.493144e+01 1.2045810 *

6403) SatDom_red_band_min..70< 0.1619375 1315 2.524249e+02 1.6912400

12806) SatDom_blue_band_var_90>=1.031272e-06 499 7.855553e+01 1.5086770 *

12807) SatDom_blue_band_var_90< 1.031272e-06 816 1.470679e+02 1.8028800

25614) SatDom_Pot_latent_heat_var_240>=1.200603e+11 620 7.760538e+01 1.7117580 *

25615) SatDom_Pot_latent_heat_var_240< 1.200603e+11 196 4.803015e+01 2.0911220 *

1601) SatDom_NIR_var..50< 9.242079e-07 230 7.825053e+01 2.2452610 *

801) SatDom_LST_night_min_190>=35.2 174 6.299261e+01 2.3481610 *

401) SatDom_LST_night_mean_60>=24.16655 1615 6.283151e+02 2.1058760

802) SatDom_blue_band_max_30< 0.03264688 395 1.161566e+02 1.7087850

1604) SatDom_Reflect_b6_mean_40>=0.1813331 291 6.287617e+01 1.5474230 *

1605) SatDom_Reflect_b6_mean_40< 0.1813331 104 2.450249e+01 2.1602880 *

803) SatDom_blue_band_max_30>=0.03264688 1220 4.297085e+02 2.2344430

1606) SatDom_LST_night_max_200< 33.00667 108 2.377722e+01 1.4525000 *

1607) SatDom_LST_night_max_200>=33.00667 1112 3.334829e+02 2.3103870

3214) SatDom_Reflect_b2_max_90< 0.368325 559 1.252279e+02 2.0647580

6428) SatDom_EVI_min_20>=0.1149563 386 5.424532e+01 1.8878500 *

6429) SatDom_EVI_min_20< 0.1149563 173 3.194785e+01 2.4594800 *

3215) SatDom_Reflect_b2_max_90>=0.368325 553 1.404365e+02 2.5586800

6430) SatDom_Reflect_b7_min_70< 0.072325 128 2.728029e+01 2.0690630 *

6431) SatDom_Reflect_b7_min_70>=0.072325 425 7.322987e+01 2.7061410 *

201) SatDom_Reflect_b2_min_120>=0.407125 2311 1.264014e+03 2.2553920

402) SatDom_GPP_mean_140< 0.058425 2087 9.814619e+02 2.1630140

804) BOMDom_min_temperature_csum_.70< 282.725 690 2.580662e+02 1.7867100

1608) SatDom_Reflect_b6_mean..70>=0.3926994 250 7.880164e+01 1.3908000 *

1609) SatDom_Reflect_b6_mean..70< 0.3926994 440 1.178135e+02 2.0116590 *

805) BOMDom_min_temperature_csum_.70>=282.725 1397 5.774295e+02 2.3488760

1610) SatDom_Pot_latent_heat_mean_120< 1.223345e+07 1257 3.729483e+02 2.2477880

3220) SatDom_FPAR_10< 19.4775 627 1.655030e+02 2.0008290

6440) METADom_Chem_rotation_minus_2_VERDICT>=-1.908196e-17 458 8.356777e+01 1.8354150

12880) BOMDom_solar_exposure_var_60< 4.117389 245 2.898762e+01 1.6126940 *

12881) BOMDom_solar_exposure_var_60>=4.117389 213 2.844806e+01 2.0915960 *

6441) METADom_Chem_rotation_minus_2_VERDICT< -1.908196e-17 169 3.544157e+01 2.4491120 *

3221) SatDom_FPAR_10>=19.4775 630 1.311475e+02 2.4935710

6442) SatDom_LAI_min_130>=1.68125 263 5.215192e+01 2.2177950 *

6443) SatDom_LAI_min_130< 1.68125 367 4.465987e+01 2.6911990 *

1611) SatDom_Pot_latent_heat_mean_120>=1.223345e+07 140 7.630698e+01 3.2565000 *

403) SatDom_GPP_mean_140>=0.058425 224 9.880954e+01 3.1160710 *

101) SatDom_LST_night_max_90>=29.92 1811 1.164849e+03 2.4953450

202) SatDom_Reflect_b2_140< 69.26393 1202 6.470320e+02 2.2876870

404) SatDom_Reflect_b5_mean..50>=0.2589431 938 3.615876e+02 2.1315990

808) SatDom_Reflect_b2_mean_20>=0.166735 745 2.255874e+02 1.9814770

1616) SatDom_PsnNet_var..70< 5.391441e-07 416 1.201748e+02 1.7757450

3232) SatDom_Reflect_b7_var_30< 0.0001905638 265 5.023244e+01 1.5820000 *

3233) SatDom_Reflect_b7_var_30>=0.0001905638 151 4.253769e+01 2.1157620 *

1617) SatDom_PsnNet_var..70>=5.391441e-07 329 6.554185e+01 2.2416110 *

809) SatDom_Reflect_b2_mean_20< 0.166735 193 5.439947e+01 2.7110880 *

405) SatDom_Reflect_b5_mean..50< 0.2589431 264 1.813942e+02 2.8422730 *

203) SatDom_Reflect_b2_140>=69.26393 609 3.636812e+02 2.9052050

406) SatDom_blue_band_min_60< 0.0214625 265 1.313722e+02 2.3977740 *

407) SatDom_blue_band_min_60>=0.0214625 344 1.115110e+02 3.2961050

814) SatDom_GPP_max..60< 0.01275 205 4.729458e+01 3.0875610 *

815) SatDom_GPP_max..60>=0.01275 139 4.215203e+01 3.6036690 *

51) SatDom_LST_day_max_170< 52.42 6348 2.993794e+03 2.5107690

102) SatDom_Reflect_b6_max_40>=0.231375 3941 1.658618e+03 2.3586300

204) SatDom_LST_night_max_220< 39.95 3659 1.376718e+03 2.3047470

408) SatDom_Reflect_b1_var_210>=0.0008046924 429 1.402534e+02 1.8164570

816) SatDom_Reflect_b2_var..80>=2.719759e-05 182 1.538710e+01 1.3585160 *

817) SatDom_Reflect_b2_var..80< 2.719759e-05 247 5.857607e+01 2.1538870 *

409) SatDom_Reflect_b1_var_210< 0.0008046924 3230 1.120594e+03 2.3696010

818) SatDom_NDVI_max_110< 0.8754 3126 9.723355e+02 2.3373610

1636) SatDom_MIR_min_250>=0.2876563 655 1.597564e+02 2.0120920

3272) SatDom_MIR_var_210< 4.827293e-05 353 3.456826e+01 1.7466570 *

3273) SatDom_MIR_var_210>=4.827293e-05 302 7.124663e+01 2.3223510 *

1637) SatDom_MIR_min_250< 0.2876563 2471 7.249106e+02 2.4235820

3274) MANDom_Trial_operatorsDAWA.Mount.Barker>=-4.255494e-19 2335 5.960014e+02 2.3811180

6548) SatDom_Emis32_mean_20< 0.98625 618 1.289150e+02 2.0751620

13096) BOMDom_max_temperature_var_.40>=8.941014 249 3.994870e+01 1.7973090 *

13097) BOMDom_max_temperature_var_.40< 8.941014 369 5.677120e+01 2.2626560 *

6549) SatDom_Emis32_mean_20>=0.98625 1717 3.884139e+02 2.4912410

13098) SatDom_Reflect_b7_var_150< 1.163851e-05 582 1.194862e+02 2.2388830

26196) SatDom_LST_day_min_80>=29.51 254 3.784047e+01 1.9587010 *

26197) SatDom_LST_day_min_80< 29.51 328 4.626516e+01 2.4558540 *

13099) SatDom_Reflect_b7_var_150>=1.163851e-05 1135 2.128578e+02 2.6206430

26198) SatDom_Latent_heat_mean_110>=5543267 141 2.360258e+01 2.1960280 *

26199) SatDom_Latent_heat_mean_110< 5543267 994 1.602271e+02 2.6808750

52398) SatDom_GPP_min..50< 0.01756875 795 9.857699e+01 2.6055970 *

52399) SatDom_GPP_min..50>=0.01756875 199 3.914749e+01 2.9816080 *

3275) MANDom_Trial_operatorsDAWA.Mount.Barker< -4.255494e-19 136 5.240985e+01 3.1526470 *

819) SatDom_NDVI_max_110>=0.8754 104 4.734701e+01 3.3386540 *

205) SatDom_LST_night_max_220>=39.95 282 1.334369e+02 3.0577660 *

103) SatDom_Reflect_b6_max_40< 0.231375 2407 1.094602e+03 2.7598670

206) BOMDom_min_temperature_max_.20>=19 172 3.954817e+01 1.8726740 *

207) BOMDom_min_temperature_max_.20< 19 2235 9.092524e+02 2.8281430

414) SatDom_Reflect_b1_max_160>=0.0479125 2005 7.221871e+02 2.7535510

828) BOMDom_min_temperature_mean_30< 7.42 1375 4.712788e+02 2.5996220

1656) SatDom_EVI_mean_200< 0.1557722 150 5.605694e+01 1.9452670 *

1657) SatDom_EVI_mean_200>=0.1557722 1225 3.431302e+02 2.6797470

3314) BOMDom_rainfall_max_.30>=2.666667 631 1.457780e+02 2.4826150 *

3315) BOMDom_rainfall_max_.30< 2.666667 594 1.467822e+02 2.8891580 *

829) BOMDom_min_temperature_mean_30>=7.42 630 1.472225e+02 3.0895080

1658) SatDom_NIR..60>=6.703837 527 8.298231e+01 2.9842880 *

1659) SatDom_NIR..60< 6.703837 103 2.855353e+01 3.6278640 *

415) SatDom_Reflect_b1_max_160< 0.0479125 230 7.866030e+01 3.4783910 *

13) METADom_sub_cropFaba.Bean>=0.5 1532 2.074569e+03 3.0774870

26) SatDom_NIR_mean_170< 0.3475594 1015 1.157413e+03 2.7251820

52) BOMDom_rainfall_csum_160< 296 233 1.515554e+02 1.9136050 *

53) BOMDom_rainfall_csum_160>=296 782 8.066644e+02 2.9669950

106) BOMDom_max_temperature_min_.20>=17.95 522 4.127768e+02 2.6838890

212) SatDom_Reflect_b5_var_40>=5.376569e-05 291 1.974914e+02 2.3462540 *

213) SatDom_Reflect_b5_var_40< 5.376569e-05 231 1.403227e+02 3.1092210 *

107) BOMDom_max_temperature_min_.20< 17.95 260 2.680525e+02 3.5353850 *

27) SatDom_NIR_mean_170>=0.3475594 517 5.438448e+02 3.7691490

54) BOMDom_solar_exposure_max_240>=32.4 238 1.621752e+02 3.2372270 *

55) BOMDom_solar_exposure_max_240< 32.4 279 2.568855e+02 4.2229030 *

7) METADom_Dicot_TF< 0.5 41410 7.543284e+04 4.0321270

14) SatDom_EVI_mean_160< 0.5025781 36299 4.871755e+04 3.8170890

28) SatDom_Reflect_b7_min_180>=0.19295 14225 1.478815e+04 3.3494140

56) SatDom_Reflect_b7_mean_150>=0.1605481 7964 6.998821e+03 3.0768370

112) SatDom_Reflect_b2_mean..30>=0.2057325 5139 3.812163e+03 2.8444110

224) SatDom_red_band_var..20>=9.28106e-06 2909 1.640923e+03 2.5768680

448) SatDom_Reflect_b3_var_80>=4.125e-07 2505 1.127948e+03 2.4391300

896) SatDom_Subzero_nights_mean_70< 0.157 2344 8.582171e+02 2.3594160

1792) SatDom_Reflect_b7_var_100>=4.004224e-06 1962 6.462197e+02 2.2456370

3584) BOMDom_max_temperature_max_230>=42.75 218 4.264786e+01 1.5910550 *

3585) BOMDom_max_temperature_max_230< 42.75 1744 4.984876e+02 2.3274600

7170) METADom_Chem_rotation_minus_1_ESTERCIDE>=-1.453915e-17 1041 2.596425e+02 2.1612970

14340) SatDom_Reflect_b6_mean..80< 0.4185512 604 1.237761e+02 1.9565400 *

14341) SatDom_Reflect_b6_mean..80>=0.4185512 437 7.554331e+01 2.4443020 *

7171) METADom_Chem_rotation_minus_1_ESTERCIDE< -1.453915e-17 703 1.675416e+02 2.5735140

14342) SatDom_LST_day_var_60>=1.743537 589 8.451990e+01 2.4699830 *

14343) SatDom_LST_day_var_60< 1.743537 114 4.409012e+01 3.1084210 *

1793) SatDom_Reflect_b7_var_100< 4.004224e-06 382 5.614520e+01 2.9437960 *

897) SatDom_Subzero_nights_mean_70>=0.157 161 3.798508e+01 3.5996890 *

449) SatDom_Reflect_b3_var_80< 4.125e-07 404 1.707746e+02 3.4309160

898) METADom_Chem_rotation_minus_2_TRIFLURALIN< 0.05586137 170 2.200351e+01 2.9635880 *

899) METADom_Chem_rotation_minus_2_TRIFLURALIN>=0.05586137 234 8.467116e+01 3.7704270 *

225) SatDom_red_band_var..20< 9.28106e-06 2230 1.691391e+03 3.1934170

450) SatDom_LST_day_mean_190>=55.839 1845 1.118194e+03 3.0007590

900) BOMDom_solar_exposure_min_160< 18.6 1518 6.834229e+02 2.8119370

1800) SatDom_LST_day_var_30>=0.7775974 1388 5.014579e+02 2.7155260

3600) SatDom_Pot_latent_heat_mean_120< 1.00225e+07 252 3.674392e+01 1.9712300 *

3601) SatDom_Pot_latent_heat_mean_120>=1.00225e+07 1136 2.941439e+02 2.8806340

7202) SatDom_GPP_var..70< 1.175951e-06 788 1.528911e+02 2.7228550

14404) SatDom_LST_night_var_70< 1.662678 147 1.977246e+01 2.2646940 *

14405) SatDom_LST_night_var_70>=1.662678 641 9.518514e+01 2.8279250 *

7203) SatDom_GPP_var..70>=1.175951e-06 348 7.721737e+01 3.2379020

14406) SatDom_GPP_var_230< 9.959427e-07 221 3.314351e+01 3.0447510 *

14407) SatDom_GPP_var_230>=9.959427e-07 127 2.148145e+01 3.5740160 *

1801) SatDom_LST_day_var_30< 0.7775974 130 3.131488e+01 3.8413080 *

901) BOMDom_solar_exposure_min_160>=18.6 327 1.294014e+02 3.8773090

1802) METADom_Chem_rotation_minus_1_STARANE< 9.237403e-18 169 1.729161e+01 3.4074560 *

1803) METADom_Chem_rotation_minus_1_STARANE>=9.237403e-18 158 3.489480e+01 4.3798730 *

451) SatDom_LST_day_mean_190< 55.839 385 1.765391e+02 4.1166750

902) METADom_Chem_rotation_minus_2_CLETHODIM< -3.371869e-18 200 4.773379e+01 3.7012500 *

903) METADom_Chem_rotation_minus_2_CLETHODIM>=-3.371869e-18 185 5.697551e+01 4.5657840 *

113) SatDom_Reflect_b2_mean..30< 0.2057325 2825 2.404023e+03 3.4996460

226) SatDom_LST_day_max_220>=66.63 1767 1.140552e+03 3.1595300

452) SatDom_Emis31_var..40< 8.45679e-07 628 3.872279e+02 2.6277710

904) SatDom_PsnNet_var_100< 3.405435e-06 295 7.698652e+01 2.0492200 *

905) SatDom_PsnNet_var_100>=3.405435e-06 333 1.240242e+02 3.1403000

1810) SatDom_MIR_mean_180< 0.2873625 232 4.370297e+01 2.8740090 *

1811) SatDom_MIR_mean_180>=0.2873625 101 2.608040e+01 3.7519800 *

453) SatDom_Emis31_var..40>=8.45679e-07 1139 4.778358e+02 3.4527220

906) SatDom_Latent_heat..10>=6.36615e+07 1007 2.758505e+02 3.3301890

1812) BOMDom_min_temperature_var_20< 20.17161 839 1.873086e+02 3.2261860

3624) SatDom_NIR_var_30>=5.338747e-05 491 1.061917e+02 3.0663750 *

3625) SatDom_NIR_var_30< 5.338747e-05 348 5.088403e+01 3.4516670 *

1813) BOMDom_min_temperature_var_20>=20.17161 168 3.414527e+01 3.8495830 *

907) SatDom_Latent_heat..10< 6.36615e+07 132 7.152288e+01 4.3875000 *

227) SatDom_LST_day_max_220< 66.63 1058 7.176846e+02 4.0676840

454) SatDom_blue_band_var_90>=1.060352e-06 486 2.615106e+02 3.6213370

908) BOMDom_rainfall_mean_190< 0.5 132 3.369917e+01 3.0115910 *

909) BOMDom_rainfall_mean_190>=0.5 354 1.604354e+02 3.8487010

1818) METADom_Chem_rotation_minus_2_ROUNDUP_READY< -1.31839e-17 162 3.738091e+01 3.5025930 *

1819) METADom_Chem_rotation_minus_2_ROUNDUP_READY>=-1.31839e-17 192 8.727450e+01 4.1407290 *

455) SatDom_blue_band_var_90< 1.060352e-06 572 2.770842e+02 4.4469230

910) SatDom_GPP_min_140< 0.0324875 328 7.835912e+01 4.1395120 *

911) SatDom_GPP_min_140>=0.0324875 244 1.260612e+02 4.8601640 *

57) SatDom_Reflect_b7_mean_150< 0.1605481 6261 6.444961e+03 3.6961320

114) BOMDom_max_temperature_var_190< 11.51094 1678 1.443233e+03 3.1697140

228) BOMDom_min_temperature_max_.50< 18.85 852 4.965599e+02 2.6569250

456) SatDom_blue_band_70>=7.492544 463 1.396844e+02 2.2024620

912) SatDom_LAI_min_40< 0.5625 347 7.990454e+01 2.0356200 *

913) SatDom_LAI_min_40>=0.5625 116 2.122612e+01 2.7015520 *

457) SatDom_blue_band_70< 7.492544 389 1.474318e+02 3.1978410

914) BOMDom_rainfall_max_230< 15.8 252 9.986785e+01 2.9990080 *

915) BOMDom_rainfall_max_230>=15.8 137 1.927575e+01 3.5635770 *

229) BOMDom_min_temperature_max_.50>=18.85 826 4.915503e+02 3.6986440

458) SatDom_EvapoTrans_min_240< 2.475 451 1.509466e+02 3.2581370

916) SatDom_Reflect_b3_mean_80>=0.02386 132 2.487270e+01 2.7282580 *

917) SatDom_Reflect_b3_mean_80< 0.02386 319 7.367594e+01 3.4773980 *

459) SatDom_EvapoTrans_min_240>=2.475 375 1.478378e+02 4.2284270

918) SatDom_Latent_heat_max_170< 1433125 116 3.392189e+01 3.6447410 *

919) SatDom_Latent_heat_max_170>=1433125 259 5.669599e+01 4.4898460 *

115) BOMDom_max_temperature_var_190>=11.51094 4583 4.366474e+03 3.8888720

230) BOMDom_solar_exposure_var_60< 2.81 996 7.047091e+02 3.1902810

460) SatDom_GPP_var_210>=3.42996e-06 312 1.465580e+02 2.4464100

920) SatDom_red_band_max_110>=0.056475 106 2.907728e+01 1.9480190 *

921) SatDom_red_band_max_110< 0.056475 206 7.760261e+01 2.7028640 *

461) SatDom_GPP_var_210< 3.42996e-06 684 3.067583e+02 3.5295910

922) SatDom_PsnNet_max_110< 0.0288 389 1.050261e+02 3.1876090

1844) SatDom_LST_night_var_110< 3.262754 143 2.393578e+01 2.8538460 *

1845) SatDom_LST_night_var_110>=3.262754 246 5.590035e+01 3.3816260 *

923) SatDom_PsnNet_max_110>=0.0288 295 9.624771e+01 3.9805420 *

231) BOMDom_solar_exposure_var_60>=2.81 3587 3.040719e+03 4.0828490

462) SatDom_Emis32_mean_90>=0.9867167 957 7.261366e+02 3.5539290

924) BOMDom_solar_exposure_var_190< 52.32539 442 2.525538e+02 2.9947740

1848) METADom_Chem_rotation_minus_2_HAMMER>=0.007344062 100 2.767754e+01 2.2881000 *

1849) METADom_Chem_rotation_minus_2_HAMMER< 0.007344062 342 1.603355e+02 3.2014040

3698) BOMDom_Latitude>=-34.5404 231 8.032086e+01 3.0196100 *

3699) BOMDom_Latitude< -34.5404 111 5.649289e+01 3.5797300 *

925) BOMDom_solar_exposure_var_190>=52.32539 515 2.167848e+02 4.0338250

1850) SatDom_NDVI_max_150< 0.6264938 341 6.169114e+01 3.7036950 *

1851) SatDom_NDVI_max_150>=0.6264938 174 4.509609e+01 4.6808050 *

463) SatDom_Emis32_mean_90< 0.9867167 2630 1.949436e+03 4.2753120

926) BOMDom_rainfall_csum_200< 429.9 1509 9.828615e+02 3.9842680

1852) METADom_Fert_rotation_minus_2_AP>=0.0005970149 487 2.233977e+02 3.4794050

3704) SatDom_Reflect_b2_var_200>=0.0002735344 200 6.161640e+01 3.0325500 *

3705) SatDom_Reflect_b2_var_200< 0.0002735344 287 9.401572e+01 3.7908010 *

1853) METADom_Fert_rotation_minus_2_AP< 0.0005970149 1022 5.761839e+02 4.2248430

3706) METADom_Chem_rotation_minus_1_WETTER< -1.3726e-17 314 9.483771e+01 3.7425160 *

3707) METADom_Chem_rotation_minus_1_WETTER>=-1.3726e-17 708 3.758999e+02 4.4387570

7414) BOMDom_rainfall_mean_0>=2.41 278 8.325697e+01 3.9983810 *

7415) BOMDom_rainfall_mean_0< 2.41 430 2.038749e+02 4.7234650

14830) METADom_sub_cropWheat>=0.5 246 5.462565e+01 4.5043500 *

14831) METADom_sub_cropWheat< 0.5 184 1.216478e+02 5.0164130 *

927) BOMDom_rainfall_csum_200>=429.9 1121 6.666877e+02 4.6670920

1854) SatDom_NIR_var_100>=1.486808e-05 808 4.157741e+02 4.4390220

3708) SatDom_red_band_var_120>=3.224184e-06 611 1.757875e+02 4.2611620

7416) SatDom_Reflect_b2_min_60>=0.279275 235 6.170767e+01 3.9409790 *

7417) SatDom_Reflect_b2_min_60< 0.279275 376 7.493099e+01 4.4612770 *

3709) SatDom_red_band_var_120< 3.224184e-06 197 1.607102e+02 4.9906600 *

1855) SatDom_NIR_var_100< 1.486808e-05 313 1.003890e+02 5.2558470

3710) BOMDom_Longitude< 148.6988 187 4.606484e+01 5.0018720 *

3711) BOMDom_Longitude>=148.6988 126 2.436033e+01 5.6327780 *

29) SatDom_Reflect_b7_min_180< 0.19295 22074 2.881311e+04 4.1184710

58) SatDom_GPP_max_210< 0.0132875 7494 6.671861e+03 3.7493270

116) BOMDom_min_temperature_var_180< 16.81367 5907 4.973321e+03 3.5974010

232) METADom_Chem_rotation_minus_1_DUAL_GOLD< 0.1059912 5778 4.260938e+03 3.5531010

464) SatDom_Reflect_b7_min_230>=0.28805 805 4.286842e+02 2.9113910

928) SatDom_Reflect_b6_mean..40< 0.3766469 329 7.896976e+01 2.3011550

1856) BOMDom_solar_exposure_var_170< 6.368056 138 3.823677e+01 1.9819570 *

1857) BOMDom_solar_exposure_var_170>=6.368056 191 1.651359e+01 2.5317800 *

929) SatDom_Reflect_b6_mean..40>=0.3766469 476 1.425187e+02 3.3331720

1858) SatDom_EVI_mean_210>=0.1233206 332 4.902589e+01 3.1352110 *

1859) SatDom_EVI_mean_210< 0.1233206 144 5.048538e+01 3.7895830 *

465) SatDom_Reflect_b7_min_230< 0.28805 4973 3.447101e+03 3.6569780

930) SatDom_Emis31_var_160< 5.5e-07 1425 7.766209e+02 3.2653470

1860) SatDom_EVI_var_240< 2.596029e-08 273 1.130076e+02 2.6021250 *

1861) SatDom_EVI_var_240>=2.596029e-08 1152 5.150731e+02 3.4225170

3722) SatDom_Pot_EvapoTrans_var..10>=6.680417 780 2.366714e+02 3.2112310

7444) SatDom_Emis32_var_170< 7.95679e-07 548 1.027362e+02 3.0196530 *

7445) SatDom_Emis32_var_170>=7.95679e-07 232 6.631504e+01 3.6637500 *

3723) SatDom_Pot_EvapoTrans_var..10< 6.680417 372 1.705696e+02 3.8655380

7446) BOMDom_rainfall_csum_100>=245.4 263 1.035043e+02 3.6823190 *

7447) BOMDom_rainfall_csum_100< 245.4 109 3.693458e+01 4.3076150 *

931) SatDom_Emis31_var_160>=5.5e-07 3548 2.364142e+03 3.8142700

1862) SatDom_Pot_EvapoTrans_var_150< 0.1637708 257 8.117434e+01 2.6204670 *

1863) SatDom_Pot_EvapoTrans_var_150>=0.1637708 3291 1.888097e+03 3.9074960

3726) BOMDom_solar_exposure_max_160< 29.45 2781 1.478534e+03 3.7958430

7452) SatDom_red_band_var_50< 2.637777e-05 2493 1.156688e+03 3.6979220

14904) BOMDom_rainfall_csum_140< 217.8 201 4.483851e+01 2.8075620 *

14905) BOMDom_rainfall_csum_140>=217.8 2292 9.385354e+02 3.7760030

29810) SatDom_Reflect_b3_min_210>=0.0536125 1974 7.074673e+02 3.6891440

59620) SatDom_blue_band_var..70>=5.983227e-06 429 1.017394e+02 3.3263170

119240) METADom_sub_cropWheat>=0.5 305 4.777629e+01 3.1814100 *

119241) METADom_sub_cropWheat< 0.5 124 3.180587e+01 3.6827420 *

59621) SatDom_blue_band_var..70< 5.983227e-06 1545 5.335715e+02 3.7898900

119242) SatDom_Pot_latent_heat_mean..40< 1.041169e+07 519 1.411917e+02 3.5156650 *

119243) SatDom_Pot_latent_heat_mean..40>=1.041169e+07 1026 3.336087e+02 3.9286060

238486) SatDom_PsnNet_mean_140< 0.03758875 915 2.301104e+02 3.8657490 *

238487) SatDom_PsnNet_mean_140>=0.03758875 111 7.008183e+01 4.4467570 *

29811) SatDom_Reflect_b3_min_210< 0.0536125 318 1.237259e+02 4.3151890

59622) METADom_Fert_rotation_minus_1_UREA>=0.1496446 205 4.554372e+01 4.1148780 *

59623) METADom_Fert_rotation_minus_1_UREA< 0.1496446 113 5.503437e+01 4.6785840 *

7453) SatDom_red_band_var_50>=2.637777e-05 288 9.102073e+01 4.6434720 *

3727) BOMDom_solar_exposure_max_160>=29.45 510 1.858458e+02 4.5163330

7454) SatDom_GPP_var_180< 5.692786e-07 270 9.741914e+01 4.2475930 *

7455) SatDom_GPP_var_180>=5.692786e-07 240 4.698957e+01 4.8186670 *

233) METADom_Chem_rotation_minus_1_DUAL_GOLD>=0.1059912 129 1.931504e+02 5.5816280 *

117) BOMDom_min_temperature_var_180>=16.81367 1587 1.054714e+03 4.3148140

234) SatDom_LST_night_var_60>=2.758261 839 5.153724e+02 3.9287130

468) SatDom_GPP_var_150>=2.370768e-06 675 3.089187e+02 3.7171560

936) SatDom_NIR_150< 64.03331 131 2.653922e+01 2.9380920 *

937) SatDom_NIR_150>=64.03331 544 1.837238e+02 3.9047610

1874) METADom_Chem_rotation_minus_2_HAMMER< 0.007344062 296 7.865641e+01 3.6488510 *

1875) METADom_Chem_rotation_minus_2_HAMMER>=0.007344062 248 6.254549e+01 4.2102020 *

469) SatDom_GPP_var_150< 2.370768e-06 164 5.190065e+01 4.7994510 *

235) SatDom_LST_night_var_60< 2.758261 748 2.739791e+02 4.7478880

470) METADom_Chem_rotation_minus_2_AMINE>=0.05643867 141 3.057672e+01 4.0851060 *

471) METADom_Chem_rotation_minus_2_AMINE< 0.05643867 607 1.670763e+02 4.9018450

942) SatDom_NDVI_max_140< 0.7109313 158 1.975585e+01 4.5205700 *

943) SatDom_NDVI_max_140>=0.7109313 449 1.162694e+02 5.0360130 *

59) SatDom_GPP_max_210>=0.0132875 14580 2.059518e+04 4.3082070

118) METADom_TEBUCONAZOLE< 36.75 10666 1.298748e+04 4.1040560

236) SatDom_PsnNet_max_160< 0.0406375 8198 8.911115e+03 3.9520870

472) SatDom_FPAR_mean_40>=0.238625 6929 7.063697e+03 3.8271710

944) SatDom_Reflect_b4_var..70>=2.114366e-05 2780 2.709783e+03 3.4748020

1888) BOMDom_rainfall_mean_.50< 0.74 1303 8.721282e+02 3.0538300

3776) SatDom_EVI_var..60>=0.0001451431 116 2.170767e+01 1.6823280 *

3777) SatDom_EVI_var..60< 0.0001451431 1187 6.108990e+02 3.1878600

7554) SatDom_Reflect_b2_max_220< 0.3383 901 3.644380e+02 2.9767480

15108) SatDom_Reflect_b7_var_90< 1.615244e-05 223 7.507214e+01 2.4685650 *

15109) SatDom_Reflect_b7_var_90>=1.615244e-05 678 2.128343e+02 3.1438940

30218) SatDom_Reflect_b7_min_240>=0.2240937 336 8.739000e+01 2.8832740 *

30219) SatDom_Reflect_b7_min_240< 0.2240937 342 8.020060e+01 3.3999420 *

7555) SatDom_Reflect_b2_max_220>=0.3383 286 7.979933e+01 3.8529370 *

1889) BOMDom_rainfall_mean_.50>=0.74 1477 1.403028e+03 3.8461810

3778) SatDom_EvapoTrans_max_170>=21.95625 626 3.959926e+02 3.1552880

7556) SatDom_Latent_heat_mean..60< 876077.6 245 7.726848e+01 2.4884080 *

7557) SatDom_Latent_heat_mean..60>=876077.6 381 1.397006e+02 3.5841210

15114) SatDom_Reflect_b4_min_30< 0.07341875 262 6.182933e+01 3.3617180 *

15115) SatDom_Reflect_b4_min_30>=0.07341875 119 3.637960e+01 4.0737820 *

3779) SatDom_EvapoTrans_max_170< 21.95625 851 4.884172e+02 4.3544070

7558) SatDom_NIR_min_60< 0.3000281 648 3.073136e+02 4.1672070

15116) METADom_Chem_rotation_minus_1_MAP< 0.002812722 504 2.037318e+02 3.9934520

30232) SatDom_Latent_heat_min_130< 3690000 123 6.779780e+01 3.4498370 *

30233) SatDom_Latent_heat_min_130>=3690000 381 8.785078e+01 4.1689500 *

15117) METADom_Chem_rotation_minus_1_MAP>=0.002812722 144 3.510958e+01 4.7753470 *

7559) SatDom_NIR_min_60>=0.3000281 203 8.590741e+01 4.9519700 *

945) SatDom_Reflect_b4_var..70< 2.114366e-05 4149 3.777456e+03 4.0632730

1890) METADom_Chem_rotation_minus_2_ROUNDUP_MAX< 0.0004147799 1688 1.232576e+03 3.6916350

3780) SatDom_Reflect_b6_var..80>=3.971897e-06 1382 7.727313e+02 3.4986110

7560) BOMDom_max_temperature_mean_.40>=23.17 886 4.240775e+02 3.2488710

15120) SatDom_NDVI_var_110>=1.30634e-05 782 2.458484e+02 3.1470590

30240) SatDom_MIR_mean_250< 0.2504812 428 1.519453e+02 2.9522660 *

30241) SatDom_MIR_mean_250>=0.2504812 354 5.802816e+01 3.3825710 *

15121) SatDom_NDVI_var_110< 1.30634e-05 104 1.091718e+02 4.0144230 *

7561) BOMDom_max_temperature_mean_.40< 23.17 496 1.946846e+02 3.9447180

15122) SatDom_red_band_var..10< 1.445057e-05 337 1.025681e+02 3.7155490 *

15123) SatDom_red_band_var..10>=1.445057e-05 159 3.690547e+01 4.4304400 *

3781) SatDom_Reflect_b6_var..80< 3.971897e-06 306 1.758027e+02 4.5633990

7562) METADom_Chem_rotation_minus_3_UNKNOWN>=0.5 130 4.935794e+01 4.1379230 *

7563) METADom_Chem_rotation_minus_3_UNKNOWN< 0.5 176 8.552794e+01 4.8776700 *

1891) METADom_Chem_rotation_minus_2_ROUNDUP_MAX>=0.0004147799 2461 2.151832e+03 4.3181800

3782) SatDom_MIR_var_120< 3.535009e-05 1893 1.606429e+03 4.1286950

7564) METADom_Chem_rotation_minus_2_DIURON< 0.07728782 1683 1.166572e+03 4.0068920

15128) SatDom_Reflect_b6_mean_160>=0.2107931 867 4.366485e+02 3.7019260

30256) SatDom_red_band_var_40>=1.543842e-06 730 3.137330e+02 3.5664930

60512) METADom_Chem_rotation_minus_2_SUMMER< -8.912142e-18 141 3.764092e+01 3.1651770 *

60513) METADom_Chem_rotation_minus_2_SUMMER>=-8.912142e-18 589 2.479472e+02 3.6625640 *

30257) SatDom_red_band_var_40< 1.543842e-06 137 3.817895e+01 4.4235770 *

15129) SatDom_Reflect_b6_mean_160< 0.2107931 816 5.636146e+02 4.3309190

30258) SatDom_GPP_var_180>=4.848882e-06 412 1.912584e+02 3.9608010

60516) METADom_Fert_rotation_minus_2_DAP>=0.1650126 153 2.667372e+01 3.6041830 *

60517) METADom_Fert_rotation_minus_2_DAP< 0.1650126 259 1.336322e+02 4.1714670 *

30259) SatDom_GPP_var_180< 4.848882e-06 404 2.583609e+02 4.7083660

60518) METADom_Chem_rotation_minus_2_ESTER< 0.01843495 275 1.124721e+02 4.5258550 *

60519) METADom_Chem_rotation_minus_2_ESTER>=0.01843495 129 1.172005e+02 5.0974420 *

7565) METADom_Chem_rotation_minus_2_DIURON>=0.07728782 210 2.147804e+02 5.1048570 *

3783) SatDom_MIR_var_120>=3.535009e-05 568 2.509199e+02 4.9496830

7566) SatDom_blue_band_max_10< 0.041875 364 1.098662e+02 4.6927200 *

7567) SatDom_blue_band_max_10>=0.041875 204 7.413283e+01 5.4081860 *

473) SatDom_FPAR_mean_40< 0.238625 1269 1.148942e+03 4.6341530

946) SatDom_Reflect_b3_var_150< 6.542535e-07 219 1.452864e+02 3.5103650 *

947) SatDom_Reflect_b3_var_150>=6.542535e-07 1050 6.693949e+02 4.8685430

1894) SatDom_Reflect_b1_min_140< 0.0898875 881 4.432897e+02 4.6953460

3788) SatDom_Pot_EvapoTrans_min..10< 25.9 374 1.609462e+02 4.3743320 *

3789) SatDom_Pot_EvapoTrans_min..10>=25.9 507 2.153722e+02 4.9321500

7578) METADom_Chem_rotation_minus_1_LOGRAN>=0.00680445 193 8.934534e+01 4.6677720 *

7579) METADom_Chem_rotation_minus_1_LOGRAN< 0.00680445 314 1.042454e+02 5.0946500 *

1895) SatDom_Reflect_b1_min_140>=0.0898875 169 6.191106e+01 5.7714200 *

237) SatDom_PsnNet_max_160>=0.0406375 2468 3.258144e+03 4.6088530

474) SatDom_Reflect_b1_mean_210< 0.1463831 1398 1.425111e+03 4.1956370

948) METADom_Chemicals_time_Chemicals_PINOXADEN>=68.80806 140 6.175329e+01 3.0667140 *

949) METADom_Chemicals_time_Chemicals_PINOXADEN< 68.80806 1258 1.165076e+03 4.3212720

1898) SatDom_Reflect_b2_var_80>=0.0002299302 388 1.586000e+02 3.8603090 *

1899) SatDom_Reflect_b2_var_80< 0.0002299302 870 8.872624e+02 4.5268510

3798) SatDom_LAI_mean_60>=0.685 632 6.330824e+02 4.3780380 *

3799) SatDom_LAI_mean_60< 0.685 238 2.030190e+02 4.9220170 *

475) SatDom_Reflect_b1_mean_210>=0.1463831 1070 1.282449e+03 5.1487380

950) SatDom_EVI_var_60< 0.0006738782 678 7.438703e+02 4.7434370

1900) SatDom_Pot_EvapoTrans_var_70< 1.378265 370 2.022387e+02 4.3013240 *

1901) SatDom_Pot_EvapoTrans_var_70>=1.378265 308 3.824306e+02 5.2745450

3802) SatDom_Reflect_b3_var_10< 3.409966e-06 149 2.614728e+02 4.9168460 *

3803) SatDom_Reflect_b3_var_10>=3.409966e-06 159 8.402799e+01 5.6097480 *

951) SatDom_EVI_var_60>=0.0006738782 392 2.345708e+02 5.8497450

1902) SatDom_LST_day_max_60>=34.29 195 8.752717e+01 5.5024100 *

1903) SatDom_LST_day_max_60< 34.29 197 1.002323e+02 6.1935530 *

119) METADom_TEBUCONAZOLE>=36.75 3914 5.951769e+03 4.8645380

238) SatDom_LST_day_mean_170>=43.46017 3451 4.234131e+03 4.6686820

476) SatDom_blue_band_var_50< 6.074349e-06 3072 3.071793e+03 4.5035290

952) SatDom_Aero_q_190>=0.5 825 7.300013e+02 3.7460730

1904) BOMDom_min_temperature_mean_230>=13.375 515 2.478498e+02 3.2440580

3808) SatDom_NDVI_70>=72.86583 168 8.934609e+01 2.7377980 *

3809) SatDom_NDVI_70< 72.86583 347 9.459866e+01 3.4891640 *

1905) BOMDom_min_temperature_mean_230< 13.375 310 1.367438e+02 4.5800650

3810) SatDom_Reflect_b2_var..60< 9.589615e-05 126 2.737235e+01 4.0183330 *

3811) SatDom_Reflect_b2_var..60>=9.589615e-05 184 4.238739e+01 4.9647280 *

953) SatDom_Aero_q_190< 0.5 2247 1.694669e+03 4.7816330

1906) METADom_Chem_rotation_minus_1_DUAL< 0.01554059 1939 1.152487e+03 4.6183700

3812) SatDom_Reflect_b5_var_30>=1.511718e-05 1681 8.351467e+02 4.4875310

7624) SatDom_Pot_EvapoTrans_mean_240< 61.39187 479 1.498562e+02 4.0528600 *

7625) SatDom_Pot_EvapoTrans_mean_240>=61.39187 1202 5.587235e+02 4.6607490

15250) BOMDom_min_temperature_var_220>=20.24806 140 4.697944e+01 3.9363570 *

15251) BOMDom_min_temperature_var_220< 20.24806 1062 4.285955e+02 4.7562430

30502) SatDom_Reflect_b6_max_70< 0.21335 725 2.589860e+02 4.5925100 *

30503) SatDom_Reflect_b6_max_70>=0.21335 337 1.083599e+02 5.1084870 *

3813) SatDom_Reflect_b5_var_30< 1.511718e-05 258 1.010680e+02 5.4708530 *

1907) METADom_Chem_rotation_minus_1_DUAL>=0.01554059 308 1.651264e+02 5.8094480

3814) SatDom_Reflect_b2_90< 48.00239 176 8.497180e+01 5.5092610 *

3815) SatDom_Reflect_b2_90>=48.00239 132 4.314859e+01 6.2096970 *

477) SatDom_blue_band_var_50>=6.074349e-06 379 3.993818e+02 6.0073350

954) SatDom_FPAR_min_10>=0.24 219 1.138246e+02 5.3831960 *

955) SatDom_FPAR_min_10< 0.24 160 8.347618e+01 6.8616250 *

239) SatDom_LST_day_mean_170< 43.46017 463 5.985644e+02 6.3243630

478) METADom_Chem_rotation_minus_2_CLETHODIM>=-4.445229e-18 282 1.668029e+02 5.7595040 *

479) METADom_Chem_rotation_minus_2_CLETHODIM< -4.445229e-18 181 2.016003e+02 7.2044200 *

15) SatDom_EVI_mean_160>=0.5025781 5111 1.311580e+04 5.5593520

30) METADom_Nitrogen_fert_repeat_2< 23.5 3583 7.285548e+03 5.1026400

60) SatDom_EvapoTrans_var_130>=0.08264779 3450 5.055707e+03 4.9748810

120) SatDom_LST_day_mean_30>=35.38345 707 8.906126e+02 4.0735500

240) SatDom_Pot_EvapoTrans_max_170< 47.24375 114 5.627145e+01 2.6750000 *

241) SatDom_Pot_EvapoTrans_max_170>=47.24375 593 5.684979e+02 4.3424110

482) SatDom_Reflect_b2_var.0< 0.0001513827 345 2.853480e+02 3.9333040

964) SatDom_Reflect_b1_mean_150< 0.03914813 110 5.436635e+01 3.5688180 *

965) SatDom_Reflect_b1_mean_150>=0.03914813 235 2.095278e+02 4.1039150 *

483) SatDom_Reflect_b2_var.0>=0.0001513827 248 1.450808e+02 4.9115320 *

121) SatDom_LST_day_mean_30< 35.38345 2743 3.442689e+03 5.2071970

242) METADom_Chem_rotation_minus_1_MCPA< 0.1362487 2436 2.515450e+03 5.0401930

484) SatDom_Emis32..20< 70.05867 2008 1.897736e+03 4.8847110

968) SatDom_Reflect_b5_mean_70>=0.3440331 369 2.738456e+02 4.2024390

1936) SatDom_red_band_mean_80>=0.04649141 106 1.238288e+02 3.6303770 *

1937) SatDom_red_band_mean_80< 0.04649141 263 1.013467e+02 4.4330040 *

969) SatDom_Reflect_b5_mean_70< 0.3440331 1639 1.413451e+03 5.0383160

1938) SatDom_Latent_heat_max_130>=5655625 1139 9.240645e+02 4.8419840

3876) METADom_Chem_rotation_minus_1_TRIF>=0.007181737 112 8.506564e+01 4.0115180 *

3877) METADom_Chem_rotation_minus_1_TRIF< 0.007181737 1027 7.533315e+02 4.9325510

7754) SatDom_LST_day_var.0< 1.92589 240 1.714968e+02 4.5752500 *

7755) SatDom_LST_day_var.0>=1.92589 787 5.418517e+02 5.0415120 *

1939) SatDom_Latent_heat_max_130< 5655625 500 3.454691e+02 5.4855600

3878) SatDom_EvapoTrans_var_150>=14.63982 167 1.139985e+02 4.9370060 *

3879) SatDom_EvapoTrans_var_150< 14.63982 333 1.560169e+02 5.7606610

7758) SatDom_EvapoTrans_max_240< 7.141667 224 7.094051e+01 5.5556250 *

7759) SatDom_EvapoTrans_max_240>=7.141667 109 5.630736e+01 6.1820180 *

485) SatDom_Emis32..20>=70.05867 428 3.414294e+02 5.7696500

970) SatDom_FPAR_var_230< 0.0002526389 122 8.506484e+01 5.1163110 *

971) SatDom_FPAR_var_230>=0.0002526389 306 1.835266e+02 6.0301310

1942) METADom_Fert_rotation_minus_1_EXTRA>=-5.499073e-17 182 6.966972e+01 5.7946150 *

1943) METADom_Fert_rotation_minus_1_EXTRA< -5.499073e-17 124 8.894482e+01 6.3758060 *

243) METADom_Chem_rotation_minus_1_MCPA>=0.1362487 307 3.202005e+02 6.5323450

486) BOMDom_rainfall_mean_.30>=0.8825 140 1.883314e+02 6.0312860 *

487) BOMDom_rainfall_mean_.30< 0.8825 167 6.725484e+01 6.9523950 *

61) SatDom_EvapoTrans_var_130< 0.08264779 133 7.127975e+02 8.4166920 *

31) METADom_Nitrogen_fert_repeat_2>=23.5 1528 3.330400e+03 6.6302950

62) METADom_Chem_rotation_minus_2_CADENCE>=0.005310412 535 5.345056e+02 5.4458500

124) SatDom_Reflect_b3_var_180< 4.185924e-06 197 1.191096e+02 4.6709140 *

125) SatDom_Reflect_b3_var_180>=4.185924e-06 338 2.281399e+02 5.8975150

250) BOMDom_max_temperature_var_190< 23.16361 199 6.231810e+01 5.5556280 *

251) BOMDom_max_temperature_var_190>=23.16361 139 1.092605e+02 6.3869780 *

63) METADom_Chem_rotation_minus_2_CADENCE< 0.005310412 993 1.640960e+03 7.2684390

126) SatDom_red_band_min_70>=0.05215 251 1.015245e+02 5.9109960 *

127) SatDom_red_band_min_70< 0.05215 742 9.204764e+02 7.7276280

254) SatDom_EVI_var..60< 6.729818e-06 295 1.457163e+02 6.9210170 *

255) SatDom_EVI_var..60>=6.729818e-06 447 4.561594e+02 8.2599550

510) SatDom_LAI_max_30< 1.05 328 2.521511e+02 8.0359760 *

511) SatDom_LAI_max_30>=1.05 119 1.421993e+02 8.8773110 *

**Wheat Regression tree: Predicted at Time of Sowing**

**Leaf values indicate node-average yield, in t/Ha**

**Model Accuracy: Missing Site Prediction R^2^ = 0.44; p<0.0001**

**n= 52362**

**node), split, n, deviance, yval**

*** denotes terminal node**

1) root 52362 1.270457e+05 3.3751660

2) METADom_Fertiliser_time_MAP< -2.744775 30869 5.644097e+04 2.9035900

4) SatDom_MIR..20>=20.35942 13975 1.761428e+04 2.3381040

8) SatDom_GPP_min.0< 0.00855 12132 1.302614e+04 2.2019530

16) METADom_Nitrogen_fert_repeat_1< 25.25 10210 9.539180e+03 2.0722000

32) SatDom_LST_night_var..60>=3.438542 5658 4.129658e+03 1.7788230

64) BOMDom_solar_exposure_var_.50>=0.9958333 5259 3.533770e+03 1.7021220

128) SatDom_FPAR_mean..20< 0.226875 5199 3.079668e+03 1.6708520

256) SatDom_EvapoTrans_var..30>=0.004807292 4982 2.534367e+03 1.6160000

512) SatDom_LST_night_max..70< 45.39 4768 2.206185e+03 1.5644880

1024) SatDom_Reflect_b7_mean.0>=0.2307531 4045 1.481878e+03 1.4669020

2048) SatDom_EvapoTrans_min..10< 2.925 647 1.490394e+02 0.9569088 *

2049) SatDom_EvapoTrans_min..10>=2.925 3398 1.132517e+03 1.5640080

4098) METADom_Chem_rotation_minus_1_VERDICT< 0.01157271 2547 7.319567e+02 1.4538910

8196) BOMDom_rainfall_csum_.10< 72.05 1892 4.547019e+02 1.3163790 *

8197) BOMDom_rainfall_csum_.10>=72.05 655 1.381360e+02 1.8510990 *

4099) METADom_Chem_rotation_minus_1_VERDICT>=0.01157271 851 2.772400e+02 1.8935840

8198) METADom_Chem_rotation_minus_1_STARANE< 0.002376174 736 1.096868e+02 1.7342260 *

8199) METADom_Chem_rotation_minus_1_STARANE>=0.002376174 115 2.924121e+01 2.9134780 *

1025) SatDom_Reflect_b7_mean.0< 0.2307531 723 4.702727e+02 2.1104560

2050) SatDom_Reflect_b3_max..40< 0.07105 244 5.239716e+01 1.3469670 *

2051) SatDom_Reflect_b3_max..40>=0.07105 479 2.031922e+02 2.4993740 *

513) SatDom_LST_night_max..70>=45.39 214 3.365058e+01 2.7636920 *

257) SatDom_EvapoTrans_var..30< 0.004807292 217 1.861670e+02 2.9301840 *

129) SatDom_FPAR_mean..20>=0.226875 60 8.520433e+00 4.4116670 *

65) BOMDom_solar_exposure_var_.50< 0.9958333 399 1.571623e+02 2.7897740 *

33) SatDom_LST_night_var..60< 3.438542 4552 4.317231e+03 2.4368590

66) METADom_Fert_rotation_minus_2_EIGHTEEN_TO_TWENTY>=-2.619432e-17 4233 3.338309e+03 2.3303430

132) METADom_Nitrogen_fert_repeat_2< 12.2 4069 2.644006e+03 2.2505460

264) SatDom_Latent_heat..30< 4.488652e+07 1182 4.842627e+02 1.7956430

528) SatDom_NIR_mean..30< 0.2329247 211 1.249245e+01 0.9846919 *

529) SatDom_NIR_mean..30>=0.2329247 971 3.028545e+02 1.9718640

1058) SatDom_MIR_min..30>=0.2859 808 1.341342e+02 1.8030690 *

1059) SatDom_MIR_min..30< 0.2859 163 3.158138e+01 2.8085890 *

265) SatDom_Latent_heat..30>=4.488652e+07 2887 1.815000e+03 2.4367930

530) METADom_Date_soil_test_10cm_numeric.test_10cm< -37 605 2.663244e+02 1.8459670 *

531) METADom_Date_soil_test_10cm_numeric.test_10cm>=-37 2282 1.281495e+03 2.5934310

1062) SatDom_Emis31_mean.0>=0.983325 840 4.134280e+02 2.1672860

2124) BOMDom_solar_exposure_mean_.70< 20.83 522 9.447939e+01 1.8177200 *

2125) BOMDom_solar_exposure_mean_.70>=20.83 318 1.504565e+02 2.7411010 *

1063) SatDom_Emis31_mean.0< 0.983325 1442 6.266629e+02 2.8416710

2126) SatDom_MIR_var.0>=4.363516e-05 782 2.859824e+02 2.5375190 *

2127) SatDom_MIR_var.0< 4.363516e-05 660 1.826249e+02 3.2020450 *

133) METADom_Nitrogen_fert_repeat_2>=12.2 164 2.555149e+01 4.3101830 *

67) METADom_Fert_rotation_minus_2_EIGHTEEN_TO_TWENTY< -2.619432e-17 319 2.936087e+02 3.8502820

134) SatDom_Reflect_b6..10>=32.70816 75 1.531842e+01 2.5225330 *

135) SatDom_Reflect_b6..10< 32.70816 244 1.054305e+02 4.2584020 *

17) METADom_Nitrogen_fert_repeat_1>=25.25 1922 2.401941e+03 2.8912230

34) SatDom_Pot_EvapoTrans_max..70< 47.225 224 5.151116e+01 1.3004460 *

35) SatDom_Pot_EvapoTrans_max..70>=47.225 1698 1.708804e+03 3.1010780

70) SatDom_Reflect_b2_min..40>=0.293225 254 9.637412e+01 1.9462990 *

71) SatDom_Reflect_b2_min..40< 0.293225 1444 1.214138e+03 3.3042040

142) SatDom_NDVI_mean.0>=0.2567766 543 2.171724e+02 2.6747880 *

143) SatDom_NDVI_mean.0< 0.2567766 901 6.522052e+02 3.6835290

286) METADom_Chem_rotation_minus_1_TERBYNE>=0.0002780193 396 1.480215e+02 3.0698230 *

287) METADom_Chem_rotation_minus_1_TERBYNE< 0.0002780193 505 2.380808e+02 4.1647720 *

9) SatDom_GPP_min.0>=0.00855 1843 2.882834e+03 3.2343520

18) METADom_Chem_rotation_minus_2_LONTREL< 0.1645563 1665 2.008914e+03 3.0324020

36) METADom_Fert_rotation_minus_2_MN>=0.002831506 403 2.042544e+02 2.0028290 *

37) METADom_Fert_rotation_minus_2_MN< 0.002831506 1262 1.241055e+03 3.3611810

74) METADom_Chem_rotation_minus_1_SELECT>=0.03011595 572 3.555412e+02 2.7717310

148) SatDom_EvapoTrans_min..30< 6.075 383 1.204089e+02 2.3722720 *

149) SatDom_EvapoTrans_min..30>=6.075 189 5.017222e+01 3.5812170 *

75) METADom_Chem_rotation_minus_1_SELECT< 0.03011595 690 5.220172e+02 3.8498260

150) BOMDom_max_temperature_max_.10< 22.95 87 1.577684e+01 2.3917240 *

151) BOMDom_max_temperature_max_.10>=22.95 603 2.945862e+02 4.0601990 *

19) METADom_Chem_rotation_minus_2_LONTREL>=0.1645563 178 1.708416e+02 5.1233710 *

5) SatDom_MIR..20< 20.35942 16894 3.066112e+04 3.3713700

10) METADom_Nitrogen_fert_repeat_3< 23.5 16639 2.648903e+04 3.3157270

20) METADom_Organic_C_10cm.test_10cm< 1.22826 9484 1.257128e+04 3.0328020

40) METADom_Chem_rotation_minus_2_GOAL>=0.009227678 1735 1.862829e+03 2.3408360

80) SatDom_LST_day_max..20< 49.24 817 3.797691e+02 1.7108080

160) SatDom_Reflect_b7_var..80< 0.0001322411 644 1.449233e+02 1.4686490 *

161) SatDom_Reflect_b7_var..80>=0.0001322411 173 5.650022e+01 2.6122540 *

81) SatDom_LST_day_max..20>=49.24 918 8.701480e+02 2.9015470

162) METADom_Chem_rotation_minus_1_BPOWER>=-1.498367e-17 312 2.008667e+02 2.0731090

324) SatDom_Reflect_b2_var..80>=5.62804e-05 156 1.466934e+01 1.4260260 *

325) SatDom_Reflect_b2_var..80< 5.62804e-05 156 5.555769e+01 2.7201920 *

163) METADom_Chem_rotation_minus_1_BPOWER< -1.498367e-17 606 3.449084e+02 3.3280690 *

41) METADom_Chem_rotation_minus_2_GOAL< 0.009227678 7749 9.691698e+03 3.1877330

82) SatDom_Pot_EvapoTrans_mean..10< 28.76 1742 1.633885e+03 2.5965670

164) METADom_Phos_fert< 14.3 930 5.970031e+02 2.1124730

328) SatDom_blue_band_var..70>=3.418763e-07 782 2.930138e+02 1.8759850

656) METADom_Chem_rotation_minus_1_ROUNDUP>=0.1852371 378 7.399315e+01 1.4519580 *

657) METADom_Chem_rotation_minus_1_ROUNDUP< 0.1852371 404 8.746660e+01 2.2727230 *

329) SatDom_blue_band_var..70< 3.418763e-07 148 2.916959e+01 3.3620270 *

165) METADom_Phos_fert>=14.3 812 5.693246e+02 3.1510100

330) METADom_Chem_rotation_minus_2_ROUNDUP_CT< 0.0008120926 207 5.418217e+01 2.2267150 *

331) METADom_Chem_rotation_minus_2_ROUNDUP_CT>=0.0008120926 605 2.777908e+02 3.4672560 *

83) SatDom_Pot_EvapoTrans_mean..10>=28.76 6007 7.272480e+03 3.3591680

166) BOMDom_Sowing_date_numeric>=15469 3002 3.163593e+03 3.0360660

332) SatDom_Reflect_b2_mean..40>=0.23491 589 2.017936e+02 2.1787610 *

333) SatDom_Reflect_b2_mean..40< 0.23491 2413 2.423232e+03 3.2453290

666) SatDom_LST_day_var..60>=5.878558 1736 1.398841e+03 2.9830130

1332) SatDom_EVI..30< 10.87218 1487 1.008090e+03 2.8180560

2664) SatDom_LST_night_min..50< 27.985 355 8.051228e+01 2.1066760 *

2665) SatDom_LST_night_min..50>=27.985 1132 6.915865e+02 3.0411480

5330) SatDom_Pot_latent_heat_var..60>=3.344876e+11 695 2.860807e+02 2.7256400 *

5331) SatDom_Pot_latent_heat_var..60< 3.344876e+11 437 2.262923e+02 3.5429290 *

1333) SatDom_EVI..30>=10.87218 249 1.086530e+02 3.9681120 *

667) SatDom_LST_day_var..60< 5.878558 677 5.986263e+02 3.9179760

1334) BOMDom_solar_exposure_min_.50< 13 385 1.662574e+02 3.3882340 *

1335) BOMDom_solar_exposure_min_.50>=13 292 1.818755e+02 4.6164380 *

167) BOMDom_Sowing_date_numeric< 15469 3005 3.482414e+03 3.6819470

334) SatDom_EVI_var.0>=4.576405e-05 806 1.075190e+03 2.8757200

668) BOMDom_rainfall_csum_.30< 77.6 540 4.051455e+02 2.3532590

1336) METADom_Fert_rotation_minus_1_N< 0.1096827 449 1.906640e+02 2.0785970 *

1337) METADom_Fert_rotation_minus_1_N>=0.1096827 91 1.348098e+01 3.7084620 *

669) BOMDom_rainfall_csum_.30>=77.6 266 2.234088e+02 3.9363530 *

335) SatDom_EVI_var.0< 4.576405e-05 2199 1.691296e+03 3.9774530

670) BOMDom_solar_exposure_mean_.80< 22.295 1442 8.880756e+02 3.7307210

1340) METADom_Fert_rotation_minus_1_SOA>=0.008581384 205 5.302592e+01 2.8348290 *

1341) METADom_Fert_rotation_minus_1_SOA< 0.008581384 1237 6.432444e+02 3.8791920 *

671) BOMDom_solar_exposure_mean_.80>=22.295 757 5.482166e+02 4.4474500

1342) SatDom_Reflect_b3_var..60>=2.51065e-06 482 2.787905e+02 4.1083200 *

1343) SatDom_Reflect_b3_var..60< 2.51065e-06 275 1.168294e+02 5.0418550 *

21) METADom_Organic_C_10cm.test_10cm>=1.22826 7155 1.215232e+04 3.6907460

42) SatDom_Reflect_b5.0>=28.84638 2036 2.864697e+03 3.0031780

84) METADom_Chem_rotation_minus_1_AVADEX_XTRA< -3.801213e-17 974 6.381435e+02 2.3658010

168) BOMDom_max_temperature_min_.10< 19.1 830 2.546666e+02 2.1385300 *

169) BOMDom_max_temperature_min_.10>=19.1 144 9.350132e+01 3.6757640 *

85) METADom_Chem_rotation_minus_1_AVADEX_XTRA>=-3.801213e-17 1062 1.467968e+03 3.5877400

170) SatDom_MixClouds..40< 1.5 924 7.094151e+02 3.2743830

340) SatDom_red_band_mean..10>=0.1148681 434 2.572810e+02 2.7300460

680) SatDom_Emis32_var..30>=7.166667e-07 276 7.990418e+01 2.3156520 *

681) SatDom_Emis32_var..30< 7.166667e-07 158 4.718957e+01 3.4539240 *

341) SatDom_red_band_mean..10< 0.1148681 490 2.096399e+02 3.7565100 *

171) SatDom_MixClouds..40>=1.5 138 6.032615e+01 5.6858700 *

43) SatDom_Reflect_b5.0< 28.84638 5119 7.942272e+03 3.9642160

86) METADom_Chem_rotation_minus_2_AMINE< 0.05179873 3725 4.536265e+03 3.6706500

172) SatDom_Pot_EvapoTrans_max..20>=36.39375 2475 2.117891e+03 3.3020970

344) SatDom_NDVI_mean..40< 0.2613728 1432 9.433761e+02 2.8864660

688) METADom_Fert_rotation_minus_2_AP>=-1.43982e-17 1267 5.810103e+02 2.7206160

1376) BOMDom_max_temperature_var_.80>=25.4305 161 5.057305e+01 1.7422980 *

1377) BOMDom_max_temperature_var_.80< 25.4305 1106 3.539120e+02 2.8630290 *

689) METADom_Fert_rotation_minus_2_AP< -1.43982e-17 165 5.990360e+01 4.1600000 *

345) SatDom_NDVI_mean..40>=0.2613728 1043 5.875004e+02 3.8727420

690) METADom_Chem_rotation_minus_1_B>=-9.725293e-18 685 2.623118e+02 3.5576930 *

691) METADom_Chem_rotation_minus_1_B< -9.725293e-18 358 1.271056e+02 4.4755590 *

173) SatDom_Pot_EvapoTrans_max..20< 36.39375 1250 1.416552e+03 4.4003840

346) SatDom_blue_band..70>=0.8397844 1034 6.817929e+02 4.1159770

692) SatDom_Reflect_b5..70>=7.191294 145 3.650268e+01 2.8340000 *

693) SatDom_Reflect_b5..70< 7.191294 889 3.681196e+02 4.3250730 *

347) SatDom_blue_band..70< 0.8397844 216 2.507455e+02 5.7618520

694) BOMDom_max_temperature_mean_.60< 30.555 187 8.274136e+01 5.4230480 *

695) BOMDom_max_temperature_mean_.60>=30.555 29 8.124255e+00 7.9465520 *

87) METADom_Chem_rotation_minus_2_AMINE>=0.05179873 1394 2.227152e+03 4.7486730

174) SatDom_Reflect_b5_var..10< 0.0001379334 724 7.642276e+02 3.9922380

348) METADom_pH_water_60cm.test_60cm>=8.45 222 3.660934e+01 2.8194590 *

349) METADom_pH_water_60cm.test_60cm< 8.45 502 2.872464e+02 4.5108760 *

175) SatDom_Reflect_b5_var..10>=0.0001379334 670 6.009982e+02 5.5660750

350) METADom_Chem_rotation_minus_1_LEXONE>=-3.335006e-17 391 1.468708e+02 4.9949100 *

351) METADom_Chem_rotation_minus_1_LEXONE< -3.335006e-17 279 1.478117e+02 6.3665230 *

11) METADom_Nitrogen_fert_repeat_3>=23.5 255 7.590051e+02 7.0021570

22) METADom_Chem_rotation_minus_1_ECLIPSE>=0.0003773585 42 1.742885e+01 3.6419050 *

23) METADom_Chem_rotation_minus_1_ECLIPSE< 0.0003773585 213 1.738309e+02 7.6647420 *

3) METADom_Fertiliser_time_MAP>=-2.744775 21493 5.388060e+04 4.0524590

6) BOMDom_Sowing_date_numeric< 16579 16446 3.478598e+04 3.7723030

12) SatDom_Reflect_b7..50>=11.14287 7126 1.130453e+04 3.1839910

24) BOMDom_rainfall_csum_0< 59.2 1811 2.638034e+03 2.2418330

48) SatDom_Pot_latent_heat_max..50< 1.462625e+07 1068 6.685221e+02 1.5424340

96) SatDom_LST_night_var..70< 11.97088 867 2.524976e+02 1.2510840 *

97) SatDom_LST_night_var..70>=11.97088 201 2.498096e+01 2.7991540 *

49) SatDom_Pot_latent_heat_max..50>=1.462625e+07 743 6.961537e+02 3.2471600

98) SatDom_PsnNet_min..10< 0.0041625 237 4.263389e+01 2.2997890 *

99) SatDom_PsnNet_min..10>=0.0041625 506 3.411803e+02 3.6908890

198) METADom_Fert_rotation_minus_2_UAN< 0.003975301 456 1.377251e+02 3.4932890 *

199) METADom_Fert_rotation_minus_2_UAN>=0.003975301 50 2.327025e+01 5.4930000 *

25) BOMDom_rainfall_csum_0>=59.2 5315 6.511198e+03 3.5050160

50) SatDom_EVI_min..60< 0.1249187 2356 2.264651e+03 3.0748600

100) SatDom_Reflect_b5_mean.0>=0.2713437 1407 9.975066e+02 2.6641510

200) SatDom_Reflect_b7_var..40>=2.223633e-06 1347 6.668057e+02 2.5666820

400) SatDom_LST_day_var..80>=14.07252 585 2.022912e+02 2.1605470 *

401) SatDom_LST_day_var..80< 14.07252 762 2.939422e+02 2.8784780 *

201) SatDom_Reflect_b7_var..40< 2.223633e-06 60 3.061547e+01 4.8523330 *

101) SatDom_Reflect_b5_mean.0< 0.2713437 949 6.779313e+02 3.6837830

202) METADom_Chem_rotation_minus_2_CADENCE< 0.007547092 797 2.994458e+02 3.4387330 *

203) METADom_Chem_rotation_minus_2_CADENCE>=0.007547092 152 7.967834e+01 4.9686840 *

51) SatDom_EVI_min..60>=0.1249187 2959 3.463504e+03 3.8475130

102) METADom_Fert_rotation_minus_1_LEGUME< -3.46511e-17 270 2.080112e+02 2.5700000 *

103) METADom_Fert_rotation_minus_1_LEGUME>=-3.46511e-17 2689 2.770597e+03 3.9757870

206) SatDom_Emis32_mean.0< 0.98695 2439 2.269419e+03 3.8583800

412) SatDom_Emis32_var..60>=8.067901e-07 909 6.127603e+02 3.4493730

824) BOMDom_rainfall_mean_.50< 0.45 492 1.612064e+02 2.9946340 *

825) BOMDom_rainfall_mean_.50>=0.45 417 2.297767e+02 3.9858990 *

413) SatDom_Emis32_var..60< 8.067901e-07 1530 1.414251e+03 4.1013790

826) SatDom_Reflect_b7_max..40< 0.369375 1296 1.067746e+03 3.9466440

1652) SatDom_Reflect_b2_mean..10>=0.2769937 119 3.490636e+01 2.8246220 *

1653) SatDom_Reflect_b2_mean..10< 0.2769937 1177 8.678796e+02 4.0600850 *

827) SatDom_Reflect_b7_max..40>=0.369375 234 1.436150e+02 4.9583760 *

207) SatDom_Emis32_mean.0>=0.98695 250 1.395654e+02 5.1212000 *

13) SatDom_Reflect_b7..50< 11.14287 9320 1.912928e+04 4.2221210

26) SatDom_EvapoTrans_mean..60>=1.76875 9069 1.596272e+04 4.1355890

52) METADom_Nitrogen_fert_repeat_3< 38.05 8897 1.291908e+04 4.0633290

104) SatDom_PsnNet..80< 0.2958688 1666 2.329826e+03 3.3545320

208) METADom_Chem_rotation_minus_2_INTERVIX< 0.1307781 1609 1.575736e+03 3.2307580

416) SatDom_Reflect_b6_var..80< 8.34542e-06 230 7.541532e+01 1.9353480 *

417) SatDom_Reflect_b6_var..80>=8.34542e-06 1379 1.049987e+03 3.4468170

834) METADom_Fert_rotation_minus_1_SULPHATE< 0.1229952 1258 7.822652e+02 3.3137200

1668) METADom_Fert_rotation_minus_1_DAP_ZN< -4.250073e-17 186 8.410769e+01 2.4063980 *

1669) METADom_Fert_rotation_minus_1_DAP_ZN>=-4.250073e-17 1072 5.184683e+02 3.4711470

3338) SatDom_Latent_heat..60< 3.394e+07 530 1.578817e+02 3.0663580 *

3339) SatDom_Latent_heat..60>=3.394e+07 542 1.888240e+02 3.8669740 *

835) METADom_Fert_rotation_minus_1_SULPHATE>=0.1229952 121 1.374646e+01 4.8305790 *

209) METADom_Chem_rotation_minus_2_INTERVIX>=0.1307781 57 3.362596e+01 6.8484210 *

105) SatDom_PsnNet..80>=0.2958688 7231 9.559430e+03 4.2266340

210) METADom_Chem_rotation_minus_2_PMAX>=0.02007657 529 6.227402e+02 3.0396790

420) SatDom_Reflect_b3_var..30>=1.225907e-06 382 1.254988e+02 2.4593190 *

421) SatDom_Reflect_b3_var..30< 1.225907e-06 147 3.422590e+01 4.5478230 *

211) METADom_Chem_rotation_minus_2_PMAX< 0.02007657 6702 8.132575e+03 4.3203220

422) METADom_Fert_rotation_minus_2_ZINC>=0.01946467 5307 5.800315e+03 4.1717170

844) METADom_Chem_rotation_minus_2_AVADEX< 0.02555631 3878 3.755200e+03 3.9816580

1688) SatDom_Latent_heat_var..40< 7.584435e+11 3581 3.078861e+03 3.8879530

3376) SatDom_Reflect_b5_var..60< 4.734435e-06 227 1.296012e+02 2.7000000 *

3377) SatDom_Reflect_b5_var..60>=4.734435e-06 3354 2.607229e+03 3.9683540

6754) SatDom_Reflect_b5_min..30>=0.298075 1037 7.518303e+02 3.5483800

13508) SatDom_Reflect_b2_min..50< 0.222675 300 9.338577e+01 2.7788000 *

13509) SatDom_Reflect_b2_min..50>=0.222675 737 4.084445e+02 3.8616420

27018) SatDom_LAI_min..40>=0.275 586 2.279166e+02 3.6414330 *

27019) SatDom_LAI_min..40< 0.275 151 4.183455e+01 4.7162250 *

6755) SatDom_Reflect_b5_min..30< 0.298075 2317 1.590633e+03 4.1563190

13510) SatDom_MixClouds_csum.0>=8.5 400 1.801427e+02 3.4379750 *

13511) SatDom_MixClouds_csum.0< 8.5 1917 1.161015e+03 4.3062080

27022) SatDom_Reflect_b1_var..30>=8.317938e-06 1552 8.003793e+02 4.1717010 *

27023) SatDom_Reflect_b1_var..30< 8.317938e-06 365 2.131641e+02 4.8781370 *

1689) SatDom_Latent_heat_var..40>=7.584435e+11 297 2.657743e+02 5.1114810 *

845) METADom_Chem_rotation_minus_2_AVADEX>=0.02555631 1429 1.524880e+03 4.6874950

1690) BOMDom_Latitude>=-34.3344 485 5.448471e+02 3.9950520

3380) SatDom_LAI_max..60< 0.8875 251 7.889636e+01 3.4118330 *

3381) SatDom_LAI_max..60>=0.8875 234 2.889958e+02 4.6206410 *

1691) BOMDom_Latitude< -34.3344 944 6.280107e+02 5.0432520

3382) METADom_Chem_rotation_minus_1_DIURON>=0.209345 343 1.381464e+02 4.5376380 *

3383) METADom_Chem_rotation_minus_1_DIURON< 0.209345 601 3.521341e+02 5.3318140 *

423) METADom_Fert_rotation_minus_2_ZINC< 0.01946467 1395 1.769206e+03 4.8856630

846) METADom_total_N_60cm.test_60cm< 7.05 167 1.012400e+02 3.2668260 *

847) METADom_total_N_60cm.test_60cm>=7.05 1228 1.170803e+03 5.1058140

1694) SatDom_EVI_var..70>=7.534334e-07 1027 8.548348e+02 4.9237290

3388) SatDom_blue_band_var.0>=5.212728e-07 709 5.765680e+02 4.6589990 *

3389) SatDom_blue_band_var.0< 5.212728e-07 318 1.177952e+02 5.5139620 *

1695) SatDom_EVI_var..70< 7.534334e-07 201 1.079406e+02 6.0361690 *

53) METADom_Nitrogen_fert_repeat_3>=38.05 172 5.941642e+02 7.8733720

106) BOMDom_min_temperature_mean_.60>=14.775 88 7.715251e+01 6.5318180 *

107) BOMDom_min_temperature_mean_.60< 14.775 84 1.927109e+02 9.2788100 *

27) SatDom_EvapoTrans_mean..60< 1.76875 251 6.450897e+02 7.3486450

54) SatDom_Reflect_b5_var..70< 0.0002486262 178 1.732119e+02 6.5024160 *

55) SatDom_Reflect_b5_var..70>=0.0002486262 73 3.360219e+01 9.4120550 *

7) BOMDom_Sowing_date_numeric>=16579 5047 1.359763e+04 4.9653680

14) BOMDom_rainfall_csum_.40>=77.2 737 8.184043e+02 3.0366350

28) BOMDom_solar_exposure_mean_.40>=16.305 214 7.988143e+01 1.8937850 *

29) BOMDom_solar_exposure_mean_.40< 16.305 523 3.446482e+02 3.5042640

58) SatDom_Reflect_b4_min..70>=0.1008938 235 7.086616e+01 2.8642980 *

59) SatDom_Reflect_b4_min..70< 0.1008938 288 9.900239e+01 4.0264580 *

15) BOMDom_rainfall_csum_.40< 77.2 4310 9.568767e+03 5.2951760

30) METADom_Chem_rotation_minus_2_LVEMCPA>=0.0002155172 3330 5.669444e+03 4.9507030

60) METADom_Chem_rotation_minus_1_LVE< 0.01510193 443 6.346111e+02 3.5435440

120) METADom_Chem_rotation_minus_2_SPINREPORTED_NAKER< 0.0007490637 187 8.200160e+01 2.4721930 *

121) METADom_Chem_rotation_minus_2_SPINREPORTED_NAKER>=0.0007490637 256 1.811861e+02 4.3261330 *

61) METADom_Chem_rotation_minus_1_LVE>=0.01510193 2887 4.023050e+03 5.1666260

122) METADom_Fert_rotation_minus_1_UAN>=0.03611111 125 7.476832e+01 2.8752000 *

123) METADom_Fert_rotation_minus_1_UAN< 0.03611111 2762 3.262249e+03 5.2703290

246) SatDom_PsnNet_min..80< 0.0163625 2584 2.407378e+03 5.1549610

492) SatDom_FPAR_mean..60>=0.125875 2237 1.629927e+03 4.9723920

984) SatDom_GPP_var.0< 2.130043e-07 896 5.030118e+02 4.5416180

1968) SatDom_Latent_heat_mean..20>=448016.7 755 2.997124e+02 4.3574300 *

1969) SatDom_Latent_heat_mean..20< 448016.7 141 4.053556e+01 5.5278720 *

985) SatDom_GPP_var.0>=2.130043e-07 1341 8.495562e+02 5.2602160

1970) METADom_ESP_10cm.test_10cm>=1.2 934 4.391527e+02 5.0049570

3940) BOMDom_min_temperature_var_.50>=11.41939 236 7.883900e+01 4.3400000 *

3941) BOMDom_min_temperature_var_.50< 11.41939 698 2.206799e+02 5.2297850 *

1971) METADom_ESP_10cm.test_10cm< 1.2 407 2.098900e+02 5.8459950 *

493) SatDom_FPAR_mean..60< 0.125875 347 2.222038e+02 6.3319310 *

247) SatDom_PsnNet_min..80>=0.0163625 178 3.212062e+02 6.9451120

494) SatDom_Reflect_b5_mean..10>=0.3402462 46 1.794055e+01 5.3052170 *

495) SatDom_Reflect_b5_mean..10< 0.3402462 132 1.364504e+02 7.5165910 *

31) METADom_Chem_rotation_minus_2_LVEMCPA< 0.0002155172 980 2.161492e+03 6.4656840

62) SatDom_Pot_latent_heat_mean..20>=6872796 583 7.907674e+02 5.6586110

124) METADom_Fert_rotation_minus_1_SULPHATE< 0.009111044 168 1.262391e+02 4.4357140 *

125) METADom_Fert_rotation_minus_1_SULPHATE>=0.009111044 415 3.115816e+02 6.1536630

250) BOMDom_solar_exposure_csum_.80>=268.6 262 1.130370e+02 5.7231300 *

251) BOMDom_solar_exposure_csum_.80< 268.6 153 6.681887e+01 6.8909150 *

63) SatDom_Pot_latent_heat_mean..20< 6872796 397 4.333146e+02 7.6508820

126) METADom_ESP_60cm.test_60cm< 4.958599 159 1.416412e+02 6.8933960 *

127) METADom_ESP_60cm.test_60cm>=4.958599 238 1.394929e+02 8.1569330 *

**Wheat Regression tree: Predicted at End of Season**

**Leaf values indicate node-average yield, in t/Ha**

**Model Accuracy: Missing Site Prediction R^2^ = 0.76; p<0.0001**

**n= 52362**

**node), split, n, deviance, yval**

*** denotes terminal node**

1) root 52362 127045.70000 3.3751660

2) SatDom_MIR_min_150>=0.1412594 28022 35811.19000 2.5318580

4) SatDom_Reflect_b6_min_130>=0.2482875 10784 10053.19000 1.8697350

8) SatDom_Reflect_b7_mean_160>=0.29908 3903	1170.56400 1.2813630

16) SatDom_MIR_mean_160>=0.36115 841	150.20120 0.8136147 *

17) SatDom_MIR_mean_160< 0.36115 3062	785.82480 1.4098330

34) BOMDom_rainfall_csum_160< 276.825 2467	447.37000 1.3032100 *

35) BOMDom_rainfall_csum_160>=276.825 595	194.12380 1.8519160 *

9) SatDom_Reflect_b7_mean_160< 0.29908 6881	6765.08800 2.2034680

18) SatDom_Reflect_b1_var_140>=7.727995e-07 6661	4489.56500 2.1088350

36) SatDom_Latent_heat_180< 6.485603e+08 5936	2898.07900 1.9697780

72) BOMDom_rainfall_csum_130< 231.4 2933	944.17690 1.6829760

144) SatDom_LST_night_mean_80< 24.7045 1445	349.98580 1.4152530 *

145) SatDom_LST_night_mean_80>=24.7045 1488	390.04020 1.9429640

290) SatDom_NIR_var..30< 3.883281e-07 214	20.53150 1.1845330 *

291) SatDom_NIR_var..30>=3.883281e-07 1274	225.73500 2.0703610 *

73) BOMDom_rainfall_csum_130>=231.4 3003	1477.01800 2.2498930

146) SatDom_Reflect_b5_min..40>=0.23145 2789	1143.00600 2.1614090

292) SatDom_GPP_max_120< 0.02969375 1106	278.02460 1.7954520 *

293) SatDom_GPP_max_120>=0.02969375 1683	619.52190 2.4019010

586) SatDom_LST_night_mean_120>=25.9315 1315	351.09120 2.2515740 *

587) SatDom_LST_night_mean_120< 25.9315 368	132.52510 2.9390760 *

147) SatDom_Reflect_b5_min..40< 0.23145 214	27.58816 3.4030840 *

37) SatDom_Latent_heat_180>=6.485603e+08 725	536.89660 3.2473790

74) SatDom_red_band_var_170>=3.561313e-05 299	69.08327 2.4850840 *

75) SatDom_red_band_var_170< 3.561313e-05 426	172.11660 3.7824180 *

19) SatDom_Reflect_b1_var_140< 7.727995e-07 220	409.79190 5.0686820

38) SatDom_Pot_EvapoTrans_max..70< 60 126	39.35132 4.1563490 *

39) SatDom_Pot_EvapoTrans_max..70>=60 94	124.98570 6.2915960 *

5) SatDom_Reflect_b6_min_130< 0.2482875 17238 18072.54000 2.9460790

10) SatDom_red_band_190>=29.1758 10757	8050.20100 2.6621160

20) BOMDom_solar_exposure_max_60< 11.05 2260	1179.95200 2.0594470

40) METADom_Chem_rotation_minus_2_SIMAZINE>=0.04749257 922	216.21930 1.5736980 *

41) METADom_Chem_rotation_minus_2_SIMAZINE< 0.04749257 1338	596.27630 2.3941700

82) SatDom_FPAR_mean_90>=0.5700833 1060	313.92880 2.1935850 *

83) SatDom_FPAR_mean_90< 0.5700833 278	77.08192 3.1589930 *

21) BOMDom_solar_exposure_max_60>=11.05 8497	5831.06700 2.8224110

42) SatDom_LST_night_max_60< 33.84 8322	5179.29000 2.7841770

84) SatDom_NIR_min_110< 0.32885 4403	2382.63900 2.5588940

168) SatDom_Reflect_b4_var_120< 1.281602e-05 2661	1111.35100 2.3423680

336) BOMDom_min_temperature_min_60< 0.35 1189	366.06180 2.0030530 *

337) BOMDom_min_temperature_min_60>=0.35 1472	497.81810 2.6164470

674) SatDom_Reflect_b5_min..80>=0.4189 183	12.02136 1.7847540 *

675) SatDom_Reflect_b5_min..80< 0.4189 1289	341.24210 2.7345230

1350) SatDom_Reflect_b1_var_120>=1.098737e-05 616	93.96020 2.3999350 *

1351) SatDom_Reflect_b1_var_120< 1.098737e-05 673	115.20140 3.0407730 *

169) SatDom_Reflect_b4_var_120>=1.281602e-05 1742	955.95670 2.8896500

338) SatDom_red_band_max_110>=0.097225 304	108.59670 2.0965130 *

339) SatDom_red_band_max_110< 0.097225 1438	615.69580 3.0573230

678) METADom_Chem_rotation_minus_1_POWER>=-3.178881e-17 881 306.63910 2.7887740 *

679) METADom_Chem_rotation_minus_1_POWER< -3.178881e-17 557 145.02600 3.4820830 *

85) SatDom_NIR_min_110>=0.32885 3919	2322.12800 3.0372820

170) SatDom_LST_day_var_20>=0.7064931 3584	1792.49000 2.9369480

340) SatDom_Pot_latent_heat_min_30< 5363750 3012	1283.65000 2.8074700

680) BOMDom_solar_exposure_mean_60< 8.825 223	37.05788 1.8415250 *

681) BOMDom_solar_exposure_mean_60>=8.825 2789	1021.88500 2.8847040

1362) SatDom_NDVI_max_170< 0.2004 158	16.86859 1.8539870 *

1363) SatDom_NDVI_max_170>=0.2004 2631	827.08100 2.9466020

2726) SatDom_NIR_var_100>=3.116294e-05 1563	362.87630 2.7602180 *

2727) SatDom_NIR_var_100< 3.116294e-05 1068	330.44410 3.2193730 *

341) SatDom_Pot_latent_heat_min_30>=5363750 572	192.45490 3.6187410 *

171) SatDom_LST_day_var_20< 0.7064931 335	107.55040 4.1107160 *

43) SatDom_LST_night_max_60>=33.84 175	61.07643 4.6406290 *

11) SatDom_red_band_190< 29.1758 6481	7715.27200 3.4173940

22) SatDom_EvapoTrans_min..80< 3.95 2073	2049.60500 2.8793490

44) BOMDom_min_temperature_min_.50< 7.8 871	528.41670 2.2321010

88) SatDom_PsnNet_max_130< 0.032575 571	192.75930 1.8559890 *

89) SatDom_PsnNet_max_130>=0.032575 300	101.14470 2.9479670 *

45) BOMDom_min_temperature_min_.50>=7.8 1202	891.89370 3.3483610

90) SatDom_Reflect_b5_var_40< 0.000106932 623	342.27560 2.8533230

180) METADom_Fert_rotation_minus_2_TWENTY_FOUR_TO_SIXTEEN< -2.480655e-17 230 66.99684 2.2175650 *

181) METADom_Fert_rotation_minus_2_TWENTY_FOUR_TO_SIXTEEN>=-2.480655e-17 393 127.90980 3.2253940 *

91) SatDom_Reflect_b5_var_40>=0.000106932 579	232.66730 3.8810190 *

23) SatDom_EvapoTrans_min..80>=3.95 4408	4783.32400 3.6704260

46) SatDom_MIR_min.0< 0.1589062 2330	1910.86300 3.2853300

92) SatDom_Reflect_b3_mean_30< 0.02486812 204	59.86032 1.9263730 *

93) SatDom_Reflect_b3_mean_30>=0.02486812 2126	1438.11200 3.4157290

186) SatDom_Reflect_b1..60>=2.540712 1736	940.55480 3.2346660

372) SatDom_LAI_var_130< 0.084125 1217	468.23160 3.0050290

744) BOMDom_solar_exposure_min_130< 7.8 433	133.98180 2.5403230 *

745) BOMDom_solar_exposure_min_130>=7.8 784	189.09960 3.2616840 *

373) SatDom_LAI_var_130>=0.084125 519	257.66020 3.7731410 *

187) SatDom_Reflect_b1..60< 2.540712 390	187.30990 4.2216920 *

47) SatDom_MIR_min.0>=0.1589062 2078	2139.48500 4.1022230

94) SatDom_Reflect_b6_mean..10>=0.3138256 505	350.87850 3.1502570

188) SatDom_Reflect_b4_mean_120< 0.07023812 247	59.95304 2.4776520 *

189) SatDom_Reflect_b4_mean_120>=0.07023812 258	72.20488 3.7941860 *

95) SatDom_Reflect_b6_mean..10< 0.3138256 1573	1184.03000 4.4078450

190) SatDom_Reflect_b6_max_180>=0.270125 1384	706.66050 4.2393570

380) SatDom_LST_day_var..30>=3.889894 869	370.39310 3.9791250 *

381) SatDom_LST_day_var..30< 3.889894 515	178.11770 4.6784660 *

191) SatDom_Reflect_b6_max_180< 0.270125 189	150.37500 5.6416400 *

3) SatDom_MIR_min_150< 0.1412594 24340 48363.22000 4.3460440

6) SatDom_EVI_mean_160< 0.5025781 19422 25875.81000 4.0202770

12) SatDom_PsnNet_max_160< 0.032275 10170 11262.00000 3.7184990

24) METADom_Chem_rotation_minus_2_ROUNDUP_POWERMAX< 0.1111326 9279 8794.61300 3.6055530

48) METADom_Chem_rotation_minus_2_ESTER< 0.1067012 8988	7377.23100 3.5443110

96) BOMDom_rainfall_csum_160< 318.05 3557	3097.71500 3.2171720

192) METADom_Chem_rotation_minus_1_GLEAN>=0.00170151 1996	1427.64700 2.8919840

384) SatDom_PsnNet_var_180< 1.103906e-07 284	65.07937 1.9126760 *

385) SatDom_PsnNet_var_180>=1.103906e-07 1712	1045.01700 3.0544390

770) METADom_Chem_rotation_minus_2_GRAMMOXONE< 0.001582709 1242 510.43620 2.8014010

1540) METADom_Fert_rotation_minus_1_SOA< 0.007542432 375	102.74670 2.2629870 *

1541) METADom_Fert_rotation_minus_1_SOA>=0.007542432 867	251.96140 3.0342790 *

771) METADom_Chem_rotation_minus_2_GRAMMOXONE>=0.001582709 470 244.91310 3.7231060

1542) SatDom_Reflect_b2..30< 13.85418 290	80.85533 3.3090340 *

1543) SatDom_Reflect_b2..30>=13.85418 180	34.22779 4.3902220 *

193) METADom_Chem_rotation_minus_1_GLEAN< 0.00170151 1561	1189.10600 3.6329790

386) SatDom_Reflect_b4_var_130< 1.514175e-05 692	453.52720 3.1476590

772) SatDom_MIR_min_90>=0.07474063 442	175.71350 2.7590270 *

773) SatDom_MIR_min_90< 0.07474063 250	93.02944 3.8347600 *

387) SatDom_Reflect_b4_var_130>=1.514175e-05 869	442.79650 4.0194480

774) SatDom_NIR_var_150< 6.335487e-06 150	39.21150 3.0308000 *

775) SatDom_NIR_var_150>=6.335487e-06 719	226.38440 4.2257020 *

97) BOMDom_rainfall_csum_160>=318.05 5431	3649.52700 3.7585690

194) SatDom_LST_night_mean_120< 27.929 3672	1968.19700 3.5708580

388) BOMDom_max_temperature_mean_170>=31.635 235	117.13890 2.4865960 *

389) BOMDom_max_temperature_mean_170< 31.635 3437	1555.89600 3.6449930

778) SatDom_Reflect_b7_mean.0>=0.179955 2334	891.38710 3.4785130 *

779) SatDom_Reflect_b7_mean.0< 0.179955 1103	462.93910 3.9972710 *

195) SatDom_LST_night_mean_120>=27.929 1759	1281.84700 4.1504260

390) SatDom_LST_day_var..30< 16.64659 1535	839.14490 3.9814200

780) BOMDom_solar_exposure_max_140< 22.95 302	94.63691 3.1653640 *

781) BOMDom_solar_exposure_max_140>=22.95 1233	494.13230 4.1812980

1562) METADom_Fert_rotation_minus_2_MES>=-1.07336e-17 722	208.02220 3.8749030 *

1563) METADom_Fert_rotation_minus_2_MES< -1.07336e-17 511	122.56350 4.6142070 *

391) SatDom_LST_day_var..30>=16.64659 224	98.40654 5.3085710 *

49) METADom_Chem_rotation_minus_2_ESTER>=0.1067012 291	342.47380 5.4971130

98) SatDom_NDVI_min..10< 0.3004594 246	116.04210 5.1299590 *

99) SatDom_NDVI_min..10>=0.3004594 45	11.98850 7.5042220 *

25) METADom_Chem_rotation_minus_2_ROUNDUP_POWERMAX>=0.1111326 891 1116.30900 4.8947250

50) SatDom_Reflect_b7_mean_30>=0.1181525 599	505.96380 4.4308350

100) SatDom_MIR_var..20< 3.572749e-06 73	36.89666 2.7613700 *

101) SatDom_MIR_var..20>=3.572749e-06 526	237.37110 4.6625290 *

51) SatDom_Reflect_b7_mean_30< 0.1181525 292	217.01980 5.8463360 *

13) SatDom_PsnNet_max_160>=0.032275 9252 12669.54000 4.3519980

26) SatDom_MIR_max_10>=0.17215 5990	6248.05800 4.0557760

52) SatDom_LAI_mean_10>=0.24375 3624	3159.64500 3.7798400

104) SatDom_NDVI_max_140< 0.7276594 1591	1021.38500 3.3887120

208) SatDom_MIR_var_190< 0.0001489201 1181	537.77950 3.1248350

416) SatDom_NDVI_min..70< 0.28735 789	289.56520 2.8902280 *

417) SatDom_NDVI_min..70>=0.28735 392	117.38000 3.5970410 *

209) SatDom_MIR_var_190>=0.0001489201 410	164.49750 4.1488050 *

105) SatDom_NDVI_max_140>=0.7276594 2033	1704.38900 4.0859320

210) SatDom_LST_day_mean_100< 33.3289 1100	806.54300 3.7213000

420) SatDom_LAI_min_100>=1.3375 611	359.80320 3.3846970 *

421) SatDom_LAI_min_100< 1.3375 489	291.01410 4.1418810 *

211) SatDom_LST_day_mean_100>=33.3289 933	579.16390 4.5158310 *

53) SatDom_LAI_mean_10< 0.24375 2366	2389.83000 4.4784280

106) SatDom_Latent_heat_70< 1.545598e+08 181	43.36204 2.4416020 *

107) SatDom_Latent_heat_70>=1.545598e+08 2185	1533.35800 4.6471530

214) SatDom_LST_night_mean_80< 24.4185 816	382.73710 4.1601350 *

215) SatDom_LST_night_mean_80>=24.4185 1369	841.71270 4.9374430

430) SatDom_Reflect_b3_min_160>=0.01685 1305	588.02550 4.8486670 *

431) SatDom_Reflect_b3_min_160< 0.01685 64	33.68235 6.7476560 *

27) SatDom_MIR_max_10< 0.17215 3262	4930.70400 4.8959500

54) SatDom_Reflect_b7_max_70>=0.0699 2877	3658.97800 4.7025510

108) SatDom_Reflect_b2_var_190>=0.0002498159 1185	1318.21300 4.1245820

216) SatDom_NIR_var_100< 3.661161e-06 194	122.14510 2.5863400 *

217) SatDom_NIR_var_100>=3.661161e-06 991	647.16450 4.4257110

434) SatDom_Reflect_b7_var_30>=1.015825e-05 823	432.12270 4.2533290 *

435) SatDom_Reflect_b7_var_30< 1.015825e-05 168	70.78089 5.2701790 *

109) SatDom_Reflect_b2_var_190< 0.0002498159 1692	1667.68500 5.1073350

218) SatDom_Reflect_b7_var_120>=9.721528e-07 1581	1288.36900 4.9896270

436) SatDom_EvapoTrans_min_70< 8.95 1009	746.51740 4.7093860

872) SatDom_GPP_var_10< 3.309696e-07 258	93.22838 4.0283330 *

873) SatDom_GPP_var_10>=3.309696e-07 751	492.50910 4.9433560 *

437) SatDom_EvapoTrans_min_70>=8.95 572	322.82810 5.4839690 *

219) SatDom_Reflect_b7_var_120< 9.721528e-07 111	45.41343 6.7838740 *

55) SatDom_Reflect_b7_max_70< 0.0699 385	359.98460 6.3411690

110) SatDom_Reflect_b2_max_50< 0.3646 307	120.92450 5.9877850 *

111) SatDom_Reflect_b2_max_50>=0.3646 78	49.82667 7.7320510 *

7) SatDom_EVI_mean_160>=0.5025781 4918 12286.50000 5.6325500

14) METADom_Nitrogen_fert_repeat_2< 23.5 3425	6719.68200 5.1698510

28) METADom_Fert_rotation_minus_1_AGFLOW>=-1.579466e-16 3305 4787.58400 5.0422300

56) SatDom_PsnNet_max_90< 0.0223 2646	3239.66300 4.7988440

112) SatDom_LST_day_mean_30>=35.38345 642	776.61820 4.1240190

224) SatDom_Pot_EvapoTrans_mean_170< 43.05487 78	31.14800 2.5988460 *

225) SatDom_Pot_EvapoTrans_mean_170>=43.05487 564	538.93770 4.3349470 *

113) SatDom_LST_day_mean_30< 35.38345 2004	2077.02500 5.0150300

226) SatDom_Pot_latent_heat..30< 9.863505e+08 1629	1446.48500 4.8113260

452) SatDom_Pot_latent_heat_var..10< 4.378695e+11 770	534.99550 4.5075450 *

453) SatDom_Pot_latent_heat_var..10>=4.378695e+11 859	776.73610 5.0836320

906) SatDom_Reflect_b4_mean_160>=0.08259812 96	125.55240 3.9909370 *

907) SatDom_Reflect_b4_mean_160< 0.08259812 763	522.13980 5.2211140 *

227) SatDom_Pot_latent_heat..30>=9.863505e+08 375	269.30850 5.8999200 *

57) SatDom_PsnNet_max_90>=0.0223 659	761.83810 6.0194690

114) SatDom_Reflect_b1_var_130< 1.439127e-05 264	136.84850 5.0278410 *

115) SatDom_Reflect_b1_var_130>=1.439127e-05 395	191.88800 6.6822280 *

29) METADom_Fert_rotation_minus_1_AGFLOW< -1.579466e-16 120	395.72720 8.6847500

58) SatDom_Reflect_b6_30>=37.40318 33	14.76199 6.1506060 *

59) SatDom_Reflect_b6_30< 37.40318 87	88.65869 9.6459770 *

15) METADom_Nitrogen_fert_repeat_2>=23.5 1493	3151.43400 6.6939990

30) METADom_Chem_rotation_minus_2_CADENCE>=0.005310412 515	543.92360 5.5259610

60) SatDom_Reflect_b3_var..20< 2.868225e-05 418	288.01180 5.2193780 *

61) SatDom_Reflect_b3_var..20>=2.868225e-05 97	47.31459 6.8471130 *

31) METADom_Chem_rotation_minus_2_CADENCE< 0.005310412 978	1534.90000 7.3090700

62) METADom_Fert_rotation_minus_1_THIRTY_TWO_TO_NINE>=0.001565996 264 109.42830 6.0908710 *

63) METADom_Fert_rotation_minus_1_THIRTY_TWO_TO_NINE< 0.001565996 714 888.83520 7.7594960

126) SatDom_LST_day_max..70>=62.235 302	216.22500 7.0305630 *

127) SatDom_LST_day_max..70< 62.235 412	394.52170 8.2938110 *

**Canola Regression tree: Predicted at Time of Sowing**

**Leaf values indicate node-average yield, in t/Ha**

**Model Accuracy: Missing Site Prediction R^2^ = 0.21; p<0.0001**

**n= 14321**

**node), split, n, deviance, yval**

*** denotes terminal node**

1) root 14321 7941.3380000 2.0151750

2) METADom_Nitrogen_fert_repeat_2< 2.37367 9248 4529.3200000 1.8442430

4) SatDom_GPP_mean..50< 0.009111875 4066 1638.1140000 1.5848110

8) BOMDom_min_temperature_mean_.50< 18.005 3360 1169.7250000 1.4835510

16) SatDom_Reflect_b5_min.0>=0.2127 3058 950.5352000 1.4172300

32) SatDom_EvapoTrans_min..70< 0.95 306	40.9912200 0.8553268

64) SatDom_NDVI_var..40< 5.10032e-05 261	16.0463800 0.7411111 *

65) SatDom_NDVI_var..40>=5.10032e-05 45	1.7921780 1.5177780 *

33) SatDom_EvapoTrans_min..70>=0.95 2752 802.1862000 1.4797090

66) METADom_Fert_rotation_minus_2_SUPREME< -2.020953e-17 383 65.6054500 1.0118800

132) SatDom_FPAR_max..40< 0.19625 245	23.4084300 0.7825306

264) METADom_Chem_rotation_minus_2_FOR>=-7.784572e-18 211	11.7003100 0.6984834 *

265) METADom_Chem_rotation_minus_2_FOR< -7.784572e-18 34	0.9678235 1.3041180 *

133) SatDom_FPAR_max..40>=0.19625 138	6.4301780 1.4190580 *

67) METADom_Fert_rotation_minus_2_SUPREME>=-2.020953e-17 2369 639.2035000 1.5553440

134) SatDom_Reflect_b5_mean..10>=0.2395594 2269 548.7863000 1.5167560

268) SatDom_Latent_heat_mean..50>=808500 1399 265.4293000 1.3566690

536) BOMDom_max_temperature_mean_.50>=30.63 184	14.9503500 0.8844565

1072) SatDom_red_band_mean..80< 0.1870728 89	2.7762880 0.6688764 *

1073) SatDom_red_band_mean..80>=0.1870728 95	4.1627830 1.0864210 *

537) BOMDom_max_temperature_mean_.50< 30.63 1215 203.2363000 1.4281810

1074) SatDom_Reflect_b7_var..30< 0.0001520474 810 104.2687000 1.3013210

2148) SatDom_Pot_latent_heat_mean..50< 1.398506e+07 541	46.0315500 1.1697040

4296) SatDom_NDVI_var..80>=7.160867e-06 118	7.3777760 0.9132203 *

4297) SatDom_NDVI_var..80< 7.160867e-06 423	28.7258400 1.2412530

8594) METADom_Chem_rotation_minus_2_ESTERCIDE>=-5.937091e-17 380 17.4413000 1.1935530 *

8595) METADom_Chem_rotation_minus_2_ESTERCIDE< -5.937091e-17 43 2.7790650 1.6627910 *

2149) SatDom_Pot_latent_heat_mean..50>=1.398506e+07 269	30.0174400 1.5660220

4298) SatDom_Emis32_var..30>=9.546667e-07 162	13.9477700 1.4067280 *

4299) SatDom_Emis32_var..30< 9.546667e-07 107	5.7353590 1.8071960 *

1075) SatDom_Reflect_b7_var..30>=0.0001520474 405	59.8604400 1.6819010

2150) METADom_Fert_rotation_minus_1_AMMONIUMNITRATE>=-5.303917e-17 302 27.2359500 1.5649340

4300) SatDom_Reflect_b4_min..20< 0.08055 79	11.1115700 1.2850630 *

4301) SatDom_Reflect_b4_min..20>=0.08055 223	7.7443870 1.6640810 *

2151) METADom_Fert_rotation_minus_1_AMMONIUMNITRATE< -5.303917e-17 103 16.3781700 2.0248540 *

269) SatDom_Latent_heat_mean..50< 808500 870 189.8496000 1.7741840

538) SatDom_GPP..50< 0.2759063 81	9.1521580 1.0817280 *

539) SatDom_GPP..50>=0.2759063 789 137.8711000 1.8452720

1078) METADom_Fert_rotation_minus_1_AP< -1.593777e-17 657	98.5259800 1.7614160

2156) SatDom_LST_night_var..30>=2.654634 549	63.1868200 1.6907100

4312) SatDom_Reflect_b6_var.0< 2.134355e-05 27	1.1340070 1.0081480 *

4313) SatDom_Reflect_b6_var.0>=2.134355e-05 522	48.8231100 1.7260150 *

2157) SatDom_LST_night_var..30< 2.654634 108	18.6430200 2.1208330 *

1079) METADom_Fert_rotation_minus_1_AP>=-1.593777e-17 132	11.7299700 2.2626520 *

135) SatDom_Reflect_b5_mean..10< 0.2395594 100	10.3788200 2.4309000 *

17) SatDom_Reflect_b5_min.0< 0.2127 302	69.5439500 2.1550990

34) METADom_Fert_rotation_minus_1_DAP< 0.1109129 182	22.8097700 1.8715380

68) SatDom_Pot_EvapoTrans_min..70>=60.925 26	1.7235120 1.3073080 *

69) SatDom_Pot_EvapoTrans_min..70< 60.925 156	11.4294500 1.9655770 *

35) METADom_Fert_rotation_minus_1_DAP>=0.1109129 120	9.9051970 2.5851670 *

9) BOMDom_min_temperature_mean_.50>=18.005 706 269.9725000 2.0667280

18) SatDom_EvapoTrans_min..60< 1.075 144	16.6063300 1.2368060

36) METADom_Fert_rotation_minus_2_DAP< 0.06941086 37	1.0722270 0.8178378 *

37) METADom_Fert_rotation_minus_2_DAP>=0.06941086 107	6.7934970 1.3816820 *

19) SatDom_EvapoTrans_min..60>=1.075 562 128.7697000 2.2793770

38) SatDom_LST_night_max..50< 41.45 190	34.6989600 1.9360000

76) METADom_Fert_rotation_minus_1_MAP>=0.2895969 149	17.2671900 1.8002680 *

77) METADom_Fert_rotation_minus_1_MAP< 0.2895969 41	4.7108780 2.4292680 *

39) SatDom_LST_night_max..50>=41.45 372	60.2260800 2.4547580

78) SatDom_Emis32_mean..50< 0.9867 280	33.9223000 2.3500360 *

79) SatDom_Emis32_mean..50>=0.9867 92	13.8874900 2.7734780 *

5) SatDom_GPP_mean..50>=0.009111875 5182 2402.8160000 2.0478040

10) SatDom_PsnNet_var..60< 6.823516e-07 2196 1035.0130000 1.8376320

20) SatDom_Pot_latent_heat_mean..20>=1432419 1922 785.7791000 1.7334340

40) SatDom_PsnNet_mean..30>=0.006528125 1757 617.4272000 1.6502450

80) METADom_Chem_rotation_minus_2_AVADEX< 0.03841744 1370 370.4852000 1.5204740

160) SatDom_FPAR_var..40>=0.001127639 292	32.4270700 0.9876370

320) METADom_Fert_rotation_minus_2_PLAIN< 0.001701985 154	2.9357850 0.7429221 *

321) METADom_Fert_rotation_minus_2_PLAIN>=0.001701985 138	9.9773280 1.2607250 *

161) SatDom_FPAR_var..40< 0.001127639 1078 232.6985000 1.6648050

322) SatDom_LAI_mean..30< 0.33375 706	93.0386500 1.4850420

644) METADom_Chem_rotation_minus_2_TOPIC< -4.466913e-18 99	9.7501780 0.9511111 *

645) METADom_Chem_rotation_minus_2_TOPIC>=-4.466913e-18 607	50.4621600 1.5721250

1290) SatDom_LST_night_var..20>=4.361125 197	8.6035800 1.3525890 *

1291) SatDom_LST_night_var..20< 4.361125 410	27.8018600 1.6776100 *

323) SatDom_LAI_mean..30>=0.33375 372	73.5479500 2.0059680

646) SatDom_FPAR..80>=1.94125 277	34.3525800 1.8366430

1292) METADom_Fert_rotation_minus_2_TWENTY_SEVEN_TO_TWELVE< 0.01666667 96 6.6163660 1.5465630 *

1293) METADom_Fert_rotation_minus_2_TWENTY_SEVEN_TO_TWELVE>=0.01666667 181 15.3736600 1.9904970 *

647) SatDom_FPAR..80< 1.94125 95	8.0966910 2.4996840 *

81) METADom_Chem_rotation_minus_2_AVADEX>=0.03841744 387 142.1973000 2.1096380

162) SatDom_blue_band_var.0< 2.068216e-07 68	6.0959220 1.1583820 *

163) SatDom_blue_band_var.0>=2.068216e-07 319	61.4524400 2.3124140

326) SatDom_MIR_min..10< 0.2271406 216	30.6483400 2.1455090 *

327) SatDom_MIR_min..10>=0.2271406 103	12.1684900 2.6624270 *

41) SatDom_PsnNet_mean..30< 0.006528125 165	26.7155100 2.6192730

82) SatDom_Latent_heat_var..70>=7.947795e+09 115	14.1950000 2.4700000 *

83) SatDom_Latent_heat_var..70< 7.947795e+09 50	4.0643620 2.9626000 *

21) SatDom_Pot_latent_heat_mean..20< 1432419 274	81.9882200 2.5685400

42) SatDom_red_band_min.0>=0.102325 101	12.6809000 2.0445540 *

43) SatDom_red_band_min.0< 0.102325 173	25.3870700 2.8744510 *

11) SatDom_PsnNet_var..60>=6.823516e-07 2986 1199.4620000 2.2023710

22) METADom_Chem_rotation_minus_2_CADENCE< 0.005275474 1796 532.1315000 2.0095160

44) SatDom_FPAR_var..20< 0.0003362847 791 159.9405000 1.7720860

88) METADom_Chem_rotation_minus_2_BRODAL< 0.05726167 711	87.8259500 1.6764140

176) SatDom_MIR_var..20>=1.75091e-05 520	40.1918300 1.5558460 *

177) SatDom_MIR_var..20< 1.75091e-05 191	19.4957500 2.0046600 *

89) METADom_Chem_rotation_minus_2_BRODAL>=0.05726167 80	7.7672490 2.6223750 *

45) SatDom_FPAR_var..20>=0.0003362847 1005 292.5042000 2.1963880

90) SatDom_Reflect_b7_max..40< 0.187425 76	11.4792900 1.3096050 *

91) SatDom_Reflect_b7_max..40>=0.187425 929 216.3704000 2.2689340

182) SatDom_LST_day_var..30< 7.240234 595 100.0143000 2.0999160

364) BOMDom_solar_exposure_min_.70>=7.45 280	35.0827600 1.8562500

728) SatDom_Reflect_b5_max..40>=0.3351125 165	16.9576300 1.7042420 *

729) SatDom_Reflect_b5_max..40< 0.3351125 115	8.8424260 2.0743480 *

365) BOMDom_solar_exposure_min_.70< 7.45 315	33.5297600 2.3165080

730) BOMDom_rainfall_max_.30>=3.8 98	6.1459030 2.0762240 *

731) BOMDom_rainfall_max_.30< 3.8 217	19.1704200 2.4250230 *

183) SatDom_LST_day_var..30>=7.240234 334	69.0787000 2.5700300

366) SatDom_FPAR_var..20>=0.001281597 186	24.0195500 2.3105380 *

367) SatDom_FPAR_var..20< 0.001281597 148	16.7943000 2.8961490 *

23) METADom_Chem_rotation_minus_2_CADENCE>=0.005275474 1190 499.7158000 2.4934370

46) METADom_pH_CaCl2_60cm.test_60cm>=7.042779 416 115.1026000 1.9660340

92) METADom_Chem_rotation_minus_2_SIMAZINE>=0.06443296 71	15.9907000 1.2539440 *

93) METADom_Chem_rotation_minus_2_SIMAZINE< 0.06443296 345	55.7006000 2.1125800

186) METADom_ESP_60cm.test_60cm>=5.840898 220	21.5730900 1.9763640 *

187) METADom_ESP_60cm.test_60cm< 5.840898 125	22.8610300 2.3523200 *

47) METADom_pH_CaCl2_60cm.test_60cm< 7.042779 774 206.7096000 2.7768990

94) SatDom_Latent_heat_min..50< 966875 250	31.4640500 2.3850000

188) BOMDom_rainfall_max_.10>=8.5 152	13.0204100 2.2269740 *

189) BOMDom_rainfall_max_.10< 8.5 98	8.7604990 2.6301020 *

95) SatDom_Latent_heat_min..50>=966875 524 118.5304000 2.9638740

190) SatDom_EvapoTrans_min..30< 9218.803 476	76.0347800 2.8915340

380) SatDom_MIR_var..10>=1.563077e-05 217	40.4519800 2.7189860 *

381) SatDom_MIR_var..10< 1.563077e-05 259	23.7091600 3.0361000 *

191) SatDom_EvapoTrans_min..30>=9218.803 48	15.3025200 3.6812500 *

3) METADom_Nitrogen_fert_repeat_2>=2.37367 5073 2649.2330000 2.3267810

6) METADom_Chem_rotation_minus_2_AGRITONE< 0.002960825 3158 1651.6420000 2.1360960

12) SatDom_EvapoTrans..50>=86193.57 1023 342.2463000 1.6764810

24) SatDom_Reflect_b6_min..10>=0.3428187 280	49.3564600 1.0938930

48) SatDom_EVI_var..50>=7.345966e-07 185	11.4131900 0.8545405 *

49) SatDom_EVI_var..50< 7.345966e-07 95	6.7054000 1.5600000 *

25) SatDom_Reflect_b6_min..10< 0.3428187 743 162.0416000 1.8960300

50) METADom_Fert_rotation_minus_1_Z>=-2.276825e-17 609	86.8809500 1.7715440

100) BOMDom_min_temperature_var_.30>=6.268222 464	46.9399900 1.6726290

200) SatDom_EvapoTrans..20< 686239.7 327	25.8587800 1.5629360 *

201) SatDom_EvapoTrans..20>=686239.7 137	7.7549840 1.9344530 *

101) BOMDom_min_temperature_var_.30< 6.268222 145	20.8738600 2.0880690 *

51) METADom_Fert_rotation_minus_1_Z< -2.276825e-17 134	22.8315700 2.4617910 *

13) SatDom_EvapoTrans..50< 86193.57 2135 989.7436000 2.3563230

26) METADom_Chem_rotation_minus_2_SPINREPORTED_NAKER< -3.009745e-17 234 38.0348200 1.5253420

52) BOMDom_solar_exposure_var_.70>=61.68989 32	1.7894000 0.7525000 *

53) BOMDom_solar_exposure_var_.70< 61.68989 202	14.1045000 1.6477720 *

27) METADom_Chem_rotation_minus_2_SPINREPORTED_NAKER>=-3.009745e-17 1901 770.2349000 2.4586110

54) SatDom_Reflect_b3_var..40< 1.86612e-07 135	24.1199900 1.4877040

108) SatDom_EVI_max..20< 0.1564281 78	2.3334990 1.1832050 *

109) SatDom_EVI_max..20>=0.1564281 57	4.6578040 1.9043860 *

55) SatDom_Reflect_b3_var..40>=1.86612e-07 1766 609.1274000 2.5328310

110) SatDom_Reflect_b1_var.0< 0.000140493 1510 428.9337000 2.4305960

220) SatDom_blue_band_var..40< 1.375977e-07 213	32.2526900 1.8309860 *

221) SatDom_blue_band_var..40>=1.375977e-07 1297 307.5242000 2.5290670

442) BOMDom_min_temperature_var_.30>=5.937778 999 214.3512000 2.4076780

884) SatDom_LAI_var..40< 0.001753472 549	98.2622600 2.2083240

1768) METADom_Fert_rotation_minus_2_THIRTY_TWO_TO_TEN>=0.002174436 28 8.0850430 1.1564290 *

1769) METADom_Fert_rotation_minus_2_THIRTY_TWO_TO_TEN< 0.002174436 521 57.5306100 2.2648560

3538) SatDom_LST_day_min..40>=49.69 112	8.0310990 1.9399110 *

3539) SatDom_LST_day_min..40< 49.69 409	34.4350700 2.3538390 *

885) SatDom_LAI_var..40>=0.001753472 450	67.6524400 2.6508890

1770) SatDom_EvapoTrans_min.0< 8.125 353	41.5679000 2.5699150 *

1771) SatDom_EvapoTrans_min.0>=8.125 97	15.3469900 2.9455670 *

443) BOMDom_min_temperature_var_.30< 5.937778 298	29.1035500 2.9360070 *

111) SatDom_Reflect_b1_var.0>=0.000140493 256	71.3186100 3.1358590

222) SatDom_blue_band..20>=4.573588 142	25.0118500 2.8386620

444) SatDom_NIR..30>=16.89849 27	0.9301407 2.2114810 *

445) SatDom_NIR..30< 16.89849 115	10.9675800 2.9859130 *

223) SatDom_blue_band..20< 4.573588 114	18.1415200 3.5060530 *

7) METADom_Chem_rotation_minus_2_AGRITONE>=0.002960825 1915 693.4022000 2.6412380

14) SatDom_NIR_var..40< 0.0001116116 1711 486.5761000 2.5473060

28) BOMDom_min_temperature_var_0>=3.662944 1220 275.1293000 2.4146150

56) SatDom_Reflect_b5_var..70< 0.0003999032 938 182.5817000 2.3011190

112) METADom_Nitrogen_fert_repeat_2>=48.3 86	17.1474500 1.6731400 *

113) METADom_Nitrogen_fert_repeat_2< 48.3 852 128.0961000 2.3645070

226) METADom_Chem_rotation_minus_1_CT>=0.001656319 444	42.0474600 2.2080180

452) SatDom_GPP_mean..60< 0.0093925 116	6.7233540 1.8761210 *

453) SatDom_GPP_mean..60>=0.0093925 328	18.0269500 2.3253960 *

227) METADom_Chem_rotation_minus_1_CT< 0.001656319 408	63.3431800 2.5348040

454) METADom_Chem_rotation_minus_2_AVADEX>=0.04340492 214	24.8788900 2.3574300 *

455) METADom_Chem_rotation_minus_2_AVADEX< 0.04340492 194	24.3046600 2.7304640 *

57) SatDom_Reflect_b5_var..70>=0.0003999032 282	40.2755200 2.7921280

114) METADom_ATRAZINE>=495 72	4.7749110 2.4161110 *

115) METADom_ATRAZINE< 495 210	21.8303700 2.9210480 *

29) BOMDom_min_temperature_var_0< 3.662944 491 136.5935000 2.8770060

58) BOMDom_rainfall_max_.20< 2.9 232	37.1458000 2.5276290

116) BOMDom_min_temperature_min_.30>=8.85 64	4.4839610 2.1342190 *

117) BOMDom_min_temperature_min_.30< 8.85 168	18.9829500 2.6775000 *

59) BOMDom_rainfall_max_.20>=2.9 259	45.7621000 3.1899610

118) SatDom_PsnNet_min..80< 0.0097625 233	26.9616000 3.1046780 *

119) SatDom_PsnNet_min..80>=0.0097625 26	1.9190350 3.9542310 *

15) SatDom_NIR_var..40>=0.0001116116 204	65.1113200 3.4290690

30) SatDom_red_band_var..50>=1.544983e-05 92	21.4240700 3.0005430 *

31) SatDom_red_band_var..50< 1.544983e-05 112	12.9154700 3.7810710 *

**Canola Regression tree: Predicted at End of Season**

**Leaf values indicate node-average yield, in t/Ha**

**Model Accuracy: Missing Site Prediction R^2^ = 0.19; p<0.0001**

**n= 14321**

**node), split, n, deviance, yval**

*** denotes terminal node**

1) root 14321 7941.3380000 2.0151750

2) SatDom_Reflect_b7_min_160>=0.101525 6346 2782.6220000 1.6122390

4) BOMDom_rainfall_csum_120< 238.45 2365 526.4854000 1.1680850

8) SatDom_NDVI_min..30< 0.2051 952 147.1319000 0.8906933

16) SatDom_PsnNet_180< 8.590025 468	22.9458900 0.6551496

32) SatDom_LST_day_min_150>=37.89 412	13.2266400 0.6037379 *

33) SatDom_LST_day_min_150< 37.89 56	0.6184554 1.0333930 *

17) SatDom_PsnNet_180>=8.590025 484	73.1143400 1.1184500

34) SatDom_Pot_EvapoTrans_var_120>=2.637609 260	15.4884800 0.8462692

68) SatDom_FPAR_mean_80< 0.6495 182	4.0871870 0.7275275 *

69) SatDom_FPAR_mean_80>=0.6495 78	2.8475330 1.1233330 *

35) SatDom_Pot_EvapoTrans_var_120< 2.637609 224	16.0073100 1.4343750 *

9) SatDom_NDVI_min..30>=0.2051 1413 256.7475000 1.3549750

18) METADom_Chemicals_time_Chemicals_PYRASULFOTOLE< 63.80805 1201 150.9780000 1.2688260

36) METADom_Chem_rotation_minus_1_ECLIPSE>=-3.590878e-17 985 95.4792100 1.1825280

72) SatDom_Reflect_b6_mean_140>=0.2153238 491	35.9568700 1.0156420

144) BOMDom_max_temperature_mean_20< 20.695 443	22.4228800 0.9663431 *

145) BOMDom_max_temperature_mean_20>=20.695 48	2.5208810 1.4706250 *

73) SatDom_Reflect_b6_mean_140< 0.2153238 494	32.2556400 1.3484010

146) SatDom_NDVI_var_120>=4.851037e-05 224	7.9822210 1.2008930 *

147) SatDom_NDVI_var_120< 4.851037e-05 270	15.3559400 1.4707780 *

37) METADom_Chem_rotation_minus_1_ECLIPSE< -3.590878e-17 216 14.7113000 1.6623610 *

19) METADom_Chemicals_time_Chemicals_PYRASULFOTOLE>=63.80805 212 46.3604700 1.8430190

38) METADom_Chem_rotation_minus_1_ESTERCIDE< -1.483189e-17 103 20.6900700 1.5779610 *

39) METADom_Chem_rotation_minus_1_ESTERCIDE>=-1.483189e-17 109 11.5960800 2.0934860 *

5) BOMDom_rainfall_csum_120>=238.45 3981 1512.4200000 1.8760990

10) SatDom_NDVI_mean_140< 0.6541383 2621 840.8975000 1.7005040

20) SatDom_Latent_heat_mean..80< 1458125 1788 438.8148000 1.5473320

40) BOMDom_rainfall_mean_90< 3.695 1649 317.5459000 1.4815950

80) SatDom_MIR_var_50< 0.0002181144 1522 242.7728000 1.4314060

160) SatDom_Reflect_b1_var..30< 4.898313e-05 731	98.3253600 1.2624080

320) SatDom_Latent_heat_var..70>=6.112759e+10 110	4.3789670 0.8014545 *

321) SatDom_Latent_heat_var..70< 6.112759e+10 621	66.4337700 1.3440580

642) SatDom_Pot_EvapoTrans_min..30< 43.49375 495	39.2560800 1.2628480

1284) SatDom_Emis31_mean_80< 0.98325 234	16.5273500 1.1008120 *

1285) SatDom_Emis31_mean_80>=0.98325 261	11.0765800 1.4081230 *

643) SatDom_Pot_EvapoTrans_min..30>=43.49375 126	11.0882900 1.6630950 *

161) SatDom_Reflect_b1_var..30>=4.898313e-05 791 104.2757000 1.5875850

322) BOMDom_max_temperature_csum_150>=5680.7 131	9.3261600 1.1718320 *

323) BOMDom_max_temperature_csum_150< 5680.7 660	67.8116900 1.6701060

646) SatDom_Reflect_b2_max..10< 0.2565125 524	37.9986500 1.5917750 *

647) SatDom_Reflect_b2_max..10>=0.2565125 136	14.2101000 1.9719120 *

81) SatDom_MIR_var_50>=0.0002181144 127	24.9941000 2.0830710

162) BOMDom_max_temperature_mean_.70< 36.68 85	8.1858750 1.8722350 *

163) BOMDom_max_temperature_mean_.70>=36.68 42	5.3830980 2.5097620 *

41) BOMDom_rainfall_mean_90>=3.695 139	29.6052100 2.3271940

82) SatDom_blue_band_min_120< 0.0323 100	8.3904510 2.1043000 *

83) SatDom_blue_band_min_120>=0.0323 39	3.5076360 2.8987180 *

21) SatDom_Latent_heat_mean..80>=1458125 833 270.0916000 2.0292800

42) SatDom_Reflect_b5_var_80< 0.0007440978 744 157.3116000 1.9135220

84) BOMDom_rainfall_mean_110< 1.46 439	53.5341900 1.7078590

168) BOMDom_max_temperature_min_190< 21.15 222	15.7238500 1.4987390 *

169) BOMDom_max_temperature_min_190>=21.15 217	18.1700000 1.9217970 *

85) BOMDom_rainfall_mean_110>=1.46 305	58.4825400 2.2095410

170) SatDom_Reflect_b2_var..20< 4.692235e-05 163	27.3474600 1.9852760 *

171) SatDom_Reflect_b2_var..20>=4.692235e-05 142	13.5266000 2.4669720 *

43) SatDom_Reflect_b5_var_80>=0.0007440978 89	19.4692800 2.9969660

86) METADom_SIMAZINE>=24.91009 51	3.3701690 2.7207840 *

87) METADom_SIMAZINE< 24.91009 38	6.9880870 3.3676320 *

11) SatDom_NDVI_mean_140>=0.6541383 1360 434.9601000 2.2145070

22) SatDom_MixClouds_140< 5 1200 287.7606000 2.1067750

44) BOMDom_max_temperature_min_40< 13.15 666 100.7379000 1.8850000

88) METADom_Chem_rotation_minus_1_CT>=0.004405336 169	15.3988600 1.5214200 *

89) METADom_Chem_rotation_minus_1_CT< 0.004405336 497	55.4022700 2.0086320

178) SatDom_Latent_heat_min_130>=4654893 250	18.9106500 1.8648800 *

179) SatDom_Latent_heat_min_130< 4654893 247	26.0965900 2.1541300 *

45) BOMDom_max_temperature_min_40>=13.15 534 113.4123000 2.3833710

90) SatDom_LAI_var_80< 0.04458333 299	39.5528700 2.1251170

180) SatDom_EVI_var_190>=2.701217e-05 78	5.6845850 1.6876920 *

181) SatDom_EVI_var_190< 2.701217e-05 221	13.6762500 2.2795020 *

91) SatDom_LAI_var_80>=0.04458333 235	28.5449000 2.7119570

182) SatDom_Reflect_b2_max_80< 0.435225 172	11.2917000 2.5731400 *

183) SatDom_Reflect_b2_max_80>=0.435225 63	4.8895430 3.0909520 *

23) SatDom_MixClouds_140>=5 160	28.8156000 3.0225000 *

3) SatDom_Reflect_b7_min_160< 0.101525 7975 3308.5340000 2.3358060

6) SatDom_GPP_max_170< 0.05075 4487 1738.3760000 2.1502630

12) SatDom_Reflect_b7_mean_50>=0.077815 4162 1386.1280000 2.0897570

24) SatDom_EvapoTrans_min_140< 11.0875 252	66.6454600 1.2646030

48) SatDom_Reflect_b5_var_150< 4.28242e-05 131	10.6508700 0.8748855 *

49) SatDom_Reflect_b5_var_150>=4.28242e-05 121	14.5577400 1.6865290 *

25) SatDom_EvapoTrans_min_140>=11.0875 3910 1136.8430000 2.1429390

50) SatDom_Reflect_b5_min..60< 0.3986812 3801 1014.0490000 2.1146590

100) SatDom_Pot_EvapoTrans..20>=3196.812 3535 864.9076000 2.0663000

200) SatDom_MIR_var_120>=6.574132e-06 1243 233.4713000 1.8322530

400) SatDom_NDVI_var_100< 0.0001963981 803 110.6459000 1.6887050

800) SatDom_Reflect_b1_var_120< 8.005453e-05 623	57.9988600 1.5797590

1600) SatDom_Latent_heat_min_60< 1818750 214	13.6269300 1.3508880 *

1601) SatDom_Latent_heat_min_60>=1818750 409	27.2969000 1.6995110 *

801) SatDom_Reflect_b1_var_120>=8.005453e-05 180	19.6593900 2.0657780 *

401) SatDom_NDVI_var_100>=0.0001963981 440	76.0813400 2.0942270

802) SatDom_Reflect_b2..20>=15.29957 376	49.1128300 2.0018350

1604) BOMDom_min_temperature_var_.30>=18.90472 54	2.0670150 1.6181480 *

1605) BOMDom_min_temperature_var_.30< 18.90472 322	37.7630000 2.0661800 *

803) SatDom_Reflect_b2..20< 15.29957 64	4.9021360 2.6370310 *

201) SatDom_MIR_var_120< 6.574132e-06 2292 526.4209000 2.1932290

402) SatDom_Reflect_b2_min_70< 0.273725 1043 199.3498000 1.9753400

804) SatDom_MIR_mean_190>=0.2400412 280	25.6387700 1.6065000

1608) SatDom_Reflect_b2_min_130>=0.4367 59	2.7774580 1.2291530 *

1609) SatDom_Reflect_b2_min_130< 0.4367 221	12.2174200 1.7072400 *

805) SatDom_MIR_mean_190< 0.2400412 763 121.6401000 2.1106950

1610) SatDom_Reflect_b6_max_30>=0.2738375 328	29.1514200 1.8592680 *

1611) SatDom_Reflect_b6_max_30< 0.2738375 435	56.1197700 2.3002760

3222) SatDom_Emis31_var_170>=9.166667e-07 81	4.1131800 1.9295060 *

3223) SatDom_Emis31_var_170< 9.166667e-07 354	38.3236500 2.3851130 *

403) SatDom_Reflect_b2_min_70>=0.273725 1249 236.2046000 2.3751800

806) SatDom_Reflect_b3_mean..70>=0.09519563 157	13.7446200 1.8586620 *

807) SatDom_Reflect_b3_mean..70< 0.09519563 1092 174.5518000 2.4494410

1614) SatDom_LAI_var_70>=0.03236458 808	92.6673200 2.3476980

3228) SatDom_Pot_EvapoTrans_mean..40>=42.25625 639	57.5126600 2.2736780 *

3229) SatDom_Pot_EvapoTrans_mean..40< 42.25625 169	18.4157100 2.6275740 *

1615) SatDom_LAI_var_70< 0.03236458 284	49.7235600 2.7389080

3230) METADom_Chem_rotation_minus_2_ROUNDUP_READY>=-8.586881e-18 130 14.8366000 2.5001540 *

3231) METADom_Chem_rotation_minus_2_ROUNDUP_READY< -8.586881e-18 154 21.2208700 2.9404550 *

101) SatDom_Pot_EvapoTrans..20< 3196.812 266	31.0094000 2.7573310

202) SatDom_LST_day_var_80< 1.627956 103	8.1497690 2.5152430 *

203) SatDom_LST_day_var_80>=1.627956 163	13.0086800 2.9103070 *

51) SatDom_Reflect_b5_min..60>=0.3986812 109	13.7537100 3.1290830 *

13) SatDom_Reflect_b7_mean_50< 0.077815 325 141.8863000 2.9251080

26) BOMDom_solar_exposure_var_90< 7.378056 234	42.3384400 2.6077780

52) METADom_Crop_rotation_minus_2_sub_cropWheat< 0.5 149	19.5495900 2.4108720 *

53) METADom_Crop_rotation_minus_2_sub_cropWheat>=0.5 85	6.8851650 2.9529410 *

27) BOMDom_solar_exposure_var_90>=7.378056 91	15.3928900 3.7410990 *

7) SatDom_GPP_max_170>=0.05075 3488 1216.9760000 2.5744900

14) SatDom_Latent_heat..30< 5.10637e+07 713 220.9346000 2.1295650

28) SatDom_Emis31_var_70>=5.012346e-07 438	87.3550800 1.8387210

56) SatDom_EvapoTrans_min..80>=1.49375 337	46.4575200 1.6855490

112) BOMDom_Longitude< 138.6855 30	1.2493870 1.1093330 *

113) BOMDom_Longitude>=138.6855 307	34.2740400 1.7418570 *

57) SatDom_EvapoTrans_min..80< 1.49375 101	6.6093960 2.3498020 *

29) SatDom_Emis31_var_70< 5.012346e-07 275	37.5177400 2.5928000 *

15) SatDom_Latent_heat..30>=5.10637e+07 2775 818.6330000 2.6888070

30) SatDom_Reflect_b7_var..80>=5.860573e-07 2706 694.3568000 2.6560680

60) SatDom_Reflect_b2_var_100>=0.000288405 1113 273.2305000 2.4432260

120) SatDom_PsnNet_var_60>=6.860156e-08 904 169.8778000 2.3167040

240) BOMDom_max_temperature_min_30>=15.1 184	21.4571900 1.9597830 *

241) BOMDom_max_temperature_min_30< 15.1 720 118.9901000 2.4079170

482) SatDom_Reflect_b7_var_60< 6.300942e-05 322	50.0923300 2.2168630

964) METADom_SIMAZINE>=510 52	9.3793440 1.8567310 *

965) METADom_SIMAZINE< 510 270	32.6699500 2.2862220 *

483) SatDom_Reflect_b7_var_60>=6.300942e-05 398	47.6352400 2.5624870 *

121) SatDom_PsnNet_var_60< 6.860156e-08 209	26.2891500 2.9904780 *

61) SatDom_Reflect_b2_var_100< 0.000288405 1593 335.4769000 2.8047770

122) METADom_Chem_rotation_minus_2_GOLD< 0.0186494 820 135.4067000 2.6176950

244) SatDom_Reflect_b6_max_140< 0.204 568	64.3968400 2.4973590

488) SatDom_Reflect_b7_min_60< 0.066975 127	10.2230000 2.2200000 *

489) SatDom_Reflect_b7_min_60>=0.066975 441	41.5904200 2.5772340 *

245) SatDom_Reflect_b6_max_140>=0.204 252	44.2458100 2.8889290 *

123) METADom_Chem_rotation_minus_2_GOLD>=0.0186494 773 140.9257000 3.0032340

246) BOMDom_rainfall_csum_100>=246.5 697 100.6532000 2.9355810

492) SatDom_EVI_var_150>=0.0001223555 328	46.9338800 2.8033230 *

493) SatDom_EVI_var_150< 0.0001223555 369	42.8819500 3.0531440 *

247) BOMDom_rainfall_csum_100< 246.5 76	7.8255680 3.6236840 *

31) SatDom_Reflect_b7_var..80< 5.860573e-07 69	7.6279770 3.9727540 *
