## Supplementary Code 1 for "Explainable machine learning models of major crop traits from satellite-monitored continent-wide field trial data"

```
###############################################################################
#

## Runs an ensemble machine learning pipeline over the National Variety
Trials

###############################################################################
#

require(plyr)
require(igraph)
require(data.table)
require(beepr)
require(gdata)
require(stringr)
require(zoo)
require(umap)
library(ggplot2)
library(scales)
#require(rminer)

## and ML packages / imputation packages
require(doMC)
require(doParallel)
require(rpart)
require(randomForest)
require(xgboost)
#require(ranger)
require(missForest)
require(inTrees)
require(caret)
require(liquidSVM)
require(pls)
#install.packages("e1071")
#require(e1071)

## Please note: to use these data, it is necessary to remove factors with a
large number of levels:
## e.g. PHENDom_Trial.Comments has >800 levels

###############################################################################
#

## Constructs multiple imputations using random-sample imputation (25) and
random forests (25)

###############################################################################
#

## loads site-wise data
All_frame<-read.csv("model_frames_and_imputations/site_frame.csv", header =
T)

## strips phenotypes out, and indices
P_frame<-All_frame[,grep("PHENDom", colnames(All_frame))]
M_frame<-All_frame[,-c(grep("PHENDom", colnames(All_frame)))]
M_frame<-M_frame[,-c(grep("_Index", colnames(M_frame)))]

## removes any Inf/NaNs
is.na(M_frame)<-is.na(M_frame)

## replaces the NAs in "_exists" vectors with zeroes/FALSE
for(i in grep("_exists", colnames(M_frame))){
  M_frame[is.na(M_frame[,i]),i]<-FALSE
}

## replaces NA-rich metadata dummy-coding with three-level factor
(t/f/Unmeasured)
for(i in grep("METADom_", colnames(M_frame))){
  if(sum(is.na(M_frame[,i]))>0){
    if(!(i %in% c(grep("_time",c(colnames(M_frame)), ignore.case = T),
                  grep("_date",c(colnames(M_frame)), ignore.case = T),
                  grep("_soil",c(colnames(M_frame)), ignore.case = T),
                  grep("_10cm",c(colnames(M_frame)), ignore.case = T),
                  grep("_60cm",c(colnames(M_frame)), ignore.case = T)))){
      if(is.character(M_frame[,i]) | is.logical(M_frame[,i]) |
is.factor(M_frame[,i])){
        M_frame[is.na(M_frame[,i]),i]<-c("UNREPORTED")

        M_frame[,i]<-as.factor(M_frame[,i])
        if(length(unique(M_frame[,i]))>10){
          print(length(unique(M_frame[,i])))
          print(i)}
        #   print(i)
        #  print(c(colnames(M_frame)[i]))

      }

    }
  }
```

```
}


## removes anything >90% missing
M_frame<-M_frame[,c(colMeans(!is.na(M_frame))>=0.1)]

## and the three sites that are 80% missing

## keeps a vector of data missingness
missing_by_site<-cbind(colnames(All_frame),colMeans(is.na(All_frame)))

## measures combined missingness for environment data: 0.7%
mean(as.numeric(missing_by_site[c(grep("SatDom", missing_by_site[,1]),
                                  grep("BOMDom", missing_by_site[,1]),
                                  grep("ENVDom", missing_by_site[,1])),2]))

mean(as.numeric(missing_by_site[grep("SatDom", missing_by_site[,1]),2]))
mean(as.numeric(missing_by_site[grep("BOMDom", missing_by_site[,1]),2]))

## measures average data missingness for metadata: 74%
mean(as.numeric(missing_by_site[grep("METADom", missing_by_site[,1]),2]))

rm(All_frame)
gc()


###############################################################################
#

## First, measures the variance in accuracy generated by imputation

###############################################################################
#

## builds models, with 2018 data as holdout set, to compare variance
inserted by imputed data

## to dramatically increase processing times, uses Ranger for assessing MI
variance
require(ranger)

## cycles through the data and assesses accuracy & variance
## across each of the multiple imputations,
## using several methods.

MI_test_random_imp<-NULL
MI_test_missforest_imp<-NULL

for(i in 1:25){

  ## loads rescaled, dummy-coded data
  mod_frame1<-read.csv(file =
paste0("model_frames_and_imputations/site_imputations/RF_imputed_rescaled",i,".csv")
, header = T)

  ## attaches yield
  mod_frame1<-data.frame(Yield = P_frame$PHENDom_MEAN.YIELD,
                         mod_frame1)

  mod_frame1<-mod_frame1[!is.na(mod_frame1$Yield),]

  ## cuts post-flowering data
  mod_frame1<-mod_frame1[,-c(grep("2[0-9]{2}$", colnames(mod_frame1)))]
  mod_frame1<-mod_frame1[,-c(grep("PostSowing", colnames(mod_frame1),
ignore.case = T))]

  ## removes 2018 as holdout sample (year is rescaled)
  holdout_2018<-
mod_frame1[mod_frame1$METADom_Year!=max(mod_frame1$METADom_Year),]
  holdout_2018<-
holdout_2018[holdout_2018$METADom_Year==max(holdout_2018$METADom_Year),]
  mod_frame1<-mod_frame1[mod_frame1$METADom_Year<-as.matrix(mod_frame1[,-
c(1)])
  mod_frame2<-xgb.DMatrix(mod_frame2, label= mod_frame1$Yield)

  ##  builds rpart model
  set.seed(8394)
  system.time(rpart_mod<-rpart(Yield~., data = mod_frame1,
                            control = rpart.control(minbucket = 25,
minsplit = 50, cp = 0.01, xval = 10)))

  rpart_predicted<-predict(rpart_mod, newdata = holdout_2018)

  ##  builds BCRF (with faster/less customised ranger implementation) and
xgBoost models
  set.seed(8394)
  system.time(rf_mod<-ranger(Yield~., mod_frame1, seed = 8394, num.trees =
1000))

  rf_predicted<-predict(rf_mod, data = holdout_2018)

  set.seed(8394)
  system.time(xg_mod<-xgboost(data = mod_frame2, nthread = 16, nrounds
```

```
=10))

  holdout_2018_mat<-as.matrix(holdout_2018[,-c(1)])

  xg_predicted<-predict(xg_mod, newdata = holdout_2018_mat)

  cor.test(xg_predicted, holdout_2018$Yield)
  RMSE(xg_predicted, holdout_2018$Yield)

  ## removes constant data
  holdout_2018_2<-holdout_2018[,which(unlist(sapply(mod_frame1,var))>0)]
  holdout_Y<-holdout_2018_2[,c(1)]
  holdout_2018_2<-holdout_2018_2[,-c(1)]
  mod_frame2<-mod_frame1[,which(unlist(sapply(mod_frame1,var))>0)]
  Yield_vals<-mod_frame2[,1]
  mod_frame2<-mod_frame2[,-c(1)]

  ## Builds linear SVMs using linearSVM
  system.time(lsvm_mod<-svmRegression(mod_frame2, Yield_vals))

  lsvm_prediction<-predict(lsvm_mod, newdata = holdout_2018_2)


  mod_frame2<-mod_frame1[,which(unlist(sapply(mod_frame1,var))>0)]

  pls.options(parallel = makeCluster(10, type = "PSOCK"))
  ## Builds a PLSR model, 10-fold internally cross-validated, 20 components
  system.time(pslr_mod<-plsr(Yield~., ncomp = 10, data = mod_frame2,
validation = "CV", segments  =10))

  plsr_prediction<-as.vector(predict(pslr_mod, newdata = holdout_2018_2,
comps = 10))

  stopCluster(pls.options()$parallel)

#  plsr_prediction<-predict(pslr_mod, newdata = holdout_2018_2)

  ## PLSR is complete merde???!?!?!?

  ## collates results
  MI_test_missforest_imp[[i]]<-data.frame(rpart_R =
c(cor.test(rpart_predicted, holdout_2018$Yield)$estimate),
                                          rpart_RMSE =
RMSE(rpart_predicted, holdout_2018$Yield),
                                          bcrf_R =
c(cor.test(rf_predicted$predictions, holdout_2018$Yield)$estimate),
                                          bcrf_RMSE =
RMSE(rf_predicted$predictions, holdout_2018$Yield),
                                          xgb_R = c(cor.test(xg_predicted,
holdout_2018$Yield)$estimate),
                                          xgb_RMSE = RMSE(xg_predicted,
holdout_2018$Yield),
                                          lsvm_R =
c(cor.test(lsvm_prediction, holdout_Y)$estimate),
                                          lsvm_RMSE = RMSE(lsvm_prediction,
holdout_Y),
                                          plsr_R =
c(cor.test(plsr_prediction, holdout_Y)$estimate),
                                          plsr_RMSE = RMSE(plsr_prediction,
holdout_Y))
  print(i)

 #beep(2)


}

MI_test_missforest_imp<-ldply(MI_test_missforest_imp, as.data.frame)

## repeats this process for random-sample imputations

for(i in 1:25){

  ## loads rescaled, dummy-coded data
  mod_frame1<-read.csv(file =
paste0("model_frames_and_imputations/site_imputations/random_sample_rescaled",i,".csv")
, header = T)

  ## attaches yield
  mod_frame1<-data.frame(Yield = P_frame$PHENDom_MEAN.YIELD,
                         mod_frame1)

  mod_frame1<-mod_frame1[!is.na(mod_frame1$Yield),]

  ## cuts post-flowering data
  mod_frame1<-mod_frame1[,-c(grep("2[0-9]{2}$", colnames(mod_frame1)))]
  mod_frame1<-mod_frame1[,-c(grep("PostSowing", colnames(mod_frame1),
ignore.case = T))]

  ## removes 2018 as holdout sample (year is rescaled)
  holdout_2018<-
mod_frame1[mod_frame1$METADom_Year!=max(mod_frame1$METADom_Year),]
  holdout_2018<-
```

```
holdout_2018[holdout_2018$METADom_Year==max(holdout_2018$METADom_Year),]
  mod_frame1<-mod_frame1[mod_frame1$METADom_Year<-as.matrix(mod_frame1[,-
c(1)])
  mod_frame2<-xgb.DMatrix(mod_frame2, label= mod_frame1$Yield)

  ##  builds rpart model
  set.seed(8394)
  system.time(rpart_mod<-rpart(Yield~., data = mod_frame1,
                             control = rpart.control(minbucket = 25,
minsplit = 50, cp = 0.01, xval = 10)))

  rpart_predicted<-predict(rpart_mod, newdata = holdout_2018)

  ##  builds BCRF (with faster/less customised ranger implementation) and
xgBoost models
  set.seed(8394)
  system.time(rf_mod<-ranger(Yield~., mod_frame1, seed = 8394, num.trees =
1000))

  rf_predicted<-predict(rf_mod, data = holdout_2018)

  set.seed(8394)
  system.time(xg_mod<-xgboost(data = mod_frame2, nthread = 16, nrounds
=10))

  holdout_2018_mat<-as.matrix(holdout_2018[,-c(1)])

  xg_predicted<-predict(xg_mod, newdata = holdout_2018_mat)

  cor.test(xg_predicted, holdout_2018$Yield)
  RMSE(xg_predicted, holdout_2018$Yield)

  ## removes constant data
  holdout_2018_2<-holdout_2018[,which(unlist(sapply(mod_frame1,var))>0)]
  holdout_Y<-holdout_2018_2[,c(1)]
  holdout_2018_2<-holdout_2018_2[,-c(1)]
  mod_frame2<-mod_frame1[,which(unlist(sapply(mod_frame1,var))>0)]
  Yield_vals<-mod_frame2[,1]
  mod_frame2<-mod_frame2[,-c(1)]

  ## Builds linear SVMs using linearSVM
  system.time(lsvm_mod<-svmRegression(mod_frame2, Yield_vals))

  lsvm_prediction<-predict(lsvm_mod, newdata = holdout_2018_2)


  mod_frame2<-mod_frame1[,which(unlist(sapply(mod_frame1,var))>0)]


  pls.options(parallel = makeCluster(10, type = "PSOCK"))
  ## Builds a PLSR model, 10-fold internally cross-validated, 20 components
  system.time(pslr_mod<-plsr(Yield~., ncomp = 10, data = mod_frame2,
validation = "CV", segments  =10))

  plsr_prediction<-as.vector(predict(pslr_mod, newdata = holdout_2018_2,
comps = 10))

  stopCluster(pls.options()$parallel)

  #  plsr_prediction<-predict(pslr_mod, newdata = holdout_2018_2)

  ## PLSR is complete merde???!?!?!?

  ## collates results
  MI_test_random_imp[[i]]<-data.frame(rpart_R = c(cor.test(rpart_predicted,
holdout_2018$Yield)$estimate),
                                          rpart_RMSE =
RMSE(rpart_predicted, holdout_2018$Yield),
                                          bcrf_R =
c(cor.test(rf_predicted$predictions, holdout_2018$Yield)$estimate),
                                          bcrf_RMSE =
RMSE(rf_predicted$predictions, holdout_2018$Yield),
                                          xgb_R = c(cor.test(xg_predicted,
holdout_2018$Yield)$estimate),
                                          xgb_RMSE = RMSE(xg_predicted,
holdout_2018$Yield),
                                          lsvm_R =
c(cor.test(lsvm_prediction, holdout_Y)$estimate),
                                          lsvm_RMSE = RMSE(lsvm_prediction,
holdout_Y),
                                          plsr_R =
c(cor.test(plsr_prediction, holdout_Y)$estimate),
                                          plsr_RMSE = RMSE(plsr_prediction,
holdout_Y))
  print(i)

 # beep(2)
}

MI_test_random_imp<-ldply(MI_test_random_imp, as.data.frame)

saveRDS(MI_test_missforest_imp, file =
"model_frames_and_imputations/missforest_imp_test.rds")
```

```
saveRDS(MI_test_random_imp, file =
"model_frames_and_imputations/random_imp_test.rds")

detach(package:ranger)

## re-loads random forest to fix/overwrite namespace
require(randomForest)

##############################################################################
#

## builds a model for all-site data, undifferentiated by species or
variety,
## predicting species-normalised (relative) yield variation

##############################################################################
#

## constructs site-wise models

## avoids the rescaled data for tree-based partitioning,
## to allow interpretation of raw values

mod_frame1<-read.csv(file =
"model_frames_and_imputations/site_imputations/RF_imputed_1.csv", header =
T)

## fixes the stupid 2018-specific "" level in soil tests
mod_frame1$METADom_P_test_class_10cm.test_10cm[mod_frame1$METADom_P_test_class_10cm.test_10cm==""]<
-"UNMEASURED"
mod_frame1$METADom_P_test_class_60cm.test_60cm[mod_frame1$METADom_P_test_class_60cm.test_60cm==""]<
-"UNMEASURED"

## substitutes the single "Other" crop rotation for "UNK"
#mod_frame1$METADom_Crop_rotation_minus_0[mod_frame1$METADom_Crop_rotation_minus_0=="Other"]<
-"UNK"

## dummy-codes crop rotation data

Dummy_crop<-
cbind(as.matrix(model.matrix(~as.factor(mod_frame1$METADom_Crop_rotation_minus_0)))
,

as.matrix(model.matrix(~as.factor(mod_frame1$METADom_Crop_rotation_minus_1)))
,

as.matrix(model.matrix(~as.factor(mod_frame1$METADom_Crop_rotation_minus_2)))
,

as.matrix(model.matrix(~as.factor(mod_frame1$METADom_Crop_rotation_minus_3)))
,

as.matrix(model.matrix(~as.factor(mod_frame1$METADom_Crop_rotation_minus_4)))
,

as.matrix(model.matrix(~as.factor(mod_frame1$METADom_Crop_rotation_minus_5)))
,

as.matrix(model.matrix(~as.factor(mod_frame1$METADom_Crop_rotation_minus_6)))
)

colnames(Dummy_crop)<-
paste0("METADom_Crop_rotation_",unlist(lapply(strsplit(colnames(Dummy_crop)
,

split = "[0-9])"), tail, 1)),
                             "_minus_")

colnames(Dummy_crop)[1:19]<-paste0(colnames(Dummy_crop)[1:19], 0)
colnames(Dummy_crop)[20:41]<-paste0(colnames(Dummy_crop)[20:41], 1)
colnames(Dummy_crop)[42:63]<-paste0(colnames(Dummy_crop)[42:63], 2)
colnames(Dummy_crop)[64:84]<-paste0(colnames(Dummy_crop)[64:84], 3)
colnames(Dummy_crop)[85:100]<-paste0(colnames(Dummy_crop)[85:100], 4)
colnames(Dummy_crop)[101:112]<-paste0(colnames(Dummy_crop)[101:112], 5)
colnames(Dummy_crop)[113:119]<-paste0(colnames(Dummy_crop)[113:119], 6)

## removes crop rotation factor
mod_frame1<-mod_frame1[,-c(grep("METADom_Crop_rotation_",
colnames(mod_frame1)))]

mod_frame1<-data.frame(mod_frame1, Dummy_crop)

is.na(mod_frame1)<-is.na(mod_frame1)


mod_frame1<-data.frame(Yield = P_frame$PHENDom_MEAN.YIELD,
                       mod_frame1)

mod_frame1<-mod_frame1[!is.na(mod_frame1$Yield),]

## changes character to factor, integer to numeric
for(i in 1:ncol(mod_frame1)){
  if(is.character(mod_frame1[,i])){mod_frame1[,i]<-
```

```
as.factor(mod_frame1[,i])}
  if(is.integer(mod_frame1[,i])){mod_frame1[,i]<-
as.numeric(mod_frame1[,i])}
}

## removes 2018 as holdout sample (year is rescaled)
holdout_2018<-mod_frame1[mod_frame1$METADom_Year>=2018,]
mod_frame1<-mod_frame1[mod_frame1$METADom_Year<2018,]

set.seed(8394)
system.time(rpart_mod<-rpart(Yield~., data = mod_frame1,
                             control = rpart.control(minbucket = 25,
minsplit = 50, cp = 0.001, xval = 10)))


cor.test(predict(rpart_mod, newdata = holdout_2018), holdout_2018$Yield)
rpart_prediction<-predict(rpart_mod, newdata = holdout_2018)

#mod_frame1<-as.matrix(mod_frame1)
#mod_frame1<-as.data.frame(mod_frame1)
## adds a crossvalidation strata - year
set.seed(8394)
system.time(rf_mod<-randomForest(Yield~., mod_frame1,
                                 strata = "METADom_Year",
                                 xval = 10, minbucket = 100, minsplit =
300, mtry = 500))

cor.test(predict(rf_mod, newdata = holdout_2018), holdout_2018$Yield)
rf_prediction<-predict(rf_mod, newdata = holdout_2018)

beep()


## loads rescaled, dummy-coded data
mod_frame2<-read.csv(file =
paste0("model_frames_and_imputations/site_imputations/rf_imputed_rescaled1.csv")
, header = T)

## attaches yield
mod_frame2<-data.frame(Yield = P_frame$PHENDom_MEAN.YIELD,
                       mod_frame2)

mod_frame2<-mod_frame2[!is.na(mod_frame2$Yield),]

## removes 2018 as holdout sample (year is rescaled)
holdout_2018_2<-
mod_frame2[mod_frame2$METADom_Year!=max(mod_frame2$METADom_Year),]
holdout_2018_2<-
holdout_2018_2[holdout_2018_2$METADom_Year==max(holdout_2018_2$METADom_Year),
]
mod_frame2<-mod_frame2[mod_frame2$METADom_Year<-as.matrix(mod_frame2[,-
c(1)])
mod_frame3<-xgb.DMatrix(mod_frame3, label= mod_frame2$Yield)

set.seed(8394)
system.time(xg_mod<-xgboost(data = mod_frame3, nthread = 16, nrounds  =10,
save_name = "xg_sitewise"))

holdout_2018_mat<-as.matrix(holdout_2018_2[,-c(1)])

xg_prediction<-predict(xg_mod, newdata = holdout_2018_mat)

cor.test(xg_prediction, holdout_2018_2$Yield)

## removes constant data
holdout_2018_3<-holdout_2018_2[,which(unlist(sapply(mod_frame2,var))>0)]
holdout_Y<-holdout_2018_3[,c(1)]
holdout_2018_3<-holdout_2018_3[,-c(1)]
mod_frame3<-mod_frame2[,which(unlist(sapply(mod_frame2,var))>0)]
Yield_vals<-mod_frame3[,1]
mod_frame3<-mod_frame3[,-c(1)]

## Builds linear SVMs using linearSVM
system.time(lsvm_mod<-svmRegression(mod_frame3, Yield_vals))

lsvm_prediction<-predict(lsvm_mod, newdata = holdout_2018_3)

cor.test(lsvm_prediction, holdout_Y)

mod_frame3<-data.frame(Yield = Yield_vals, mod_frame3)

pls.options(parallel = makeCluster(10, type = "PSOCK"))
## Builds a PLSR model, 10-fold internally cross-validated, 20 components
system.time(pslr_mod<-plsr(Yield~., ncomp = 10, data = mod_frame3,
validation = "CV", segments  =10))

plsr_prediction<-as.vector(predict(pslr_mod, newdata = holdout_2018_3,
ncomp = 10))

stopCluster(pls.options()$parallel)

#  plsr_prediction<-predict(pslr_mod, newdata = holdout_2018_2)
```

```
## attaches the predictions to the target data
xobj<-data.frame(Yield = holdout_2018$Yield,
  rpart_prediction,
         rf_prediction,
         xg_prediction,
         lsvm_prediction,
         plsr_prediction)

## saves models, target data
saveRDS(xobj,file =
"model_frames_and_imputations/site_models/site_allTimes_PredObs.rds")
saveRDS(rpart_mod,file =
"model_frames_and_imputations/site_models/site_allTimes_rpart.rds")
saveRDS(rf_mod,file =
"model_frames_and_imputations/site_models/site_allTimes_bcrf.rds")
saveRDS(xg_mod,file =
"model_frames_and_imputations/site_models/site_allTimes_xgb.rds")
saveRDS(lsvm_mod,file =
"model_frames_and_imputations/site_models/site_allTimes_lsvm.rds")
saveRDS(pslr_mod,file =
"model_frames_and_imputations/site_models/site_allTimes_plsr.rds")

## cleans up workspace
rm(xobj, rpart_mod, rf_mod, xg_mod, lsvm_mod, pslr_mod)
gc()

## Now constructs the time-adjusted data: removes all data to 90DAS, and
all data after sowing.

## removes data after days 90-99, including post-sowing

mod_frame1<-mod_frame1[,-c(grep("2[0-9]{2}$", colnames(mod_frame1)))]
mod_frame1<-mod_frame1[,-c(grep("1[0-9]{2}$", colnames(mod_frame1)))]
mod_frame1<-mod_frame1[,-c(grep("PostSowing", colnames(mod_frame1),
ignore.case = T))]

holdout_2018<-holdout_2018[,-c(grep("2[0-9]{2}$", colnames(holdout_2018)))]
holdout_2018<-holdout_2018[,-c(grep("1[0-9]{2}$", colnames(holdout_2018)))]
holdout_2018<-holdout_2018[,-c(grep("PostSowing", colnames(holdout_2018),
ignore.case = T))]

#holdout_2018<-mod_frame1[mod_frame1$METADom_Year==2018,]
#mod_frame1<-mod_frame1[mod_frame1$METADom_Year<2018,]

## rebuilds models
set.seed(8394)
system.time(rpart_mod<-rpart(Yield~., data = mod_frame1,
                             control = rpart.control(minbucket = 25,
minsplit = 50, cp = 0.001, xval = 10)))


cor.test(predict(rpart_mod, newdata = holdout_2018), holdout_2018$Yield)
rpart_prediction<-predict(rpart_mod, newdata = holdout_2018)

set.seed(8394)
system.time(rf_mod<-randomForest(Yield~., mod_frame1, xval = 10, minbucket
= 100, minsplit = 300, mtry = 500))

cor.test(predict(rf_mod, newdata = holdout_2018), holdout_2018$Yield)
rf_prediction<-predict(rf_mod, newdata = holdout_2018)

beep()

mod_frame2<-mod_frame2[,-c(grep("2[0-9]{2}$", colnames(mod_frame2)))]
mod_frame2<-mod_frame2[,-c(grep("1[0-9]{2}$", colnames(mod_frame2)))]
mod_frame2<-mod_frame2[,-c(grep("PostSowing", colnames(mod_frame2),
ignore.case = T))]

holdout_2018_2<-holdout_2018_2[,-c(grep("2[0-9]{2}$",
colnames(holdout_2018_2)))]
holdout_2018_2<-holdout_2018_2[,-c(grep("1[0-9]{2}$",
colnames(holdout_2018_2)))]
holdout_2018_2<-holdout_2018_2[,-c(grep("PostSowing",
colnames(holdout_2018_2), ignore.case = T))]

mod_frame3<-as.matrix(mod_frame2[,-c(1)])
mod_frame3<-xgb.DMatrix(mod_frame3, label= mod_frame2$Yield)

set.seed(8394)
system.time(xg_mod<-xgboost(data = mod_frame3, nthread = 16, nrounds  =10,
save_name = "xg_sitewise"))

holdout_2018_mat<-as.matrix(holdout_2018_2[,-c(1)])

xg_prediction<-predict(xg_mod, newdata = holdout_2018_mat)
cor.test(xg_prediction, holdout_2018_2$Yield)

## removes constant data
holdout_2018_3<-holdout_2018_2[,which(unlist(sapply(mod_frame2,var))>0)]
holdout_Y<-holdout_2018_3[,c(1)]
holdout_2018_3<-holdout_2018_3[,-c(1)]
mod_frame3<-mod_frame2[,which(unlist(sapply(mod_frame2,var))>0)]
Yield_vals<-mod_frame3[,1]
```

```
mod_frame3<-mod_frame3[,-c(1)]

## Builds linear SVMs using linearSVM
system.time(lsvm_mod<-svmRegression(mod_frame3, Yield_vals))

lsvm_prediction<-predict(lsvm_mod, newdata = holdout_2018_3)

cor.test(lsvm_prediction, holdout_Y)

mod_frame3<-data.frame(Yield = Yield_vals, mod_frame3)

pls.options(parallel = makeCluster(10, type = "PSOCK"))
## Builds a PLSR model, 10-fold internally cross-validated, 20 components
system.time(pslr_mod<-plsr(Yield~., ncomp = 10, data = mod_frame3,
validation = "CV", segments  =10))

plsr_prediction<-as.vector(predict(pslr_mod, newdata = holdout_2018_3,
ncomp = 10))

stopCluster(pls.options()$parallel)

## attaches the predictions to the target data
xobj<-data.frame(Yield = holdout_2018$Yield,
                 rpart_prediction,
                 rf_prediction,
                 xg_prediction,
                 lsvm_prediction,
                 plsr_prediction)

## saves models, target data
saveRDS(xobj,file =
"model_frames_and_imputations/site_models/site_90DAS_PredObs.rds")
saveRDS(rpart_mod,file =
"model_frames_and_imputations/site_models/site_90DAS_rpart.rds")
saveRDS(rf_mod,file =
"model_frames_and_imputations/site_models/site_90DAS_bcrf.rds")
saveRDS(xg_mod,file =
"model_frames_and_imputations/site_models/site_90DAS_xgb.rds")
saveRDS(lsvm_mod,file =
"model_frames_and_imputations/site_models/site_90DAS_lsvm.rds")
saveRDS(pslr_mod,file =
"model_frames_and_imputations/site_models/site_90DAS_plsr.rds")

## cleans up workspace
rm(xobj, rpart_mod, rf_mod, xg_mod, lsvm_mod, pslr_mod)
gc()

## another tedious repeat:
## removes data after sowing

mod_frame1<-mod_frame1[,-c(grep("_[0-9]{2}$", colnames(mod_frame1)))]
holdout_2018<-holdout_2018[,-c(grep("_[0-9]{2}$", colnames(holdout_2018)))]

## rebuilds models
set.seed(8394)
system.time(rpart_mod<-rpart(Yield~., data = mod_frame1,
                             control = rpart.control(minbucket = 25,
minsplit = 50, cp = 0.001, xval = 10)))


cor.test(predict(rpart_mod, newdata = holdout_2018), holdout_2018$Yield)
rpart_prediction<-predict(rpart_mod, newdata = holdout_2018)

set.seed(8394)
system.time(rf_mod<-randomForest(Yield~., mod_frame1, xval = 10, minbucket
= 100, minsplit = 300, mtry = 500))

cor.test(predict(rf_mod, newdata = holdout_2018), holdout_2018$Yield)
rf_prediction<-predict(rf_mod, newdata = holdout_2018)

beep()

mod_frame2<-mod_frame2[,-c(grep("_[0-9]{2}$", colnames(mod_frame2)))]
holdout_2018_2<-holdout_2018_2[,-c(grep("_[0-9]{2}$",
colnames(holdout_2018_2)))]

mod_frame3<-as.matrix(mod_frame2[,-c(1)])
mod_frame3<-xgb.DMatrix(mod_frame3, label= mod_frame2$Yield)

set.seed(8394)
system.time(xg_mod<-xgboost(data = mod_frame3, nthread = 16, nrounds  =10,
save_name = "xg_sitewise"))

holdout_2018_mat<-as.matrix(holdout_2018_2[,-c(1)])

xg_prediction<-predict(xg_mod, newdata = holdout_2018_mat)
cor.test(xg_prediction, holdout_2018_2$Yield)

## removes constant data
holdout_2018_3<-holdout_2018_2[,which(unlist(sapply(mod_frame2,var))>0)]
holdout_Y<-holdout_2018_3[,c(1)]
holdout_2018_3<-holdout_2018_3[,-c(1)]
mod_frame3<-mod_frame2[,which(unlist(sapply(mod_frame2,var))>0)]
```

```
Yield_vals<-mod_frame3[,1]
mod_frame3<-mod_frame3[,-c(1)]

## Builds linear SVMs using linearSVM
system.time(lsvm_mod<-svmRegression(mod_frame3, Yield_vals))

lsvm_prediction<-predict(lsvm_mod, newdata = holdout_2018_3)

cor.test(lsvm_prediction, holdout_Y)

mod_frame3<-data.frame(Yield = Yield_vals, mod_frame3)

pls.options(parallel = makeCluster(10, type = "PSOCK"))
## Builds a PLSR model, 10-fold internally cross-validated, 20 components
system.time(pslr_mod<-plsr(Yield~., ncomp = 10, data = mod_frame3,
validation = "CV", segments  =10))

plsr_prediction<-as.vector(predict(pslr_mod, newdata = holdout_2018_3,
ncomp = 10))

stopCluster(pls.options()$parallel)

## attaches the predictions to the target data
xobj<-data.frame(Yield = holdout_2018$Yield,
                  rpart_prediction,
                  rf_prediction,
                  xg_prediction,
                  lsvm_prediction,
                  plsr_prediction)

## saves models, target data
saveRDS(xobj,file =
"model_frames_and_imputations/site_models/site_atSowing_PredObs.rds")
saveRDS(rpart_mod,file =
"model_frames_and_imputations/site_models/site_atSowing_rpart.rds")
saveRDS(rf_mod,file =
"model_frames_and_imputations/site_models/site_atSowing_bcrf.rds")
saveRDS(xg_mod,file =
"model_frames_and_imputations/site_models/site_atSowing_xgb.rds")
saveRDS(lsvm_mod,file =
"model_frames_and_imputations/site_models/site_atSowing_lsvm.rds")
saveRDS(pslr_mod,file =
"model_frames_and_imputations/site_models/site_atSowing_plsr.rds")

## cleans up workspace
rm(xobj, rpart_mod, rf_mod, xg_mod, lsvm_mod, pslr_mod,
   mod_frame1, mod_frame2, mod_frame3,
   holdout_2018, holdout_2018_2,holdout_2018_3,holdout_2018_mat,
   rpart_prediction,rf_prediction, xg_prediction,
lsvm_prediction,plsr_prediction, holdout_Y)
gc()

##############################################################################
#

## Now for the core target: constructs the large, variety-differentiated
dataset

##############################################################################
#

## switches to readr
#require(readr)

## gets the first imputation, aligns with the extensive data
#mod_frame1<-read_csv(file =
"model_frames_and_imputations/site_imputations/RF_imputed_1.csv")
mod_frame1<-read.csv(file =
"model_frames_and_imputations/site_imputations/RF_imputed_1.csv", header =
T)

## adds the P-frame data back on (as imputations only included E/M data)
mod_frame1<-data.frame(mod_frame1,P_frame)

mod_frame2<-read.csv(file =
"model_frames_and_imputations/variety_frame.csv", header = T)

#mod_frame1<-as.data.frame(mod_frame1)
#mod_frame2<-as.data.frame(mod_frame2)

## removes sites with no BOM or Satellite data, due to missing sowing dates
(N=11 sites, 104 variety-years)
mod_frame2<-mod_frame2[-
c(which(is.na(mod_frame2$BOMDom_Sowing_date_numeric))),]

## removes (non-imputed, na-rich) columns from variety frame that are in
the site frame
sum(!(colnames(mod_frame2) %in% colnames(mod_frame1)))

head(mod_frame2[,1:10])
head(mod_frame1[,1:10])

index_1<-
```

```
paste(mod_frame1$BOMDom_Sowing_date_numeric,mod_frame1$BOMDom_Latitude,mod_frame1$BOMDom_Longitude
, sep = "_")
index_2<-
paste(mod_frame2$BOMDom_Sowing_date_numeric,mod_frame2$BOMDom_Latitude,mod_frame2$BOMDom_Longitude
, sep = "_")

mod_frame1$index_1<-index_1
mod_frame2$index_2<-index_2

## data that was dummy-coded
colnames(mod_frame2)[which(!(colnames(mod_frame2) %in%
colnames(mod_frame1)))]

## trims to the variables that are unique to the variety data frame
## keeps the site-year index
mod_frame2<-mod_frame2[,c(1,which(!(colnames(mod_frame2) %in%
colnames(mod_frame1))))]

## and merges
mod_frame1<-mod_frame1[match(mod_frame2$index_2, mod_frame1$index_1),]
mod_frame1<-data.frame(mod_frame1, mod_frame2)

## Fills the dummy-coded data, adds "unknowns"

## breeders
mod_frame1$MANDom_BreederUNKNOWN[is.na(mod_frame1$MANDom_BreederUNKNOWN)]<-
1

for(i in unique(grep("MANDom_Breeder", colnames(mod_frame1)))){
  if(sum(is.na(mod_frame1[,i]))>0){
    mod_frame1[is.na(mod_frame1[,i]),i]<-0
  }
}

## Takes factors with NAs and replaces with 1/true in "UNKNOWN" dummy
coding variables

## starts with management/series data
mod_frame1$MANDom_Series_nameUNKNOWN[is.na(mod_frame1$MANDom_Series_nameUNKNOWN)]<
-1

for(i in unique(grep("MANDom_Series", colnames(mod_frame1)))){
  if(sum(is.na(mod_frame1[,i]))>0){
    mod_frame1[is.na(mod_frame1[,i]),i]<-0
  }
}

mod_frame1$Trial_seriesUNKNOWN[is.na(mod_frame1$Trial_seriesUNKNOWN)]<-1
for(i in unique(grep("Trial_series", colnames(mod_frame1)))){
  if(sum(is.na(mod_frame1[,i]))>0){
    mod_frame1[is.na(mod_frame1[,i]),i]<-0
  }
}

mod_frame1$MANDom_Trial_operatorsUNKNOWN[is.na(mod_frame1$MANDom_Trial_operatorsUNKNOWN)]<
-1
for(i in unique(grep("MANDom_Trial_operators", colnames(mod_frame1)))){
  if(sum(is.na(mod_frame1[,i]))>0){
    mod_frame1[is.na(mod_frame1[,i]),i]<-0
  }
}

## fixes chem/fertiliser/crop rotations, adds "unknown" dummy-coding
sub_frame<-mod_frame1[,grep("_rotation_minus", colnames(mod_frame1))]

sub_frame$METADom_Chem_rotation_minus_0_UNKNOWN<-
is.na(sub_frame$METADom_Chem_rotation_minus_0_BOXERGOLD)
sub_frame$METADom_Chem_rotation_minus_1_UNKNOWN<-
is.na(sub_frame$METADom_Chem_rotation_minus_1_BOXERGOLD)
sub_frame$METADom_Chem_rotation_minus_2_UNKNOWN<-
is.na(sub_frame$METADom_Chem_rotation_minus_2_BOXERGOLD)
sub_frame$METADom_Chem_rotation_minus_3_UNKNOWN<-
is.na(sub_frame$METADom_Chem_rotation_minus_3_BOXERGOLD)
sub_frame$METADom_Chem_rotation_minus_4_UNKNOWN<-
is.na(sub_frame$METADom_Chem_rotation_minus_4_N)
sub_frame$METADom_Chem_rotation_minus_5_UNKNOWN<-
is.na(sub_frame$METADom_Chem_rotation_minus_5_N)
sub_frame$METADom_Chem_rotation_minus_6_UNKNOWN<-
is.na(sub_frame$METADom_Chem_rotation_minus_6_N)

sub_frame$METADom_Fert_rotation_minus_0_UNKNOWN<-
is.na(sub_frame$METADom_Fert_rotation_minus_0_PHOS)
sub_frame$METADom_Fert_rotation_minus_1_UNKNOWN<-
is.na(sub_frame$METADom_Fert_rotation_minus_1_PHOS)
sub_frame$METADom_Fert_rotation_minus_2_UNKNOWN<-
is.na(sub_frame$METADom_Fert_rotation_minus_2_PHOS)
sub_frame$METADom_Fert_rotation_minus_3_UNKNOWN<-
is.na(sub_frame$METADom_Fert_rotation_minus_3_PHOS)
sub_frame$METADom_Fert_rotation_minus_4_UNKNOWN<-
is.na(sub_frame$METADom_Fert_rotation_minus_4_PHOS)
sub_frame$METADom_Fert_rotation_minus_5_UNKNOWN<-
is.na(sub_frame$METADom_Fert_rotation_minus_5_N)
sub_frame$METADom_Fert_rotation_minus_6_UNKNOWN<-
```

```
is.na(sub_frame$METADom_Fert_rotation_minus_6_N)

## fills all NAs with zeroes
for(i in 1:ncol(sub_frame)){
  if(sum(is.na(sub_frame[,i]))>0){
    sub_frame[is.na(sub_frame[,i]),i]<-0
  }
}

sum(is.na(sub_frame))

## replaces NA-rich dummy-coded data with properly coded data
mod_frame1<-mod_frame1[,-c(grep("_rotation_minus", colnames(mod_frame1)))]
mod_frame1<-data.frame(mod_frame1, sub_frame)

rm(sub_frame)

## removes the nearly-absent chemical timing and fertiliser timing data
sub_frame<-mod_frame1[,c(grep("METADom_Fertiliser_time",
colnames(mod_frame1)),
                         grep("METADom_Chemicals_time",
colnames(mod_frame1)))]

peanutCluster<-makeCluster(5)
registerDoParallel(peanutCluster)

## only retains the 25 columns that actually contain sufficient values:
## the rest is largely going to be imputation junk...
sub_frame<-sub_frame[,colMeans(is.na(sub_frame))<0.9]

## runs missForest imputation on these data
set.seed(38485)
sub_Miss<-missForest(sub_frame, maxiter = 5, mtry = 25, ntree = 10,
maxnodes =10, parallelize = c('variables'))

#saveRDS(sub_Miss, "temp_miss.rds")
sub_frame<-sub_Miss$ximp

## integrates back into the data frame, removes redundant timings
mod_frame1<-mod_frame1[,-c(grep("METADom_Fertiliser_time",
colnames(mod_frame1)),
                          grep("METADom_Chemicals_time",
colnames(mod_frame1)))]

mod_frame1<-data.frame(mod_frame1, sub_frame)

## removes the 6k-level factor "Trial ID"
#mod_frame1<-mod_frame1[,-c(grep("MANDom_TrialCode",
colnames(mod_frame1)))]

## removes the repeated soil test data, whjich are almost completely
missing
mod_frame1<-mod_frame1[,-c(grep("cm_repeat_1$", colnames(mod_frame1)))]

## and removes the "germination rain date", which is 99% missing and
embedded in both the BOM and satellite data anyway
mod_frame1<-mod_frame1[,-c(grep("METADom_GERMINATION.RAIN.DATE$",
colnames(mod_frame1)))]

## finally, adds negative values to the integer/mixed environment scores
## observed in the field
mod_frame1$MANDom_Farm_machinery[is.na(mod_frame1$MANDom_Farm_machinery)]<-
c(-1)
mod_frame1$ENVDom_Bird_or_Animal_damage_score[is.na(mod_frame1$ENVDom_Bird_or_Animal_damage_score)]<
-c(-1)
mod_frame1$ENVDom_Bird_or_Animal_damage_score_obs_2[is.na(mod_frame1$ENVDom_Bird_or_Animal_damage_score_obs_2)]<
-c(-1)
mod_frame1$ENVDom_Frost_Damage_score[is.na(mod_frame1$ENVDom_Frost_Damage_score)]<
-c(-1)
mod_frame1$ENVDom_Herbicide_damage[is.na(mod_frame1$ENVDom_Herbicide_damage)]<
-c(-1)
mod_frame1$ENVDom_Grazing_damage[is.na(mod_frame1$ENVDom_Grazing_damage)]<-
c(-1)

## rescales these yet again to zero mean, unit variance
mod_frame1$MANDom_Farm_machinery<-scale(mod_frame1$MANDom_Farm_machinery)
mod_frame1$ENVDom_Bird_or_Animal_damage_score<-
scale(mod_frame1$ENVDom_Bird_or_Animal_damage_score)
mod_frame1$ENVDom_Bird_or_Animal_damage_score_obs_2<-
scale(mod_frame1$ENVDom_Bird_or_Animal_damage_score_obs_2)
mod_frame1$ENVDom_Frost_Damage_score<-
scale(mod_frame1$ENVDom_Frost_Damage_score)
mod_frame1$ENVDom_Herbicide_damage<-
scale(mod_frame1$ENVDom_Herbicide_damage)
mod_frame1$ENVDom_Grazing_damage<-scale(mod_frame1$ENVDom_Grazing_damage)

## and collapses the crop type into a single variable
Crop_all<-as.character(mod_frame1$PHENDom_Crop.Type)
mean(Crop_all==as.character(mod_frame1$PHENDom_Crop), na.rm = T)
Crop_all[is.na(Crop_all)]<-
as.character(mod_frame1$PHENDom_Crop[is.na(Crop_all)])

#Crop_all[is.na(Crop_all)]<-"UNREPORTED"
```

```
mod_frame1$PHENDom_Crop_alldata<-as.factor(Crop_all)

## removes the indices
mod_frame1<-mod_frame1[,-c(grep("index",colnames(mod_frame1), ignore.case =
T))]

## Keeps a vector where "-" value yields are retained as zeroes
mod_frame1$PHENDom_zeroInflated_yield_t_ha<-
as.character(mod_frame1$PHENDom_yield_t_ha)
mod_frame1$PHENDom_zeroInflated_yield_t_ha[mod_frame1$PHENDom_zeroInflated_yield_t_ha=="
-"]<-0
mod_frame1$PHENDom_zeroInflated_yield_t_ha<-
as.numeric(mod_frame1$PHENDom_zeroInflated_yield_t_ha)

sum(mod_frame1$PHENDom_yield_t_ha=="-", na.rm = T)
mean(mod_frame1$PHENDom_yield_t_ha=="-", na.rm = T)

## removes the "-" values from yield phenotypes, replaces with NAs
mod_frame1$PHENDom_yield_t_ha[mod_frame1$PHENDom_yield_t_ha=="-"]<-NA
mod_frame1$PHENDom_yield_t_ha<-
as.numeric(as.character(mod_frame1$PHENDom_yield_t_ha))

## removes the redundant "SowYear" variable
mod_frame1<-mod_frame1[,-c(grep("METADom_Sow_Year", colnames(mod_frame1)))]

## remove a couple sites with no sowdate
mod_frame1<-mod_frame1[!is.na(mod_frame1$BOMDom_Sowing_date_numeric),]

## corrects some variables marked as phenotypes, such as row number per
plot, and places them into management
mod_frame1$MANDom_Rows_per_Plot<-
as.numeric(as.character(mod_frame1$PHENDom_Rows_per_Plot))
mod_frame1$MANDom_Fungicide_flutriafol_infert_presowing<-
mod_frame1$PHENDom_Fungicide_flutriafol_infert_presowing

mod_frame1<-mod_frame1[,-c(grep("PHENDom_Rows_per_Plot",
colnames(mod_frame1)),

grep("PHENDom_Fungicide_flutriafol_infert_presowing",
colnames(mod_frame1)))]

## fixes rows per plot to integers <=10, dummy-codes
mod_frame1$MANDom_Rows_per_Plot[!(mod_frame1$MANDom_Rows_per_Plot %in%
c(1:10))]<-c("UNK")

Rows_per_plot_<-factor(as.factor(mod_frame1$MANDom_Rows_per_Plot),
                 levels =
levels(as.factor(mod_frame1$MANDom_Rows_per_Plot))[c(1,3:10,2,11)])
Rows_per_plot_<-model.matrix(~Rows_per_plot_)
Rows_per_plot_<-Rows_per_plot_[,-c(1)]

colnames(Rows_per_plot_)<-paste0("MANDom_",colnames(Rows_per_plot_))

Fungicide_flutriafol_infert_presowing<-
as.character(mod_frame1$MANDom_Fungicide_flutriafol_infert_presowing)
Fungicide_flutriafol_infert_presowing[Fungicide_flutriafol_infert_presowing=="1"]<
-c("TRUE")
Fungicide_flutriafol_infert_presowing[Fungicide_flutriafol_infert_presowing=="0"]<
-c("FALSE")
Fungicide_flutriafol_infert_presowing[is.na(Fungicide_flutriafol_infert_presowing)]<
-c("UNK")

Fungicide_flutriafol_infert_presowing<-
factor(as.factor(Fungicide_flutriafol_infert_presowing),
                       levels =
levels(as.factor(Fungicide_flutriafol_infert_presowing))[c(3,1,2)])
Fungicide_flutriafol_infert_presowing<-
model.matrix(~Fungicide_flutriafol_infert_presowing)
Fungicide_flutriafol_infert_presowing<-
Fungicide_flutriafol_infert_presowing[,-c(1)]
colnames(Fungicide_flutriafol_infert_presowing)<-
paste0("MANDom_",colnames(Fungicide_flutriafol_infert_presowing))

## adds rows per plot, dummy-coded
mod_frame1<-data.frame(mod_frame1,
                       Rows_per_plot_,
                       Fungicide_flutriafol_infert_presowing)

mod_frame1<-mod_frame1[,-c(grep("MANDom_Rows_per_Plot$",
colnames(mod_frame1)))]
mod_frame1<-mod_frame1[,-
c(grep("MANDom_Fungicide_flutriafol_infert_presowing$",
colnames(mod_frame1)))]

rm(Fungicide_flutriafol_infert_presowing, Rows_per_plot_)


## adds the prefix to the trial series dummy-coding
colnames(mod_frame1)[grep("Trial_series", colnames(mod_frame1))]<-
paste0("METADom_",

c(colnames(mod_frame1)[grep("Trial_series", colnames(mod_frame1))]))
```

```
## removes the EnvDom intercept variable
mod_frame1<-mod_frame1[,-c(which(colnames(mod_frame1)=="X.Intercept."))]

## dummy-codes the species
sub_crop<-as.factor(Phen_data$PHENDom_Crop_alldata)
sub_crop<-model.matrix(~sub_crop)
sub_crop<-sub_crop[,-c(1)]
#colnames(sub_crop)[1]<-c("Crop_intercept")
colnames(sub_crop)<-paste0("METADom_",colnames(sub_crop))

## adds species, dummy-coded
mod_frame1<-data.frame(mod_frame1,
                       sub_crop)

## dummy codes crop rotations minus 0-6
sub_crop<-as.factor(mod_frame1$METADom_Crop_rotation_minus_0)
sub_crop<-model.matrix(~sub_crop)
sub_crop<-sub_crop[,-c(1)]
#colnames(sub_crop)[1]<-c("Crop_intercept")
colnames(sub_crop)<-
paste0("METADom_Crop_rotation_minus_0_",colnames(sub_crop))

## adds species, dummy-coded
mod_frame1<-data.frame(mod_frame1,
                       sub_crop)

## dummy codes crop rotations minus 0-6
sub_crop<-as.factor(mod_frame1$METADom_Crop_rotation_minus_1)
sub_crop<-model.matrix(~sub_crop)
sub_crop<-sub_crop[,-c(1)]
#colnames(sub_crop)[1]<-c("Crop_intercept")
colnames(sub_crop)<-
paste0("METADom_Crop_rotation_minus_1_",colnames(sub_crop))

## adds species, dummy-coded
mod_frame1<-data.frame(mod_frame1,
                       sub_crop)

## dummy codes crop rotations minus 1 and 2
sub_crop<-as.factor(mod_frame1$METADom_Crop_rotation_minus_2)
sub_crop<-model.matrix(~sub_crop)
sub_crop<-sub_crop[,-c(1)]
#colnames(sub_crop)[1]<-c("Crop_intercept")
colnames(sub_crop)<-
paste0("METADom_Crop_rotation_minus_2_",colnames(sub_crop))

## adds species, dummy-coded
mod_frame1<-data.frame(mod_frame1,
                       sub_crop)

## dummy codes crop rotations minus 1 and 2
sub_crop<-as.factor(mod_frame1$METADom_Crop_rotation_minus_3)
sub_crop<-model.matrix(~sub_crop)
sub_crop<-sub_crop[,-c(1)]
#colnames(sub_crop)[1]<-c("Crop_intercept")
colnames(sub_crop)<-
paste0("METADom_Crop_rotation_minus_3_",colnames(sub_crop))

## adds species, dummy-coded
mod_frame1<-data.frame(mod_frame1,
                       sub_crop)

## dummy codes crop rotations minus 1 and 2
sub_crop<-as.factor(mod_frame1$METADom_Crop_rotation_minus_4)
sub_crop<-model.matrix(~sub_crop)
sub_crop<-sub_crop[,-c(1)]
#colnames(sub_crop)[1]<-c("Crop_intercept")
colnames(sub_crop)<-
paste0("METADom_Crop_rotation_minus_4_",colnames(sub_crop))

## adds species, dummy-coded
mod_frame1<-data.frame(mod_frame1,
                       sub_crop)

## dummy codes crop rotations minus 1 and 2
sub_crop<-as.factor(mod_frame1$METADom_Crop_rotation_minus_5)
sub_crop<-model.matrix(~sub_crop)
sub_crop<-sub_crop[,-c(1)]
#colnames(sub_crop)[1]<-c("Crop_intercept")
colnames(sub_crop)<-
paste0("METADom_Crop_rotation_minus_5_",colnames(sub_crop))

## adds species, dummy-coded
mod_frame1<-data.frame(mod_frame1,
                       sub_crop)

sub_crop<-as.factor(mod_frame1$METADom_Crop_rotation_minus_6)
sub_crop<-model.matrix(~sub_crop)
sub_crop<-sub_crop[,-c(1)]
#colnames(sub_crop)[1]<-c("Crop_intercept")
colnames(sub_crop)<-
paste0("METADom_Crop_rotation_minus_6_",colnames(sub_crop))
```

```
## adds species, dummy-coded
mod_frame1<-data.frame(mod_frame1,
                       sub_crop)

sub_crop<-as.factor(mod_frame1$MANDom_Orientation)
sub_crop<-model.matrix(~sub_crop)
sub_crop<-sub_crop[,-c(1)]
#colnames(sub_crop)[1]<-c("Crop_intercept")
colnames(sub_crop)<-paste0("MANDom_Orientation_",colnames(sub_crop))

## adds orientation, dummy-coded
mod_frame1<-data.frame(mod_frame1,
                       sub_crop)


sub_crop<-as.factor(mod_frame1$METADom_previous_crop_same)
sub_crop<-model.matrix(~sub_crop)
sub_crop<-sub_crop[,-c(1)]
#colnames(sub_crop)[1]<-c("Crop_intercept")
colnames(sub_crop)<-
paste0("METADom_previous_crop_same_",colnames(sub_crop))

## adds species, dummy-coded
mod_frame1<-data.frame(mod_frame1,
                       sub_crop)

## removes the original factor crop rotations
mod_frame1<-mod_frame1[,-c(grep("METADom_Crop_rotation_minus_0$",
colnames(mod_frame1)))]
mod_frame1<-mod_frame1[,-c(grep("METADom_Crop_rotation_minus_1$",
colnames(mod_frame1)))]
mod_frame1<-mod_frame1[,-c(grep("METADom_Crop_rotation_minus_2$",
colnames(mod_frame1)))]
mod_frame1<-mod_frame1[,-c(grep("METADom_Crop_rotation_minus_3$",
colnames(mod_frame1)))]
mod_frame1<-mod_frame1[,-c(grep("METADom_Crop_rotation_minus_4$",
colnames(mod_frame1)))]
mod_frame1<-mod_frame1[,-c(grep("METADom_Crop_rotation_minus_5$",
colnames(mod_frame1)))]
mod_frame1<-mod_frame1[,-c(grep("METADom_Crop_rotation_minus_6$",
colnames(mod_frame1)))]

## removes the original factor orientation
mod_frame1<-mod_frame1[,-c(grep("MANDom_Orientation$",
colnames(mod_frame1)))]
## removes the original factor previous crop same
mod_frame1<-mod_frame1[,-c(grep("METADom_previous_crop_same$",
colnames(mod_frame1)))]

## dummy-codes the soil test types

Soil_test_class_10cm_<-
as.factor(mod_frame1$METADom_P_test_class_10cm.test_10cm)
Soil_test_class_10cm_<-model.matrix(~Soil_test_class_10cm_)
Soil_test_class_10cm_<-Soil_test_class_10cm_[,-c(1)]
colnames(Soil_test_class_10cm_)<-
paste0("METADom_",colnames(Soil_test_class_10cm_))

Soil_test_class_60cm_<-
as.factor(mod_frame1$METADom_P_test_class_60cm.test_60cm)
## replaces "" with unmeasured
Soil_test_class_60cm_[Soil_test_class_60cm_==""]<-"UNMEASURED"
Soil_test_class_60cm_<-model.matrix(~Soil_test_class_60cm_)
Soil_test_class_60cm_<-Soil_test_class_60cm_[,-c(1)]
colnames(Soil_test_class_60cm_)<-
paste0("METADom_",colnames(Soil_test_class_60cm_))

## adds species, dummy-coded
mod_frame1<-data.frame(mod_frame1,
                       Soil_test_class_10cm_, Soil_test_class_60cm_)

mod_frame1<-mod_frame1[,-c(grep("METADom_P_test_class_10cm.test_10cm$",
colnames(mod_frame1)))]
mod_frame1<-mod_frame1[,-c(grep("METADom_P_test_class_60cm.test_60cm$",
colnames(mod_frame1)))]

rm(Soil_test_class_60cm_, Soil_test_class_10cm_, sub_crop)

## fixes the SatDom names, replacing single periods with underscores
colnames(mod_frame1)[c(grep("[a-z]\\.[0-9]{3}$", colnames(mod_frame1)))]<-
gsub("\\.", "_",colnames(mod_frame1)[c(grep("[a-z]\\.[0-9]{3}$",
colnames(mod_frame1)))])
colnames(mod_frame1)[c(grep("[a-z]\\.[0-9]{2}$", colnames(mod_frame1)))]<-
gsub("\\.", "_",colnames(mod_frame1)[c(grep("[a-z]\\.[0-9]{2}$",
colnames(mod_frame1)))])

colnames(mod_frame1)[c(grep("[A-Z]\\.[0-9]{3}$", colnames(mod_frame1)))]<-
gsub("\\.", "_",colnames(mod_frame1)[c(grep("[A-Z]\\.[0-9]{3}$",
colnames(mod_frame1)))])
colnames(mod_frame1)[c(grep("[A-Z]\\.[0-9]{2}$", colnames(mod_frame1)))]<-
gsub("\\.", "_",colnames(mod_frame1)[c(grep("[A-Z]\\.[0-9]{2}$",
colnames(mod_frame1)))])
```

```
colnames(mod_frame1)[c(grep("[0-9]\\.[0-9]{3}$", colnames(mod_frame1)))]<-
gsub("\\.", "_",colnames(mod_frame1)[c(grep("[0-9]\\.[0-9]{3}$",
colnames(mod_frame1)))])
colnames(mod_frame1)[c(grep("[0-9]\\.[0-9]{2}$", colnames(mod_frame1)))]<-
gsub("\\.", "_",colnames(mod_frame1)[c(grep("[0-9]\\.[0-9]{2}$",
colnames(mod_frame1)))])

## fixes/collates some phenotype distributions
Flowering<-mod_frame1$PHENDom_Flowering_50pct
## screens out 'scores'
Flowering[Flowering<=10]<-NA
## as most (but not all) flowering time data are given as Julian Days,
## checks whether flowering happened after sowing
Flowering<-c(Flowering-mod_frame1$MANDom_JTS)
## selects 30 days as an arbitrary cutoff
Flowering[Flowering<30]<-NA
## attaches QC'd data
mod_frame1$PHENDom_Flowering_QC<-Flowering

## fixes Glucosinolates
mod_frame1$PHENDom_Glucosinolates<-
as.numeric(as.character(mod_frame1$PHENDom_Glucosinolate_2))

## Saves the resulting data frame
write.csv(mod_frame1, file =
"model_frames_and_imputations/merged_variety_data_impute1.csv", row.names =
F)

## saves an RDS for faster loading to memory
saveRDS(mod_frame1, file = "model_frames_and_imputations/mod_frame1.rds")

## Rescales the data frame, re-writes

## first, separates phenotypes
Phen_data<-mod_frame1[,grep("PHENDom", colnames(mod_frame1))]
mod_frame2<-mod_frame1[,-c(grep("PHENDom", colnames(mod_frame1)))]

## removes invariant columns
mod_frame2<-mod_frame2[,which(apply(mod_frame2, 2, var, na.rm=TRUE)!=0)]

## rescales
mod_frame2<-as.matrix(mod_frame2)

## loops to prevent vector memory limits being exhausted
for(i in 1:ncol(mod_frame2)){
  mod_frame2[,i]<-scale(mod_frame2[,i])
}
#mod_frame2<-scale(mod_frame2)

## re-combines these data frames
mod_frame2<-data.frame(Phen_data,mod_frame2)

write.csv(mod_frame2, file =
"model_frames_and_imputations/merged_variety_data_rescaled_impute1.csv",
row.names = F)

## saves an RDS for faster loading to memory
saveRDS(mod_frame2, file =
"model_frames_and_imputations/mod_frame1_rescaled.rds")

rm(sub_frame,sub_Miss,Phen_data, mod_frame2)


###########################################################################
#

## Constructs ML models across all targets for variety-site-year data
frames

###########################################################################
#

predobs_frame_unscaled<-NULL
predobs_frame_scaled<-NULL

## registers cluster
peanutCluster<-makeCluster(detectCores()-2)
registerDoParallel(peanutCluster)

## loads data
mod_frame1<-readRDS("model_frames_and_imputations/mod_frame1.rds")

## separates phenotypes
Phen_data<-mod_frame1[,grep("PHENDom", colnames(mod_frame1))]
mod_frame1<-mod_frame1[,-c(grep("PHENDom", colnames(mod_frame1)))]

## attaches the target phenotype
mod_frame1$Yield<-Phen_data$PHENDom_yield_t_ha

## removes na data
mod_frame_Y<-mod_frame1[!is.na(mod_frame1$Yield),]

## removes "TRIAL ID" vectors (used for crossvalidation)
```

```
mod_frame_Y<-mod_frame_Y[,-c(grep("Trial_ID", colnames(mod_frame_Y),
ignore.case = T))]

## splits off 2018 data
test_frame_Y<-mod_frame_Y[mod_frame_Y$METADom_Year>=2018,]
mod_frame_Y<-mod_frame_Y[mod_frame_Y$METADom_Year<2018,]

Site_index_vec<-paste0(mod_frame_Y$BOMDom_Longitude,
mod_frame_Y$BOMDom_Latitude, mod_frame_Y$BOMDom_Sowing_date_numeric)

## subsamples 100 experiments from the pre-2018 data
set.seed(8394)
Site_subset<-sample(Site_index_vec, 100)
test_frame_Y_site<-mod_frame_Y[c(Site_index_vec %in% Site_subset),]
mod_frame_Y<-mod_frame_Y[!(Site_index_vec %in% Site_subset),]

## keeps track of which species is attached to which prediction, for later
Species_test_df1<-data.frame(test_frame_Y[,grep("METADom_sub_crop",
colnames(test_frame_Y))])

require(ranger)
system.time(Rrf_mod<-ranger(Yield~., mod_frame_Y, seed = 8394, num.trees =
10000))
detach(package:ranger)

pred_Rrf<-predict(Rrf_mod, data = test_frame_Y)
Rrf_prediction<-pred_Rrf$predictions

pred_Rrf_site<-predict(Rrf_mod, data = test_frame_Y_site)

tmp_pred<-predict(Rrf_mod, data = mod_frame_Y)
## keeps internal accuracy
predobs_frame_unscaled[[1]]<-data.frame(Yield = mod_frame_Y$Yield, Rrf_200
=  tmp_pred$predictions)
### and forward-projection accuracy
predobs_frame_unscaled[[2]]<-data.frame(Yield = test_frame_Y$Yield, Rrf_200
= pred_Rrf$predictions)
### and missing-site accuracy
predobs_frame_unscaled[[3]]<-data.frame(Yield = test_frame_Y_site$Yield,
Rrf_200 = pred_Rrf_site$predictions)
rm(tmp_pred)

## constructs rpart model
system.time(rpart_mod<-rpart(Yield~., mod_frame_Y, control =
rpart.control(cp = 0.0001, minbucket = 100, minsplit = 300)))
rpart_prediction<-predict(rpart_mod, newdata = test_frame_Y)
cor.test(rpart_prediction,test_frame_Y$Yield)

## keeps internal accuracy
predobs_frame_unscaled[[1]]<-data.frame(predobs_frame_unscaled[[1]], rp_200
=  predict(rpart_mod))
### and forward-projection accuracy
predobs_frame_unscaled[[2]]<-data.frame(predobs_frame_unscaled[[2]], rp_200
=  predict(rpart_mod, newdata = test_frame_Y))
### and missing-site accuracy
predobs_frame_unscaled[[3]]<-data.frame(predobs_frame_unscaled[[3]], rp_200
= predict(rpart_mod, newdata = test_frame_Y_site))

## re-loads random forest to fix/overwrite namespace
require(randomForest)

## builds another bcrf model, cross-validated by unique year
clusterSetRNGStream(cl = peanutCluster, 7745)
set.seed(8394)
system.time(rf_mod<-randomForest(Yield~., mod_frame_Y,
                                  strata = "METADom_Year",
                                  xval = 10, minbucket = 100, minsplit =
300, mtry = 100, ntree = 1001, sampsize = 10000))
beep()

rf_prediction<-predict(rf_mod, newdata = test_frame_Y)
cor.test(predict(rf_mod, newdata = test_frame_Y_site),
test_frame_Y_site$Yield)
cor.test(rf_prediction, test_frame_Y$Yield)

## keeps internal accuracy
predobs_frame_unscaled[[1]]<-data.frame(predobs_frame_unscaled[[1]],
xvrf_200 =  predict(rf_mod))
### and forward-projection accuracy
predobs_frame_unscaled[[2]]<-data.frame(predobs_frame_unscaled[[2]],
xvrf_200 =  predict(rf_mod, newdata = test_frame_Y))
### and missing-site accuracy
predobs_frame_unscaled[[3]]<-data.frame(predobs_frame_unscaled[[3]],
xvrf_200 = predict(rf_mod, newdata = test_frame_Y_site))

## Saves models
saveRDS(rpart_mod,file =
"model_frames_and_imputations/siteVariety_models/rpart_allTimes.rds")
saveRDS(Rrf_mod,file =
"model_frames_and_imputations/siteVariety_models/Rrf_allTimes.rds")
saveRDS(rf_mod,file =
"model_frames_and_imputations/siteVariety_models/bcrf_allTimes.rds")
```

```
## ensures no carry-over
rm(rf_mod, Rrf_mod, rpart_mod)

####################################################

## Runs these models again, using data to 100 DAS, Day of Sowing

####################################################

## removes na data,"TRIAL ID" vectors (used for crossvalidation), adds
species
mod_frame_Y<-mod_frame1[!is.na(mod_frame1$Yield),]
mod_frame_Y<-mod_frame_Y[,-c(grep("Trial_ID", colnames(mod_frame_Y),
ignore.case = T))]
#mod_frame_Y<-data.frame(mod_frame_Y,
#                         Crop =
as.factor(P_frame$PHENDom_Crop[!is.na(P_frame$PHENDom_MEAN.YIELD)]))

## Filters data
mod_frame_Y<-mod_frame_Y[,-c(grep("2[0-9]{2}$", colnames(mod_frame_Y)))]
mod_frame_Y<-mod_frame_Y[,-c(grep("1[0-9]{2}$", colnames(mod_frame_Y)))]
mod_frame_Y<-mod_frame_Y[,-c(grep("PostSowing", colnames(mod_frame_Y),
ignore.case = T))]

## converts all chemical, fertiliser and soil tests with dates >=100DAS
## to zero dose values
Chem_subvector<-
unlist(lapply(strsplit(colnames(mod_frame_Y)[grep("Chemicals_time_Chemicals_"
,

colnames(mod_frame_Y))], split = "time_Chemicals_"),tail,1))

for(i in 1:length(grep("Chemicals_time_Chemicals_",
colnames(mod_frame_Y)))){
  ## picks appropriate column name
  sub_index<-grep(c(colnames(mod_frame_Y)[grep(paste0(Chem_subvector[i],
"$"),
                                               colnames(mod_frame_Y))])[1],
colnames(mod_frame_Y))
  ## replaces doses after 100DAS with zero
  sub_vec<-mod_frame_Y[,c(sub_index[[1]])]
  sub_vec[mod_frame_Y[,c(grep("Chemicals_time_Chemicals_",
colnames(mod_frame_Y))[i])]>=100]<-0
  mod_frame_Y[,sub_index]<-sub_vec
  rm(sub_vec, sub_index)
  print(i)
}

## replaces the timings themselves with NAs
for(i in grep("Chemicals_time_Chemicals_", colnames(mod_frame_Y))){
  mod_frame_Y[which(mod_frame_Y[,i]>=100),i]<-0
}

Fert_subvector<-
unlist(lapply(strsplit(colnames(mod_frame_Y)[grep("METADom_Fertiliser_time_"
,

colnames(mod_frame_Y))], split = "Fertiliser_time_"),tail,1))

for(i in 1:length(grep("METADom_Fertiliser_time_",
colnames(mod_frame_Y)))){
  ## picks appropriate column name
  sub_index<-grep(c(colnames(mod_frame_Y)[grep(paste0(Fert_subvector[i],
"$"),
                                               colnames(mod_frame_Y))])[1],
colnames(mod_frame_Y))
  ## replaces doses after 100DAS with zero
  sub_vec<-mod_frame_Y[,c(sub_index[[1]])]
  sub_vec[mod_frame_Y[,c(grep("Fertiliser_time_",
colnames(mod_frame_Y))[i])]>=100]<-0
  mod_frame_Y[,sub_index]<-sub_vec
  rm(sub_vec, sub_index)
}

## replaces the timings themselves with NAs
for(i in grep("Fertiliser_time_", colnames(mod_frame_Y))){
  mod_frame_Y[which(mod_frame_Y[,i]>=100),i]<-0
}

## splits off 2018 data
test_frame_Y<-mod_frame_Y[mod_frame_Y$METADom_Year>=2018,]
mod_frame_Y<-mod_frame_Y[mod_frame_Y$METADom_Year<2018,]

## subsamples 100 experiments from the pre-2018 data
test_frame_Y_site<-mod_frame_Y[c(Site_index_vec %in% Site_subset),]
mod_frame_Y<-mod_frame_Y[!(Site_index_vec %in% Site_subset),]

##  adjusts the columns in test_frame to match
#test_frame_Y<-test_frame_Y[,c(colnames(test_frame_Y) %in%
colnames(mod_frame_Y))]

##########
```

```
require(ranger)
system.time(Rrf_mod<-ranger(Yield~., mod_frame_Y, seed = 8394, num.trees =
10000))
detach(package:ranger)

pred_Rrf<-predict(Rrf_mod, data = test_frame_Y)
Rrf_prediction_100<-pred_Rrf$predictions

pred_Rrf_site<-predict(Rrf_mod, data = test_frame_Y_site)

tmp_pred<-predict(Rrf_mod, data = mod_frame_Y)
## keeps internal accuracy
predobs_frame_unscaled[[1]]<-data.frame(predobs_frame_unscaled[[1]],
Rrf_100 =  tmp_pred$predictions)
### and forward-projection accuracy
predobs_frame_unscaled[[2]]<-data.frame(predobs_frame_unscaled[[2]],
Rrf_100 = pred_Rrf$predictions)
### and missing-site accuracy
predobs_frame_unscaled[[3]]<-data.frame(predobs_frame_unscaled[[3]],
Rrf_100 = pred_Rrf_site$predictions)
rm(tmp_pred)

## constructs rpart model
system.time(rpart_mod<-rpart(Yield~., mod_frame_Y, control =
rpart.control(cp = 0.0001, minbucket = 100, minsplit = 300)))
rpart_prediction_100<-predict(rpart_mod, newdata = test_frame_Y)

cor.test(rpart_prediction_100,test_frame_Y$Yield)

## keeps internal accuracy
predobs_frame_unscaled[[1]]<-data.frame(predobs_frame_unscaled[[1]], rp_100
=  predict(rpart_mod))
### and forward-projection accuracy
predobs_frame_unscaled[[2]]<-data.frame(predobs_frame_unscaled[[2]], rp_100
=  predict(rpart_mod, newdata = test_frame_Y))
### and missing-site accuracy
predobs_frame_unscaled[[3]]<-data.frame(predobs_frame_unscaled[[3]], rp_100
= predict(rpart_mod, newdata = test_frame_Y_site))

## re-loads random forest to fix/overwrite namespace
require(randomForest)

## builds another bcrf model, cross-validated by unique year
## keeps ntree odd to break ties
clusterSetRNGStream(cl = peanutCluster, 7745)
set.seed(8394)
system.time(rf_mod<-randomForest(Yield~., mod_frame_Y,
                                 strata = "METADom_Year",
                                 xval = 10, minbucket = 100, minsplit =
300, mtry = 100, ntree = 1001, sampsize = 10000))
beep()

rf_prediction_100<-predict(rf_mod, newdata = test_frame_Y)
cor.test(rf_prediction_100, test_frame_Y$Yield)

## keeps internal accuracy
predobs_frame_unscaled[[1]]<-data.frame(predobs_frame_unscaled[[1]],
xvrf_100 =  predict(rf_mod))
### and forward-projection accuracy
predobs_frame_unscaled[[2]]<-data.frame(predobs_frame_unscaled[[2]],
xvrf_100 =  predict(rf_mod, newdata = test_frame_Y))
### and missing-site accuracy
predobs_frame_unscaled[[3]]<-data.frame(predobs_frame_unscaled[[3]],
xvrf_100 = predict(rf_mod, newdata = test_frame_Y_site))

## Saves models
saveRDS(rpart_mod,file =
"model_frames_and_imputations/siteVariety_models/rpart_100DAS.rds")
saveRDS(Rrf_mod,file =
"model_frames_and_imputations/siteVariety_models/Rrf_100DAS.rds")
saveRDS(rf_mod,file =
"model_frames_and_imputations/siteVariety_models/bcrf_100DAS.rds")

rm(rf_mod, Rrf_mod, rpart_mod)

######################################################

## Prediction at Time of Sowing (TOS)

######################################################

## removes na data,"TRIAL ID" vectors (used for crossvalidation), adds
species
mod_frame_Y<-mod_frame1[!is.na(mod_frame1$Yield),]
mod_frame_Y<-mod_frame_Y[,-c(grep("Trial_ID", colnames(mod_frame_Y),
ignore.case = T))]

#mod_frame_Y<-data.frame(mod_frame_Y,
#                        Crop =
as.factor(P_frame$PHENDom_Crop[!is.na(P_frame$PHENDom_MEAN.YIELD)]))

## Filters data
mod_frame_Y<-mod_frame_Y[,-c(grep("_[0-9]{3}$", colnames(mod_frame_Y)))]
```

```
mod_frame_Y<-mod_frame_Y[,-c(grep("_[0-9]{2}$", colnames(mod_frame_Y)))]

mod_frame_Y<-mod_frame_Y[,-c(grep("PostSowing", colnames(mod_frame_Y),
ignore.case = T))]

## converts all chemical, fertiliser and soil tests with dates >=0
## to zero dose values
Chem_subvector<-
unlist(lapply(strsplit(colnames(mod_frame_Y)[grep("Chemicals_time_Chemicals_"
,

colnames(mod_frame_Y))], split = "time_Chemicals_"),tail,1))

for(i in 1:length(grep("Chemicals_time_Chemicals_",
colnames(mod_frame_Y)))){
  ## picks appropriate column name
  sub_index<-grep(c(colnames(mod_frame_Y)[grep(paste0(Chem_subvector[i],
"$"),
                                               colnames(mod_frame_Y))])[1],
colnames(mod_frame_Y))
  ## replaces doses after 100DAS with zero
  sub_vec<-mod_frame_Y[,c(sub_index[[1]])]
  sub_vec[mod_frame_Y[,c(grep("Chemicals_time_Chemicals_",
colnames(mod_frame_Y))[i])]>=0]<-0
  mod_frame_Y[,sub_index]<-sub_vec
  rm(sub_vec, sub_index)
  print(i)
}

## replaces the timings themselves with NAs
for(i in grep("Chemicals_time_Chemicals_", colnames(mod_frame_Y))){
  mod_frame_Y[which(mod_frame_Y[,i]>=0),i]<-0
}

Fert_subvector<-
unlist(lapply(strsplit(colnames(mod_frame_Y)[grep("METADom_Fertiliser_time_"
,

colnames(mod_frame_Y))], split = "Fertiliser_time_"),tail,1))

for(i in 1:length(grep("METADom_Fertiliser_time_",
colnames(mod_frame_Y)))){
  ## picks appropriate column name
  sub_index<-grep(c(colnames(mod_frame_Y)[grep(paste0(Fert_subvector[i],
"$"),
                                               colnames(mod_frame_Y))])[1],
colnames(mod_frame_Y))
  ## replaces doses after 100DAS with zero
  sub_vec<-mod_frame_Y[,c(sub_index[[1]])]
  sub_vec[mod_frame_Y[,c(grep("Fertiliser_time_",
colnames(mod_frame_Y))[i])]>=0]<-0
  mod_frame_Y[,sub_index]<-sub_vec
  rm(sub_vec, sub_index)
}

## replaces the timings themselves with NAs
for(i in grep("Fertiliser_time_", colnames(mod_frame_Y))){
  mod_frame_Y[which(mod_frame_Y[,i]>=0),i]<-0
}
##  adjusts the columns in test_frame to match
#test_frame_Y<-test_frame_Y[,c(colnames(test_frame_Y) %in%
colnames(mod_frame_Y))]

## splits off 2018 data
test_frame_Y<-mod_frame_Y[mod_frame_Y$METADom_Year>=2018,]
mod_frame_Y<-mod_frame_Y[mod_frame_Y$METADom_Year<2018,]

## subsamples 100 experiments from the pre-2018 data
test_frame_Y_site<-mod_frame_Y[c(Site_index_vec %in% Site_subset),]
mod_frame_Y<-mod_frame_Y[!(Site_index_vec %in% Site_subset),]

####################################################################

require(ranger)
system.time(Rrf_mod<-ranger(Yield~., mod_frame_Y, seed = 8394, num.trees =
10000))
detach(package:ranger)

pred_Rrf<-predict(Rrf_mod, data = test_frame_Y)
Rrf_prediction_TOS<-pred_Rrf$predictions

pred_Rrf_site<-predict(Rrf_mod, data = test_frame_Y_site)

tmp_pred<-predict(Rrf_mod, data = mod_frame_Y)
## keeps internal accuracy
predobs_frame_unscaled[[1]]<-data.frame(predobs_frame_unscaled[[1]],
Rrf_TOS =  tmp_pred$predictions)
### and forward-projection accuracy
predobs_frame_unscaled[[2]]<-data.frame(predobs_frame_unscaled[[2]],
Rrf_TOS = pred_Rrf$predictions)
### and missing-site accuracy
predobs_frame_unscaled[[3]]<-data.frame(predobs_frame_unscaled[[3]],
Rrf_TOS = pred_Rrf_site$predictions)
```

```
rm(tmp_pred)

## constructs rpart model
system.time(rpart_mod<-rpart(Yield~., mod_frame_Y, control =
rpart.control(cp = 0.0001, minbucket = 100, minsplit = 300)))
rpart_prediction_TOS<-predict(rpart_mod, newdata = test_frame_Y)

cor.test(rpart_prediction_TOS,test_frame_Y$Yield)

## keeps internal accuracy
predobs_frame_unscaled[[1]]<-data.frame(predobs_frame_unscaled[[1]], rp_TOS
=  predict(rpart_mod))
### and forward-projection accuracy
predobs_frame_unscaled[[2]]<-data.frame(predobs_frame_unscaled[[2]], rp_TOS
=  predict(rpart_mod, newdata = test_frame_Y))
### and missing-site accuracy
predobs_frame_unscaled[[3]]<-data.frame(predobs_frame_unscaled[[3]], rp_TOS
= predict(rpart_mod, newdata = test_frame_Y_site))

## re-loads random forest to fix/overwrite namespace
require(randomForest)

## builds another bcrf model, cross-validated by unique year
clusterSetRNGStream(cl = peanutCluster, 7745)
set.seed(8394)
system.time(rf_mod<-randomForest(Yield~., mod_frame_Y,
                                 strata = "METADom_Year",
                                 xval = 10, minbucket = 100, minsplit =
300, mtry = 100, ntree = 1001, sampsize = 10000))
beep()

rf_prediction_TOS<-predict(rf_mod, newdata = test_frame_Y)
cor.test(rf_prediction_TOS, test_frame_Y$Yield)

## keeps internal accuracy
predobs_frame_unscaled[[1]]<-data.frame(predobs_frame_unscaled[[1]],
xvrf_TOS =  predict(rf_mod))
### and forward-projection accuracy
predobs_frame_unscaled[[2]]<-data.frame(predobs_frame_unscaled[[2]],
xvrf_TOS =  predict(rf_mod, newdata = test_frame_Y))
### and missing-site accuracy
predobs_frame_unscaled[[3]]<-data.frame(predobs_frame_unscaled[[3]],
xvrf_TOS = predict(rf_mod, newdata = test_frame_Y_site))

## Saves models
saveRDS(rpart_mod,file =
"model_frames_and_imputations/siteVariety_models/rpart_TOS.rds")
saveRDS(Rrf_mod,file =
"model_frames_and_imputations/siteVariety_models/Rrf_TOS.rds")
saveRDS(rf_mod,file =
"model_frames_and_imputations/siteVariety_models/bcrf_TOS.rds")

saveRDS(predobs_frame_unscaled, file =
"model_frames_and_imputations/siteVariety_models/accuracy_predobs_alldata1.rds"
)

rm(rf_mod, Rrf_mod, rpart_mod)

########################################################################

## to allow filtering of the (rescaled) chemical and fertiliser timings,
## pulls down the non-rescaled versions of these variables

mod_frame1<-readRDS("model_frames_and_imputations/mod_frame1.rds")

ChemFert_times<-mod_frame1[,c(grep("Chemicals_time_Chemicals_",
colnames(mod_frame1)),
              grep("METADom_Fertiliser_time_", colnames(mod_frame1)))]

## returns the rescaled data
#mod_frame1<-read.csv(file =
"model_frames_and_imputations/merged_variety_data_rescaled_impute1.csv",
header = T)
mod_frame1<-readRDS("model_frames_and_imputations/mod_frame1_rescaled.rds")

## separates phenotypes
Phen_data<-mod_frame1[,grep("PHENDom", colnames(mod_frame1))]
mod_frame1<-mod_frame1[,-c(grep("PHENDom", colnames(mod_frame1)))]

## attaches the phenotype
mod_frame1$Yield<-Phen_data$PHENDom_yield_t_ha

## filters the chemical/fertiliser times
ChemFert_times<-ChemFert_times[!is.na(mod_frame1$Yield),]

## removes na data
mod_frame_Y<-mod_frame1[!is.na(mod_frame1$Yield),]

## splits off 2018 data (rescaled remember)
test_frame_Y<-mod_frame_Y[c(mod_frame_Y$METADom_Year %in%
tail(unique(mod_frame_Y$METADom_Year),2)),]
mod_frame_Y<-mod_frame_Y[!(mod_frame_Y$METADom_Year %in%
tail(unique(mod_frame_Y$METADom_Year),2)),]
```

```
## once again, subsets 100 random holdout sites
Site_index_vec<-paste0(mod_frame_Y$BOMDom_Longitude,
mod_frame_Y$BOMDom_Latitude, mod_frame_Y$BOMDom_Sowing_date_numeric)

## subsamples 100 experiments from the pre-2018 data
set.seed(8394)
Site_subset<-sample(Site_index_vec, 100)
test_frame_Y_site<-mod_frame_Y[c(Site_index_vec %in% Site_subset),]
mod_frame_Y<-mod_frame_Y[!(Site_index_vec %in% Site_subset),]

## keeps track of which species is attached to which prediction, for later
Species_test_df2<-data.frame(test_frame_Y[,grep("METADom_sub_crop",
colnames(test_frame_Y))])

test_Phen<-test_frame_Y$Yield
test_frame_Y<-as.matrix(test_frame_Y[,-
c(grep("Yield",colnames(test_frame_Y)))])

test_Phen_site<-test_frame_Y_site$Yield
test_frame_Y_site<-as.matrix(test_frame_Y_site[,-
c(grep("Yield",colnames(test_frame_Y_site)))])

mod_Phen<-mod_frame_Y$Yield
mod_frame_Y<-as.matrix(mod_frame_Y[,-
c(grep("Yield",colnames(mod_frame_Y)))])

## Fits a linear SVM
system.time(lsvm_mod<-svm(x = mod_frame_Y, y = mod_Phen, #scale = F,
threads = c(detectCores()-4),
                          grid_choice = 2,
                          folds = 10,
                          threads  = -1, random_seed = 8394))

lsvm_prediction<-predict(lsvm_mod, newdata = test_frame_Y)

lsvm_prediction2<-predict(lsvm_mod, newdata = test_frame_Y_site)
cor.test(lsvm_prediction, test_Phen)

## keeps internal accuracy
predobs_frame_scaled[[1]]<-data.frame(Yield = mod_Phen, lsvm_200 =
predict(lsvm_mod, newdata = mod_frame_Y))
### and forward-projection accuracy
predobs_frame_scaled[[2]]<-data.frame(Yield = test_Phen, lsvm_200 =
lsvm_prediction)
### and missing site accuracy
predobs_frame_scaled[[3]]<-data.frame(Yield = test_Phen_site, lsvm_200 =
lsvm_prediction2)

## Fits an xgBoost model
mod_frame3<-xgb.DMatrix(mod_frame_Y, label= mod_Phen)

set.seed(8394)
system.time(xg_mod<-xgboost(data = mod_frame3, nthread = 16, nrounds  = 30,
save_name = "xg_sitewise"))

xg_prediction<-predict(xg_mod, newdata = test_frame_Y)

xg_prediction2<-predict(xg_mod, newdata = test_frame_Y_site)
cor.test(xg_prediction, test_Phen)

## keeps internal accuracy
predobs_frame_scaled[[1]]<-data.frame(predobs_frame_scaled[[1]], xg_200 =
predict(xg_mod, newdata = mod_frame_Y))
### and forward-projection accuracy
predobs_frame_scaled[[2]]<-data.frame(predobs_frame_scaled[[2]], xg_200 =
xg_prediction)
## and missing site accuracy
predobs_frame_scaled[[3]]<-data.frame(predobs_frame_scaled[[3]], xg_200 =
xg_prediction2)

## and the PLSR
mod_frame3<-data.frame(Yield = mod_Phen, mod_frame_Y)

pls.options(parallel = makeCluster(10, type = "PSOCK"))
## Builds a PLSR model, 10-fold internally cross-validated, 20 components
system.time(plsr_mod<-plsr(Yield~., ncomp = 10, data = mod_frame3,
validation = "CV", segments  =10))

## select ncomp
which.min(plsr_mod$validation$adj)
plsr_prediction<-as.vector(predict(plsr_mod, newdata = test_frame_Y, ncomp
= 10))
plsr_prediction_2<-as.vector(predict(plsr_mod, newdata = test_frame_Y_site,
ncomp = 10))

cor.test(plsr_prediction, test_Phen)

## keeps internal accuracy
predobs_frame_scaled[[1]]<-data.frame(predobs_frame_scaled[[1]], plsr_200 =
as.vector(predict(plsr_mod, newdata =  mod_frame_Y, ncomp = 10)))
### and forward-projection accuracy
predobs_frame_scaled[[2]]<-data.frame(predobs_frame_scaled[[2]], plsr_200 =
```

```
plsr_prediction)
## and missing site accuracy
predobs_frame_scaled[[3]]<-data.frame(predobs_frame_scaled[[3]], plsr_200 =
plsr_prediction_2)

stopCluster(pls.options()$parallel)

# rpart_prediction,rf_prediction, xg_prediction,
lsvm_prediction,plsr_prediction,

## Saves these models

## saves models, target data
saveRDS(xg_mod,file =
"model_frames_and_imputations/siteVariety_models/xgb_allTimes.rds")
saveRDS(lsvm_mod,file =
"model_frames_and_imputations/siteVariety_models/lsvm_allTimes.rds")
saveRDS(plsr_mod,file =
"model_frames_and_imputations/siteVariety_models/plsr_allTimes.rds")

rm(xg_mod,lsvm_mod,pslr_mod, mod_frame3)

######################################################

## Runs these models again, using data to 100 DAS, Day of Sowing

######################################################

## removes na data
mod_frame_Y<-mod_frame1[!is.na(mod_frame1$Yield),]

## Filters data
mod_frame_Y<-mod_frame_Y[,-c(grep("2[0-9]{2}$", colnames(mod_frame_Y)))]
mod_frame_Y<-mod_frame_Y[,-c(grep("1[0-9]{2}$", colnames(mod_frame_Y)))]
mod_frame_Y<-mod_frame_Y[,-c(grep("PostSowing", colnames(mod_frame_Y),
ignore.case = T))]

## converts all chemical, fertiliser and soil tests with dates >=100DAS
## to zero dose values
Chem_subvector<-
unlist(lapply(strsplit(colnames(mod_frame_Y)[grep("Chemicals_time_Chemicals_"
,

colnames(mod_frame_Y))], split = "time_Chemicals_"),tail,1))

for(i in 1:length(grep("Chemicals_time_Chemicals_",
colnames(mod_frame_Y)))){
  ## picks appropriate column name
  sub_index<-grep(c(colnames(mod_frame_Y)[grep(paste0(Chem_subvector[i],
"$"),
                                               colnames(mod_frame_Y))])[1],
colnames(mod_frame_Y))
  ## replaces doses after 100DAS with zero
  sub_vec<-mod_frame_Y[,c(sub_index[[1]])]
  ## indexes by the correct (unscaled) times

sub_vec[unlist(ChemFert_times[which(colnames(ChemFert_times)==c(colnames(mod_frame_Y)[grep("Chemicals_time_Chemicals_"
, colnames(mod_frame_Y))[i]]))])>=100]<-0
  #sub_vec[mod_frame_Y[,c(grep("Chemicals_time_Chemicals_",
colnames(mod_frame_Y))[i])]>=100]<-0
  mod_frame_Y[,sub_index]<-sub_vec
  rm(sub_vec, sub_index)
  print(i)
}

## replaces the timings themselves with NAs
for(i in grep("Chemicals_time_Chemicals_", colnames(mod_frame_Y))){
  mod_frame_Y[which(mod_frame_Y[,i]>=100),i]<-0
}

Fert_subvector<-
unlist(lapply(strsplit(colnames(mod_frame_Y)[grep("METADom_Fertiliser_time_"
,

colnames(mod_frame_Y))], split = "Fertiliser_time_"),tail,1))

for(i in 1:length(grep("METADom_Fertiliser_time_",
colnames(mod_frame_Y)))){
  ## picks appropriate column name
  sub_index<-grep(c(colnames(mod_frame_Y)[grep(paste0(Fert_subvector[i],
"$"),
                                               colnames(mod_frame_Y))])[1],
colnames(mod_frame_Y))
  ## replaces doses after 100DAS with zero
  sub_vec<-mod_frame_Y[,c(sub_index[[1]])]

sub_vec[unlist(ChemFert_times[which(colnames(ChemFert_times)==c(colnames(mod_frame_Y)[grep("Fertiliser_time_"
, colnames(mod_frame_Y))[i]]))])>=100]<-0

 # sub_vec[mod_frame_Y[,c(grep("Fertiliser_time_",
colnames(mod_frame_Y))[i])]>=100]<-0
  mod_frame_Y[,sub_index]<-sub_vec
  rm(sub_vec, sub_index)
```

```
}

## replaces the timings themselves with NAs
for(i in grep("Fertiliser_time_", colnames(mod_frame_Y))){
  mod_frame_Y[which(mod_frame_Y[,i]>=100),i]<-0
}

## splits off 2018 data (rescaled remember)
test_frame_Y<-mod_frame_Y[c(mod_frame_Y$METADom_Year %in%
tail(unique(mod_frame_Y$METADom_Year),2)),]
mod_frame_Y<-mod_frame_Y[!(mod_frame_Y$METADom_Year %in%
tail(unique(mod_frame_Y$METADom_Year),2)),]

## splits off random sites
test_frame_Y_site<-mod_frame_Y[c(Site_index_vec %in% Site_subset),]
mod_frame_Y<-mod_frame_Y[!(Site_index_vec %in% Site_subset),]

test_Phen_site<-test_frame_Y_site$Yield
test_frame_Y_site<-as.matrix(test_frame_Y_site[,-
c(grep("Yield",colnames(test_frame_Y_site)))])

test_Phen<-test_frame_Y$Yield
test_frame_Y<-as.matrix(test_frame_Y[,-
c(grep("Yield",colnames(test_frame_Y)))])

mod_Phen<-mod_frame_Y$Yield
mod_frame_Y<-as.matrix(mod_frame_Y[,-
c(grep("Yield",colnames(mod_frame_Y)))])

## Fits a linear SVM
system.time(lsvm_mod<-svm(x = mod_frame_Y, y = mod_Phen, #scale = F,
                          grid_choice = 2,
                          folds = 10,
                          threads  = -1, random_seed = 8394))

lsvm_prediction_100<-predict(lsvm_mod, newdata = test_frame_Y)
lsvm_prediction_100_2<-predict(lsvm_mod, newdata = test_frame_Y_site)

## keeps internal accuracy
predobs_frame_scaled[[1]]<-data.frame(predobs_frame_scaled[[1]], lsvm_100 =
predict(lsvm_mod, newdata = mod_frame_Y))
### and forward-projection accuracy
predobs_frame_scaled[[2]]<-data.frame(predobs_frame_scaled[[2]], lsvm_100 =
lsvm_prediction_100)
### and forward-projection accuracy
predobs_frame_scaled[[3]]<-data.frame(predobs_frame_scaled[[3]], lsvm_100 =
lsvm_prediction_100_2)


## Fits an xgBoost model
mod_frame3<-xgb.DMatrix(mod_frame_Y, label= mod_Phen)

set.seed(8394)
system.time(xg_mod<-xgboost(data = mod_frame3, nthread = 16, nrounds  =10,
save_name = "xg_sitewise"))

xg_prediction_100<-predict(xg_mod, newdata = test_frame_Y)
xg_prediction_100_2<-predict(xg_mod, newdata = test_frame_Y_site)
cor.test(xg_prediction_100, test_Phen)

## keeps internal accuracy
predobs_frame_scaled[[1]]<-data.frame(predobs_frame_scaled[[1]], xg_100 =
predict(xg_mod, newdata = mod_frame_Y))
### and forward-projection accuracy
predobs_frame_scaled[[2]]<-data.frame(predobs_frame_scaled[[2]], xg_100 =
xg_prediction_100)
### and forward-projection accuracy
predobs_frame_scaled[[3]]<-data.frame(predobs_frame_scaled[[3]], xg_100 =
xg_prediction_100_2)

## and the PLSR
mod_frame3<-data.frame(Yield = mod_Phen, mod_frame_Y)

pls.options(parallel = makeCluster(10, type = "PSOCK"))
## Builds a PLSR model, 10-fold internally cross-validated, 20 components
system.time(pslr_mod<-plsr(Yield~., ncomp = 10, data = mod_frame3,
validation = "CV", segments  =10))
plsr_prediction_100<-as.vector(predict(pslr_mod, newdata = test_frame_Y,
ncomp = 10))
plsr_prediction_100_2<-as.vector(predict(pslr_mod, newdata =
test_frame_Y_site, ncomp = 10))

## keeps internal accuracy
predobs_frame_scaled[[1]]<-data.frame(predobs_frame_scaled[[1]], plsr_100 =
as.vector(predict(pslr_mod, newdata =  mod_frame_Y, ncomp = 10)))
### and forward-projection accuracy
predobs_frame_scaled[[2]]<-data.frame(predobs_frame_scaled[[2]], plsr_100 =
plsr_prediction_100)
## and missing site accuracy
predobs_frame_scaled[[3]]<-data.frame(predobs_frame_scaled[[3]], plsr_100 =
plsr_prediction_100_2)

stopCluster(pls.options()$parallel)
```

```
## saves models
saveRDS(xg_mod,file =
"model_frames_and_imputations/siteVariety_models/xgb_100DAS.rds")
saveRDS(lsvm_mod,file =
"model_frames_and_imputations/siteVariety_models/lsvm_100DAS.rds")
saveRDS(pslr_mod,file =
"model_frames_and_imputations/siteVariety_models/plsr_100DAS.rds")

rm(xg_mod,lsvm_mod,pslr_mod, mod_frame3)

#####################################################

## Runs these models again, using TOS data

#####################################################

## removes na data
mod_frame_Y<-mod_frame1[!is.na(mod_frame1$Yield),]

# Filters data
mod_frame_Y<-mod_frame_Y[,-c(grep("_[0-9]{3}$", colnames(mod_frame_Y)))]
mod_frame_Y<-mod_frame_Y[,-c(grep("_[0-9]{2}$", colnames(mod_frame_Y)))]
mod_frame_Y<-mod_frame_Y[,-c(grep("_[0-9]{1}$", colnames(mod_frame_Y)))]

mod_frame_Y<-mod_frame_Y[,-c(grep("PostSowing", colnames(mod_frame_Y),
ignore.case = T))]


## converts all chemical, fertiliser and soil tests with dates >=100DAS
## to zero dose values
Chem_subvector<-
unlist(lapply(strsplit(colnames(mod_frame_Y)[grep("Chemicals_time_Chemicals_"
,

colnames(mod_frame_Y))], split = "time_Chemicals_"),tail,1))

for(i in 1:length(grep("Chemicals_time_Chemicals_",
colnames(mod_frame_Y)))){
  ## picks appropriate column name
  sub_index<-grep(c(colnames(mod_frame_Y)[grep(paste0(Chem_subvector[i],
"$"),
                                               colnames(mod_frame_Y))])[1],
colnames(mod_frame_Y))
  ## replaces doses after 100DAS with zero
  sub_vec<-mod_frame_Y[,c(sub_index[[1]])]
  ## indexes by the correct (unscaled) times

sub_vec[unlist(ChemFert_times[which(colnames(ChemFert_times)==c(colnames(mod_frame_Y)[grep("Chemicals_time_Chemicals_"
, colnames(mod_frame_Y))[i]]))])>=0]<-0
  #sub_vec[mod_frame_Y[,c(grep("Chemicals_time_Chemicals_",
colnames(mod_frame_Y))[i])]>=100]<-0
  mod_frame_Y[,sub_index]<-sub_vec
  rm(sub_vec, sub_index)
  print(i)
}

## replaces the timings themselves with NAs
for(i in grep("Chemicals_time_Chemicals_", colnames(mod_frame_Y))){
  mod_frame_Y[which(mod_frame_Y[,i]>=0),i]<-0
}

Fert_subvector<-
unlist(lapply(strsplit(colnames(mod_frame_Y)[grep("METADom_Fertiliser_time_"
,

colnames(mod_frame_Y))], split = "Fertiliser_time_"),tail,1))

for(i in 1:length(grep("METADom_Fertiliser_time_",
colnames(mod_frame_Y)))){
  ## picks appropriate column name
  sub_index<-grep(c(colnames(mod_frame_Y)[grep(paste0(Fert_subvector[i],
"$"),
                                               colnames(mod_frame_Y))])[1],
colnames(mod_frame_Y))
  ## replaces doses after 100DAS with zero
  sub_vec<-mod_frame_Y[,c(sub_index[[1]])]

sub_vec[unlist(ChemFert_times[which(colnames(ChemFert_times)==c(colnames(mod_frame_Y)[grep("Fertiliser_time_"
, colnames(mod_frame_Y))[i]]))])>0]<-0
  mod_frame_Y[,sub_index]<-sub_vec
  rm(sub_vec, sub_index)
}

## replaces the timings themselves with NAs
for(i in grep("Fertiliser_time_", colnames(mod_frame_Y))){
  mod_frame_Y[which(mod_frame_Y[,i]>=0),i]<-0
}

## splits off 2018 data (rescaled remember)
test_frame_Y<-mod_frame_Y[c(mod_frame_Y$METADom_Year %in%
tail(unique(mod_frame_Y$METADom_Year),2)),]
mod_frame_Y<-mod_frame_Y[!(mod_frame_Y$METADom_Year %in%
```

```
tail(unique(mod_frame_Y$METADom_Year),2)),]

## splits off random sites
test_frame_Y_site<-mod_frame_Y[c(Site_index_vec %in% Site_subset),]
mod_frame_Y<-mod_frame_Y[!(Site_index_vec %in% Site_subset),]

test_Phen_site<-test_frame_Y_site$Yield
test_frame_Y_site<-as.matrix(test_frame_Y_site[,-
c(grep("Yield",colnames(test_frame_Y_site)))])

test_Phen<-test_frame_Y$Yield
test_frame_Y<-as.matrix(test_frame_Y[,-
c(grep("Yield",colnames(test_frame_Y)))])

mod_Phen<-mod_frame_Y$Yield
mod_frame_Y<-as.matrix(mod_frame_Y[,-
c(grep("Yield",colnames(mod_frame_Y)))])

## Fits a linear SVM
system.time(lsvm_mod<-svm(x = mod_frame_Y, y = mod_Phen, #scale = F,
                          grid_choice = 2,
                          folds = 10,
                          #max_gamma = 25,
                          threads  =-1, random_seed = 8394))

lsvm_prediction_TOS<-predict(lsvm_mod, newdata = test_frame_Y)
lsvm_prediction_TOS_2<-predict(lsvm_mod, newdata = test_frame_Y_site)
cor.test(lsvm_prediction_TOS, test_Phen)

## keeps internal accuracy
predobs_frame_scaled[[1]]<-data.frame(predobs_frame_scaled[[1]], lsvm_TOS =
predict(lsvm_mod, newdata = mod_frame_Y))
### and forward-projection accuracy
predobs_frame_scaled[[2]]<-data.frame(predobs_frame_scaled[[2]], lsvm_TOS =
lsvm_prediction_TOS)
## and missing site accuracy
predobs_frame_scaled[[3]]<-data.frame(predobs_frame_scaled[[3]], lsvm_TOS =
lsvm_prediction_TOS_2)

## Fits an xgBoost model
mod_frame3<-xgb.DMatrix(mod_frame_Y, label= mod_Phen)

set.seed(8394)
system.time(xg_mod<-xgboost(data = mod_frame3, nthread = 16, nrounds  =10,
save_name = "xg_sitewise"))

xg_prediction_TOS<-predict(xg_mod, newdata = test_frame_Y)
xg_prediction_TOS_2<-predict(xg_mod, newdata = test_frame_Y_site)
cor.test(xg_prediction_TOS, test_Phen)

## keeps internal accuracy
predobs_frame_scaled[[1]]<-data.frame(predobs_frame_scaled[[1]], xg_TOS =
predict(xg_mod, newdata = mod_frame_Y))
### and forward-projection accuracy
predobs_frame_scaled[[2]]<-data.frame(predobs_frame_scaled[[2]], xg_TOS =
xg_prediction_TOS)
## and missing site accuracy
predobs_frame_scaled[[3]]<-data.frame(predobs_frame_scaled[[3]], xg_TOS =
xg_prediction_TOS_2)

## and the PLSR
mod_frame3<-data.frame(Yield = mod_Phen, mod_frame_Y)

pls.options(parallel = makeCluster(10, type = "PSOCK"))
## Builds a PLSR model, 10-fold internally cross-validated, 20 components
system.time(pslr_mod<-plsr(Yield~., ncomp = 10, data = mod_frame3,
validation = "CV", segments  =10))

plsr_prediction_TOS<-as.vector(predict(pslr_mod, newdata = test_frame_Y,
ncomp = 10))
plsr_prediction_TOS_2<-as.vector(predict(pslr_mod, newdata =
test_frame_Y_site, ncomp = 10))
cor.test(plsr_prediction_TOS, test_Phen)

## keeps internal accuracy
predobs_frame_scaled[[1]]<-data.frame(predobs_frame_scaled[[1]], plsr_TOS =
as.vector(predict(pslr_mod, newdata =  mod_frame_Y, ncomp = 10)))
### and forward-projection accuracy
predobs_frame_scaled[[2]]<-data.frame(predobs_frame_scaled[[2]], plsr_TOS =
plsr_prediction_TOS)
## and missing site accuracy
predobs_frame_scaled[[3]]<-data.frame(predobs_frame_scaled[[3]], plsr_TOS =
plsr_prediction_TOS_2)

stopCluster(pls.options()$parallel)

## saves models, target data
saveRDS(xg_mod,file =
"model_frames_and_imputations/siteVariety_models/xgb_TOS.rds")
saveRDS(lsvm_mod,file =
"model_frames_and_imputations/siteVariety_models/lsvm_TOS.rds")
saveRDS(pslr_mod,file =
"model_frames_and_imputations/siteVariety_models/plsr_TOS.rds")
```

```
saveRDS(predobs_frame_scaled, file =
"model_frames_and_imputations/siteVariety_models/accuracy_predobs_alldata2.rds"
)


## tidy up the workspace
rm(rpart_prediction,rf_prediction,
Rrf_prediction,xg_prediction,lsvm_prediction, plsr_prediction,
     rpart_prediction_100,rf_prediction_100,Rrf_prediction_100,
xg_prediction_100,lsvm_prediction_100, plsr_prediction_100,
     rpart_prediction_TOS, rf_prediction_TOS, Rrf_prediction_TOS,
xg_prediction_TOS, lsvm_prediction_TOS, plsr_prediction_TOS)

rm(xg_mod,lsvm_mod, plsr_mod, rpart_mod, xg_mod, rf_mod, Rrf_mod)
rm(mod_frame3)

gc()
## now moves on to targeted models, ignores plsr, xgb, ranger RF.

###############################################################################
#

## constructs a rolling forecast from wheat and canola data,
## relative to TOS.

###############################################################################
#

## to allow filtering of the (rescaled) chemical and fertiliser timings,
## pulls down the non-rescaled versions of these variables

mod_frame1<-readRDS("model_frames_and_imputations/mod_frame1.rds")

## writes out a small data frame for Zhen He
write.csv(mod_frame1[1:1000,], file = "Zhen_data.csv", row.names = F)
## and another with lat-longs and flowering/emergence for brendan
mod_frame_X<-data.frame(mod_frame1$BOMDom_Sowing_date_numeric,
                        mod_frame1$BOMDom_Latitude,
                        mod_frame1$BOMDom_Longitude,
                        mod_frame1$METADom_sub_cropCanola,
                        mod_frame1$METADom_sub_cropWheat,
                        mod_frame1$METADom_sub_cropField.Pea,
                        mod_frame1$METADom_sub_cropLupin,
                        mod_frame1$METADom_sub_cropChickpea,
                        mod_frame1$METADom_sub_cropFaba.Bean,
                        mod_frame1$PHENDom_Flowering_QC,
                        mod_frame1$PHENDom_Establishment,
                        mod_frame1$PHENDom_Early_Growth)

write.csv(mod_frame_X, file = "Brendan_data.csv", row.names = F)

ChemFert_times<-mod_frame1[,c(grep("Chemicals_time_Chemicals_",
colnames(mod_frame1)),
                              grep("METADom_Fertiliser_time_",
colnames(mod_frame1)))]

## loads rescaled data...
#mod_frame1<-read.csv(file =
"model_frames_and_imputations/merged_variety_data_rescaled_impute1.csv",
header = T)
mod_frame1<-readRDS("model_frames_and_imputations/mod_frame1_rescaled.rds")

## separates phenotypes
Phen_data<-mod_frame1[,grep("PHENDom", colnames(mod_frame1))]
mod_frame1<-mod_frame1[,-c(grep("PHENDom", colnames(mod_frame1)))]

## attaches the phenotype
mod_frame1$Yield<-Phen_data$PHENDom_yield_t_ha

## filters the chemical/fertiliser times
ChemFert_times<-ChemFert_times[!is.na(mod_frame1$Yield),]

## removes na data
mod_frame_Y<-mod_frame1[!is.na(mod_frame1$Yield),]

## filters out +200DAS and postsowing data
mod_frame_Y<-mod_frame_Y[,-c(grep("_2[0-9]{2}$", colnames(mod_frame_Y)))]
mod_frame_Y<-mod_frame_Y[,-c(grep("PostSowing", colnames(mod_frame_Y),
ignore.case = T))]

mod_frame_Y2<-mod_frame_Y

## builds rolling forecasts, keeps just the accuracy.
forecast_frame_W<-NULL
forecast_frame_C<-NULL

for(j in c(1:19)){

  mod_frame_Y<-mod_frame_Y2

  # Filters data
  for(x in j:19){
```

```
    mod_frame_Y<-mod_frame_Y[,-c(grep(paste0("_",x,"[0-9]$"),
colnames(mod_frame_Y)))]
  }


  ## converts all chemical, fertiliser and soil tests with dates >=jth DAS
  ## to zero dose values
  Chem_subvector<-
unlist(lapply(strsplit(colnames(mod_frame_Y)[grep("Chemicals_time_Chemicals_"
,

colnames(mod_frame_Y))], split = "time_Chemicals_"),tail,1))

  for(i in 1:length(grep("Chemicals_time_Chemicals_",
colnames(mod_frame_Y)))){
    ## picks appropriate column name
    sub_index<-grep(c(colnames(mod_frame_Y)[grep(paste0(Chem_subvector[i],
"$"),

colnames(mod_frame_Y))])[1], colnames(mod_frame_Y))
    ## replaces doses after 100DAS with zero
    sub_vec<-mod_frame_Y[,c(sub_index[[1]])]
    ## indexes by the correct (unscaled) times

sub_vec[unlist(ChemFert_times[which(colnames(ChemFert_times)==c(colnames(mod_frame_Y)[grep("Chemicals_time_Chemicals_"
, colnames(mod_frame_Y))[i]]))])>=c(j*10)]<-0
    #sub_vec[mod_frame_Y[,c(grep("Chemicals_time_Chemicals_",
colnames(mod_frame_Y))[i])]>=100]<-0
    mod_frame_Y[,sub_index]<-sub_vec
    rm(sub_vec, sub_index)
  }

  ## replaces the timings themselves with NAs
  for(i in grep("Chemicals_time_Chemicals_", colnames(mod_frame_Y))){
    mod_frame_Y[which(mod_frame_Y[,i]>=c(j*10)),i]<-0
  }

  Fert_subvector<-
unlist(lapply(strsplit(colnames(mod_frame_Y)[grep("METADom_Fertiliser_time_"
,

colnames(mod_frame_Y))], split = "Fertiliser_time_"),tail,1))

  for(i in 1:length(grep("METADom_Fertiliser_time_",
colnames(mod_frame_Y)))){
    ## picks appropriate column name
    sub_index<-grep(c(colnames(mod_frame_Y)[grep(paste0(Fert_subvector[i],
"$"),

colnames(mod_frame_Y))])[1], colnames(mod_frame_Y))
    ## replaces doses after 100DAS with zero
    sub_vec<-mod_frame_Y[,c(sub_index[[1]])]

sub_vec[unlist(ChemFert_times[which(colnames(ChemFert_times)==c(colnames(mod_frame_Y)[grep("Fertiliser_time_"
, colnames(mod_frame_Y))[i]]))])>=c(j*10)]<-0
    mod_frame_Y[,sub_index]<-sub_vec
    rm(sub_vec, sub_index)
  }

  ## replaces the timings themselves with NAs
  for(i in grep("Fertiliser_time_", colnames(mod_frame_Y))){
    mod_frame_Y[which(mod_frame_Y[,i]>=c(j*10)),i]<-0
  }

  ## splits off 2018 data, which has been rescaled
  test_frame_Y<-
mod_frame_Y[mod_frame_Y$METADom_Year>=c(tail(unique(mod_frame_Y$METADom_Year)[order(unique(mod_frame_Y$METADom_Year))],2)[1]),
]
  mod_frame_Y<-mod_frame_Y[mod_frame_Y$METADom_Year<-
mod_frame_Y[mod_frame_Y$METADom_sub_cropWheat==max(mod_frame_Y$METADom_sub_cropWheat),
]
  test_frame_W<-
test_frame_Y[test_frame_Y$METADom_sub_cropWheat==max(test_frame_Y$METADom_sub_cropWheat),
]

  mod_frame_C<-
mod_frame_Y[mod_frame_Y$METADom_sub_cropCanola==max(mod_frame_Y$METADom_sub_cropCanola),
]
  test_frame_C<-
test_frame_Y[test_frame_Y$METADom_sub_cropCanola==max(test_frame_Y$METADom_sub_cropCanola),
]

  rm(mod_frame_Y,test_frame_Y)

  ## builds a naive bcrf model


#  set.seed(8394)
#  system.time(rf_mod<-randomForest(Yield~., mod_frame_W,
#                                   strata = "METADom_Year",
#                                   xval = 10, minbucket = 100, minsplit =
150, mtry = 75))
#  beep()
```

```
 # rf_prediction<-predict(rf_mod, newdata = test_frame_W)
  require(ranger)
  system.time(rf_mod<-ranger(Yield~., mod_frame_W, seed = 8394, num.trees =
1001))
  detach(package:ranger)

  rf_prediction<-predict(rf_mod, data = test_frame_W)

  test_Phen<-test_frame_W$Yield
  mod_Phen<-mod_frame_W$Yield

  mod_frame_W<-as.matrix(mod_frame_W[,-
c(grep("Yield",colnames(mod_frame_W)))])
  test_frame_W<-as.matrix(test_frame_W[,-
c(grep("Yield",colnames(test_frame_W)))])

  ## Fits a linear SVM
  system.time(lsvm_mod<-svm(x = mod_frame_W, y = mod_Phen, #scale = F,
                            grid_choice = 2,
                            threads  =-1, random_seed = 8394))
  lsvm_prediction<-predict(lsvm_mod, newdata = test_frame_W)

  cor.test(lsvm_prediction, test_Phen)

  ## and the PLSR
  mod_frame3<-data.frame(Yield = mod_Phen, mod_frame_W)

  pls.options(parallel = makeCluster(10, type = "PSOCK"))
  ## Builds a PLSR model, 10-fold internally cross-validated, 20 components
  system.time(pslr_mod<-plsr(Yield~., ncomp = 10, data = mod_frame3,
validation = "CV", segments  =10))

  plsr_prediction<-as.vector(predict(pslr_mod, newdata = test_frame_W,
ncomp = 10))
  cor.test(plsr_prediction, test_Phen)

  stopCluster(pls.options()$parallel)

  forecast_frame_W<-rbind(forecast_frame_W,
                      c(cor.test(rf_prediction$predictions,
test_Phen)$estimate,
                      cor.test(lsvm_prediction, test_Phen)$estimate,
                      cor.test(plsr_prediction, test_Phen)$estimate))

  print(paste0(j, "wheat"))
  rm(rf_mod, lsvm_mod, pslr_mod, rf_prediction,lsvm_prediction,
plsr_prediction,mod_frame3)


  require(ranger)
  system.time(rf_mod<-ranger(Yield~., mod_frame_C, seed = 8394, num.trees =
1001))
  detach(package:ranger)

  rf_prediction<-predict(rf_mod, data = test_frame_C)

  test_Phen<-test_frame_C$Yield
  mod_Phen<-mod_frame_C$Yield

  mod_frame_C<-as.matrix(mod_frame_C[,-
c(grep("Yield",colnames(mod_frame_C)))])
  test_frame_C<-as.matrix(test_frame_C[,-
c(grep("Yield",colnames(test_frame_C)))])

  ## Fits a linear SVM
  system.time(lsvm_mod<-svm(x = mod_frame_C, y = mod_Phen, #scale = F,
                            grid_choice = 2,
                            threads  =-1, random_seed = 8394))

  lsvm_prediction<-predict(lsvm_mod, newdata = test_frame_C)

  cor.test(lsvm_prediction, test_Phen)

  ## and the PLSR
  mod_frame3<-data.frame(Yield = mod_Phen, mod_frame_C)

  pls.options(parallel = makeCluster(10, type = "PSOCK"))
  ## Builds a PLSR model, 10-fold internally cross-validated, 20 components
  system.time(pslr_mod<-plsr(Yield~., ncomp = 10, data = mod_frame3,
validation = "CV", segments  =10))

  plsr_prediction<-as.vector(predict(pslr_mod, newdata = test_frame_C,
ncomp = 10))
  cor.test(plsr_prediction, test_Phen)

  stopCluster(pls.options()$parallel)

  forecast_frame_C<-rbind(forecast_frame_C,
                          c(cor.test(rf_prediction$predictions,
test_Phen)$estimate,
                            cor.test(lsvm_prediction, test_Phen)$estimate,
                            cor.test(plsr_prediction, test_Phen)$estimate))
```

```
  print(paste0(j, "canola"))

  rm(rf_mod, lsvm_mod, pslr_mod, rf_prediction,lsvm_prediction,
plsr_prediction,mod_frame3)
  gc()

}

## Saves these results
saveRDS(forecast_frame_W,file =
"model_frames_and_imputations/siteVariety_models/rolling_Wheat_PredObs.rds"
)
saveRDS(forecast_frame_C,file =
"model_frames_and_imputations/siteVariety_models/rolling_Canola_PredObs.rds"
)


## compares to the accuracy of wheat yield in 2018, from the omnibus models
predobs_frame_unscaled<-readRDS(file =
"model_frames_and_imputations/siteVariety_models/accuracy_predobs_alldata1.rds"
)
predobs_frame_scaled<-readRDS(file =
"model_frames_and_imputations/siteVariety_models/accuracy_predobs_alldata2.rds"
)

## Partitions the prediction accuracy by species


## Accuracy of the 100DAS forecast, BCRF, wheat and canola only
cor.test(predobs_frame_unscaled[[2]]$Yield[Species_test_df1$METADom_sub_cropWheat==1]
,

predobs_frame_unscaled[[2]]$Rrf_100[Species_test_df1$METADom_sub_cropWheat==1])$estimat
e

cor.test(predobs_frame_scaled[[2]]$Yield[Species_test_df2$METADom_sub_cropWheat>0]
,

predobs_frame_scaled[[2]]$lsvm_100[c(Species_test_df2$METADom_sub_cropWheat>0)])$estimat
e

cor.test(predobs_frame_scaled[[2]]$Yield[Species_test_df2$METADom_sub_cropWheat>0]
,

predobs_frame_scaled[[2]]$plsr_100[Species_test_df2$METADom_sub_cropWheat>0])$estimat
e

## shows the accuracy of wheat-only and canola-only model forecasts
forecast_frame_W[10,]

## Accuracy of the 100DAS forecast, BCRF, wheat and canola only
cor.test(predobs_frame_unscaled[[2]]$Yield[Species_test_df1$METADom_sub_cropCanola==1]
,

predobs_frame_unscaled[[2]]$Rrf_100[Species_test_df1$METADom_sub_cropCanola==1])$estimat
e

cor.test(predobs_frame_scaled[[2]]$Yield[Species_test_df2$METADom_sub_cropCanola>0]
,

predobs_frame_scaled[[2]]$lsvm_100[c(Species_test_df2$METADom_sub_cropCanola>0)])$estimat
e

cor.test(predobs_frame_scaled[[2]]$Yield[Species_test_df2$METADom_sub_cropCanola>0]
,

predobs_frame_scaled[[2]]$plsr_100[Species_test_df2$METADom_sub_cropCanola>0])$estimat
e

## shows the accuracy of wheat-only and canola-only model forecasts
forecast_frame_C[10,]

## Accuracy of the TOS forecast, BCRF, wheat and canola only
cor.test(predobs_frame_unscaled[[2]]$Yield[Species_test_df1$METADom_sub_cropWheat==1]
,

predobs_frame_unscaled[[2]]$Rrf_TOS[Species_test_df1$METADom_sub_cropWheat==1])$estimat
e

cor.test(predobs_frame_scaled[[2]]$Yield[Species_test_df2$METADom_sub_cropWheat>0]
,

predobs_frame_scaled[[2]]$lsvm_TOS[c(Species_test_df2$METADom_sub_cropWheat>0)])$estimat
e

cor.test(predobs_frame_scaled[[2]]$Yield[Species_test_df2$METADom_sub_cropWheat>0]
,

predobs_frame_scaled[[2]]$plsr_TOS[Species_test_df2$METADom_sub_cropWheat>0])$estimat
e

## shows the accuracy of wheat-only and canola-only model forecasts
forecast_frame_W[1,]
```

```
## Accuracy of the 100DAS forecast, BCRF, wheat and canola only
cor.test(predobs_frame_unscaled[[2]]$Yield[Species_test_df1$METADom_sub_cropCanola==1]
,

predobs_frame_unscaled[[2]]$Rrf_TOS[Species_test_df1$METADom_sub_cropCanola==1])$estimat
e

cor.test(predobs_frame_scaled[[2]]$Yield[Species_test_df2$METADom_sub_cropCanola>0]
,

predobs_frame_scaled[[2]]$lsvm_TOS[c(Species_test_df2$METADom_sub_cropCanola>0)])$estimat
e

cor.test(predobs_frame_scaled[[2]]$Yield[Species_test_df2$METADom_sub_cropCanola>0]
,

predobs_frame_scaled[[2]]$plsr_TOS[Species_test_df2$METADom_sub_cropCanola>0])$estimat
e

## shows the accuracy of wheat-only and canola-only model forecasts
forecast_frame_C[1,]

#Species_test_df2

rm(mod_frame_C, mod_frame_W)

###############################################################################
#

## constructs a rolling forecast from wheat and canola data,
## by year

###############################################################################
#

## to allow filtering of the (rescaled) chemical and fertiliser timings,
## pulls down the non-rescaled versions of these variables

## loads rescaled data...
mod_frame1<-readRDS("model_frames_and_imputations/mod_frame1_rescaled.rds")

## separates phenotypes
Phen_data<-mod_frame1[,grep("PHENDom", colnames(mod_frame1))]
mod_frame1<-mod_frame1[,-c(grep("PHENDom", colnames(mod_frame1)))]

## attaches the phenotype
mod_frame1$Yield<-Phen_data$PHENDom_yield_t_ha

## filters the chemical/fertiliser times
ChemFert_times<-ChemFert_times[!is.na(mod_frame1$Yield),]

## removes na data
mod_frame_Y<-mod_frame1[!is.na(mod_frame1$Yield),]

## filters out +200DAS and postsowing data
mod_frame_Y<-mod_frame_Y[,-c(grep("_2[0-9]{2}$", colnames(mod_frame_Y)))]
mod_frame_Y<-mod_frame_Y[,-c(grep("PostSowing", colnames(mod_frame_Y),
ignore.case = T))]


## builds rolling forecasts, keeps just the accuracy.
Annual_forecast_frame_W<-NULL
Annual_forecast_frame_C<-NULL

for(j in c(10:1)){

  ## splits off <-
mod_frame_Y[mod_frame_Y$METADom_Year==c(tail(unique(mod_frame_Y$METADom_Year)[order(unique(mod_frame_Y$METADom_Year))],j)[1]),
]
  mod_frame_Y2<-mod_frame_Y[mod_frame_Y$METADom_Year<-
mod_frame_Y2[mod_frame_Y2$METADom_sub_cropWheat==max(mod_frame_Y2$METADom_sub_cropWheat),
]
  test_frame_W<-
test_frame_Y[test_frame_Y$METADom_sub_cropWheat==max(test_frame_Y$METADom_sub_cropWheat),
]

  mod_frame_C<-
mod_frame_Y2[mod_frame_Y2$METADom_sub_cropCanola==max(mod_frame_Y2$METADom_sub_cropCanola),
]
  test_frame_C<-
test_frame_Y[test_frame_Y$METADom_sub_cropCanola==max(test_frame_Y$METADom_sub_cropCanola),
]

  rm(mod_frame_Y2,test_frame_Y)

  Sample_sizes<-rbind(c(nrow(mod_frame_W), nrow(mod_frame_C)))

  ## builds a naive bcrf model
  require(ranger)
  system.time(rf_mod<-ranger(Yield~., mod_frame_W, seed = 8394, num.trees =
1000))
  detach(package:ranger)
```

```
  rf_prediction<-predict(rf_mod, data = test_frame_W)

  test_Phen<-test_frame_W$Yield
  mod_Phen<-mod_frame_W$Yield

  mod_frame_W<-as.matrix(mod_frame_W[,-
c(grep("Yield",colnames(mod_frame_W)))])
  test_frame_W<-as.matrix(test_frame_W[,-
c(grep("Yield",colnames(test_frame_W)))])

  ## Fits a linear SVM
  system.time(lsvm_mod<-svm(x = mod_frame_W, y = mod_Phen, #scale = F,
                            grid_choice = 2,
                            threads  =-1, random_seed = 8394))
  lsvm_prediction<-predict(lsvm_mod, newdata = test_frame_W)

  cor.test(lsvm_prediction, test_Phen)

  ## and the PLSR
  mod_frame3<-data.frame(Yield = mod_Phen, mod_frame_W)

  pls.options(parallel = makeCluster(10, type = "PSOCK"))
  ## Builds a PLSR model, 10-fold internally cross-validated, 20 components
  system.time(pslr_mod<-plsr(Yield~., ncomp = 10, data = mod_frame3,
validation = "CV", segments  =10))

  plsr_prediction<-as.vector(predict(pslr_mod, newdata = test_frame_W,
ncomp = 10))
  cor.test(plsr_prediction, test_Phen)

  stopCluster(pls.options()$parallel)

  Annual_forecast_frame_W<-rbind(Annual_forecast_frame_W,
                          c(nrow(mod_frame_W),nrow(test_frame_W),
                            cor.test(rf_prediction$predictions,
test_Phen)$estimate,

cor.test(rf_prediction$predictions, test_Phen)$conf.int,
                                                cor.test(lsvm_prediction,
test_Phen)$estimate,
                                                cor.test(lsvm_prediction,
test_Phen)$conf.int,
                                                cor.test(plsr_prediction,
test_Phen)$estimate,
                                                cor.test(plsr_prediction,
test_Phen)$conf.int))

  print(paste0(j, "wheat"))
  rm(rf_mod, lsvm_mod, pslr_mod, rf_prediction,lsvm_prediction,
plsr_prediction,mod_frame3)


  require(ranger)
  system.time(rf_mod<-ranger(Yield~., mod_frame_C, seed = 8394, num.trees =
1000))
  detach(package:ranger)

  rf_prediction<-predict(rf_mod, data = test_frame_C)

  test_Phen<-test_frame_C$Yield
  mod_Phen<-mod_frame_C$Yield

  mod_frame_C<-as.matrix(mod_frame_C[,-
c(grep("Yield",colnames(mod_frame_C)))])
  test_frame_C<-as.matrix(test_frame_C[,-
c(grep("Yield",colnames(test_frame_C)))])

  ## Fits a linear SVM
  system.time(lsvm_mod<-svm(x = mod_frame_C, y = mod_Phen, #scale = F,
                            grid_choice = 2,
                            threads  =-1, random_seed = 8394))

  lsvm_prediction<-predict(lsvm_mod, newdata = test_frame_C)

  cor.test(lsvm_prediction, test_Phen)

  ## and the PLSR
  mod_frame3<-data.frame(Yield = mod_Phen, mod_frame_C)

  pls.options(parallel = makeCluster(10, type = "PSOCK"))
  ## Builds a PLSR model, 10-fold internally cross-validated, 20 components
  system.time(pslr_mod<-plsr(Yield~., ncomp = 10, data = mod_frame3,
validation = "CV", segments  =10))

  plsr_prediction<-as.vector(predict(pslr_mod, newdata = test_frame_C,
ncomp = 10))
  cor.test(plsr_prediction, test_Phen)

  stopCluster(pls.options()$parallel)

  Annual_forecast_frame_C<-rbind(Annual_forecast_frame_C,
                                 c(nrow(mod_frame_C),nrow(test_frame_C),
```

```
                                    cor.test(rf_prediction$predictions,
test_Phen)$estimate,
                                    cor.test(rf_prediction$predictions,
test_Phen)$conf.int,
                                    cor.test(lsvm_prediction,
test_Phen)$estimate,
                                    cor.test(lsvm_prediction,
test_Phen)$conf.int,
                                    cor.test(plsr_prediction,
test_Phen)$estimate,
                                    cor.test(plsr_prediction,
test_Phen)$conf.int))
  print(paste0(j, "canola"))

  rm(rf_mod, lsvm_mod, pslr_mod, rf_prediction,lsvm_prediction,
plsr_prediction,mod_frame3)
  gc()

}

## Saves these results
saveRDS(Annual_forecast_frame_W,file =
"model_frames_and_imputations/siteVariety_models/Annual_rolling_Wheat_PredObs.rds"
)
saveRDS(Annual_forecast_frame_C,file =
"model_frames_and_imputations/siteVariety_models/Annual_rolling_Canola_PredObs.rds"
)

rm(mod_frame_C, mod_frame_W)

## constructs the same models, but with NO climate data, to assess the role
of climate stationarity in accuracy disruptions

## removes na data
mod_frame_Y<-mod_frame1[!is.na(mod_frame1$Yield),]

## filters out +200DAS and postsowing data
mod_frame_Y<-mod_frame_Y[,-c(grep("_2[0-9]{2}$", colnames(mod_frame_Y)))]
mod_frame_Y<-mod_frame_Y[,-c(grep("PostSowing", colnames(mod_frame_Y),
ignore.case = T))]
mod_frame_Y<-mod_frame_Y[,-c(grep("^SatDom_", colnames(mod_frame_Y)))]
mod_frame_Y<-mod_frame_Y[,-c(grep("^ENVDom_", colnames(mod_frame_Y)))]
mod_frame_Y<-mod_frame_Y[,-c(grep("^BOMDom_", c(colnames(mod_frame_Y)[-
c(1:3)])))]

## builds rolling forecasts, keeps just the accuracy.
Annual_forecast_frame_noE_W<-NULL
Annual_forecast_frame_noE_C<-NULL

for(j in c(10:1)){

  ## splits off <-
mod_frame_Y[mod_frame_Y$METADom_Year==c(tail(unique(mod_frame_Y$METADom_Year)[order(unique(mod_frame_Y$METADom_Year))],j)[1]),
]
  mod_frame_Y2<-mod_frame_Y[mod_frame_Y$METADom_Year<-
mod_frame_Y2[mod_frame_Y2$METADom_sub_cropWheat==max(mod_frame_Y2$METADom_sub_cropWheat),
]
  test_frame_W<-
test_frame_Y[test_frame_Y$METADom_sub_cropWheat==max(test_frame_Y$METADom_sub_cropWheat),
]

  mod_frame_C<-
mod_frame_Y2[mod_frame_Y2$METADom_sub_cropCanola==max(mod_frame_Y2$METADom_sub_cropCanola),
]
  test_frame_C<-
test_frame_Y[test_frame_Y$METADom_sub_cropCanola==max(test_frame_Y$METADom_sub_cropCanola),
]

  rm(mod_frame_Y2,test_frame_Y)

  Sample_sizes<-rbind(c(nrow(mod_frame_W), nrow(mod_frame_C)))

  ## builds a naive bcrf model
  require(ranger)
  system.time(rf_mod<-ranger(Yield~., mod_frame_W, seed = 8394, num.trees =
1000))
  detach(package:ranger)

  rf_prediction<-predict(rf_mod, data = test_frame_W)

  test_Phen<-test_frame_W$Yield
  mod_Phen<-mod_frame_W$Yield

  mod_frame_W<-as.matrix(mod_frame_W[,-
c(grep("Yield",colnames(mod_frame_W)))])
  test_frame_W<-as.matrix(test_frame_W[,-
c(grep("Yield",colnames(test_frame_W)))])

  ## Fits a linear SVM
  system.time(lsvm_mod<-svm(x = mod_frame_W, y = mod_Phen, #scale = F,
                            grid_choice = 2,
                            threads  =-1, random_seed = 8394))
  lsvm_prediction<-predict(lsvm_mod, newdata = test_frame_W)
```

```
  cor.test(lsvm_prediction, test_Phen)

  ## and the PLSR
  mod_frame3<-data.frame(Yield = mod_Phen, mod_frame_W)

  pls.options(parallel = makeCluster(10, type = "PSOCK"))
  ## Builds a PLSR model, 10-fold internally cross-validated, 20 components
  system.time(pslr_mod<-plsr(Yield~., ncomp = 10, data = mod_frame3,
validation = "CV", segments  =10))

  plsr_prediction<-as.vector(predict(pslr_mod, newdata = test_frame_W,
ncomp = 10))
  cor.test(plsr_prediction, test_Phen)

  stopCluster(pls.options()$parallel)

  Annual_forecast_frame_noE_W<-rbind(Annual_forecast_frame_noE_W,
                                c(nrow(mod_frame_W),nrow(test_frame_W),
                                  cor.test(rf_prediction$predictions,
test_Phen)$estimate,
                                  cor.test(rf_prediction$predictions,
test_Phen)$conf.int,
                                  cor.test(lsvm_prediction,
test_Phen)$estimate,
                                  cor.test(lsvm_prediction,
test_Phen)$conf.int,
                                  cor.test(plsr_prediction,
test_Phen)$estimate,
                                  cor.test(plsr_prediction,
test_Phen)$conf.int))

  print(paste0(j, "wheat"))
  rm(rf_mod, lsvm_mod, pslr_mod, rf_prediction,lsvm_prediction,
plsr_prediction,mod_frame3)


  require(ranger)
  system.time(rf_mod<-ranger(Yield~., mod_frame_C, seed = 8394, num.trees =
1000))
  detach(package:ranger)

  rf_prediction<-predict(rf_mod, data = test_frame_C)

  test_Phen<-test_frame_C$Yield
  mod_Phen<-mod_frame_C$Yield

  mod_frame_C<-as.matrix(mod_frame_C[,-
c(grep("Yield",colnames(mod_frame_C)))])
  test_frame_C<-as.matrix(test_frame_C[,-
c(grep("Yield",colnames(test_frame_C)))])

  ## Fits a linear SVM
  system.time(lsvm_mod<-svm(x = mod_frame_C, y = mod_Phen, #scale = F,
                            grid_choice = 2,
                            threads  =-1, random_seed = 8394))

  lsvm_prediction<-predict(lsvm_mod, newdata = test_frame_C)

  cor.test(lsvm_prediction, test_Phen)

  ## and the PLSR
  mod_frame3<-data.frame(Yield = mod_Phen, mod_frame_C)

  pls.options(parallel = makeCluster(10, type = "PSOCK"))
  ## Builds a PLSR model, 10-fold internally cross-validated, 20 components
  system.time(pslr_mod<-plsr(Yield~., ncomp = 10, data = mod_frame3,
validation = "CV", segments  =10))

  plsr_prediction<-as.vector(predict(pslr_mod, newdata = test_frame_C,
ncomp = 10))
  cor.test(plsr_prediction, test_Phen)

  stopCluster(pls.options()$parallel)

  Annual_forecast_frame_noE_C<-rbind(Annual_forecast_frame_noE_C,
                                c(nrow(mod_frame_C),nrow(test_frame_C),
                                  cor.test(rf_prediction$predictions,
test_Phen)$estimate,
                                  cor.test(rf_prediction$predictions,
test_Phen)$conf.int,
                                  cor.test(lsvm_prediction,
test_Phen)$estimate,
                                  cor.test(lsvm_prediction,
test_Phen)$conf.int,
                                  cor.test(plsr_prediction,
test_Phen)$estimate,
                                  cor.test(plsr_prediction,
test_Phen)$conf.int))
  print(paste0(j, "canola"))

  rm(rf_mod, lsvm_mod, pslr_mod, rf_prediction,lsvm_prediction,
plsr_prediction,mod_frame3)
```

```
  gc()

}

## Saves these results
saveRDS(Annual_forecast_frame_noE_W,file =
"model_frames_and_imputations/siteVariety_models/Annual_rolling_Wheat_noE_PredObs.rds"
)
saveRDS(Annual_forecast_frame_noE_C,file =
"model_frames_and_imputations/siteVariety_models/Annual_rolling_Canola_noE_PredObs.rds"
)

rm(mod_frame_C, mod_frame_W)

## Builds the same kind of model, but for protein (wheat) and
glucosinolates (canola)

## to allow filtering of the (rescaled) chemical and fertiliser timings,
## pulls down the non-rescaled versions of these variables

## loads rescaled data...
mod_frame1<-readRDS("model_frames_and_imputations/mod_frame1_rescaled.rds")

## separates phenotypes
Phen_data<-mod_frame1[,grep("PHENDom", colnames(mod_frame1))]
mod_frame1<-mod_frame1[,-c(grep("PHENDom", colnames(mod_frame1)))]

## attaches the phenotype
mod_frame1$Protein<-Phen_data$PHENDom_Protein

## removes na data
mod_frame_Y<-mod_frame1[!is.na(mod_frame1$Protein),]

## filters out +200DAS and postsowing data
mod_frame_Y<-mod_frame_Y[,-c(grep("_2[0-9]{2}$", colnames(mod_frame_Y)))]
mod_frame_Y<-mod_frame_Y[,-c(grep("PostSowing", colnames(mod_frame_Y),
ignore.case = T))]


## builds rolling forecasts, keeps just the accuracy.
Annual_forecast_frame_Prot_W<-NULL
Annual_forecast_frame_Prot_C<-NULL

for(j in c(7:1)){

  ## splits off <-
mod_frame_Y[mod_frame_Y$METADom_Year==c(tail(unique(mod_frame_Y$METADom_Year)[order(unique(mod_frame_Y$METADom_Year))],j)[1]),
]
  mod_frame_Y2<-mod_frame_Y[mod_frame_Y$METADom_Year<-
mod_frame_Y2[mod_frame_Y2$METADom_sub_cropWheat==max(mod_frame_Y2$METADom_sub_cropWheat),
]
  test_frame_W<-
test_frame_Y[test_frame_Y$METADom_sub_cropWheat==max(test_frame_Y$METADom_sub_cropWheat),
]

  mod_frame_C<-
mod_frame_Y2[mod_frame_Y2$METADom_sub_cropCanola==max(mod_frame_Y2$METADom_sub_cropCanola),
]
  test_frame_C<-
test_frame_Y[test_frame_Y$METADom_sub_cropCanola==max(test_frame_Y$METADom_sub_cropCanola),
]

  rm(mod_frame_Y2,test_frame_Y)

  Sample_sizes<-rbind(c(nrow(mod_frame_W), nrow(mod_frame_C)))

  ## builds a naive bcrf model
  require(ranger)
  system.time(rf_mod<-ranger(Protein~., mod_frame_W, seed = 8394, num.trees
= 1000))
  detach(package:ranger)

  rf_prediction<-predict(rf_mod, data = test_frame_W)

  test_Phen<-test_frame_W$Protein
  mod_Phen<-mod_frame_W$Protein

  mod_frame_W<-as.matrix(mod_frame_W[,-
c(grep("Protein",colnames(mod_frame_W)))])
  test_frame_W<-as.matrix(test_frame_W[,-
c(grep("Protein",colnames(test_frame_W)))])

  ## Fits a linear SVM
  system.time(lsvm_mod<-svm(x = mod_frame_W, y = mod_Phen, #scale = F,
                            grid_choice = 2,
                            threads  =-1, random_seed = 8394))
  lsvm_prediction<-predict(lsvm_mod, newdata = test_frame_W)

  cor.test(lsvm_prediction, test_Phen)

  ## and the PLSR
  mod_frame3<-data.frame(Protein = mod_Phen, mod_frame_W)
```

```
  pls.options(parallel = makeCluster(10, type = "PSOCK"))
  ## Builds a PLSR model, 10-fold internally cross-validated, 20 components
  system.time(pslr_mod<-plsr(Protein~., ncomp = 10, data = mod_frame3,
validation = "CV", segments  =10))

  plsr_prediction<-as.vector(predict(pslr_mod, newdata = test_frame_W,
ncomp = 10))
  cor.test(plsr_prediction, test_Phen)

  stopCluster(pls.options()$parallel)

  Annual_forecast_frame_Prot_W<-rbind(Annual_forecast_frame_Prot_W,
                                  c(nrow(mod_frame_W),nrow(test_frame_W),
                                    cor.test(rf_prediction$predictions,
test_Phen)$estimate,
                                    cor.test(rf_prediction$predictions,
test_Phen)$conf.int,
                                    cor.test(lsvm_prediction,
test_Phen)$estimate,
                                    cor.test(lsvm_prediction,
test_Phen)$conf.int,
                                    cor.test(plsr_prediction,
test_Phen)$estimate,
                                    cor.test(plsr_prediction,
test_Phen)$conf.int))

  print(paste0(j, "wheat"))
  rm(rf_mod, lsvm_mod, pslr_mod, rf_prediction,lsvm_prediction,
plsr_prediction,mod_frame3)


  require(ranger)
  system.time(rf_mod<-ranger(Protein~., mod_frame_C, seed = 8394, num.trees
= 1000))
  detach(package:ranger)

  rf_prediction<-predict(rf_mod, data = test_frame_C)

  test_Phen<-test_frame_C$Protein
  mod_Phen<-mod_frame_C$Protein

  mod_frame_C<-as.matrix(mod_frame_C[,-
c(grep("Protein",colnames(mod_frame_C)))])
  test_frame_C<-as.matrix(test_frame_C[,-
c(grep("Protein",colnames(test_frame_C)))])

  ## Fits a linear SVM
  system.time(lsvm_mod<-svm(x = mod_frame_C, y = mod_Phen, #scale = F,
                            grid_choice = 2,
                            threads  =-1, random_seed = 8394))

  lsvm_prediction<-predict(lsvm_mod, newdata = test_frame_C)

  cor.test(lsvm_prediction, test_Phen)

  ## and the PLSR
  mod_frame3<-data.frame(Protein = mod_Phen, mod_frame_C)

  pls.options(parallel = makeCluster(10, type = "PSOCK"))
  ## Builds a PLSR model, 10-fold internally cross-validated, 20 components
  system.time(pslr_mod<-plsr(Protein~., ncomp = 10, data = mod_frame3,
validation = "CV", segments  =10))

  plsr_prediction<-as.vector(predict(pslr_mod, newdata = test_frame_C,
ncomp = 10))
  cor.test(plsr_prediction, test_Phen)

  stopCluster(pls.options()$parallel)

  Annual_forecast_frame_Prot_C<-rbind(Annual_forecast_frame_Prot_C,
                                  c(nrow(mod_frame_C),nrow(test_frame_C),
                                    cor.test(rf_prediction$predictions,
test_Phen)$estimate,
                                    cor.test(rf_prediction$predictions,
test_Phen)$conf.int,
                                    cor.test(lsvm_prediction,
test_Phen)$estimate,
                                    cor.test(lsvm_prediction,
test_Phen)$conf.int,
                                    cor.test(plsr_prediction,
test_Phen)$estimate,
                                    cor.test(plsr_prediction,
test_Phen)$conf.int))
  print(paste0(j, "canola"))

  rm(rf_mod, lsvm_mod, pslr_mod, rf_prediction,lsvm_prediction,
plsr_prediction,mod_frame3)
  gc()

}

## Saves these results
saveRDS(Annual_forecast_frame_Prot_W,file =
```

```
"model_frames_and_imputations/siteVariety_models/Annual_rolling_protein_Wheat_PredObs.rds"
)
saveRDS(Annual_forecast_frame_Prot_C,file =
"model_frames_and_imputations/siteVariety_models/Annual_rolling_protein_Canola_PredObs.rds"
)


## loads rescaled data...
mod_frame1<-readRDS("model_frames_and_imputations/mod_frame1_rescaled.rds")

## separates phenotypes
Phen_data<-mod_frame1[,grep("PHENDom", colnames(mod_frame1))]
mod_frame1<-mod_frame1[,-c(grep("PHENDom", colnames(mod_frame1)))]

## attaches the phenotype
mod_frame1$Protein<-Phen_data$PHENDom_Protein

## removes na data
mod_frame_Y<-mod_frame1[!is.na(mod_frame1$Protein),]

## filters out +200DAS and postsowing data
mod_frame_Y<-mod_frame_Y[,-c(grep("_2[0-9]{2}$", colnames(mod_frame_Y)))]
mod_frame_Y<-mod_frame_Y[,-c(grep("PostSowing", colnames(mod_frame_Y),
ignore.case = T))]


## builds rolling forecasts, keeps just the accuracy.
Annual_forecast_frame_Prot_W<-NULL
Annual_forecast_frame_Prot_C<-NULL

for(j in c(7:1)){

  ## splits off <-
mod_frame_Y[mod_frame_Y$METADom_Year==c(tail(unique(mod_frame_Y$METADom_Year)[order(unique(mod_frame_Y$METADom_Year))],j)[1]),
]
  mod_frame_Y2<-mod_frame_Y[mod_frame_Y$METADom_Year<-
mod_frame_Y2[mod_frame_Y2$METADom_sub_cropWheat==max(mod_frame_Y2$METADom_sub_cropWheat),
]
  test_frame_W<-
test_frame_Y[test_frame_Y$METADom_sub_cropWheat==max(test_frame_Y$METADom_sub_cropWheat),
]

  mod_frame_C<-
mod_frame_Y2[mod_frame_Y2$METADom_sub_cropCanola==max(mod_frame_Y2$METADom_sub_cropCanola),
]
  test_frame_C<-
test_frame_Y[test_frame_Y$METADom_sub_cropCanola==max(test_frame_Y$METADom_sub_cropCanola),
]

  rm(mod_frame_Y2,test_frame_Y)

  Sample_sizes<-rbind(c(nrow(mod_frame_W), nrow(mod_frame_C)))

  ## builds a naive bcrf model
  require(ranger)
  system.time(rf_mod<-ranger(Protein~., mod_frame_W, seed = 8394, num.trees
= 1000))
  detach(package:ranger)

  rf_prediction<-predict(rf_mod, data = test_frame_W)

  test_Phen<-test_frame_W$Protein
  mod_Phen<-mod_frame_W$Protein

  mod_frame_W<-as.matrix(mod_frame_W[,-
c(grep("Protein",colnames(mod_frame_W)))])
  test_frame_W<-as.matrix(test_frame_W[,-
c(grep("Protein",colnames(test_frame_W)))])

  ## Fits a linear SVM
  system.time(lsvm_mod<-svm(x = mod_frame_W, y = mod_Phen, #scale = F,
                            grid_choice = 2,
                            threads  =-1, random_seed = 8394))
  lsvm_prediction<-predict(lsvm_mod, newdata = test_frame_W)

  cor.test(lsvm_prediction, test_Phen)

  ## and the PLSR
  mod_frame3<-data.frame(Protein = mod_Phen, mod_frame_W)

  pls.options(parallel = makeCluster(10, type = "PSOCK"))
  ## Builds a PLSR model, 10-fold internally cross-validated, 20 components
  system.time(pslr_mod<-plsr(Protein~., ncomp = 10, data = mod_frame3,
validation = "CV", segments  =10))

  plsr_prediction<-as.vector(predict(pslr_mod, newdata = test_frame_W,
ncomp = 10))
  cor.test(plsr_prediction, test_Phen)

  stopCluster(pls.options()$parallel)

  Annual_forecast_frame_Prot_W<-rbind(Annual_forecast_frame_Prot_W,
```

```
c(nrow(mod_frame_W),nrow(test_frame_W),
                                        cor.test(rf_prediction$predictions,
test_Phen)$estimate,
                                        cor.test(rf_prediction$predictions,
test_Phen)$conf.int,
                                        cor.test(lsvm_prediction,
test_Phen)$estimate,
                                        cor.test(lsvm_prediction,
test_Phen)$conf.int,
                                        cor.test(plsr_prediction,
test_Phen)$estimate,
                                        cor.test(plsr_prediction,
test_Phen)$conf.int))

  print(paste0(j, "wheat"))
  rm(rf_mod, lsvm_mod, pslr_mod, rf_prediction,lsvm_prediction,
plsr_prediction,mod_frame3)


  require(ranger)
  system.time(rf_mod<-ranger(Protein~., mod_frame_C, seed = 8394, num.trees
= 1000))
  detach(package:ranger)

  rf_prediction<-predict(rf_mod, data = test_frame_C)

  test_Phen<-test_frame_C$Protein
  mod_Phen<-mod_frame_C$Protein

  mod_frame_C<-as.matrix(mod_frame_C[,-
c(grep("Protein",colnames(mod_frame_C)))])
  test_frame_C<-as.matrix(test_frame_C[,-
c(grep("Protein",colnames(test_frame_C)))])

  ## Fits a linear SVM
  system.time(lsvm_mod<-svm(x = mod_frame_C, y = mod_Phen, #scale = F,
                            grid_choice = 2,
                            threads  =-1, random_seed = 8394))

  lsvm_prediction<-predict(lsvm_mod, newdata = test_frame_C)

  cor.test(lsvm_prediction, test_Phen)

  ## and the PLSR
  mod_frame3<-data.frame(Protein = mod_Phen, mod_frame_C)

  pls.options(parallel = makeCluster(10, type = "PSOCK"))
  ## Builds a PLSR model, 10-fold internally cross-validated, 20 components
  system.time(pslr_mod<-plsr(Protein~., ncomp = 10, data = mod_frame3,
validation = "CV", segments  =10))

  plsr_prediction<-as.vector(predict(pslr_mod, newdata = test_frame_C,
ncomp = 10))
  cor.test(plsr_prediction, test_Phen)

  stopCluster(pls.options()$parallel)

  Annual_forecast_frame_Prot_C<-rbind(Annual_forecast_frame_Prot_C,

c(nrow(mod_frame_C),nrow(test_frame_C),
                                        cor.test(rf_prediction$predictions,
test_Phen)$estimate,
                                        cor.test(rf_prediction$predictions,
test_Phen)$conf.int,
                                        cor.test(lsvm_prediction,
test_Phen)$estimate,
                                        cor.test(lsvm_prediction,
test_Phen)$conf.int,
                                        cor.test(plsr_prediction,
test_Phen)$estimate,
                                        cor.test(plsr_prediction,
test_Phen)$conf.int))
  print(paste0(j, "canola"))

  rm(rf_mod, lsvm_mod, pslr_mod, rf_prediction,lsvm_prediction,
plsr_prediction,mod_frame3)
  gc()

}

## Saves these results
saveRDS(Annual_forecast_frame_Prot_C,file =
"model_frames_and_imputations/siteVariety_models/Annual_rolling_protein_Glucosinolates_PredObs.rds"
)


rm(mod_frame_C,
mod_frame_W,Annual_forecast_frame_Prot_W,Annual_forecast_frame_Prot_C)

###############################################################################
#

## Generates best-case phenotype prediction models at +100DAS and +200DAS
```

```
## for key phenotypes, using only data from WITHIN a species

##############################################################################
#

## loads rescaled data for lsvms...
#mod_frame1<-read.csv(file =
"model_frames_and_imputations/merged_variety_data_rescaled_impute1.csv",
header = T)
mod_frame1<-readRDS("model_frames_and_imputations/mod_frame1_rescaled.rds")

## separates phenotypes
Phen_data<-mod_frame1[,grep("PHENDom", colnames(mod_frame1))]
mod_frame1<-mod_frame1[,-c(grep("PHENDom", colnames(mod_frame1)))]

#######################################################################

## builds yield models for each species

## filters out +200DAS and postsowing data
mod_frame_Y<-mod_frame1[,-c(grep("_2[0-9]{2}$", colnames(mod_frame1)))]
mod_frame_Y<-mod_frame_Y[,-c(grep("PostSowing", colnames(mod_frame_Y),
ignore.case = T))]

spp_list<-unique(Phen_data$PHENDom_Crop_alldata)[-c(1,9)]

Yield_predobs_200<-NULL

## builds yield models
for(i in 1:9){

  ## depending on the species, and therefore sample sizes available,
  ## partitions data by year so that the test data contain enough
observations
  ## to evaluate models (N>250)

  ## splits off 2018+ data (rescaled)
  test_phen<-
Phen_data[c(mod_frame_Y$METADom_Year>=c(tail(unique(mod_frame_Y$METADom_Year),2)[1])),
]
  mod_phen<-Phen_data[c(mod_frame_Y$METADom_Year<-
mod_frame_Y[c(mod_frame_Y$METADom_Year>=c(tail(unique(mod_frame_Y$METADom_Year),2)[1])),
]
  mod_frame_sub<-mod_frame_Y[c(mod_frame_Y$METADom_Year<-
Phen_data[c(mod_frame_Y$METADom_Year>=c(tail(unique(mod_frame_Y$METADom_Year),4)[1])),
]
    mod_phen<-Phen_data[c(mod_frame_Y$METADom_Year<-
mod_frame_Y[c(mod_frame_Y$METADom_Year>=c(tail(unique(mod_frame_Y$METADom_Year),4)[1])),
]
    mod_frame_sub<-mod_frame_Y[c(mod_frame_Y$METADom_Year<-
Phen_data[c(mod_frame_Y$METADom_Year>=c(tail(unique(mod_frame_Y$METADom_Year),3)[1])),
]
    mod_phen<-Phen_data[c(mod_frame_Y$METADom_Year<-
mod_frame_Y[c(mod_frame_Y$METADom_Year>=c(tail(unique(mod_frame_Y$METADom_Year),3)[1])),
]
    mod_frame_sub<-mod_frame_Y[c(mod_frame_Y$METADom_Year<-
Phen_data[c(mod_frame_Y$METADom_Year>=c(tail(unique(mod_frame_Y$METADom_Year),5)[1])),
]
    mod_phen<-Phen_data[c(mod_frame_Y$METADom_Year<-
mod_frame_Y[c(mod_frame_Y$METADom_Year>=c(tail(unique(mod_frame_Y$METADom_Year),5)[1])),
]
    mod_frame_sub<-mod_frame_Y[c(mod_frame_Y$METADom_Year<-
mod_frame_sub[mod_phen$PHENDom_Crop_alldata==spp_list[[i]],]
  test_frame_sub<-
test_frame_Y[test_phen$PHENDom_Crop_alldata==spp_list[[i]],]

  test_phen_sub<-test_phen[test_phen$PHENDom_Crop_alldata==spp_list[[i]],]
  mod_phen_sub<-mod_phen[mod_phen$PHENDom_Crop_alldata==spp_list[[i]],]

  ## removes NAs in the phenotype, and all corresponding rows in model test
and train data frames
  mod_frame_sub<-mod_frame_sub[!is.na(mod_phen_sub$PHENDom_yield_t_ha),]
  test_frame_sub<-test_frame_sub[!is.na(test_phen_sub$PHENDom_yield_t_ha),]

  mod_phen_sub<-mod_phen_sub[!is.na(mod_phen_sub$PHENDom_yield_t_ha),]
  test_phen_sub<-test_phen_sub[!is.na(test_phen_sub$PHENDom_yield_t_ha),]

  ## attaches Yield to model frame
  mod_frame_sub$Yield<-mod_phen_sub$PHENDom_yield_t_ha

  ## makes shorthand copies of phenotype
  mod_Phen_Y<-mod_frame_sub$Yield
  test_phen_Y<-test_phen_sub$PHENDom_yield_t_ha

  ## builds a bcrf model, cross-validated by unique year
  set.seed(8394)
  system.time(rf_mod<-randomForest(Yield~., mod_frame_sub,
                                  strata = "METADom_Year",
                                  xval = 10, minbucket = 100, minsplit =
300, mtry = 100))
  beep()

  rf_prediction<-predict(rf_mod, newdata = test_frame_sub)
```

```
  cor.test(rf_prediction, test_phen_Y)
  print(paste0("bcrf trained",i))

  ## removes the phenotype values from the model frame
  mod_frame_sub<-as.matrix(mod_frame_sub[,-
c(grep("Yield",colnames(mod_frame_sub)))])

  ## Fits a linear SVM
  system.time(lsvm_mod<-svm(x = mod_frame_sub, y = mod_Phen_Y, #scale = F,
                            grid_choice = 2,
                            #max_gamma = 25,
                            threads  =-1, random_seed = 8394))
  lsvm_prediction<-predict(lsvm_mod, newdata = test_frame_sub)

  cor.test(lsvm_prediction, test_phen_Y)
  print(paste0("lsvm trained",i))

  ## saves models, pred/obs data as list
  saveRDS(lsvm_mod, file =
paste0("model_frames_and_imputations/siteVariety_models/rf_Yield_200DAS",
                                  spp_list[[i]],".rds"))
  saveRDS(rf_mod, file =
paste0("model_frames_and_imputations/siteVariety_models/lsvm_Yield_200DAS",
                                  spp_list[[i]],".rds"))
  Yield_predobs_200[[i]]<-rbind(test_phen_Y, rf_prediction,
lsvm_prediction)


}

## filters out +100DAS and postsowing data
mod_frame_Y<-mod_frame1[,-c(grep("_[0-9]{3}$", colnames(mod_frame1)))]
mod_frame_Y<-mod_frame_Y[,-c(grep("PostSowing", colnames(mod_frame_Y),
ignore.case = T))]

spp_list<-unique(Phen_data$PHENDom_Crop_alldata)[-c(1,9)]

Yield_predobs_100<-NULL

## builds yield models
for(i in 1:9){

  ## depending on the species, and therefore sample sizes available,
  ## partitions data by year so that the test data contain enough
observations
  ## to evaluate models (N>250)

  ## splits off 2018+ data (rescaled)
  test_phen<-
Phen_data[c(mod_frame_Y$METADom_Year>=c(tail(unique(mod_frame_Y$METADom_Year),2)[1])),
]
  mod_phen<-Phen_data[c(mod_frame_Y$METADom_Year<-
mod_frame_Y[c(mod_frame_Y$METADom_Year>=c(tail(unique(mod_frame_Y$METADom_Year),2)[1])),
]
  mod_frame_sub<-mod_frame_Y[c(mod_frame_Y$METADom_Year<-
Phen_data[c(mod_frame_Y$METADom_Year>=c(tail(unique(mod_frame_Y$METADom_Year),4)[1])),
]
    mod_phen<-Phen_data[c(mod_frame_Y$METADom_Year<-
mod_frame_Y[c(mod_frame_Y$METADom_Year>=c(tail(unique(mod_frame_Y$METADom_Year),4)[1])),
]
    mod_frame_sub<-mod_frame_Y[c(mod_frame_Y$METADom_Year<-
Phen_data[c(mod_frame_Y$METADom_Year>=c(tail(unique(mod_frame_Y$METADom_Year),3)[1])),
]
    mod_phen<-Phen_data[c(mod_frame_Y$METADom_Year<-
mod_frame_Y[c(mod_frame_Y$METADom_Year>=c(tail(unique(mod_frame_Y$METADom_Year),3)[1])),
]
    mod_frame_sub<-mod_frame_Y[c(mod_frame_Y$METADom_Year<-
Phen_data[c(mod_frame_Y$METADom_Year>=c(tail(unique(mod_frame_Y$METADom_Year),5)[1])),
]
    mod_phen<-Phen_data[c(mod_frame_Y$METADom_Year<-
mod_frame_Y[c(mod_frame_Y$METADom_Year>=c(tail(unique(mod_frame_Y$METADom_Year),5)[1])),
]
    mod_frame_sub<-mod_frame_Y[c(mod_frame_Y$METADom_Year<-
mod_frame_sub[mod_phen$PHENDom_Crop_alldata==spp_list[[i]],]
  test_frame_sub<-
test_frame_Y[test_phen$PHENDom_Crop_alldata==spp_list[[i]],]

  test_phen_sub<-test_phen[test_phen$PHENDom_Crop_alldata==spp_list[[i]],]
  mod_phen_sub<-mod_phen[mod_phen$PHENDom_Crop_alldata==spp_list[[i]],]

  ## removes NAs in the phenotype, and all corresponding rows in model test
and train data frames
  mod_frame_sub<-mod_frame_sub[!is.na(mod_phen_sub$PHENDom_yield_t_ha),]
  test_frame_sub<-test_frame_sub[!is.na(test_phen_sub$PHENDom_yield_t_ha),]

  mod_phen_sub<-mod_phen_sub[!is.na(mod_phen_sub$PHENDom_yield_t_ha),]
  test_phen_sub<-test_phen_sub[!is.na(test_phen_sub$PHENDom_yield_t_ha),]

  ## attaches Yield to model frame
  mod_frame_sub$Yield<-mod_phen_sub$PHENDom_yield_t_ha

  ## makes shorthand copies of phenotype
```

```
  mod_Phen_Y<-mod_frame_sub$Yield
  test_phen_Y<-test_phen_sub$PHENDom_yield_t_ha

  ## builds a bcrf model, cross-validated by unique year
  set.seed(8394)
  system.time(rf_mod<-randomForest(Yield~., mod_frame_sub,
                                   strata = "METADom_Year",
                                   xval = 10, minbucket = 100, minsplit =
300, mtry = 100))
  beep()

  rf_prediction<-predict(rf_mod, newdata = test_frame_sub)

  cor.test(rf_prediction, test_phen_Y)
  print(paste0("bcrf trained",i))

  ## removes the phenotype values from the model frame
  mod_frame_sub<-as.matrix(mod_frame_sub[,-
c(grep("Yield",colnames(mod_frame_sub)))])

  ## Fits a linear SVM
  system.time(lsvm_mod<-svm(x = mod_frame_sub, y = mod_Phen_Y, #scale = F,
                            grid_choice = 2,
                            #max_gamma = 25,
                            threads  =-1, random_seed = 8394))
  lsvm_prediction<-predict(lsvm_mod, newdata = test_frame_sub)

  cor.test(lsvm_prediction, test_phen_Y)
  print(paste0("lsvm trained",i))

  ## saves models, pred/obs data as list
  saveRDS(lsvm_mod, file =
paste0("model_frames_and_imputations/siteVariety_models/rf_Yield_100DAS",
                                  spp_list[[i]],".rds"))
  saveRDS(rf_mod, file =
paste0("model_frames_and_imputations/siteVariety_models/lsvm_Yield_100DAS",
                                spp_list[[i]],".rds"))

  Yield_predobs_100[[i]]<-rbind(test_phen_Y, rf_prediction,
lsvm_prediction)


}

## builds (metadata) protein models for wheat, oats, triticale
table(is.na(Phen_data$PHENDom_Protein_4), by = Phen_data$PHENDom_Crop.Type)

## converts protein (from source 4) to numeric
Phen_data$PHENDom_Protein_4<-
as.numeric(as.character(Phen_data$PHENDom_Protein_4))

## filters out +200DAS and postsowing data
mod_frame_Y<-mod_frame1[,-c(grep("_2[0-9]{2}$", colnames(mod_frame1)))]
mod_frame_Y<-mod_frame_Y[,-c(grep("PostSowing", colnames(mod_frame_Y),
ignore.case = T))]

spp_list<-unique(Phen_data$PHENDom_Crop_alldata)[-c(1:5,7:9)]

Protein_predobs_200<-NULL

for(i in 1:3){

  ## removes NAs in the phenotype, and all corresponding rows in model test
and train data frames
  mod_phen<-Phen_data[!is.na(Phen_data$PHENDom_Protein_4),]
  mod_frame_sub<-mod_frame_Y[!is.na(Phen_data$PHENDom_Protein_4),]

  ## splits out the required species
  mod_frame_sub<-
mod_frame_sub[mod_phen$PHENDom_Crop_alldata==spp_list[[i]],]
  mod_phen<-mod_phen[mod_phen$PHENDom_Crop_alldata==spp_list[[i]],]

  ## depending on the species, and therefore sample sizes available,
  ## partitions data by year so that the test data contain enough
observations
  ## to evaluate models (N>250)

  ## splits off last two years
  test_phen<-
mod_phen[c(mod_frame_sub$METADom_Year>=c(tail(unique(mod_frame_sub$METADom_Year),2)[1])),
]
  mod_phen<-mod_phen[c(mod_frame_sub$METADom_Year<-
mod_frame_sub[c(mod_frame_sub$METADom_Year>=c(tail(unique(mod_frame_sub$METADom_Year),2)[1])),
]
  mod_frame_sub<-mod_frame_sub[c(mod_frame_sub$METADom_Year<-
mod_phen$PHENDom_Protein_4

  ## makes shorthand copies of phenotype
  mod_Phen_Y<-mod_frame_sub$Protein
  test_phen_Y<-test_phen$PHENDom_Protein_4

  ## builds a bcrf model, cross-validated by unique year
  set.seed(8394)
```

```
  system.time(rf_mod<-randomForest(Protein~., mod_frame_sub,
                                   strata = "METADom_Year",
                                   xval = 10, minbucket = 100, minsplit =
300, mtry = 100))
  beep()

  rf_prediction<-predict(rf_mod, newdata = test_frame_sub)

  cor.test(rf_prediction, test_phen_Y)
  print(paste0("bcrf trained",i))


  ## removes the phenotype values from the model frame
  mod_frame_sub<-as.matrix(mod_frame_sub[,-
c(grep("Protein",colnames(mod_frame_sub)))])

  ## Fits a linear SVM
  system.time(lsvm_mod<-svm(x = mod_frame_sub, y = mod_Phen_Y, #scale = F,
                            grid_choice = 2,
                            #max_gamma = 25,
                            threads  =-1, random_seed = 8394))
  lsvm_prediction<-predict(lsvm_mod, newdata = test_frame_sub)

  cor.test(lsvm_prediction, test_phen_Y)
  print(paste0("lsvm trained",i))

  ## saves models, pred/obs data as list
  saveRDS(lsvm_mod, file =
paste0("model_frames_and_imputations/siteVariety_models/rf_Protein_200DAS",
                                  spp_list[[i]],".rds"))
  saveRDS(rf_mod, file =
paste0("model_frames_and_imputations/siteVariety_models/lsvm_Protein_200DAS"
,
                                spp_list[[i]],".rds"))
  Protein_predobs_200[[i]]<-rbind(test_phen_Y, rf_prediction,
lsvm_prediction)

}


## filters out +100DAS and postsowing data
mod_frame_Y<-mod_frame1[,-c(grep("_[0-9]{3}$", colnames(mod_frame1)))]
mod_frame_Y<-mod_frame_Y[,-c(grep("PostSowing", colnames(mod_frame_Y),
ignore.case = T))]

Protein_predobs_100<-NULL

for(i in 1:3){

  ## removes NAs in the phenotype, and all corresponding rows in model test
and train data frames
  mod_phen<-Phen_data[!is.na(Phen_data$PHENDom_Protein_4),]
  mod_frame_sub<-mod_frame_Y[!is.na(Phen_data$PHENDom_Protein_4),]

  ## splits out the required species
  mod_frame_sub<-
mod_frame_sub[mod_phen$PHENDom_Crop_alldata==spp_list[[i]],]
  mod_phen<-mod_phen[mod_phen$PHENDom_Crop_alldata==spp_list[[i]],]

  ## depending on the species, and therefore sample sizes available,
  ## partitions data by year so that the test data contain enough
observations
  ## to evaluate models (N>250)

  ## splits off last two years
  test_phen<-
mod_phen[c(mod_frame_sub$METADom_Year>=c(tail(unique(mod_frame_sub$METADom_Year),2)[1])),
]
  mod_phen<-mod_phen[c(mod_frame_sub$METADom_Year<-
mod_frame_sub[c(mod_frame_sub$METADom_Year>=c(tail(unique(mod_frame_sub$METADom_Year),2)[1])),
]
  mod_frame_sub<-mod_frame_sub[c(mod_frame_sub$METADom_Year<-
mod_phen$PHENDom_Protein_4

  ## makes shorthand copies of phenotype
  mod_Phen_Y<-mod_frame_sub$Protein
  test_phen_Y<-test_phen$PHENDom_Protein_4

  ## builds a bcrf model, cross-validated by unique year
  set.seed(8394)
  system.time(rf_mod<-randomForest(Protein~., mod_frame_sub,
                                   strata = "METADom_Year",
                                   xval = 10, minbucket = 100, minsplit =
300, mtry = 100))
  beep()

  rf_prediction<-predict(rf_mod, newdata = test_frame_sub)

  cor.test(rf_prediction, test_phen_Y)
  print(paste0("bcrf trained",i))

  ## removes the phenotype values from the model frame
  mod_frame_sub<-as.matrix(mod_frame_sub[,-
```

```
c(grep("Protein",colnames(mod_frame_sub)))])

  ## Fits a linear SVM
  system.time(lsvm_mod<-svm(x = mod_frame_sub, y = mod_Phen_Y, #scale = F,
                            grid_choice = 2,
                            #max_gamma = 25,
                            threads  =-1, random_seed = 8394))
  lsvm_prediction<-predict(lsvm_mod, newdata = test_frame_sub)

  cor.test(lsvm_prediction, test_phen_Y)
  print(paste0("lsvm trained",i))

  ## saves models, pred/obs data as list
  saveRDS(lsvm_mod, file =
paste0("model_frames_and_imputations/siteVariety_models/rf_Protein_100DAS",
                                  spp_list[[i]],".rds"))
  saveRDS(rf_mod, file =
paste0("model_frames_and_imputations/siteVariety_models/lsvm_Protein_100DAS"
,
                                spp_list[[i]],".rds"))
  Protein_predobs_100[[i]]<-rbind(test_phen_Y, rf_prediction,
lsvm_prediction)

}


saveRDS(Yield_predobs_200,file =
"model_frames_and_imputations/siteVariety_models/Yield_predobs_200.rds")
saveRDS(Yield_predobs_100,file =
"model_frames_and_imputations/siteVariety_models/Yield_predobs_100.rds")

saveRDS(Protein_predobs_200,file =
"model_frames_and_imputations/siteVariety_models/Protein_predobs_200.rds")
saveRDS(Protein_predobs_100,file =
"model_frames_and_imputations/siteVariety_models/Protein_predobs_100.rds")

##########################################################################################################################
#

## Compares accuracy of species-targeted annual forecast models to
"omnibus" annual forecast species model.

## returns accuracy of yield+protein models across species where terminal
year is 2018 (other comparisons are not valid)
## crops not tested in 2018 are Lupins, Lentils, Triticale
crop_grep<-c("METADom_sub_cropCanola",
             "METADom_sub_cropChickpea",
             "METADom_sub_cropFaba.Bean",
             "METADom_sub_cropLupin",
             "METADom_sub_cropOat",
             "METADom_sub_cropField.Pea",
             "METADom_sub_cropLentil",
             "METADom_sub_cropTriticale",
             "METADom_sub_cropWheat")


delta_accuracy_omnibus<-NULL
for(i in c(1:7, 9)){

  ## adds the omnibus model accuracy for this species at 200DAS
  delta_accuracy_omnibus[[i]]<-c(length(Yield_predobs_200[[i]][1,]),
                                 ## the xvBCRF prediction

cor.test(Yield_predobs_200[[i]][2,],Yield_predobs_200[[i]][1,])$estimate,

rmserr(Yield_predobs_200[[i]][2,],Yield_predobs_200[[i]][1,])$rmse,
                                 ## the lsvm prediction

cor.test(Yield_predobs_200[[i]][3,],Yield_predobs_200[[i]][1,])$estimate,

rmserr(Yield_predobs_200[[i]][3,],Yield_predobs_200[[i]][1,])$rmse,
                                 ## BCRF omnibus models

cor.test(predobs_frame_unscaled[[2]]$Yield[Species_test_df1[,c(colnames(Species_test_df1)==crop_grep[[i]])]==1]
,

predobs_frame_unscaled[[2]]$xvrf_200[Species_test_df1[,c(colnames(Species_test_df1)==crop_grep[[i]])]==1])$estimate
,

rmserr(predobs_frame_unscaled[[2]]$Yield[Species_test_df1[,c(colnames(Species_test_df1)==crop_grep[[i]])]==1]
,

predobs_frame_unscaled[[2]]$xvrf_200[Species_test_df1[,c(colnames(Species_test_df1)==crop_grep[[i]])]==1])$rmse
,
                                 ## LSVM omnibus models

cor.test(predobs_frame_scaled[[2]]$Yield[Species_test_df1[,c(colnames(Species_test_df1)==crop_grep[[i]])]==1]
,

predobs_frame_scaled[[2]]$lsvm_200[Species_test_df1[,c(colnames(Species_test_df1)==crop_grep[[i]])]==1])$estimate
,

rmserr(predobs_frame_scaled[[2]]$Yield[Species_test_df1[,c(colnames(Species_test_df1)==crop_grep[[i]])]==1]
```

```
,

predobs_frame_scaled[[2]]$lsvm_200[Species_test_df1[,c(colnames(Species_test_df1)==crop_grep[[i]])]==1])$rmse
,
                                 ## the 100DAS data
                                 length(Yield_predobs_100[[i]][1,]),
                                 ## the xvBCRF prediction

cor.test(Yield_predobs_100[[i]][2,],Yield_predobs_100[[i]][1,])$estimate,

rmserr(Yield_predobs_100[[i]][2,],Yield_predobs_100[[i]][1,])$rmse,
                                 ## the lsvm prediction

cor.test(Yield_predobs_100[[i]][3,],Yield_predobs_100[[i]][1,])$estimate,

rmserr(Yield_predobs_100[[i]][3,],Yield_predobs_100[[i]][1,])$rmse,
                                 ## BCRF omnibus models

cor.test(predobs_frame_unscaled[[2]]$Yield[Species_test_df1[,c(colnames(Species_test_df1)==crop_grep[[i]])]==1]
,

predobs_frame_unscaled[[2]]$xvrf_100[Species_test_df1[,c(colnames(Species_test_df1)==crop_grep[[i]])]==1])$estimate
,

rmserr(predobs_frame_unscaled[[2]]$Yield[Species_test_df1[,c(colnames(Species_test_df1)==crop_grep[[i]])]==1]
,

predobs_frame_unscaled[[2]]$xvrf_100[Species_test_df1[,c(colnames(Species_test_df1)==crop_grep[[i]])]==1])$rmse
,
                                 ## LSVM omnibus models

cor.test(predobs_frame_scaled[[2]]$Yield[Species_test_df1[,c(colnames(Species_test_df1)==crop_grep[[i]])]==1]
,

predobs_frame_scaled[[2]]$lsvm_100[Species_test_df1[,c(colnames(Species_test_df1)==crop_grep[[i]])]==1])$estimate
,

rmserr(predobs_frame_scaled[[2]]$Yield[Species_test_df1[,c(colnames(Species_test_df1)==crop_grep[[i]])]==1]
,

predobs_frame_scaled[[2]]$lsvm_100[Species_test_df1[,c(colnames(Species_test_df1)==crop_grep[[i]])]==1])$rmse
)
  ## prints target sample size
}

## returns accuracy of yield+protein models across species where terminal
year is 2018 (other comparisons are not valid)
delta_accuracy_omnibus<-ldply(as.data.frame(delta_accuracy_omnibus[-
c(4,7,8)]))
colnames(delta_accuracy_omnibus)<-c("junk","n_sample",
                                    "BCRF_targeted_rsq",
"BCRF_targeted_RMSE",
                                    "LSVM_targeted_rsq",
"LSVM_targeted_RMSE",
                                    "BCRF_omni_rsq", "BCRF_omni_RMSE",
                                    "LSVM_omni_rsq", "LSVM_omni_RMSE",
                                    "n_sample_100",
                                    "BCRF_targeted_rsq_100",
"BCRF_targeted_RMSE_100",
                                    "LSVM_targeted_rsq_100",
"LSVM_targeted_RMSE_100",
                                    "BCRF_omni_rsq_100",
"BCRF_omni_RMSE_100",
                                    "LSVM_omni_rsq_100",
"LSVM_omni_RMSE_100")

delta_accuracy_omnibus<-delta_accuracy_omnibus[-c(1)]

rownames(delta_accuracy_omnibus)<-c("METADom_sub_cropCanola",
                                    "METADom_sub_cropChickpea",
                                    "METADom_sub_cropFaba.Bean",
                                    #  "METADom_sub_cropLupin",
                                    "METADom_sub_cropOat",
                                    "METADom_sub_cropField.Pea",
                                    #  "METADom_sub_cropLentil",
                                    # "METADom_sub_cropTriticale",
                                    "METADom_sub_cropWheat")
write.csv(delta_accuracy_omnibus, "omnibus_versus_targeted.csv", row.names
= T)

## That is, omnibus models have marginal benefits sometimes,
## large costs often.

## what's going on? IDK.

###############################################################################
#

## Builds BCRF models for wheat only, 100DAS, all phenotypes w. >5000
observations
## then generates rulesets for all of them

###############################################################################
```

```
#

## loads NON-rescaled data
#mod_frame1<-read.csv(file =
"model_frames_and_imputations/merged_variety_data_rescaled_impute1.csv",
header = T)
mod_frame1<-readRDS("model_frames_and_imputations/mod_frame1.rds")

## separates phenotypes
Phen_data<-mod_frame1[,grep("PHENDom", colnames(mod_frame1))]
mod_frame1<-mod_frame1[,-c(grep("PHENDom", colnames(mod_frame1)))]

Common_phens<-colnames(Phen_data)[colSums(!is.na(Phen_data))>25000]

## generates a common (numeric) phenotype frame, in which I am
interested....
Common_phens<-Common_phens[c(20,24,27,28,35)]
Common_phens<-c(Common_phens,
                "PHENDom_yield_pct_of_average",
                "PHENDom_zeroInflated_yield_t_ha",
                "PHENDom_Flowering_QC",
                "PHENDom_Glucosinolates")

## converts zero-inflated yield to pass/fail flag
Phen_data$PHENDom_FailedYield<-
c(as.numeric(as.character(Phen_data$PHENDom_zeroInflated_yield_t_ha))>0)

Common_phens<-c(Common_phens,"PHENDom_FailedYield")

## Fixes a prob
Phen_data$PHENDom_yield_pct_of_average<-
as.numeric(as.character(Phen_data$PHENDom_yield_pct_of_average))
Phen_data$PHENDom_FailedYield<-as.factor(Phen_data$PHENDom_FailedYield)

## converts protein (from source 4) to numeric
Phen_data$PHENDom_Protein_4<-
as.numeric(as.character(Phen_data$PHENDom_Protein_4))

## generates a naive random forest for each, and crossvalidates in 100
random holdout sites
set.seed(8394)
CommonPhen_predictions<-NULL

for(i in 1:length(Common_phens)){

  mod_frame_sub<-mod_frame1

  ## removes NAs in the phenotype, and all corresponding rows in model test
and train data frames
  mod_phen<-Phen_data[,c(colnames(Phen_data)==Common_phens[i])]

  ## screens a few junk characters
  if(is.character(mod_phen)){
    mod_phen<-as.numeric(mod_phen)
  }

  mod_frame_sub<-mod_frame_sub[!is.na(mod_phen),]
  mod_phen<-mod_phen[!is.na(mod_phen)]

  mod_frame_sub$Phenotype<-mod_phen

  ## Selects 100 random holdout sites
  Site_vec<-sample(paste0(mod_frame_sub$BOMDom_Sowing_date_numeric,
                mod_frame_sub$BOMDom_Latitude,
                mod_frame_sub$BOMDom_Longitude), size = 100, replace = F)

  test_frame_sub<-
mod_frame_sub[c(paste0(mod_frame_sub$BOMDom_Sowing_date_numeric,
                                       mod_frame_sub$BOMDom_Latitude,
                                       mod_frame_sub$BOMDom_Longitude) %in%
Site_vec),]

  mod_frame_sub<-
mod_frame_sub[!(paste0(mod_frame_sub$BOMDom_Sowing_date_numeric,
                                         mod_frame_sub$BOMDom_Latitude,
                                         mod_frame_sub$BOMDom_Longitude)
%in% Site_vec),]

  require(ranger)
  system.time(rf_mod<-ranger(Phenotype~., mod_frame_sub, seed = 8394,
num.trees = 1001))
  detach(package:ranger)

  rf_prediction<-predict(rf_mod, data = test_frame_sub)

  ## retains phenotype and prediction
  CommonPhen_predictions[[i]]<-cbind(test_frame_sub$Phenotype,
rf_prediction$predictions)
  rm(mod_frame_sub, test_frame_sub)
}

names(CommonPhen_predictions)<-Common_phens
```

```
saveRDS(CommonPhen_predictions, file =
"model_frames_and_imputations/site_models/CommonPhen_predictions.rds")

## All done for S
```
