## Supplementary Code 2 for "Explainable machine learning models of major crop traits from satellite-monitored continent-wide field trial data"

```
## Analysis to break the "black box" of ML into an interpretable model,
with clear heuristic rulesets

require(inTrees)
require(randomForest)
#require(rminer)

#############

##  Just before we start, compares the predictive value of XXX with and
without Dicot T/F variable

#############

## loads data
mod_frame1<-readRDS("model_frames_and_imputations/mod_frame1.rds")

peanutCluster<-makeCluster(detectCores()-2)
registerDoParallel(peanutCluster)

## first, separates phenotypes
Phen_data<-mod_frame1[,grep("PHENDom", colnames(mod_frame1))]
mod_frame1<-mod_frame1[,-c(grep("PHENDom", colnames(mod_frame1)))]

## attaches the phenotype
mod_frame1$Yield<-Phen_data$PHENDom_yield_t_ha

mod_frame_Y<-mod_frame1[!is.na(mod_frame1$Yield),]

require(ranger)
system.time(rf_mod<-ranger(Yield~., mod_frame_Y, seed = 8394, num.trees =
1000, importance = "impurity"))
detach(package:ranger)

## removes the Dicot t/f and Fabaceae t/f variables
mod_frame_Y<-mod_frame_Y[,-c(grep("Dicot", colnames(mod_frame_Y)))]
mod_frame_Y<-mod_frame_Y[,-c(grep("Fabac", colnames(mod_frame_Y)))]

require(ranger)
system.time(rf_mod2<-ranger(Yield~., mod_frame_Y, seed = 8394, num.trees =
1000, importance = "impurity"))
detach(package:ranger)

## puts in a "CLETHODIM applicable crops" variable
mod_frame_Y<-mod_frame1[!is.na(mod_frame1$Yield),]

CLACs<-rep(0, times = nrow(mod_frame_Y))
CLACs[mod_frame_Y$METADom_sub_cropCanola==1]<-1
CLACs[mod_frame_Y$METADom_sub_cropFaba.Bean==1]<-1
CLACs[mod_frame_Y$METADom_sub_cropChickpea==1]<-1
CLACs[mod_frame_Y$METADom_sub_cropField.Pea==1]<-1
CLACs[mod_frame_Y$METADom_sub_cropLupin==1]<-1
CLACs[mod_frame_Y$METADom_sub_cropLentil==1]<-1

mod_frame_Y$METADom_CLACs<-CLACs

require(ranger)
system.time(rf_mod3<-ranger(Yield~., mod_frame_Y, seed = 8394, num.trees =
1000, importance = "impurity"))
detach(package:ranger)

## Identifies importance of this grouping
head(rf_mod3$variable.importance[order(rf_mod3$variable.importance,
decreasing =  T)],30)

## Clethodim importance is maintained.

############################################################################
#

## Generates a Wheat yield forecast model, interprets trees

############################################################################
#

## loads the model
rf_mod_allTimes<-readRDS(file =
"model_frames_and_imputations/siteVariety_models/bcrf_allTimes.rds")

## loads data
mod_frame1<-readRDS("model_frames_and_imputations/mod_frame1.rds")

peanutCluster<-makeCluster(detectCores()-2)
registerDoParallel(peanutCluster)

## first, separates phenotypes
Phen_data<-mod_frame1[,grep("PHENDom", colnames(mod_frame1))]
mod_frame1<-mod_frame1[,-c(grep("PHENDom", colnames(mod_frame1)))]

## attaches the phenotype
mod_frame1$Yield<-Phen_data$PHENDom_yield_t_ha

####################################

## filters out +200DAS and postsowing data
mod_frame_Y<-mod_frame1[,-c(grep("_2[0-9]{2}$", colnames(mod_frame1)))]
mod_frame_Y<-mod_frame_Y[,-c(grep("PostSowing", colnames(mod_frame_Y),
ignore.case = T))]

## removes "TRIAL ID" vectors (used for crossvalidation)
#mod_frame_Y<-mod_frame_Y[,-c(grep("Trial_ID", colnames(mod_frame_Y),
ignore.case = T))]

## attaches the phenotype
mod_frame_Y$Yield<-Phen_data$PHENDom_yield_t_ha

## gets wheat only
mod_frame_Y<-mod_frame_Y[Phen_data$PHENDom_Crop_alldata=="Wheat",]

## removes na data
mod_frame_Y<-mod_frame_Y[!is.na(mod_frame_Y$Yield),]

## splits off 2018 data
test_frame_Y<-mod_frame_Y[mod_frame_Y$METADom_Year>=2018,]
mod_frame_Y<-mod_frame_Y[mod_frame_Y$METADom_Year<2018,]
```

```
## builds an rpart model, cross-validated by unique year
set.seed(8394)
system.time(rpart_mod_all<-rpart(Yield~., mod_frame_Y,
                            control = rpart.control(minbucket = 25, minsplit
= 50, cp = 0.001, xval = 10)))


beep()

## builds a reference xgboost tree
for(i in 1:ncol(mod_frame_Y)){mod_frame_Y[,i]<-as.numeric(mod_frame_Y[,i])}

mod_frame2<-mod_frame_Y[,-c(grep("Yield",colnames(mod_frame_Y)))]
#mod_frame2<-sapply(mod_frame2, as.numeric)
mod_frame2<-xgb.DMatrix(as.matrix(mod_frame2), label= mod_frame_Y$Yield)

system.time(xg_mod_all<-xgboost(data = mod_frame2, nthread = 16, nrounds
=10))


## accuracy under forward prediction (note: this is different to the
crossvalidation accuracy reported in Supp. Figs below)
for(i in 1:ncol(test_frame_Y)){test_frame_Y[,i]<-
as.numeric(test_frame_Y[,i])}

cor.test(predict(rpart_mod_all, newdata = test_frame_Y),
test_frame_Y$Yield)

## allmod accuracy Rsq = 0.748658

## accuracy of xgb model
throwaway<-test_frame_Y
for(i in 1:ncol(throwaway)){throwaway[,i]<-as.numeric(throwaway[,i])}
throwaway<-throwaway[,-c(grep("Yield",colnames(throwaway)))]
throwaway<-as.matrix(throwaway)

cor.test(predict(xg_mod_all, newdata = throwaway), test_frame_Y$Yield)

## allmod accuracy Rsq = 0.7486631

###########################################################################
#

## Explores domain-driven model accuracy, by leaving out entire domains

###########################################################################
#


## builds a model with no BOM data (not actually different)

mod_frame_BOM<-mod_frame_Y[,-c(c(grep("BOMDom_", colnames(mod_frame_Y))[-
c(1:3)]))]

set.seed(8394)
system.time(rpart_mod_noBOM<-rpart(Yield~., mod_frame_BOM,
                               control = rpart.control(minbucket = 25,
minsplit = 50, cp = 0.01, xval = 10)))


beep()

## accuracy improves very slightly: 0.753255
cor.test(predict(rpart_mod_noBOM, newdata = test_frame_Y),
test_frame_Y$Yield)$estimate-0.748658

#system.time(rf_mod_noBOM<-randomForest(Yield~., mod_frame_BOM,
#                                   strata = "METADom_Year",
#                                   xval = 10, minbucket = 100, minsplit =
300, mtry = 200))

for(i in 1:ncol(mod_frame_Y)){mod_frame_Y[,i]<-as.numeric(mod_frame_Y[,i])}

mod_frame2<-as.matrix(mod_frame_BOM[,-
c(grep("Yield",colnames(mod_frame_BOM)))])
mod_frame2<-xgb.DMatrix(mod_frame2, label= mod_frame_BOM$Yield)

system.time(xg_mod_noBOM<-xgboost(data = mod_frame2, nthread = 16, nrounds
=10))


## Builds a model with no satellite data
mod_frame_BOM<-mod_frame_Y[,-c(c(grep("SatDom_", colnames(mod_frame_Y))[-
c(1:3)]))]

set.seed(8394)
system.time(rpart_mod_noSAT<-rpart(Yield~., mod_frame_BOM,
                                   control = rpart.control(minbucket = 25,
minsplit = 50, cp = 0.005, xval = 10)))


beep()

mod_frame2<-as.matrix(mod_frame_BOM[,-
c(grep("Yield",colnames(mod_frame_BOM)))])
mod_frame2<-xgb.DMatrix(mod_frame2, label= mod_frame_BOM$Yield)

system.time(xg_mod_noSAT<-xgboost(data = mod_frame2, nthread = 16, nrounds
=10))

## Builds a model with no management data
mod_frame_BOM<-mod_frame_Y[,-c(grep("MANDom_", colnames(mod_frame_Y)))]

set.seed(8394)
system.time(rpart_mod_noMAN<-rpart(Yield~., mod_frame_BOM,
                                   control = rpart.control(minbucket = 25,
minsplit = 50, cp = 0.01, xval = 10)))

mod_frame2<-as.matrix(mod_frame_BOM[,-
c(grep("Yield",colnames(mod_frame_BOM)))])
mod_frame2<-xgb.DMatrix(mod_frame2, label= mod_frame_BOM$Yield)

system.time(xg_mod_noMAN<-xgboost(data = mod_frame2, nthread = 16, nrounds
=10))

beep()
```

```
## removes metadata
mod_frame_BOM<-mod_frame_Y[,-c(grep("METADom_", colnames(mod_frame_Y)))]

set.seed(8394)
system.time(rpart_mod_noMETA<-rpart(Yield~., mod_frame_BOM,
                                    control = rpart.control(minbucket = 25,
minsplit = 50, cp = 0.01, xval = 10)))

mod_frame2<-as.matrix(mod_frame_BOM[,-
c(grep("Yield",colnames(mod_frame_BOM)))])
mod_frame2<-xgb.DMatrix(mod_frame2, label= mod_frame_BOM$Yield)

system.time(xg_mod_noMETA<-xgboost(data = mod_frame2, nthread = 16, nrounds
=10))

beep()



## Shows the delta forward-projection accuracy of each LOO model

## for the rpart model:

## Alldata accuracy is rsq = 0.748658

## accuracy improves very slightly: 0.753255
cor.test(predict(rpart_mod_noBOM, newdata = test_frame_Y),
test_frame_Y$Yield)
## accuracy falls a LOT: 0.3564833
cor.test(predict(rpart_mod_noSAT, newdata = test_frame_Y),
test_frame_Y$Yield)
## accuracy falls very slightly: 0.7092213
cor.test(predict(rpart_mod_noMETA, newdata = test_frame_Y),
test_frame_Y$Yield)
## accuracy improves very slightly: 0.753255
cor.test(predict(rpart_mod_noMAN, newdata = test_frame_Y),
test_frame_Y$Yield)

## the internal accuracy estimates (not forward-projection)
cbind(xg_mod_all$evaluation_log,
      xg_mod_noBOM$evaluation_log,
      xg_mod_noSAT$evaluation_log,
      xg_mod_noMETA$evaluation_log,
      xg_mod_noMAN$evaluation_log)

#################

## instead of leave-one-out, keeps just one domain each time

#################

## builds a model with BOM only
mod_frame_BOM<-mod_frame_Y[,-c(grep("SatDom_", colnames(mod_frame_Y)),
                               grep("MANDom_", colnames(mod_frame_Y)),
                               grep("METADom_", colnames(mod_frame_Y)),
                               grep("ENVDom_", colnames(mod_frame_Y)))]

set.seed(8394)
system.time(rpart_mod_BOMOnly<-rpart(Yield~., mod_frame_BOM,
                                   control = rpart.control(minbucket = 25,
minsplit = 50, cp = 0.01, xval = 10)))

mod_frame2<-as.matrix(mod_frame_BOM[,-
c(grep("Yield",colnames(mod_frame_BOM)))])
mod_frame2<-xgb.DMatrix(mod_frame2, label= mod_frame_BOM$Yield)

system.time(xg_mod_BOMOnly<-xgboost(data = mod_frame2, nthread = 16,
nrounds  =10))


beep()


## builds a model with satellite only
mod_frame_BOM<-mod_frame_Y[,-c(c(grep("BOMDom_", colnames(mod_frame_Y))[-
c(1:3)]),
                               grep("MANDom_", colnames(mod_frame_Y)),
                               grep("METADom_", colnames(mod_frame_Y)),
                               grep("ENVDom_", colnames(mod_frame_Y)))]

set.seed(8394)
system.time(rpart_mod_SatOnly<-rpart(Yield~., mod_frame_BOM,
                                     control = rpart.control(minbucket =
25, minsplit = 50, cp = 0.01, xval = 10)))

mod_frame2<-as.matrix(mod_frame_BOM[,-
c(grep("Yield",colnames(mod_frame_BOM)))])
mod_frame2<-xgb.DMatrix(mod_frame2, label= mod_frame_BOM$Yield)

system.time(xg_mod_SatOnly<-xgboost(data = mod_frame2, nthread = 16,
nrounds  =10))


beep()

## builds a model with Metadata only
mod_frame_BOM<-mod_frame_Y[,-c(c(grep("BOMDom_", colnames(mod_frame_Y))[-
c(1:3)]),
                               grep("SatDom_", colnames(mod_frame_Y)),
                               grep("MANDom_", colnames(mod_frame_Y)),
                              # grep("METADom_", colnames(mod_frame_Y)),
                               grep("ENVDom_", colnames(mod_frame_Y)))]

set.seed(8394)
system.time(rpart_mod_METAOnly<-rpart(Yield~., mod_frame_BOM,
                                     control = rpart.control(minbucket =
25, minsplit = 50, cp = 0.01, xval = 10)))

mod_frame2<-as.matrix(mod_frame_BOM[,-
c(grep("Yield",colnames(mod_frame_BOM)))])
mod_frame2<-xgb.DMatrix(mod_frame2, label= mod_frame_BOM$Yield)

system.time(xg_mod_METAOnly<-xgboost(data = mod_frame2, nthread = 16,
nrounds  =10))


beep()
```

```
## builds a model with management data only
mod_frame_BOM<-mod_frame_Y[,-c(c(grep("BOMDom_", colnames(mod_frame_Y))[-
c(1:3)]),
                              grep("SatDom_", colnames(mod_frame_Y)),
                             # grep("MANDom_", colnames(mod_frame_Y)),
                              grep("METADom_", colnames(mod_frame_Y)),
                              grep("ENVDom_", colnames(mod_frame_Y)))]

set.seed(8394)
system.time(rpart_mod_MANOnly<-rpart(Yield~., mod_frame_BOM,
                                     control = rpart.control(minbucket =
25, minsplit = 50, cp = 0.01, xval = 10)))

mod_frame2<-as.matrix(mod_frame_BOM[,-
c(grep("Yield",colnames(mod_frame_BOM)))])
mod_frame2<-xgb.DMatrix(mod_frame2, label= mod_frame_BOM$Yield)

system.time(xg_mod_MANOnly<-xgboost(data = mod_frame2, nthread = 16,
nrounds  =10))


beep()


### Again compares accuracy

## Alldata accuracy is rsq = 0.748658

## Rsq = 0.3370216
cor.test(predict(rpart_mod_BOMOnly, newdata = test_frame_Y),
test_frame_Y$Yield)
## Rsq = 0.7071491
cor.test(predict(rpart_mod_SatOnly, newdata = test_frame_Y),
test_frame_Y$Yield)
## Rsq = 0.587962
cor.test(predict(rpart_mod_METAOnly, newdata = test_frame_Y),
test_frame_Y$Yield)
## Rsq = 0.3731411
cor.test(predict(rpart_mod_MANOnly, newdata = test_frame_Y),
test_frame_Y$Yield)

cbind(xg_mod_all$evaluation_log,
      xg_mod_BOMOnly$evaluation_log,
      xg_mod_SatOnly$evaluation_log,
      xg_mod_METAOnly$evaluation_log,
      xg_mod_MANOnly$evaluation_log)

############################################################

## builds a figure for comparison

############################################################


tiff(filename  = "Publication_docs/Figure_domain_improvement.tiff",
     width  =7, height = 5, units = "in", res  =330)

par(mfrow = c(1,2), bty = "n", mar = c(5.1,4.1,1.1,1.1))

plot(rpart_mod_all$cptable[,4], type = "l",
     xlab = "Model complexity", ylab = "Missing-Plot CV Error Rate",
     ylim = c(0,1), xlim = c(0,10))


points(rpart_mod_BOMOnly$cptable[,4], type = "l", col = "orange")
points(rpart_mod_BOMOnly$cptable[,4], pch = 16, col = "orange")

points(rpart_mod_MANOnly$cptable[,4], type = "l", col = "darkgreen")
points(rpart_mod_MANOnly$cptable[,4], pch = 16, col = "darkgreen")

points(rpart_mod_METAOnly$cptable[,4], type = "l", col = "pink", lwd = 2)
points(rpart_mod_METAOnly$cptable[,4], pch = 16, col = "pink")

points(rpart_mod_SatOnly$cptable[,4], type = "l", col = "lightblue")
points(rpart_mod_SatOnly$cptable[,4], pch = 20, col = "lightblue")

points(rpart_mod_all$cptable[,4])

abline(h = seq(0,1, by = 0.2), lty = 3)

mtext("a", side  =3, adj = 0)

## now the xgboost models
plot(xg_mod_all$evaluation_log,
     type = "l",
     ylab = "RMSE", xlab = "Number of Boosting Iterations",
     xlim = c(0,10), ylim = c(0,2.5))

points(xg_mod_BOMOnly$evaluation_log, type = "l", col = "orange")
points(xg_mod_BOMOnly$evaluation_log, pch = 16, col = "orange")

points(xg_mod_MANOnly$evaluation_log, type = "l", col = "darkgreen")
points(xg_mod_MANOnly$evaluation_log, pch = 16, col = "darkgreen")

points(xg_mod_METAOnly$evaluation_log, type = "l", col = "pink")
points(xg_mod_METAOnly$evaluation_log, pch = 16, col = "pink")

points(xg_mod_SatOnly$evaluation_log, type = "l", col = "lightblue")
points(xg_mod_SatOnly$evaluation_log, pch = 16, col = "lightblue")

points(xg_mod_all$evaluation_log)

abline(h = seq(0,3, by = 0.5), lty = 3)
mtext("b", side  =3, adj = 0)

dev.off()



tiff(filename  =
"Publication_docs/Figure_domain_improvement_LeaveOneOut.tiff",
     width  =7, height = 5, units = "in", res  =330)


par(mfrow = c(1,2), bty = "n", mar = c(5.1,4.1,1.1,1.1))

plot(rpart_mod_all$cptable[,4], type = "l",
```

```
     xlab = "Model complexity", ylab = "Missing-Plot CV Error Rate",
     ylim = c(0,1), xlim = c(0,10))


points(rpart_mod_noBOM$cptable[,4], type = "l", col = "orange")
points(rpart_mod_noBOM$cptable[,4], pch = 16, col = "orange")

points(rpart_mod_noMAN$cptable[,4], type = "l", col = "darkgreen")
points(rpart_mod_noMAN$cptable[,4], pch = 16, col = "darkgreen", cex = 0.6)

points(rpart_mod_noMETA$cptable[,4], type = "l", col = "pink", lty = 2)
points(rpart_mod_noMETA$cptable[,4], pch = 20, col = "pink", cex = 0.6)

points(rpart_mod_noSAT$cptable[,4], type = "l", col = "lightblue")
points(rpart_mod_noSAT$cptable[,4], pch = 20, col = "lightblue")

points(rpart_mod_all$cptable[,4])

abline(h = seq(0,1, by = 0.2), lty = 3)

mtext("a", side  =3, adj = 0)

## now the xgboost models
plot(xg_mod_all$evaluation_log,
     type = "l",
     ylab = "RMSE", xlab = "Number of Boosting Iterations",
     xlim = c(0,10), ylim = c(0,2.5))

points(xg_mod_noBOM$evaluation_log, type = "l", col = "orange")
points(xg_mod_noBOM$evaluation_log, pch = 16, col = "orange")

points(xg_mod_noMAN$evaluation_log, type = "l", col = "darkgreen")
points(xg_mod_noMAN$evaluation_log, pch = 16, col = "darkgreen")

points(xg_mod_noMETA$evaluation_log, type = "l", col = "pink", lty = 2)
points(xg_mod_noMETA$evaluation_log, pch = 20, col = "pink", cex = 0.6)

points(xg_mod_noSAT$evaluation_log, type = "l", col = "lightblue")
points(xg_mod_noSAT$evaluation_log, pch = 16, col = "lightblue")

points(xg_mod_all$evaluation_log)

abline(h = seq(0,3, by = 0.5), lty = 3)
mtext("b", side  =3, adj = 0)

dev.off()


par(mfrow = c(1,1), bty = "n", mar = c(5.1,4.1,2.1,2.1))
## cleans up the workspace
rm(xg_mod_all, xg_mod_BOMOnly, xg_mod_MANOnly, xg_mod_METAOnly)

rm(rpart_mod_all, rpart_mod_BOMOnly, rpart_mod_MANOnly, rpart_mod_METAOnly)
rm(rpart_mod_noBOM, rpart_mod_noSAT, rpart_mod_noMETA)

############################################################

## builds a BCRF model for interpretation

############################################################


## loads data
mod_frame1<-readRDS("model_frames_and_imputations/mod_frame1.rds")

## separates phenotypes
Phen_data<-mod_frame1[,grep("PHENDom", colnames(mod_frame1))]
mod_frame1<-mod_frame1[,-c(grep("PHENDom", colnames(mod_frame1)))]

## attaches the target phenotype
mod_frame1$Yield<-Phen_data$PHENDom_yield_t_ha

## removes na data
mod_frame_Y<-mod_frame1[!is.na(mod_frame1$Yield),]


## removes "TRIAL ID" vectors (used for crossvalidation)
mod_frame_Y<-mod_frame_Y[,-c(grep("Trial_ID", colnames(mod_frame_Y),
ignore.case = T))]

## splits off 2018 data
test_frame_Y<-mod_frame_Y[mod_frame_Y$METADom_Year>=2018,]
mod_frame_Y<-mod_frame_Y[mod_frame_Y$METADom_Year<2018,]

Site_index_vec<-paste0(mod_frame_Y$BOMDom_Longitude,
mod_frame_Y$BOMDom_Latitude, mod_frame_Y$BOMDom_Sowing_date_numeric)

## subsamples 100 experiments from the pre-2018 data
set.seed(8394)
Site_subset<-sample(Site_index_vec, 100)
test_frame_Y_site<-mod_frame_Y[c(Site_index_vec %in% Site_subset),]
mod_frame_Y<-mod_frame_Y[!(Site_index_vec %in% Site_subset),]

## keeps track of which species is attached to which prediction, for later
Species_test_df1<-data.frame(test_frame_Y[,grep("METADom_sub_crop",
colnames(test_frame_Y))])


rf_mod<-readRDS(file =
"model_frames_and_imputations/siteVariety_models/bcrf_allTimes.rds")

## builds defined ruleset
set.seed(37491)
rfTreelist<-RF2List(rf_mod)
test_Phen<-test_frame_Y$Yield
train_Phen<-mod_frame_Y$Yield

RF_list_obj<-RF2List(rf_mod)

## generates most robust rule set for common decision cascades of length
<=5
xobj_rules<-extractRules(RF_list_obj,mod_frame_Y, 5, ntree = 5000, random =
FALSE)

## trims out unique rule cascades only
xobj_rules<-unique(xobj_rules)

## renders into a matrix
```

```
newrules<-getRuleMetric(xobj_rules,mod_frame_Y, train_Phen)

## trims the set
newrules<-pruneRule(newrules,mod_frame_Y, train_Phen)

## presents these as a readable set of heuristics
clean_rules<-presentRules(newrules,colnames(mod_frame_Y))

## Crudely filters by error rates, removing above-median error rates
clean_rules_sub<-
newrules[c(as.numeric(as.character(newrules[,3]))<=quantile(as.numeric(as.character(newrules[,3]))
, 0.1)),]

## random-samples data frame to prevent overwhelming vector-memory limits
set.seed(29374)
samp_vec<-sample(c(1:nrow(mod_frame_Y)), replace = F, size = 10000)

mod_frame_Y<-mod_frame_Y[c(samp_vec),]
train_Phen<-train_Phen[samp_vec]

system.time(clean_rules_select<-selectRuleRRF(newrules,mod_frame_Y,
train_Phen))

## writes them out
#write.csv(clean_rules, "Table_SXX_wheat_rules_250DAS.csv", row.names = F)

## uses feature selection, not error, to get rules
clean_rules_select<-selectRuleRRF(newrules,mod_frame_Y, train_Phen)

saveRDS(clean_rules_select,"clean_wheat_rules_250DAS.rds")

## keeps lowest error 100 rules
clean_rules_tail<-head(clean_rules[order(as.numeric(clean_rules[,3])),],
100)

## the most important 'ranked' rules
clean_rules_tail<-
tail(clean_rules_select[order(as.numeric(clean_rules_select[,6])),], 50)

clean_rules_tail<-presentRules(clean_rules_tail,colnames(mod_frame_Y))
## keeps most frequent 100 rules
#clean_rules_tail<-tail(clean_rules[order(clean_rules[,2]),], 100)

write.csv(clean_rules_tail, file = "clean_ruleset.csv")

## These rules don't clarify much, UNLESS you know what the species is (and
hence where the relative yield falls)
## E.g. the FPAR_max_150<0.03 =2.9 t yild could be a great chickpea yield,
or average wheat yield.


#################

## Rebuilds this approach, using species-specific yield models

################

## sets up data frames, loads models

## loads rescaled data for lsvms...
#mod_frame1<-read.csv(file =
"model_frames_and_imputations/merged_variety_data_rescaled_impute1.csv",
header = T)
mod_frame1<-readRDS("model_frames_and_imputations/mod_frame1.rds")

## separates phenotypes
Phen_data<-mod_frame1[,grep("PHENDom", colnames(mod_frame1))]
mod_frame1<-mod_frame1[,-c(grep("PHENDom", colnames(mod_frame1)))]

######################################################################


## removes "TRIAL ID" vectors (used for crossvalidation)
mod_frame1<-mod_frame1[,-c(grep("Trial_ID", colnames(mod_frame1),
ignore.case = T))]

## Yield models for each species

## filters out +200DAS and postsowing data
mod_frame_Y<-mod_frame1[,-c(grep("_2[0-9]{2}$", colnames(mod_frame1)))]
mod_frame_Y<-mod_frame_Y[,-c(grep("PostSowing", colnames(mod_frame_Y),
ignore.case = T))]

spp_list<-unique(Phen_data$PHENDom_Crop_alldata)[-c(1,9)]

Prediction_rules<-NULL
Prediction_rules2<-NULL

## builds yield models
for(i in 1:9){

  ## depending on the species, and therefore sample sizes available,
  ## partitions data by year so that the test data contain enough
observations
  ## to evaluate models (N>250)

  ## splits off 2018+ data (rescaled)
  test_phen<-
Phen_data[c(mod_frame_Y$METADom_Year>=c(tail(unique(mod_frame_Y$METADom_Year),2)[1])),
]
  mod_phen<-Phen_data[c(mod_frame_Y$METADom_Year<-
mod_frame_Y[c(mod_frame_Y$METADom_Year>=c(tail(unique(mod_frame_Y$METADom_Year),2)[1])),
]
  mod_frame_sub<-mod_frame_Y[c(mod_frame_Y$METADom_Year<-
Phen_data[c(mod_frame_Y$METADom_Year>=c(tail(unique(mod_frame_Y$METADom_Year),4)[1])),
]
    mod_phen<-Phen_data[c(mod_frame_Y$METADom_Year<-
mod_frame_Y[c(mod_frame_Y$METADom_Year>=c(tail(unique(mod_frame_Y$METADom_Year),4)[1])),
]
    mod_frame_sub<-mod_frame_Y[c(mod_frame_Y$METADom_Year<-
Phen_data[c(mod_frame_Y$METADom_Year>=c(tail(unique(mod_frame_Y$METADom_Year),3)[1])),
]
    mod_phen<-Phen_data[c(mod_frame_Y$METADom_Year<-
mod_frame_Y[c(mod_frame_Y$METADom_Year>=c(tail(unique(mod_frame_Y$METADom_Year),3)[1])),
]
    mod_frame_sub<-mod_frame_Y[c(mod_frame_Y$METADom_Year<-
Phen_data[c(mod_frame_Y$METADom_Year>=c(tail(unique(mod_frame_Y$METADom_Year),5)[1])),
]
```

```
    mod_phen<-Phen_data[c(mod_frame_Y$METADom_Year<-
mod_frame_Y[c(mod_frame_Y$METADom_Year>=c(tail(unique(mod_frame_Y$METADom_Year),5)[1])),
]
    mod_frame_sub<-mod_frame_Y[c(mod_frame_Y$METADom_Year<-
mod_frame_sub[mod_phen$PHENDom_Crop_alldata==spp_list[[i]],]
  test_frame_sub<-
test_frame_Y[test_phen$PHENDom_Crop_alldata==spp_list[[i]],]

  test_phen_sub<-test_phen[test_phen$PHENDom_Crop_alldata==spp_list[[i]],]
  mod_phen_sub<-mod_phen[mod_phen$PHENDom_Crop_alldata==spp_list[[i]],]

  ## removes NAs in the phenotype, and all corresponding rows in model test
and train data frames
  mod_frame_sub<-mod_frame_sub[!is.na(mod_phen_sub$PHENDom_yield_t_ha),]
  test_frame_sub<-test_frame_sub[!is.na(test_phen_sub$PHENDom_yield_t_ha),]

  mod_phen_sub<-mod_phen_sub[!is.na(mod_phen_sub$PHENDom_yield_t_ha),]
  test_phen_sub<-test_phen_sub[!is.na(test_phen_sub$PHENDom_yield_t_ha),]

  ## attaches Yield to model frame
  mod_frame_sub$Yield<-mod_phen_sub$PHENDom_yield_t_ha

  ## makes shorthand copies of phenotype
  train_Phen<-mod_frame_sub$Yield
  test_Phen<-test_phen_sub$PHENDom_yield_t_ha

#  mod_frame_sub<-as.matrix(mod_frame_sub)

  ## Loads the appropriate bcrf model
#  rf_mod<-readRDS(file =
paste0("model_frames_and_imputations/siteVariety_models/lsvm_Yield_200DAS",spp_list[[i]],".rds")
)

  ## builds a bcrf model, cross-validated by unique year
  ## note the difference here between the other reported model,
  ## which was built using RESCALED data
  if(nrow(mod_frame_sub)>10000){
    set.seed(8394)
    system.time(rf_mod<-randomForest(Yield~., mod_frame_sub,
                                    strata = "METADom_Year",
                                    xval = 10, minbucket = 100, minsplit =
300, mtry = 100, ntree = 1001, sampsize = 10000))
    beep()
  }

  if(nrow(mod_frame_sub)<=10000){
    set.seed(8394)
    system.time(rf_mod<-randomForest(Yield~., mod_frame_sub,
                                    strata = "METADom_Year",
                                    xval = 10, minbucket = 100, minsplit =
300, mtry = 100, ntree = 1001))
    beep()
  }

  ## builds defined ruleset
  set.seed(37491)
  rfTreelist<-RF2List(rf_mod)
#  test_Phen<-mod_frame_sub$Yield
#  train_Phen<-mod_frame_sub$Yield

  RF_list_obj<-RF2List(rf_mod)

  ## generates most robust rule set for common decision cascades of length
<=5
  xobj_rules<-extractRules(RF_list_obj,mod_frame_sub, 5, ntree = 5000,
random = FALSE)

  ## trims out unique rule cascades only
  xobj_rules<-unique(xobj_rules)
  ## renders into a matrix
  newrules<-getRuleMetric(xobj_rules,mod_frame_sub, train_Phen)
  ## trims the set
  newrules<-pruneRule(newrules,mod_frame_sub, train_Phen)

  ## presents these as a readable set of heuristics
  clean_rules<-presentRules(newrules,colnames(mod_frame_sub))

  ## removes yield
  mod_frame_sub<-mod_frame_sub[,-c(grep("Yield", colnames(mod_frame_sub)))]

  ## if this is yield, randomly subsamples 20k rows to prevent vector
memory exhaustion
  set.seed(8394)

  if(i==9){
    sample_vec<-sample(seq(1,nrow(mod_frame_sub)), 20000, replace = F)
    mod_frame_sub<-mod_frame_sub[sample_vec,]
    train_Phen<-train_Phen[sample_vec]
    }

  ## uses feature selection, not error, to get rules
  clean_rules_select<-selectRuleRRF(newrules,mod_frame_sub, train_Phen)

  ## keeps lowest error 100 rules
  clean_rules_tail_Errors<-
head(clean_rules[order(as.numeric(clean_rules[,3])),], 100)

  ## the top 100 (or less) most important 'ranked' rules
  clean_rules_tail<-
tail(clean_rules_select[order(as.numeric(clean_rules_select[,6])),], 100)

  clean_rules_tail<-presentRules(clean_rules_tail,colnames(mod_frame_sub))

  Prediction_rules[[i]]<-clean_rules_tail
  Prediction_rules2[[i]]<-clean_rules_tail_Errors

  rm(clean_rules_tail,clean_rules_tail_Errors, rf_mod)
  gc()

  print(paste0(i, " doneburgers"))
}


rule_mat<-NULL
for(i in 1:length(spp_list)){
  ## strings together the prediction rules
  rule_mat<-rbind(rule_mat,cbind(Prediction_rules[[i]],
                                 rep(as.character(spp_list[[i]]), times =
```

```
nrow(Prediction_rules[[i]]))))
}
write.csv(rule_mat, file = "pred_rules_clean.csv", row.names = F)


rule_mat<-NULL
for(i in 1:length(spp_list)){
  ## strings together the prediction rules
  rule_mat<-rbind(rule_mat,cbind(Prediction_rules2[[i]],
                                 rep(as.character(spp_list[[i]]), times =
nrow(Prediction_rules2[[i]]))))
}
write.csv(rule_mat, file = "pred_rules_100.csv", row.names = F)


####################################

## Variety-specific wheat models

## filters out +200DAS and postsowing data
mod_frame_Y<-mod_frame1[,-c(grep("_[0-9]{3}$", colnames(mod_frame1)))]
mod_frame_Y<-mod_frame_Y[,-c(grep("PostSowing", colnames(mod_frame_Y),
ignore.case = T))]

variety_list<-
names(which(colSums(mod_frame_Y[,grep("METADom_Variety",colnames(mod_frame_Y))])>2000)
)

Prediction_rulesV<-NULL
Prediction_rulesV2<-NULL

## builds yield models
for(i in 1:7){

  ## gets the appropriate variety
  Variety_vec<-mod_frame_Y[,colnames(mod_frame_Y)==variety_list[[i]]]

  ## splits off 100 random site-variety combos
  set.seed(8394)
  Test_vec<-sample(which(Variety_vec==1), 100, replace = F)
  Train_vec<-c(which(Variety_vec==1)[!(which(Variety_vec==1) %in%
Test_vec)])

  test_phen_sub<-Phen_data[Test_vec,]
  mod_phen_sub<-Phen_data[Train_vec,]

  test_frame_sub<-mod_frame_Y[Test_vec,]
  mod_frame_sub<-mod_frame_Y[Train_vec,]

  ## removes NAs in the phenotype, and all corresponding rows in model test
and train data frames
  mod_frame_sub<-mod_frame_sub[!is.na(mod_phen_sub$PHENDom_yield_t_ha),]
  test_frame_sub<-test_frame_sub[!is.na(test_phen_sub$PHENDom_yield_t_ha),]

  mod_phen_sub<-mod_phen_sub[!is.na(mod_phen_sub$PHENDom_yield_t_ha),]
  test_phen_sub<-test_phen_sub[!is.na(test_phen_sub$PHENDom_yield_t_ha),]

  ## attaches Yield to model frame
  mod_frame_sub$Yield<-mod_phen_sub$PHENDom_yield_t_ha

  ## makes shorthand copies of phenotype
  train_Phen<-mod_frame_sub$Yield
  test_Phen<-test_phen_sub$PHENDom_yield_t_ha

  ## Builds an Rpart model
  set.seed(8394)
  system.time(rpart_mod<-rpart(Yield~., data = mod_frame_sub,
                               control = rpart.control(minbucket = 25,
minsplit = 50, cp = 0.01, xval = 10)))

  ## builds a bcrf model, cross-validated by unique year
  ## note the difference here between the other reported model,
  ## which was built using RESCALED data
  set.seed(8394)
  system.time(rf_mod<-randomForest(Yield~., mod_frame_sub,
                                   strata = "METADom_Year",
                                   xval = 10, minbucket = 100, minsplit =
300, mtry = 100, ntree = 1001))
  beep()

  ## keeps the models
  saveRDS(rf_mod, file =
paste0("model_frames_and_imputations/siteVariety_models/xvBCRF_Yield_200DAS",variety_list[[i]],".rds")
)
  saveRDS(rpart_mod, file =
paste0("model_frames_and_imputations/siteVariety_models/rp_Yield_200DAS",variety_list[[i]],".rds")
)


  ## keeps the prediction target and accuracies

  ## builds defined ruleset
  set.seed(37491)
  rfTreelist<-RF2List(rf_mod)

  RF_list_obj<-RF2List(rf_mod)

  ## generates most robust rule set for common decision cascades of length
<=5
  xobj_rules<-extractRules(RF_list_obj,mod_frame_sub, 5, ntree = 5000,
random = FALSE)

  ## trims out unique rule cascades only
  xobj_rules<-unique(xobj_rules)
  ## renders into a matrix
  newrules<-getRuleMetric(xobj_rules,mod_frame_sub, train_Phen)
  ## trims the set
  newrules<-pruneRule(newrules,mod_frame_sub, train_Phen)

  ## presents these as a readable set of heuristics
  clean_rules<-presentRules(newrules,colnames(mod_frame_sub))

  ## removes yield
  mod_frame_sub<-mod_frame_sub[,-c(grep("Yield", colnames(mod_frame_sub)))]

  ## if this is yield, randomly subsamples 20k rows to prevent vector
memory exhaustion
  set.seed(8394)
```

```
  if(i==9){
    sample_vec<-sample(seq(1,nrow(mod_frame_sub)), 20000, replace = F)
    mod_frame_sub2<-mod_frame_sub[sample_vec,]
    train_Phen2<-train_Phen[sample_vec]}

  ## uses feature selection, not error, to get rules
  clean_rules_select<-selectRuleRRF(newrules,mod_frame_sub2, train_Phen2)

  ## keeps lowest error 100 rules
  clean_rules_tail_Errors<-
head(clean_rules[order(as.numeric(clean_rules[,3])),], 100)

  ## the top 100 (or less) most important 'ranked' rules
  clean_rules_tail<-
tail(clean_rules_select[order(as.numeric(clean_rules_select[,6])),], 100)

  clean_rules_tail<-presentRules(clean_rules_tail,colnames(mod_frame_sub))

  Prediction_rulesV[[i]]<-clean_rules_tail
  Prediction_rulesV2[[i]]<-clean_rules_tail_Errors

  rm(clean_rules_tail,clean_rules_tail_Errors, rf_mod)
  gc()

  print(paste0(i, " doneburgers"))
}




###########################################

## Builds an Rpart model for canola and wheat yield, using all data and TOS
data

###########################################


## loads unscaled data...
#mod_frame1<-read.csv(file =
"model_frames_and_imputations/merged_variety_data_rescaled_impute1.csv",
header = T)
mod_frame1<-readRDS("model_frames_and_imputations/mod_frame1.rds")

mod_frame1<-mod_frame1[,-c(grep("Trial_ID", colnames(mod_frame1),
ignore.case = T))]

## separates phenotypes
Phen_data<-mod_frame1[,grep("PHENDom", colnames(mod_frame1))]
mod_frame1<-mod_frame1[,-c(grep("PHENDom", colnames(mod_frame1)))]

######################################################################


## filters out +200DAS and postsowing data
mod_frame_Y<-mod_frame1[,-c(grep("_2[0-9]{2}$", colnames(mod_frame1)))]
mod_frame_Y<-mod_frame_Y[,-c(grep("PostSowing", colnames(mod_frame_Y),
ignore.case = T))]

## splits off 2018+ data (rescaled)
test_phen<-
Phen_data[c(mod_frame_Y$METADom_Year>=c(tail(unique(mod_frame_Y$METADom_Year),2)[1])),
]
mod_phen<-Phen_data[c(mod_frame_Y$METADom_Year<-
mod_frame_Y[c(mod_frame_Y$METADom_Year>=c(tail(unique(mod_frame_Y$METADom_Year),2)[1])),
]
mod_frame_sub<-mod_frame_Y[c(mod_frame_Y$METADom_Year<-
mod_frame_sub[mod_phen$PHENDom_Crop_alldata=="Wheat",]
test_frame_sub<-test_frame_Y[test_phen$PHENDom_Crop_alldata=="Wheat",]

test_phen_sub<-test_phen[test_phen$PHENDom_Crop_alldata=="Wheat",]
mod_phen_sub<-mod_phen[mod_phen$PHENDom_Crop_alldata=="Wheat",]

## removes NAs in the phenotype, and all corresponding rows in model test
and train data frames
mod_frame_sub<-mod_frame_sub[!is.na(mod_phen_sub$PHENDom_yield_t_ha),]
test_frame_sub<-test_frame_sub[!is.na(test_phen_sub$PHENDom_yield_t_ha),]

mod_phen_sub<-mod_phen_sub[!is.na(mod_phen_sub$PHENDom_yield_t_ha),]
test_phen_sub<-test_phen_sub[!is.na(test_phen_sub$PHENDom_yield_t_ha),]

## attaches Yield to model frame
mod_frame_sub$Yield<-mod_phen_sub$PHENDom_yield_t_ha

## makes shorthand copies of phenotype
mod_Phen_Y<-mod_frame_sub$Yield
test_phen_Y<-test_phen_sub$PHENDom_yield_t_ha

## Builds an Rpart model

system.time(Wheat_rpart<-rpart(Yield~., data = mod_frame_sub,
                               control = rpart.control(minbucket = 25,
minsplit = 50, cp = 0.001, xval = 10)))


## measures the accuracy of pruned models (plotted), and non-pruned models
prune_wheat<-prune(Wheat_rpart, cp = 0.005)

cor.test(predict(prune_wheat, newdata = test_frame_sub),test_phen_Y)
rmserr(predict(prune_wheat, newdata = test_frame_sub),test_phen_Y)

cor.test(predict(Wheat_rpart, newdata = test_frame_sub),test_phen_Y)
rmserr(predict(Wheat_rpart, newdata = test_frame_sub),test_phen_Y)

## filters out +200DAS and postsowing data
mod_frame_Y<-mod_frame1[,-c(grep("_2[0-9]{2}$", colnames(mod_frame1)))]
mod_frame_Y<-mod_frame_Y[,-c(grep("PostSowing", colnames(mod_frame_Y),
ignore.case = T))]

## splits off 2018+ data (rescaled)
test_phen<-
Phen_data[c(mod_frame_Y$METADom_Year>=c(tail(unique(mod_frame_Y$METADom_Year),2)[1])),
]
mod_phen<-Phen_data[c(mod_frame_Y$METADom_Year<-
mod_frame_Y[c(mod_frame_Y$METADom_Year>=c(tail(unique(mod_frame_Y$METADom_Year),2)[1])),
]
mod_frame_sub<-mod_frame_Y[c(mod_frame_Y$METADom_Year<-
```

```
mod_frame_sub[mod_phen$PHENDom_Crop_alldata=="Canola",]
test_frame_sub<-test_frame_Y[test_phen$PHENDom_Crop_alldata=="Canola",]

test_phen_sub<-test_phen[test_phen$PHENDom_Crop_alldata=="Canola",]
mod_phen_sub<-mod_phen[mod_phen$PHENDom_Crop_alldata=="Canola",]

## removes NAs in the phenotype, and all corresponding rows in model test
and train data frames
mod_frame_sub<-mod_frame_sub[!is.na(mod_phen_sub$PHENDom_yield_t_ha),]
test_frame_sub<-test_frame_sub[!is.na(test_phen_sub$PHENDom_yield_t_ha),]

mod_phen_sub<-mod_phen_sub[!is.na(mod_phen_sub$PHENDom_yield_t_ha),]
test_phen_sub<-test_phen_sub[!is.na(test_phen_sub$PHENDom_yield_t_ha),]

## attaches Yield to model frame
mod_frame_sub$Yield<-mod_phen_sub$PHENDom_yield_t_ha

## makes shorthand copies of phenotype
mod_Phen_Y<-mod_frame_sub$Yield
test_phen_Y<-test_phen_sub$PHENDom_yield_t_ha

## Builds an Rpart model

system.time(Canola_rpart<-rpart(Yield~., data = mod_frame_sub,
                                control = rpart.control(minbucket = 25,
minsplit = 50, cp = 0.001, xval = 10)))

predict(Canola_rpart, data = test_frame_sub)

## measures the accuracy of pruned models (plotted), and non-pruned models
prune_canola<-prune(Canola_rpart, cp = 0.005)
cor.test(predict(prune_canola, newdata = test_frame_sub),test_phen_Y)
rmserr(predict(prune_canola, newdata = test_frame_sub),test_phen_Y)

cor.test(predict(Canola_rpart, newdata = test_frame_sub),test_phen_Y)
rmserr(predict(Canola_rpart, newdata = test_frame_sub),test_phen_Y)

####################################################

## Prediction at Time of Sowing (TOS)

####################################################

## registers cluster
peanutCluster<-makeCluster(detectCores()-2)
registerDoParallel(peanutCluster)

## loads data
mod_frame1<-readRDS("model_frames_and_imputations/mod_frame1.rds")

## separates phenotypes
Phen_data<-mod_frame1[,grep("PHENDom", colnames(mod_frame1))]
mod_frame1<-mod_frame1[,-c(grep("PHENDom", colnames(mod_frame1)))]

## attaches the target phenotype
mod_frame1$Yield<-Phen_data$PHENDom_yield_t_ha

## removes na data,"TRIAL ID" vectors (used for crossvalidation), adds
species
mod_frame_Y<-mod_frame1[!is.na(mod_frame1$Yield),]
mod_frame_Y<-mod_frame_Y[,-c(grep("Trial_ID", colnames(mod_frame_Y),
ignore.case = T))]

#mod_frame_Y<-data.frame(mod_frame_Y,
#                        Crop =
as.factor(P_frame$PHENDom_Crop[!is.na(P_frame$PHENDom_MEAN.YIELD)]))

## Filters data
mod_frame_Y<-mod_frame_Y[,-c(grep("_[0-9]{3}$", colnames(mod_frame_Y)))]
mod_frame_Y<-mod_frame_Y[,-c(grep("_[0-9]{2}$", colnames(mod_frame_Y)))]

mod_frame_Y<-mod_frame_Y[,-c(grep("PostSowing", colnames(mod_frame_Y),
ignore.case = T))]

## converts all chemical, fertiliser and soil tests with dates >=0
## to zero dose values
Chem_subvector<-
unlist(lapply(strsplit(colnames(mod_frame_Y)[grep("Chemicals_time_Chemicals_"
,

colnames(mod_frame_Y))], split = "time_Chemicals_"),tail,1))

for(i in 1:length(grep("Chemicals_time_Chemicals_",
colnames(mod_frame_Y)))){
  ## picks appropriate column name
  sub_index<-grep(c(colnames(mod_frame_Y)[grep(paste0(Chem_subvector[i],
"$"),
                                              colnames(mod_frame_Y))])[1],
colnames(mod_frame_Y))
  ## replaces doses after 100DAS with zero
  sub_vec<-mod_frame_Y[,c(sub_index[[1]])]
  sub_vec[mod_frame_Y[,c(grep("Chemicals_time_Chemicals_",
colnames(mod_frame_Y))[i])]>=0]<-0
  mod_frame_Y[,sub_index]<-sub_vec
  rm(sub_vec, sub_index)
  print(i)
}

## replaces the timings themselves with NAs
for(i in grep("Chemicals_time_Chemicals_", colnames(mod_frame_Y))){
  mod_frame_Y[which(mod_frame_Y[,i]>=0),i]<-0
}

Fert_subvector<-
unlist(lapply(strsplit(colnames(mod_frame_Y)[grep("METADom_Fertiliser_time_"
,

colnames(mod_frame_Y))], split = "Fertiliser_time_"),tail,1))

for(i in 1:length(grep("METADom_Fertiliser_time_",
colnames(mod_frame_Y)))){
  ## picks appropriate column name
  sub_index<-grep(c(colnames(mod_frame_Y)[grep(paste0(Fert_subvector[i],
"$"),
                                              colnames(mod_frame_Y))])[1],
colnames(mod_frame_Y))
  ## replaces doses after 100DAS with zero
  sub_vec<-mod_frame_Y[,c(sub_index[[1]])]
```

```
  sub_vec[mod_frame_Y[,c(grep("Fertiliser_time_",
colnames(mod_frame_Y))[i])]>=0]<-0
  mod_frame_Y[,sub_index]<-sub_vec
  ## replaces the timings themselves with NAs
  mod_frame_Y[,grep(paste0("Fertiliser_time_",Chem_subvector[i]),
colnames(mod_frame_Y))>=0]<-0
  rm(sub_vec, sub_index)
}

## replaces the timings themselves with NAs
for(i in grep("Fertiliser_time_", colnames(mod_frame_Y))){
  mod_frame_Y[which(mod_frame_Y[,i]>=0),i]<-0
}

## splits off 2018 data
test_frame_Y<-mod_frame_Y[mod_frame_Y$METADom_Year>=2018,]
mod_frame_Y<-mod_frame_Y[mod_frame_Y$METADom_Year<2018,]


## splits into wheat and canola only
mod_frame_C<-mod_frame_Y[mod_frame_Y$METADom_sub_cropCanola==1,]
mod_frame_W<-mod_frame_Y[mod_frame_Y$METADom_sub_cropWheat==1,]

test_frame_C<-test_frame_Y[test_frame_Y$METADom_sub_cropCanola==1,]
test_frame_W<-test_frame_Y[test_frame_Y$METADom_sub_cropWheat==1,]

## subsamples 100 experiments from the pre-2018 data
#test_frame_Y_site<-mod_frame_Y[c(Site_index_vec %in% Site_subset),]
#mod_frame_Y<-mod_frame_Y[!(Site_index_vec %in% Site_subset),]


system.time(Wheat_rpart_TOS<-rpart(Yield~., data = mod_frame_W,
                                    control = rpart.control(minbucket = 25,
minsplit = 50, cp = 0.001, xval = 10)))


system.time(Canola_rpart_TOS<-rpart(Yield~., data = mod_frame_C,
                                control = rpart.control(minbucket = 25,
minsplit = 50, cp = 0.001, xval = 10)))

## the canola model is OK, just
cor.test(predict(Canola_rpart_TOS, newdata =
test_frame_C),test_frame_C$Yield)
rmserr(predict(Canola_rpart_TOS, newdata =
test_frame_C),test_frame_C$Yield)

## The wheat model, good.
cor.test(predict(Wheat_rpart_TOS, newdata =
test_frame_W),test_frame_W$Yield)
rmserr(predict(Wheat_rpart_TOS, newdata = test_frame_W),test_frame_W$Yield)



########################################################################################
#
########################################################################################
#

## Figures

########################################################################################
#
########################################################################################
#

#devtools::install_github('thomasp85/gganimate')

require(ggplot2)
require(gganimate)
require(ggthemes)
require(leaflet)
require(viridis)
require(mapview)
require(umap)
require(pracma)
require(rpart.plot)

# webshot::install_phantomjs()
library(animation)
library(png)
library(htmlwidgets)
library(webshot)
library(ggmap)
require(gridExtra)
require(beeswarm)
require(rnaturalearth)
require(rnaturalearthdata)
require(ggspatial)
require(sf)

##########################################

## Prints the rpart model across all species as plain text

rpart_mod<-readRDS(file =
"model_frames_and_imputations/siteVariety_models/rpart_allTimes.rds")

rpart_mod

## gets the fraction of data that is satellite data
length(grep("SatDom",
names(rpart_mod$variable.importance)))/length(names(rpart_mod$variable.importance)
)

rpart_mod<-readRDS(file =
"model_frames_and_imputations/siteVariety_models/rpart_TOS.rds")

rpart_mod



####################

## Builds a site clustering based on BOM data, Satellite data, and combined

####################

## loads phenotype data
```

```
P_frame<-read.csv("model_frames_and_imputations/site_frame.csv", header =
T)
P_frame<-P_frame[,grep("PHENDom", colnames(P_frame))]

## just keeps the species, a few key variables, and yield
P_frame<-data.frame(Yield = P_frame$PHENDom_MEAN.YIELD,
                    Yield2 = P_frame$PHENDom_Pins_Site_Mean_Yield,
                    DTH = P_frame$PHENDom_Days_to_Harvest,
                    Crop = P_frame$PHENDom_Crop)

## loads the first random-sample imputation
Site_Modelframe<-read.csv(file =
"model_frames_and_imputations/site_imputations/random_sample_1.csv", header
= T)


## gets environment data, performs a umap clustering
sub_mat<-Site_Modelframe[,c(grep("BOMDom_", colnames(Site_Modelframe)),
                            grep("SatDom_", colnames(Site_Modelframe)))]

## trims data past 200 days after sowing, around the latest date you could
expect to harvest most crops
sub_mat<-sub_mat[,-c(grep("2[0-9]{2}$", colnames(sub_mat)))]
P_frame<-P_frame[!duplicated(paste(Site_Modelframe[,1],Site_Modelframe[,2],
sep = "_")),]
P_frame$Year<-
Site_Modelframe$METADom_Year[!duplicated(paste(Site_Modelframe[,1],Site_Modelframe[,2]
, sep = "_"))]
P_frame$JTS<-
Site_Modelframe$MANDom_JTS[!duplicated(paste(Site_Modelframe[,1],Site_Modelframe[,2]
, sep = "_"))]
P_frame$Lat<-
Site_Modelframe$BOMDom_Latitude[!duplicated(paste(Site_Modelframe[,1],Site_Modelframe[,2]
, sep = "_"))]
P_frame$Long<-
Site_Modelframe$BOMDom_Longitude[!duplicated(paste(Site_Modelframe[,1],Site_Modelframe[,2]
, sep = "_"))]
sub_mat<-sub_mat[!duplicated(paste(Site_Modelframe[,1],Site_Modelframe[,2],
sep = "_")),]
rownames(sub_mat)<-paste(sub_mat[,1],sub_mat[,2], sep = "_")

sub_mat<-as.matrix(sub_mat[,-c(1)])

## Captures principal components
system.time(env_PCA<-prcomp(sub_mat))


qplot(env_PCA$x[,2], env_PCA$x[,4], col = P_frame$Crop)


## and builds a UMAP
ENV_umap<-umap(sub_mat,config = umap.defaults, method = "naive")

umap_plot_E<-data.frame(ENV_umap$layout,P_frame)
colnames(umap_plot_E)[1:2]<-c("Dimension_1", "Dimension_2")

Umap_alldata_figure<-qplot(umap_plot_E$Dimension_1,umap_plot_E$Dimension_2,
                           col = c(umap_plot_E$Year),
                           xlim = c(-8,9), ylim = c(-8.5,7.5),
                           xlab  = "Dimension 1", ylab  =" Dimension 2",
                           alpha = I(0))+
  geom_point(size = 0.5)+
  scale_color_viridis("Year",c(umap_plot_E$Year), option  ="D")+
  theme_bw()+
  theme(legend.position = c(0.9,0.2))


ggsave(Umap_alldata_figure,file = "Publication_docs/Fig1c.tiff", device =
"tiff", height = 6, width = 6,dpi = 330)



ggplot(aes(Dimension_1,Dimension_2,col = Year), data = umap_plot_E,
       col = Year)+
  geom_point()

qplot(umap_plot_E$Dimension_1,umap_plot_E$Dimension_2,col =
umap_plot_E$Crop,
      xlim = c(-10,10), ylim = c(-8,8), alpha = I(0.5))+theme_bw()

## generates one for wheat only



ENV_Wheat_umap<-umap(sub_mat[which(P_frame$Crop=="Wheat"),],config =
umap.defaults, method = "naive")

umap_plot_E_Wheat<-
data.frame(ENV_Wheat_umap$layout,P_frame[which(P_frame$Crop=="Wheat"),])
colnames(umap_plot_E_Wheat)[1:2]<-c("Dimension_1", "Dimension_2")


qplot(umap_plot_E_Wheat$Dimension_2,umap_plot_E_Wheat$Dimension_1,
      col = umap_plot_E_Wheat$Long,
      ylim = c(-5,5), xlim = c(-8,8),
      alpha = I(0.5))+theme_bw()

Umap_alldata_figure_Wheat<-
qplot(umap_plot_E_Wheat$Dimension_2,umap_plot_E_Wheat$Dimension_1,
                                 col = c(umap_plot_E_Wheat$Long),
                                 ylim = c(-5,5), xlim = c(-8,8),
                                 xlab  = "", ylab  ="",
                                 alpha = I(0))+
  theme_bw()+
  geom_point(size = 1)+
  scale_color_viridis(c(umap_plot_E_Wheat$Yield2), discrete  =F)


## gets only the BOM data
sub_mat<-Site_Modelframe[,grep("BOMDom_", colnames(Site_Modelframe))]

## trims data past 200 days after sowing, around the latest date you could
expect to harvest most crops
sub_mat<-sub_mat[,-c(grep("2[0-9]{2}$", colnames(sub_mat)))]
P_frame<-P_frame[!duplicated(paste(Site_Modelframe[,1],Site_Modelframe[,2],
sep = "_")),]
P_frame$Year<-
Site_Modelframe$METADom_Year[!duplicated(paste(Site_Modelframe[,1],Site_Modelframe[,2]
```

```
, sep = "_"))]
P_frame$JTS<-
Site_Modelframe$MANDom_JTS[!duplicated(paste(Site_Modelframe[,1],Site_Modelframe[,2]
, sep = "_"))]
P_frame$Lat<-
Site_Modelframe$BOMDom_Latitude[!duplicated(paste(Site_Modelframe[,1],Site_Modelframe[,2]
, sep = "_"))]
P_frame$Long<-
Site_Modelframe$BOMDom_Longitude[!duplicated(paste(Site_Modelframe[,1],Site_Modelframe[,2]
, sep = "_"))]
sub_mat<-sub_mat[!duplicated(paste(Site_Modelframe[,1],Site_Modelframe[,2],
sep = "_")),]
rownames(sub_mat)<-paste(sub_mat[,1],sub_mat[,2], sep = "_")

sub_mat<-as.matrix(sub_mat[,-c(1)])

BOM_umap<-umap(sub_mat,config = umap.defaults, method = "naive")

umap_plot_BOM<-data.frame(BOM_umap$layout,P_frame)
colnames(umap_plot_BOM)[1:2]<-c("Dimension_1", "Dimension_2")

qplot(aes(Dimension_1,Dimension_2, data = umap_plot_BOM), xlim = c(0,35),
ylim = c(-250,250), col = umap_plot_BOM$Year)+
  geom_point(col = umap_plot_BOM$Year)

qplot(umap_plot_BOM$Dimension_1,umap_plot_BOM$Dimension_2, xlim = c(0,35),
ylim = c(-250,250), col = umap_plot_BOM$Year)

## now just the BOM figure
Umap_BOM_figure<-qplot(umap_plot_BOM$Dimension_2,umap_plot_BOM$Dimension_1,
                       col = c(umap_plot_BOM$Year),
                       ylim = c(-50,50), xlim = c(-80,80),
                       xlab  = "Dimension 2", ylab  =" Dimension 1",
                       alpha = I(0.5))+
  scale_color_viridis("Year",c(umap_plot_BOM$Year), option  ="B")+
  theme_bw()+
  theme(legend.position = c(0.9,0.5))


## gets only the Satellite data
sub_mat<-Site_Modelframe[,grep("SatDom_", colnames(Site_Modelframe))]

## trims data past 200 days after sowing, around the latest date you could
expect to harvest most crops
sub_mat<-sub_mat[,-c(grep("2[0-9]{2}$", colnames(sub_mat)))]
sub_mat<-sub_mat[!duplicated(paste(Site_Modelframe[,1],Site_Modelframe[,2],
sep = "_")),]
rownames(sub_mat)<-c(paste(Site_Modelframe[,1],Site_Modelframe[,2], sep =
"_")[!duplicated(paste(Site_Modelframe[,1],Site_Modelframe[,2], sep =
"_"))])

sub_mat<-as.matrix(sub_mat[,-c(1)])

Sat_umap<-umap(sub_mat,config = umap.defaults, method = "naive")

umap_plot_SAT<-data.frame(Sat_umap$layout,P_frame)
colnames(umap_plot_SAT)[1:2]<-c("Dimension_1", "Dimension_2")

qplot(umap_plot_SAT$Dimension_1,umap_plot_SAT$Dimension_2,
      xlim = c(-15,15),
      ylim = c(-12,12),
      alpha = I(0.5),
      col = c(umap_plot_SAT$Year))

#sub_mat<-sub_mat[,-c(grep("1[0-9]{2}$", colnames(sub_mat)))]

############################################################

## Fig. SXX Plots satellite data at a single location-year

############################################################

One_site<-list.files("single_site_plots")

xobj<-read.csv(paste0("single_site_plots/",One_site[[1]]), header = T)

out_frame_single<-data.frame(Calendar_date = xobj$calendar_date,
                             Value = xobj$value)

colnames(out_frame_single)[colnames(out_frame_single)=="Value"]<-
as.character(xobj$band[[1]])

for(i in 2:length(One_site)){

  xobj<-read.csv(paste0("single_site_plots/",One_site[[i]]), header = T)

  xobj2<-data.frame(Calendar_date = xobj$calendar_date,
                    Value = xobj$value)

  colnames(xobj2)[colnames(xobj2)=="Value"]<-as.character(xobj$band[[1]])

  out_frame_single<-merge(out_frame_single, xobj2, by.x = "Calendar_date",
by.y = "Calendar_date", all = T)
}

out_frame_single$Calendar_date<-as.Date(out_frame_single$Calendar_date)

#out_frame_single<-out_frame_single[order(out_frame_single),]

## weeds out the fire mask
out_frame_single_fire<-
out_frame_single[!is.na(out_frame_single$Burn_Date),]

out_frame_single<-out_frame_single[is.na(out_frame_single$Burn_Date),]

## trial was sown 21/12/2010 (14964) and sown on 14 Jun 2010 (14774)

## 50% flowering dates were at 292-295 Julian days, 14902-14905

tiff(filename  = "Publication_docs/Single_site_plot.tiff",
     width  =7.5, height = 7, units = "in", res  =330)

## plots raw values
par(mfrow = c(5,5), mar = c(3.1,4.1,0.1,1.1), bty = "n")


plot(out_frame_single$Calendar_date, out_frame_single$PsnNet_500m.x,
     xlab = "",
```

```
     ylab = "Net Photosynth", pch = 20, las = 2)

abline(v = 14774, col = "red")
abline(v = 14964, col = "orange")
abline(v = 14903, col = rgb(0,0,1,0.5), lwd  =3)

plot(out_frame_single$Calendar_date[out_frame_single$ET_500m.x<1000],
out_frame_single$ET_500m.x[out_frame_single$ET_500m.x<1000],
     xlab = "",
     ylab = "EvapoTrans", pch = 20, las = 2)

abline(v = 14774, col = "red")
abline(v = 14964, col = "orange")
abline(v = 14903, col = rgb(0,0,1,0.5), lwd  =3)


plot(out_frame_single$Calendar_date[out_frame_single$PET_500m.x<1000],
out_frame_single$PET_500m.x[out_frame_single$PET_500m.x<1000],
     xlab = "",
     ylab = "Pot. EvapoTrans", pch = 20, las = 2)

abline(v = 14774, col = "red")
abline(v = 14964, col = "orange")
abline(v = 14903, col = rgb(0,0,1,0.5), lwd  =3)


plot(out_frame_single$Calendar_date, out_frame_single$Fpar_500m,
     xlab = "",
     ylab = "FPAR", pch = 20, las = 2)

abline(v = 14774, col = "red")
abline(v = 14964, col = "orange")
abline(v = 14903, col = rgb(0,0,1,0.5), lwd  =3)

plot(out_frame_single$Calendar_date, out_frame_single$Lai_500m,
     xlab = "",
     ylab = "LAI", pch = 20, las = 2)

abline(v = 14774, col = "red")
abline(v = 14964, col = "orange")
abline(v = 14903, col = rgb(0,0,1,0.5), lwd  =3)


plot(out_frame_single$Calendar_date, out_frame_single$`250m_16_days_EVI`,
     xlab = "",
     ylab = "EVI", pch = 20, las = 2)

abline(v = 14774, col = "red")
abline(v = 14964, col = "orange")
abline(v = 14903, col = rgb(0,0,1,0.5), lwd  =3)

plot(out_frame_single$Calendar_date, out_frame_single$`250m_16_days_NDVI`,
     xlab = "",
     ylab = "NDVI", pch = 20, las = 2)

abline(v = 14774, col = "red")
abline(v = 14964, col = "orange")
abline(v = 14903, col = rgb(0,0,1,0.5), lwd  =3)


plot(out_frame_single$Calendar_date, out_frame_single$Gpp_500m.x,
     xlab = "",
     ylab = "GPP", pch = 20, las = 2)

abline(v = 14774, col = "red")
abline(v = 14964, col = "orange")
abline(v = 14903, col = rgb(0,0,1,0.5), lwd  =3)

plot(out_frame_single$Calendar_date[out_frame_single$LST_Day_1km>10],
c((out_frame_single$LST_Day_1km*0.02)-
273.15)[out_frame_single$LST_Day_1km>10],
     xlab = "",
     ylab = "LST day", pch = 20, las = 2)

abline(v = 14774, col = "red")
abline(v = 14964, col = "orange")
abline(v = 14903, col = rgb(0,0,1,0.5), lwd  =3)


plot(out_frame_single$Calendar_date[out_frame_single$LST_Night_1km>10],
c((out_frame_single$LST_Night_1km*0.02)-
273.15)[out_frame_single$LST_Night_1km>10],
     xlab = "",
     ylab = "LST night", pch = 20, las = 2)

abline(v = 14774, col = "red")
abline(v = 14964, col = "orange")
abline(v = 14903, col = rgb(0,0,1,0.5), lwd  =3)


plot(out_frame_single$Calendar_date, out_frame_single$Clear_sky_days,
     xlab = "",
     ylab = "Clear day", pch = 20, las = 2)

abline(v = 14774, col = "red")
abline(v = 14964, col = "orange")
abline(v = 14903, col = rgb(0,0,1,0.5), lwd  =3)

plot(out_frame_single$Calendar_date, out_frame_single$Clear_sky_days,
     xlab = "",
     ylab = "Clear Night", pch = 20, las = 2)

abline(v = 14774, col = "red")
abline(v = 14964, col = "orange")
abline(v = 14903, col = rgb(0,0,1,0.5), lwd  =3)

plot(out_frame_single$Calendar_date, out_frame_single$sur_refl_b01,
     xlab = "",
     ylab = "Surf_ref_wave01", pch = 20, ylim = c(250,2500), las = 2)

abline(v = 14774, col = "red")
abline(v = 14964, col = "orange")
abline(v = 14903, col = rgb(0,0,1,0.5), lwd  =3)

plot(out_frame_single$Calendar_date, out_frame_single$sur_refl_b02,
     xlab = "",
     ylab = "Surf_ref_wave02", pch = 20, las = 2)
```

```
abline(v = 14774, col = "red")
abline(v = 14964, col = "orange")
abline(v = 14903, col = rgb(0,0,1,0.5), lwd  =3)

plot(out_frame_single$Calendar_date, out_frame_single$sur_refl_b03,
     xlab = "",
     ylab = "Surf_ref_wave03", pch = 20, ylim = c(125,1250), las = 2)

abline(v = 14774, col = "red")
abline(v = 14964, col = "orange")
abline(v = 14903, col = rgb(0,0,1,0.5), lwd  =3)

plot(out_frame_single$Calendar_date, out_frame_single$sur_refl_b04,
     xlab = "",
     ylab = "Surf_ref_wave04", pch = 20, ylim = c(500,1500))

abline(v = 14774, col = "red")
abline(v = 14964, col = "orange")
abline(v = 14903, col = rgb(0,0,1,0.5), lwd  =3)

plot(out_frame_single$Calendar_date, out_frame_single$sur_refl_b05,
     xlab = "",
     ylab = "Surf_ref_wave05", pch = 20, las = 2)

abline(v = 14774, col = "red")
abline(v = 14964, col = "orange")
abline(v = 14903, col = rgb(0,0,1,0.5), lwd  =3)


plot(out_frame_single$Calendar_date, out_frame_single$sur_refl_b06,
     xlab = "",
     ylab = "Surf_ref_wave06", pch = 20, las = 2)

abline(v = 14774, col = "red")
abline(v = 14964, col = "orange")
abline(v = 14903, col = rgb(0,0,1,0.5), lwd  =3)

plot(out_frame_single$Calendar_date, out_frame_single$sur_refl_b07,
     xlab = "",
     ylab = "Surf_ref_wave07", pch = 20, las = 2)

abline(v = 14774, col = "red")
abline(v = 14964, col = "orange")
abline(v = 14903, col = rgb(0,0,1,0.5), lwd  =3)

#plot(out_frame_single$Calendar_date, out_frame_single$Emis_31,
#     xlab = "",
#     ylab = "Emis 31", pch = 20)

#abline(v = 14708, col = "red", lty = 3)

plot(out_frame_single$Calendar_date, out_frame_single$Emis_32,
     xlab = "",
     ylab = "Emis 32", pch = 20, las = 2)

abline(v = 14774, col = "red")
abline(v = 14964, col = "orange")
abline(v = 14903, col = rgb(0,0,1,0.5), lwd  =3)

plot(out_frame_single$Calendar_date,
out_frame_single$`250m_16_days_blue_reflectance`,
     xlab = "",
     ylab = "Blue", pch = 20, las = 2)

abline(v = 14774, col = "red")
abline(v = 14964, col = "orange")
abline(v = 14903, col = rgb(0,0,1,0.5), lwd  =3)


plot(out_frame_single$Calendar_date,
out_frame_single$`250m_16_days_MIR_reflectance`,
     xlab = "",
     ylab = "MIR", pch = 20, las = 2)

abline(v = 14774, col = "red")
abline(v = 14964, col = "orange")
abline(v = 14903, col = rgb(0,0,1,0.5), lwd  =3)

plot(out_frame_single$Calendar_date,
out_frame_single$`250m_16_days_NIR_reflectance`,
     xlab = "",
     ylab = "NIR", pch = 20, las = 2)

abline(v = 14774, col = "red")
abline(v = 14964, col = "orange")
abline(v = 14903, col = rgb(0,0,1,0.5), lwd  =3)

plot(out_frame_single$Calendar_date[out_frame_single$LE_500m.x<1000],
out_frame_single$LE_500m.x[out_frame_single$LE_500m.x<1000],
     xlab = "",
     ylab = "L.Heat flux", pch = 20, las = 2)

abline(v = 14774, col = "red")
abline(v = 14964, col = "orange")
abline(v = 14903, col = rgb(0,0,1,0.5), lwd  =3)

plot(out_frame_single_fire$Calendar_date, out_frame_single_fire$Burn_Date,
     xlab = "",
     ylab = "Fires", pch = 20, las = 2)

abline(v = 14774, col = "red")
abline(v = 14964, col = "orange")
abline(v = 14903, col = rgb(0,0,1,0.5), lwd  =3)

par(mfrow = c(1,1), mar = c(5.1,4.1,4.1,2.1))

dev.off()


## Splits into vegetation products and other

tiff(filename  = "Publication_docs/Single_site_plot_vegetation.tiff",
     width  =7.5, height = 4, units = "in", res  =330)

## plots raw values
par(mfrow = c(2,5), mar = c(3.1,4.1,2.1,1.1), bty = "n")

plot(out_frame_single$Calendar_date, out_frame_single$PsnNet_500m.x,
```

```
     xlab = "",
     ylim = c(0, 500),
     ylab = "Net Photosynthesis", pch = 20, las = 2)

mtext("a", adj = 0)
abline(v = 14774, col = "red")
abline(v = 14964, col = "orange")
abline(v = 14903, col = rgb(0,0,1,0.5), lwd  =3)

plot(out_frame_single$Calendar_date[out_frame_single$ET_500m.x<1000],
out_frame_single$ET_500m.x[out_frame_single$ET_500m.x<1000],
     xlab = "",
     ylim = c(0,210),
     ylab = "Evapotranspiration", pch = 20, las = 2)

mtext("b", adj = 0)
abline(v = 14774, col = "red")
abline(v = 14964, col = "orange")
abline(v = 14903, col = rgb(0,0,1,0.5), lwd  =3)


plot(out_frame_single$Calendar_date[out_frame_single$PET_500m.x<1000],
out_frame_single$PET_500m.x[out_frame_single$PET_500m.x<1000],
     xlab = "",
     ylim = c(0,680),
     ylab = "Potential Evapotrans.", pch = 20, las = 2)

mtext("c", adj = 0)
abline(v = 14774, col = "red")
abline(v = 14964, col = "orange")
abline(v = 14903, col = rgb(0,0,1,0.5), lwd  =3)


plot(out_frame_single$Calendar_date, out_frame_single$Fpar_500m,
     xlab = "",
     ylim = c(0,100),
     ylab = "FPAR", pch = 20, las = 2)

mtext("d", adj = 0)
abline(v = 14774, col = "red")
abline(v = 14964, col = "orange")
abline(v = 14903, col = rgb(0,0,1,0.5), lwd  =3)

plot(out_frame_single$Calendar_date, out_frame_single$Lai_500m,
     xlab = "",
     ylab = "Leaf Area Index", pch = 20, las = 2)

mtext("d", adj = 0)
abline(v = 14774, col = "red")
abline(v = 14964, col = "orange")
abline(v = 14903, col = rgb(0,0,1,0.5), lwd  =3)


plot(out_frame_single$Calendar_date, out_frame_single$`250m_16_days_EVI`,
     xlab = "",
     ylim = c(0,7200),
     ylab = "Enhanced Vegetation Index", pch = 20, las = 2)

mtext("f", adj = 0)
abline(v = 14774, col = "red")
abline(v = 14964, col = "orange")
abline(v = 14903, col = rgb(0,0,1,0.5), lwd  =3)

plot(out_frame_single$Calendar_date, out_frame_single$`250m_16_days_NDVI`,
     xlab = "",
     ylab = "NDVI", pch = 20, las = 2)

mtext("g", adj = 0)
abline(v = 14774, col = "red")
abline(v = 14964, col = "orange")
abline(v = 14903, col = rgb(0,0,1,0.5), lwd  =3)


plot(out_frame_single$Calendar_date, out_frame_single$Gpp_500m.x,
     xlab = "",
     ylim = c(0,560),
     ylab = "Gross Primary Productivity", pch = 20, las = 2)

mtext("h", adj = 0)
abline(v = 14774, col = "red")
abline(v = 14964, col = "orange")
abline(v = 14903, col = rgb(0,0,1,0.5), lwd  =3)


plot(out_frame_single$Calendar_date[out_frame_single$LE_500m.x<1000],
out_frame_single$LE_500m.x[out_frame_single$LE_500m.x<1000],
     xlab = "",
     ylim = c(0,480),
     ylab = "Latent Heat Flux", pch = 20, las = 2)

mtext("i", adj = 0)
abline(v = 14774, col = "red")
abline(v = 14964, col = "orange")
abline(v = 14903, col = rgb(0,0,1,0.5), lwd  =3)

plot(out_frame_single_fire$Calendar_date, out_frame_single_fire$Burn_Date,
     xlab = "",
     ylim = c(0,1),
     ylab = "Fires", pch = 20, las = 2)

mtext("j", adj = 0)
abline(v = 14774, col = "red")
abline(v = 14964, col = "orange")
abline(v = 14903, col = rgb(0,0,1,0.5), lwd  =3)

par(mfrow = c(1,1), mar = c(5.1,4.1,4.1,2.1))

dev.off()



## Also plots Net photosynthesis separately for Figure 1.

## trial was harvested 21/12/2010 (14964) and sown on 14 Jun 2010 (14774)

Photosynth_Fig1b<-qplot(out_frame_single$Calendar_date,
out_frame_single$PsnNet_500m.x/8,
                        xlab = "",
```

```
                         ylab = "Net Photosynthesis\n (kg C/mÂ² per day)",
alpha = I(0))+
  geom_smooth(col = "black", bg = "grey", span = 0.3)+
  geom_point(cex = 0.5)+
  theme_bw()+
  geom_vline(xintercept=14774, col = "red")+
  geom_vline(xintercept=14964, col = "orange")+
  geom_vline(xintercept = 14903, col = "lightblue", alpha = I(0.5), lwd =
3)+
  annotate("text", label = "Sowing", x = as.Date(14765), y = 350/8, colour
= "red", angle = 90)+
  annotate("text", label = "Anthesis", x = as.Date(14891), y = 50/8, colour
= "darkblue", angle = 90)+
  annotate("text", label = "Harvest", x = as.Date(14955), y = 350/8, colour
= "orange", angle = 90)


ggsave(Photosynth_Fig1b,file = "Publication_docs/Fig1b.tiff", device =
"tiff", height = 6, width = 5,dpi = 330)


Photosynth_Fig1b+theme(text = element_text(size=10),
                        axis.text = element_text(size=10))

#############################################################

## Figure 2. Makes a figure of ML model accuracy, by method, for yield

#############################################################

predobs_frame_unscaled<-readRDS(file =
"model_frames_and_imputations/siteVariety_models/accuracy_predobs_alldata1.rds"
)
predobs_frame_scaled<-readRDS(file =
"model_frames_and_imputations/siteVariety_models/accuracy_predobs_alldata2.rds"
)

bestfit_reg<-odregress(predobs_frame_unscaled[[3]][,1],
                        predobs_frame_unscaled[[3]]$Rrf_200)

## plots internal accuracy vs OOB accuracy
Fig_Rrf1<-qplot(predobs_frame_unscaled[[3]][,1],
jitter(predobs_frame_unscaled[[3]]$Rrf_200, factor =200),
                alpha = I(0),
                xlim = c(0,10), ylim = c(0,10),
                ylab = "Predicted Yield (t/Ha)", xlab = "Observed Yield
(t/Ha)")+
  geom_point(pch = 20, cex = 0.5, alpha = 0.6, col = "orange")+
  geom_abline(slope = 1, intercept = 0, col = "darkgrey")+
  geom_density2d()+
  geom_abline(slope = bestfit_reg$coeff[1], intercept =
bestfit_reg$coeff[2], col = "black")+
  theme_bw()+
  annotate("text", x = 4, y = 9, cex = 3, label = "RÂ² = 0.82; RMSE =
0.94")

cor.test(predobs_frame_unscaled[[3]][,1],
predobs_frame_unscaled[[3]]$Rrf_200)
rmserr(predobs_frame_unscaled[[3]][,1],
predobs_frame_unscaled[[3]]$Rrf_200)$rmse

bestfit_reg<-odregress(predobs_frame_unscaled[[2]][,1],
                        predobs_frame_unscaled[[2]]$Rrf_200)

Fig_Rrf2<-qplot(predobs_frame_unscaled[[2]][,1],
predobs_frame_unscaled[[2]]$Rrf_200,
                alpha = I(0),
                xlim = c(0,12), ylim = c(0,12),
                ylab = "Predicted Yield (t/Ha)", xlab = "Observed Yield
(t/Ha)")+
  geom_abline(slope = 1, intercept = 0, col = "darkgrey")+
  geom_hex(bins = 50)+
  geom_abline(slope = bestfit_reg$coeff[1], intercept =
bestfit_reg$coeff[2], col = "orange", alpha = I(0.8))+
  theme_bw()+
  scale_fill_viridis("Observations",option = "B", limits=c(0, 130))+
  theme(legend.position = "none")+
  annotate("text", x = 4, y = 9, cex = 3, label = "RÂ² = 0.70; RMSE =
1.24")

round(rmserr(predobs_frame_unscaled[[2]][,1],
predobs_frame_unscaled[[2]]$Rrf_200)$rmse, digits = 2)

bestfit_reg<-odregress(predobs_frame_unscaled[[3]][,1],
                        predobs_frame_unscaled[[3]]$rp_200)

## plots internal accuracy vs OOB accuracy
Fig_rp1<-qplot(predobs_frame_unscaled[[3]][,1],
jitter(predobs_frame_unscaled[[3]]$rp_200, factor =200),
               alpha = I(0),
               xlim = c(0,10), ylim = c(0,10),
               ylab = "Predicted Yield (t/Ha)", xlab = "Observed Yield
(t/Ha)")+
  geom_point(pch = 20, cex = 0.5, alpha = 0.6, col = "orange")+
  geom_abline(slope = 1, intercept = 0, col = "darkgrey")+
  geom_density2d()+
  geom_abline(slope = bestfit_reg$coeff[1], intercept =
bestfit_reg$coeff[2], col = "black")+
  theme_bw()+
  annotate("text", x = 4, y = 9, cex = 3, label = "RÂ² = 0.76; RMSE =
1.11")

cor.test(predobs_frame_unscaled[[3]][,1],predobs_frame_unscaled[[3]]$rp_200
)
round(rmserr(predobs_frame_unscaled[[3]][,1],predobs_frame_unscaled[[3]]$rp_200)$rmse
, digits = 2)

bestfit_reg<-odregress(predobs_frame_unscaled[[2]][,1],
                        predobs_frame_unscaled[[2]]$rp_200)

Fig_rp2<-qplot(predobs_frame_unscaled[[2]][,1],
predobs_frame_unscaled[[2]]$rp_200,
               alpha = I(0),
               xlim = c(0,12), ylim = c(0,12),
               ylab = "Predicted Yield (t/Ha)", xlab = "Observed Yield
(t/Ha)")+
  geom_abline(slope = 1, intercept = 0, col = "darkgrey")+
  geom_hex(bins = 50)+
```

```
  geom_abline(slope = bestfit_reg$coeff[1], intercept =
bestfit_reg$coeff[2], col = "orange", alpha = I(0.8))+
  theme_bw()+
  scale_fill_viridis("Observations",option = "B", limits=c(0, 130))+
  theme(legend.position = "none")+
  annotate("text", x = 4, y = 9, cex = 3, label = "RÂ² = 0.59, RMSE =
1.42")

round(rmserr(predobs_frame_unscaled[[2]][,1],
predobs_frame_unscaled[[2]]$rp_200)$rmse, digits= 2)

bestfit_reg<-odregress(predobs_frame_unscaled[[3]][,1],
                       predobs_frame_unscaled[[3]]$xvrf_200)

## plots internal accuracy vs OOB accuracy
Fig_xvrf1<-qplot(predobs_frame_unscaled[[3]][,1],
jitter(predobs_frame_unscaled[[3]]$xvrf_200, factor =200),
                 alpha = I(0),
                 xlim = c(0,10), ylim = c(0,10),
                 ylab = "Predicted Yield (t/Ha)", xlab = "Observed Yield
(t/Ha)")+
  geom_point(pch = 20, cex = 0.5, alpha = 0.6, col = "orange")+
  geom_abline(slope = 1, intercept = 0, col = "darkgrey")+
  geom_density2d()+
  geom_abline(slope = bestfit_reg$coeff[1], intercept =
bestfit_reg$coeff[2], col = "black")+
  theme_bw()+
  annotate("text", x = 4, y = 9, cex = 3, label = "RÂ² = 0.84; RMSE =
0.91")

cor.test(predobs_frame_unscaled[[3]][,1],
predobs_frame_unscaled[[3]]$xvrf_200)
round(rmserr(predobs_frame_unscaled[[3]][,1],predobs_frame_unscaled[[3]]$xvrf_200)$rmse
, digits = 2)

## the older design
qplot(predobs_frame_unscaled[[3]][,1],
predobs_frame_unscaled[[3]]$xvrf_200,
      alpha = I(0),
      xlim = c(0,10), ylim = c(0,10),
      ylab = "Predicted Yield (t/Ha)", xlab = "Observed Yield (t/Ha)")+
  geom_abline(slope = 1, intercept = 0, col = "darkgrey")+
  geom_hex(bins = 50)+
  geom_abline(slope = bestfit_reg$coeff[1], intercept =
bestfit_reg$coeff[2], col = "orange", alpha = I(0.8))+
  theme_bw()+
  scale_fill_viridis("Observations",option = "B", limits=c(0, 90))+
  theme(legend.position = "none")+
  annotate("text", x = 4, y = 9, cex = 3, label = "RÂ² = 0.84")

bestfit_reg<-odregress(predobs_frame_unscaled[[2]][,1],
                       predobs_frame_unscaled[[2]]$xvrf_200)

Fig_xvrf2<-qplot(predobs_frame_unscaled[[2]][,1],
predobs_frame_unscaled[[2]]$xvrf_200,
                 alpha = I(0),
                 xlim = c(0,12), ylim = c(0,12),
                 ylab = "Predicted Yield (t/Ha)", xlab = "Observed Yield
(t/Ha)")+
  geom_abline(slope = 1, intercept = 0, col = "darkgrey")+
  geom_hex(bins = 50)+
  geom_abline(slope = bestfit_reg$coeff[1], intercept =
bestfit_reg$coeff[2], color = "orange", alpha = I(0.8))+
  theme_bw()+
  scale_fill_viridis("Observations",option = "B", limits=c(0, 130))+
  theme(legend.position = "none")+
  annotate("text", x = 4, y = 9, cex = 3, label = "RÂ² = 0.69; RMSE =
1.23")

cor.test(predobs_frame_unscaled[[2]][,1],
predobs_frame_unscaled[[2]]$xvrf_200)
round(rmserr(predobs_frame_unscaled[[2]][,1],predobs_frame_unscaled[[2]]$xvrf_200)$rmse
, digits = 2)

bestfit_reg<-odregress(predobs_frame_scaled[[3]][,1],
                       predobs_frame_scaled[[3]]$xg_200)

## plots internal accuracy vs OOB accuracy
Fig_xg1<-qplot(predobs_frame_scaled[[3]][,1],
jitter(predobs_frame_scaled[[3]]$xg_200, factor =200),
               alpha = I(0),
               xlim = c(0,10), ylim = c(0,10),
               ylab = "Predicted Yield (t/Ha)", xlab = "Observed Yield
(t/Ha)")+
  geom_point(pch = 20, cex = 0.5, alpha = 0.6, col = "orange")+
  geom_abline(slope = 1, intercept = 0, col = "darkgrey")+
  geom_density2d()+
  geom_abline(slope = bestfit_reg$coeff[1], intercept =
bestfit_reg$coeff[2], col = "black")+
  theme_bw()+
  annotate("text", x = 4, y = 9, cex = 3, label = "RÂ² = 0.78; RMSE =
0.90")

cor.test(predobs_frame_scaled[[3]][,1], predobs_frame_scaled[[3]]$xg_200)
round(rmserr(predobs_frame_scaled[[3]][,1],predobs_frame_scaled[[3]]$xg_200)$rmse
, digits = 2)

bestfit_reg<-odregress(predobs_frame_scaled[[2]][,1],
                       predobs_frame_scaled[[2]]$xg_200)

Fig_xg2<-qplot(predobs_frame_scaled[[2]][,1],
predobs_frame_scaled[[2]]$xg_200,
               alpha = I(0),
               xlim = c(0,12), ylim = c(0,12),
               ylab = "Predicted Yield (t/Ha)", xlab = "Observed Yield
(t/Ha)")+
  geom_abline(slope = 1, intercept = 0, col = "darkgrey")+
  geom_hex(bins = 50)+
  geom_abline(slope = bestfit_reg$coeff[1], intercept =
bestfit_reg$coeff[2], color = "orange", alpha = I(0.8))+
  theme_bw()+
  scale_fill_viridis("Observations",option = "B", limits=c(0, 130))+
  theme(legend.position = "none")+
  annotate("text", x = 4, y = 9, cex = 3, label =  "RÂ² = 0.73; RMSE =
1.10")

cor.test(predobs_frame_scaled[[2]][,1], predobs_frame_scaled[[2]]$xg_200)
round(rmserr(predobs_frame_scaled[[2]][,1],predobs_frame_scaled[[2]]$xg_200)$rmse
```

```
, digits = 2)

bestfit_reg<-odregress(predobs_frame_scaled[[3]][,1],
                       predobs_frame_scaled[[3]]$lsvm_200)

## plots internal accuracy vs OOB accuracy
Fig_lsvm1<-qplot(predobs_frame_scaled[[3]][,1],
jitter(predobs_frame_scaled[[3]]$lsvm_200, factor =200),
                 alpha = I(0),
                 xlim = c(0,10), ylim = c(0,10),
                 ylab = "Predicted Yield (t/Ha)", xlab = "Observed Yield
(t/Ha)")+
  geom_point(pch = 20, cex = 0.5, alpha = 0.6, col = "orange")+
  geom_abline(slope = 1, intercept = 0, col = "darkgrey")+
  geom_density2d()+
  geom_abline(slope = bestfit_reg$coeff[1], intercept =
bestfit_reg$coeff[2], col = "black")+
  theme_bw()+
  annotate("text", x = 4, y = 9, cex = 3, label = "RÂ² = 0.68; RMSE =
1.09")

cor.test(predobs_frame_scaled[[3]][,1], predobs_frame_scaled[[3]]$lsvm_200)
round(rmserr(predobs_frame_scaled[[3]][,1],predobs_frame_scaled[[3]]$lsvm_200)$rmse
, digits = 2)

bestfit_reg<-odregress(predobs_frame_scaled[[2]][,1],
                       predobs_frame_scaled[[2]]$lsvm_200)

Fig_lsvm2<-qplot(predobs_frame_scaled[[2]][,1],
predobs_frame_scaled[[2]]$lsvm_200,
                 alpha = I(0),
                 xlim = c(0,12), ylim = c(0,12),
                 ylab = "Predicted Yield (t/Ha)", xlab = "Observed Yield
(t/Ha)")+
  geom_abline(slope = 1, intercept = 0, col = "darkgrey")+
  geom_hex(bins = 50)+
  geom_abline(slope = bestfit_reg$coeff[1], intercept =
bestfit_reg$coeff[2], col = "orange", alpha = I(0.8))+
  theme_bw()+
  scale_fill_viridis("Observations",option = "B", limits=c(0, 130))+
  theme(legend.position = "none")+
  annotate("text", x = 4, y = 9, cex = 3, label = "RÂ² = 0.45; RMSE =
1.43")

cor.test(predobs_frame_scaled[[2]][,1], predobs_frame_scaled[[2]]$lsvm_200)
round(rmserr(predobs_frame_scaled[[2]][,1],predobs_frame_scaled[[2]]$lsvm_200)$rmse
, digits = 2)

bestfit_reg<-odregress(predobs_frame_scaled[[3]][,1],
                       predobs_frame_scaled[[3]]$plsr_200)

## plots internal accuracy vs OOB accuracy
Fig_plsr1<-qplot(predobs_frame_scaled[[3]][,1],
jitter(predobs_frame_scaled[[3]]$plsr_200, factor =200), alpha = I(0),
                 xlim = c(0,10), ylim = c(0,10),
                 ylab = "Predicted Yield (t/Ha)", xlab = "Observed Yield
(t/Ha)")+
  geom_point(pch = 20, cex = 0.5, alpha = 0.6, col = "orange")+
  geom_abline(slope = 1, intercept = 0, col = "darkgrey")+
  geom_density2d()+
  geom_abline(slope = bestfit_reg$coeff[1], intercept =
bestfit_reg$coeff[2], col = "black")+
  theme_bw()+
  annotate("text", x = 4, y = 9, cex = 3, label = "RÂ² = 0.76; RMSE =
0.94")

cor.test(predobs_frame_scaled[[3]][,1], predobs_frame_scaled[[3]]$plsr_200)
round(rmserr(predobs_frame_scaled[[3]][,1],predobs_frame_scaled[[3]]$plsr_200)$rmse
, digits = 2)

#  scale_fill_viridis("Observations",option = "B", limits=c(0, 90))+
#  theme(legend.position = c(0.2, 0.7))+
#  annotate("text", x = 7.5, y = 0.5, label = "RÂ² = 0.76")

bestfit_reg<-odregress(predobs_frame_scaled[[2]][,1],
                       predobs_frame_scaled[[2]]$plsr_200)

Fig_plsr2<-qplot(predobs_frame_scaled[[2]][,1],
predobs_frame_scaled[[2]]$plsr_200,
                 alpha = I(0),
                 xlim = c(0,12), ylim = c(0,12),
                 ylab = "Predicted Yield (t/Ha)", xlab = "Observed Yield
(t/Ha)")+
  geom_abline(slope = 1, intercept = 0, col = "darkgrey")+
  geom_hex(bins = 50)+
  geom_abline(slope = bestfit_reg$coeff[1], intercept =
bestfit_reg$coeff[2], col = "orange")+
  theme_bw()+
  scale_fill_viridis("Observations",option = "B", limits=c(0, 130))+
  theme(legend.position = c(0.2, 0.7))+
  annotate("text",  x = 4, y = 9, cex = 3,  label = "RÂ² = 0.74; RMSE =
1.09")

cor.test(predobs_frame_scaled[[2]][,1], predobs_frame_scaled[[2]]$plsr_200)
round(rmserr(predobs_frame_scaled[[2]][,1],predobs_frame_scaled[[2]]$plsr_200)$rmse
, digits = 2)


## plots internal accuracy vs OOB accuracy
Fig_plsr1<-qplot(predobs_frame_scaled[[3]][,1],
jitter(predobs_frame_scaled[[3]]$plsr_200, factor =100),
                 alpha = I(0),
                 xlim = c(0,10), ylim = c(0,10),
                 ylab = "Predicted Yield (t/Ha)", xlab = "Observed Yield
(t/Ha)")+
  geom_point(pch = 20, cex = 0.5, alpha = 0.6, col = "orange")+
  geom_abline(slope = 1, intercept = 0, col = "darkgrey")+
  geom_density2d()+
  geom_abline(slope = bestfit_reg$coeff[1], intercept =
bestfit_reg$coeff[2], col = "black")+
  theme_bw()+
  annotate("text", x = 4, y = 9, cex = 3, label = "RÂ² = 0.76; RMSE =
0.94")

cor.test(predobs_frame_scaled[[3]][,1], predobs_frame_scaled[[3]]$plsr_200)
round(rmserr(predobs_frame_scaled[[3]][,1],predobs_frame_scaled[[3]]$plsr_200)$rmse
, digits = 2)
```

```
## too compact
tworow_acc<-grid.arrange(Fig_rp1, Fig_Rrf1,  Fig_xvrf1, Fig_xg1, Fig_lsvm1,
Fig_plsr1,
                         Fig_rp2, Fig_Rrf2, Fig_xvrf2, Fig_xg2, Fig_lsvm2,
Fig_plsr2, nrow = 2)

set.seed(8364)
## better...
tworow_acc<-grid.arrange(Fig_rp1+labs(title = "a"),
                         Fig_Rrf1+labs(title = "b"),
                         Fig_xvrf1+labs(title = "c"),
                         Fig_xg1+labs(title = "d"),
                         Fig_lsvm1+labs(title = "e"),
                         Fig_plsr1+labs(title = "f"),
                         nrow = 2)

ggsave(tworow_acc,file = "Publication_docs/Fig_2.tiff", device = "tiff",
height = 5, width = 7,dpi = 330)

# RÂ²
cor.test(predobs_frame_unscaled[[3]][,1],
predobs_frame_unscaled[[3]]$rp_200)
cor.test(predobs_frame_unscaled[[3]][,1],
predobs_frame_unscaled[[3]]$Rrf_200)
cor.test(predobs_frame_unscaled[[3]][,1],
predobs_frame_unscaled[[3]]$xvrf_200)
cor.test(predobs_frame_scaled[[3]][,1], predobs_frame_scaled[[3]]$lsvm_200)
cor.test(predobs_frame_scaled[[3]][,1], predobs_frame_scaled[[3]]$xg_200)
cor.test(predobs_frame_scaled[[3]][,1], predobs_frame_scaled[[3]]$plsr_200)

tworow_acc<-grid.arrange(Fig_rp2+labs(title = "a"), Fig_Rrf2+labs(title =
"b"),  Fig_xvrf2+labs(title = "c"),
                         Fig_xg2+labs(title = "d"), Fig_lsvm2+labs(title =
"e"), Fig_plsr2+labs(title = "f"),
                         nrow = 2)
## Saves figure 2
ggsave(tworow_acc,file = "Publication_docs/Fig_2_forward_projection.tiff",
device = "tiff", height = 6, width = 7.5,dpi = 330)


## prints R2 for missing trial site predictions
for(i in 1:ncol(predobs_frame_unscaled[[3]])){
  print(c(colnames(predobs_frame_unscaled[[3]])[i]))
  print(cor.test(predobs_frame_unscaled[[3]]$Yield,
predobs_frame_unscaled[[3]][,i])$estimate)}

for(i in 1:ncol(predobs_frame_scaled[[3]])){
  print(c(colnames(predobs_frame_scaled[[3]])[i]))
  print(cor.test(predobs_frame_scaled[[3]]$Yield,
predobs_frame_scaled[[3]][,i])$estimate)}


## prints R2 for annual forecasts
for(i in 1:ncol(predobs_frame_unscaled[[2]])){
  print(c(colnames(predobs_frame_unscaled[[2]])[i]))
  print(round(cor.test(predobs_frame_unscaled[[2]]$Yield,
predobs_frame_unscaled[[2]][,i])$estimate, digits  =2))}

for(i in 1:ncol(predobs_frame_scaled[[2]])){
  print(c(colnames(predobs_frame_scaled[[2]])[i]))
  print(round(cor.test(predobs_frame_scaled[[2]]$Yield,
predobs_frame_scaled[[2]][,i])$estimate, digits  =2))}


## prints R2 for (somewhat bullshit) random-sample holdout accuracy
for(i in 1:ncol(predobs_frame_unscaled[[1]])){
  print(c(colnames(predobs_frame_unscaled[[1]])[i]))
  print(round(cor.test(predobs_frame_unscaled[[1]]$Yield,
predobs_frame_unscaled[[1]][,i])$estimate, digits  =2))}

for(i in 1:ncol(predobs_frame_scaled[[1]])){
  print(c(colnames(predobs_frame_scaled[[1]])[i]))
  print(round(cor.test(predobs_frame_scaled[[1]]$Yield,
predobs_frame_scaled[[1]][,i])$estimate, digits  =2))}


## saves a different plot for the training schematic in supp fig 2

bestfit_reg<-odregress(predobs_frame_unscaled[[3]][,1],
                       predobs_frame_unscaled[[3]]$rp_200)

Schematic1<-qplot(predobs_frame_unscaled[[3]][,1],
jitter(predobs_frame_unscaled[[3]]$rp_200, factor =200),
                  alpha = I(0),
                  xlim = c(0,10), ylim = c(0,10),
                  ylab = "Predicted Yield (t/Ha)", xlab = "Observed Yield
(t/Ha)")+
  geom_point(pch = 20, cex = 0.5, alpha = 0.6, col = "orange")+
  geom_abline(slope = 1, intercept = 0, col = "darkgrey")+
  # geom_density2d()+
  geom_abline(slope = bestfit_reg$coeff[1], intercept =
bestfit_reg$coeff[2], col = "darkgreen", lwd = 2, alpha = 0.7)+
  theme_bw()+
  annotate("text", x = 7.5, y = 0.5, label = "RÂ² = 0.76")

cor.test(predobs_frame_unscaled[[3]][,1],predobs_frame_unscaled[[3]]$rp_200
)

bestfit_reg<-odregress(predobs_frame_unscaled[[2]][,1],
                       predobs_frame_unscaled[[2]]$rp_200)
## and the annual forecast data
Schematic2<-qplot(predobs_frame_unscaled[[2]][,1],
jitter(predobs_frame_unscaled[[2]]$rp_200, factor =200),
                  alpha = I(0),
                  xlim = c(0,10), ylim = c(0,10),
                  ylab = "Predicted Yield (t/Ha)", xlab = "Observed Yield
(t/Ha)")+
  geom_point(pch = 20, cex = 0.5,col = "lightblue")+
  geom_abline(slope = 1, intercept = 0, col = "darkgrey")+
  # geom_density2d()+
  geom_abline(slope = bestfit_reg$coeff[1], intercept =
bestfit_reg$coeff[2], col = "darkgreen", lwd = 2, alpha = 0.7)+
  theme_bw()+
  annotate("text", x = 7.5, y = 0.5, label = "RÂ² = 0.59")

cor.test(predobs_frame_unscaled[[2]][,1],predobs_frame_unscaled[[2]]$rp_200
)
```

```
Schematic3<-grid.arrange(Schematic1, Schematic2, nrow = 1)

ggsave(Schematic3,file = "Publication_docs/Fig_S2XX_Schematic.tiff", device
= "tiff", height = 4, width = 7.5,dpi = 330)


grid.arrange(Fig_rp1, Fig_Rrf1,  Fig_xvrf1,
             Fig_xg1, Fig_lsvm1, Fig_plsr1,
             Fig_rp2, Fig_Rrf2, Fig_xvrf2,
             Fig_xg2, Fig_lsvm2, Fig_plsr2, nrow = 4)

grid.arrange(Fig_rp1, Fig_Rrf1,  Fig_xvrf1,
             Fig_xg1, Fig_lsvm1, Fig_plsr1, nrow = 2)

rm(rf_mod, Rrf_mod, rpart_mod,xg_mod,lsvm_mod,pslr_mod, mod_frame3)


#xg_mod<-readRDS(file =
"model_frames_and_imputations/siteVariety_models/xgb_TOS.rds")
#lsvm_mod<-readRDS(file =
"model_frames_and_imputations/siteVariety_models/lsvm_TOS.rds")
#pslr_mod<-readRDS(file =
"model_frames_and_imputations/siteVariety_models/plsr_TOS.rds")

## and the yield data for prediction accuracy
mod_frame1<-readRDS(file = "model_frames_and_imputations/mod_frame1.rds")
beep(5)

peanutCluster<-makeCluster(detectCores()-2)
registerDoParallel(peanutCluster)

## first, separates phenotypes
Phen_data<-mod_frame1[,grep("PHENDom", colnames(mod_frame1))]
mod_frame1<-mod_frame1[,-c(grep("PHENDom", colnames(mod_frame1)))]

## produces a lookup table of phenotypes by species
Phen_summary_table<-data.frame(All = colSums(!is.na(Phen_data)),
                               Barley =
colSums(!is.na(Phen_data[Phen_data$PHENDom_Crop.Type=="Barley",])),
                               Canola =
colSums(!is.na(Phen_data[Phen_data$PHENDom_Crop.Type=="Canola",])),
                               Chickpea =
colSums(!is.na(Phen_data[Phen_data$PHENDom_Crop.Type=="Chickpea",])),
                               Faba =
colSums(!is.na(Phen_data[Phen_data$PHENDom_Crop.Type=="Faba Bean",])),
                               Lupin =
colSums(!is.na(Phen_data[Phen_data$PHENDom_Crop.Type=="Lupin",])),
                               Oat =
colSums(!is.na(Phen_data[Phen_data$PHENDom_Crop.Type=="Oat",])),
                               FieldPea =
colSums(!is.na(Phen_data[Phen_data$PHENDom_Crop.Type=="Field Pea",])),
                               Lentil =
colSums(!is.na(Phen_data[Phen_data$PHENDom_Crop.Type=="Lentil",])),
                               Triticale =
colSums(!is.na(Phen_data[Phen_data$PHENDom_Crop.Type=="Triticale",])),
                               Wheat =
colSums(!is.na(Phen_data[Phen_data$PHENDom_Crop.Type=="Wheat",])))

write.csv(Phen_summary_table, "Phen_summary_table.csv")
rm(Phen_summary_table)



############################################################

## Figure SXX. Imputation noise under different methods

############################################################

## Makes a figure of model accuracy


#(file =
"model_frames_and_imputations/siteVariety_models/xgb_allTimes.rds")


alldata<-read.csv(file = "data/constructed_csv_files/alldata.csv", header =
T)

results<-readRDS("data/results_list.rds")

## sets up dev

tiff(filename  = "Publication_docs/Imputation_variability.tiff",
     width  =7.5, height = 5, units = "in", res  =330)

par(bty = "n", mar = c(5.1, 7.1,4.1,2.1))

## Makes a figure of imputation noise introduced
boxplot(MI_test_random_imp[,9],MI_test_missforest_imp[,9],
        MI_test_random_imp[,1],MI_test_missforest_imp[,1],
        MI_test_random_imp[,3],MI_test_missforest_imp[,3],
        MI_test_random_imp[,5],MI_test_missforest_imp[,5],
        MI_test_random_imp[,7],MI_test_missforest_imp[,7],
        names = c("plsr random", "plsr",
                  "rpart random", "rpart",
                  "BCRF random", "BCRF",
                  "xgBoost random", "xgBoost",
                  "lsvm random", "lsvm"),
        las = 2, ylim = c(0,1),
        col = c("orange", "lightblue"), horizontal = T, xlab = "Forecast
accuracy (RÂ²)")

abline(v = seq(0,1,by = 0.2), lty = 3)
abline(h = seq(0,10), col = "grey", lty = 3)

dev.off()

par(mfrow = c(1,1), mar = c(5.1, 4.1,4.1,2.1))

rm(alldata)

############################################################

## Fig. SXX Plots some common non-yield phenotypes, with sample size and
BCRF accuracy

############################################################
```

```
CommonPhen_predictions<-readRDS(file =
"model_frames_and_imputations/site_models/CommonPhen_predictions.rds")

## [1] "PHENDom_Hectolitre_weight"       "PHENDom_Pct_below_2_0_mm"
"PHENDom_Protein"                 "PHENDom_Protein_4"
## [5] "PHENDom_Thousand_grain_weight"   "PHENDom_Early_Growth"
"PHENDom_yield_pct_of_average"    "PHENDom_zeroInflated_yield_t_ha"
## [9] "PHENDom_Flowering_QC"            "PHENDom_Glucosinolates"

## plots accuracy of phenotype models
namelist<-NULL
for(i in c(1:10)){
  bestfit_reg<-odregress(CommonPhen_predictions[[i]][,1],
                        CommonPhen_predictions[[i]][,2])

  namelist[[i]]<-ggplot(data.frame(Predicted =
CommonPhen_predictions[[i]][,2],
                                  Observed =
CommonPhen_predictions[[i]][,1]),
                        aes(x = Observed,y = Predicted)) +
    geom_abline(intercept = 0, slope = 1, col = "darkgrey")+
    geom_point(alpha = I(0.6), col = "orange", pch  =20, cex= 0.5)+
    geom_density2d()+
    geom_abline(slope = bestfit_reg$coeff[1], intercept =
bestfit_reg$coeff[2], col = "black", alpha = 0.7)+
    coord_fixed(xlim =
c(min(CommonPhen_predictions[[i]][,1]),max(CommonPhen_predictions[[i]][,1]))
,
                ylim =
c(min(CommonPhen_predictions[[i]][,1]),max(CommonPhen_predictions[[i]][,1]))
, ratio = 1)+
    annotate("text",
             x =
c(min(CommonPhen_predictions[[i]][,1])+c(max(CommonPhen_predictions[[i]][,1])/5))
,
             y = c(max(CommonPhen_predictions[[i]][,1])*0.9),
             label = paste0("RÂ² = ",
round(cor.test(CommonPhen_predictions[[i]][,1],

CommonPhen_predictions[[i]][,2])$estimate, digits = 2),
                            "\nN =
",length(CommonPhen_predictions[[i]][,1])))+
    theme_bw()
  ## prints target sample size
  print(length(CommonPhen_predictions[[i]][,1]))

}
names(CommonPhen_predictions)

# Canola    Chickpea  Faba Bean Lupin     Oat       Field Pea Lentil
Triticale Wheat
Pheno_grid<-grid.arrange(namelist[[1]]+labs(title = paste0("a   \n",
Common_phens[1])),
                         namelist[[2]]+labs(title = paste0("a   \n",
Common_phens[2])),
                         namelist[[3]]+labs(title = paste0("a   \n",
Common_phens[3])),
                         namelist[[4]]+labs(title = paste0("a   \n",
Common_phens[4])),
                         namelist[[5]]+labs(title = paste0("a   \n",
Common_phens[5])),
                         namelist[[6]]+labs(title = paste0("a   \n",
Common_phens[6])),
                         namelist[[7]]+labs(title = paste0("a   \n",
Common_phens[7])),
                         namelist[[8]]+labs(title = paste0("a   \n",
Common_phens[8])),
                         namelist[[9]]+labs(title = paste0("a   \n",
Common_phens[9])),
                         nrow =3)

Pheno_grid<-grid.arrange(namelist[[4]]+labs(title = "a   \nProtein %,
metadata"),
                         namelist[[8]]+labs(title = "b   \nFlowering Time
(Days)"),
                         namelist[[9]]+labs(title = "c   \nGlucosinolates
%"),
                         namelist[[1]]+labs(title = "d   \nHectolitre
Weight (Kg/hL)"),
                         namelist[[2]]+labs(title = "e   \n% Grain Below
2.0mm"),
                         namelist[[5]]+labs(title = "f   \n1000 Grain
Weight"),
                         nrow =2)

# Canola    Chickpea  Faba Bean Lupin     Oat       Field Pea Lentil
Triticale Wheat
#Pheno_grid<-grid.arrange(namelist[[4]]+labs(title = "a   \nProtein,
Metadata"),
#                         namelist[[6]]+labs(title = "b   \nFlowering Time
(Days)"),
#                         namelist[[7]]+labs(title = "c   \nGlucosinolate
Pct"),
#                         namelist[[1]]+labs(title = "d   \nHectolitre
Weight (Kg/hL)"),
#                         namelist[[2]]+labs(title = "e   \nPct Grain Below
2.0mm"),
#                         namelist[[5]]+labs(title = "f   \n1000 Grain
Weight"),
#                         nrow =2)

ggsave(Pheno_grid,file = "Publication_docs/Fig_SXX_Multiphenotypes.tiff",
device = "tiff", height = 5, width = 7.5,dpi = 330)

############################################################

## Fig. SXX Plots the yield rpart tree, with explanations

############################################################

Canola_rpart_TOS

plot(Wheat_rpart)
text(Wheat_rpart,cex= 0.4)

rpart.plot(Wheat_rpart)
```

```
rpart.plot(Canola_rpart)

rpart.rules(Canola_rpart)

prune_wheat<-prune(Wheat_rpart, cp = 0.005)
prune_canola<-prune(Canola_rpart, cp = 0.005)

rpart.plot(prune_wheat)
rpart.plot(prune_canola)

rpart.plot(Wheat_rpart_TOS)
rpart.rules(Wheat_rpart_TOS)

TOS_pruned<-prune(Wheat_rpart_TOS, cp = 0.02)
## NB gives better accuracy than unpruned model
cor.test(predict(TOS_pruned, newdata = test_frame_W),test_frame_W$Yield)
rmserr(predict(TOS_pruned, newdata = test_frame_W),test_frame_W$Yield)

TOS_pruned_C<-prune(Canola_rpart_TOS, cp = 0.01)

## NB accuracy suuuucks, r2= 0
cor.test(predict(TOS_pruned_C, newdata = test_frame_C),test_frame_C$Yield)
rmserr(predict(TOS_pruned_C, newdata = test_frame_C),test_frame_C$Yield)



## prnts a couple with no labels, for manual re-labeling

rpart.plot(prune_wheat,box.palette=scale_color_viridis_d(begin = 0, end =
7, option = "A"), fallen.leaves = F, branch = .4, tweak  =2)

rpart.plot(prune_wheat,box.palette=c("#ffffff","#faf398","#afd179","#68ba52","#09911a")
,
           fallen.leaves = F, branch = .4, tweak  =2, split.cex = 0.8,
branch.col = "darkgreen")

## Manual viridis type D pallette (Counterintuitive because >yellowing is
higher yield...)

rpart.plot(prune_wheat,box.palette=c("#26828EFF","#1F9E89FF","#35B779FF","#6DCD59FF")
,
           fallen.leaves = F, branch = .4, tweak  =2, split.cex = 0.8,
branch.col = "darkgreen")

## builds a dev.for the tree

tiff(filename  = "Publication_docs/FigS8_tree.tiff",
     width  =8, height = 8, units = "in", res  =330)

rpart.plot(prune_wheat,

box.palette=c("#faf398","#deffb3","#afd179","#68ba52","#09911a"),
           fallen.leaves = F, branch = .1, tweak  =2, split.cex = 0.75,
extra = 1, border.col = "darkgreen")

dev.off()


tiff(filename  = "Publication_docs/FigS9_tree.tiff",
     width  =5, height = 6, units = "in", res  =330)

rpart.plot(TOS_pruned,

box.palette=c("#faf398","#deffb3","#afd179","#68ba52","#09911a"),
           fallen.leaves = F, branch = .1, tweak  =1, split.cex = 0.75,
extra = 1, border.col = "darkgreen")

dev.off()


tiff(filename  = "Publication_docs/FigS10_tree.tiff",
     width  =8, height = 8, units = "in", res  =330)

rpart.plot(prune_canola,

box.palette=c("#faf398","#deffb3","#afd179","#68ba52","#09911a"),
           fallen.leaves = F, branch = .1, tweak  =2, split.cex = 0.75,
extra = 1, border.col = "darkgreen")

dev.off()

## And another for the TOS tree

tiff(filename  = "Publication_docs/FigS9_TOS_tree.tiff",
     width  =7.5, height = 6, units = "in", res  =330)

rpart.plot(TOS_pruned,

box.palette=c("#faf398","#deffb3","#afd179","#68ba52","#09911a"),
           fallen.leaves = F, branch = .4, tweak  =2, split.cex = 0.75,
extra = 1, split.col = "white")

dev.off()


rpart.plot(Wheat_rpart,

box.palette=c("#faf398","#deffb3","#afd179","#68ba52","#09911a"),
           fallen.leaves = F, branch = .4, tweak  =2, split.cex = 0.9,
extra = 1)


rpart.plot(prune_wheat,box.palette=c("YlGn"), fallen.leaves = F, branch =
.4, tweak  =2)



par(mfrow = c(2,1))

rpart.plot(prune_canola, cex = 0.4, box.palette="Greens", fallen.leaves =
F)

rpart.plot(prune_wheat, cex = 0.4, box.palette="Greens", fallen.leaves = F)

par(mfrow =c(1,1))

rpart.plot(prune_wheat, cex = 0.4, box.palette="Greens", fallen.leaves = F)
```

```
rpart.plot(Wheat_rpart_TOS, cex = 0.4, box.palette="Greens", fallen.leaves
= F)

rpart.plot(TOS_pruned, cex = 0.4, box.palette="Greens", fallen.leaves = F)

par(mfrow =c(1,1))

par(mfrow =c(2,1))

rpart.plot(prune_wheat, cex = 0.4, box.palette="Greens", fallen.leaves = F)

rpart.plot(TOS_pruned, cex = 0.4, box.palette="Greens", fallen.leaves = F)

par(mfrow =c(1,1))


Canola_rules<-rpart.rules(prune_canola)


rpart_mod<-readRDS(file =
"model_frames_and_imputations/siteVariety_models/rpart_allTimes.rds")

rpart_mod

tiff(filename  = "Publication_docs/unlabeled_tree.tiff",
     width  =7.5, height = 5, units = "in", res  =330)
plot(rpart_mod)
## saves this image, without text, for manual color-coding in figure
dev.off()

## writes the actual info, for manual coding
tiff(filename  = "Publication_docs/labeled_tree.tiff",
     width  =7.5, height = 5, units = "in", res  =1500)
plot(rpart_mod)
text(rpart_mod, cex = 0.05, col = "orange")
dev.off()

## prunes it down somewhat, to plot the top-level complexity
pruned_rp<-prune(rpart_mod, cp = 0.005)

plot(pruned_rp)
text(pruned_rp, cex = 0.5)
pruned_rp$cptable

## this accounts for 58% of yield

## prunes it down somewhat, to plot the top-level complexity
pruned_rp<-prune(rpart_mod, cp = 0.01)

plot(pruned_rp)
text(pruned_rp, cex = 0.5)

## this tree accounts for half the variation in yield across sites &
species

## this model accounts for


pruned_rp<-prune(rpart_mod, cp = 0.1)


############################################################

## Figure SXX plots wheat yield against frost severity

############################################################


mod_frame1<-readRDS("model_frames_and_imputations/mod_frame1.rds")

WheatYields<-data.frame(Yield = mod_frame1$PHENDom_yield_t_ha,
                        mod_frame1$ENVDom_Frost_Damage_score,
                        MinTemp =
mod_frame1$BOMDom_min_temperature_min_110,
                        MinTemp2 =
mod_frame1$BOMDom_min_temperature_min_120,
                        MinTemp3 =
mod_frame1$BOMDom_min_temperature_min_130,
                        MinTemp4 =
mod_frame1$BOMDom_min_temperature_min_140,
                        ZeroYield =
mod_frame1$PHENDom_zeroInflated_yield_t_ha)

WheatYields<-WheatYields[mod_frame1$METADom_sub_cropWheat==1,]

WheatYields$FlowerMinT<-unlist(lapply(seq(1:nrow(WheatYields)), function(i)
min(WheatYields[i,c(3:6)])))

Temp_anomaly<-ggplot(aes(FlowerMinT, Yield),data = WheatYields,
                     xlim = c(-10,20))+
  # geom_point(col = "orange", alpha = I(0.5))+
  geom_hex(bins = 50)+
  scale_fill_viridis("Observed Yields", option = "A")+
  #  geom_density2d()+
  geom_smooth(col = "lightblue")+
  geom_vline(xintercept = 0, col = "white", lty = 3)+
  geom_hline(yintercept = mean(WheatYields$Yield, na.rm = T), col =
"white", lty = 3)+
  theme_bw()+
  theme(legend.position = c(0.75,0.75))+
  labs(x = "Minimum Temperature (Deg. C) 110-140DAS",y = "Yield (t/Ha)")


## the sample size?
cor.test(WheatYields$FlowerMinT, WheatYields$Yield)
sum(!is.na(WheatYields$FlowerMinT+WheatYields$Yield))

## Saves this as a supp figure
ggsave(Temp_anomaly,file = "Publication_docs/Fig_SXX_FloweringTemps.tiff",
       device = "tiff", height = 7, width = 7.5,dpi = 330)

rm(Temp_anomaly, WheatYields)

############################################################

## Figure 3XX. Gains in rolling forecast accuracy, w. nonstationarity vibes
```

```
##########################################################

forecast_frame_W<-readRDS(file =
"model_frames_and_imputations/siteVariety_models/Annual_rolling_Wheat_PredObs.rds"
)
forecast_frame_C<-readRDS(file =
"model_frames_and_imputations/siteVariety_models/Annual_rolling_Canola_PredObs.rds"
)

mod_frame1<-readRDS("model_frames_and_imputations/mod_frame1.rds")

## the mean yields
mean_wheat<-
aggregate(mod_frame1$PHENDom_yield_t_ha[mod_frame1$METADom_sub_cropWheat>0]
, mean, by =
list(mod_frame1$METADom_Year[mod_frame1$METADom_sub_cropWheat>0]), na.rm =
T)
mean_canola<-
aggregate(mod_frame1$PHENDom_yield_t_ha[mod_frame1$METADom_sub_cropCanola>0]
, mean, by =
list(mod_frame1$METADom_Year[mod_frame1$METADom_sub_cropCanola>0]), na.rm =
T)

## the variance in yields
var_wheat<-
aggregate(mod_frame1$PHENDom_yield_t_ha[mod_frame1$METADom_sub_cropWheat>0]
, var, by =
list(mod_frame1$METADom_Year[mod_frame1$METADom_sub_cropWheat>0]), na.rm =
T)
var_canola<-
aggregate(mod_frame1$PHENDom_yield_t_ha[mod_frame1$METADom_sub_cropCanola>0]
, var, by =
list(mod_frame1$METADom_Year[mod_frame1$METADom_sub_cropCanola>0]), na.rm =
T)

## sets up dev

tiff(filename  = "Publication_docs/Fig3.tiff",
     width  =7.5, height = 6.5, units = "in", res  =330)

## change as appropriate
Year_seq<-c(2009:2018)

#par(mfrow = c(1,2), bty = "n")

layout.matrix <- matrix(c(1,2,3,4), nrow = 2, ncol = 2)

layout(mat = layout.matrix,
       heights = c(2, 1), # Heights of the two rows
       widths = c(2, 2))

par(mar = c(0.1,4.1,2.1,2.1), bty = "n")

plot(Year_seq-0.1,
     forecast_frame_W[,3], type = "l", col = "grey",
     xlim = c(2008.5, 2018.5),
     ylim = c(-0.1,0.85),
     ylab = "Forecast accuracy RÂ²",
     xlab = "", las  =2, axes = F)
axis(2)

abline(h = seq(-0.2,0.8, by = 0.2), lty = 3, col = "grey")
abline(v = seq(2009,2018, by = 2), lty = 3, col = "grey")

points(Year_seq-0.1,forecast_frame_W[,3], pch = 20)
## adds CIs
for(i in 1:length(Year_seq)){
  points(c(Year_seq[[i]]-0.1,Year_seq[[i]]-0.1),
c(forecast_frame_W[i,4],forecast_frame_W[i,5]),
         type = "l")
}

## now lsvms
points(Year_seq,
       forecast_frame_W[,6], type = "l",
       col = "orange")
points(Year_seq,forecast_frame_W[,6], pch = 20, col = "orange")
## adds CIs
for(i in 1:length(Year_seq)){
  points(c(Year_seq[[i]],Year_seq[[i]]),
c(forecast_frame_W[i,7],forecast_frame_W[i,8]),
         type = "l", col = "orange")
}

## now PLSR
points(Year_seq+0.1,
       forecast_frame_W[,9], type = "l",
       col = "lightblue")
points(Year_seq+0.1,forecast_frame_W[,9], pch = 20, col = "lightblue")
## adds CIs
for(i in 1:length(Year_seq)){
  points(c(Year_seq[[i]]+0.1,Year_seq[[i]]+0.1),
c(forecast_frame_W[i,10],forecast_frame_W[i,11]),
         type = "l", col = "lightblue")
}
mtext("a", adj = 0)


par(mar = c(5.1,4.1,0.1,2.1))

boxplot(mod_frame1$PHENDom_yield_t_ha[which(mod_frame1$METADom_sub_cropWheat>
0 &
mod_frame1$METADom_Year>2008)]~mod_frame1$METADom_Year[which(mod_frame1$METADom_sub_cropWheat>
0 & mod_frame1$METADom_Year>2008)],
        las = 2, pch = "", ylim = c(0,8.5), ylab = "Yield (t/Ha)", xlab =
"Year")

par(mar = c(0.1,4.1,2.1,2.1))

plot(Year_seq-0.1,
     forecast_frame_C[,3], type = "l", col = "grey",
     xlim = c(2008.5, 2018.5),
     ylim = c(-0.1,0.85),
     ylab = "Forecast accuracy RÂ²",
     xlab = "", las = 2, axes = F)

axis(2)
```

```
abline(h = seq(-0.2,0.8, by = 0.2), lty = 3, col = "grey")
abline(v = seq(2009,2018, by = 2), lty = 3, col = "grey")

points(Year_seq-0.1,forecast_frame_C[,3], pch = 20)
## adds CIs
for(i in 1:length(Year_seq)){
  points(c(Year_seq[[i]]-0.1,Year_seq[[i]]-0.1),
c(forecast_frame_C[i,4],forecast_frame_C[i,5]),
         type = "l")
}

## now lsvms
points(Year_seq,
       forecast_frame_C[,6], type = "l",
       col = "orange")
points(Year_seq,forecast_frame_C[,6], pch = 20, col = "orange")
## adds CIs
for(i in 1:length(Year_seq)){
  points(c(Year_seq[[i]],Year_seq[[i]]),
c(forecast_frame_C[i,7],forecast_frame_C[i,8]),
         type = "l", col = "orange")
}

## now PLSR
points(Year_seq+0.1,
       forecast_frame_C[,9], type = "l",
       col = "lightblue")
points(Year_seq+0.1,forecast_frame_C[,9], pch = 20, col = "lightblue")
## adds CIs
for(i in 1:length(Year_seq)){
  points(c(Year_seq[[i]]+0.1,Year_seq[[i]]+0.1),
c(forecast_frame_C[i,10],forecast_frame_C[i,11]),
         type = "l", col = "lightblue")
}
mtext("b", adj = 0)


par(mar = c(5.1,4.1,0.1,2.1))

boxplot(mod_frame1$PHENDom_yield_t_ha[which(mod_frame1$METADom_sub_cropCanola>
0 &
mod_frame1$METADom_Year>2008)]~mod_frame1$METADom_Year[which(mod_frame1$METADom_sub_cropCanola>
0 & mod_frame1$METADom_Year>2008)],
        las = 2, pch = "", ylim = c(0,5), ylab = "Yield (t/Ha)", xlab =
"Year")

par(mfrow = c(1,1))

dev.off()

par(mar = c(5.1,4.1,2.1,2.1))

## plots the deltas (all are correlated, but data is super limited)

splom(cbind(diff(forecast_frame_W[,3]),diff(forecast_frame_W[,6]),diff(forecast_frame_W[,9]))
)
splom(cbind(diff(forecast_frame_C[,3]),diff(forecast_frame_C[,6]),diff(forecast_frame_C[,9]))
)

cor.test(diff(forecast_frame_W[,3]),diff(forecast_frame_W[,6]))
cor.test(diff(forecast_frame_W[,3]),diff(forecast_frame_W[,9]))
cor.test(diff(forecast_frame_W[,6]),diff(forecast_frame_W[,9]))

cor.test(diff(forecast_frame_C[,3]),diff(forecast_frame_C[,6]))
cor.test(diff(forecast_frame_C[,3]),diff(forecast_frame_C[,9]))
cor.test(diff(forecast_frame_C[,6]),diff(forecast_frame_C[,9]))

## mean rsq
c(0.782853+0.9316459+0.7910698)/3

##########################################################

## Fig. S3XX the gain in model accuracy throughout the season

##########################################################

forecast_frame_W<-readRDS(file =
"model_frames_and_imputations/siteVariety_models/rolling_Wheat_PredObs.rds"
)
forecast_frame_C<-readRDS(file =
"model_frames_and_imputations/siteVariety_models/rolling_Canola_PredObs.rds"
)

## plot the gain in model accuracy through the season

## black are Naive random forests, blue are plsr models, orange is linear
SVMs

tiff(filename  = "Publication_docs/Rolling forecasts.tiff",
     width  =7.5, height = 6, units = "in", res  =330)

par(mfrow = c(1,2), bty = "n")
plot(seq(1, 190, by = 10),forecast_frame_W[,1], type = "l",
     xlim = c(0,200), ylim = c(0,0.8),
     ylab = "Forecast accuracy RÂ²",
     xlab = "Days after Sowing")
points(seq(1, 190, by = 10),forecast_frame_W[,1], pch = 20)

points(seq(1, 190, by = 10),forecast_frame_W[,2], pch = 20, col = "orange")
points(seq(1, 190, by = 10),forecast_frame_W[,2], type = "l", col =
"orange")

points(seq(1, 190, by = 10),forecast_frame_W[,3], pch = 20, col =
"lightblue")
points(seq(1, 190, by = 10),forecast_frame_W[,3], type = "l", col =
"lightblue")

abline(h = seq(-1,1, by = 0.2), lty = 3)
mtext("a", adj = 0)


plot(seq(1, 190, by = 10),forecast_frame_C[,1], type = "l",
     xlim = c(0,200), ylim = c(0,0.8),
     ylab = "Forecast accuracy RÂ²",
     xlab = "Days after Sowing")
points(seq(1, 190, by = 10),forecast_frame_C[,1], pch = 20)

points(seq(1, 190, by = 10),forecast_frame_C[,2], pch = 20, col = "orange")
```

```
points(seq(1, 190, by = 10),forecast_frame_C[,2], type = "l", col =
"orange")

points(seq(1, 190, by = 10),forecast_frame_C[,3], pch = 20, col =
"lightblue")
points(seq(1, 190, by = 10),forecast_frame_C[,3], type = "l", col =
"lightblue")

abline(h = seq(-1,1, by = 0.2), lty = 3)
mtext("b", adj = 0)

par(mfrow = c(1,1))
dev.off()

rm(forecast_frame_W, forecast_frame_C)

############################################################

## Figure 5 Plots reproducibility of variable importance scores

############################################################

rpart_mod<-readRDS(file =
"model_frames_and_imputations/siteVariety_models/rpart_allTimes.rds")
rf_mod<-readRDS(file =
"model_frames_and_imputations/siteVariety_models/bcrf_allTimes.rds")
plsr_mod<-readRDS(file =
"model_frames_and_imputations/siteVariety_models/plsr_allTimes.rds")

imp_frame_comparison<-data.frame(varNames =
names(rpart_mod$variable.importance),
                                 rpart_imp = rpart_mod$variable.importance)
## merges on rf importance
imp_frame_comparison<-data.frame(imp_frame_comparison,
                                 rf_imp =
rf_mod$importance[match(imp_frame_comparison$varNames,rownames(rf_mod$importance))]
)

## gets the coefficients for important features under PLSR, rescaled to 0-1
p_coefs<-plsr_mod$coefficients/sum(sapply(plsr_mod$coefficients, abs))

## merges on plsr importance
imp_frame_comparison<-data.frame(imp_frame_comparison,
                                 plsr_imp =
p_coefs[match(imp_frame_comparison$varNames,rownames(p_coefs))])
#require(rminer)
#lsvm_imp<-Importance(lsvm_mod)

## the most important 100 plsr weights
tail(imp_frame_comparison[order(imp_frame_comparison$plsr_imp, na.last =
F),], 100)

qplot(c(imp_frame_comparison$rpart_imp), c(imp_frame_comparison$rf_imp))
qplot(c(imp_frame_comparison$plsr_imp), c(imp_frame_comparison$rf_imp))
qplot(c(imp_frame_comparison$plsr_imp), c(imp_frame_comparison$rpart_imp),
col = c(imp_frame_comparison$rf_imp))


imp_frame_comparison2<-imp_frame_comparison
imp_frame_comparison2$plsr_imp<-abs(imp_frame_comparison$plsr_imp)

imp_frame_comparison2[is.na(imp_frame_comparison2)]<-0

Imp_plot1<-qplot(plsr_imp, rf_imp, data = imp_frame_comparison2, alpha =
I(0),
                 ylab = "BCRF Importance", xlab = "PLSR Coefficient
Absolute Values")+
  geom_point(aes(y = rf_imp, x = plsr_imp), cex = 0.4, pch  =20)+
  ## adds the Latent Heat predictors as red circles
  geom_point(aes(y = rf_imp, x = plsr_imp),data =
imp_frame_comparison2[grep("Latent_heat",
imp_frame_comparison2$varNames),],
             col = "#ff0066", pch  =20, alpha = I(0.7))+
  ## adds the cumulative rainfall as blue droplets
  geom_point(aes(y = rf_imp, x = plsr_imp),data =
imp_frame_comparison2[grep("BOMDom_rainfall_csum_",
imp_frame_comparison2$varNames),],
             col = "lightblue", alpha = I(0.7), cex = 1.5)+
  ## adds the FPAR as a green circle
  geom_point(aes(y = rf_imp, x = plsr_imp),data =
imp_frame_comparison2[grep("SatDom_FPAR",
imp_frame_comparison2$varNames),],
             col = "darkgreen", alpha = I(0.7), pch = 20)+
  ## adds the EVI as a green open square
  geom_point(aes(y = rf_imp, x = plsr_imp),data =
imp_frame_comparison2[grep("SatDom_EVI", imp_frame_comparison2$varNames),],
             col = "darkgreen", alpha = I(0.7), pch = 20)+
  ## adds the NDVI as a green open triangle
  geom_point(aes(y = rf_imp, x = plsr_imp),data =
imp_frame_comparison2[grep("SatDom_NDVI",
imp_frame_comparison2$varNames),],
             col = "darkgreen", alpha = I(0.7), pch = 20)+
  ##Direct labels some points
  geom_text(aes(y = rf_imp, x = plsr_imp),data =
imp_frame_comparison2[c(1),], label = "Dicot t/f", nudge_x = 0.0000017,
nudge_y = 0, size=3)+
  geom_point(aes(y = rf_imp, x = plsr_imp),data =
imp_frame_comparison2[c(1),], pch  =20)+
  geom_text(aes(y = rf_imp, x = plsr_imp),data =
imp_frame_comparison2[c(9),], label = "Canola t/f", nudge_x = 0.0000017,
nudge_y = 0, size=3)+
  geom_point(aes(y = rf_imp, x = plsr_imp),data =
imp_frame_comparison2[c(9),], pch  =20)+
  geom_text(aes(y = rf_imp, x = plsr_imp),data =
imp_frame_comparison2[c(8),], label = "Clethodim", nudge_x = -0.0000023,
nudge_y = 0, size=3)+
  geom_point(aes(y = rf_imp, x = plsr_imp),data =
imp_frame_comparison2[c(8),], pch  =20)+
  # geom_text(aes(y = rf_imp, x = plsr_imp),data =
imp_frame_comparison2[c(10),], label = "Fabaceae t/f", nudge_x = 0.0000017,
nudge_y = 0, size=3)+
  #  geom_point(aes(y = rf_imp, x = plsr_imp),data =
imp_frame_comparison2[c(10),], pch  =20)+
  geom_text(aes(y = rf_imp, x = plsr_imp),data =
imp_frame_comparison2[c(11),], label = "Applied Sulphur", nudge_x =
0.0000027, nudge_y = 0, size=3)+
  geom_point(aes(y = rf_imp, x = plsr_imp),data =
imp_frame_comparison2[c(11),], pch  =20)+
```

```
  geom_text(aes(y = rf_imp, x = plsr_imp),data =
imp_frame_comparison2[c(13),], label = "Wheat t/f", nudge_x = 0.0000017,
nudge_y = 0, size=3)+
  geom_point(aes(y = rf_imp, x = plsr_imp),data =
imp_frame_comparison2[c(13),], pch  =20)+
  geom_point(aes(y = rf_imp, x = plsr_imp),data =
imp_frame_comparison2[c(13),], pch  =20)+
  theme_bw()


Imp_plot2<-qplot(plsr_imp, rpart_imp, data = imp_frame_comparison2, alpha =
I(0),
                 ylab = "RPRM Importance", xlab = "PLSR Coefficient
Absolute Values")+
  geom_point(aes(y = rpart_imp, x = plsr_imp), cex = 0.4, pch  =20)+
  ## adds the Latent Heat predictors as red circles
  geom_point(aes(y = rpart_imp, x = plsr_imp),data =
imp_frame_comparison2[grep("Latent_heat",
imp_frame_comparison2$varNames),],
             col = "#ff0066", pch  =20, alpha = I(0.7))+
  ## adds the cumulative rainfall as blue droplets
  geom_point(aes(y = rpart_imp, x = plsr_imp),data =
imp_frame_comparison2[grep("BOMDom_rainfall_csum_",
imp_frame_comparison2$varNames),],
             col = "lightblue", alpha = I(0.7), cex = 1.5)+
  ## adds the FPAR as a green circle
  geom_point(aes(y = rpart_imp, x = plsr_imp),data =
imp_frame_comparison2[grep("SatDom_FPAR",
imp_frame_comparison2$varNames),],
             col = "darkgreen", alpha = I(0.7), pch = 20)+
  ## adds the EVI as a green open square
  geom_point(aes(y = rpart_imp, x = plsr_imp),data =
imp_frame_comparison2[grep("SatDom_EVI", imp_frame_comparison2$varNames),],
             col = "darkgreen", alpha = I(0.7), pch = 20)+
  ## adds the NDVI as a green open triangle
  geom_point(aes(y = rpart_imp, x = plsr_imp),data =
imp_frame_comparison2[grep("SatDom_NDVI",
imp_frame_comparison2$varNames),],
             col = "darkgreen", alpha = I(0.7), pch = 20)+
  ##Direct labels some points
  geom_text(aes(y = rpart_imp, x = plsr_imp),data =
imp_frame_comparison2[c(1),], label = "Dicot t/f", nudge_x = 0.0000017,
nudge_y = 0, size=3)+
  geom_point(aes(y = rpart_imp, x = plsr_imp),data =
imp_frame_comparison2[c(1),], pch  =20)+
  geom_text(aes(y = rpart_imp, x = plsr_imp),data =
imp_frame_comparison2[c(9),], label = "Canola t/f", nudge_x = 0.0000023,
nudge_y = 0, size=3)+
  geom_point(aes(y = rpart_imp, x = plsr_imp),data =
imp_frame_comparison2[c(9),], pch  =20)+
  geom_text(aes(y = rpart_imp, x = plsr_imp),data =
imp_frame_comparison2[c(8),], label = "Clethodim", nudge_x = 0.0000024,
nudge_y = 0, size=3)+
  geom_point(aes(y = rpart_imp, x = plsr_imp),data =
imp_frame_comparison2[c(8),], pch  =20)+
  geom_text(aes(y = rpart_imp, x = plsr_imp),data =
imp_frame_comparison2[c(10),], label = "Fabaceae t/f", nudge_x = 0.0000026,
nudge_y = 0, size=3)+
  geom_point(aes(y = rpart_imp, x = plsr_imp),data =
imp_frame_comparison2[c(10),], pch  =20)+
  geom_text(aes(y = rpart_imp, x = plsr_imp),data =
imp_frame_comparison2[c(11),], label = "Applied Sulphur", nudge_x =
0.0000026, nudge_y = 1000, size=3)+
  geom_point(aes(y = rpart_imp, x = plsr_imp),data =
imp_frame_comparison2[c(11),], pch  =20)+
  geom_text(aes(y = rpart_imp, x = plsr_imp),data =
imp_frame_comparison2[c(13),], label = "Wheat t/f", nudge_x = 0.0000017,
nudge_y = 0, size=3)+
  geom_point(aes(y = rpart_imp, x = plsr_imp),data =
imp_frame_comparison2[c(13),], pch  =20)+
  theme_bw()

Imp_plot3<-qplot(rpart_imp, rf_imp,data = imp_frame_comparison2, alpha =
I(0),
                 xlab = "RPRM Importance", ylab = "BCRF Importance")+
  geom_point(aes(x = rpart_imp, y = rf_imp), cex = 0.4, pch  =20)+
  ## adds the Latent Heat predictors as pink circles
  geom_point(aes(x = rpart_imp, y = rf_imp),data =
imp_frame_comparison2[grep("Latent_heat",
imp_frame_comparison2$varNames),],
             col = "#ff0066", pch  =20, alpha = I(0.7))+
  ## adds the cumulative rainfall as blue droplets
  geom_point(aes(x = rpart_imp, y = rf_imp),data =
imp_frame_comparison2[grep("BOMDom_rainfall_csum_",
imp_frame_comparison2$varNames),],
             col = "lightblue", alpha = I(0.7), cex = 1.5)+
  ## adds the raw reflectance bands as an orange square
  # geom_point(aes(x = rpart_imp, y = rf_imp),data =
imp_frame_comparison2[grep("SatDom_Reflect_b",
imp_frame_comparison2$varNames),],
  #             col = "orange", alpha = I(0.7), pch  =15)+
  ## adds the FPAR as a green circle
  geom_point(aes(x = rpart_imp, y = rf_imp),data =
imp_frame_comparison2[grep("SatDom_FPAR",
imp_frame_comparison2$varNames),],
             col = "darkgreen", alpha = I(0.7), pch = 20)+
  ## adds the EVI as a green open square
  geom_point(aes(x = rpart_imp, y = rf_imp),data =
imp_frame_comparison2[grep("SatDom_EVI", imp_frame_comparison2$varNames),],
             col = "darkgreen", alpha = I(0.7), pch = 20)+
  ## adds the NDVI as a green open triangle
  geom_point(aes(x = rpart_imp, y = rf_imp),data =
imp_frame_comparison2[grep("SatDom_NDVI",
imp_frame_comparison2$varNames),],
             col = "darkgreen", alpha = I(0.7), pch = 20)+
  ##Direct labels some points
  geom_text(aes(x = rpart_imp, y = rf_imp),data =
imp_frame_comparison2[c(1),], label = "Dicot t/f", nudge_x = -5000, nudge_y
= -9, size=3)+
  geom_point(aes(x = rpart_imp, y = rf_imp),data =
imp_frame_comparison2[c(1),], pch  =20)+
  geom_text(aes(x = rpart_imp, y = rf_imp),data =
imp_frame_comparison2[c(9),], label = "Canola t/f", nudge_x = 0.000001,
nudge_y = +7, size=3)+
  geom_point(aes(x = rpart_imp, y = rf_imp),data =
imp_frame_comparison2[c(9),], pch  =20)+
  geom_text(aes(x = rpart_imp, y = rf_imp),data =
imp_frame_comparison2[c(8),], label = "Clethodim", nudge_x = -0.000001,
```

```
nudge_y = -7, size=3)+
  geom_point(aes(x = rpart_imp, y = rf_imp),data =
imp_frame_comparison2[c(8),], pch  =20)+
  geom_text(aes(x = rpart_imp, y = rf_imp),data =
imp_frame_comparison2[c(10),], label = "Fabaceae t/f", nudge_x = 0.0000017,
nudge_y = -7, size=3)+
  geom_point(aes(x = rpart_imp, y = rf_imp),data =
imp_frame_comparison2[c(10),], pch  =20)+
  geom_text(aes(x = rpart_imp, y = rf_imp),data =
imp_frame_comparison2[c(11),], label = "Applied Sulphur", nudge_x =
0.0000017, nudge_y = +7, size=3)+
  geom_point(aes(x = rpart_imp, y = rf_imp),data =
imp_frame_comparison2[c(11),], pch  =20)+
  geom_text(aes(x = rpart_imp, y = rf_imp),data =
imp_frame_comparison2[c(13),], label = "Wheat t/f", nudge_x = 0.0000017,
nudge_y = -7, size=3)+
  geom_point(aes(x = rpart_imp, y = rf_imp),data =
imp_frame_comparison2[c(13),], pch  =20)+
  ## annotates
  geom_text(x = 30000, y = 75, label = "Latent Heat Flux", col = "#ff0066",
size=3)+
  geom_text(x = 15000, y = 33, label = "Cumulative Rainfall", col =
"lightblue", size=3)+
  geom_text(x = 23000, y = 8, label = "Vegetation Indices", col =
"darkgreen", size=3)+
  theme_bw()

Imp_plot<-grid.arrange(Imp_plot3+labs(title =
"a")+theme(text=element_text(size=9)),
                       Imp_plot1+labs(title =
"b")+theme(text=element_text(size=9)),
                       Imp_plot2+labs(title =
"c")+theme(text=element_text(size=9)),
                       qplot(c(0,0), c(10,10)),
                       nrow = 2)

#ggsave(Imp_plot,file = "Publication_docs/Fig_SXX3_Importance.tiff", device
= "tiff", height = 3, width = 7.5,dpi = 330)


## Generates a plot of cumulative seasonal rainfall importance, etc.
imp_frame_comparison2<-imp_frame_comparison
imp_frame_comparison2$plsr_imp<-abs(imp_frame_comparison$plsr_imp)

imp_frame_comparison2[is.na(imp_frame_comparison2)]<-0

## Strips the timing off the end of all variables

## gets negative values
Var_Timings<-lapply(as.character(imp_frame_comparison$varNames), strsplit,
split = "\\.\\.")
Var_Timings<-lapply(Var_Timings, unlist)
Var_Timings<-lapply(Var_Timings, tail, 1)
Var_Timings<-as.numeric(unlist(Var_Timings))
Var_Timings<-c(-(Var_Timings))

Var_Timings2<-lapply(as.character(imp_frame_comparison$varNames), strsplit,
split = "_\\.")
Var_Timings2<-lapply(Var_Timings2, unlist)
Var_Timings2<-lapply(Var_Timings2, tail, 1)
Var_Timings2<-as.numeric(unlist(Var_Timings2))
Var_Timings2<-c(-Var_Timings2)

Var_Timings[is.na(Var_Timings)]<-Var_Timings2[is.na(Var_Timings)]

Var_Timings2<-lapply(as.character(imp_frame_comparison$varNames), strsplit,
split = "_")
Var_Timings2<-lapply(Var_Timings2, unlist)
Var_Timings2<-lapply(Var_Timings2, tail, 1)
Var_Timings2<-as.numeric(unlist(Var_Timings2))
Var_Timings2[which(Var_Timings2<10 & Var_Timings2!=0)]<-NA

Var_Timings[is.na(Var_Timings)]<-Var_Timings2[is.na(Var_Timings)]
Var_Timings[Var_Timings==c(-42)]<-NA

## attaches to importance frame
imp_frame_comparison2<-data.frame(Var_Timings,imp_frame_comparison2)
## orders all data by these timing values
imp_frame_comparison2<-
imp_frame_comparison2[order(imp_frame_comparison2$Var_Timings),]
## removes NAs
imp_frame_comparison2<-
imp_frame_comparison2[!is.na(imp_frame_comparison2$Var_Timings),]

imp_frame_comparison3<-imp_frame_comparison2

imp_frame_comparison3$Water<-c(imp_frame_comparison2$varNames %in%
imp_frame_comparison2$varNames[grep("rainfall_csum",

imp_frame_comparison2$varNames)])
imp_frame_comparison3$rpart_imp<-log(imp_frame_comparison3$rpart_imp)
imp_frame_comparison3$rf_imp<-log(imp_frame_comparison3$rf_imp)
imp_frame_comparison3$plsr_imp<-log(imp_frame_comparison3$plsr_imp)

imp_frame_comparison3<-imp_frame_comparison3[-
c(which(imp_frame_comparison3$Var_Timings>300)),]

imp_frame_comparison3<-
imp_frame_comparison3[imp_frame_comparison3$rf_imp>=0,]

Importance_timings<-ggplot(imp_frame_comparison3, aes(Var_Timings,
rf_imp,group = Var_Timings))+
  geom_boxplot(outlier.size = 0)+
  ## adds the cumulative rainfall as blue droplets
  geom_point(aes(x = Var_Timings, y = rf_imp),
             data = imp_frame_comparison3[grep("BOMDom_rainfall_csum_",
imp_frame_comparison3$varNames),],
             col = "lightblue")+
  geom_point(aes(x = Var_Timings, y = rf_imp),
             data = imp_frame_comparison3[grep("BOMDom_rainfall_csum_",
imp_frame_comparison3$varNames),],
             col = "black", pch = 1)+
  geom_point(aes(x = Var_Timings, y = rf_imp),
             data = imp_frame_comparison3[grep("Latent_heat",
imp_frame_comparison3$varNames),],
             col = "#ff0066", alpha = I(0.7))+
  geom_smooth(col = "black", method = "loess", span = 0.1)+
  geom_vline(xintercept = 0, col = "red", alpha =I(0.5))+
```

```
  theme_bw()+
  theme(legend.position = c(0.1,0.8))+
  labs(x = "Days After Sowing", y = "Variable Importance")

Importance_timings

## Finished Figure 5

Imp_plot<-grid.arrange(Imp_plot3+labs(title =
"a")+theme(text=element_text(size=9)),
                       Imp_plot1+labs(title =
"b")+theme(text=element_text(size=9)),
                       Imp_plot2+labs(title =
"c")+theme(text=element_text(size=9)),
                       Importance_timings+labs(title =
"d")+theme(text=element_text(size=9)),
                       nrow = 2)

ggsave(Imp_plot,file = "Publication_docs/Fig_5_Importance.tiff", device =
"tiff", height = 8, width = 7.5,dpi = 330)




imp_frame_comparison3<-imp_frame_comparison2
imp_frame_comparison3$varNames<-
as.character(imp_frame_comparison3$varNames)
imp_frame_comparison3$varNames<-
gsub("\\.","_",imp_frame_comparison3$varNames)
imp_frame_comparison3$varNames<-
lapply(as.character(imp_frame_comparison3$varNames), strsplit, split =
"_[0-9]")
imp_frame_comparison3$varNames<-lapply(imp_frame_comparison3$varNames,
unlist)
imp_frame_comparison3$varNames<-
unlist(lapply(imp_frame_comparison3$varNames, head, 1))

imp_frame_comparison3<-reshape(imp_frame_comparison3, direction = "wide",
timevar = "Var_Timings", idvar = "varNames")

## Plots the importance score of variables relative to sowing dates
#imp_frame_comparison3<-
imp_frame_comparison2[grep("Latent_heat",imp_frame_comparison2$varNames),]

imp_frame_comparison3<-
imp_frame_comparison2[grep("rainfall_csum",imp_frame_comparison2$varNames),
]

## Plots the importance score of variables relative to sowing dates
imp_frame_comparison3$rpart_imp<-
c(imp_frame_comparison3$rpart_imp/max(imp_frame_comparison3$rpart_imp,
na.rm  =T))
imp_frame_comparison3$rf_imp<-
c(imp_frame_comparison3$rf_imp/max(imp_frame_comparison3$rf_imp, na.rm
=T))
imp_frame_comparison3$plsr_imp<-
c(imp_frame_comparison3$plsr_imp/max(imp_frame_comparison3$plsr_imp, na.rm
=T))

## merges on all times, coerces zeroes
imp_frame_comparison3<-merge(data.frame(Times_vec = seq(-80, 250, by =
10)), imp_frame_comparison3,
                             by.x = "Times_vec", by.y = "Var_Timings", all
= T)

#imp_frame_comparison3[is.na(imp_frame_comparison3)]<-0

## Plots the importance score of variables relative to sowing dates
ggplot(data = imp_frame_comparison3,aes(Times_vec,rpart_imp))+
  geom_point(col = "lightblue")+
  geom_line(col = "lightblue")+
  geom_point(aes(Times_vec,rf_imp), col = "orange")+
  geom_line(aes(Times_vec,rf_imp), col = "orange")+
  geom_point(aes(Times_vec,plsr_imp))+
  geom_line(aes(Times_vec,plsr_imp))+
  theme_bw()

## Does the same with latent heat flux
imp_frame_comparison3<-imp_frame_comparison2[grep("Latent_heat_[0-
9]",imp_frame_comparison2$varNames),]

imp_frame_comparison3$rpart_imp<-
c(imp_frame_comparison3$rpart_imp/max(imp_frame_comparison3$rpart_imp,
na.rm  =T))
imp_frame_comparison3$rf_imp<-
c(imp_frame_comparison3$rf_imp/max(imp_frame_comparison3$rf_imp, na.rm
=T))
imp_frame_comparison3$plsr_imp<-
c(imp_frame_comparison3$plsr_imp/max(imp_frame_comparison3$plsr_imp, na.rm
=T))

## Plots the importance score of variables relative to sowing dates
qplot(Var_Timings,rpart_imp,data = imp_frame_comparison3, xlim = c(-
50,250))+
  geom_line(aes(Var_Timings,rpart_imp), col = "black")+
  geom_point(aes(Var_Timings,rf_imp),data = imp_frame_comparison3, col =
"orange")+
  geom_line(aes(Var_Timings,rf_imp),data = imp_frame_comparison3, col =
"orange")+
  geom_point(aes(Var_Timings,plsr_imp),data = imp_frame_comparison3, col =
"lightblue")+
  geom_line(aes(Var_Timings,plsr_imp),data = imp_frame_comparison3, col =
"lightblue")+
  theme_bw()

## And the vegetation Indices
imp_frame_comparison3<-imp_frame_comparison2[c(grep("VI_[0-
9]",imp_frame_comparison2$varNames),grep("FPAR_[0-
9]",imp_frame_comparison2$varNames)),]

imp_frame_comparison3$rpart_imp<-
c(imp_frame_comparison3$rpart_imp/max(imp_frame_comparison3$rpart_imp,
na.rm  =T))
imp_frame_comparison3$rf_imp<-
c(imp_frame_comparison3$rf_imp/max(imp_frame_comparison3$rf_imp, na.rm
=T))
imp_frame_comparison3$plsr_imp<-
c(imp_frame_comparison3$plsr_imp/max(imp_frame_comparison3$plsr_imp, na.rm
```

```
=T))

## Plots the importance score of variables relative to sowing dates
qplot(Var_Timings,rpart_imp,data = imp_frame_comparison3, xlim = c(-
50,250))+
  geom_line(aes(Var_Timings,rpart_imp), col = "black")+
  geom_point(aes(Var_Timings,rf_imp),data = imp_frame_comparison3, col =
"orange")+
  geom_line(aes(Var_Timings,rf_imp),data = imp_frame_comparison3, col =
"orange")+
  geom_point(aes(Var_Timings,plsr_imp),data = imp_frame_comparison3, col =
"lightblue")+
  geom_line(aes(Var_Timings,plsr_imp),data = imp_frame_comparison3, col =
"lightblue")+
  theme_bw()

qplot(Var_Timings,c(rpart_imp/max(rpart_imp)),data = imp_frame_comparison3,
xlim = c(-50,250))+
  geom_line(aes(Var_Timings,c(rpart_imp/max(rpart_imp))), col = "black")+
  geom_point(aes(Var_Timings,c(rf_imp/max(rf_imp))),data =
imp_frame_comparison3, col = "orange")+
  geom_line(aes(Var_Timings,c(rf_imp/max(rf_imp))),data =
imp_frame_comparison3, col = "orange")+
  geom_point(aes(Var_Timings,c(plsr_imp/max(plsr_imp, na.rm = T))),data =
imp_frame_comparison3, col = "lightblue")+
  geom_line(aes(Var_Timings,c(plsr_imp/max(plsr_imp, na.rm = T))),data =
imp_frame_comparison3, col = "lightblue")+
  theme_bw()



#######################################


## Figure 1a


#############################################################

## Fig 1

#############################################################

## makes a leaflet map of locations, coloured by crop
lat_longs<-read.csv(file =
"data/constructed_csv_files/geolocs_output1.csv", header = T)

lat_longs<-lat_longs[!is.na(rowMeans(lat_longs[,c(3:4)])),]

## merges on site-average trial yields and other phenotypes

results<-readRDS("data/results_list.rds")

results$Trial_ID<-toupper(results$`Trial ID`)

xobj<-merge(lat_longs, results, by.x = "Trial_ID", by.y = "Trial_ID", all =
T)

xobj<-xobj[!duplicated(paste0(xobj$Trial_ID, xobj$`Sowing Date`,
xobj$Latitude, xobj$`Site Mean (t/ha)`)),]

xobj<-xobj[!is.na(xobj$Trial_ID),]

xobj$`Site Mean (t/ha)`<-as.numeric(as.character(xobj$`Site Mean (t/ha)`))

xobj_canola<-xobj[which(xobj$`Crop Type`=="Canola"),]

ggplot(data = ne_countries(country = "australia", scale = "medium",
returnclass = "sf")) +
  geom_sf(color = "black", fill = "black") +
  annotation_scale(location = "bl", width_hint = 0.5) +
  annotation_north_arrow(location = "bl", which_north = "true",
                         pad_x = unit(0.75, "in"), pad_y = unit(0.5, "in"),
                         style = north_arrow_fancy_orienteering) +
  coord_sf(ylim = c(-10, -43), xlim = c(115, 155))+
  theme_bw()+
  scale_color_viridis(xobj$Sowing_date_numeric.x, option = "D")+
  geom_point(data = xobj, aes(Longitude,Latitude, col =
Sowing_date_numeric.x), pch  =20, alpha = I(0.5), cex = 0.5)

AusMap<-ggplot(data = ne_countries(country = "australia", scale = "medium",
returnclass = "sf")) +
  geom_sf(color = "black", fill = "white") +
  annotation_scale(location = "bl", width_hint = 0.5) +
  annotation_north_arrow(location = "bl", which_north = "true",
                         pad_x = unit(0.4, "npc"), pad_y = unit(0.4,
"npc"),
                         style = north_arrow_nautical) +
  coord_sf(ylim = c(-10, -43), xlim = c(112, 153))+
  theme_bw()+
  scale_color_viridis(xobj$Sowing_date_numeric.x, option = "D")+
  geom_point(data = xobj, aes(Longitude,Latitude, col =
Sowing_date_numeric.x),  pch  =20, alpha = I(0.8), cex = 0.5)+
  theme(legend.position = "none")


## lays out figure 1
gridMatrix<-matrix(c(1,1,1,2,2,2,2,
                     1,1,1,2,2,2,2,
                     1,1,1,2,2,2,2,
                     3,3,3,3,3,3,3,
                     3,3,3,3,3,3,3,
                     3,3,3,3,3,3,3,
                     3,3,3,3,3,3,3,
                     3,3,3,3,3,3,3), nrow = 8, byrow = T)

Fig_1<-grid.arrange(AusMap+labs(title = "a")+
                      theme(text = element_text(size=9)),
                    Photosynth_Fig1b+labs(title = "b")+
                      theme(text = element_text(size=9)),
                    Umap_alldata_figure+labs(title = "c")+
                      theme(text = element_text(size=9)),
                    nrow = 2, layout_matrix = gridMatrix)

#Fig_1<-grid.arrange(AusMap, Photosynth_Fig1b, Umap_alldata_figure, nrow =
1)

ggsave(Fig_1,file = "Publication_docs/Fig1.tiff", device = "tiff", height =
```

```
8, width = 7,dpi = 330)



LeafMap = leaflet() %>%
  addProviderTiles("CartoDB.DarkMatter") %>%
  addCircleMarkers(lat_longs$Longitude, lat_longs$Latitude,
                   radius  =0.5, fillOpacity = 0.2,
                   color = heat.colors(lat_longs$Sowing_date_numeric))

LeafMap



############################################################

## Figure SXX: distribution of select phenotypes

############################################################


mod_frame1<-readRDS("model_frames_and_imputations/mod_frame1.rds")

## separates phenotypes
Phen_data<-mod_frame1[,grep("PHENDom", colnames(mod_frame1))]
mod_frame1<-mod_frame1[,-c(grep("PHENDom", colnames(mod_frame1)))]

## sets up dev
tiff(filename  = "Publication_docs/Agronomic_trait_boxplots.tiff",
     width  =7.5, height = 7, units = "in", res  =330)

par(mfrow = c(2,3), bty = "n")

## wheat yield and protein
boxplot(Phen_data$PHENDom_yield_t_ha[Phen_data$PHENDom_Crop_alldata=="Wheat"]~mod_frame1$METADom_Year[Phen_data$PHENDom_Crop_alldata=="Wheat"]
,
        xlab = "Year", ylab = "Yield, t/Ha", pch = ".", main  ="a", las =
2, varwidth  =T)
abline(h = seq(0,14, by  =2), lty = 3, col = "lightgrey")

boxplot(Phen_data$PHENDom_Flowering_QC[Phen_data$PHENDom_Crop_alldata=="Wheat"]~mod_frame1$METADom_Year[Phen_data$PHENDom_Crop_alldata=="Wheat"]
,
        xlab = "Year", ylab = "Days to Flowering", pch = ".", main  ="b",
las = 2, varwidth  =T)
abline(h = seq(50,150, by  =50), lty = 3, col = "lightgrey")

boxplot(Phen_data$PHENDom_Protein[Phen_data$PHENDom_Crop_alldata=="Wheat"]~mod_frame1$METADom_Year[Phen_data$PHENDom_Crop_alldata=="Wheat"]
,
        xlab = "Year", ylab = "Protein, % dry weight", pch = ".", main
="c", las = 2, varwidth  =T)
abline(h = seq(0,40, by  =10), lty = 3, col = "lightgrey")

boxplot(Phen_data$PHENDom_yield_t_ha[Phen_data$PHENDom_Crop_alldata=="Canola"]~mod_frame1$METADom_Year[Phen_data$PHENDom_Crop_alldata=="Canola"]
,
        xlab = "Year", ylab = "Yield, t/Ha", pch = ".", main  ="d", las =
2, varwidth  =T)
abline(h = seq(0,5, by  =1), lty = 3, col = "lightgrey")

boxplot(as.numeric(Phen_data$PHENDom_Flowering_QC)[Phen_data$PHENDom_Crop_alldata=="Canola"]~mod_frame1$METADom_Year[Phen_data$PHENDom_Crop_alldata=="Canola"]
,
        xlab = "Year", ylab = "Days to Flowering", pch = ".", main  ="e",
las = 2, varwidth  =T)
abline(h = seq(50,200, by  =50), lty = 3, col = "lightgrey")

boxplot(as.numeric(Phen_data$PHENDom_Glucosinolates)[Phen_data$PHENDom_Crop_alldata=="Canola"]~mod_frame1$METADom_Year[Phen_data$PHENDom_Crop_alldata=="Canola"]
,
        xlab = "Year", ylab = "Glucosinolates, %", pch = ".", main  ="f",
las = 2, varwidth  =T)
abline(h = seq(0,50, by  =10), lty = 3, col = "lightgrey")

par(mfrow = c(1,1))

dev.off()


tiff(filename  = "Publication_docs/Species_Yields_boxplots.tiff",
     width  =7.5, height = 7, units = "in", res  =330)

par(mfrow = c(3,3), bty = "n", mar = c(4.1,3.1,2.1,2.1))

boxplot(Phen_data$PHENDom_yield_t_ha[Phen_data$PHENDom_Crop_alldata=="Wheat"]~mod_frame1$METADom_Year[Phen_data$PHENDom_Crop_alldata=="Wheat"]
,
        main = "Wheat, N=54658",
        xlab = "Year", ylab = "yield, t/Ha", pch = ".", las = 2, varwidth
=T)

abline(h = seq(0,14, by  =2), lty = 3, col = "lightgrey")

boxplot(Phen_data$PHENDom_yield_t_ha[Phen_data$PHENDom_Crop_alldata=="Oat"]~mod_frame1$METADom_Year[Phen_data$PHENDom_Crop_alldata=="Oat"]
,
        main = "Oat, N=3873",
        xlab = "Year", ylab = "yield, t/Ha", pch = ".", las = 2, varwidth
=T)

abline(h = seq(0,13, by  =1), lty = 3, col = "lightgrey")

boxplot(Phen_data$PHENDom_yield_t_ha[Phen_data$PHENDom_Crop_alldata=="Triticale"]~mod_frame1$METADom_Year[Phen_data$PHENDom_Crop_alldata=="Triticale"]
,
        main = "Triticale, N=2081",
        xlab = "Year", ylab = "yield, t/Ha", pch = ".", las = 2, varwidth
=T)

abline(h = seq(0,13, by  =1), lty = 3, col = "lightgrey")

boxplot(Phen_data$PHENDom_yield_t_ha[Phen_data$PHENDom_Crop_alldata=="Canola"]~mod_frame1$METADom_Year[Phen_data$PHENDom_Crop_alldata=="Canola"]
,
        main = "Canola, N=14929",
        xlab = "Year", ylab = "yield, t/Ha", pch = ".", las = 2, varwidth
=T)

abline(h = seq(0,13, by  =1), lty = 3, col = "lightgrey")

boxplot(Phen_data$PHENDom_yield_t_ha[Phen_data$PHENDom_Crop_alldata=="Chickpea"]~mod_frame1$METADom_Year[Phen_data$PHENDom_Crop_alldata=="Chickpea"]
,
        main = "Chickpea, N=3677",
        xlab = "Year", ylab = "yield, t/Ha", pch = ".", las = 2, varwidth
=T)
```

```
abline(h = seq(0,13, by  =1), lty = 3, col = "lightgrey")

boxplot(Phen_data$PHENDom_yield_t_ha[Phen_data$PHENDom_Crop_alldata=="Field
Pea"]~mod_frame1$METADom_Year[Phen_data$PHENDom_Crop_alldata=="Field Pea"],
        main = "Field Pea, N=3896",
        xlab = "Year", ylab = "yield, t/Ha", pch = ".", las = 2, varwidth
=T)

abline(h = seq(0,13, by  =1), lty = 3, col = "lightgrey")

boxplot(Phen_data$PHENDom_yield_t_ha[Phen_data$PHENDom_Crop_alldata=="Faba
Bean"]~mod_frame1$METADom_Year[Phen_data$PHENDom_Crop_alldata=="Faba
Bean"],
        main = "Faba Bean, N=2224",
        xlab = "Year", ylab = "yield, t/Ha", pch = ".", las = 2, varwidth
=T)

abline(h = seq(0,13, by  =1), lty = 3, col = "lightgrey")

boxplot(Phen_data$PHENDom_yield_t_ha[Phen_data$PHENDom_Crop_alldata=="Lentil"]~mod_frame1$METADom_Year[Phen_data$PHENDom_Crop_alldata=="Lentil"]
,
        main = "Lentil, N=2260",
        xlab = "Year", ylab = "yield, t/Ha", pch = ".", las = 2, varwidth
=T)

abline(h = seq(0,13, by  =1), lty = 3, col = "lightgrey")

boxplot(Phen_data$PHENDom_yield_t_ha[Phen_data$PHENDom_Crop_alldata=="Lupin"]~mod_frame1$METADom_Year[Phen_data$PHENDom_Crop_alldata=="Lupin"]
,
        main = "Lupin, N=2747",
        xlab = "Year", ylab = "yield, t/Ha", pch = ".", las = 2, varwidth
=T)

abline(h = seq(0,13, by  =1), lty = 3, col = "lightgrey")

par(mfrow = c(1,1))

dev.off()


############################################################

## Figure SXX-SXX. Accuracy of yield and protein models by species

############################################################


## Yield models at +100DAS or +200DAS
Yield_predobs_100<-readRDS(file =
"model_frames_and_imputations/siteVariety_models/Yield_predobs_100.rds")
Protein_predobs_100<-readRDS(file =
"model_frames_and_imputations/siteVariety_models/Protein_predobs_100.rds")

Yield_predobs_200<-readRDS(file =
"model_frames_and_imputations/siteVariety_models/Yield_predobs_200.rds")
Protein_predobs_200<-readRDS(file =
"model_frames_and_imputations/siteVariety_models/Protein_predobs_200.rds")

## makes a figure of model forecast accuracy, versus internal (missing
plot) accuracy
# Canola    Chickpea  Faba Bean Lupin     Oat       Field Pea Lentil
Triticale Wheat

#saveRDS(lsvm_mod, file =
paste0("model_frames_and_imputations/siteVariety_models/rf_Yield_100DAS",
#                               spp_list[[i]],".rds"))
#saveRDS(rf_mod, file =
paste0("model_frames_and_imputations/siteVariety_models/lsvm_Yield_100DAS",
#                              spp_list[[i]],".rds"))

qplot(Yield_predobs_100[[1]][1,],
      jitter(Yield_predobs_100[[1]][3,], factor = 50),
      # xlim = c(0,12), ylim = c(0,12),
      alpha = I(0),
      xlab = "Predicted Yield, t/Ha", ylab = "Observed Yield, t/Ha")+
  geom_point(col = "orange", alpha = I(0.7))+
  geom_smooth(method = "lm", col = "black")+
  theme_bw()


## plots accuracy of yield+protein models across species
namelist<-NULL
for(i in 1:9){
  bestfit_reg<-odregress(Yield_predobs_200[[i]][1,],
                         Yield_predobs_200[[i]][2,])

  namelist[[i]]<-ggplot(data.frame(Predicted =
jitter(Yield_predobs_200[[i]][2,], factor = 50),
                                   Observed = Yield_predobs_200[[i]][1,]),
                        aes(x = Observed,y = Predicted)) +
    geom_abline(intercept = 0, slope = 1, col  ="darkgrey")+
    geom_point(alpha = I(0.6), col = "orange", pch  =20, cex= 0.5)+
    geom_abline(slope = bestfit_reg$coeff[1], intercept =
bestfit_reg$coeff[2], col = "black", alpha = 0.7)+
    coord_fixed(xlim = c(0,max(Yield_predobs_200[[i]][1,])), ylim =
c(0,max(Yield_predobs_200[[i]][1,])), ratio = 1)+
    annotate("text",
             x = c(max(Yield_predobs_200[[i]][1,])/2),
             y = 0.5,
             label = paste0("RÂ² = ",
round(cor.test(Yield_predobs_200[[i]][1,],

Yield_predobs_200[[i]][2,])$estimate, digits = 2)))+
    theme_bw()
  ## prints target sample size
  print(length(Yield_predobs_200[[i]][1,]))

}


# Canola    Chickpea  Faba Bean Lupin     Oat       Field Pea Lentil
Triticale Wheat
Yield_grid<-grid.arrange(namelist[[1]]+labs(title = "a   Canola"),
                         namelist[[2]]+labs(title = "b   Chickpea"),
                         namelist[[3]]+labs(title = "c   Faba Bean"),
                         namelist[[4]]+labs(title = "d   Lupin"),
                         namelist[[5]]+labs(title = "e   Oat"),
                         namelist[[6]]+labs(title = "f   Field Pea"),
```

```
                         namelist[[7]]+labs(title = "g   Lentil"),
                         namelist[[8]]+labs(title = "h   Triticale"),
                         namelist[[9]]+labs(title = "i   Wheat"), nrow =3)

# Canola    Chickpea  Faba Bean Lupin     Oat       Field Pea Lentil
Triticale Wheat
Yield_grid<-grid.arrange(namelist[[1]]+labs(title = "a")+
                           annotate("text", label ="Canola",
                                    x =
c(max(Yield_predobs_200[[1]][1,])/2),
                                    y =
c(max(Yield_predobs_200[[1]][1,])*0.9)),
                         namelist[[2]]+labs(title = "b")+
                           annotate("text", label ="Chickpea",
                                    x =
c(max(Yield_predobs_200[[2]][1,])/2),
                                    y =
c(max(Yield_predobs_200[[2]][1,])*0.9)),
                         namelist[[3]]+labs(title = "c")+
                           annotate("text", label ="Faba Bean",
                                    x =
c(max(Yield_predobs_200[[3]][1,])/2),
                                    y =
c(max(Yield_predobs_200[[3]][1,])*0.9)),
                         namelist[[4]]+labs(title = "d ")+
                           annotate("text", label ="Lupin",
                                    x =
c(max(Yield_predobs_200[[4]][1,])/2),
                                    y =
c(max(Yield_predobs_200[[4]][1,])*0.9)),
                         namelist[[5]]+labs(title = "e ")+
                           annotate("text", label ="Oat",
                                    x =
c(max(Yield_predobs_200[[5]][1,])/2),
                                    y =
c(max(Yield_predobs_200[[5]][1,])*0.9)),
                         namelist[[6]]+labs(title = "f")+
                           annotate("text", label ="Field Pea",
                                    x =
c(max(Yield_predobs_200[[6]][1,])/2),
                                    y =
c(max(Yield_predobs_200[[6]][1,])*0.9)),
                         namelist[[7]]+labs(title = "g")+
                           annotate("text", label ="Lentil",
                                    x =
c(max(Yield_predobs_200[[7]][1,])/2),
                                    y =
c(max(Yield_predobs_200[[7]][1,])*0.9)),
                         namelist[[8]]+labs(title = "h")+
                           annotate("text", label ="Triticale",
                                    x =
c(max(Yield_predobs_200[[8]][1,])/2),
                                    y =
c(max(Yield_predobs_200[[8]][1,])*0.9)),
                         namelist[[9]]+labs(title = "i")+
                           annotate("text", label ="Wheat",
                                    x =
c(max(Yield_predobs_200[[9]][1,])/2),
                                    y =
c(max(Yield_predobs_200[[9]][1,])*0.9)), nrow =3)



ggsave(Yield_grid,file = "Publication_docs/Fig_SXX_YieldGrid200DAS.tiff",
device = "tiff", height = 7, width = 7,dpi = 330)


Yield_grid<-grid.arrange(namelist[[1]]+labs(title = "a   Canola"),
                         namelist[[2]]+labs(title = "b   Chickpea"),
                         namelist[[3]]+labs(title = "c   Faba Bean"),
                         namelist[[4]]+labs(title = "d   Lupin"),
                         namelist[[5]]+labs(title = "e   Oat"),
                         namelist[[6]]+labs(title = "f   Field Pea"),
                         namelist[[7]]+labs(title = "g   Lentil"),
                         namelist[[8]]+labs(title = "h   Triticale"),
                         namelist[[9]]+labs(title = "i   Wheat"), nrow =3)


## Repeats for Protein predictions
namelist<-NULL
for(i in 1:3){
  bestfit_reg<-odregress(Protein_predobs_200[[i]][1,],
                         Protein_predobs_200[[i]][2,])

  namelist[[i]]<-ggplot(data.frame(Predicted =
jitter(Protein_predobs_200[[i]][2,], factor = 50),
                                   Observed =
Protein_predobs_200[[i]][1,]),
                        aes(x = Observed,y = Predicted)) +
    geom_abline(intercept = 0, slope = 1, col = "darkgrey")+
    geom_point(alpha = I(0.6), col = "orange", pch  =20, cex= 0.5)+
    geom_abline(slope = bestfit_reg$coeff[1], intercept =
bestfit_reg$coeff[2], col = "black", alpha = 0.7)+
    coord_fixed(xlim = c(5,max(Protein_predobs_200[[i]][1,])), ylim =
c(5,max(Protein_predobs_200[[i]][1,])), ratio = 1)+
    geom_density2d(alpha = I(0.7))+
    annotate("text",
             x = c(max(Protein_predobs_200[[i]][1,])/2)+2.5,
             y = 5.5,
             label = paste0("RÂ² = ",
round(cor.test(Protein_predobs_200[[i]][1,],

Protein_predobs_200[[i]][2,])$estimate, digits = 2)))+
    theme_bw()
  ## prints target sample size
  print(length(Protein_predobs_200[[i]][1,]))

}

## and for protein 100DAS
## Repeats for Protein predictions
namelist2<-NULL
for(i in 1:3){
  bestfit_reg<-odregress(Protein_predobs_100[[i]][1,],
                         Protein_predobs_100[[i]][2,])

  namelist2[[i]]<-ggplot(data.frame(Predicted =
jitter(Protein_predobs_100[[i]][2,], factor = 50),
```

```
                                    Observed =
Protein_predobs_100[[i]][1,]),
                         aes(x = Observed,y = Predicted)) +
    geom_abline(intercept = 0, slope = 1, col = "darkgrey")+
    geom_point(alpha = I(0.6), col = "orange", pch  =20, cex= 0.5)+
    geom_abline(slope = bestfit_reg$coeff[1], intercept =
bestfit_reg$coeff[2], col = "black", alpha = 0.7)+
    coord_fixed(xlim = c(5,max(Protein_predobs_100[[i]][1,])), ylim =
c(5,max(Protein_predobs_100[[i]][1,])), ratio = 1)+
    geom_density2d(alpha = I(0.7))+
    annotate("text",
             x = c(max(Protein_predobs_100[[i]][1,])/2)+2.5,
             y = 5.5,
             label = paste0("RÂ² = ",
round(cor.test(Protein_predobs_100[[i]][1,],

Protein_predobs_100[[i]][2,])$estimate, digits = 2)))+
    theme_bw()
  ## prints target sample size
  print(length(Protein_predobs_100[[i]][1,]))

}

#Protein_grid<-grid.arrange(namelist[[1]]+labs(title = "a   Oat"),
#                          namelist[[2]]+labs(title = "b   Triticale"),
#                          namelist[[3]]+labs(title = "c   Wheat"), nrow =1)

# Canola    Chickpea  Faba Bean Lupin     Oat       Field Pea Lentil
Triticale Wheat
Protein_grid<-grid.arrange(namelist[[1]]+labs(title = "a   Oat 200DAS"),
                           namelist[[2]]+labs(title = "b   Triticale
200DAS"),
                           namelist[[3]]+labs(title = "c   Wheat 200DAS"),
                           namelist2[[1]]+labs(title = "d   Oat 100DAS"),
                           namelist2[[2]]+labs(title = "e   Triticale
100DAS"),
                           namelist2[[3]]+labs(title = "f   Wheat 100DAS"),
nrow =2)

ggsave(Protein_grid,file = "Publication_docs/Fig_SXX_ProteinGrid.tiff",
device = "tiff", height = 5, width = 7,dpi = 330)

lsvm_100DAS_wheat<-ggplot(data.frame(Predicted_yield =
Yield_predobs_100[[9]][3,],
                                     Observed_yield =
Yield_predobs_100[[9]][1,]),
                          aes(x = Predicted_yield, y = Observed_yield),
                          alpha = I(0)) +
  geom_hex(bins = 50)+
  geom_abline(intercept = 0, slope = 1)+
  geom_smooth(method = "lm", col = "orange")+
  coord_fixed(xlim = c(0,12), ylim = c(0,12), ratio = 1)+
  scale_fill_viridis()+
  theme_bw()

cor.test(Yield_predobs_100[[9]][3,],Yield_predobs_100[[9]][1,])

bcrf_100DAS_wheat<-ggplot(data.frame(Predicted_yield =
Yield_predobs_100[[9]][2,],
                                     Observed_yield =
Yield_predobs_100[[9]][1,]),
                          aes(x = Predicted_yield, y = Observed_yield),
                          alpha = I(0)) +
  geom_hex(bins = 50)+
  geom_abline(intercept = 0, slope = 1)+
  geom_smooth(method = "lm", col = "orange")+
  coord_fixed(xlim = c(0,12), ylim = c(0,12), ratio = 1)+
  scale_fill_viridis()+
  theme_bw()


cor.test(Yield_predobs_100[[9]][2,],Yield_predobs_100[[9]][1,])

grid.arrange(bcrf_100DAS_wheat, lsvm_100DAS_wheat, nrow =1)



lsvm_100DAS_canola<-ggplot(data.frame(Predicted_yield =
Yield_predobs_100[[1]][3,],
                                      Observed_yield =
Yield_predobs_100[[1]][1,]),
                           aes(x = Predicted_yield, y = Observed_yield),
                           alpha = I(0)) +
  geom_hex(bins = 50)+
  geom_abline(intercept = 0, slope = 1)+
  geom_smooth(method = "lm", col = "orange")+
  coord_fixed(xlim = c(0,3), ylim = c(0,3), ratio = 1)+
  scale_fill_viridis()+
  theme_bw()

cor.test(Yield_predobs_100[[1]][3,],Yield_predobs_100[[1]][1,])

bcrf_100DAS_canola<-ggplot(data.frame(Predicted_yield =
Yield_predobs_100[[1]][2,],
                                      Observed_yield =
Yield_predobs_100[[1]][1,]),
                           aes(x = Predicted_yield, y = Observed_yield),
                           alpha = I(0)) +
  geom_hex(bins = 50)+
  geom_abline(intercept = 0, slope = 1)+
  geom_smooth(method = "lm", col = "orange")+
  coord_fixed(xlim = c(0,3), ylim = c(0,3), ratio = 1)+
  scale_fill_viridis()+
  theme_bw()

cor.test(Yield_predobs_100[[1]][2,],Yield_predobs_100[[1]][1,])

require(gridExtra)
grid.arrange(bcrf_100DAS_wheat, lsvm_100DAS_wheat,
             bcrf_100DAS_canola, lsvm_100DAS_canola, nrow =2)

namelist<-NULL
for(i in 1:9){

  namelist[[i]]<-ggplot(data.frame(Predicted = Yield_predobs_100[[i]][2,],
                                   Observed = Yield_predobs_100[[i]][1,]),
                        aes(x = Predicted, y = Observed),
                        alpha = I(0.6)) +
```

```
    geom_hex(bins = 50)+
    geom_abline(intercept = 0, slope = 1)+
    geom_smooth(method = "lm", col = "orange")+
    coord_fixed(xlim = c(0,max(Yield_predobs_100[[i]][1,])), ylim =
c(0,max(Yield_predobs_100[[i]][1,])), ratio = 1)+
    scale_fill_viridis()+
    theme_bw()
}

grid.arrange(namelist[[1]],
             namelist[[2]],
             namelist[[3]],
             namelist[[4]],
             namelist[[5]],
             namelist[[6]],
             namelist[[7]],
             namelist[[8]],
             namelist[[9]], nrow =3)


namelist<-NULL
for(i in 1:9){

  namelist[[i]]<-ggplot(data.frame(Predicted = Yield_predobs_100[[i]][3,],
                                   Observed = Yield_predobs_100[[i]][1,]),
                        aes(x = Predicted, y = Observed),
                        alpha = I(0.6)) +
    geom_hex(bins = 50)+
    geom_abline(intercept = 0, slope = 1)+
    geom_smooth(method = "lm", col = "orange")+
    coord_fixed(xlim = c(0,max(Yield_predobs_100[[i]][1,])), ylim =
c(0,max(Yield_predobs_100[[i]][1,])), ratio = 1)+
    scale_fill_viridis()+
    theme_bw()
}
grid.arrange(namelist[[1]],
             namelist[[2]],
             namelist[[3]],
             namelist[[4]],
             namelist[[5]],
             namelist[[6]],
             namelist[[7]],
             namelist[[8]],
             namelist[[9]], nrow =3)

#########################################################################
#########################################################################
#########################################################################
#########################################################################

## The code required to reproduce all figures from the paper:
## requires running S1 and S2 code in order, in the same workspace.

## Not necessary to reproduce the analysis

#########################################################################
#########################################################################
#########################################################################




## figures

#devtools::install_github('thomasp85/gganimate')

require(ggplot2)
require(gganimate)
require(ggthemes)
require(leaflet)
require(viridis)
require(mapview)
require(umap)
require(pracma)
require(rpart.plot)

# webshot::install_phantomjs()
library(animation)
library(png)
library(htmlwidgets)
library(webshot)
library(ggmap)
require(gridExtra)
require(beeswarm)
require(rnaturalearth)
require(rnaturalearthdata)
require(ggspatial)
require(sf)

##########################################

## Prints the rpart model across all species as plain text

rpart_mod<-readRDS(file =
"model_frames_and_imputations/siteVariety_models/rpart_allTimes.rds")

rpart_mod

cor.test(predobs_frame_unscaled[[2]]$Rrf_200,
predobs_frame_unscaled[[2]]$Yield)

## gets the fraction of data that is satellite data
length(grep("SatDom",
names(rpart_mod$variable.importance)))/length(names(rpart_mod$variable.importance)
)

rpart_mod<-readRDS(file =
"model_frames_and_imputations/siteVariety_models/rpart_TOS.rds")

rpart_mod

cor.test(predobs_frame_unscaled[[2]]$Rrf_TOS,
predobs_frame_unscaled[[2]]$Yield)



###################
```

```
## Builds a site clustering based on BOM data, Satellite data, and combined

###################

## loads phenotype data
P_frame<-read.csv("model_frames_and_imputations/site_frame.csv", header =
T)
P_frame<-P_frame[,grep("PHENDom", colnames(P_frame))]

## just keeps the species, a few key variables, and yield
P_frame<-data.frame(Yield = P_frame$PHENDom_MEAN.YIELD,
                    Yield2 = P_frame$PHENDom_Pins_Site_Mean_Yield,
                    DTH = P_frame$PHENDom_Days_to_Harvest,
                    Crop = P_frame$PHENDom_Crop)

## loads the first random-sample imputation
Site_Modelframe<-read.csv(file =
"model_frames_and_imputations/site_imputations/random_sample_1.csv", header
= T)


## gets environment data, performs a umap clustering
sub_mat<-Site_Modelframe[,c(grep("BOMDom_", colnames(Site_Modelframe)),
                            grep("SatDom_", colnames(Site_Modelframe)))]

## trims data past 200 days after sowing, around the latest date you could
expect to harvest most crops
sub_mat<-sub_mat[,-c(grep("2[0-9]{2}$", colnames(sub_mat)))]
P_frame<-P_frame[!duplicated(paste(Site_Modelframe[,1],Site_Modelframe[,2],
sep = "_")),]
P_frame$Year<-
Site_Modelframe$METADom_Year[!duplicated(paste(Site_Modelframe[,1],Site_Modelframe[,2]
, sep = "_"))]
P_frame$JTS<-
Site_Modelframe$MANDom_JTS[!duplicated(paste(Site_Modelframe[,1],Site_Modelframe[,2]
, sep = "_"))]
P_frame$Lat<-
Site_Modelframe$BOMDom_Latitude[!duplicated(paste(Site_Modelframe[,1],Site_Modelframe[,2]
, sep = "_"))]
P_frame$Long<-
Site_Modelframe$BOMDom_Longitude[!duplicated(paste(Site_Modelframe[,1],Site_Modelframe[,2]
, sep = "_"))]
sub_mat<-sub_mat[!duplicated(paste(Site_Modelframe[,1],Site_Modelframe[,2],
sep = "_")),]
rownames(sub_mat)<-paste(sub_mat[,1],sub_mat[,2], sep = "_")

sub_mat<-as.matrix(sub_mat[,-c(1)])

## Captures principal components
system.time(env_PCA<-prcomp(sub_mat))


qplot(env_PCA$x[,2], env_PCA$x[,4], col = P_frame$Crop)


## and builds a UMAP
ENV_umap<-umap(sub_mat,config = umap.defaults, method = "naive")

umap_plot_E<-data.frame(ENV_umap$layout,P_frame)
colnames(umap_plot_E)[1:2]<-c("Dimension_1", "Dimension_2")

Umap_alldata_figure<-qplot(umap_plot_E$Dimension_1,umap_plot_E$Dimension_2,
                           col = c(umap_plot_E$Year),
                           xlim = c(-8,9), ylim = c(-8.5,7.5),
                           xlab  = "Dimension 1", ylab  =" Dimension 2",
                           alpha = I(0))+
  geom_point(size = 0.5)+
  scale_color_viridis("Year",c(umap_plot_E$Year), option  ="D")+
  theme_bw()+
  theme(legend.position = c(0.9,0.2))


ggsave(Umap_alldata_figure,file = "Publication_docs/Fig1c.tiff", device =
"tiff", height = 6, width = 6,dpi = 330)



ggplot(aes(Dimension_1,Dimension_2,col = Year), data = umap_plot_E,
       col = Year)+
  geom_point()

qplot(umap_plot_E$Dimension_1,umap_plot_E$Dimension_2,col =
umap_plot_E$Crop,
      xlim = c(-10,10), ylim = c(-8,8), alpha = I(0.5))+theme_bw()

## generates one for wheat only



ENV_Wheat_umap<-umap(sub_mat[which(P_frame$Crop=="Wheat"),],config =
umap.defaults, method = "naive")

umap_plot_E_Wheat<-
data.frame(ENV_Wheat_umap$layout,P_frame[which(P_frame$Crop=="Wheat"),])
colnames(umap_plot_E_Wheat)[1:2]<-c("Dimension_1", "Dimension_2")


qplot(umap_plot_E_Wheat$Dimension_2,umap_plot_E_Wheat$Dimension_1,
      col = umap_plot_E_Wheat$Long,
      ylim = c(-5,5), xlim = c(-8,8),
      alpha = I(0.5))+theme_bw()

Umap_alldata_figure_Wheat<-
qplot(umap_plot_E_Wheat$Dimension_2,umap_plot_E_Wheat$Dimension_1,
                                 col = c(umap_plot_E_Wheat$Long),
                                 ylim = c(-5,5), xlim = c(-8,8),
                                 xlab  = "", ylab  ="",
                                 alpha = I(0))+
  theme_bw()+
  geom_point(size = 1)+
  scale_color_viridis(c(umap_plot_E_Wheat$Yield2), discrete  =F)


## gets only the BOM data
sub_mat<-Site_Modelframe[,grep("BOMDom_", colnames(Site_Modelframe))]

## trims data past 200 days after sowing, around the latest date you could
expect to harvest most crops
```

```
sub_mat<-sub_mat[,-c(grep("2[0-9]{2}$", colnames(sub_mat)))]
P_frame<-P_frame[!duplicated(paste(Site_Modelframe[,1],Site_Modelframe[,2],
sep = "_")),]
P_frame$Year<-
Site_Modelframe$METADom_Year[!duplicated(paste(Site_Modelframe[,1],Site_Modelframe[,2]
, sep = "_"))]
P_frame$JTS<-
Site_Modelframe$MANDom_JTS[!duplicated(paste(Site_Modelframe[,1],Site_Modelframe[,2]
, sep = "_"))]
P_frame$Lat<-
Site_Modelframe$BOMDom_Latitude[!duplicated(paste(Site_Modelframe[,1],Site_Modelframe[,2]
, sep = "_"))]
P_frame$Long<-
Site_Modelframe$BOMDom_Longitude[!duplicated(paste(Site_Modelframe[,1],Site_Modelframe[,2]
, sep = "_"))]
sub_mat<-sub_mat[!duplicated(paste(Site_Modelframe[,1],Site_Modelframe[,2],
sep = "_")),]
rownames(sub_mat)<-paste(sub_mat[,1],sub_mat[,2], sep = "_")

sub_mat<-as.matrix(sub_mat[,-c(1)])

BOM_umap<-umap(sub_mat,config = umap.defaults, method = "naive")

umap_plot_BOM<-data.frame(BOM_umap$layout,P_frame)
colnames(umap_plot_BOM)[1:2]<-c("Dimension_1", "Dimension_2")

qplot(aes(Dimension_1,Dimension_2, data = umap_plot_BOM), xlim = c(0,35),
ylim = c(-250,250), col = umap_plot_BOM$Year)+
  geom_point(col = umap_plot_BOM$Year)

qplot(umap_plot_BOM$Dimension_1,umap_plot_BOM$Dimension_2, xlim = c(0,35),
ylim = c(-250,250), col = umap_plot_BOM$Year)

## now just the BOM figure
Umap_BOM_figure<-qplot(umap_plot_BOM$Dimension_2,umap_plot_BOM$Dimension_1,
                       col = c(umap_plot_BOM$Year),
                       ylim = c(-50,50), xlim = c(-80,80),
                       xlab  = "Dimension 2", ylab  =" Dimension 1",
                       alpha = I(0.5))+
  scale_color_viridis("Year",c(umap_plot_BOM$Year), option  ="B")+
  theme_bw()+
  theme(legend.position = c(0.9,0.5))


## gets only the Satellite data
sub_mat<-Site_Modelframe[,grep("SatDom_", colnames(Site_Modelframe))]

## trims data past 200 days after sowing, around the latest date you could
expect to harvest most crops
sub_mat<-sub_mat[,-c(grep("2[0-9]{2}$", colnames(sub_mat)))]
sub_mat<-sub_mat[!duplicated(paste(Site_Modelframe[,1],Site_Modelframe[,2],
sep = "_")),]
rownames(sub_mat)<-c(paste(Site_Modelframe[,1],Site_Modelframe[,2], sep =
"_")[!duplicated(paste(Site_Modelframe[,1],Site_Modelframe[,2], sep =
"_"))])

sub_mat<-as.matrix(sub_mat[,-c(1)])

Sat_umap<-umap(sub_mat,config = umap.defaults, method = "naive")

umap_plot_SAT<-data.frame(Sat_umap$layout,P_frame)
colnames(umap_plot_SAT)[1:2]<-c("Dimension_1", "Dimension_2")

qplot(umap_plot_SAT$Dimension_1,umap_plot_SAT$Dimension_2,
      xlim = c(-15,15),
      ylim = c(-12,12),
      alpha = I(0.5),
      col = c(umap_plot_SAT$Year))

#sub_mat<-sub_mat[,-c(grep("1[0-9]{2}$", colnames(sub_mat)))]

############################################################

## Fig. SXX Plots satellite data at a single location-year

############################################################

One_site<-list.files("single_site_plots")

xobj<-read.csv(paste0("single_site_plots/",One_site[[1]]), header = T)

out_frame_single<-data.frame(Calendar_date = xobj$calendar_date,
                             Value = xobj$value)

colnames(out_frame_single)[colnames(out_frame_single)=="Value"]<-
as.character(xobj$band[[1]])

for(i in 2:length(One_site)){

  xobj<-read.csv(paste0("single_site_plots/",One_site[[i]]), header = T)

  xobj2<-data.frame(Calendar_date = xobj$calendar_date,
                    Value = xobj$value)

  colnames(xobj2)[colnames(xobj2)=="Value"]<-as.character(xobj$band[[1]])

  out_frame_single<-merge(out_frame_single, xobj2, by.x = "Calendar_date",
by.y = "Calendar_date", all = T)
}

out_frame_single$Calendar_date<-as.Date(out_frame_single$Calendar_date)

#out_frame_single<-out_frame_single[order(out_frame_single),]

## weeds out the fire mask
out_frame_single_fire<-
out_frame_single[!is.na(out_frame_single$Burn_Date),]

out_frame_single<-out_frame_single[is.na(out_frame_single$Burn_Date),]

## trial was sown 21/12/2010 (14964) and sown on 14 Jun 2010 (14774)

## 50% flowering dates were at 292-295 Julian days, 14902-14905

tiff(filename  = "Publication_docs/Single_site_plot.tiff",
     width  =7.5, height = 7, units = "in", res  =330)

## plots raw values
```

```
par(mfrow = c(5,5), mar = c(3.1,4.1,0.1,1.1), bty = "n")


plot(out_frame_single$Calendar_date, out_frame_single$PsnNet_500m.x,
     xlab = "",
     ylab = "Net Photosynth", pch = 20, las = 2)

abline(v = 14774, col = "red")
abline(v = 14964, col = "orange")
abline(v = 14903, col = rgb(0,0,1,0.5), lwd  =3)

plot(out_frame_single$Calendar_date[out_frame_single$ET_500m.x<1000],
out_frame_single$ET_500m.x[out_frame_single$ET_500m.x<1000],
     xlab = "",
     ylab = "EvapoTrans", pch = 20, las = 2)

abline(v = 14774, col = "red")
abline(v = 14964, col = "orange")
abline(v = 14903, col = rgb(0,0,1,0.5), lwd  =3)


plot(out_frame_single$Calendar_date[out_frame_single$PET_500m.x<1000],
out_frame_single$PET_500m.x[out_frame_single$PET_500m.x<1000],
     xlab = "",
     ylab = "Pot. EvapoTrans", pch = 20, las = 2)

abline(v = 14774, col = "red")
abline(v = 14964, col = "orange")
abline(v = 14903, col = rgb(0,0,1,0.5), lwd  =3)


plot(out_frame_single$Calendar_date, out_frame_single$Fpar_500m,
     xlab = "",
     ylab = "FPAR", pch = 20, las = 2)

abline(v = 14774, col = "red")
abline(v = 14964, col = "orange")
abline(v = 14903, col = rgb(0,0,1,0.5), lwd  =3)

plot(out_frame_single$Calendar_date, out_frame_single$Lai_500m,
     xlab = "",
     ylab = "LAI", pch = 20, las = 2)

abline(v = 14774, col = "red")
abline(v = 14964, col = "orange")
abline(v = 14903, col = rgb(0,0,1,0.5), lwd  =3)


plot(out_frame_single$Calendar_date, out_frame_single$`250m_16_days_EVI`,
     xlab = "",
     ylab = "EVI", pch = 20, las = 2)

abline(v = 14774, col = "red")
abline(v = 14964, col = "orange")
abline(v = 14903, col = rgb(0,0,1,0.5), lwd  =3)

plot(out_frame_single$Calendar_date, out_frame_single$`250m_16_days_NDVI`,
     xlab = "",
     ylab = "NDVI", pch = 20, las = 2)

abline(v = 14774, col = "red")
abline(v = 14964, col = "orange")
abline(v = 14903, col = rgb(0,0,1,0.5), lwd  =3)


plot(out_frame_single$Calendar_date, out_frame_single$Gpp_500m.x,
     xlab = "",
     ylab = "GPP", pch = 20, las = 2)

abline(v = 14774, col = "red")
abline(v = 14964, col = "orange")
abline(v = 14903, col = rgb(0,0,1,0.5), lwd  =3)

plot(out_frame_single$Calendar_date[out_frame_single$LST_Day_1km>10],
c((out_frame_single$LST_Day_1km*0.02)-
273.15)[out_frame_single$LST_Day_1km>10],
     xlab = "",
     ylab = "LST day", pch = 20, las = 2)

abline(v = 14774, col = "red")
abline(v = 14964, col = "orange")
abline(v = 14903, col = rgb(0,0,1,0.5), lwd  =3)


plot(out_frame_single$Calendar_date[out_frame_single$LST_Night_1km>10],
c((out_frame_single$LST_Night_1km*0.02)-
273.15)[out_frame_single$LST_Night_1km>10],
     xlab = "",
     ylab = "LST night", pch = 20, las = 2)

abline(v = 14774, col = "red")
abline(v = 14964, col = "orange")
abline(v = 14903, col = rgb(0,0,1,0.5), lwd  =3)


plot(out_frame_single$Calendar_date, out_frame_single$Clear_sky_days,
     xlab = "",
     ylab = "Clear day", pch = 20, las = 2)

abline(v = 14774, col = "red")
abline(v = 14964, col = "orange")
abline(v = 14903, col = rgb(0,0,1,0.5), lwd  =3)

plot(out_frame_single$Calendar_date, out_frame_single$Clear_sky_days,
     xlab = "",
     ylab = "Clear Night", pch = 20, las = 2)

abline(v = 14774, col = "red")
abline(v = 14964, col = "orange")
abline(v = 14903, col = rgb(0,0,1,0.5), lwd  =3)

plot(out_frame_single$Calendar_date, out_frame_single$sur_refl_b01,
     xlab = "",
     ylab = "Surf_ref_wave01", pch = 20, ylim = c(250,2500), las = 2)

abline(v = 14774, col = "red")
abline(v = 14964, col = "orange")
abline(v = 14903, col = rgb(0,0,1,0.5), lwd  =3)
```

```
plot(out_frame_single$Calendar_date, out_frame_single$sur_refl_b02,
     xlab = "",
     ylab = "Surf_ref_wave02", pch = 20, las = 2)

abline(v = 14774, col = "red")
abline(v = 14964, col = "orange")
abline(v = 14903, col = rgb(0,0,1,0.5), lwd  =3)

plot(out_frame_single$Calendar_date, out_frame_single$sur_refl_b03,
     xlab = "",
     ylab = "Surf_ref_wave03", pch = 20, ylim = c(125,1250), las = 2)

abline(v = 14774, col = "red")
abline(v = 14964, col = "orange")
abline(v = 14903, col = rgb(0,0,1,0.5), lwd  =3)

plot(out_frame_single$Calendar_date, out_frame_single$sur_refl_b04,
     xlab = "",
     ylab = "Surf_ref_wave04", pch = 20, ylim = c(500,1500))

abline(v = 14774, col = "red")
abline(v = 14964, col = "orange")
abline(v = 14903, col = rgb(0,0,1,0.5), lwd  =3)

plot(out_frame_single$Calendar_date, out_frame_single$sur_refl_b05,
     xlab = "",
     ylab = "Surf_ref_wave05", pch = 20, las = 2)

abline(v = 14774, col = "red")
abline(v = 14964, col = "orange")
abline(v = 14903, col = rgb(0,0,1,0.5), lwd  =3)


plot(out_frame_single$Calendar_date, out_frame_single$sur_refl_b06,
     xlab = "",
     ylab = "Surf_ref_wave06", pch = 20, las = 2)

abline(v = 14774, col = "red")
abline(v = 14964, col = "orange")
abline(v = 14903, col = rgb(0,0,1,0.5), lwd  =3)

plot(out_frame_single$Calendar_date, out_frame_single$sur_refl_b07,
     xlab = "",
     ylab = "Surf_ref_wave07", pch = 20, las = 2)

abline(v = 14774, col = "red")
abline(v = 14964, col = "orange")
abline(v = 14903, col = rgb(0,0,1,0.5), lwd  =3)

#plot(out_frame_single$Calendar_date, out_frame_single$Emis_31,
#     xlab = "",
#     ylab = "Emis 31", pch = 20)

#abline(v = 14708, col = "red", lty = 3)

plot(out_frame_single$Calendar_date, out_frame_single$Emis_32,
     xlab = "",
     ylab = "Emis 32", pch = 20, las = 2)

abline(v = 14774, col = "red")
abline(v = 14964, col = "orange")
abline(v = 14903, col = rgb(0,0,1,0.5), lwd  =3)

plot(out_frame_single$Calendar_date,
out_frame_single$`250m_16_days_blue_reflectance`,
     xlab = "",
     ylab = "Blue", pch = 20, las = 2)

abline(v = 14774, col = "red")
abline(v = 14964, col = "orange")
abline(v = 14903, col = rgb(0,0,1,0.5), lwd  =3)


plot(out_frame_single$Calendar_date,
out_frame_single$`250m_16_days_MIR_reflectance`,
     xlab = "",
     ylab = "MIR", pch = 20, las = 2)

abline(v = 14774, col = "red")
abline(v = 14964, col = "orange")
abline(v = 14903, col = rgb(0,0,1,0.5), lwd  =3)

plot(out_frame_single$Calendar_date,
out_frame_single$`250m_16_days_NIR_reflectance`,
     xlab = "",
     ylab = "NIR", pch = 20, las = 2)

abline(v = 14774, col = "red")
abline(v = 14964, col = "orange")
abline(v = 14903, col = rgb(0,0,1,0.5), lwd  =3)

plot(out_frame_single$Calendar_date[out_frame_single$LE_500m.x<1000],
out_frame_single$LE_500m.x[out_frame_single$LE_500m.x<1000],
     xlab = "",
     ylab = "L.Heat flux", pch = 20, las = 2)

abline(v = 14774, col = "red")
abline(v = 14964, col = "orange")
abline(v = 14903, col = rgb(0,0,1,0.5), lwd  =3)

plot(out_frame_single_fire$Calendar_date, out_frame_single_fire$Burn_Date,
     xlab = "",
     ylab = "Fires", pch = 20, las = 2)

abline(v = 14774, col = "red")
abline(v = 14964, col = "orange")
abline(v = 14903, col = rgb(0,0,1,0.5), lwd  =3)

par(mfrow = c(1,1), mar = c(5.1,4.1,4.1,2.1))

dev.off()


## Splits into vegetation products and other

tiff(filename  = "Publication_docs/Single_site_plot_vegetation.tiff",
     width  =7.5, height = 4, units = "in", res  =330)
```

```
## plots raw values
par(mfrow = c(2,5), mar = c(3.1,4.1,2.1,1.1), bty = "n")

plot(out_frame_single$Calendar_date, out_frame_single$PsnNet_500m.x,
     xlab = "",
     ylim = c(0, 500),
     ylab = "Net Photosynthesis", pch = 20, las = 2)

mtext("a", adj = 0)
abline(v = 14774, col = "red")
abline(v = 14964, col = "orange")
abline(v = 14903, col = rgb(0,0,1,0.5), lwd  =3)

plot(out_frame_single$Calendar_date[out_frame_single$ET_500m.x<1000],
out_frame_single$ET_500m.x[out_frame_single$ET_500m.x<1000],
     xlab = "",
     ylim = c(0,210),
     ylab = "Evapotranspiration", pch = 20, las = 2)

mtext("b", adj = 0)
abline(v = 14774, col = "red")
abline(v = 14964, col = "orange")
abline(v = 14903, col = rgb(0,0,1,0.5), lwd  =3)


plot(out_frame_single$Calendar_date[out_frame_single$PET_500m.x<1000],
out_frame_single$PET_500m.x[out_frame_single$PET_500m.x<1000],
     xlab = "",
     ylim = c(0,680),
     ylab = "Potential Evapotrans.", pch = 20, las = 2)

mtext("c", adj = 0)
abline(v = 14774, col = "red")
abline(v = 14964, col = "orange")
abline(v = 14903, col = rgb(0,0,1,0.5), lwd  =3)


plot(out_frame_single$Calendar_date, out_frame_single$Fpar_500m,
     xlab = "",
     ylim = c(0,100),
     ylab = "FPAR", pch = 20, las = 2)

mtext("d", adj = 0)
abline(v = 14774, col = "red")
abline(v = 14964, col = "orange")
abline(v = 14903, col = rgb(0,0,1,0.5), lwd  =3)

plot(out_frame_single$Calendar_date, out_frame_single$Lai_500m,
     xlab = "",
     ylab = "Leaf Area Index", pch = 20, las = 2)

mtext("d", adj = 0)
abline(v = 14774, col = "red")
abline(v = 14964, col = "orange")
abline(v = 14903, col = rgb(0,0,1,0.5), lwd  =3)


plot(out_frame_single$Calendar_date, out_frame_single$`250m_16_days_EVI`,
     xlab = "",
     ylim = c(0,7200),
     ylab = "Enhanced Vegetation Index", pch = 20, las = 2)

mtext("f", adj = 0)
abline(v = 14774, col = "red")
abline(v = 14964, col = "orange")
abline(v = 14903, col = rgb(0,0,1,0.5), lwd  =3)

plot(out_frame_single$Calendar_date, out_frame_single$`250m_16_days_NDVI`,
     xlab = "",
     ylab = "NDVI", pch = 20, las = 2)

mtext("g", adj = 0)
abline(v = 14774, col = "red")
abline(v = 14964, col = "orange")
abline(v = 14903, col = rgb(0,0,1,0.5), lwd  =3)


plot(out_frame_single$Calendar_date, out_frame_single$Gpp_500m.x,
     xlab = "",
     ylim = c(0,560),
     ylab = "Gross Primary Productivity", pch = 20, las = 2)

mtext("h", adj = 0)
abline(v = 14774, col = "red")
abline(v = 14964, col = "orange")
abline(v = 14903, col = rgb(0,0,1,0.5), lwd  =3)


plot(out_frame_single$Calendar_date[out_frame_single$LE_500m.x<1000],
out_frame_single$LE_500m.x[out_frame_single$LE_500m.x<1000],
     xlab = "",
     ylim = c(0,480),
     ylab = "Latent Heat Flux", pch = 20, las = 2)

mtext("i", adj = 0)
abline(v = 14774, col = "red")
abline(v = 14964, col = "orange")
abline(v = 14903, col = rgb(0,0,1,0.5), lwd  =3)

plot(out_frame_single_fire$Calendar_date, out_frame_single_fire$Burn_Date,
     xlab = "",
     ylim = c(0,1),
     ylab = "Fires", pch = 20, las = 2)

mtext("j", adj = 0)
abline(v = 14774, col = "red")
abline(v = 14964, col = "orange")
abline(v = 14903, col = rgb(0,0,1,0.5), lwd  =3)

par(mfrow = c(1,1), mar = c(5.1,4.1,4.1,2.1))

dev.off()



## Also plots Net photosynthesis separately for Figure 1.
```

```
## trial was harvested 21/12/2010 (14964) and sown on 14 Jun 2010 (14774)

Photosynth_Fig1b<-qplot(out_frame_single$Calendar_date,
out_frame_single$PsnNet_500m.x/8,
                        xlab = "",
                        ylab = "Net Photosynthesis\n (kg C/mÂ² per day)",
alpha = I(0))+
  geom_smooth(col = "black", bg = "grey", span = 0.3)+
  geom_point(cex = 0.5)+
  theme_bw()+
  geom_vline(xintercept=14774, col = "red")+
  geom_vline(xintercept=14964, col = "orange")+
  geom_vline(xintercept = 14903, col = "lightblue", alpha = I(0.5), lwd =
3)+
  annotate("text", label = "Sowing", x = as.Date(14765), y = 350/8, colour
= "red", angle = 90)+
  annotate("text", label = "Anthesis", x = as.Date(14891), y = 50/8, colour
= "darkblue", angle = 90)+
  annotate("text", label = "Harvest", x = as.Date(14955), y = 350/8, colour
= "orange", angle = 90)


ggsave(Photosynth_Fig1b,file = "Publication_docs/Fig1b.tiff", device =
"tiff", height = 6, width = 5,dpi = 330)


Photosynth_Fig1b+theme(text = element_text(size=10),
                       axis.text = element_text(size=10))

##########################################################

## Figure 2. Makes a figure of ML model accuracy, by method, for yield

##########################################################

predobs_frame_unscaled<-readRDS(file =
"model_frames_and_imputations/siteVariety_models/accuracy_predobs_alldata1.rds"
)
predobs_frame_scaled<-readRDS(file =
"model_frames_and_imputations/siteVariety_models/accuracy_predobs_alldata2.rds"
)

bestfit_reg<-odregress(predobs_frame_unscaled[[3]][,1],
                       predobs_frame_unscaled[[3]]$Rrf_200)

## plots internal accuracy vs OOB accuracy
Fig_Rrf1<-qplot(predobs_frame_unscaled[[3]][,1],
jitter(predobs_frame_unscaled[[3]]$Rrf_200, factor =200),
                alpha = I(0),
                xlim = c(0,10), ylim = c(0,10),
                ylab = "Predicted Yield (t/Ha)", xlab = "Observed Yield
(t/Ha)")+
  geom_point(pch = 20, cex = 0.5, alpha = 0.6, col = "orange")+
  geom_abline(slope = 1, intercept = 0, col = "darkgrey")+
  geom_density2d()+
  geom_abline(slope = bestfit_reg$coeff[1], intercept =
bestfit_reg$coeff[2], col = "black")+
  theme_bw()+
  annotate("text", x = 4, y = 9, cex = 3, label = "RÂ² = 0.82; RMSE =
0.94")

cor.test(predobs_frame_unscaled[[3]][,1],
predobs_frame_unscaled[[3]]$Rrf_200)
rmserr(predobs_frame_unscaled[[3]][,1],
predobs_frame_unscaled[[3]]$Rrf_200)$rmse

bestfit_reg<-odregress(predobs_frame_unscaled[[2]][,1],
                       predobs_frame_unscaled[[2]]$Rrf_200)

Fig_Rrf2<-qplot(predobs_frame_unscaled[[2]][,1],
predobs_frame_unscaled[[2]]$Rrf_200,
                alpha = I(0),
                xlim = c(0,12), ylim = c(0,12),
                ylab = "Predicted Yield (t/Ha)", xlab = "Observed Yield
(t/Ha)")+
  geom_abline(slope = 1, intercept = 0, col = "darkgrey")+
  geom_hex(bins = 50)+
  geom_abline(slope = bestfit_reg$coeff[1], intercept =
bestfit_reg$coeff[2], col = "orange", alpha = I(0.8))+
  theme_bw()+
  scale_fill_viridis("Observations",option = "B", limits=c(0, 130))+
  theme(legend.position = "none")+
  annotate("text", x = 4, y = 9, cex = 3, label = "RÂ² = 0.70; RMSE =
1.24")

round(rmserr(predobs_frame_unscaled[[2]][,1],
predobs_frame_unscaled[[2]]$Rrf_200)$rmse, digits = 2)

bestfit_reg<-odregress(predobs_frame_unscaled[[3]][,1],
                       predobs_frame_unscaled[[3]]$rp_200)

## plots internal accuracy vs OOB accuracy
Fig_rp1<-qplot(predobs_frame_unscaled[[3]][,1],
jitter(predobs_frame_unscaled[[3]]$rp_200, factor =200),
               alpha = I(0),
               xlim = c(0,10), ylim = c(0,10),
               ylab = "Predicted Yield (t/Ha)", xlab = "Observed Yield
(t/Ha)")+
  geom_point(pch = 20, cex = 0.5, alpha = 0.6, col = "orange")+
  geom_abline(slope = 1, intercept = 0, col = "darkgrey")+
  geom_density2d()+
  geom_abline(slope = bestfit_reg$coeff[1], intercept =
bestfit_reg$coeff[2], col = "black")+
  theme_bw()+
  annotate("text", x = 4, y = 9, cex = 3, label = "RÂ² = 0.76; RMSE =
1.11")

cor.test(predobs_frame_unscaled[[3]][,1],predobs_frame_unscaled[[3]]$rp_200
)
round(rmserr(predobs_frame_unscaled[[3]][,1],predobs_frame_unscaled[[3]]$rp_200)$rmse
, digits = 2)

bestfit_reg<-odregress(predobs_frame_unscaled[[2]][,1],
                       predobs_frame_unscaled[[2]]$rp_200)

Fig_rp2<-qplot(predobs_frame_unscaled[[2]][,1],
predobs_frame_unscaled[[2]]$rp_200,
               alpha = I(0),
```

```
               xlim = c(0,12), ylim = c(0,12),
               ylab = "Predicted Yield (t/Ha)", xlab = "Observed Yield
(t/Ha)")+
  geom_abline(slope = 1, intercept = 0, col = "darkgrey")+
  geom_hex(bins = 50)+
  geom_abline(slope = bestfit_reg$coeff[1], intercept =
bestfit_reg$coeff[2], col = "orange", alpha = I(0.8))+
  theme_bw()+
  scale_fill_viridis("Observations",option = "B", limits=c(0, 130))+
  theme(legend.position = "none")+
  annotate("text", x = 4, y = 9, cex = 3, label = "RÂ² = 0.59, RMSE =
1.42")

round(rmserr(predobs_frame_unscaled[[2]][,1],
predobs_frame_unscaled[[2]]$rp_200)$rmse, digits= 2)

bestfit_reg<-odregress(predobs_frame_unscaled[[3]][,1],
                       predobs_frame_unscaled[[3]]$xvrf_200)

## plots internal accuracy vs OOB accuracy
Fig_xvrf1<-qplot(predobs_frame_unscaled[[3]][,1],
jitter(predobs_frame_unscaled[[3]]$xvrf_200, factor =200),
                 alpha = I(0),
                 xlim = c(0,10), ylim = c(0,10),
                 ylab = "Predicted Yield (t/Ha)", xlab = "Observed Yield
(t/Ha)")+
  geom_point(pch = 20, cex = 0.5, alpha = 0.6, col = "orange")+
  geom_abline(slope = 1, intercept = 0, col = "darkgrey")+
  geom_density2d()+
  geom_abline(slope = bestfit_reg$coeff[1], intercept =
bestfit_reg$coeff[2], col = "black")+
  theme_bw()+
  annotate("text", x = 4, y = 9, cex = 3, label = "RÂ² = 0.84; RMSE =
0.91")

cor.test(predobs_frame_unscaled[[3]][,1],
predobs_frame_unscaled[[3]]$xvrf_200)
round(rmserr(predobs_frame_unscaled[[3]][,1],predobs_frame_unscaled[[3]]$xvrf_200)$rmse
, digits = 2)

## the older design
qplot(predobs_frame_unscaled[[3]][,1],
predobs_frame_unscaled[[3]]$xvrf_200,
      alpha = I(0),
      xlim = c(0,10), ylim = c(0,10),
      ylab = "Predicted Yield (t/Ha)", xlab = "Observed Yield (t/Ha)")+
  geom_abline(slope = 1, intercept = 0, col = "darkgrey")+
  geom_hex(bins = 50)+
  geom_abline(slope = bestfit_reg$coeff[1], intercept =
bestfit_reg$coeff[2], col = "orange", alpha = I(0.8))+
  theme_bw()+
  scale_fill_viridis("Observations",option = "B", limits=c(0, 90))+
  theme(legend.position = "none")+
  annotate("text", x = 4, y = 9, cex = 3, label = "RÂ² = 0.84")

bestfit_reg<-odregress(predobs_frame_unscaled[[2]][,1],
                       predobs_frame_unscaled[[2]]$xvrf_200)

Fig_xvrf2<-qplot(predobs_frame_unscaled[[2]][,1],
predobs_frame_unscaled[[2]]$xvrf_200,
                 alpha = I(0),
                 xlim = c(0,12), ylim = c(0,12),
                 ylab = "Predicted Yield (t/Ha)", xlab = "Observed Yield
(t/Ha)")+
  geom_abline(slope = 1, intercept = 0, col = "darkgrey")+
  geom_hex(bins = 50)+
  geom_abline(slope = bestfit_reg$coeff[1], intercept =
bestfit_reg$coeff[2], color = "orange", alpha = I(0.8))+
  theme_bw()+
  scale_fill_viridis("Observations",option = "B", limits=c(0, 130))+
  theme(legend.position = "none")+
  annotate("text", x = 4, y = 9, cex = 3, label = "RÂ² = 0.69; RMSE =
1.23")

cor.test(predobs_frame_unscaled[[2]][,1],
predobs_frame_unscaled[[2]]$xvrf_200)
round(rmserr(predobs_frame_unscaled[[2]][,1],predobs_frame_unscaled[[2]]$xvrf_200)$rmse
, digits = 2)

bestfit_reg<-odregress(predobs_frame_scaled[[3]][,1],
                       predobs_frame_scaled[[3]]$xg_200)

## plots internal accuracy vs OOB accuracy
Fig_xg1<-qplot(predobs_frame_scaled[[3]][,1],
jitter(predobs_frame_scaled[[3]]$xg_200, factor =200),
               alpha = I(0),
               xlim = c(0,10), ylim = c(0,10),
               ylab = "Predicted Yield (t/Ha)", xlab = "Observed Yield
(t/Ha)")+
  geom_point(pch = 20, cex = 0.5, alpha = 0.6, col = "orange")+
  geom_abline(slope = 1, intercept = 0, col = "darkgrey")+
  geom_density2d()+
  geom_abline(slope = bestfit_reg$coeff[1], intercept =
bestfit_reg$coeff[2], col = "black")+
  theme_bw()+
  annotate("text", x = 4, y = 9, cex = 3, label = "RÂ² = 0.78; RMSE =
0.90")

cor.test(predobs_frame_scaled[[3]][,1], predobs_frame_scaled[[3]]$xg_200)
round(rmserr(predobs_frame_scaled[[3]][,1],predobs_frame_scaled[[3]]$xg_200)$rmse
, digits = 2)

bestfit_reg<-odregress(predobs_frame_scaled[[2]][,1],
                       predobs_frame_scaled[[2]]$xg_200)

Fig_xg2<-qplot(predobs_frame_scaled[[2]][,1],
predobs_frame_scaled[[2]]$xg_200,
               alpha = I(0),
               xlim = c(0,12), ylim = c(0,12),
               ylab = "Predicted Yield (t/Ha)", xlab = "Observed Yield
(t/Ha)")+
  geom_abline(slope = 1, intercept = 0, col = "darkgrey")+
  geom_hex(bins = 50)+
  geom_abline(slope = bestfit_reg$coeff[1], intercept =
bestfit_reg$coeff[2], color = "orange", alpha = I(0.8))+
  theme_bw()+
  scale_fill_viridis("Observations",option = "B", limits=c(0, 130))+
  theme(legend.position = "none")+
```

```
  annotate("text", x = 4, y = 9, cex = 3, label =  "RÂ² = 0.73; RMSE =
1.10")

cor.test(predobs_frame_scaled[[2]][,1], predobs_frame_scaled[[2]]$xg_200)
round(rmserr(predobs_frame_scaled[[2]][,1],predobs_frame_scaled[[2]]$xg_200)$rmse
, digits = 2)

bestfit_reg<-odregress(predobs_frame_scaled[[3]][,1],
                       predobs_frame_scaled[[3]]$lsvm_200)

## plots internal accuracy vs OOB accuracy
Fig_lsvm1<-qplot(predobs_frame_scaled[[3]][,1],
jitter(predobs_frame_scaled[[3]]$lsvm_200, factor =200),
                 alpha = I(0),
                 xlim = c(0,10), ylim = c(0,10),
                 ylab = "Predicted Yield (t/Ha)", xlab = "Observed Yield
(t/Ha)")+
  geom_point(pch = 20, cex = 0.5, alpha = 0.6, col = "orange")+
  geom_abline(slope = 1, intercept = 0, col = "darkgrey")+
  geom_density2d()+
  geom_abline(slope = bestfit_reg$coeff[1], intercept =
bestfit_reg$coeff[2], col = "black")+
  theme_bw()+
  annotate("text", x = 4, y = 9, cex = 3, label = "RÂ² = 0.68; RMSE =
1.09")

cor.test(predobs_frame_scaled[[3]][,1], predobs_frame_scaled[[3]]$lsvm_200)
round(rmserr(predobs_frame_scaled[[3]][,1],predobs_frame_scaled[[3]]$lsvm_200)$rmse
, digits = 2)

bestfit_reg<-odregress(predobs_frame_scaled[[2]][,1],
                       predobs_frame_scaled[[2]]$lsvm_200)

Fig_lsvm2<-qplot(predobs_frame_scaled[[2]][,1],
predobs_frame_scaled[[2]]$lsvm_200,
                 alpha = I(0),
                 xlim = c(0,12), ylim = c(0,12),
                 ylab = "Predicted Yield (t/Ha)", xlab = "Observed Yield
(t/Ha)")+
  geom_abline(slope = 1, intercept = 0, col = "darkgrey")+
  geom_hex(bins = 50)+
  geom_abline(slope = bestfit_reg$coeff[1], intercept =
bestfit_reg$coeff[2], col = "orange", alpha = I(0.8))+
  theme_bw()+
  scale_fill_viridis("Observations",option = "B", limits=c(0, 130))+
  theme(legend.position = "none")+
  annotate("text", x = 4, y = 9, cex = 3, label = "RÂ² = 0.45; RMSE =
1.43")

cor.test(predobs_frame_scaled[[2]][,1], predobs_frame_scaled[[2]]$lsvm_200)
round(rmserr(predobs_frame_scaled[[2]][,1],predobs_frame_scaled[[2]]$lsvm_200)$rmse
, digits = 2)

bestfit_reg<-odregress(predobs_frame_scaled[[3]][,1],
                       predobs_frame_scaled[[3]]$plsr_200)

## plots internal accuracy vs OOB accuracy
Fig_plsr1<-qplot(predobs_frame_scaled[[3]][,1],
jitter(predobs_frame_scaled[[3]]$plsr_200, factor =200), alpha = I(0),
                 xlim = c(0,10), ylim = c(0,10),
                 ylab = "Predicted Yield (t/Ha)", xlab = "Observed Yield
(t/Ha)")+
  geom_point(pch = 20, cex = 0.5, alpha = 0.6, col = "orange")+
  geom_abline(slope = 1, intercept = 0, col = "darkgrey")+
  geom_density2d()+
  geom_abline(slope = bestfit_reg$coeff[1], intercept =
bestfit_reg$coeff[2], col = "black")+
  theme_bw()+
  annotate("text", x = 4, y = 9, cex = 3, label = "RÂ² = 0.76; RMSE =
0.94")

cor.test(predobs_frame_scaled[[3]][,1], predobs_frame_scaled[[3]]$plsr_200)
round(rmserr(predobs_frame_scaled[[3]][,1],predobs_frame_scaled[[3]]$plsr_200)$rmse
, digits = 2)

#  scale_fill_viridis("Observations",option = "B", limits=c(0, 90))+
#  theme(legend.position = c(0.2, 0.7))+
#  annotate("text", x = 7.5, y = 0.5, label = "RÂ² = 0.76")

bestfit_reg<-odregress(predobs_frame_scaled[[2]][,1],
                       predobs_frame_scaled[[2]]$plsr_200)

Fig_plsr2<-qplot(predobs_frame_scaled[[2]][,1],
predobs_frame_scaled[[2]]$plsr_200,
                 alpha = I(0),
                 xlim = c(0,12), ylim = c(0,12),
                 ylab = "Predicted Yield (t/Ha)", xlab = "Observed Yield
(t/Ha)")+
  geom_abline(slope = 1, intercept = 0, col = "darkgrey")+
  geom_hex(bins = 50)+
  geom_abline(slope = bestfit_reg$coeff[1], intercept =
bestfit_reg$coeff[2], col = "orange")+
  theme_bw()+
  scale_fill_viridis("Observations",option = "B", limits=c(0, 130))+
  theme(legend.position = c(0.2, 0.7))+
  annotate("text",  x = 4, y = 9, cex = 3,  label = "RÂ² = 0.74; RMSE =
1.09")

cor.test(predobs_frame_scaled[[2]][,1], predobs_frame_scaled[[2]]$plsr_200)
round(rmserr(predobs_frame_scaled[[2]][,1],predobs_frame_scaled[[2]]$plsr_200)$rmse
, digits = 2)


## plots internal accuracy vs OOB accuracy
Fig_plsr1<-qplot(predobs_frame_scaled[[3]][,1],
jitter(predobs_frame_scaled[[3]]$plsr_200, factor =100),
                 alpha = I(0),
                 xlim = c(0,10), ylim = c(0,10),
                 ylab = "Predicted Yield (t/Ha)", xlab = "Observed Yield
(t/Ha)")+
  geom_point(pch = 20, cex = 0.5, alpha = 0.6, col = "orange")+
  geom_abline(slope = 1, intercept = 0, col = "darkgrey")+
  geom_density2d()+
  geom_abline(slope = bestfit_reg$coeff[1], intercept =
bestfit_reg$coeff[2], col = "black")+
  theme_bw()+
  annotate("text", x = 4, y = 9, cex = 3, label = "RÂ² = 0.76; RMSE =
0.94")
```

```
cor.test(predobs_frame_scaled[[3]][,1], predobs_frame_scaled[[3]]$plsr_200)
round(rmserr(predobs_frame_scaled[[3]][,1],predobs_frame_scaled[[3]]$plsr_200)$rmse
, digits = 2)

## too compact
tworow_acc<-grid.arrange(Fig_rp1, Fig_Rrf1,  Fig_xvrf1, Fig_xg1, Fig_lsvm1,
Fig_plsr1,
                         Fig_rp2, Fig_Rrf2, Fig_xvrf2, Fig_xg2, Fig_lsvm2,
Fig_plsr2, nrow = 2)

set.seed(8364)
## better...
tworow_acc<-grid.arrange(Fig_rp1+labs(title = "a"),
                         Fig_Rrf1+labs(title = "b"),
                         Fig_xvrf1+labs(title = "c"),
                         Fig_xg1+labs(title = "d"),
                         Fig_lsvm1+labs(title = "e"),
                         Fig_plsr1+labs(title = "f"),
                         nrow = 2)

ggsave(tworow_acc,file = "Publication_docs/Fig_2.tiff", device = "tiff",
height = 5, width = 7,dpi = 330)

# RÂ²
cor.test(predobs_frame_unscaled[[3]][,1],
predobs_frame_unscaled[[3]]$rp_200)
cor.test(predobs_frame_unscaled[[3]][,1],
predobs_frame_unscaled[[3]]$Rrf_200)
cor.test(predobs_frame_unscaled[[3]][,1],
predobs_frame_unscaled[[3]]$xvrf_200)
cor.test(predobs_frame_scaled[[3]][,1], predobs_frame_scaled[[3]]$lsvm_200)
cor.test(predobs_frame_scaled[[3]][,1], predobs_frame_scaled[[3]]$xg_200)
cor.test(predobs_frame_scaled[[3]][,1], predobs_frame_scaled[[3]]$plsr_200)

tworow_acc<-grid.arrange(Fig_rp2+labs(title = "a"), Fig_Rrf2+labs(title =
"b"),  Fig_xvrf2+labs(title = "c"),
                         Fig_xg2+labs(title = "d"), Fig_lsvm2+labs(title =
"e"), Fig_plsr2+labs(title = "f"),
                         nrow = 2)
## Saves figure 2
ggsave(tworow_acc,file = "Publication_docs/Fig_2_forward_projection.tiff",
device = "tiff", height = 6, width = 7.5,dpi = 330)


## prints R2 for missing trial site predictions
for(i in 1:ncol(predobs_frame_unscaled[[3]])){
  print(c(colnames(predobs_frame_unscaled[[3]])[i]))
  print(cor.test(predobs_frame_unscaled[[3]]$Yield,
predobs_frame_unscaled[[3]][,i])$estimate)}

for(i in 1:ncol(predobs_frame_scaled[[3]])){
  print(c(colnames(predobs_frame_scaled[[3]])[i]))
  print(cor.test(predobs_frame_scaled[[3]]$Yield,
predobs_frame_scaled[[3]][,i])$estimate)}


## prints R2 for annual forecasts
for(i in 1:ncol(predobs_frame_unscaled[[2]])){
  print(c(colnames(predobs_frame_unscaled[[2]])[i]))
  print(round(cor.test(predobs_frame_unscaled[[2]]$Yield,
predobs_frame_unscaled[[2]][,i])$estimate, digits  =2))}

for(i in 1:ncol(predobs_frame_scaled[[2]])){
  print(c(colnames(predobs_frame_scaled[[2]])[i]))
  print(round(cor.test(predobs_frame_scaled[[2]]$Yield,
predobs_frame_scaled[[2]][,i])$estimate, digits  =2))}


## prints R2 for (somewhat bullshit) random-sample holdout accuracy
for(i in 1:ncol(predobs_frame_unscaled[[1]])){
  print(c(colnames(predobs_frame_unscaled[[1]])[i]))
  print(round(cor.test(predobs_frame_unscaled[[1]]$Yield,
predobs_frame_unscaled[[1]][,i])$estimate, digits  =2))}

for(i in 1:ncol(predobs_frame_scaled[[1]])){
  print(c(colnames(predobs_frame_scaled[[1]])[i]))
  print(round(cor.test(predobs_frame_scaled[[1]]$Yield,
predobs_frame_scaled[[1]][,i])$estimate, digits  =2))}


## saves a different plot for the training schematic in supp fig 2

bestfit_reg<-odregress(predobs_frame_unscaled[[3]][,1],
                       predobs_frame_unscaled[[3]]$rp_200)

Schematic1<-qplot(predobs_frame_unscaled[[3]][,1],
jitter(predobs_frame_unscaled[[3]]$rp_200, factor =200),
                  alpha = I(0),
                  xlim = c(0,10), ylim = c(0,10),
                  ylab = "Predicted Yield (t/Ha)", xlab = "Observed Yield
(t/Ha)")+
  geom_point(pch = 20, cex = 0.5, alpha = 0.6, col = "orange")+
  geom_abline(slope = 1, intercept = 0, col = "darkgrey")+
  # geom_density2d()+
  geom_abline(slope = bestfit_reg$coeff[1], intercept =
bestfit_reg$coeff[2], col = "darkgreen", lwd = 2, alpha = 0.7)+
  theme_bw()+
  annotate("text", x = 7.5, y = 0.5, label = "RÂ² = 0.76")

cor.test(predobs_frame_unscaled[[3]][,1],predobs_frame_unscaled[[3]]$rp_200
)

bestfit_reg<-odregress(predobs_frame_unscaled[[2]][,1],
                       predobs_frame_unscaled[[2]]$rp_200)
## and the annual forecast data
Schematic2<-qplot(predobs_frame_unscaled[[2]][,1],
jitter(predobs_frame_unscaled[[2]]$rp_200, factor =200),
                  alpha = I(0),
                  xlim = c(0,10), ylim = c(0,10),
                  ylab = "Predicted Yield (t/Ha)", xlab = "Observed Yield
(t/Ha)")+
  geom_point(pch = 20, cex = 0.5,col = "lightblue")+
  geom_abline(slope = 1, intercept = 0, col = "darkgrey")+
  # geom_density2d()+
  geom_abline(slope = bestfit_reg$coeff[1], intercept =
bestfit_reg$coeff[2], col = "darkgreen", lwd = 2, alpha = 0.7)+
  theme_bw()+
```

```
  annotate("text", x = 7.5, y = 0.5, label = "RÂ² = 0.59")

cor.test(predobs_frame_unscaled[[2]][,1],predobs_frame_unscaled[[2]]$rp_200
)

Schematic3<-grid.arrange(Schematic1, Schematic2, nrow = 1)

ggsave(Schematic3,file = "Publication_docs/Fig_S2XX_Schematic.tiff", device
= "tiff", height = 4, width = 7.5,dpi = 330)


grid.arrange(Fig_rp1, Fig_Rrf1,  Fig_xvrf1,
             Fig_xg1, Fig_lsvm1, Fig_plsr1,
             Fig_rp2, Fig_Rrf2, Fig_xvrf2,
             Fig_xg2, Fig_lsvm2, Fig_plsr2, nrow = 4)

grid.arrange(Fig_rp1, Fig_Rrf1,  Fig_xvrf1,
             Fig_xg1, Fig_lsvm1, Fig_plsr1, nrow = 2)

rm(rf_mod, Rrf_mod, rpart_mod,xg_mod,lsvm_mod,pslr_mod, mod_frame3)


#xg_mod<-readRDS(file =
"model_frames_and_imputations/siteVariety_models/xgb_TOS.rds")
#lsvm_mod<-readRDS(file =
"model_frames_and_imputations/siteVariety_models/lsvm_TOS.rds")
#pslr_mod<-readRDS(file =
"model_frames_and_imputations/siteVariety_models/plsr_TOS.rds")

## and the yield data for prediction accuracy
mod_frame1<-readRDS(file = "model_frames_and_imputations/mod_frame1.rds")
beep(5)

peanutCluster<-makeCluster(detectCores()-2)
registerDoParallel(peanutCluster)

## first, separates phenotypes
Phen_data<-mod_frame1[,grep("PHENDom", colnames(mod_frame1))]
mod_frame1<-mod_frame1[,-c(grep("PHENDom", colnames(mod_frame1)))]

## produces a lookup table of phenotypes by species
Phen_summary_table<-data.frame(All = colSums(!is.na(Phen_data)),
                               Barley =
colSums(!is.na(Phen_data[Phen_data$PHENDom_Crop.Type=="Barley",])),
                               Canola =
colSums(!is.na(Phen_data[Phen_data$PHENDom_Crop.Type=="Canola",])),
                               Chickpea =
colSums(!is.na(Phen_data[Phen_data$PHENDom_Crop.Type=="Chickpea",])),
                               Faba =
colSums(!is.na(Phen_data[Phen_data$PHENDom_Crop.Type=="Faba Bean",])),
                               Lupin =
colSums(!is.na(Phen_data[Phen_data$PHENDom_Crop.Type=="Lupin",])),
                               Oat =
colSums(!is.na(Phen_data[Phen_data$PHENDom_Crop.Type=="Oat",])),
                               FieldPea =
colSums(!is.na(Phen_data[Phen_data$PHENDom_Crop.Type=="Field Pea",])),
                               Lentil =
colSums(!is.na(Phen_data[Phen_data$PHENDom_Crop.Type=="Lentil",])),
                               Triticale =
colSums(!is.na(Phen_data[Phen_data$PHENDom_Crop.Type=="Triticale",])),
                               Wheat =
colSums(!is.na(Phen_data[Phen_data$PHENDom_Crop.Type=="Wheat",])))

write.csv(Phen_summary_table, "Phen_summary_table.csv")
rm(Phen_summary_table)



############################################################

## Figure SXX. Imputation noise under different methods

############################################################

## Makes a figure of model accuracy


#(file =
"model_frames_and_imputations/siteVariety_models/xgb_allTimes.rds")


alldata<-read.csv(file = "data/constructed_csv_files/alldata.csv", header =
T)

results<-readRDS("data/results_list.rds")

## sets up dev

tiff(filename  = "Publication_docs/Imputation_variability.tiff",
     width  =7.5, height = 5, units = "in", res  =330)

par(bty = "n", mar = c(5.1, 7.1,4.1,2.1))

## Makes a figure of imputation noise introduced
boxplot(MI_test_random_imp[,9],MI_test_missforest_imp[,9],
        MI_test_random_imp[,1],MI_test_missforest_imp[,1],
        MI_test_random_imp[,3],MI_test_missforest_imp[,3],
        MI_test_random_imp[,5],MI_test_missforest_imp[,5],
        MI_test_random_imp[,7],MI_test_missforest_imp[,7],
        names = c("plsr random", "plsr",
                  "rpart random", "rpart",
                  "BCRF random", "BCRF",
                  "xgBoost random", "xgBoost",
                  "lsvm random", "lsvm"),
        las = 2, ylim = c(0,1),
        col = c("orange", "lightblue"), horizontal = T, xlab = "Forecast
accuracy (RÂ²)")

abline(v = seq(0,1,by = 0.2), lty = 3)
abline(h = seq(0,10), col = "grey", lty = 3)

dev.off()

par(mfrow = c(1,1), mar = c(5.1, 4.1,4.1,2.1))

rm(alldata)

############################################################
```

```
## Fig. SXX Plots some common non-yield phenotypes, with sample size and
BCRF accuracy

##########################################################

CommonPhen_predictions<-readRDS(file =
"model_frames_and_imputations/site_models/CommonPhen_predictions.rds")

## [1] "PHENDom_Hectolitre_weight"       "PHENDom_Pct_below_2_0_mm"
"PHENDom_Protein"                 "PHENDom_Protein_4"
## [5] "PHENDom_Thousand_grain_weight"   "PHENDom_Early_Growth"
"PHENDom_yield_pct_of_average"    "PHENDom_zeroInflated_yield_t_ha"
## [9] "PHENDom_Flowering_QC"            "PHENDom_Glucosinolates"

## plots accuracy of phenotype models
namelist<-NULL
for(i in c(1:10)){
  bestfit_reg<-odregress(CommonPhen_predictions[[i]][,1],
                         CommonPhen_predictions[[i]][,2])

  namelist[[i]]<-ggplot(data.frame(Predicted =
CommonPhen_predictions[[i]][,2],
                                   Observed =
CommonPhen_predictions[[i]][,1]),
                        aes(x = Observed,y = Predicted)) +
    geom_abline(intercept = 0, slope = 1, col = "darkgrey")+
    geom_point(alpha = I(0.6), col = "orange", pch  =20, cex= 0.5)+
    geom_density2d()+
    geom_abline(slope = bestfit_reg$coeff[1], intercept =
bestfit_reg$coeff[2], col = "black", alpha = 0.7)+
    coord_fixed(xlim =
c(min(CommonPhen_predictions[[i]][,1]),max(CommonPhen_predictions[[i]][,1]))
,
                ylim =
c(min(CommonPhen_predictions[[i]][,1]),max(CommonPhen_predictions[[i]][,1]))
, ratio = 1)+
    annotate("text",
             x =
c(min(CommonPhen_predictions[[i]][,1])+c(max(CommonPhen_predictions[[i]][,1])/5))
,
             y = c(max(CommonPhen_predictions[[i]][,1])*0.9),
             label = paste0("RÂ² = ",
round(cor.test(CommonPhen_predictions[[i]][,1],

CommonPhen_predictions[[i]][,2])$estimate, digits = 2),
                            "\nN =
",length(CommonPhen_predictions[[i]][,1])))+
    theme_bw()
  ## prints target sample size
  print(length(CommonPhen_predictions[[i]][,1]))

}
names(CommonPhen_predictions)

# Canola    Chickpea  Faba Bean Lupin     Oat       Field Pea Lentil
Triticale Wheat
Pheno_grid<-grid.arrange(namelist[[1]]+labs(title = paste0("a   \n",
Common_phens[1])),
                         namelist[[2]]+labs(title = paste0("a   \n",
Common_phens[2])),
                         namelist[[3]]+labs(title = paste0("a   \n",
Common_phens[3])),
                         namelist[[4]]+labs(title = paste0("a   \n",
Common_phens[4])),
                         namelist[[5]]+labs(title = paste0("a   \n",
Common_phens[5])),
                         namelist[[6]]+labs(title = paste0("a   \n",
Common_phens[6])),
                         namelist[[7]]+labs(title = paste0("a   \n",
Common_phens[7])),
                         namelist[[8]]+labs(title = paste0("a   \n",
Common_phens[8])),
                         namelist[[9]]+labs(title = paste0("a   \n",
Common_phens[9])),
                         nrow =3)

Pheno_grid<-grid.arrange(namelist[[4]]+labs(title = "a   \nProtein %,
metadata"),
                         namelist[[8]]+labs(title = "b   \nFlowering Time
(Days)"),
                         namelist[[9]]+labs(title = "c   \nGlucosinolates
%"),
                         namelist[[1]]+labs(title = "d   \nHectolitre
Weight (Kg/hL)"),
                         namelist[[2]]+labs(title = "e   \n% Grain Below
2.0mm"),
                         namelist[[5]]+labs(title = "f   \n1000 Grain
Weight"),
                         nrow =2)

# Canola    Chickpea  Faba Bean Lupin     Oat       Field Pea Lentil
Triticale Wheat
#Pheno_grid<-grid.arrange(namelist[[4]]+labs(title = "a   \nProtein,
Metadata"),
#                         namelist[[6]]+labs(title = "b   \nFlowering Time
(Days)"),
#                         namelist[[7]]+labs(title = "c   \nGlucosinolate
Pct"),
#                         namelist[[1]]+labs(title = "d   \nHectolitre
Weight (Kg/hL)"),
#                         namelist[[2]]+labs(title = "e   \nPct Grain Below
2.0mm"),
#                         namelist[[5]]+labs(title = "f   \n1000 Grain
Weight"),
#                         nrow =2)

ggsave(Pheno_grid,file = "Publication_docs/Fig_SXX_Multiphenotypes.tiff",
device = "tiff", height = 5, width = 7.5,dpi = 330)

##########################################################

## Fig. SXX Plots the yield rpart tree, with explanations

##########################################################

Canola_rpart_TOS
```

```
plot(Wheat_rpart)
text(Wheat_rpart,cex= 0.4)

rpart.plot(Wheat_rpart)


rpart.plot(Canola_rpart)

rpart.rules(Canola_rpart)

prune_wheat<-prune(Wheat_rpart, cp = 0.005)
prune_canola<-prune(Canola_rpart, cp = 0.005)

rpart.plot(prune_wheat)
rpart.plot(prune_canola)

rpart.plot(Wheat_rpart_TOS)
rpart.rules(Wheat_rpart_TOS)

TOS_pruned<-prune(Wheat_rpart_TOS, cp = 0.02)
## NB gives better accuracy than unpruned model
cor.test(predict(TOS_pruned, newdata = test_frame_W),test_frame_W$Yield)
rmserr(predict(TOS_pruned, newdata = test_frame_W),test_frame_W$Yield)

TOS_pruned_C<-prune(Canola_rpart_TOS, cp = 0.01)

## NB accuracy suuuucks, r2= 0
cor.test(predict(TOS_pruned_C, newdata = test_frame_C),test_frame_C$Yield)
rmserr(predict(TOS_pruned_C, newdata = test_frame_C),test_frame_C$Yield)



## prnts a couple with no labels, for manual re-labeling

rpart.plot(prune_wheat,box.palette=scale_color_viridis_d(begin = 0, end =
7, option = "A"), fallen.leaves = F, branch = .4, tweak  =2)

rpart.plot(prune_wheat,box.palette=c("#ffffff","#faf398","#afd179","#68ba52","#09911a")
,
           fallen.leaves = F, branch = .4, tweak  =2, split.cex = 0.8,
branch.col = "darkgreen")

## Manual viridis type D pallette (Counterintuitive because >yellowing is
higher yield...)

rpart.plot(prune_wheat,box.palette=c("#26828EFF","#1F9E89FF","#35B779FF","#6DCD59FF")
,
           fallen.leaves = F, branch = .4, tweak  =2, split.cex = 0.8,
branch.col = "darkgreen")

## builds a dev.for the tree

tiff(filename  = "Publication_docs/FigS8_tree.tiff",
     width  =8, height = 8, units = "in", res  =330)

rpart.plot(prune_wheat,

box.palette=c("#faf398","#deffb3","#afd179","#68ba52","#09911a"),
           fallen.leaves = F, branch = .1, tweak  =2, split.cex = 0.75,
extra = 1, border.col = "darkgreen")

dev.off()


tiff(filename  = "Publication_docs/FigS9_tree.tiff",
     width  =5, height = 6, units = "in", res  =330)

rpart.plot(TOS_pruned,

box.palette=c("#faf398","#deffb3","#afd179","#68ba52","#09911a"),
           fallen.leaves = F, branch = .1, tweak  =1, split.cex = 0.75,
extra = 1, border.col = "darkgreen")

dev.off()


tiff(filename  = "Publication_docs/FigS10_tree.tiff",
     width  =8, height = 8, units = "in", res  =330)

rpart.plot(prune_canola,

box.palette=c("#faf398","#deffb3","#afd179","#68ba52","#09911a"),
           fallen.leaves = F, branch = .1, tweak  =2, split.cex = 0.75,
extra = 1, border.col = "darkgreen")

dev.off()

## And another for the TOS tree

tiff(filename  = "Publication_docs/FigS9_TOS_tree.tiff",
     width  =7.5, height = 6, units = "in", res  =330)

rpart.plot(TOS_pruned,

box.palette=c("#faf398","#deffb3","#afd179","#68ba52","#09911a"),
           fallen.leaves = F, branch = .4, tweak  =2, split.cex = 0.75,
extra = 1, split.col = "white")

dev.off()


rpart.plot(Wheat_rpart,

box.palette=c("#faf398","#deffb3","#afd179","#68ba52","#09911a"),
           fallen.leaves = F, branch = .4, tweak  =2, split.cex = 0.9,
extra = 1)


rpart.plot(prune_wheat,box.palette=c("YlGn"), fallen.leaves = F, branch =
.4, tweak  =2)



par(mfrow = c(2,1))

rpart.plot(prune_canola, cex = 0.4, box.palette="Greens", fallen.leaves =
F)
```

```
rpart.plot(prune_wheat, cex = 0.4, box.palette="Greens", fallen.leaves = F)

par(mfrow =c(1,1))

rpart.plot(prune_wheat, cex = 0.4, box.palette="Greens", fallen.leaves = F)


rpart.plot(Wheat_rpart_TOS, cex = 0.4, box.palette="Greens", fallen.leaves
= F)

rpart.plot(TOS_pruned, cex = 0.4, box.palette="Greens", fallen.leaves = F)

par(mfrow =c(1,1))

par(mfrow =c(2,1))

rpart.plot(prune_wheat, cex = 0.4, box.palette="Greens", fallen.leaves = F)

rpart.plot(TOS_pruned, cex = 0.4, box.palette="Greens", fallen.leaves = F)

par(mfrow =c(1,1))


Canola_rules<-rpart.rules(prune_canola)


rpart_mod<-readRDS(file =
"model_frames_and_imputations/siteVariety_models/rpart_allTimes.rds")

rpart_mod

tiff(filename  = "Publication_docs/unlabeled_tree.tiff",
     width  =7.5, height = 5, units = "in", res  =330)
plot(rpart_mod)
## saves this image, without text, for manual color-coding in figure
dev.off()

## writes the actual info, for manual coding
tiff(filename  = "Publication_docs/labeled_tree.tiff",
     width  =7.5, height = 5, units = "in", res  =1500)
plot(rpart_mod)
text(rpart_mod, cex = 0.05, col = "orange")
dev.off()

## prunes it down somewhat, to plot the top-level complexity
pruned_rp<-prune(rpart_mod, cp = 0.005)

plot(pruned_rp)
text(pruned_rp, cex = 0.5)
pruned_rp$cptable

## this accounts for 58% of yield

## prunes it down somewhat, to plot the top-level complexity
pruned_rp<-prune(rpart_mod, cp = 0.01)

plot(pruned_rp)
text(pruned_rp, cex = 0.5)

## this tree accounts for half the variation in yield across sites &
species

## this model accounts for


pruned_rp<-prune(rpart_mod, cp = 0.1)


############################################################

## Figure SXX plots wheat yield against frost severity

############################################################


mod_frame1<-readRDS("model_frames_and_imputations/mod_frame1.rds")

WheatYields<-data.frame(Yield = mod_frame1$PHENDom_yield_t_ha,
                        mod_frame1$ENVDom_Frost_Damage_score,
                        MinTemp =
mod_frame1$BOMDom_min_temperature_min_110,
                        MinTemp2 =
mod_frame1$BOMDom_min_temperature_min_120,
                        MinTemp3 =
mod_frame1$BOMDom_min_temperature_min_130,
                        MinTemp4 =
mod_frame1$BOMDom_min_temperature_min_140,
                        ZeroYield =
mod_frame1$PHENDom_zeroInflated_yield_t_ha)

WheatYields<-WheatYields[mod_frame1$METADom_sub_cropWheat==1,]

WheatYields$FlowerMinT<-unlist(lapply(seq(1:nrow(WheatYields)), function(i)
min(WheatYields[i,c(3:6)])))

Temp_anomaly<-ggplot(aes(FlowerMinT, Yield),data = WheatYields,
                     xlim = c(-10,20))+
  # geom_point(col = "orange", alpha = I(0.5))+
  geom_hex(bins = 50)+
  scale_fill_viridis("Observed Yields", option = "A")+
  #  geom_density2d()+
  geom_smooth(col = "lightblue")+
  geom_vline(xintercept = 0, col = "white", lty = 3)+
  geom_hline(yintercept = mean(WheatYields$Yield, na.rm = T), col =
"white", lty = 3)+
  theme_bw()+
  theme(legend.position = c(0.75,0.75))+
  labs(x = "Minimum Temperature (Deg. C) 110-140DAS",y = "Yield (t/Ha)")


## the sample size?
cor.test(WheatYields$FlowerMinT, WheatYields$Yield)
sum(!is.na(WheatYields$FlowerMinT+WheatYields$Yield))

## Saves this as a supp figure
ggsave(Temp_anomaly,file = "Publication_docs/Fig_SXX_FloweringTemps.tiff",
       device = "tiff", height = 7, width = 7.5,dpi = 330)
```

```
rm(Temp_anomaly, WheatYields)

###########################################################

## Figure 3XX. Gains in rolling forecast accuracy, w. nonstationarity vibes

###########################################################

forecast_frame_W<-readRDS(file =
"model_frames_and_imputations/siteVariety_models/Annual_rolling_Wheat_PredObs.rds"
)
forecast_frame_C<-readRDS(file =
"model_frames_and_imputations/siteVariety_models/Annual_rolling_Canola_PredObs.rds"
)

mod_frame1<-readRDS("model_frames_and_imputations/mod_frame1.rds")

## the mean yields
mean_wheat<-
aggregate(mod_frame1$PHENDom_yield_t_ha[mod_frame1$METADom_sub_cropWheat>0]
, mean, by =
list(mod_frame1$METADom_Year[mod_frame1$METADom_sub_cropWheat>0]), na.rm =
T)
mean_canola<-
aggregate(mod_frame1$PHENDom_yield_t_ha[mod_frame1$METADom_sub_cropCanola>0]
, mean, by =
list(mod_frame1$METADom_Year[mod_frame1$METADom_sub_cropCanola>0]), na.rm =
T)

## the variance in yields
var_wheat<-
aggregate(mod_frame1$PHENDom_yield_t_ha[mod_frame1$METADom_sub_cropWheat>0]
, var, by =
list(mod_frame1$METADom_Year[mod_frame1$METADom_sub_cropWheat>0]), na.rm =
T)
var_canola<-
aggregate(mod_frame1$PHENDom_yield_t_ha[mod_frame1$METADom_sub_cropCanola>0]
, var, by =
list(mod_frame1$METADom_Year[mod_frame1$METADom_sub_cropCanola>0]), na.rm =
T)

## sets up dev

tiff(filename  = "Publication_docs/Fig3.tiff",
     width  =7.5, height = 6.5, units = "in", res  =330)

## change as appropriate
Year_seq<-c(2009:2018)

#par(mfrow = c(1,2), bty = "n")

layout.matrix <- matrix(c(1,2,3,4), nrow = 2, ncol = 2)

layout(mat = layout.matrix,
       heights = c(2, 1), # Heights of the two rows
       widths = c(2, 2))

par(mar = c(0.1,4.1,2.1,2.1), bty = "n")

plot(Year_seq-0.1,
     forecast_frame_W[,3], type = "l", col = "grey",
     xlim = c(2008.5, 2018.5),
     ylim = c(-0.1,0.85),
     ylab = "Forecast accuracy RÂ²",
     xlab = "", las  =2, axes = F)
axis(2)

abline(h = seq(-0.2,0.8, by = 0.2), lty = 3, col = "grey")
abline(v = seq(2009,2018, by = 2), lty = 3, col = "grey")

points(Year_seq-0.1,forecast_frame_W[,3], pch = 20)
## adds CIs
for(i in 1:length(Year_seq)){
  points(c(Year_seq[[i]]-0.1,Year_seq[[i]]-0.1),
c(forecast_frame_W[i,4],forecast_frame_W[i,5]),
         type = "l")
}

## now lsvms
points(Year_seq,
       forecast_frame_W[,6], type = "l",
       col = "orange")
points(Year_seq,forecast_frame_W[,6], pch = 20, col = "orange")
## adds CIs
for(i in 1:length(Year_seq)){
  points(c(Year_seq[[i]],Year_seq[[i]]),
c(forecast_frame_W[i,7],forecast_frame_W[i,8]),
         type = "l", col = "orange")
}

## now PLSR
points(Year_seq+0.1,
       forecast_frame_W[,9], type = "l",
       col = "lightblue")
points(Year_seq+0.1,forecast_frame_W[,9], pch = 20, col = "lightblue")
## adds CIs
for(i in 1:length(Year_seq)){
  points(c(Year_seq[[i]]+0.1,Year_seq[[i]]+0.1),
c(forecast_frame_W[i,10],forecast_frame_W[i,11]),
         type = "l", col = "lightblue")
}
mtext("a", adj = 0)


par(mar = c(5.1,4.1,0.1,2.1))

boxplot(mod_frame1$PHENDom_yield_t_ha[which(mod_frame1$METADom_sub_cropWheat>
0 &
mod_frame1$METADom_Year>2008)]~mod_frame1$METADom_Year[which(mod_frame1$METADom_sub_cropWheat>
0 & mod_frame1$METADom_Year>2008)],
        las = 2, pch = "", ylim = c(0,8.5), ylab = "Yield (t/Ha)", xlab =
"Year")

par(mar = c(0.1,4.1,2.1,2.1))

plot(Year_seq-0.1,
     forecast_frame_C[,3], type = "l", col = "grey",
     xlim = c(2008.5, 2018.5),
```

```
     ylim = c(-0.1,0.85),
     ylab = "Forecast accuracy RÂ²",
     xlab = "", las = 2, axes = F)

axis(2)
abline(h = seq(-0.2,0.8, by = 0.2), lty = 3, col = "grey")
abline(v = seq(2009,2018, by = 2), lty = 3, col = "grey")

points(Year_seq-0.1,forecast_frame_C[,3], pch = 20)
## adds CIs
for(i in 1:length(Year_seq)){
  points(c(Year_seq[[i]]-0.1,Year_seq[[i]]-0.1),
c(forecast_frame_C[i,4],forecast_frame_C[i,5]),
         type = "l")
}

## now lsvms
points(Year_seq,
       forecast_frame_C[,6], type = "l",
       col = "orange")
points(Year_seq,forecast_frame_C[,6], pch = 20, col = "orange")
## adds CIs
for(i in 1:length(Year_seq)){
  points(c(Year_seq[[i]],Year_seq[[i]]),
c(forecast_frame_C[i,7],forecast_frame_C[i,8]),
         type = "l", col = "orange")
}

## now PLSR
points(Year_seq+0.1,
       forecast_frame_C[,9], type = "l",
       col = "lightblue")
points(Year_seq+0.1,forecast_frame_C[,9], pch = 20, col = "lightblue")
## adds CIs
for(i in 1:length(Year_seq)){
  points(c(Year_seq[[i]]+0.1,Year_seq[[i]]+0.1),
c(forecast_frame_C[i,10],forecast_frame_C[i,11]),
         type = "l", col = "lightblue")
}
mtext("b", adj = 0)


par(mar = c(5.1,4.1,0.1,2.1))

boxplot(mod_frame1$PHENDom_yield_t_ha[which(mod_frame1$METADom_sub_cropCanola>
0 &
mod_frame1$METADom_Year>2008)]~mod_frame1$METADom_Year[which(mod_frame1$METADom_sub_cropCanola>
0 & mod_frame1$METADom_Year>2008)],
        las = 2, pch = "", ylim = c(0,5), ylab = "Yield (t/Ha)", xlab =
"Year")

par(mfrow = c(1,1))

dev.off()

par(mar = c(5.1,4.1,2.1,2.1))

## plots the deltas (all are correlated, but data is super limited)

splom(cbind(diff(forecast_frame_W[,3]),diff(forecast_frame_W[,6]),diff(forecast_frame_W[,9]))
)
splom(cbind(diff(forecast_frame_C[,3]),diff(forecast_frame_C[,6]),diff(forecast_frame_C[,9]))
)

cor.test(diff(forecast_frame_W[,3]),diff(forecast_frame_W[,6]))
cor.test(diff(forecast_frame_W[,3]),diff(forecast_frame_W[,9]))
cor.test(diff(forecast_frame_W[,6]),diff(forecast_frame_W[,9]))

cor.test(diff(forecast_frame_C[,3]),diff(forecast_frame_C[,6]))
cor.test(diff(forecast_frame_C[,3]),diff(forecast_frame_C[,9]))
cor.test(diff(forecast_frame_C[,6]),diff(forecast_frame_C[,9]))

## mean rsq
c(0.782853+0.9316459+0.7910698)/3

##########################################################

## Fig. S3XX the gain in model accuracy throughout the season

##########################################################

forecast_frame_W<-readRDS(file =
"model_frames_and_imputations/siteVariety_models/rolling_Wheat_PredObs.rds"
)
forecast_frame_C<-readRDS(file =
"model_frames_and_imputations/siteVariety_models/rolling_Canola_PredObs.rds"
)

## plot the gain in model accuracy through the season

## black are Naive random forests, blue are plsr models, orange is linear
SVMs

tiff(filename  = "Publication_docs/Rolling forecasts.tiff",
     width  =7.5, height = 6, units = "in", res  =330)

par(mfrow = c(1,2), bty = "n")
plot(seq(1, 190, by = 10),forecast_frame_W[,1], type = "l",
     xlim = c(0,200), ylim = c(0,0.8),
     ylab = "Forecast accuracy RÂ²",
     xlab = "Days after Sowing")
points(seq(1, 190, by = 10),forecast_frame_W[,1], pch = 20)

points(seq(1, 190, by = 10),forecast_frame_W[,2], pch = 20, col = "orange")
points(seq(1, 190, by = 10),forecast_frame_W[,2], type = "l", col =
"orange")

points(seq(1, 190, by = 10),forecast_frame_W[,3], pch = 20, col =
"lightblue")
points(seq(1, 190, by = 10),forecast_frame_W[,3], type = "l", col =
"lightblue")

abline(h = seq(-1,1, by = 0.2), lty = 3)
mtext("a", adj = 0)


plot(seq(1, 190, by = 10),forecast_frame_C[,1], type = "l",
     xlim = c(0,200), ylim = c(0,0.8),
```

```
     ylab = "Forecast accuracy RÂ²",
     xlab = "Days after Sowing")
points(seq(1, 190, by = 10),forecast_frame_C[,1], pch = 20)

points(seq(1, 190, by = 10),forecast_frame_C[,2], pch = 20, col = "orange")
points(seq(1, 190, by = 10),forecast_frame_C[,2], type = "l", col =
"orange")

points(seq(1, 190, by = 10),forecast_frame_C[,3], pch = 20, col =
"lightblue")
points(seq(1, 190, by = 10),forecast_frame_C[,3], type = "l", col =
"lightblue")

abline(h = seq(-1,1, by = 0.2), lty = 3)
mtext("b", adj = 0)

par(mfrow = c(1,1))
dev.off()

rm(forecast_frame_W, forecast_frame_C)

#########################################################

## Figure 5 Plots reproducibility of variable importance scores

#########################################################

rpart_mod<-readRDS(file =
"model_frames_and_imputations/siteVariety_models/rpart_allTimes.rds")
rf_mod<-readRDS(file =
"model_frames_and_imputations/siteVariety_models/bcrf_allTimes.rds")
plsr_mod<-readRDS(file =
"model_frames_and_imputations/siteVariety_models/plsr_allTimes.rds")

imp_frame_comparison<-data.frame(varNames =
names(rpart_mod$variable.importance),
                                 rpart_imp = rpart_mod$variable.importance)
## merges on rf importance
imp_frame_comparison<-data.frame(imp_frame_comparison,
                                 rf_imp =
rf_mod$importance[match(imp_frame_comparison$varNames,rownames(rf_mod$importance))]
)

## gets the coefficients for important features under PLSR, rescaled to 0-1
p_coefs<-plsr_mod$coefficients/sum(sapply(plsr_mod$coefficients, abs))

## merges on plsr importance
imp_frame_comparison<-data.frame(imp_frame_comparison,
                                 plsr_imp =
p_coefs[match(imp_frame_comparison$varNames,rownames(p_coefs))])
#require(rminer)
#lsvm_imp<-Importance(lsvm_mod)

## the most important 100 plsr weights
tail(imp_frame_comparison[order(imp_frame_comparison$plsr_imp, na.last =
F),], 100)

qplot(c(imp_frame_comparison$rpart_imp), c(imp_frame_comparison$rf_imp))
qplot(c(imp_frame_comparison$plsr_imp), c(imp_frame_comparison$rf_imp))
qplot(c(imp_frame_comparison$plsr_imp), c(imp_frame_comparison$rpart_imp),
col = c(imp_frame_comparison$rf_imp))


imp_frame_comparison2<-imp_frame_comparison
imp_frame_comparison2$plsr_imp<-abs(imp_frame_comparison$plsr_imp)

imp_frame_comparison2[is.na(imp_frame_comparison2)]<-0

Imp_plot1<-qplot(plsr_imp, rf_imp, data = imp_frame_comparison2, alpha =
I(0),
                 ylab = "BCRF Importance", xlab = "PLSR Coefficient
Absolute Values")+
  geom_point(aes(y = rf_imp, x = plsr_imp), cex = 0.4, pch  =20)+
  ## adds the Latent Heat predictors as red circles
  geom_point(aes(y = rf_imp, x = plsr_imp),data =
imp_frame_comparison2[grep("Latent_heat",
imp_frame_comparison2$varNames),],
             col = "#ff0066", pch  =20, alpha = I(0.7))+
  ## adds the cumulative rainfall as blue droplets
  geom_point(aes(y = rf_imp, x = plsr_imp),data =
imp_frame_comparison2[grep("BOMDom_rainfall_csum_",
imp_frame_comparison2$varNames),],
             col = "lightblue", alpha = I(0.7), cex = 1.5)+
  ## adds the FPAR as a green circle
  geom_point(aes(y = rf_imp, x = plsr_imp),data =
imp_frame_comparison2[grep("SatDom_FPAR",
imp_frame_comparison2$varNames),],
             col = "darkgreen", alpha = I(0.7), pch = 20)+
  ## adds the EVI as a green open square
  geom_point(aes(y = rf_imp, x = plsr_imp),data =
imp_frame_comparison2[grep("SatDom_EVI", imp_frame_comparison2$varNames),],
             col = "darkgreen", alpha = I(0.7), pch = 20)+
  ## adds the NDVI as a green open triangle
  geom_point(aes(y = rf_imp, x = plsr_imp),data =
imp_frame_comparison2[grep("SatDom_NDVI",
imp_frame_comparison2$varNames),],
             col = "darkgreen", alpha = I(0.7), pch = 20)+
  ##Direct labels some points
  geom_text(aes(y = rf_imp, x = plsr_imp),data =
imp_frame_comparison2[c(1),], label = "Dicot t/f", nudge_x = 0.0000017,
nudge_y = 0, size=3)+
  geom_point(aes(y = rf_imp, x = plsr_imp),data =
imp_frame_comparison2[c(1),], pch  =20)+
  geom_text(aes(y = rf_imp, x = plsr_imp),data =
imp_frame_comparison2[c(9),], label = "Canola t/f", nudge_x = 0.0000017,
nudge_y = 0, size=3)+
  geom_point(aes(y = rf_imp, x = plsr_imp),data =
imp_frame_comparison2[c(9),], pch  =20)+
  geom_text(aes(y = rf_imp, x = plsr_imp),data =
imp_frame_comparison2[c(8),], label = "Clethodim", nudge_x = -0.0000023,
nudge_y = 0, size=3)+
  geom_point(aes(y = rf_imp, x = plsr_imp),data =
imp_frame_comparison2[c(8),], pch  =20)+
  # geom_text(aes(y = rf_imp, x = plsr_imp),data =
imp_frame_comparison2[c(10),], label = "Fabaceae t/f", nudge_x = 0.0000017,
nudge_y = 0, size=3)+
  #  geom_point(aes(y = rf_imp, x = plsr_imp),data =
imp_frame_comparison2[c(10),], pch  =20)+
```

```
  geom_text(aes(y = rf_imp, x = plsr_imp),data =
imp_frame_comparison2[c(11),], label = "Applied Sulphur", nudge_x =
0.0000027, nudge_y = 0, size=3)+
  geom_point(aes(y = rf_imp, x = plsr_imp),data =
imp_frame_comparison2[c(11),], pch  =20)+
  geom_text(aes(y = rf_imp, x = plsr_imp),data =
imp_frame_comparison2[c(13),], label = "Wheat t/f", nudge_x = 0.0000017,
nudge_y = 0, size=3)+
  geom_point(aes(y = rf_imp, x = plsr_imp),data =
imp_frame_comparison2[c(13),], pch  =20)+
  geom_point(aes(y = rf_imp, x = plsr_imp),data =
imp_frame_comparison2[c(13),], pch  =20)+
  theme_bw()


Imp_plot2<-qplot(plsr_imp, rpart_imp, data = imp_frame_comparison2, alpha =
I(0),
                 ylab = "RPRM Importance", xlab = "PLSR Coefficient
Absolute Values")+
  geom_point(aes(y = rpart_imp, x = plsr_imp), cex = 0.4, pch  =20)+
  ## adds the Latent Heat predictors as red circles
  geom_point(aes(y = rpart_imp, x = plsr_imp),data =
imp_frame_comparison2[grep("Latent_heat",
imp_frame_comparison2$varNames),],
             col = "#ff0066", pch  =20, alpha = I(0.7))+
  ## adds the cumulative rainfall as blue droplets
  geom_point(aes(y = rpart_imp, x = plsr_imp),data =
imp_frame_comparison2[grep("BOMDom_rainfall_csum_",
imp_frame_comparison2$varNames),],
             col = "lightblue", alpha = I(0.7), cex = 1.5)+
  ## adds the FPAR as a green circle
  geom_point(aes(y = rpart_imp, x = plsr_imp),data =
imp_frame_comparison2[grep("SatDom_FPAR",
imp_frame_comparison2$varNames),],
             col = "darkgreen", alpha = I(0.7), pch = 20)+
  ## adds the EVI as a green open square
  geom_point(aes(y = rpart_imp, x = plsr_imp),data =
imp_frame_comparison2[grep("SatDom_EVI", imp_frame_comparison2$varNames),],
             col = "darkgreen", alpha = I(0.7), pch = 20)+
  ## adds the NDVI as a green open triangle
  geom_point(aes(y = rpart_imp, x = plsr_imp),data =
imp_frame_comparison2[grep("SatDom_NDVI",
imp_frame_comparison2$varNames),],
             col = "darkgreen", alpha = I(0.7), pch = 20)+
  ##Direct labels some points
  geom_text(aes(y = rpart_imp, x = plsr_imp),data =
imp_frame_comparison2[c(1),], label = "Dicot t/f", nudge_x = 0.0000017,
nudge_y = 0, size=3)+
  geom_point(aes(y = rpart_imp, x = plsr_imp),data =
imp_frame_comparison2[c(1),], pch  =20)+
  geom_text(aes(y = rpart_imp, x = plsr_imp),data =
imp_frame_comparison2[c(9),], label = "Canola t/f", nudge_x = 0.0000023,
nudge_y = 0, size=3)+
  geom_point(aes(y = rpart_imp, x = plsr_imp),data =
imp_frame_comparison2[c(9),], pch  =20)+
  geom_text(aes(y = rpart_imp, x = plsr_imp),data =
imp_frame_comparison2[c(8),], label = "Clethodim", nudge_x = 0.0000024,
nudge_y = 0, size=3)+
  geom_point(aes(y = rpart_imp, x = plsr_imp),data =
imp_frame_comparison2[c(8),], pch  =20)+
  geom_text(aes(y = rpart_imp, x = plsr_imp),data =
imp_frame_comparison2[c(10),], label = "Fabaceae t/f", nudge_x = 0.0000026,
nudge_y = 0, size=3)+
  geom_point(aes(y = rpart_imp, x = plsr_imp),data =
imp_frame_comparison2[c(10),], pch  =20)+
  geom_text(aes(y = rpart_imp, x = plsr_imp),data =
imp_frame_comparison2[c(11),], label = "Applied Sulphur", nudge_x =
0.0000026, nudge_y = 1000, size=3)+
  geom_point(aes(y = rpart_imp, x = plsr_imp),data =
imp_frame_comparison2[c(11),], pch  =20)+
  geom_text(aes(y = rpart_imp, x = plsr_imp),data =
imp_frame_comparison2[c(13),], label = "Wheat t/f", nudge_x = 0.0000017,
nudge_y = 0, size=3)+
  geom_point(aes(y = rpart_imp, x = plsr_imp),data =
imp_frame_comparison2[c(13),], pch  =20)+
  theme_bw()

Imp_plot3<-qplot(rpart_imp, rf_imp,data = imp_frame_comparison2, alpha =
I(0),
                 xlab = "RPRM Importance", ylab = "BCRF Importance")+
  geom_point(aes(x = rpart_imp, y = rf_imp), cex = 0.4, pch  =20)+
  ## adds the Latent Heat predictors as pink circles
  geom_point(aes(x = rpart_imp, y = rf_imp),data =
imp_frame_comparison2[grep("Latent_heat",
imp_frame_comparison2$varNames),],
             col = "#ff0066", pch  =20, alpha = I(0.7))+
  ## adds the cumulative rainfall as blue droplets
  geom_point(aes(x = rpart_imp, y = rf_imp),data =
imp_frame_comparison2[grep("BOMDom_rainfall_csum_",
imp_frame_comparison2$varNames),],
             col = "lightblue", alpha = I(0.7), cex = 1.5)+
  ## adds the raw reflectance bands as an orange square
  # geom_point(aes(x = rpart_imp, y = rf_imp),data =
imp_frame_comparison2[grep("SatDom_Reflect_b",
imp_frame_comparison2$varNames),],
  #             col = "orange", alpha = I(0.7), pch  =15)+
  ## adds the FPAR as a green circle
  geom_point(aes(x = rpart_imp, y = rf_imp),data =
imp_frame_comparison2[grep("SatDom_FPAR",
imp_frame_comparison2$varNames),],
             col = "darkgreen", alpha = I(0.7), pch = 20)+
  ## adds the EVI as a green open square
  geom_point(aes(x = rpart_imp, y = rf_imp),data =
imp_frame_comparison2[grep("SatDom_EVI", imp_frame_comparison2$varNames),],
             col = "darkgreen", alpha = I(0.7), pch = 20)+
  ## adds the NDVI as a green open triangle
  geom_point(aes(x = rpart_imp, y = rf_imp),data =
imp_frame_comparison2[grep("SatDom_NDVI",
imp_frame_comparison2$varNames),],
             col = "darkgreen", alpha = I(0.7), pch = 20)+
  ##Direct labels some points
  geom_text(aes(x = rpart_imp, y = rf_imp),data =
imp_frame_comparison2[c(1),], label = "Dicot t/f", nudge_x = -5000, nudge_y
= -9, size=3)+
  geom_point(aes(x = rpart_imp, y = rf_imp),data =
imp_frame_comparison2[c(1),], pch  =20)+
  geom_text(aes(x = rpart_imp, y = rf_imp),data =
imp_frame_comparison2[c(9),], label = "Canola t/f", nudge_x = 0.000001,
```

```
nudge_y = +7, size=3)+
  geom_point(aes(x = rpart_imp, y = rf_imp),data =
imp_frame_comparison2[c(9),], pch  =20)+
  geom_text(aes(x = rpart_imp, y = rf_imp),data =
imp_frame_comparison2[c(8),], label = "Clethodim", nudge_x = -0.000001,
nudge_y = -7, size=3)+
  geom_point(aes(x = rpart_imp, y = rf_imp),data =
imp_frame_comparison2[c(8),], pch  =20)+
  geom_text(aes(x = rpart_imp, y = rf_imp),data =
imp_frame_comparison2[c(10),], label = "Fabaceae t/f", nudge_x = 0.0000017,
nudge_y = -7, size=3)+
  geom_point(aes(x = rpart_imp, y = rf_imp),data =
imp_frame_comparison2[c(10),], pch  =20)+
  geom_text(aes(x = rpart_imp, y = rf_imp),data =
imp_frame_comparison2[c(11),], label = "Applied Sulphur", nudge_x =
0.0000017, nudge_y = +7, size=3)+
  geom_point(aes(x = rpart_imp, y = rf_imp),data =
imp_frame_comparison2[c(11),], pch  =20)+
  geom_text(aes(x = rpart_imp, y = rf_imp),data =
imp_frame_comparison2[c(13),], label = "Wheat t/f", nudge_x = 0.0000017,
nudge_y = -7, size=3)+
  geom_point(aes(x = rpart_imp, y = rf_imp),data =
imp_frame_comparison2[c(13),], pch  =20)+
  ## annotates
  geom_text(x = 30000, y = 75, label = "Latent Heat Flux", col = "#ff0066",
size=3)+
  geom_text(x = 15000, y = 33, label = "Cumulative Rainfall", col =
"lightblue", size=3)+
  geom_text(x = 23000, y = 8, label = "Vegetation Indices", col =
"darkgreen", size=3)+
  theme_bw()

Imp_plot<-grid.arrange(Imp_plot3+labs(title =
"a")+theme(text=element_text(size=9)),
                       Imp_plot1+labs(title =
"b")+theme(text=element_text(size=9)),
                       Imp_plot2+labs(title =
"c")+theme(text=element_text(size=9)),
                       qplot(c(0,0), c(10,10)),
                       nrow = 2)

#ggsave(Imp_plot,file = "Publication_docs/Fig_SXX3_Importance.tiff", device
= "tiff", height = 3, width = 7.5,dpi = 330)


## Generates a plot of cumulative seasonal rainfall importance, etc.
imp_frame_comparison2<-imp_frame_comparison
imp_frame_comparison2$plsr_imp<-abs(imp_frame_comparison$plsr_imp)

imp_frame_comparison2[is.na(imp_frame_comparison2)]<-0

## Strips the timing off the end of all variables

## gets negative values
Var_Timings<-lapply(as.character(imp_frame_comparison$varNames), strsplit,
split = "\\.\\.")
Var_Timings<-lapply(Var_Timings, unlist)
Var_Timings<-lapply(Var_Timings, tail, 1)
Var_Timings<-as.numeric(unlist(Var_Timings))
Var_Timings<-c(-(Var_Timings))

Var_Timings2<-lapply(as.character(imp_frame_comparison$varNames), strsplit,
split = "_\\.")
Var_Timings2<-lapply(Var_Timings2, unlist)
Var_Timings2<-lapply(Var_Timings2, tail, 1)
Var_Timings2<-as.numeric(unlist(Var_Timings2))
Var_Timings2<-c(-Var_Timings2)

Var_Timings[is.na(Var_Timings)]<-Var_Timings2[is.na(Var_Timings)]

Var_Timings2<-lapply(as.character(imp_frame_comparison$varNames), strsplit,
split = "_")
Var_Timings2<-lapply(Var_Timings2, unlist)
Var_Timings2<-lapply(Var_Timings2, tail, 1)
Var_Timings2<-as.numeric(unlist(Var_Timings2))
Var_Timings2[which(Var_Timings2<10 & Var_Timings2!=0)]<-NA

Var_Timings[is.na(Var_Timings)]<-Var_Timings2[is.na(Var_Timings)]
Var_Timings[Var_Timings==c(-42)]<-NA

## attaches to importance frame
imp_frame_comparison2<-data.frame(Var_Timings,imp_frame_comparison2)
## orders all data by these timing values
imp_frame_comparison2<-
imp_frame_comparison2[order(imp_frame_comparison2$Var_Timings),]
## removes NAs
imp_frame_comparison2<-
imp_frame_comparison2[!is.na(imp_frame_comparison2$Var_Timings),]

imp_frame_comparison3<-imp_frame_comparison2

imp_frame_comparison3$Water<-c(imp_frame_comparison2$varNames %in%
imp_frame_comparison2$varNames[grep("rainfall_csum",

imp_frame_comparison2$varNames)])
imp_frame_comparison3$rpart_imp<-log(imp_frame_comparison3$rpart_imp)
imp_frame_comparison3$rf_imp<-log(imp_frame_comparison3$rf_imp)
imp_frame_comparison3$plsr_imp<-log(imp_frame_comparison3$plsr_imp)

imp_frame_comparison3<-imp_frame_comparison3[-
c(which(imp_frame_comparison3$Var_Timings>300)),]

imp_frame_comparison3<-
imp_frame_comparison3[imp_frame_comparison3$rf_imp>=0,]

Importance_timings<-ggplot(imp_frame_comparison3, aes(Var_Timings,
rf_imp,group = Var_Timings))+
  geom_boxplot(outlier.size = 0)+
  ## adds the cumulative rainfall as blue droplets
  geom_point(aes(x = Var_Timings, y = rf_imp),
             data = imp_frame_comparison3[grep("BOMDom_rainfall_csum_",
imp_frame_comparison3$varNames),],
             col = "lightblue")+
  geom_point(aes(x = Var_Timings, y = rf_imp),
             data = imp_frame_comparison3[grep("BOMDom_rainfall_csum_",
imp_frame_comparison3$varNames),],
             col = "black", pch = 1)+
  geom_point(aes(x = Var_Timings, y = rf_imp),
```

```
            data = imp_frame_comparison3[grep("Latent_heat",
imp_frame_comparison3$varNames),],
            col = "#ff0066", alpha = I(0.7))+
  geom_smooth(col = "black", method = "loess", span = 0.1)+
  geom_vline(xintercept = 0, col = "red", alpha =I(0.5))+
  theme_bw()+
  theme(legend.position = c(0.1,0.8))+
  labs(x = "Days After Sowing", y = "Variable Importance")

Importance_timings

## Finished Figure 5

Imp_plot<-grid.arrange(Imp_plot3+labs(title =
"a")+theme(text=element_text(size=9)),
                      Imp_plot1+labs(title =
"b")+theme(text=element_text(size=9)),
                      Imp_plot2+labs(title =
"c")+theme(text=element_text(size=9)),
                      Importance_timings+labs(title =
"d")+theme(text=element_text(size=9)),
                      nrow = 2)

ggsave(Imp_plot,file = "Publication_docs/Fig_5_Importance.tiff", device =
"tiff", height = 8, width = 7.5,dpi = 330)




imp_frame_comparison3<-imp_frame_comparison2
imp_frame_comparison3$varNames<-
as.character(imp_frame_comparison3$varNames)
imp_frame_comparison3$varNames<-
gsub("\\.","_",imp_frame_comparison3$varNames)
imp_frame_comparison3$varNames<-
lapply(as.character(imp_frame_comparison3$varNames), strsplit, split =
"_[0-9]")
imp_frame_comparison3$varNames<-lapply(imp_frame_comparison3$varNames,
unlist)
imp_frame_comparison3$varNames<-
unlist(lapply(imp_frame_comparison3$varNames, head, 1))

imp_frame_comparison3<-reshape(imp_frame_comparison3, direction = "wide",
timevar = "Var_Timings", idvar = "varNames")

## Plots the importance score of variables relative to sowing dates
#imp_frame_comparison3<-
imp_frame_comparison2[grep("Latent_heat",imp_frame_comparison2$varNames),]

imp_frame_comparison3<-
imp_frame_comparison2[grep("rainfall_csum",imp_frame_comparison2$varNames),
]

## Plots the importance score of variables relative to sowing dates
imp_frame_comparison3$rpart_imp<-
c(imp_frame_comparison3$rpart_imp/max(imp_frame_comparison3$rpart_imp,
na.rm  =T))
imp_frame_comparison3$rf_imp<-
c(imp_frame_comparison3$rf_imp/max(imp_frame_comparison3$rf_imp, na.rm
=T))
imp_frame_comparison3$plsr_imp<-
c(imp_frame_comparison3$plsr_imp/max(imp_frame_comparison3$plsr_imp, na.rm
=T))

## merges on all times, coerces zeroes
imp_frame_comparison3<-merge(data.frame(Times_vec = seq(-80, 250, by =
10)), imp_frame_comparison3,
                             by.x = "Times_vec", by.y = "Var_Timings", all
= T)

#imp_frame_comparison3[is.na(imp_frame_comparison3)]<-0

## Plots the importance score of variables relative to sowing dates
ggplot(data = imp_frame_comparison3,aes(Times_vec,rpart_imp))+
  geom_point(col = "lightblue")+
  geom_line(col = "lightblue")+
  geom_point(aes(Times_vec,rf_imp), col = "orange")+
  geom_line(aes(Times_vec,rf_imp), col = "orange")+
  geom_point(aes(Times_vec,plsr_imp))+
  geom_line(aes(Times_vec,plsr_imp))+
  theme_bw()

## Does the same with latent heat flux
imp_frame_comparison3<-imp_frame_comparison2[grep("Latent_heat_[0-
9]",imp_frame_comparison2$varNames),]

imp_frame_comparison3$rpart_imp<-
c(imp_frame_comparison3$rpart_imp/max(imp_frame_comparison3$rpart_imp,
na.rm  =T))
imp_frame_comparison3$rf_imp<-
c(imp_frame_comparison3$rf_imp/max(imp_frame_comparison3$rf_imp, na.rm
=T))
imp_frame_comparison3$plsr_imp<-
c(imp_frame_comparison3$plsr_imp/max(imp_frame_comparison3$plsr_imp, na.rm
=T))

## Plots the importance score of variables relative to sowing dates
qplot(Var_Timings,rpart_imp,data = imp_frame_comparison3, xlim = c(-
50,250))+
  geom_line(aes(Var_Timings,rpart_imp), col = "black")+
  geom_point(aes(Var_Timings,rf_imp),data = imp_frame_comparison3, col =
"orange")+
  geom_line(aes(Var_Timings,rf_imp),data = imp_frame_comparison3, col =
"orange")+
  geom_point(aes(Var_Timings,plsr_imp),data = imp_frame_comparison3, col =
"lightblue")+
  geom_line(aes(Var_Timings,plsr_imp),data = imp_frame_comparison3, col =
"lightblue")+
  theme_bw()

## And the vegetation Indices
imp_frame_comparison3<-imp_frame_comparison2[c(grep("VI_[0-
9]",imp_frame_comparison2$varNames),grep("FPAR_[0-
9]",imp_frame_comparison2$varNames)),]

imp_frame_comparison3$rpart_imp<-
c(imp_frame_comparison3$rpart_imp/max(imp_frame_comparison3$rpart_imp,
na.rm  =T))
```

```
imp_frame_comparison3$rf_imp<-
c(imp_frame_comparison3$rf_imp/max(imp_frame_comparison3$rf_imp, na.rm
=T))
imp_frame_comparison3$plsr_imp<-
c(imp_frame_comparison3$plsr_imp/max(imp_frame_comparison3$plsr_imp, na.rm
=T))

## Plots the importance score of variables relative to sowing dates
qplot(Var_Timings,rpart_imp,data = imp_frame_comparison3, xlim = c(-
50,250))+
  geom_line(aes(Var_Timings,rpart_imp), col = "black")+
  geom_point(aes(Var_Timings,rf_imp),data = imp_frame_comparison3, col =
"orange")+
  geom_line(aes(Var_Timings,rf_imp),data = imp_frame_comparison3, col =
"orange")+
  geom_point(aes(Var_Timings,plsr_imp),data = imp_frame_comparison3, col =
"lightblue")+
  geom_line(aes(Var_Timings,plsr_imp),data = imp_frame_comparison3, col =
"lightblue")+
  theme_bw()

qplot(Var_Timings,c(rpart_imp/max(rpart_imp)),data = imp_frame_comparison3,
xlim = c(-50,250))+
  geom_line(aes(Var_Timings,c(rpart_imp/max(rpart_imp))), col = "black")+
  geom_point(aes(Var_Timings,c(rf_imp/max(rf_imp))),data =
imp_frame_comparison3, col = "orange")+
  geom_line(aes(Var_Timings,c(rf_imp/max(rf_imp))),data =
imp_frame_comparison3, col = "orange")+
  geom_point(aes(Var_Timings,c(plsr_imp/max(plsr_imp, na.rm = T))),data =
imp_frame_comparison3, col = "lightblue")+
  geom_line(aes(Var_Timings,c(plsr_imp/max(plsr_imp, na.rm = T))),data =
imp_frame_comparison3, col = "lightblue")+
  theme_bw()



########################################


## Figure 1a


##############################################################

## Fig 1

##############################################################

## makes a leaflet map of locations, coloured by crop
lat_longs<-read.csv(file =
"data/constructed_csv_files/geolocs_output1.csv", header = T)

lat_longs<-lat_longs[!is.na(rowMeans(lat_longs[,c(3:4)])),]

## merges on site-average trial yields and other phenotypes

results<-readRDS("data/results_list.rds")

results$Trial_ID<-toupper(results$`Trial ID`)

xobj<-merge(lat_longs, results, by.x = "Trial_ID", by.y = "Trial_ID", all =
T)

xobj<-xobj[!duplicated(paste0(xobj$Trial_ID, xobj$`Sowing Date`,
xobj$Latitude, xobj$`Site Mean (t/ha)`)),]

xobj<-xobj[!is.na(xobj$Trial_ID),]

xobj$`Site Mean (t/ha)`<-as.numeric(as.character(xobj$`Site Mean (t/ha)`))

xobj_canola<-xobj[which(xobj$`Crop Type`=="Canola"),]

ggplot(data = ne_countries(country = "australia", scale = "medium",
returnclass = "sf")) +
  geom_sf(color = "black", fill = "black") +
  annotation_scale(location = "bl", width_hint = 0.5) +
  annotation_north_arrow(location = "bl", which_north = "true",
                         pad_x = unit(0.75, "in"), pad_y = unit(0.5, "in"),
                         style = north_arrow_fancy_orienteering) +
  coord_sf(ylim = c(-10, -43), xlim = c(115, 155))+
  theme_bw()+
  scale_color_viridis(xobj$Sowing_date_numeric.x, option = "D")+
  geom_point(data = xobj, aes(Longitude,Latitude, col =
Sowing_date_numeric.x), pch  =20, alpha = I(0.5), cex = 0.5)

AusMap<-ggplot(data = ne_countries(country = "australia", scale = "medium",
returnclass = "sf")) +
  geom_sf(color = "black", fill = "white") +
  annotation_scale(location = "bl", width_hint = 0.5) +
  annotation_north_arrow(location = "bl", which_north = "true",
                         pad_x = unit(0.4, "npc"), pad_y = unit(0.4,
"npc"),
                         style = north_arrow_nautical) +
  coord_sf(ylim = c(-10, -43), xlim = c(112, 153))+
  theme_bw()+
  scale_color_viridis(xobj$Sowing_date_numeric.x, option = "D")+
  geom_point(data = xobj, aes(Longitude,Latitude, col =
Sowing_date_numeric.x),  pch  =20, alpha = I(0.8), cex = 0.5)+
  theme(legend.position = "none")


## lays out figure 1
gridMatrix<-matrix(c(1,1,1,2,2,2,2,
                     1,1,1,2,2,2,2,
                     1,1,1,2,2,2,2,
                     3,3,3,3,3,3,3,
                     3,3,3,3,3,3,3,
                     3,3,3,3,3,3,3,
                     3,3,3,3,3,3,3,
                     3,3,3,3,3,3,3), nrow = 8, byrow = T)

Fig_1<-grid.arrange(AusMap+labs(title = "a")+
                      theme(text = element_text(size=9)),
                    Photosynth_Fig1b+labs(title = "b")+
                      theme(text = element_text(size=9)),
                    Umap_alldata_figure+labs(title = "c")+
                      theme(text = element_text(size=9)),
                    nrow = 2, layout_matrix = gridMatrix)
```

```
#Fig_1<-grid.arrange(AusMap, Photosynth_Fig1b, Umap_alldata_figure, nrow =
1)

ggsave(Fig_1,file = "Publication_docs/Fig1.tiff", device = "tiff", height =
8, width = 7,dpi = 330)



LeafMap = leaflet() %>%
  addProviderTiles("CartoDB.DarkMatter") %>%
  addCircleMarkers(lat_longs$Longitude, lat_longs$Latitude,
                   radius  =0.5, fillOpacity = 0.2,
                   color = heat.colors(lat_longs$Sowing_date_numeric))

LeafMap



############################################################

## Figure SXX: distribution of select phenotypes

############################################################


mod_frame1<-readRDS("model_frames_and_imputations/mod_frame1.rds")

## separates phenotypes
Phen_data<-mod_frame1[,grep("PHENDom", colnames(mod_frame1))]
mod_frame1<-mod_frame1[,-c(grep("PHENDom", colnames(mod_frame1)))]

## sets up dev
tiff(filename  = "Publication_docs/Agronomic_trait_boxplots.tiff",
     width  =7.5, height = 7, units = "in", res  =330)

par(mfrow = c(2,3), bty = "n")

## wheat yield and protein
boxplot(Phen_data$PHENDom_yield_t_ha[Phen_data$PHENDom_Crop_alldata=="Wheat"]~mod_frame1$METADom_Year[Phen_data$PHENDom_Crop_alldata=="Wheat"]
,
        xlab = "Year", ylab = "Yield, t/Ha", pch = ".", main  ="a", las =
2, varwidth  =T)
abline(h = seq(0,14, by  =2), lty = 3, col = "lightgrey")

boxplot(Phen_data$PHENDom_Flowering_QC[Phen_data$PHENDom_Crop_alldata=="Wheat"]~mod_frame1$METADom_Year[Phen_data$PHENDom_Crop_alldata=="Wheat"]
,
        xlab = "Year", ylab = "Days to Flowering", pch = ".", main  ="b",
las = 2, varwidth  =T)
abline(h = seq(50,150, by  =50), lty = 3, col = "lightgrey")

boxplot(Phen_data$PHENDom_Protein[Phen_data$PHENDom_Crop_alldata=="Wheat"]~mod_frame1$METADom_Year[Phen_data$PHENDom_Crop_alldata=="Wheat"]
,
        xlab = "Year", ylab = "Protein, % dry weight", pch = ".", main
="c", las = 2, varwidth  =T)
abline(h = seq(0,40, by  =10), lty = 3, col = "lightgrey")

boxplot(Phen_data$PHENDom_yield_t_ha[Phen_data$PHENDom_Crop_alldata=="Canola"]~mod_frame1$METADom_Year[Phen_data$PHENDom_Crop_alldata=="Canola"]
,
        xlab = "Year", ylab = "Yield, t/Ha", pch = ".", main  ="d", las =
2, varwidth  =T)
abline(h = seq(0,5, by  =1), lty = 3, col = "lightgrey")

boxplot(as.numeric(Phen_data$PHENDom_Flowering_QC)[Phen_data$PHENDom_Crop_alldata=="Canola"]~mod_frame1$METADom_Year[Phen_data$PHENDom_Crop_alldata=="Canola"]
,
        xlab = "Year", ylab = "Days to Flowering", pch = ".", main  ="e",
las = 2, varwidth  =T)
abline(h = seq(50,200, by  =50), lty = 3, col = "lightgrey")

boxplot(as.numeric(Phen_data$PHENDom_Glucosinolates)[Phen_data$PHENDom_Crop_alldata=="Canola"]~mod_frame1$METADom_Year[Phen_data$PHENDom_Crop_alldata=="Canola"]
,
        xlab = "Year", ylab = "Glucosinolates, %", pch = ".", main  ="f",
las = 2, varwidth  =T)
abline(h = seq(0,50, by  =10), lty = 3, col = "lightgrey")

par(mfrow = c(1,1))

dev.off()


tiff(filename  = "Publication_docs/Species_Yields_boxplots.tiff",
     width  =7.5, height = 7, units = "in", res  =330)

par(mfrow = c(3,3), bty = "n", mar = c(4.1,3.1,2.1,2.1))

boxplot(Phen_data$PHENDom_yield_t_ha[Phen_data$PHENDom_Crop_alldata=="Wheat"]~mod_frame1$METADom_Year[Phen_data$PHENDom_Crop_alldata=="Wheat"]
,
        main = "Wheat, N=54658",
        xlab = "Year", ylab = "yield, t/Ha", pch = ".", las = 2, varwidth
=T)

abline(h = seq(0,14, by  =2), lty = 3, col = "lightgrey")

boxplot(Phen_data$PHENDom_yield_t_ha[Phen_data$PHENDom_Crop_alldata=="Oat"]~mod_frame1$METADom_Year[Phen_data$PHENDom_Crop_alldata=="Oat"]
,
        main = "Oat, N=3873",
        xlab = "Year", ylab = "yield, t/Ha", pch = ".", las = 2, varwidth
=T)

abline(h = seq(0,13, by  =1), lty = 3, col = "lightgrey")

boxplot(Phen_data$PHENDom_yield_t_ha[Phen_data$PHENDom_Crop_alldata=="Triticale"]~mod_frame1$METADom_Year[Phen_data$PHENDom_Crop_alldata=="Triticale"]
,
        main = "Triticale, N=2081",
        xlab = "Year", ylab = "yield, t/Ha", pch = ".", las = 2, varwidth
=T)

abline(h = seq(0,13, by  =1), lty = 3, col = "lightgrey")

boxplot(Phen_data$PHENDom_yield_t_ha[Phen_data$PHENDom_Crop_alldata=="Canola"]~mod_frame1$METADom_Year[Phen_data$PHENDom_Crop_alldata=="Canola"]
,
        main = "Canola, N=14929",
        xlab = "Year", ylab = "yield, t/Ha", pch = ".", las = 2, varwidth
=T)

abline(h = seq(0,13, by  =1), lty = 3, col = "lightgrey")

boxplot(Phen_data$PHENDom_yield_t_ha[Phen_data$PHENDom_Crop_alldata=="Chickpea"]~mod_frame1$METADom_Year[Phen_data$PHENDom_Crop_alldata=="Chickpea"]
```

```
,
        main = "Chickpea, N=3677",
        xlab = "Year", ylab = "yield, t/Ha", pch = ".", las = 2, varwidth
=T)

abline(h = seq(0,13, by  =1), lty = 3, col = "lightgrey")

boxplot(Phen_data$PHENDom_yield_t_ha[Phen_data$PHENDom_Crop_alldata=="Field
Pea"]~mod_frame1$METADom_Year[Phen_data$PHENDom_Crop_alldata=="Field Pea"],
        main = "Field Pea, N=3896",
        xlab = "Year", ylab = "yield, t/Ha", pch = ".", las = 2, varwidth
=T)

abline(h = seq(0,13, by  =1), lty = 3, col = "lightgrey")

boxplot(Phen_data$PHENDom_yield_t_ha[Phen_data$PHENDom_Crop_alldata=="Faba
Bean"]~mod_frame1$METADom_Year[Phen_data$PHENDom_Crop_alldata=="Faba
Bean"],
        main = "Faba Bean, N=2224",
        xlab = "Year", ylab = "yield, t/Ha", pch = ".", las = 2, varwidth
=T)

abline(h = seq(0,13, by  =1), lty = 3, col = "lightgrey")

boxplot(Phen_data$PHENDom_yield_t_ha[Phen_data$PHENDom_Crop_alldata=="Lentil"]~mod_frame1$METADom_Year[Phen_data$PHENDom_Crop_alldata=="Lentil"]
,
        main = "Lentil, N=2260",
        xlab = "Year", ylab = "yield, t/Ha", pch = ".", las = 2, varwidth
=T)

abline(h = seq(0,13, by  =1), lty = 3, col = "lightgrey")

boxplot(Phen_data$PHENDom_yield_t_ha[Phen_data$PHENDom_Crop_alldata=="Lupin"]~mod_frame1$METADom_Year[Phen_data$PHENDom_Crop_alldata=="Lupin"]
,
        main = "Lupin, N=2747",
        xlab = "Year", ylab = "yield, t/Ha", pch = ".", las = 2, varwidth
=T)

abline(h = seq(0,13, by  =1), lty = 3, col = "lightgrey")

par(mfrow = c(1,1))

dev.off()


############################################################

## Figure SXX-SXX. Accuracy of yield and protein models by species

############################################################


## Yield models at +100DAS or +200DAS
Yield_predobs_100<-readRDS(file =
"model_frames_and_imputations/siteVariety_models/Yield_predobs_100.rds")
Protein_predobs_100<-readRDS(file =
"model_frames_and_imputations/siteVariety_models/Protein_predobs_100.rds")

Yield_predobs_200<-readRDS(file =
"model_frames_and_imputations/siteVariety_models/Yield_predobs_200.rds")
Protein_predobs_200<-readRDS(file =
"model_frames_and_imputations/siteVariety_models/Protein_predobs_200.rds")

## makes a figure of model forecast accuracy, versus internal (missing
plot) accuracy
# Canola    Chickpea  Faba Bean Lupin     Oat       Field Pea Lentil
Triticale Wheat

#saveRDS(lsvm_mod, file =
paste0("model_frames_and_imputations/siteVariety_models/rf_Yield_100DAS",
#                                spp_list[[i]],".rds"))
#saveRDS(rf_mod, file =
paste0("model_frames_and_imputations/siteVariety_models/lsvm_Yield_100DAS",
#                               spp_list[[i]],".rds"))

qplot(Yield_predobs_100[[1]][1,],
      jitter(Yield_predobs_100[[1]][3,], factor = 50),
      # xlim = c(0,12), ylim = c(0,12),
      alpha = I(0),
      xlab = "Predicted Yield, t/Ha", ylab = "Observed Yield, t/Ha")+
  geom_point(col = "orange", alpha = I(0.7))+
  geom_smooth(method = "lm", col = "black")+
  theme_bw()


## plots accuracy of yield+protein models across species
namelist<-NULL
for(i in 1:9){
  bestfit_reg<-odregress(Yield_predobs_200[[i]][1,],
                         Yield_predobs_200[[i]][2,])

  namelist[[i]]<-ggplot(data.frame(Predicted =
jitter(Yield_predobs_200[[i]][2,], factor = 50),
                                   Observed = Yield_predobs_200[[i]][1,]),
                        aes(x = Observed,y = Predicted)) +
    geom_abline(intercept = 0, slope = 1, col  ="darkgrey")+
    geom_point(alpha = I(0.6), col = "orange", pch  =20, cex= 0.5)+
    geom_abline(slope = bestfit_reg$coeff[1], intercept =
bestfit_reg$coeff[2], col = "black", alpha = 0.7)+
    coord_fixed(xlim = c(0,max(Yield_predobs_200[[i]][1,])), ylim =
c(0,max(Yield_predobs_200[[i]][1,])), ratio = 1)+
    annotate("text",
             x = c(max(Yield_predobs_200[[i]][1,])/2),
             y = 0.5,
             label = paste0("RÂ² = ",
round(cor.test(Yield_predobs_200[[i]][1,],

Yield_predobs_200[[i]][2,])$estimate, digits = 2)))+
    theme_bw()
  ## prints target sample size
  print(length(Yield_predobs_200[[i]][1,]))

}


# Canola    Chickpea  Faba Bean Lupin     Oat       Field Pea Lentil
Triticale Wheat
Yield_grid<-grid.arrange(namelist[[1]]+labs(title = "a   Canola"),
```

```
                         namelist[[2]]+labs(title = "b   Chickpea"),
                         namelist[[3]]+labs(title = "c   Faba Bean"),
                         namelist[[4]]+labs(title = "d   Lupin"),
                         namelist[[5]]+labs(title = "e   Oat"),
                         namelist[[6]]+labs(title = "f   Field Pea"),
                         namelist[[7]]+labs(title = "g   Lentil"),
                         namelist[[8]]+labs(title = "h   Triticale"),
                         namelist[[9]]+labs(title = "i   Wheat"), nrow =3)

# Canola    Chickpea  Faba Bean Lupin     Oat       Field Pea Lentil
Triticale Wheat
Yield_grid<-grid.arrange(namelist[[1]]+labs(title = "a")+
                           annotate("text", label ="Canola",
                                    x =
c(max(Yield_predobs_200[[1]][1,])/2),
                                    y =
c(max(Yield_predobs_200[[1]][1,])*0.9)),
                         namelist[[2]]+labs(title = "b")+
                           annotate("text", label ="Chickpea",
                                    x =
c(max(Yield_predobs_200[[2]][1,])/2),
                                    y =
c(max(Yield_predobs_200[[2]][1,])*0.9)),
                         namelist[[3]]+labs(title = "c")+
                           annotate("text", label ="Faba Bean",
                                    x =
c(max(Yield_predobs_200[[3]][1,])/2),
                                    y =
c(max(Yield_predobs_200[[3]][1,])*0.9)),
                         namelist[[4]]+labs(title = "d ")+
                           annotate("text", label ="Lupin",
                                    x =
c(max(Yield_predobs_200[[4]][1,])/2),
                                    y =
c(max(Yield_predobs_200[[4]][1,])*0.9)),
                         namelist[[5]]+labs(title = "e ")+
                           annotate("text", label ="Oat",
                                    x =
c(max(Yield_predobs_200[[5]][1,])/2),
                                    y =
c(max(Yield_predobs_200[[5]][1,])*0.9)),
                         namelist[[6]]+labs(title = "f")+
                           annotate("text", label ="Field Pea",
                                    x =
c(max(Yield_predobs_200[[6]][1,])/2),
                                    y =
c(max(Yield_predobs_200[[6]][1,])*0.9)),
                         namelist[[7]]+labs(title = "g")+
                           annotate("text", label ="Lentil",
                                    x =
c(max(Yield_predobs_200[[7]][1,])/2),
                                    y =
c(max(Yield_predobs_200[[7]][1,])*0.9)),
                         namelist[[8]]+labs(title = "h")+
                           annotate("text", label ="Triticale",
                                    x =
c(max(Yield_predobs_200[[8]][1,])/2),
                                    y =
c(max(Yield_predobs_200[[8]][1,])*0.9)),
                         namelist[[9]]+labs(title = "i")+
                           annotate("text", label ="Wheat",
                                    x =
c(max(Yield_predobs_200[[9]][1,])/2),
                                    y =
c(max(Yield_predobs_200[[9]][1,])*0.9)), nrow =3)



ggsave(Yield_grid,file = "Publication_docs/Fig_SXX_YieldGrid200DAS.tiff",
device = "tiff", height = 7, width = 7,dpi = 330)


Yield_grid<-grid.arrange(namelist[[1]]+labs(title = "a   Canola"),
                         namelist[[2]]+labs(title = "b   Chickpea"),
                         namelist[[3]]+labs(title = "c   Faba Bean"),
                         namelist[[4]]+labs(title = "d   Lupin"),
                         namelist[[5]]+labs(title = "e   Oat"),
                         namelist[[6]]+labs(title = "f   Field Pea"),
                         namelist[[7]]+labs(title = "g   Lentil"),
                         namelist[[8]]+labs(title = "h   Triticale"),
                         namelist[[9]]+labs(title = "i   Wheat"), nrow =3)


## Repeats for Protein predictions
namelist<-NULL
for(i in 1:3){
  bestfit_reg<-odregress(Protein_predobs_200[[i]][1,],
                         Protein_predobs_200[[i]][2,])

  namelist[[i]]<-ggplot(data.frame(Predicted =
jitter(Protein_predobs_200[[i]][2,], factor = 50),
                                   Observed =
Protein_predobs_200[[i]][1,]),
                        aes(x = Observed,y = Predicted)) +
    geom_abline(intercept = 0, slope = 1, col = "darkgrey")+
    geom_point(alpha = I(0.6), col = "orange", pch  =20, cex= 0.5)+
    geom_abline(slope = bestfit_reg$coeff[1], intercept =
bestfit_reg$coeff[2], col = "black", alpha = 0.7)+
    coord_fixed(xlim = c(5,max(Protein_predobs_200[[i]][1,])), ylim =
c(5,max(Protein_predobs_200[[i]][1,])), ratio = 1)+
    geom_density2d(alpha = I(0.7))+
    annotate("text",
             x = c(max(Protein_predobs_200[[i]][1,])/2)+2.5,
             y = 5.5,
             label = paste0("RÂ² = ",
round(cor.test(Protein_predobs_200[[i]][1,],

Protein_predobs_200[[i]][2,])$estimate, digits = 2)))+
    theme_bw()
  ## prints target sample size
  print(length(Protein_predobs_200[[i]][1,]))

}

## and for protein 100DAS
## Repeats for Protein predictions
namelist2<-NULL
for(i in 1:3){
```

```
  bestfit_reg<-odregress(Protein_predobs_100[[i]][1,],
                         Protein_predobs_100[[i]][2,])

  namelist2[[i]]<-ggplot(data.frame(Predicted =
jitter(Protein_predobs_100[[i]][2,], factor = 50),
                                    Observed =
Protein_predobs_100[[i]][1,]),
                         aes(x = Observed,y = Predicted)) +
    geom_abline(intercept = 0, slope = 1, col = "darkgrey")+
    geom_point(alpha = I(0.6), col = "orange", pch  =20, cex= 0.5)+
    geom_abline(slope = bestfit_reg$coeff[1], intercept =
bestfit_reg$coeff[2], col = "black", alpha = 0.7)+
    coord_fixed(xlim = c(5,max(Protein_predobs_100[[i]][1,])), ylim =
c(5,max(Protein_predobs_100[[i]][1,])), ratio = 1)+
    geom_density2d(alpha = I(0.7))+
    annotate("text",
             x = c(max(Protein_predobs_100[[i]][1,])/2)+2.5,
             y = 5.5,
             label = paste0("RÂ² = ",
round(cor.test(Protein_predobs_100[[i]][1,],

Protein_predobs_100[[i]][2,])$estimate, digits = 2)))+
    theme_bw()
  ## prints target sample size
  print(length(Protein_predobs_100[[i]][1,]))

}

#Protein_grid<-grid.arrange(namelist[[1]]+labs(title = "a   Oat"),
#                          namelist[[2]]+labs(title = "b   Triticale"),
#                          namelist[[3]]+labs(title = "c   Wheat"), nrow =1)

# Canola    Chickpea  Faba Bean Lupin     Oat       Field Pea Lentil
Triticale Wheat
Protein_grid<-grid.arrange(namelist[[1]]+labs(title = "a   Oat 200DAS"),
                           namelist[[2]]+labs(title = "b   Triticale
200DAS"),
                           namelist[[3]]+labs(title = "c   Wheat 200DAS"),
                           namelist2[[1]]+labs(title = "d   Oat 100DAS"),
                           namelist2[[2]]+labs(title = "e   Triticale
100DAS"),
                           namelist2[[3]]+labs(title = "f   Wheat 100DAS"),
nrow =2)

ggsave(Protein_grid,file = "Publication_docs/Fig_SXX_ProteinGrid.tiff",
device = "tiff", height = 5, width = 7,dpi = 330)

lsvm_100DAS_wheat<-ggplot(data.frame(Predicted_yield =
Yield_predobs_100[[9]][3,],
                                     Observed_yield =
Yield_predobs_100[[9]][1,]),
                          aes(x = Predicted_yield, y = Observed_yield),
                          alpha = I(0)) +
  geom_hex(bins = 50)+
  geom_abline(intercept = 0, slope = 1)+
  geom_smooth(method = "lm", col = "orange")+
  coord_fixed(xlim = c(0,12), ylim = c(0,12), ratio = 1)+
  scale_fill_viridis()+
  theme_bw()

cor.test(Yield_predobs_100[[9]][3,],Yield_predobs_100[[9]][1,])

bcrf_100DAS_wheat<-ggplot(data.frame(Predicted_yield =
Yield_predobs_100[[9]][2,],
                                     Observed_yield =
Yield_predobs_100[[9]][1,]),
                          aes(x = Predicted_yield, y = Observed_yield),
                          alpha = I(0)) +
  geom_hex(bins = 50)+
  geom_abline(intercept = 0, slope = 1)+
  geom_smooth(method = "lm", col = "orange")+
  coord_fixed(xlim = c(0,12), ylim = c(0,12), ratio = 1)+
  scale_fill_viridis()+
  theme_bw()


cor.test(Yield_predobs_100[[9]][2,],Yield_predobs_100[[9]][1,])

grid.arrange(bcrf_100DAS_wheat, lsvm_100DAS_wheat, nrow =1)



lsvm_100DAS_canola<-ggplot(data.frame(Predicted_yield =
Yield_predobs_100[[1]][3,],
                                      Observed_yield =
Yield_predobs_100[[1]][1,]),
                           aes(x = Predicted_yield, y = Observed_yield),
                           alpha = I(0)) +
  geom_hex(bins = 50)+
  geom_abline(intercept = 0, slope = 1)+
  geom_smooth(method = "lm", col = "orange")+
  coord_fixed(xlim = c(0,3), ylim = c(0,3), ratio = 1)+
  scale_fill_viridis()+
  theme_bw()

cor.test(Yield_predobs_100[[1]][3,],Yield_predobs_100[[1]][1,])

bcrf_100DAS_canola<-ggplot(data.frame(Predicted_yield =
Yield_predobs_100[[1]][2,],
                                      Observed_yield =
Yield_predobs_100[[1]][1,]),
                           aes(x = Predicted_yield, y = Observed_yield),
                           alpha = I(0)) +
  geom_hex(bins = 50)+
  geom_abline(intercept = 0, slope = 1)+
  geom_smooth(method = "lm", col = "orange")+
  coord_fixed(xlim = c(0,3), ylim = c(0,3), ratio = 1)+
  scale_fill_viridis()+
  theme_bw()

cor.test(Yield_predobs_100[[1]][2,],Yield_predobs_100[[1]][1,])

require(gridExtra)
grid.arrange(bcrf_100DAS_wheat, lsvm_100DAS_wheat,
             bcrf_100DAS_canola, lsvm_100DAS_canola, nrow =2)

namelist<-NULL
for(i in 1:9){
```

```
  namelist[[i]]<-ggplot(data.frame(Predicted = Yield_predobs_100[[i]][2,],
                                   Observed = Yield_predobs_100[[i]][1,]),
                        aes(x = Predicted, y = Observed),
                        alpha = I(0.6)) +
    geom_hex(bins = 50)+
    geom_abline(intercept = 0, slope = 1)+
    geom_smooth(method = "lm", col = "orange")+
    coord_fixed(xlim = c(0,max(Yield_predobs_100[[i]][1,])), ylim =
c(0,max(Yield_predobs_100[[i]][1,])), ratio = 1)+
    scale_fill_viridis()+
    theme_bw()
}

grid.arrange(namelist[[1]],
             namelist[[2]],
             namelist[[3]],
             namelist[[4]],
             namelist[[5]],
             namelist[[6]],
             namelist[[7]],
             namelist[[8]],
             namelist[[9]], nrow =3)


namelist<-NULL
for(i in 1:9){

  namelist[[i]]<-ggplot(data.frame(Predicted = Yield_predobs_100[[i]][3,],
                                   Observed = Yield_predobs_100[[i]][1,]),
                        aes(x = Predicted, y = Observed),
                        alpha = I(0.6)) +
    geom_hex(bins = 50)+
    geom_abline(intercept = 0, slope = 1)+
    geom_smooth(method = "lm", col = "orange")+
    coord_fixed(xlim = c(0,max(Yield_predobs_100[[i]][1,])), ylim =
c(0,max(Yield_predobs_100[[i]][1,])), ratio = 1)+
    scale_fill_viridis()+
    theme_bw()
}
grid.arrange(namelist[[1]],
             namelist[[2]],
             namelist[[3]],
             namelist[[4]],
             namelist[[5]],
             namelist[[6]],
             namelist[[7]],
             namelist[[8]],
             namelist[[9]], nrow =3)
```
